# The infection-tolerant mammalian reservoir of Lyme disease and other zoonoses broadly counters the inflammatory effects of endotoxin

**DOI:** 10.1101/2020.12.13.422519

**Authors:** Gabriela Balderrama-Gutierrez, Ana Milovic, Vanessa J. Cook, M. Nurul Islam, Youwen Zhang, Hippokratis Kiaris, John T. Belisle, Ali Mortazavi, Alan G. Barbour

## Abstract

Animals that are competent natural reservoirs of zoonotic diseases commonly suffer little morbidity from the pathogens they persistently harbor. The mechanisms of this infection tolerance and the trade-off costs are poorly understood. We used exposure to a single dose of lipopolysaccharide (LPS) endotoxin as an experimental model of inflammation to compare the responses of the cricentine rodent *Peromyscus leucopus*, the white-footed deermouse, to that of *Mus musculus*, the standard laboratory model for pathogenesis studies. Four hours after injection with either LPS or saline, blood and spleen and liver tissues were collected postmortem and subjected to RNA-seq, untargeted metabolomics, and specific RT-qPCR. This was followed by analysis of differential expression at the gene, pathway, and empirical network levels. The deermice showed the same signs of sickness as the mice with LPS exposure, and in addition demonstrated comparable increases in levels of corticosterone and expression of interleukin (IL)-6, tumor necrosis factor, IL-1β, and acute phase reactants, including C-reactive protein. But whereas the *M. musculus* response to LPS was best-characterized by network analysis as cytokine-associated, the *P. leucopus* response was dominated by pathway terms associated with neutrophil activity. Dichotomies between the species in expression profiles of arginase 1 and nitric oxide synthase 2, as well as the ratios of IL-10 to IL-12, were consistent with a type M1 polarized macrophage response in the mice and a type M2 or alternatively-activated response in the deermice. Analysis of metabolites in the plasma and RNA in the tissues revealed differences between the two species in tryptophan metabolism during response to LPS. Two up-regulated genes in particular signified the difference between the species: Slpi (secretory leukocyte proteinase inhibitor) and Ibsp (integrin-binding protein sialoprotein). The latter was previously unrecognized in the context of inflammation or infection. Key RNA-seq findings in *P. leucopus* were replicated in a second LPS experiment with older animals, in a systemic bacterial infection model, and with cultivated fibroblasts. Taken together, the results indicate that the deermouse possesses several adaptive traits to moderate effects of inflammation and oxidative stress ensuing from infection. This seems to be at the cost of infection persistence and that is to the benefit of the pathogen.

## INTRODUCTION

*Peromyscus leucopus*, the white-footed deermouse, is a major reservoir for several infectious diseases of humans (reviewed in [1]). These include Lyme disease [2], as well as varieties of anaplasmosis, babesiosis, relapsing fever, ehrlichiosis, and viral encephalitis. *P. leucopus* owes this distinction to its broad distribution across the eastern and central U.S. [3, 4], its abundance in a variety of habitats, and its role as an important host for the tick vectors of disease [5]. This rodent’s immune system and other defenses may keep the pathogens at bay, but the infections persist nevertheless [6, 7], thereby increasing the likelihood of a tick acquiring the microbe during its blood meal on that host [8]. Where enzootic transmission is pervasive the majority of *P. leucopus* live out their lives parasitized by one or more of these pathogens [9–11]. If there is a fitness cost, scrutiny has not revealed it in the field [11] or laboratory [1, 12–14].

The term applied to this phenomenon of persistent infection with little or no morbidity is infection tolerance [15–18], and was first described in plants [19, 20]. This usage is distinguished from the “tolerance” by the immune system for self-antigens [21]. But in both contexts the word conveys a moderation of the host response and avoidance of injury. Infection tolerance mainly manifests at the organismal level, for instance, by signs of illness, measures of disability, and various systemic biomarkers, such as cytokines in the blood. These are likely explained by events at the cellular and molecular levels [22, 23], but these have not been fully explored.

*P. leucopus’* tolerance of infection is matched by another deermouse, *P. maniculatus*, a major reservoir for a hantavirus [24, 25]. *P. maniculatus* has also been infected with the COVID-19 coronavirus and transmits it to cagemates. Yet infected animals displayed only moderate pathology and recovered within a few day [26, 27]. Other examples of the tolerance phenomenon are found among bat species implicated as reservoirs for SARS-CoV, Ebola, Nipah, and Hendra viruses [28, 29]. These deermice and bats exhibit a trade-off between defensive processes that check pathogen proliferation and processes that limit collateral damage from those defenses. The result is a state of persistent low-grade infection with limited pathology.

The innate and adaptive host defenses that concertedly provide resistance against pathogens are well known. Less understood are the mechanisms on the other side of the trade-off [30], namely, those that curb sickness and illness due to disordered inflammation [22, 31]. Plausible accounts for the phenomenon are many and invoke a variety of pathways. An important consideration for any host-pathogen system is whether resistance and tolerance mechanisms are separable, tightly linked, or partially co-dependent [16].

*P. leucopus* is well-suited as a model for investigations of infection tolerance in a genetically diverse population [32–34]. While colloquially known as “mice”, the genus *Peromyscus*, together with hamsters and voles, belong to the family Cricetidae and not the family Muridae, which includes Old World rats and mice, like the house mouse, *Mus musculus* [35]. We have previously sequenced the genome of *P. leucopus*, annotated its transcriptome using RNA-seq, and characterized its gut microbiomes [36–38]. There was little difference by RNA-seq between the blood of *P. leucopus* experimentally infected with the Lyme disease agent *Borreliella burgdorferi* and that of uninfected controls [36], a finding consistent with the mildness of deermouse infections with this pathogen in comparison to the pathology and disability observed in infected *M. musculus* [39]. Accordingly, we looked to alternatives that would more robustly elicit inflammation.

The study compared the responses to a single dose of bacterial lipopolysaccharide (LPS) of *P. leucopus* to that of *M. musculus*. This endotoxin leads to inflammation through its binding to a pattern recognition molecule and complex ensuing cascades [40]. Tolerance of endotoxin under experimental conditions is a long-recognized phenomenon [41], but this usually refers to a diminution in the severity of the response through cumulative previous exposures of animals or isolated cells to LPS [42, 43], rather than an inherent disposition of a naïve animal to survive a toxic dose. Some spontaneous and engineered Mendelian traits do render a mouse less susceptible to LPS through an alteration of the Toll-like receptor TLR4 for LPS or in key downstream mediators, such as MyD88, in the cascade [44, 45]. A single gene, however, is unlikely to account for what appears to be more nuanced interplays between complex contending forces in *Peromyscus* and other examples.

LPS exposure is a useful place to begin. This simple experimental system is a proxy for acute infection with a bacterial pathogen but without the confounding variables of changing numbers of the microbe and the inevitable appearance of acquired immunity. Our working assumption is that early events in the response are determinants of the eventual outcomes of the infection, whether assessed from the perspective of pathogen burden or host disability. In the course of the study we identified several pathways involved with inflammation, oxidative stress, phagocytosis, and metabolism that distinguished *P. leucopus* and *M. musculus* in their responses to LPS.

## RESULTS

### Susceptibility of *P. leucopus* to LPS

There are no published studies of responses of any *Peromyscus* species to different doses of LPS or the consequences, but there are several on *M. musculus* and its susceptibility to LPS [46–53]. The LD_50_ for *M. musculus*, i.e. the extrapolated dose at which on average 50% of the animals die, was reported in the range of 5 to 20 µg of LPS per gm of body weight. The average across different strains of inbred and outbred mice was ~15 µg/gm. Death usually occurred between 12 and 48 hours after injection. Our prior study of adult BALB/c mice, found that animals injected intraperitoneally with 10 µg LPS per gm body weight survived for at least 24 h. At 4 h they manifested, as signs of illness, reduced activity and ruffled fur and concomitantly had marked elevations in serum concentrations of tumor necrosis factor alpha (TNF), interleukin-6 (IL-6), and IL-10 and the chemokines C-C motif ligand 4 (CCL4) and CCL2 [54].

We began with a study of the effect of single doses of purified *E. coli* LPS administered intraperitoneally to adult *P. leucopus* and then monitored for seven days. Panel A of Figure 1 includes the survival curve for animals in groups of six receiving doses of 10, 50, 100, 200, or 300 µg per gm and examined continuously for the first 8 h and then at 12 h intervals thereafter. Death or moribund state occurred in at least one animal at all dosage groups except the 10 µg per gm group. Five of 6 in the 50 µg dose group survived. At higher doses the fatality rate was higher, with death occurring between days 2 and 5. Remarkably, 3 of 6 of the animals receiving the highest dose of 300 µg per gm, or a total dose of 6 mg on average per animal, survived. Survivors among the *P. leucopus* at that and the 100 µg and 200 µg per gm doses appeared to have fully recovered by 7 days after the injection. From these data we could not calculate a precise LD_50_, possibly because of the genetic heterogeneity of the outbred population [36]. But we estimated it to between 100 and 300 µg/gm from this experiment.

**Figure 1.**
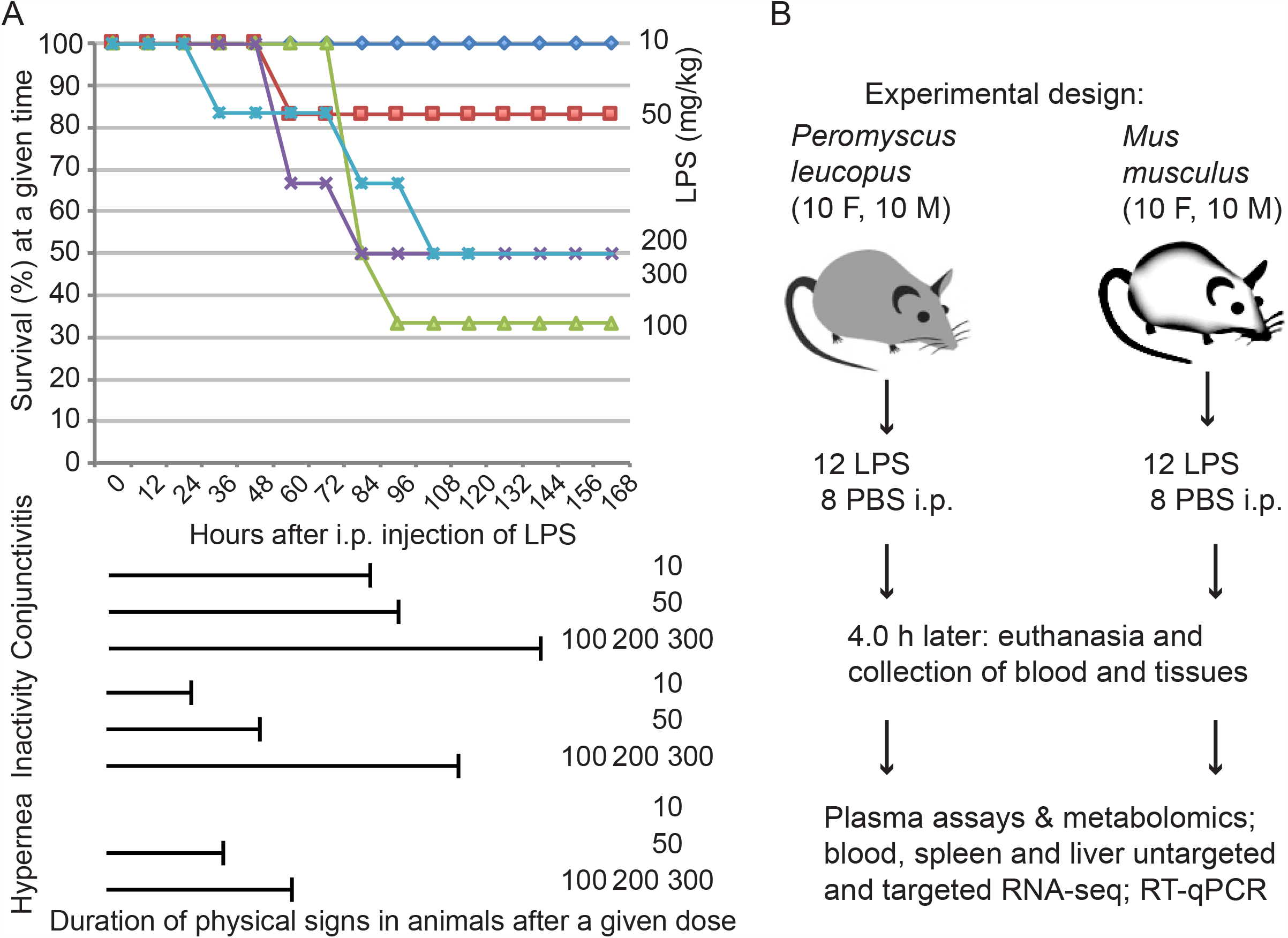
Studies of lipopolysaccharide (LPS) effects on *Peromyscus leucopus* alone and in comparison with *Mus musculus*. Panel A. Dose response of *P. leucopus* to LPS. Groups of 6 adult animals received different intraperitoneal (i.p.) doses on a mg per gm body weight basis of *Escherichia coli* LPS at time 0 and then monitored for physical signs of sickness (conjunctivitis, inactivity, and hyperpnea) and survival over the succeeding 7 days (168 h). Panel B. Experimental design of comparative study of short-term effects of LPS on *P. leucopus* and *Mus musculus*. Animals received 10 mg/gm body weight LPS in phosphate-buffered saline (PBS) or PBS alone.

There were no descriptions of exudative inflammation of the conjunctivae or conjunctivitis (Figure S1) in the literature on single parenteral doses of LPS in laboratory mice, and we did not observe this disease sign in the *M. musculus* in this experiment. In contrast, we observed this sign in *P. leucopus* at all doses by 12 h and its persistence for up to 6 days in the highest dose group (Figure 1A). All animals displayed reduced activity, as defined by the behavior of huddling in groups, moving only for eating and drinking. The durations of this sign, as well as for the sign of hyperpnea, or rapid breathing, in animals receiving doses of 50 µg/gm or higher, correlated with the dose amounts.

These results indicated that *P. leucopus* was susceptible to the physical and behavioral effects of LPS at single doses having similar consequences for *M. musculus*. Even at the lowest dose all the animals displayed ill-effects of this treatment. But in comparison to reported findings in *M. musculus*, a substantial proportion of deermice receiving doses that were 10-20 fold higher than the reported LD_50_ for the house mouse, did not further deteriorate and then recovered within a few days. This suggested to us that the LPS model recapitulated in a simple model some of the effects of an acute infection. Furthermore, it exemplifies this species’ tolerance of inflammation during infection.

### Experimental design and effects of LPS on mice and deermice

We compared the short-term responses of the outbred *P. leucopus* with the inbred BALB/c strain of *M. musculus* and of both sexes to a single dose of LPS. The comparison of animals from a heterogeneous close colony with an inbred population provided was a means to gauge the diversity of response among the deermice in the experiment. Panel B of Figure 1 summarizes the experimental design and Table S1 lists the characteristics of the animals in the experiment and selected parameters. The same animals were also the subject of gut microbiome analysis [38], and alpha diversity values of the microbiota from the earlier study are provided. The animals of each species were sexually-mature adults and comparable in size and age, though *P. leucopus* tended to be smaller and older by 1-2 weeks. With the exception of one pair of siblings split between the treatment and control goups (Table S1), the *P. leucopus* were offspring of different mating pairs. We alternated the administration of the LPS by species and by sex within a species to control for time of day effects. Animals were euthanized in the same order, and the intervals between animals were kept within a strict limit.

All animals of both species displayed effects of the LPS within an hour of the injection: reduced activity and the sign of ruffed fur. But only among the 12 LPS-treated *P. leucopus* did we again observe conjunctivitis, in this case in six animals (exact Likelihood Ratio test *p* = 0.025), equally distributed between females and males (Table S1). In a pilot study we found that the specific antibodies used for commercially-available immunoassays for various cytokines, chemokines, and acute phase reactants of *M. musculus* were insufficiently cross-reactive with orthologous proteins of *P. leucopus* to be useful. Accordingly, we applied two assays of plasma for substances that were identical in composition between *Peromyscus* and *Mus*: the steroid corticosterone, which serves the same function in rodents as cortisol in primates, and nitric oxide.

For corticosterone concentrations the mean (95% confidence intervals) for *M. musculus* controls (n = 7) and LPS-treated (n = 12) were 77 (34-120) and 624 (580-667), respectively (*p* < 10^−11^). Corresponding values for *P. leucopus* control (n = 7) and LPS-treated (n =12) were 186 (79-293) and 699 (670-727), respectively (*p* < 10^−8^). There were marginally higher baseline levels of corticosterone in *Peromyscus* than in *Mus* (*p* =0.10), but they had similar levels after LPS. The assay for nitric oxide demonstrated higher levels in 11 *M. musculus* treated with LPS (mean 29 [20–37]) than in seven controls (mean of 7 [3–12]) (*p* = 0.008). But there was not an elevation in nine *P. leucopus* after LPS compared with six controls: 7 (3-11) vs 7 (3-12) (*p* = 0.9).

### Metabolomics of plasma

We carried out untargeted metabolomics by LC/MS of the plasma of the LPS-treated and control *P. leucopus* and *M. musculus*. Of the 8125 molecular features (MF) identified in the *M. musculus* plasma samples, 123 (1.5%) differed between LPS-treated and control animals with a FDR < 0.05 and an absolute fold change of > 2.0. Of the 7714 MF identified in the *P. leucopus* plasma samples, 215 (2.8%) correspondingly differed between treated and control animals. Subsequent analysis focused on pathway enrichment, which allowed cross-species comparison of differences in the identifiable metabolites between treated and untreated animals. For the *M. musculus* and the *P. leucopus* samples the number of identified pathways with at least one KEGG-defineable compound were 76 and 73, respectively, of which 73 were in common (Table S2). Figure 2A plots pathway enrichment in deermice against that in mice for each of the pathways in common. In both species there was significant enrichment of the steroid hormone biosynthesis pathway in animals 4 h after LPS injection. This finding was consistent with the results of the immunoassays of corticosterone in the plasma of both species. This similarity in response held for most of the other pathways in which enrichment in a pathway could be determined. But there were exceptions, and we highlight three of these.

**Figure 2.**
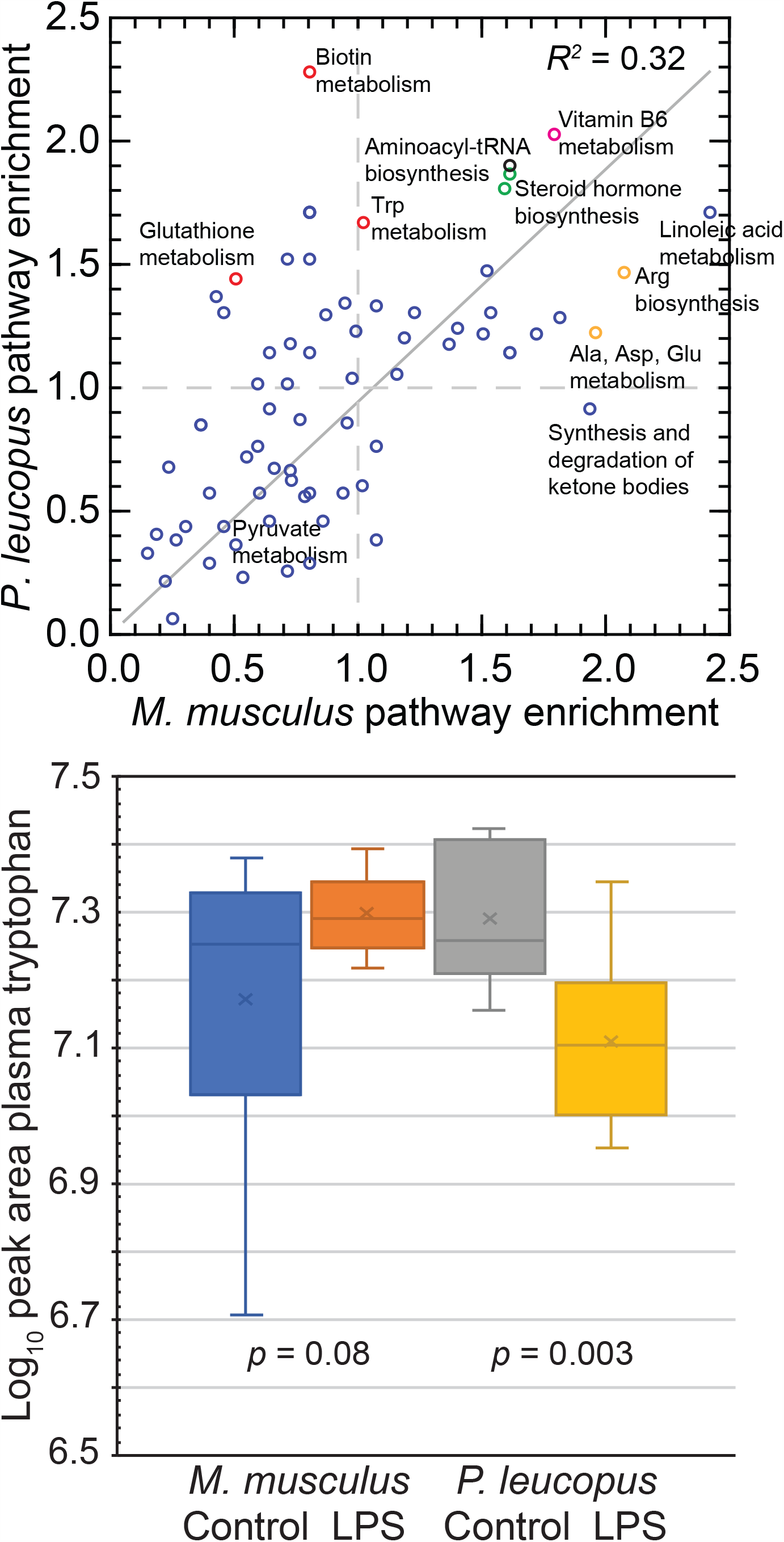
Untargeted metabolomics of plasma of *P. leucopus* and *M. musculus* with or without LPS treatment 4 h prior. Upper panel. Scatter plot of pathway enrichments in LPS-treated *P. leucopus* (*y*-axis) versus those in LPS-treated *M. musculus* (*x*-axis). An enrichment value of 1.0 means no difference in number of genes in a given pathway between treated and untreated conditions for a species. Data are given in Table S2. Selected pathways are labeled. The color of the symbols indicate the following findings for false discovery rate (FDR) *p* values of < 0.05: green, both species; red, *P. leucopus*; and orange, *M. musculus*. The coefficient of determination (*R*^*2*^) shown is for an unspecified intercept. For consistency with the dashed lines indicating enrichment values of 1.0, the regression line for an intercept of 0.0 is shown. Lower panel. Box-plots of log-transformed plasma tryptophan levels estimated as peak areas in LPS-treated and untreated animals of each species. Two-tailed *t*-test *p* values between the two conditions for each species are shown. Data for tryptophan and several its metabolites are given in Table S3.

Of the nine putative constituents asigned to the KEGG vitamin B6 metabolism pathway, eight were significant hits in *P. leucopus* plasma and five in *M. musculus* samples. The three additional metabolites detected in deermice were pyridoxine (C00314), pyridoxamine (C00534), and 2-oxo-3-hydroxy-4-phosphobutanoate (C06054). Pyridoxal (C00847) and pyridoxal 5’-phosphate (C00250) were enriched in both species of LPS-treated animals. The latter two vitamers were reported to have anti-inflammatory effects in an LPS model by reducing expression of IL-1b and suppressing NF-kB activation in macrophages and, moreover, protected mice from lethal endotoxic shock [55]. Among patients with sepsis, higher concentrations in vitamin B6 metabolite biomarkers in the blood were associated with greater likelihood of survival in two clinical research studies [56, 57].

There was overall enrichment of the complex tryptophan metabolism in LPS-treated animals of both species, but the magnitude was greatest among the *P. leucopus*, which had 30 significant hits out of a possible 41 compounds in the tryptophan metabolism pathway (Figure 2A; Table S2). Abundances of specific tryptophan metabolites were determined from peak areas. Tryptophan itself was at a significantly lower abundance in plasma of LPS-treated *P. leucopus* compared untreated animals but at a marginally higher level in *M. musuculus* after LPS treatment compared to controls (Figure 2B, Table S3, Figure S2). Two metabolites of the indole 2,3-dioxygenase (Ido1) pathway, kynurenine and anthranilic acid, were similar in concentration between the two species under both conditions. In the melatonin pathway the concentration of serotonin was significantly lower in treated deermice than in untreated counterparts.

The third pathway, biotin metabolism, demonstrated the greatest magnitude difference between species (Figure 2A), with only biotin (KEGG C00120) itself identified in mice, and three additional metabolites enriched in the plasma of LPS-treated deermice: biotinyl-5’-AMP (C05921), biocytin (C05552), and L-lysine (C00047). Biotin metabolism was reported as one of the pathways, along with vitamin B6 metabolism and tryptophan metabolism, associated with early Lyme disease in a study of urine metabolites of humans [58]. But since biotin is not synthesized by mammals, this difference between the groups may be attributable to differences in food intake during four hours of the experiment.

### Differentially expressed genes in each species

We performed RNA-seq for 3 tissues (blood, spleen, and liver) of each species 4 h after injection of LPS or saline alone to explore the species- and tissue-specific responses. After calculating gene expression levels, we performed differential gene expression analysis using 24,295 and 35,805 annotated transcripts for *P. leucopus* or *M. musculus*, respectively, as reference sets in separate analyses for each species (Tables S4-9). The volcano plots of Figure 3 and Figure S3 show hundreds of genes that were either up- or down-regulated in the blood and two organs of each species by the criteria of ≥4 fold change and FDR of <0.05. With the exception of the spleens of *P. leucopus*, there were more up-regulated than down-regulated genes in each of the comparisons. The ratio of up-to down-regulated genes ranged from ~1x for mouse blood to ~5x for *P. leucopus* blood. The highest total numbers of DEGs were in the livers of each species: 1553 for *P. leucopus* and 3250 for *M. musculus*.

**Figure 3.**
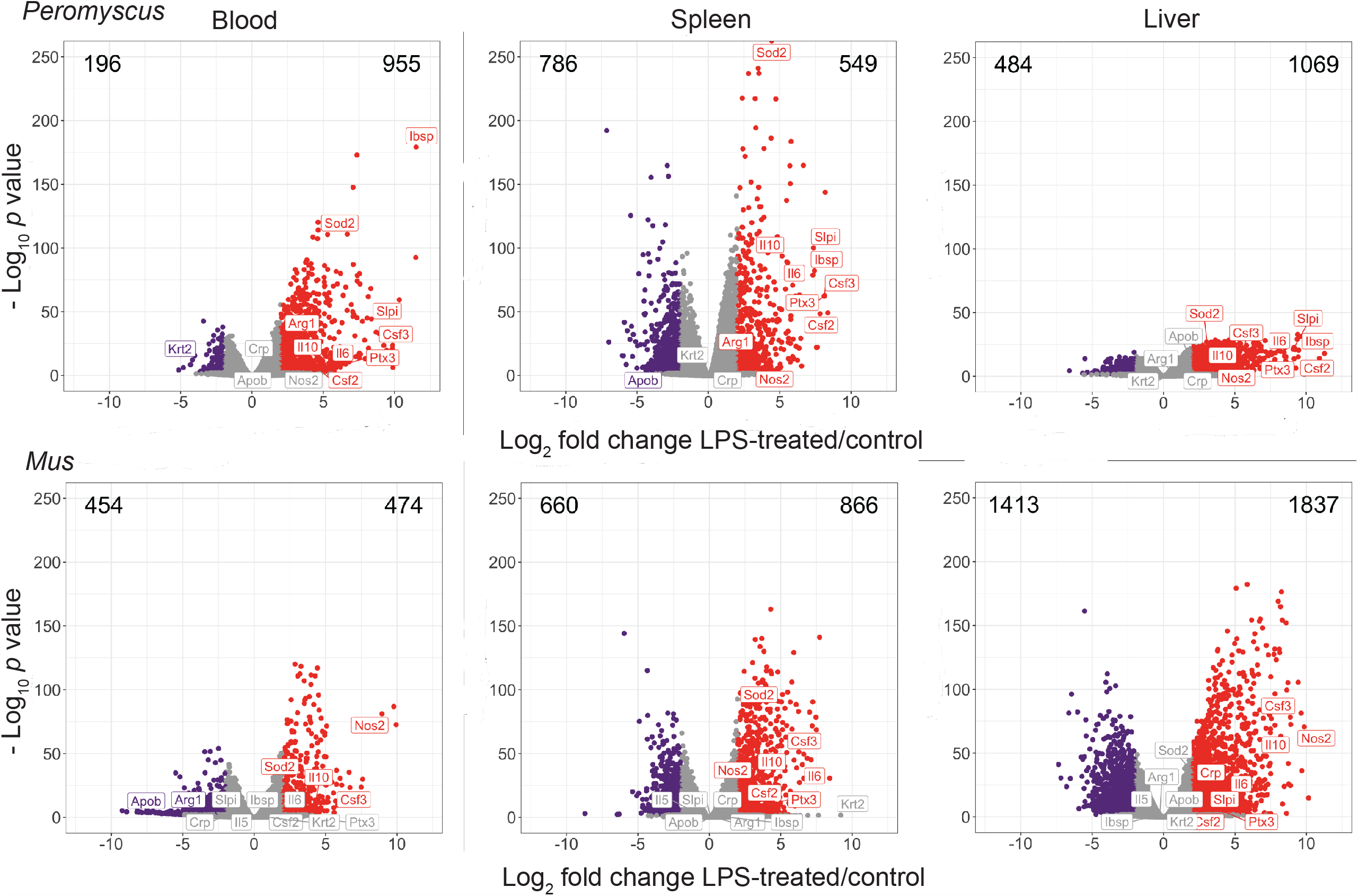
Species- and tissue-specific responses to LPS. Independent differential gene expression analysis of RNA-seq data were performed for blood, spleen, and liver tissues of *P. leucopus* and *M. musculus* collected 4 h after injection with LPS or buffer alsone as control. These are represented as volcano plots with constant scales for the log_2_-transformed fold-changes on *x*-axes and log_10_-transformed FDR *p* values on *y*-axes. Colors of symbols denote the following: red, up-regulated gene with absolute fold-change > 4.0 and *p* value < 0.05; purple, down-regulated gene with absolute fold-change > 4.0 and *p* value < 0.05; and gray, all others. Numbers at the top left and right corners in each plot respresent numbers of down- and upregulated genes, respectively. Figure S2 is same data with scales for *x*- and *y*-axes adjusted for ranges of values in each dataset. Numerical values for each gene in the 6 datasets are provided in Tables S4-S9.

Examples of up-regulated DEGs shared between the two species in their responses to LPS were IL-10 (Il10) and granulocyte colony-stimulating factor (Csf3) in all tissues, IL-6 (Il6), granulocyte-macrophage colony-stimulating factor (Csf2), and pentraxin 3 (Ptx3) in the spleen and liver, and superoxide dismutase 2 (Sod2) in the spleen. A notable difference in the responses of *P. leucopus* and *M. musculus* was consistent with the finding of the nitric oxide assays of the plasma of these animals. Inducible nitric oxide synthase or nitric oxide synthase 2 (Nos2) transcript levels were a mean of 493x higher in the blood of LPS-treated mice than in the controls (FDR *p* = 10^−78^). But in the blood of deermice Nos2 expression was barely detectable (TPM of 0.08 vs. 4.8 in mice), and expression of Nos2 was indistinguishable between the two conditions (FDR *p* = 0.34). In contrast, arginase 1 (Arg1), which by its action reduces the amount of arginine available for Nos2 to produce nitric oxide, was 21x higher in expression in the blood of LPS-treated *P. leucopus* than in controls (FDR *p* = 10^−47^), while in *M. musculus* blood Arg1 expression 4 h after LPS injection was 6x lower than baseline expression (FDR *p* = 0.04). This reciprocal expression profile for Nos2 and Arg1 between the two species was also observed in the spleen. The products of Nos2 and Arg1 are informative biomarkers for categorizing polarized macrophage responses [59]. The disposition of *P. leucopus* to M2 polarization, characterized by high Arg1 and low Nos2 expression in activated macrophages, is consistent with the ability of deermice to moderate inflammation.

Two other genes whose expression profiles served to distinguish the two species in their responses to LPS were integrin-binding sialoprotein (Ibsp) and secretory leukocyte peptidase inhibitor (Slpi). These were first and fourth ranked of the up-regulated DEGs in the blood for *P. leucopus* (Table S4), but they ranked numbers 6232 and 33,541, respectively, among measured transcripts in the blood for *M. musculus* (Table S7). The fold differences between Ibsp and Slpi in the LPS-treated deermice and control animals were 2903x (FDR *p* = 10^−174^) and 1280x (FDR *p* = 10^−57^) higher, respectively. In contrast, Ibsp was lowly expressed in mice under both conditions (TPM of 0.001 and FDR *p* = 1.0), and Slpi in expression was decreased by 3x in the treated animals (FDR *p* = 0.04).

### Sex-specific responses to LPS in *P. leucopus*

For the DEGs highlighted to this point the sex of the animal was not discernibly a major co-variable. To identify sex-specific responses the DEG analysis was applied to the RNA-seq of the blood and spleen of *P. leucopus* treated with LPS. The comparison groups were the 6 females and 6 males, as well as blood for the 4 females and 4 males that were controls. There was little overlap between the 71 sex-specific DEGs in LPS blood, the 56 in LPS spleen, and the 39 in control blood (Table S10). But the few DEGs that were in common between at least two of the groups were notable. In all groups, including the controls, a non-coding RNA (XR_003736827.1) was expressed at 500-1000x higher levels in females than in males in all tissues. This ncRNA was revealed as the Xist or “inactive X specific transcript”, which functions to inactivate genes on the second X chromosome of females and is orthologous to NR_001570.2 of *M. musculus*. D1Pas1, an autosomal DEAD-box RNA helicase, which is expressed in the testis in mice [60], was markedly lower in expression in both treated and control females. Two genes that were more highly expressed by~1000-fold in the blood of females that received LPS over both their male counterparts and control females were the X-linked lysine-specific demethylase 6A (Kdm6a; XM_028854130.1) and adiponectin receptor 2 (Adipor2; XR_003735732.1). Kdm6a has been implicated in risk of autoimmunity and reported to regulate multiple immune response genes [61]. Adiponectin, an adipokine, has an anti-inflammatory effect [62] and expression of its receptor is reportedly affected by macrophage polarization phenotype [63]. Adipor2 was also highly expressed in the spleen of LPS-treated females.

### Functional processes distinguishing and shared between species

To further define similarities and difference between the responses of the two species to LPS we identified gene groups categorized by GO terms as *P. leucopus*-specific, *M. musculus*-specific, or shared, meaning that within each pairing the enrichment groups that were up- or down-regulated. For this analysis 14,685 one-to-one orthologs were used for the comparisons. Figure 4 summarizes this analysis for blood, spleen, and liver. The genes associated with each GO term for which there was significant enrichment are listed in Table S11.

**Figure 4.**
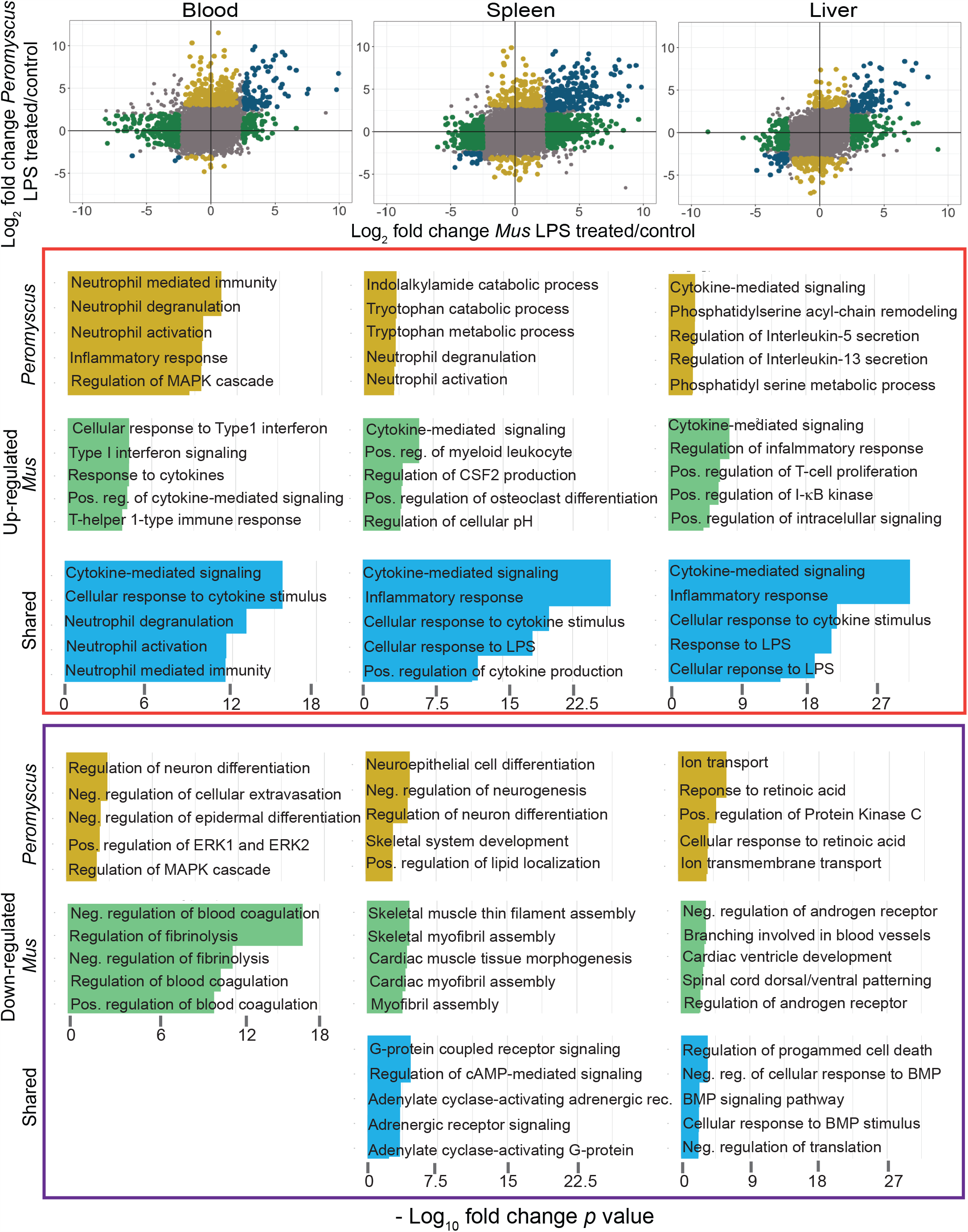
Comparison of *P. leucopus* and *M. musculus* in their responses to LPS by RNA-seq and categorization of DEGs by Gene Ontology (GO) term enrichment. In the three scatter plots for blood, spleen, and liver tissues at the top of the figure log_2_ values for fold change of *P. leucopus* are plotted against corresponding values for *M. musculus* for each gene in the dataset. DEGs specific for *P. leucopus* are indicated by gold symbols, while DEGs specific for *M. musculus* are labeled green. Genes shared between species among the DEGs are labeled in blue. Gray color are for all others. For *P. leucopus*-specific, *M. musculus*-specific, and shared genes up-regulated genes are in upper half, right half, or right upper-quadrant, respectively, of the plot. GO term enrichment was performed for each group of genes, separating up-regulated and down-regulated genes for each one of the tissues, accordingly. The colors in the horizontal bar graphs correspond with the colors indicated above. The genes that constitute each of the listed GO terms in the bottom part of the figure are given in Table S11.

In the blood both species showed up-regulation of some components of cytokine-mediated signaling and cellular responses to cytokines, as well as neutrophil activities. For cytokine-associated genes these included aconitate decarboxylase 1 (Acod1), chemokine C-X-C motif ligand 10 (Cxcl10), Cxcl11, Il1a, Il10, interleukin 1 receptor (Il1r), interleukin 1 receptor antagonist (Il1rn), superoxide dismutase 2 (Sod2), and TNF. Neutrophil-associated genes up-regulated in the blood of both species, including CD14 antigen (Cd14), Cd177, formyl peptide receptor 1 (Fpr1), Fpr2, lipocalin 2 (Lcn2), matrix metallopeptidase 8 (Mmp8), Ptx3, and toll-like receptor 2 (Tlr2). The *Peromyscus*-specific up-regulated profile in the blood featured an expanded set of genes constituting GO terms for neutrophil activities, including arachidonate 5-lipoxygenase (Alox5), Cd33, C-X-C motif receptor 2 (Cxcr2), leucine-rich alpha-2 glycoprotein (Lrg), Mmp9, resistin (Retn), and Slpi. The blood samples of *M. musculus* uniquely featured GO terms related to the following: (1) blood coagulation and fibrinolysis, including apolipoprotein H (Apoh), coagulation factor II (F2), coagulation factor XII (F12), histidine-rich glycoprotein (Hrg), plasminogen (Plg), protein C (Proc), and thrombomodulin (Thbd), and (2) type 1 interferon signalling and cellular responses, including bone marrow stromal cell antigen 2 (Bst2), signal transducer and activator of transcription 2 (Stat2), and XIAP-associated factor 1 (Xaf1).

There were 297 DEGs for the spleen that were shared between *Peromyscus* and *Mus*, a further indication that the deermice were responding in many ways similarly to mice in their early responses to this dose of LPS (Table S11). But there were distinguishing features as well. Up-regulated GO terms for the *P. leucopus* spleen were tryptophan metabolic and catabolic processes, specifically indoleamine 2,3-dioxygenase 1 (Ido1) and Ido2. For *M. musculus* uniquely-associated GO terms that were up-regulated in the spleen dealt with production and differentiation of myeloid cells, macrophages, and osteoclasts, and included the interleukins IL-12b (Il12b) and interleukin 17 members IL-17A (Il17a), Il17f, and IL-23 (Il23a). Down-regulated genes populating GO terms related to assembly of skeletal and cardiac myofibrils included cardiac muscle actin 1 (Actc1), myomesin 2 (Myom2), and Myom3 in *M. musculus*.

In the liver samples four up-regulated GO terms that distinguished the deermice from the mice were phosphatidylserine metabolic process and phosphatidylserine acyl-chain remodeling, which included phospholipase A2, group IIA (Pla2g2a), Pla2g5, and Pla1a, regulation of IL-5 and IL-13 secretion, both of which included the transcription factor GATA binding protein 3 (Gata3), [64], and tumor necrosis factor receptor superfamily, member 21 (Tnfrsf21), which reportedly is a determinant of influenza A virus susceptibility in mice [65]. Two distinguishing GO terms associated with down-regulated genes in the livers of LPS-treated *P. leucopus* were response to retinoic acid and cellular response to retinoic acid, both sets of which included wingless-type MMTV integration site family, member 11 (Wnt11), Wnt5A, and frizzled class receptor 10 (Fzd10), which mediates Wnt signaling in neuron development [66].

A distinguishing GO term for a set of up-regulated genes in the liver of *M. musculus* was positive regulation of I-kB kinase/NF-kB signaling, which included NLR family, apoptosis inhibitory protein 5 or Naip5 (Birc), Ccl19, inhibitor of kappaB kinase epsilon (Ikbke), Il12b, Il1a, LPS-induced TN factor (Litaf), myeloid differentiation primary response gene 88 (Myd88), receptor tyrosine-kinase orphan receptor 1 (Ror1), Tlr2, and tumor necrosis factor (ligand) superfamily, member 10 (Tnfsf10). Distinguishing GO terms for a set of down-regulated genes in the liver of the LPS-treated mice regard the androgen receptor signaling pathway and include the transcription factor forkhead box H1 (Foxh1), hairy/enhancer-of-split related with YRPW motif-like (Heyl), which is associated with repression of transforming growth factor beta (Tgfb) signaling [67], and Nodal (Nodal), which bind various TGFb receptors during development [68].

In summary, this layering of established GO terms over the DEG analysis provided consolidation and further evidence that *P. leucopus* and *M. musculus* have much in common in how they respond in the blood, spleen, and liver in the first few hours after exposure to LPS. But there were also several functional processes in which the species differed. One feature of the *P. leucopus* response that particularly stood out in this analysis was the activation of neutrophils and other phagocytes. Included in the lists of specific genes under all the neutrophil-associated GO terms was Slpi, which was first identified as an inhibitor of serine proteases such as neutrophil elastase [69]. In *M. musculus* the notable distinguishing GO terms concerned blood coagulation and fibrinolysis, processes that are important in the pathophysiology of sepsis [70].

### Gene network patterns across species

To further delineate the gene expression networks without a prior categorization, we performed WGCNA to empirically identify groups of genes (modules) that distinguished specific species and/or treatments in responses to LPS. Gene network analysis allowed identification of patterns across the three tissues. Since blood and the two organs inherently differed in their expression profiles across the genome, and one tissue could overshadow the patterns in other tissues, we made a matrix in which each column represented a sample and each row represented a gene per tissue. Each gene had a suffix to identify the tissue of origin. Twenty-four modules were identified, and GO term enrichment was assessed for unique genes for each module. Figure 5 summarizes the results for four modules: darkorange2, darkseagreen3, brown4, and lightblue. The associated GO terms and constituent genes of the four modules are listed in Table S12.

**Figure 5.**
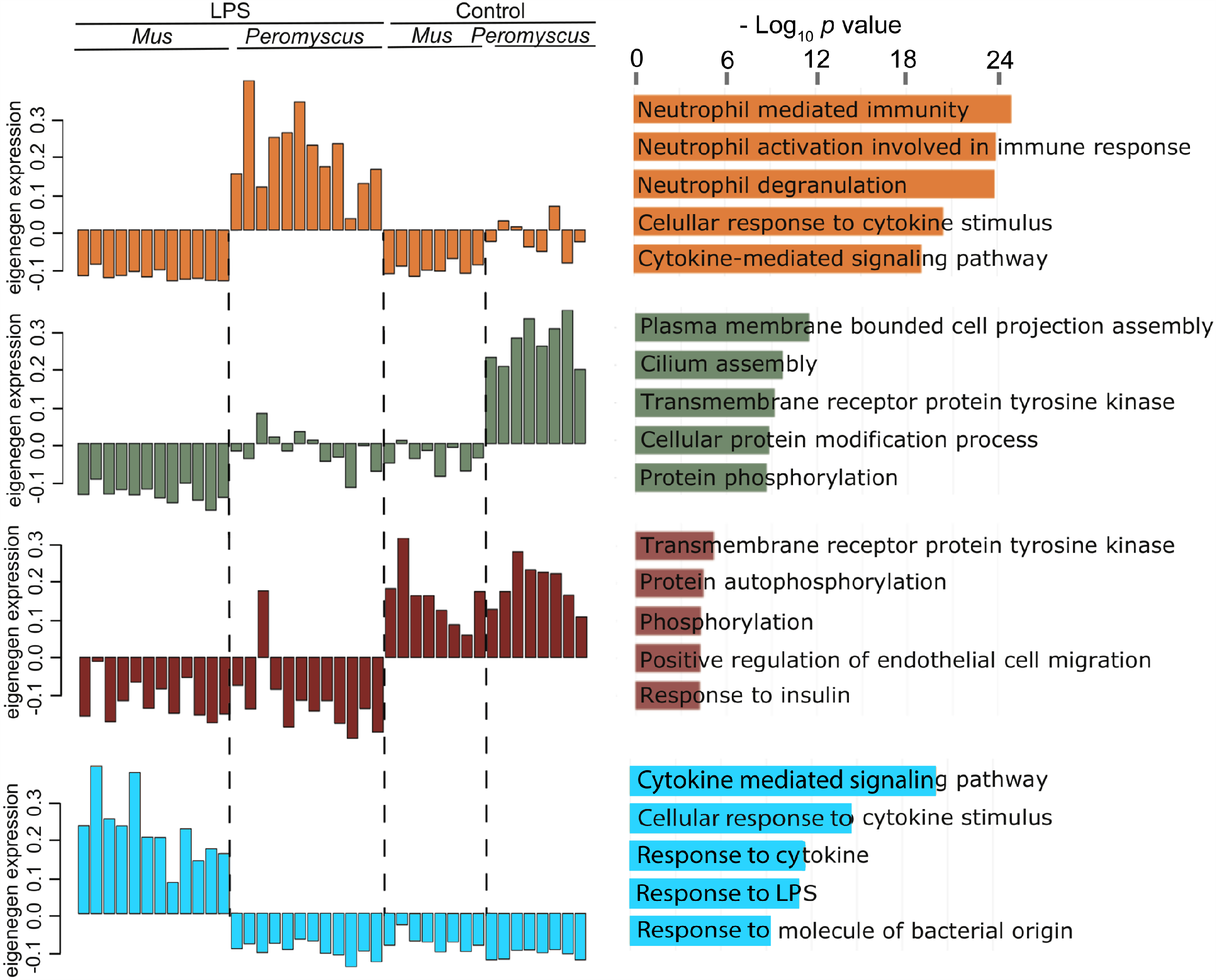
Four selected eigengene modules by network analysis of differential responses of *P. leucopus* or *M. musculus* to LPS. The different modules are distinguished by color of the hexadecimal scheme: darkorange2 for up-regulated in LPS-treated *P. leucopus*, darkseagreen4 for comparatively higher expression in untreated *P. leucopus* than other three groups, brown4 for down-regulated in both species after LPS treatment, and lightblue for up-regulated in LPS-treated *M. musculus*. The top 5 GO terms by adjusted *p* value are shown for each module. The DEGs constituting each of the GO term sets from this analysis are listed in Table S12 along with *p* values and odds ratios. The other 20 modules from this analysis are available at the Dryad dataset repository (https://doi.org/7280/D1B38G).

The darkorange module comprised genes that were up-regulated in LPS-treated *P. leucopus* and not in either control *P. leucopus* or LPS-treated or control *M. musculus*. The blood made the largest contribution with 1472 genes to this module, followed by liver and spleen with 933 and 776, respectively, each. The highest three ranked GO terms related to neutrophils: neutrophil mediated immunity (accession 0002446), neutrophil activation involved in immune response (0002283), and neutrophil degranulation (0043312). The three GO terms share several neutrophil-associated genes that were up-regulated, including those for lysosomal cysteine proteases cathepsin S (Ctss) and cathepsin B (Ctsb), and lysosomal membrane protein 2 (Lamp2). Slpi was also among the contributing genes for each of the three neutrophil GO terms.

The lightblue module comprised genes that were up-regulated in LPS-treated *M. musculus* but not in either control mice or in LPS-treated or control *P. leucopus*. In contrast to the darkorange module, the liver made the largest contribution to the lightblue module with 1170 genes, followed by spleen with 566 genes and blood with 33 genes. The three highest ranked GO terms for this module were cytokine-associated: cytokine-mediated signaling pathway (0019221), cellular response to cytokine stimulus (0071345), and response to cytokine (0034097). While the darkorange module also included two cytokine-associated GO terms at a lower rank than the three neutrophil-associated GO terms, only a minority of genes (41 of 170) in the combined list overlapped between the two species for the cytokine-mediated signaling pathway GO term. Selected up-regulated genes unique to LPS-treated *M. musculus* under this GO term included the following: cytokines Il1a, Il12, Il17, Il22, Il27, Il33, and Ifng; the chemokines Ccl5 (RANTES), Cxcl9, and Cxcl13; macrophage markers Cd86 and Cd80; and transcription factors Jak3 and Stat4. Selected DEGs associated with LPS-treated *P. leucopus* for this GO term included the following: Alox5, annexin 1 (Anxa1), Casp3, leptin (Lep), Sod2, Tgfb1, and tissue inhibitor of metalloproteinases (Timp1); the cytokines Il1b, Il10, Il19, and Interferon 1 beta (Ifnb1); and the transcription factors Gata3, Irf8, Stat1, and Stat3.

The two other modules highlighted in Figure 5 and Table S12 are the darkseagreen module, which distinguished control *P. leucopus* from the other 3 groups, and the brown module, which featured GO terms that were down-regulated in comparison to controls in both sets of LPS-treated animals. For the dark sea green module the spleen was the major contributor with 4791 genes followed by liver with 3189 genes and blood with 73 genes. The five GO terms contained from 241 to 1001 genes. The two with most genes in common were plasma membrane bounded cell projection assembly (0120031) and cilium assembly (0060271). For the brown module the liver RNA-seq with a contribution of 876 genes far out-numbered the contributions of spleen with 6 genes or blood with 3 genes. Genes common to the top three GO terms in this module were all kinases: tyrosine-protein kinase ABL (Abl1), tyrosine-protein kinase CSK (Csk), mitogen-activated protein kinase 3 or ERK1 (Mapk3), and megakaryocyte-associated tyrosine kinase (Matk). Mapk3 is a determinant of macrophage polarization [71].

### Selected genes in blood samples

Results from untargeted investigations led us to examine coding sequences that represented processes that either typified one or the other of the species’ distinguishing responses to LPS or were shared for one or more of the three tissues. Some these genes were identified in the individual DEG analyses summarized in Figure 3. These included several genes, such as for IL-6, Nos2, and Sod2, commonly up-regulated after exposure to LPS or during sepsis in mice and humans. Other genes were chosen for the well-characterized roles in the pathways and other functional processes, including regulatory, that were highlighted for us by the GO term analysis and module-based analysis summarized in Figure 4 and Figure 5, respectively. A few were selected because in *P. leucopus* they were either DEGs that were novel in this context or remarkable for the scale of the changes in expression in this species. An example of the former was the integrin-binding sialoprotein and its Ibsp gene. An example of the latter was the secretory leukocyte peptidase inhibitor and its Slpi gene.

For this cross-species differential expression analysis there was a single reference set that contained the pairs of orthologous protein coding sequences of the mRNAs for both *P. leucopus* and *M. musculus*. Figure 6A shows results for 9 genes of blood samples from 8 *M. musculus* controls (MC), 12 *M. musculus* LPS-treated (ML), 8 *P. leucopus* controls (PC), and 12 *P. leucopus* LPS-treated (PL). A preliminary analysis did not reveal significant differences between females and males of either species for this, as well as for the other two tissues, so the results are not stratified by sex. The deermice and mice were similar in the direction of the changes in expression for Il1rn, Lcn2, and Sod2, but the fold changes were greater in *P. leucopus* for Lcn2 and Sod2 than in *M. musculus* (Table S13). Four other genes were observed to decrease in expression in the blood of mice 4 h after LPS and increase, by several orders of magnitude in some cases, in the deermice: Alox5, which transforms essential fatty acids into leukotrienes, Slpi, which is commonly associated with neutrophil GO terms, transforming growth factor beta (Tgfb), which can be activated when Slpi cleaves an inhibitor [72], and Ibsp, which is associated with known of the GO terms featured in Figure 4 and has been little noted with regard to inflammation. We confirmed the markedly higher expression of Slpi in *P. leucopus* blood after LPS by RT-qPCR (Table 1).

**Table 1.** RT-qPCR of selected transcripts in blood of *P. leucopus* with and without LPS treatment

|  |  |  |  |
| --- | --- | --- | --- |
| mRNA | Control (n = 8) mean<br>copies (95% CI) | LPS (n = 12) mean<br>copies (95% CI) | *p* value |
| Glyceraldehyde 3-phosphate<br>dehydrogenase (Gapdh) | 55,888<br>(32,512-96,070) | 135,970<br>(75,934-243,471) | > 0.05 *,† |
| Nitric oxide synthase 2 (Nos2) | 0<br>(0-0) | 5<br>(1-9) | > 0.05 * |
| Arginase 1 (Arg1) | 310<br>(199-481) | 8779<br>(5185-14,864) | 4 x 10<sup>-8</sup> * |
| Secretory leukocyte peptidase<br>inhibitor (Slpi) | 5 (<br>3-9) | 43,524<br>(29,039-65,235) | 3 x 10<sup>-16</sup> * |
| * 2-tailed *t* test |  |  |  |
| † Kruskal-Wallis test |  |  |  |

**Figure 6.**
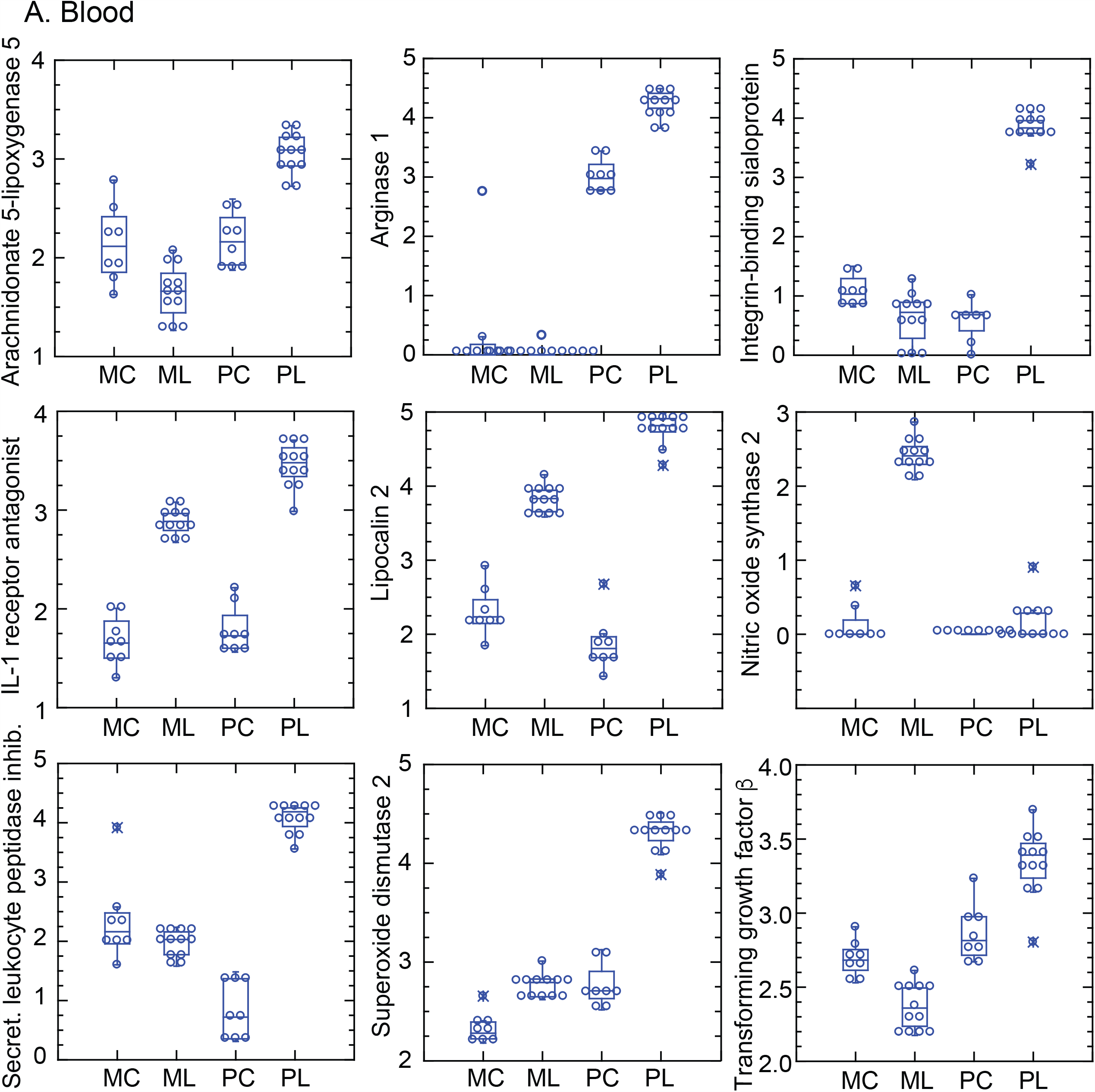
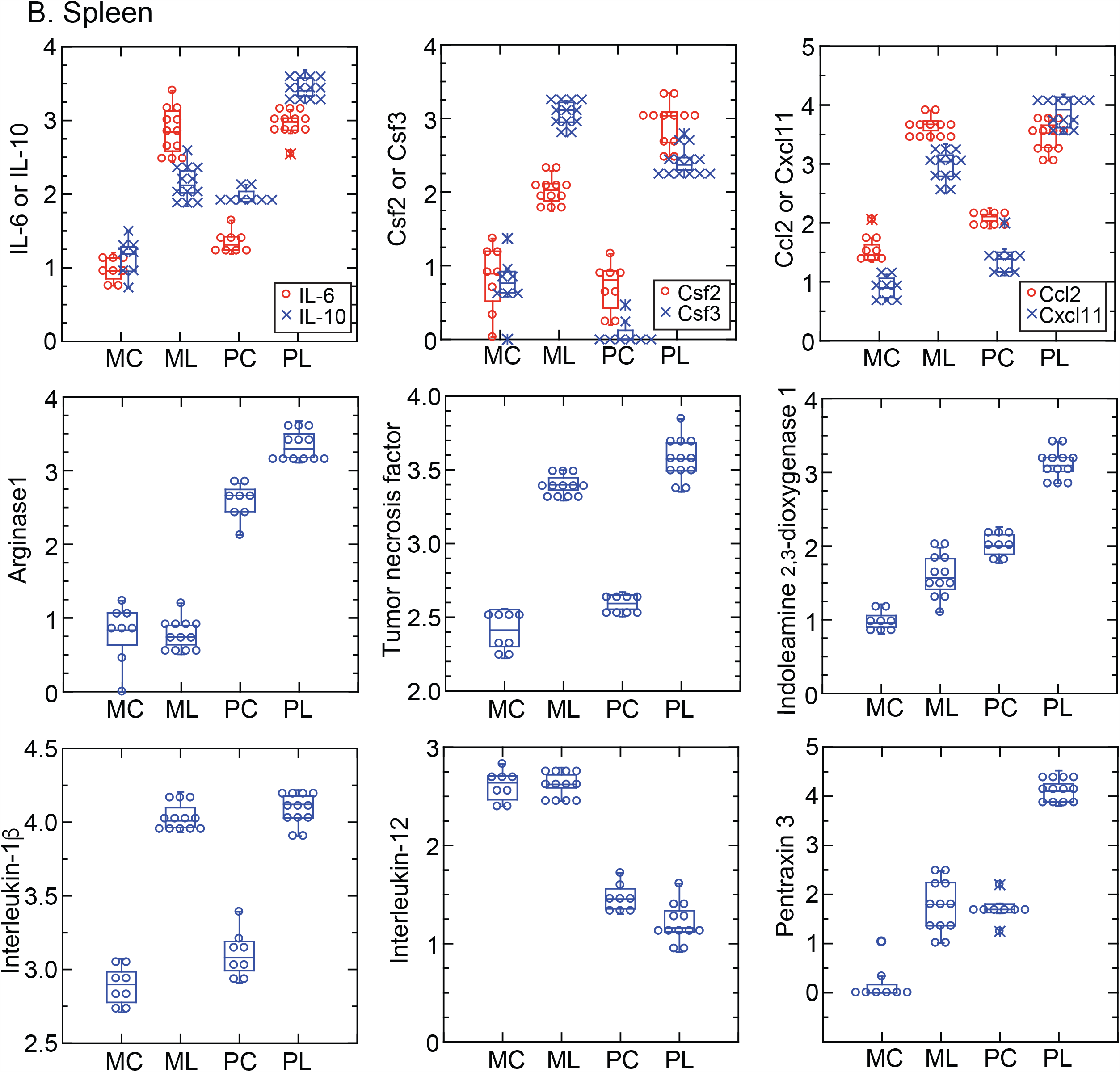
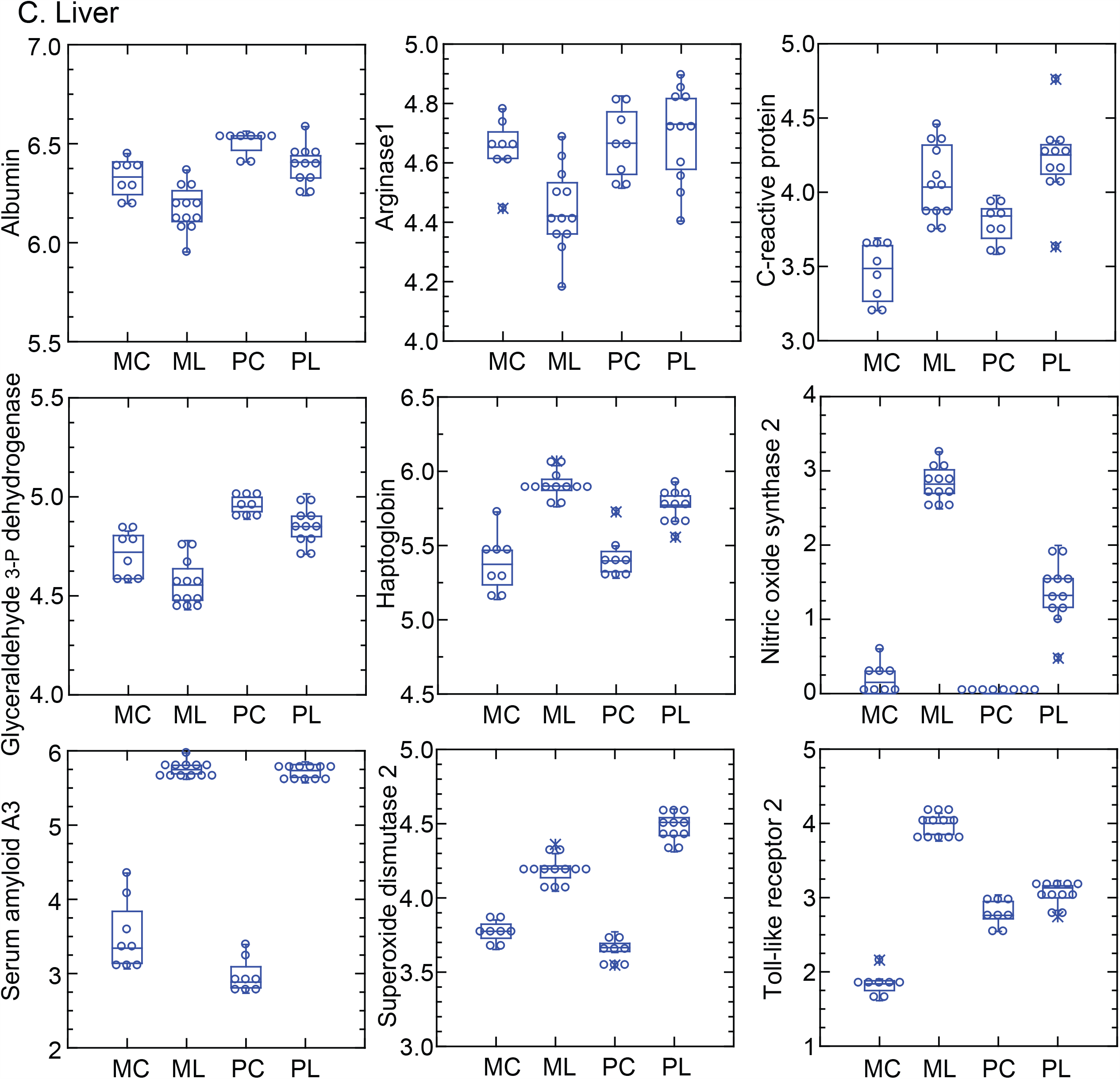
DEGs of RNA-seq with a limited dataset of orthologous protein coding sequences of *P. leucopus* and *M. musculus* for blood, spleen, and liver. The figure comprises panels A, B, and C, each with 9 box-plots for blood, spleen, and liver and for representative genes. The *x*-axes indicate the four different experiment groups: 8 *M. musculus* control (MC), 12 *M. musculus* LPS-treated (ML), 8 *P. leucopus* control (PC), and 12 *P. leucopus* LPS-treated (PL). The *y*-axes are log_10_-transformed, normalized unique reads per coding sequence. In panel B (spleen) two different genes are included in the same graph when their ranges of values across both species and conditions were commensurate. Data values by individual coding sequence are given in Table S13-S15.

The most remarkable difference between the species was in their comparative expression at baseline and after LPS of Nos2 and Arg1, a phenomenon we had also noted in the previous RNA-seq analysis. With the exception of one animal, we detected little expression of Arg1 in the blood of mice either in controls or LPS-treated animals. In contrast, Nos2 expression was many fold higher in *M. musculus* after LPS than in controls. The opposite occurred in *P. leucopus*: expression of Nos2 was undetectable in controls and marginally increased in some treated animals. At the same time Arg1 was already a high level of expression in control animals and was on average ~10-fold higher in those that received LPS. The Nos2 findings in mice and deermice were consistent with our measurements of nitric oxide in the plasma. The RNA-seq results of Nos2 and Arg1 were confirmed by species-specific RT-qPCR assays of the same samples but with different cDNA libraries from those for RNA-seq (Table 1).

### Selected genes in spleen samples

Figure 6B shows box-whisker plots of 9 selected genes for RNA samples from the spleens of the 40 animals. This study confirmed the low expression of Arg1 before and after LPS in *M. musculus* and the high baseline expression of this gene in *P. leucopus* and a further ~10-fold elevation in LPS-treated animals (Table S14). Genes that increased in the spleen in both mice and deermice with LPS exposure, albeit with different degrees of fold-change, were Il6, Il10, Il1b, Csf2, Csf3, CCl2, Cxcl11, and Tnf. These were indications that *P. leucopus* was affected by LPS and in several relevant pathways was responding similarly to *M. musculus*.

There were also increases in both species with LPS of expression of pentraxin 3 (Ptx3), which is a pattern-recognition molecule involved in innate immune responses, and indoleamine 2,3-dioxygenase 1 (Ido1), which is the first and rate-limiting enzyme of tryptophan catabolism through the kynurenine pathway [73]. But the baseline levels in *P. leucopus* were as high as those of LPS-treated *M. musculus* and were even higher in the deermice treated with LPS. In the case of Ptx3, this was more than a hundred-fold. Expression of interleukin-12 (IL-12) was approximately the same in normalized reads in control *P. leucopus* and *M. musculus*. It declined in *P. leucopus* treated with LPS but not in similarly exposed *M. musculus*. The relationship between IL-10 and IL-12 expression in the spleen in individual animals is plotted in Figure S4 of Supplementary Materials.

### Diversity of responses within study populations

To investigate whether the outbred deermouse population was more heterogeneous or diverse in responses than the inbred BALB/c strain mice, we carried out a pairwise analysis of coefficients of determination (*R*^*2*^) within each of the two species (12 animals each) as well as for all 24 animals that received LPS. When self pairings were excluded, this represented 66 pairs each for *P. leucopus* and *M. musculus* and 144 pairs of each of LPS-treated *P. leucopus* paired with each of the LPS-treated *M. musculus*. To control for the wide differences in expression noted between species for some mRNAs, the 17 chosen genes were ones for which the inter-species fold differences were within an order of magnitude and represented those that were more highly expressed in mice as well as those more highly expressed in deermice. The median fold difference between *P. leucopus* and *M. musculus* for these genes was 1.2. We used spleen because of longer reads for this sample and similar numbers of up- and down-regulated DEGs in both species (Figure 3). The genes were the following: Alox5, Ccl2, Csf2, Csf3, Cxcl11, Gapdh, Hmox1, Il1b, Il1rn, Il6, Lcn2, Mt-Co1, Nfe2l2, Slpi, Sod2, Tgfb1, and Tnf. Figure S5 shows the distributions of *R*^*2*^ values of the sets of pairs. As expected correlations were lower between mixed species pairs than intra-species pairs (upper panel); the mean and median *R*^*2*^ were 0.699 and 0.701. The seven highest intra-species pairwise *R*^*2*^ values (0.987-0.994) were observed among the *M. musculus*, and the mean and median of the 66 *R*^*2*^ values were marginally higher at 0.940 and 0.954 for the mice than the 0.937 and 0.948 for the deermice. But, overall, *P. peromyscus* animals in this experiment were not notably more diverse in their responses for this set of genes than were the sample of inbred mice under the same experimental conditions (*p* = 0.68; Mann-Whitney *U p* = 0.19). This was evidence that the DEGs, enrichment GO terms, and modules that distinguished between the two species were likely not accounted for by greater variances or a divergent sub-group among the deermice.

### Selected genes in liver samples

Figure 6C comprises analogous plots of expression of 9 selected genes in the livers of LPS-treated and untreated animals (Table S15). In both species albumin (Alb) and glyceraldehyde 3-phosphate dehydrogenase (Gapdh), a commonly used “housekeeping” gene control for transcription assays, declined about the same extent 4 h after the LPS injection. Sod2 was also elevated in the both sets of treated animals, but the fold-change was greater in the deermice. In contrast to Sod2, Tlr2 increased more than a hundred-fold in LPS-treated *M. musculus* but only marginally from its comparatively higher baseline level in *P. leucopus*.

Other evidence that *P. leucopus* was responding similarly to the endotoxin as mice was the comparable rise in both sets of treated animals in what are considered “acute phase reactants” for evaluating infection and inflammation: C-reactive protein (Crp), haptoglobin (Hp), and a serum amyloid A (Saa) [74]. In liver tissue Arg1 expression was comparatively high in both species among the controls, but, as in the blood and spleen, the levels were elevated in *P. leucopus* and declined in *M. musculus*. Nos2 expression was marginally higher in the livers of LPS-treated deermice than in their blood samples, but this was still many fold lower than observed in the treated mice.

### Pairwise analysis of selected DEGs in both species

Paired relationships between a sub-set of these genes in each of the 3 tissues are depicted in Figure 7. The responses of the two species are largely indistinguishable for Crp, Hp, Alb, and Saa3 among the liver proteins. Both species also displayed correlated responses between Tnf and Il1b and between Il1rn and Il1b, another indication that *P. leucopus* like *M. musculus* “recognizes” the LPS. Up to a certain point events after exposure to LPS in both species appear similar. What distinguished between species was the higher ratio of Il10 to Tnf expression in *P. leucopus* under both conditions, but in particular among the LPS-treated animals. Other pairs of biomarkers that uniquely characterized the responses of *P. leucopus* to endotoxin were Alox5 and Sod2, Ido1 and Ptx3, and, most starkly, Arg1 and Nos2. The profile of high Nos2-low Arg1 expression in response to LPS typifies a type M1 macrophage response in the mouse, while the low Nos2-high Arg1 profile we observed in *P. leucopus* is more characteristic of a type M2 or alternatively-activated macrophage response [59]. Further evidence of a dichotomy between species that corresponded to polarized macrophage categories was the Il10 to Il12 ratio in the spleen [75]. In control *M. musculus* the mean ratio (95% CI) was 0.036 (0.026-0.046). It was ten-fold higher in LPS-treated mice: 0.361 (0.251-0.471). At baseline the Il10 to Il12 ratio was a hundred fold higher in control deermice--3.47 (2.32-4.63)--compared to what was observed in mice. Among the *P. leucopus* treated with LPS the ratio was even higher at 191 (123-259) than the corresponding group of *M. musculus*.

**Figure 7.**
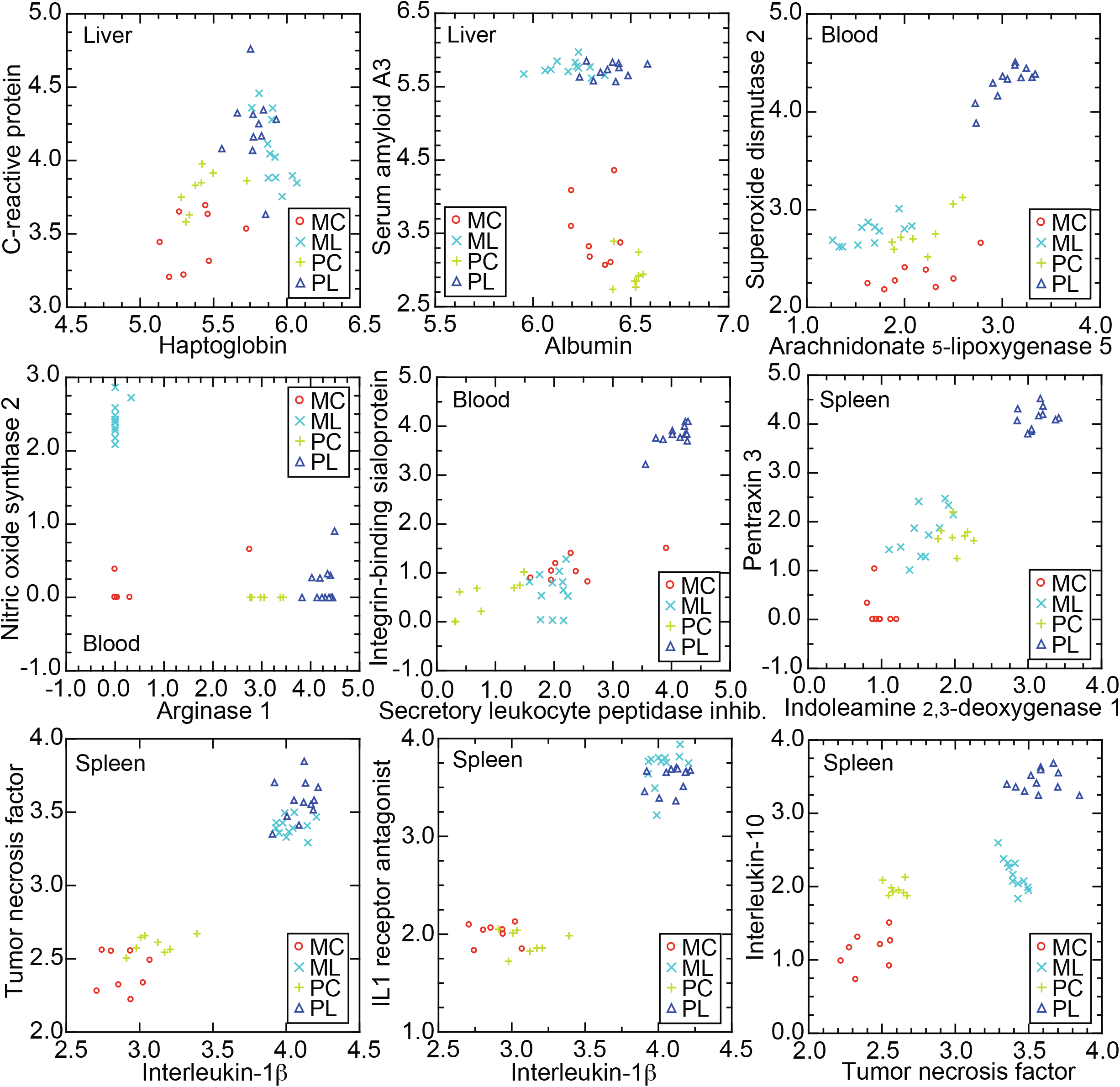
Correlations of pairs of selected genes of *P. leucopus* and *M. musculus* from the RNA-seq of Figure 6. The 9 scatter plots are log_10_ values of normalized unique reads of one coding sequence against another for each of the four groups, as defined in the legend for Figure 6 and indicated by different symbols.

We found similar profiles in blood and spleen samples of older animals in an experiment that was carried out before the deermouse-mouse comparison but under identical experimental conditions, i.e. 10 µg per gm dose or buffer alone as control and samples taken after 4 h. The 16 animals (12 females) had mean and median ages of 79 and 81 weeks, respectively, with a range of 54 to 94 weeks (Table S1). Nine animals received LPS and 7 buffer alone. Figure S6 comprises box-plots of log-transformed normalized unique reads for pairs of genes in either the blood or spleen of these animals. We observed again in this second experiment low baseline and after treatment expression of Nos2 in the blood but high baseline expression of Arg1 with a further increase in the LPS-treated (Table S16). Both Slpi and Ibsp increased more than a hundred-fold in expression in the blood after LPS. In the spleen Il1b, Tnf, Il6, Il10, Ccl2, Cxcl11, Csf2, and Csf3 all were increased in the LPS treated animals, as were two genes that notably more up-regulated in the younger sample of *P. leucopus* than in comparably-aged *M. musculus*: Ido1 and Ptx3.

### Systemic bacterial infection

Five *P. leucopus* were infected with the relapsing fever agent *Borrelia hermsii* on day 0. Another 3 animals received buffer alone. Infection of the blood was directly confirmed by microscopy on day 4. The animals were euthanized on day 5, just before the appearance of neutralizing antibody was anticipated [76]. The infected animals had enlarged spleens, as well as large numbers of bacteria in the spleens, as assessed by quantitative PCR (Table S17). We used previously collected PE100 reads for the blood of these animals [37] for first measuring unique reads mapping to mRNA sequences of selected genes. As we observed in the blood of LPS-treated deermice, Nos2 expression was not or barely detectable in either the controls and infected animals. Arg1 expression was high at baseline and was further elevated at 5 days into infection. Two genes of *P. leucopus* that clearly distinguished this species from *M. musculus* in its short-term response to LPS were Slpi and Ibsp; both of these were about a hundred-fold more highly expressed in infected animals than in controls.

RNA-seq was carried out with the same reference set and settings as for the samples in comparative response study. Given the smaller sample size of the infection study, we focused on more highly expressed genes. Of the 46,154 transcripts in the full reference set, 1773 had mean TPM values of ≥ 50 in at least one of the two groups in each experiment. This latter set was used as the basis for determining and then comparing fold changes between the LPS condition with a duration in hours and the systemic bacterial infection condition with a duration in days (Table S18). Figure 8 summarizes the pair-wise fold-changes for the individual genes. More than two-thirds of the DEGs in the infection experiment were also represented among the DEGs in the LPS experiment among the *P. leucopus*. There was a high correlation in the direction and magnitude of the fold-changes between two experiments. We noted again the substantial up-regulation of Slpi and Ibsp in the shorter term LPS experiment and the longer term infection experiment. Other coding sequences up-regulated under both conditions were Acod1, Alox5, Arg1, Cxcl2, Il1rn, Il1b, Lcn2, and Mmp8. A discordant DEG was Dusp (dual specificity phosphatase; also known as mitogen-activated protein kinase phosphatase or “Mkp-1”), which was up-regulated after LPS but down-regulated in the blood of infected animals. Dusp has been implicated in regulating lipid metabolism during sepsis [77].

**Figure 8.**
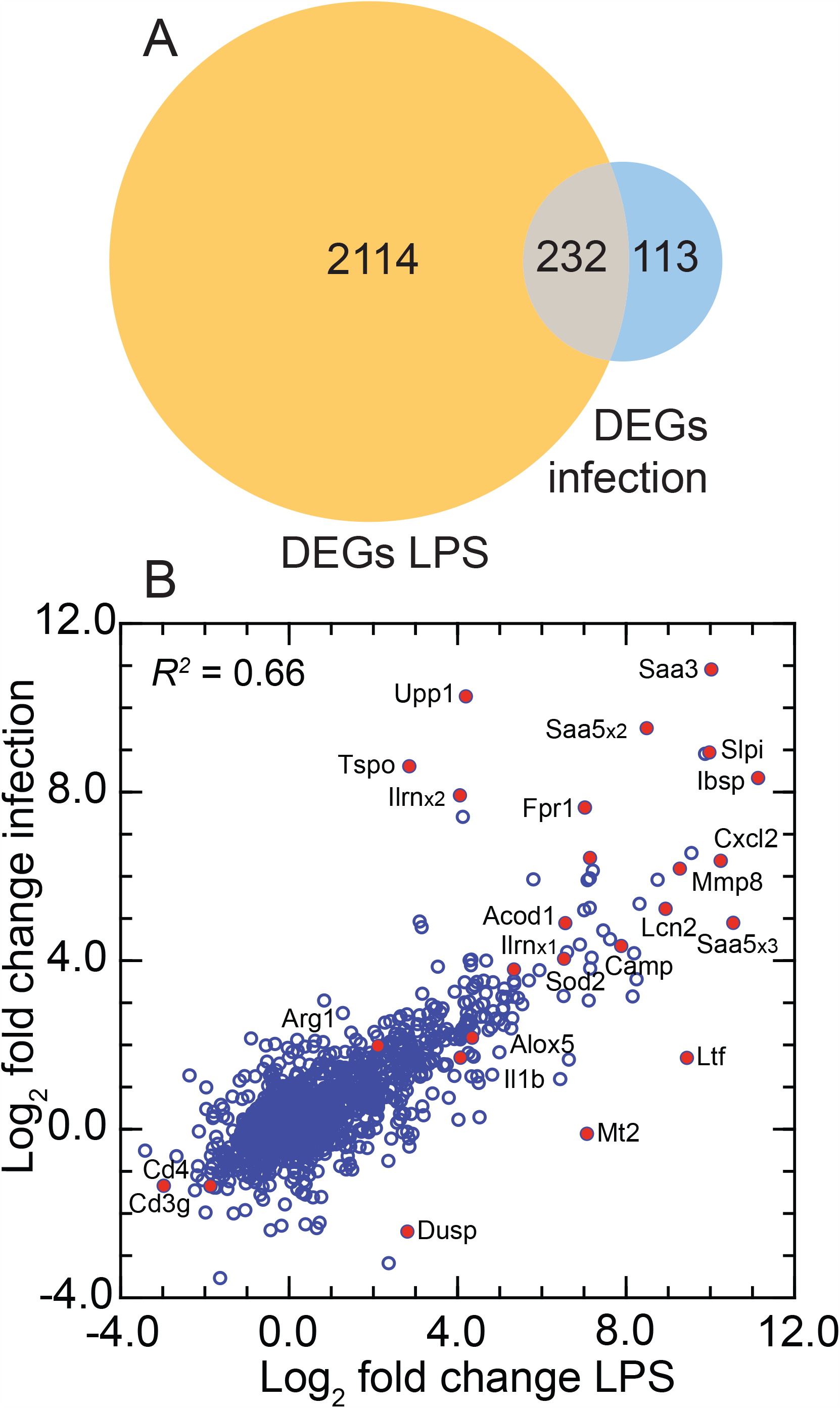
Comparison of DEGs of RNA-seq of *P. leucopus* treated with LPS and *P. leucopus* systemically infected with bacterial agent *Borrelia hermsii*. There were 12 LPS-treated animals with 8 controls, and 5 infected animals with 3 controls (Tables S17 and S18). Panel A. Venn diagram of numbers of DEGs in each experiment and the overlap between them. Panel B. The scatter plot is of log_2_-transformed fold-changes between study and control conditions for infection experiment (*y*-axis) versus short-term LPS experiment (*x*-axis). Selected genes are indicated by label adjacent to a red symbol for the datapoint.

### LPS treatment of *P. leucopus* fibroblasts

Low-passage cultures of fibroblasts isolated from ear tissue from five LL stock deermice were split into pairs, and then one member of each pair was treated with LPS for 4 h and the other member with saline alone. Of the 46,141 transcripts in the reference set, 18,462 had a mean TPM of > 1 in either the control or LPS group, and these were used the DEG analysis (Table S19). For protein coding sequences the 10 highest TPMs in order among control samples were ferritin heavy chain (Fth), Slpi, secreted protein acidic and cysteine rich (Sparc), eukaryotic translation elongation factor 1 alpha (Eef1a1), ferritin light chain (Ftl), vimentin (Vim), serpin family H member 1 (Serpinh1), ribosomal protein lateral stalk subunit P1 (Rplp1), collagen type I alpha 1 chain (Col1a1), and Gapdh. There were 324 genes that were up-regulated by criterion of fold-change ≥ 4 and FDR < 0.05, and 17 genes that were down-regulated, with an additional 80 down-regulated by criterion of fold-change ≥ 2 and FDR < 0.5 (Figure 9). Among those displaying the marked increases in expression between the control and LPS conditions were two subunits of nuclear factor kappa B (Nfkb), two forms of Saa, Nos2, Csf3, Sod2, and phospholipase A2 group IIA (Pla2g2a), which is associated with inflammation during system bacterial infection [78]. Slpi transcripts were not only surprisingly abundant in the fibroblast cultures under usual cultivation conditions, but by RNA-seq and pairwise comparisons expression further increased by 1.5 (1.2-1.9) fold in those pair members exposed to LPS (*p* = 0.02). By the same type of pair analysis with the coding sequence of mRNA as the reference, expression of Nos2 increased 792 (124-5073) fold in the LPS treated fibroblasts (*p* = 10^−5^). This was evidence that the absent or scant expression of inducible nitric oxide synthase we observed in both control and LPS-treated *P. leucopus* animals was not attributable to a genotypic incapacity to transcribe the Nos2 gene.

**Figure 9.**
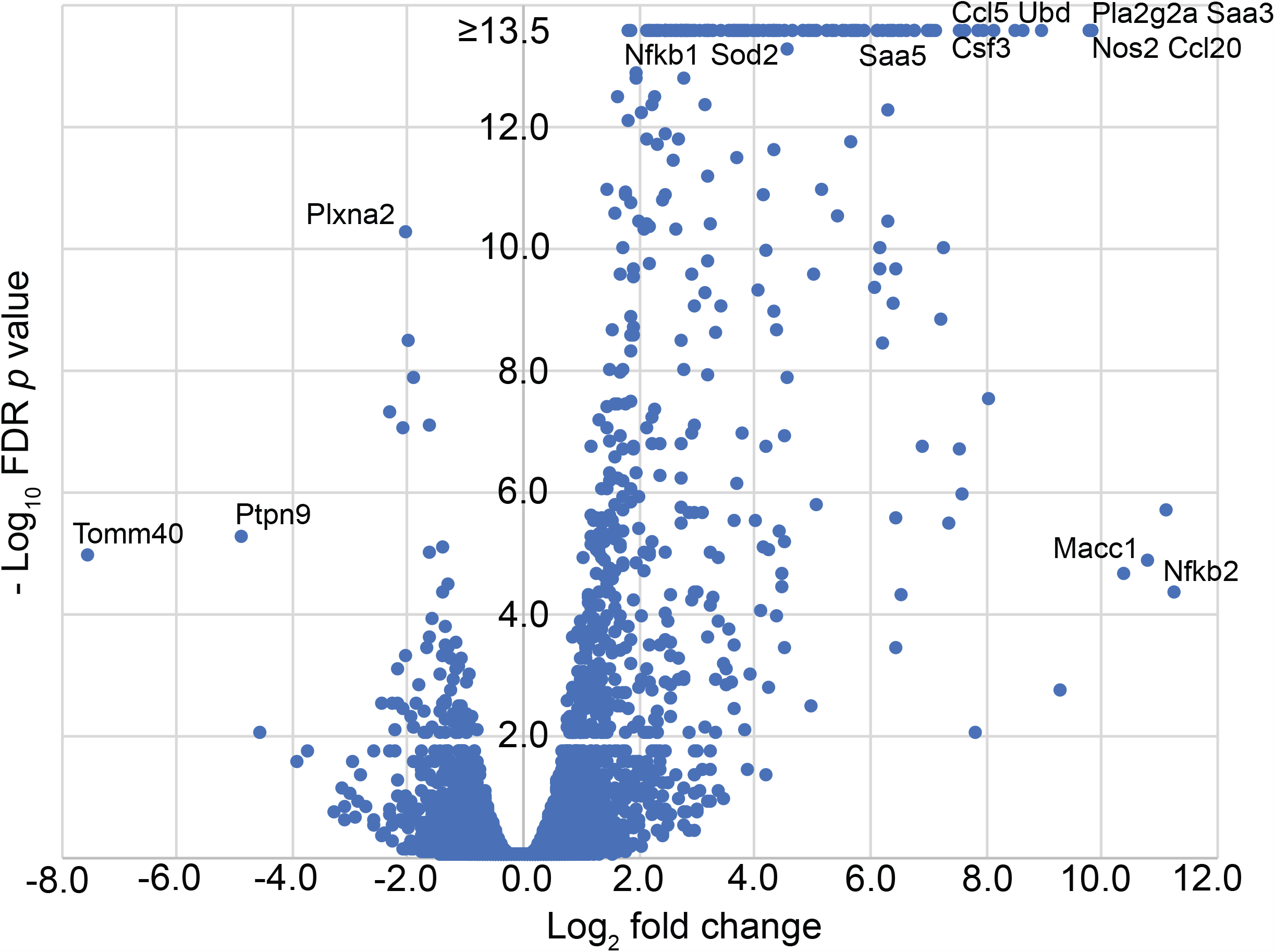
Volcano plot of RNA-seq results for pairs of *P. leucopus* fibroblast cell cultures with or without exposure to LPS. Fold changes for 544 genes (Table S19) are given on *x*-axis and false discovery rate *p* values are given on y-axis. For conciseness the upper limit for the −log_10_ values for this graph was 13.5. The exact or approximate locations of selected differentially-expressed genes are shown.

### Conjunctivitis

After identification of a variety of DEGs under conditions of LPS treatment of whole animals and isolated cells or of systemic infection, we returned to the disease sign we observed in *P. leucopus* but not in *M. musculus*, namely, conjunctivitis severe enough to cause eyelid closure within 2-4 hours of the LPS injection. Six of 12 animals receiving the LPS overtly manifested conjunctivitis during the experiment. Table 2 lists DEGs in the blood or spleen from RNA-seq with absolute fold-changes between animals with conjunctivitis and those without of >2 and FDR values of ≤0.05. Of note in the blood of deermice with conjunctivitis were 21x higher expression of Cxcl13, 12x higher expression of Ccl6, but 46x lower expression of lysozyme (with FDR of < 10^−4^), an antimicrobial enzyme and constituent of tears that coat and protect the conjunctiva [79]. In spleen samples the conjunctivitis animals were distinguished by 6-7 fold higher expression (with FDR <0.005) of two forms of carbonic anhydrase, an enzyme of relevance for the eye disorder glaucoma, because of its role in producing aqueous humor in the eye [80].

**Table 2.** Differentially-expressed genes between 6 *P. leucopus* with and 6 without conjunctivitis after LPS treatment

|  |  |  |  |  |
| --- | --- | --- | --- | --- |
| Specimen | Gene | Fold change | FDR *p*-value | Product |
| Blood | Cxcl13 | 20.7 | 4.0E-02 | C-X-C motif chemokine ligand 13 |
|  | Serpine1 | 17.2 | 4.0E-02 | serpin family E member 1 |
|  | Fn1 | 13.0 | 5.0E-02 | fibronectin 1 |
|  | Cd5l | 12.9 | 5.0E-02 | CD5 molecule like |
|  | Ccl6 | 11.7 | 2.0E-02 | C-C motif chemokine 6 |
|  | Serpinb2 | 9.4 | 4.0E-02 | serine peptidase inhibitor, clade B, member 2 |
|  | Ecm1 | 7.5 | 4.8E-03 | extracellular matrix protein 1 |
|  | Cyb5r3 | 5.5 | 1.3E-05 | NADH-cytochrome b5 reductase 3 |
|  | Ecm1 | 5.2 | 2.0E-02 | extracellular matrix protein 1 |
|  | Lyz2 | -45.8 | 5.6E-05 | lysozyme C-2 |
| Spleen | Ahsp | 14.0 | 4.4E-04 | alpha hemoglobin stabilizing protein |
|  | Slc4a1 | 12.5 | 2.0E-03 | solute carrier family 4 member 1 |
|  | Hemgn | 11.6 | 2.0E-03 | hemogen |
|  | Hmbs | 9.0 | 9.5E-04 | hydroxymethylbilane synthase |
|  | Epb42 | 8.8 | 2.7E-03 | erythrocyte membrane protein band 4.2 |
|  | Tmcc2 | 8.3 | 2.6E-03 | transmembrane & coiled-coil domain family 2 |
|  | Ca2 | 6.7 | 9.0E-04 | carbonic anhydrase 2 |
|  | Gypa | 6.0 | 5.9E-03 | glycophorin A |
|  | Rrm2 | 5.6 | 3.0E-04 | ribonucleotide reductase regulatory sub. M2 |
|  | Ca1 | 5.5 | 2.5E-03 | carbonic anhydrase 1 |
|  | Dhrs11 | 5.5 | 2.7E-03 | dehydrogenase/reductase 11 |
|  | Slc25a37 | 5.2 | 9.7E-03 | solute carrier family 25 member 37 |
|  | Alox15 | 4.9 | 5.0E-02 | arachidonate 15 |
|  | Fech | 4.9 | 3.8E-03 | ferrochelataase |
|  | Cpox | 4.2 | 3.7E-03 | coproporphyrinogen oxidase |
|  | Prxl2a | 4.0 | 4.0E-02 | peroxiredoxin like 2A |
|  | Aqp1 | 3.9 | 1.5E-03 | aquaporin 1 |
|  | Urod | 3.9 | 5.3E-03 | uroporphyrinogen decarboxylase |
|  | Selenbp1 | 3.8 | 2.0E-03 | selenium binding protein 1 |
|  | Ranbp10 | 3.7 | 6.0E-03 | RAN binding protein 10 |
|  | Gstm3 | 3.6 | 4.0E-03 | glutathione S-transferase mu 3 |
|  | Pnpo | 3.5 | 6.9E-03 | pyridoxamine 5'-phosphate oxidase |
|  | Xpo7 | 3.5 | 8.6E-03 | exportin 7 |
|  | Cat | 3.4 | 6.3E-03 | catalase |
|  | Mkrn1 | 3.3 | 5.0E-02 | makorin ring finger protein 1 |
|  | Glrx5 | 2.8 | 3.0E-02 | glutaredoxin 5 |
|  | Rad23a | 2.6 | 1.0E-02 | RAD23 homolog A, nucleotide excision repair |
|  | Tfrc | 2.4 | 4.0E-02 | transferrin receptor |
|  | Hmgb2 | 2.4 | 2.0E-02 | high mobility group box 2 |
|  | Gypcc | 2.2 | 2.0E-02 | glycophorin C |
|  | Prdx2 | 2.0 | 4.0E-02 | peroxiredoxin 2 |
|  | Serpina3n | -2.0 | 7.5E-04 | serine protease inhibitor A3N |
|  | Sirpb1b | -2.2 | 5.0E-02 | signal-regulatory protein beta-1 |
|  | H2-Q10 | -2.3 | 6.2E-03 | H-2 class I histocompatibility, Q10 alpha |

## DISCUSSION

The study yielded abundant information about a small mammal that is emerging as an informative model organism, not only for the study of pathogenesis and immunology of infectious diseases but also for the fields of aging, behavior, ecology, and reproductive biology [33]. The question driving this investigation was how do deermice largely avoid morbidity and mortality when infected with pathogens that otherwise are disabling if not fatal for humans? “Sickness” has various definitions [81], including one operationally based on behavior for animal studies [82]. *P. leucopus* and *M. musculus* became equivalently sick by the behavioral criteria of reduced activity, mutual huddling, and lower food intake. Those *P. leucopus* that had conjunctivitis, which rendered them functionally blind, arguably were even sicker than the *M. musculus*. Nevertheless, under conditions that frequently led to death or the moribund state in various strains of *M. musculus* in several studies, *P. leucopus* receiving the same or even much higher doses of LPS pulled through and recovered.

To address this central question, we mainly depended on RNA-seq of whole blood, spleen, and liver. The study did not specifically examine differences in mRNA isoforms or small non-coding RNAs. These results were complemented by analysis of a limited set of coding sequences, specific RT-qPCR assays, direct detection of the compounds in the blood, and metabolomics. The most extensive and best documented experiment was on the response among several young adult animals of both sexes to a single dose of LPS at approximately the LD_50_ for mice. But we obtained similar results with *P. leucopus* of considerably older age under the same conditions and with animals that had systemic bacterial infection of 5 days’ duration. While we noted some differences between sexes of *P. leucopus* in the experiment, for the most part females and males responded the same to LPS, at least in the short term. The experiment with *P. leucopus* fibroblast cells in culture exposed to LPS showed that the some of the findings, such as high expression of Slpi, are reproduced in vitro and, thus, could be exploited by transgenic and silencing technologies.

Integration of the metabolomics results for the plasma with the transcriptomics results for the three tissues was limited. The sources of some of the metabolites in the blood would likely have been organs, e.g. the adrenal glands, or gut microbiota not subjected to RNA-seq. Nevertheless, the metabolomics confirmed that *P. leucopus* responded similarly to *M. musculus* in many respects at the same dose of LPS. Consequently, the overall milder outcome of LPS challenge in the deermice is probably not attributable to something as pleiotropic as deficiency of the receptor for LPS or the proximate signal transduction step.

One of the striking differences between the two species in their responses to LPS was overall predominance in the GO term and de novo module analyses of neutrophil-associated genes for *P. leucopus* and cytokine-associated genes for *M. musculus* (Figures 4 and 5). The blood was the greatest contributor to the set of genes that distinguished LPS-treated *P. leucopus* from treated and untreated *M. musculus* and from control deermice, while the liver provide the majority of genes that distinguished LPS-treated *M. musculus* from deermice and from control mice. The comparatively heightened neutrophil transcriptional activity in *P. leucopus* paralleled the appearance of conjunctivitis with purulent exudates in the eyes of deermice but not mice. While increased activity of neutrophils could lead to tissue damage from elastase and other proteases and by reactive oxygen species, this may be ameliorated in the deermice by such factors as the leukocyte protease inhibitor Slpi and Sod2, a defense against oxygen radicals.

Slpi is a non-glycosylated 12 kDa cationic, cysteine-rich protein, which was known to be increased in expression in response to LPS and lipoteichoic acids and under the stimulus of TNF and IL-1b (reviewed in [83]). It inhibits neutrophil proteases like cathepsins and elastase, but also prevents degradation of the NF*κ*B inhibitory proteins I*κ*Bα and I*κ*Bβ through its anti-protease activity [84]. Slpi knockout or deficient animlas had impaired wound healing and increased inflammation [85], elevated Nos2 activity in macrophages [86], and increased susceptibility to LPS-induced shock [87].

While LPS-treated *P. leucopus* increased expression of TLR signaling adapter MyD88 and downstream inflammatory cytokines, such as IL-6, TNF, and IL-1β, to approximately the same degree as in *M. musculus*, this was countered by greater expression of cytokines associated with M2 polarization, such as IL-10 and TGFβ, as well as other anti-inflammatory mediators, like Anax1, in the context of lesser expression of IL-12 and IFNγ. The GO term cytokine-mediated signaling pathway served to distinguish both LPS-treated *P. leucopus* and LPS-treated *M. musculus* from their untreated counterparts, but the sets of constituent genes that populated the GO term for each species overlapped for only a minority (Figure 5).

Another notable difference was the contrasting expression of arginase 1 and inducible nitric oxide synthase, or Nos2, in the two species. This can be mapped as well onto macrophage polarization, where type M1 macrophages in a simplified scheme feature high Nos2 and low Arg1 and most varieties of type M2, or alternatively activated macrophages, are characterized by low Nos2 and high Arg1 [59]. The differences between species in their IL-10 to IL-12 ratios before and after LPS [75] was also consistent with a greater disposition of *P. leucopus* to a type M2 response (Figure S4). Further categorization into the M2 sub-types recognized in mice was inconclusive at this point. Nos2 gene knock-out demonstrate reduced susceptiblity to the toxicity of LPS but at the cost of greater susceptibility to *Listeria monocytogenes* or to *Leishmania major* infections [48, 88]. While experimental infections with these particular intracellular pathogens have not been reported for *P. leucopus*, this species does not seem to be particularly at risk of morbidity or mortality from the obligate intracellular bacterial pathogen *Anaplasma granulocytophila* or the Powassan encephalitis virus [89], that exploit *P. leucopus* as a competent reservoir [1].

Integrin-binding sialoprotein (Ibsp) or bone sialoprotein 2 (BSP2) is a special and revelatory case in this study. There were marked increases in the expression of this gene in *P. leucopus* but not in *M. musculus* upon exposure to LPS. The Ibsp protein has predominantly been associated with bone and tooth morphogenesis with no reported association with inflammation or innate immunity. Consequently, it would not be expected to be included in many of the innate immunity and inflammation pathways and GO terms that distinguished LPS-treated from untreated animals of either or both species. But there is justification for viewing Ibsp as *P. leucopus*’ functional substitute of osteopontin (Spp1), another small integrin-binding glycoprotein that is heavily modified post-translationally [90]. Both BSP2 and osteopontin (BSP1) bind to alpha v beta 3 integrins, and their genes are within a hundred kilobases of each other on chromosome 4 of humans and chromosome 22 of *P. leucopus* [36]. Osteopontin is a biomarker of sepsis in humans [91] and reportedly acts through Stat1 degradation to inhibit Nos2 transcription [92].

Could observed differences between the two species in these experiments be attributable in part to differences in their microbiomes? The gut metagenomes were determined from pre-experiment fecal pellets from the animals in this study, and the results have been reported [38]. In general, the deermice and mice had similar distributions and frequencies of bacteria at the taxonomic level of family and in the representation and proportions of different biosynthetic, metabolic, catabolic, and regulatory functions. But there were also substantive differences in the gut microbiota between the species that plausibly could account for some of the distinguishing responses to LPS. The first was the abundant presence in the intestine in *P. leucopus*, but not in *M. musculus*, of a protozoan, identified as a new species of the parabasalid genus *Tritrichomonas*. The presence of *Tritrichomonas muris* in some populations of the same inbred strains of *M. musculus* altered their cellular immune responses to other microorganisms and antigens [93]. Further distinguishing the gut microbiota was the greater abundance and diversity in *P. leucopus* of *Lactobacillus* spp., the primary niche for which was the forestomach [38]. In two studies of the use of *Lactobacillus* spp. as probiotics, feeding of mice with *L. paracasei* or *L. plantarum* resulted in less inflammation in comparision to controls when challenged with an influenza virus or *Klebsiella pnemonia*, respectively [94, 95]. In the present study a tryptophan metabolite that distinguished the plasma samples of the two species was indolepropionic acid (Table S3), which at baseline was three-fold higher in concentration in deermice than in mice but was nine-fold lower concentration in LPS-treated *P. leucopus*. It was only marginally lower between the corresponding sets of *M. musculus*. This compound is the metabolic product of certain intestinal microbes and reportedly has anti-inflammatory and anti-oxidant properties [96, 97].

In summary, *P. leucopus* differed in many respects from *M. musculus* in these experiments in its response to LPS. Under conditions in which about half the mice would be expected to die within 48 h, the deermice survived the acute insult and recovered, a phenomenon that modeled the tolerance of these animals to infection. In their outcomes after LPS exposure the naïve *P. leucopus* resembled *M. musculus* that had become accustomed to LPS by prior exposure to a low dose of LPS [98]. A characteristic profile of these pre-treated mice is comparatively lesser expression of proinflammatory mediators and greater expression of genes of phagoctyes [99], similarly to what we observed in naïve *P. leucopus*. The phenotype of Slpi knockout mice resembles that of animals that have been tolerized to LPS in their suppression by administered IFNγ [100], a cytokine that is an alternative to LPS as an inducer of Nos2 expression in macrophages [101].

Whether this points to a single pathway or even a single gene remains to be determined. But we doubt that the phenomenon of infection tolerance can be reduced to a simple explanation. The evidence rather is of multiple adaptive traits representing different aspects of innate immunity, metabolism, oxidative stress management, and perhaps the microbiome that serve to sustain *P. leucopus* populations amidst the varied infectious agents they face. This is also to the benefit of these agents in this trade-off, because it renders *P. leucopus* a competent vertebrate reservoir for them and one of impact for human populations in endemic areas.

## METHODS

### Animals

Adult outbred *P. leucopus* of the LL stock were obtained from the Peromyscus Genetic Stock Center (PGSC) of the University of South Carolina [102]. The LL stock colony was founded with 38 animals captured near Linville, NC between 1982 and 1985 and has been closed since 1985. Sib-matings are avoided, and complete pedigree records are kept. Animals of the LL stock closed colony had mitochondria of the same genome sequence with little or no heteroplasmy [37]. Adult BALB/cAnNCrl (BALB/c) and C.B.17 strain Severe Combined Immundeficiency (SCID) *M. musculus* were purchased from Charles River. All animals in the LPS experiments spent at least two weeks at the UCI facilities before the experiment.

Animals were maintained in the AAALAC-accredited U.C. Irvine vivarium with 2-5 animals per cage according to sex and on 12 hours light-12 hours dark lighting schedule, temperature of 21-23° C, humidity of 30-70%, water *ad libitum*, and a diet of 8604 Teklad Rodent (Harlan Laboratories). For prior to injections animals were lightly anesthetized with 2.5% isoflurane in presence of 2 liters/min oxygen. The rodents were euthanized by carbon dioxide overdose and intracardiac exsanguination at the termination of the experiment or if they were moribund, unable to eat or drink, or otherwise distressed. Dissection was carried out immediately. Instruments were cleaned first and then sterilized between dissections.

The study was carried out in accordance with the recommendations in the Guide for the Care and Use of Laboratory Animals of the National Institutes of Health. University of California Irvine protocol AUP-18-020 was approved by the Institutional Animal Care and Use Committee (IACUC). The protocol for the comparative study of *P. leucopus* and *M. musculus* for responses to LPS after 4 h was in addition approved by the Animal Care and Use Review Office of the United States Army Medical Research and Materiel Command. P. leucopus studied at the PGSC were under IACUC-approved protocol 2349-101211-041917 of the University of South Carolina.

Tables S1 and S21 of Supplementary Materials respectively provide information on each of the animals in the LPS experiments and corresponding National Center for Biotechnology Information (NCBI) (http://ncbi.nlm.nih.gov) BioProject and BioSample identifying numbers and descriptions for these samples. The gut metagenomes from feces collected from the 20 *P. leucopus* and 20 *M. musculus* animals 1-2 days before the comparative experiment have been described [38] and are available from MG-RAST database (https://www.mg-rast.org) under accession numbers mgm4832531.3-mgm4832578.

### LPS susceptibility and dose responses

Thirty adult *P. leucopus*, divided into five groups of three females and three males, were each injected intraperitoneally (i.p.) on day 0 with a 50 µl volume of *Escherichia coli* O111:B4 LPS purified by ion exchange and with <1% protein and <1% RNA (Sigma Aldrich; catalog L3024), which was diluted in sterile, endotoxin-free 0.9% saline (Sigma Aldrich) to achieve the following doses in µg/gm (mg/kg) body weight: 10, 50, 100, 200, and 300. The doses were administered in randomized order over the period 1400-1700 h of a single day. Animals were returned to their cages with ad libitum food and water and then monitored every 12 h for the following signs: reduced activity by criterion of huddling with little or movement for > 5 min, ruffled fur, hyperpnea or rapid respiration rate, and conjunctivitis by the criterion of closed eyes with crusting observable on the eyelids. The primary endpoint was death during the period between observations or the moribund state (immobility, rapid respiration, and not able to feed or drink) at the scheduled monitoring time or when notified in the interim by vivarium attendants.

### LPS effects at 4 h

Animals were anesthetized with isoflurane and injected i.p. with a single dose of E. coli O111:B4 LPS at a concentration of 10 µg/gm body weight in a 50 µl volume as described above. The control group was anesthetized and then injected with the 0.9% saline alone. The experiment started at 0800 h with 10 min intervals between animals and with alteration of LPS and control injections. At 4.0 hr after their injection the animals were euthanized. After opening the chest, exsanguination was performed by cardiac puncture and blood was transferred to a heparin sulfate coated tubes (Becton-Dickinson Microtainer). Anticoagulated blood was centrifuged to pellet blood cells for 3 min at 4600 x g at 4° C. Plasma and blood pellet was kept separately at −80° C until further analysis. Liver and spleen were extracted, flash-frozen in liquid nitrogen, and stored at −80 °C until RNA extraction.

### Experimental infection

Infection of a group of adult *P. leucopus* LL stock with the relapsing fever agent *Borrelia hermsii* strain MTW of genomic group II was described by Barbour et al. [37]. *Peromyscus* species are natural hosts of genomic group II strains of *B. hermsii* [103]. In brief, animals were anesthetized and then injected on day 0 with 10^3^ bacteria by each of i.p. and subcutaneous routes in 50 µl volumes of PBS and diluted plasma from infected SCID mice, as described [104]. On day 4 a drop of tail vein blood was mixed with PBS an examined as a wet mount by phase microscopy to confirm infection. On day 5 animals were euthanized with carbon dioxide and terminal exsanguination. Whole blood was dispensed into heparin-coated tubes, and the spleens were removed by dissection, weighed, and then flash frozen in liquid nitrogen. Confirmation of infection and quantitation of bacterial burdens in the spleens was carried by quantitative PCR, as described [105].

### Fibroblast cell culture and LPS treatment

Fresh ear punches were collected from five LL stock *P. leucopus* animals (2 females and 3 males) during routine marking procedures at the time of weaning at ~3 weeks of age. The PGSC identification numbers (and mating pair for each animal) were 22608 (H-1075), 22609 (H-1075), 22610 (H-1121), 22611 (H-1121), and 22614 (H-1127). After the ear punch tissue was bathed in 70% ethanol for 2 min, it was placed in RPMI 1640 medium supplemented with 10% fetal bovine serum (HyClone FetalClone II; Thermo Scientific), minced, and then treated with 5 mg/ml collagenase type I (Millipore) for 1 h. After undigested debris was removed, the disassociated cells were cultivated in the same medium at 37° C and in 5% CO2. Cells were passed when adherent layers reached 45-90% confluency and for no more than 7 passages. For the experiment individual cultures were split into pairs and they were incubated at initial concentrations of 3 × 10^5^ cells per well for 24 h. LPS or saline alone was added medium for final LPS concentration of 1 µg/ml and then the incubation was continued for 4 h. After disassociation of the fibroblast layer with trypsin and then addition of RNAlater (Thermo Scientific), the cells were harvested and stored at a concentration of ~10^6^/ml in −80 °Cuntil RNA extraction.

### Analyses of blood

Nitric oxide was analyzed from frozen plasma samples using Nitric Oxide Assay Kit (Invitrogen) according to manufacturer’s instructions and nitrite concertation was calculated based on standard curve measuring optical density at 540 nm on a plate reader (BioTek Synergy2). Corticosterone was measured colorimetrically in plasma using DetectX Corticosterone Enzyme Immunoassay Kit K014-H1 (Arbor Assays) based on corticosterone-peroxidase conjugate. After incubation, the reaction is read at 450nm on a BioTek Synergy2 plate reader.

### Metabolomics

Untargeted detection and analysis of metabolites in plasma of *P. leucopus* and *M. musculus* were carried out essentially as described [106]. In brief, 40 µl volumes of plasma, which had been stored frozen at −80° C, were extracted with 120 µl methanol containing as internal standards phenylalanine d5 (175 ng/mL), 1-methyl tryptophan (37.5 ng/mL), and arachnidonoyl amide (30 ng/mL). After precipitated proteins were removed by centrifugation, the supernatant was dried under vacuum and then suspended in 50% methanol. Aliquots of 10 µl were subjected to high pressure liquid chromatography (HPLC) and quadrupole (Q) time-of-flight (TOF) mass spectrometry (MS) with periodic inclusion of pooled samples for quality control. Metabolites were separated on an Agilent Technologies Poroshell C8 column (100 x 2.1 mm, 2.7 µm) with a gradient of acetonitrile and water both containing 0.1% formic acid with an Agilent 1260 HPLC pump under conditions previously described [106]. The eluent was introduced into an Agilent Technologies 6520 Q-TOF-MS instrument equipped with an electrospray ionization source. The parameters for the analysis were a capillary voltage of 4000 V, fragmenter voltage of 120 V, gas at 310° C, gas flow of 10 L/min, and nebulizer pressure of 45 psig. Data were acquired in the positive ion mode at a scan range of 75-1700 for the mass-to-charge ratio (*m/z*) and at a rate of 1.67 spectra per second. The raw data files were converted to the XML-based mzML format with Proteowizard [107]. The files were then processed for peak picking, grouping, and retention time correction with the XCMS suite of software [108]. The *centWave* algorithm was applied to detect chromatographic peaks with these parameter settings: ≤ 30 ppm for *m/z* deviation in consecutive peaks, signal to noise ratio of 10, a prefilter of 3 scans with peak intensity of ≥ 750, and 10 to 45 s for peak width. Molecular features (MF), defined by *m/z* and retention time, were grouped across samples using a bandwidth of 15 and overlapping *m/z* slice of 0.02. Retention time correction was performed with the *retcor* algorithm of XCMS. Finally, peak area was normalized by median fold normalization method as previously described [109]. Pathway enrichment analysis was performed with *Mummichog* network analysis software v. 2 [110], which predicts functional activity from spectral feature tables without a priori identification of metabolites. This was implemented on-line at Metaboanalyst (http://www.metaboanalyst.ca). The parameter settings were mass accuracy of 20 ppm, positive ion mode, and cut-off *p* value of 0.05 for the Fisher’s exact test of observed and expected hits of a given pathway. Pathway enrichment in Kyoto Encyclopedia of Genes and Genomes (KEGG; https://www.genome.jp/kegg/) terms was determined separately for the *P. leucopus* and *M. musculus* sets of LPS-treated and control animals. A cross-species comparison of enrichments for pathways in common was plotted. Identifiable metabolites constituting the pathways that distinguished between LPS-treated and control animals were listed for each species. As a measure of abundance, the peak areas of selected metabolites were extracted from raw data set using Skyline software [111]. Molecular features data for the 40 individual plasma samples was deposited with the Dryad data repository (https://doi.org/10.7280/D12M4N).

### RNA extractions

RNA from blood was extracted using NucleoSpin RNA Blood Mini kit (Macherey-Nagel). Lysis buffer and proteinase K were added to the frozen pellet and shaken while thawing. RNA from liver and spleen was extracted from 30 mg of frozen tissues, mechanically homogenized and further lysed in RLT buffer with 2-mercaptoethanol on TissueLyser (Qiagen) with 3 mm beads then extracted according to the protocol using RNeasy Mini Kit (Qiagen).

Extraction of total RNA from blood of *P. leucopus* infected with *B. burgdorferi* was described by Long et al. [36]. For extraction of total RNA from freshly obtained and chilled anticoagulated blood was mixed with an equal volume of Buffer EL of the QIAamp RNA Blood Mini Kit (Qiagen) and then processed with the kit and Qiagen QIAshredder spin columns according to the manufacturer’s protocol. For extraction of RNA from frozen liver the Qiagen TissueLyser instrument with 3 mm stainless steel beads and the RNeasy Mini Kit (Qiagen) was used. Nucleic acid concentrations were determined with a Qubit fluorometer (ThermoFisher Scientific). The quality of the extracted RNA assessed by Agilent 2100 BioAnalyser with the Nano RNA chip. The RNA was stored in RNase-free distilled water at −80° C.

### Reverse transcriptase quantitative PCR

RT-qPCR assays were developed and implemented for measurement of transcripts of genes for the following proteins of *P. leucopus*: nitric oxide synthase 2 (Nos2), arginase 1 (Arg1), secretory leukocyte peptidase inhibitor (Slpi), and glyceraldehyde 3-phosphate dehydrogenase (Gapdh). cDNA was synthesized from extracted RNA using iScript Reverse Transcription kit and iScript Supermix (BioRad) for qPCR, according to the manufacturer’s instructions. The reaction mix was incubated in a thermal cycler for 5 min at 25^°^ C, 20 min at 46^°^ C, and 1 min of 95^°^ C. PCR of cDNA was carried out with qPCR PowerUp™ SYBR™ Green Master Mix (Applied Biosystems) and performed in 96-well plates in a StepOnePlus Real-Time PCR System (Applied Biosystems) instrument. The forward and reverse primers, synthesized by Integrated DNA Technologies (San Diego, CA), were, respectively, the following: Arg1, 5’-TCCGCTGACAACCAACTCTG and 5’-GACAGGTGTGCCAGTAGATG; Nos2, 5’-GACTGGATTTGGCTGGTCCC and 5’-GAACACCACTTTCACCAAGAC; Slpi, 5’-TCCCATCAGCAGACCAGTG and 5’-TTGGGAGGATTCAGCATCATACA; and Gapdh, 5’-TCACCACCATGGAGAAGGC and 5’-GCTAAGCAGTTGGTGGTGCA. The product sizes were 352, 192, 81, and 169 bp, respectively. For all assays the initial step was 95° C for 10 min followed by 40 cycles. The cycle conditions for Arg1 and Nos2 were 95^°^ C for 15 s, 52^°^ C for 30 s, and 72^°^ C for 60 s. For Slpi and Gapdh they were 95^°^ C for 15 s and 52^°^ C for 30 s. Standards were the corresponding PCR products cloned into the *E. coli* plasmid vector pUC57 and stored frozen at −80° C in single-use aliquots after plasmid purification, as described.

### RNA-seq

Library preparation with the Illumina TruSeq mRNA stranded kit was carried out as described [36]. The libraries were normalized and then multiplexed to achieve 12 samples per flow cell on an Illumina HiSeq 4000 instrument and 100 or 150 cycles of paired-end read chemistry at the U.C. Irvine Genomic High Throughput Facility. The quality of sequencing reads was analyzed using FastQC (Babraham Bioinformatics). The reads were trimmed of low-quality reads (Phred score of <15) and adapter sequences, and corrected for poor-quality bases using Trimmomatic [112]. Paired-end reads in fastq files were quantified using *kallisto* v. 0.46.1 ([113] using the GENCODE annotation v. 21 (https://www.gencodegenes.org) for mouse and GCF_004664715.1_Pero_o.1 _rna from NCBI for *P. leucopus* [36]. The mean (95% confidence interval) coefficients of determination (*R*^*2*^) between paired replicates of 4 RNA extracts (blood of 4 LPS-treated *P. leucopus*), but with independent cDNA libraries and sequencing lanes was 0.994 (0.990-0.997) for 8055 genes with mean transcripts-per-million (TPM) >1, while the mean *R*^*2*^ for the 24 discordant pairs for these samples was 0.975 (0.973-0.977). Sequencing reads were deposited with Sequence Read Archive (Table 3).

**Table 3.** BioProjects, BioSamples, and Sequence Read Archive accession numbers for RNA-seq

|  |  |  |  |
| --- | --- | --- | --- |
| Subject | BioProjects | BioSamples | Sequence Read Archive |
| Blood, spleen, and liver of 2-3 month old *Peromyscus leucopus* LL stock after injection of lipopolysaccharide (LPS) or buffer alone (control) | PRJNA643534 | SAMN15445639-SAMN15445698<br>(20 animals x 3 tissues = 60 samples) | SRR12781556-SRR12781615 |
| Blood, spleen, and liver of 2-3 month old *Mus musculus* BALB/c after treatment with LPS or buffer alone | PRJNA643535 | SAMN15445905-SAMN15445964<br>(20 animals x 3 tissues = 60 samples) | SRR12782328-SRR12782387 |
| Blood and/or spleen of 1-2 year old *P. leucopus* LL stock after injection of LPS or buffer alone. | PRJNA644403 | SAMN15469591-SAMN15469598;<br>SAMN16439936-SAMN16439945<br>(18 samples) | SRR15769470-SRR15769487 |
| Blood of *P. leucopus* LL stock infected with *Borrelia hermsii* strain MTW | PRJNA508222 | SAMN10522571-SAMN10522573;<br>SAMN10522575-SAMN10522578<br>(7 samples) | SRR8283810-SRR8283816 |
| Fibroblast cells from the ears of *P. leucopus* LL stock cultivated with or without LPS | PRJNA672217 | SAMN16563388-SAMN16563397<br>(5 pairs) | SRR13021354-SRR13021363 |

### Differential expression

We used *edgeR* v. 3.28.1 [114] for differential gene expression (DEG) analysis. Genes were called differentially expressed if their absolute fold-change between conditions was > 4.0 and the false-discovery rate (FDR) was <0.05. To compare fold-changes across species, we merged the output tables from the DEG analyses and retained 14,685 orthologous genes that were synonymously annotated between both species, out of total 24,295 annotated genes for *P. leucopus* and 35,805 for *M. musculus* (Tables S4-S9 of Supplementary Materials). To screen for DEGs that varied in magnitude by species, we required an absolute fold change of > 5.0 (log_2_ = 2.5) in one species and < 5.0 in the other. For DEGs designated as “shared” between the species, the absolute fold change was > 5.0 in both. Enrichment analysis was done for each one of the gene groups, separated by up or done regulation. Enrichment of Gene Ontology (GO; http://geneontology.org) terms for biological processes was computed using EnrichR (https://amp.pharm.mssm.edu/Enrichr) [115] and plotted using *ggplot* (v. 3.3.2) of the *R* package [116]. The GO terms were sorted by ascending *p* value.

RNA-seq of a limited set of protein coding sequences (CDS) of both species in the same DEG analysis was carried out using CLC Genomics Workbench v. 20 (Qiagen). Paired-end reads were mapped with a length fraction of 0.4, similarity fraction of 0.9, and penalties of 3 for mismatch, insertion, or deletion to the CDS of sets of corresponding orthologous mRNAs of *P. leucopus* and *M. musculus*. Accession numbers are given in Table S20. Counted reads were those that were species-specific. For example, for Arg1 the coding sequences for both *P. leucopus* and *M. musculus* were in the reference set but for *P. leucopus* only the unique reads for Arg1 for that species were tabulated. The cross-hybridization of reads, e.g. number of *M. musculus* reads mapping to *P. leucopus* Arg1, was no more than 5% of that of the homologously mapped gene, e.g. *M. musculus* Arg1, and usually <1%. Expression values were unique reads normalized for total reads across all the samples without adjustment for reference sequence length. These were log_10_-transformed.

### Weighted correlation network analysis

We used WGCNA [117] to identify densely interconnected genes (modules) for the 6 datasets across species and tissues by building a matrix of gene expression with genes TPM >1 in one or more individual. We selected a power (@) of 13 for a soft threshold for the weighted network and specified a minimum of 100 genes per module. To merge modules with similar gene expression profiles, we carried out a dynamic tree cut, with an eigengene dissimilarity threshold of 0.2 that generated the final eigengene profiles, where eigengene is the first principal component of module expression matrix [118]. The inferred modules were distinguished by assigned hexadecimal color code (https://www.color-hex.com/color-names.html), e.g. darkorange2. Modules, associated GO terms, and constituent genes that are not presented in this paper have been deposited with Dryad (http://datadryad.org) under the dataset name “Peromyscus_WGCNA_supplement” (https://doi.org/7280/D1B38G).

### Statistics

Means are presented with 95% confidence intervals. Parametric (*t* test) and non-parametric (Mann-Whitney) tests of significance were 2-tailed. Unless otherwise stated, the *t* test *p* value is given. Adjustment of *p* values for multiple testing was by the Benjamini-Hochberg method [119], as implemented in *edgeR*, CLC Genomics Workbench (see above), or False Discovery Rate Online Calculator (https://tools.carbocation.com/FDR). For categorical data an exact Likelihood Ratio Test was performed with StatExact v. 6 (Cytel Statistical Software). Other methods are given in software programs or suites cited above.

## ACKNOWLEDGEMENTS

We thank Khalil Bassam, Angelique Cortez, and Hanjuan Shao for their assistance in these studies. The research was supported by NIH grant AI136523 (to A.G.B.) and NSF grant OIA-1736150 (to H.K.), This research was also supported by the Office of the Assistant Secretary of Defense for Health Affairs through the Tick Borne Disease Research Program under award no. W81XWH-17-1-0481 (to A.G.B.). Opinions, interpretations, conclusions, and recommendations are those of the authors and are not necessarily endorsed by the Department of Defense. This work was made possible, in part, through access to the Genomics High Throughput Facility Shared Resource of the Cancer Center Support Grant (P30CA-062203) at the University of California, Irvine and NIH shared instrumentation grants 1S10RR025496-01, 1S10OD010794-01, and 1S10OD021718-01. The PGSC was supported by NSF grant DBI1755670.

