## Supplemental Figures S1-S6 and Tables S1-S20 for "The infection-tolerant mammalian reservoir of Lyme disease and other zoonoses broadly counters the inflammatory effects of endotoxin"

### Supplemental Materials

Figures S1-S6

Tables S1-S20

Figure S1. Digital photograph of two adult *Peromyscus leucopus* with exudative conjunctivitis and huddled together. The animals had received 10 mg/gm body of *Escherichia coli* lipopolysaccharide intraperitoneally the day before.

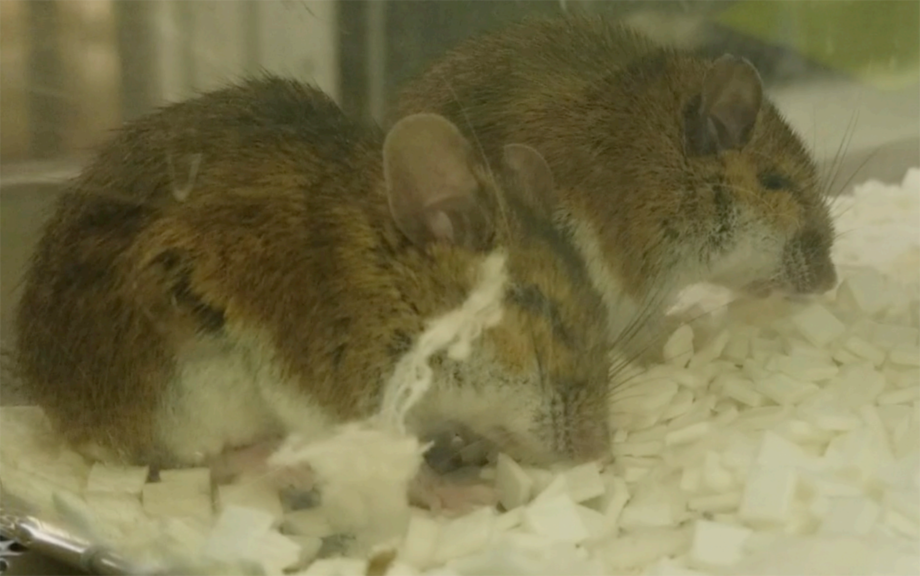

Figure S2. Box-whisker plots of tryptophan metabolites in plasma of *M. musculus* and *P. leucopus* four hours after intraperitoneal injection of LPS at a dose of 10 µg/gm body weigth or buffer alone. The abundances as estimated from peak areas of LC/MS are indicated on the *y*-axis.

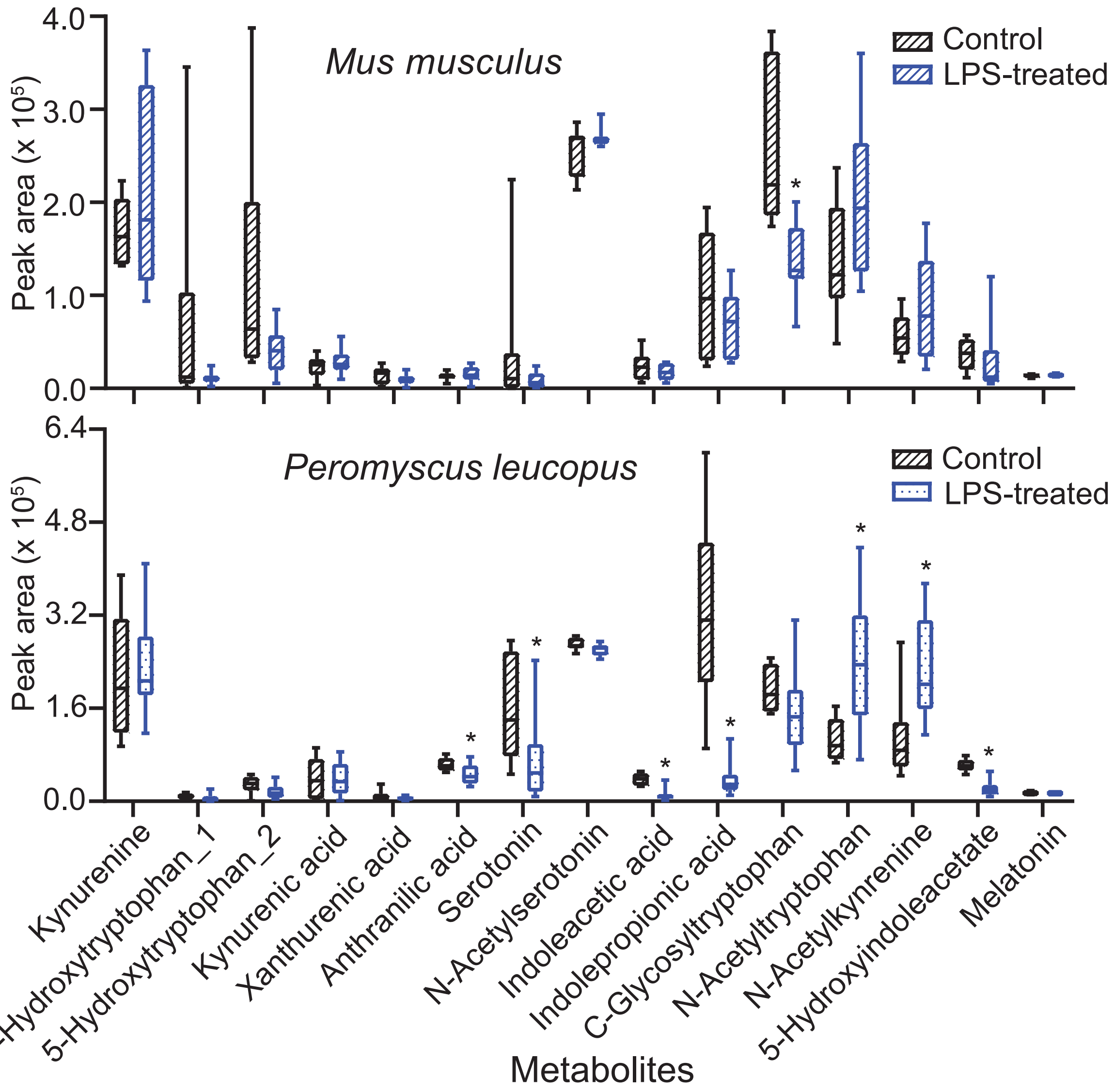

Mus musculus
Peromyscus leucopus
Control
LPS-treated
Peak area (x 10^5)
4.0
3.0
2.0
1.0
0.0
6.4
4.8
3.2
1.6
0.0
*
Kynurenine
5-Hydroxytryptophan_1
5-Hydroxytryptophan_2
Kynurenic acid
Xanthurenic acid
Anthranilic acid
Serotonin
N-Acetylserotonin
Indoleacetic acid
Indolepropionic acid
C-Glycosyltryptophan
N-Acetyltryptophan
N-Acetylkynrenine
5-Hydroxyindoleacetate
Melatonin
Metabolites

Figure S3. Species- and tissue-specific responses to LPS. Independent differential gene expression analysis of RNA-seq data were performed for blood, spleen, and liver tissues of *P. leucopus* and *M. musculus* collected 4 h after injection with LPS or buffer alsone as control. These are represented as volcano plots with range-adjusted scales for the $\log_2$-transformed fold-changes on *x*-axes and $\log_{10}$-transformed FDR *p* values on *y*-axes. Colors of symbols denote the following: red, up-regulated gene with absolute fold-change > 4.0 and *p* value < 0.05; purple, down-regulated gene with absolute fold-change > 4.0 and *p* value < 0.05; and gray, all others. Numbers at the top left and right corners in each plot respresent numbers of down- and up-regulated genes, respectively. Figure 3 is same data with constant scales for *x*- and *y*-axes across the plots. Numerical values for each gene in the 6 datasets are provided in Tables S4-S9.

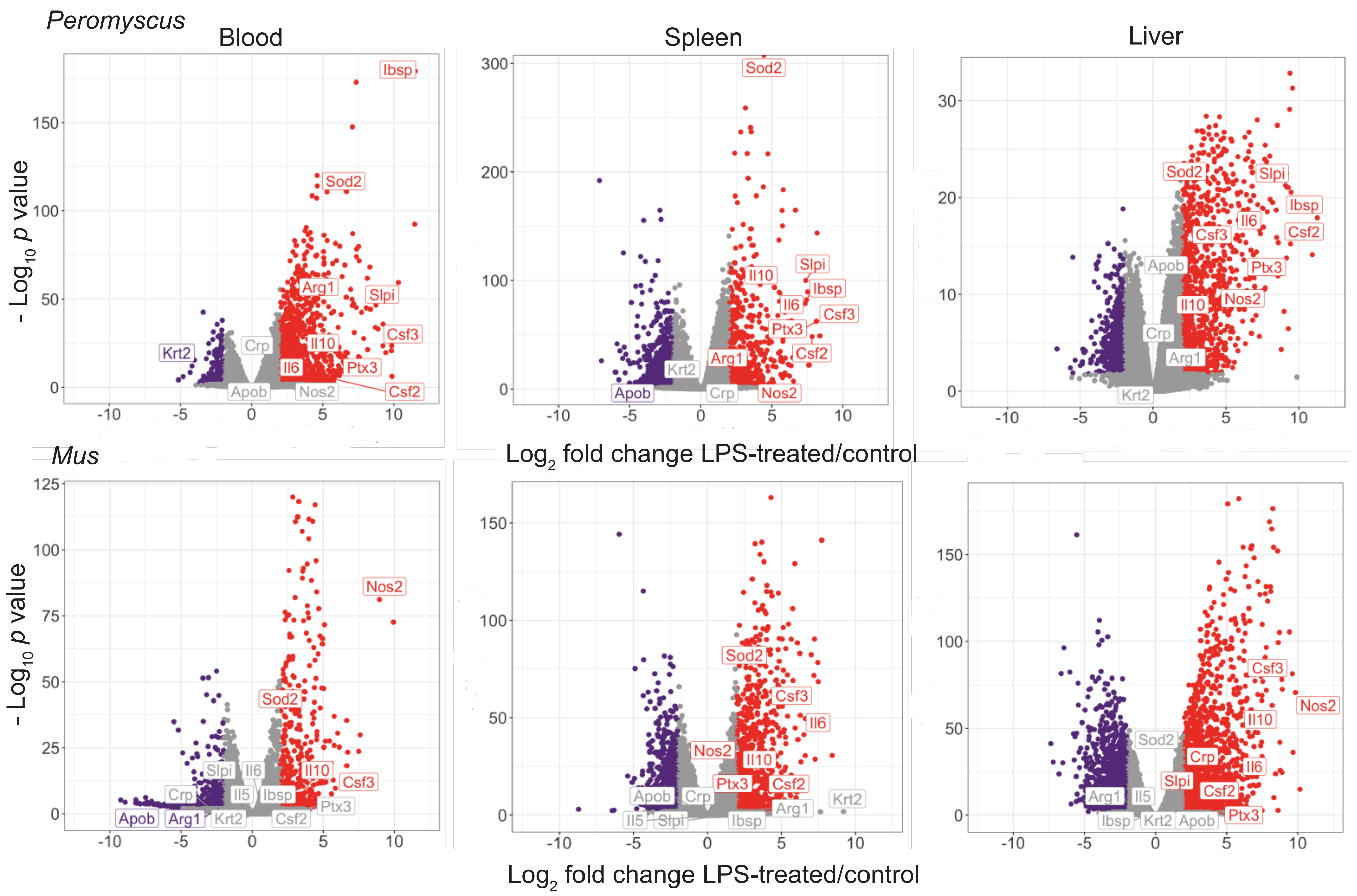

Peromyscus
Blood
Spleen
Liver
- Log10 p value
Log2 fold change LPS-treated/control
Mus
- Log10 p value
Log2 fold change LPS-treated/control
Ibsp
Sod2
Arg1
Slpi
Csf3
Il10
Il6
Ptx3
Csf2
Krt2
Crp
Apob
Nos2
Sod2
Il10
Slpi
Ibsp
Il6
Csf3
Ptx3
Csf2
Arg1
Krt2
Apob
Crp
Nos2
Sod2
Slpi
Ibsp
Il6
Csf3
Csf2
Apob
Ptx3
Nos2
Il10
Crp
Arg1
Krt2
Nos2
Sod2
Slpi
Il6
Il10
Csf3
Crp
Il5
Ibsp
Ptx3
Apob
Arg1
Krt2
Csf2
Sod2
Csf3
Il6
Nos2
Il10
Ptx3
Csf2
Apob
Crp
Krt2
Arg1
Il5
Slpi
Ibsp
Csf3
Nos2
Il10
Sod2
Crp
Il6
Slpi
Csf2
Arg1
Il5
Ptx3
Ibsp
Krt2
Apob

Figure S4. Correlation of IL-10 and IL-10 *P. leucopus* and *M. musculus* from RNA-seq of spleen and of Figure 6B and Table S14. The scatter plot is $\log_{10}$ values of normalized unique reads of one coding sequence against another for each of the four groups, as defined in the legend for Figure 6 and indicated by different symbols.

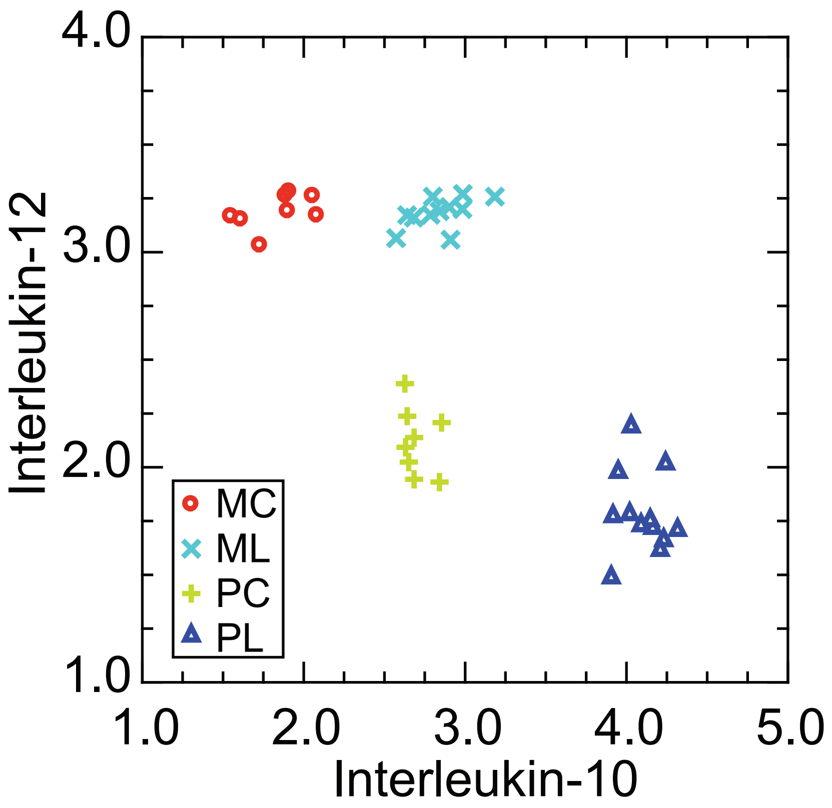

4.0
3.0
2.0
1.0
Interleukin-12
MC
ML
PC
PL
1.0
2.0
3.0
4.0
5.0
Interleukin-10

Figure S5. Assessment of diversity among individual animals by species in transcriptional responses to LPS for 17 genes. Pairwise coefficients of determination ($R^2$) were calculated for the 66 intraspecies pairs for LPS-treated *P. leucopus*, the 66 intraspecies pairs for LPS-treated *M. musculus*, and the 144 interspecies pairs for all LPS-treated animals. Data were drawn from selective RNA-seq for the spleen (Table S14). The top panel is a frequency distribution of $R^2$ values for all 3 sets of determinations. The bottom panel shows just the distributions of intraspecies pairwise determinations.

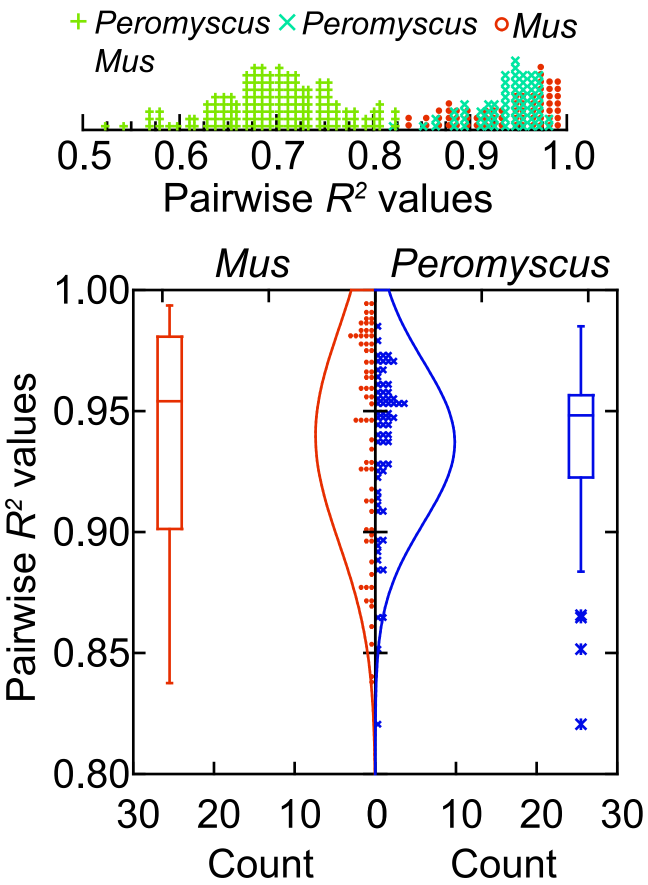

+ Peromyscus
Mus
× Peromyscus
○ Mus
0.5
0.6
0.7
0.8
0.9
1.0
Pairwise $R^2$ values
Mus
Peromyscus
1.00
0.95
0.90
0.85
0.80
Pairwise $R^2$ values
30
20
10
0
10
20
30
Count
Count

Figure S6. DEGs of RNA-seq of selected protein coding sequences for blood or spleen samples from *P. leucopus* of age 1-2 years. The figure comprises 9 box-plots for representative genes, singly or in pairs. The *x*-axes indicate the treatment groups: control or LPS-treated. The *y*-axes are $log_{10}$-transformed, normalized unique reads per coding sequence. Data values by individual coding sequence are given in Table S13.

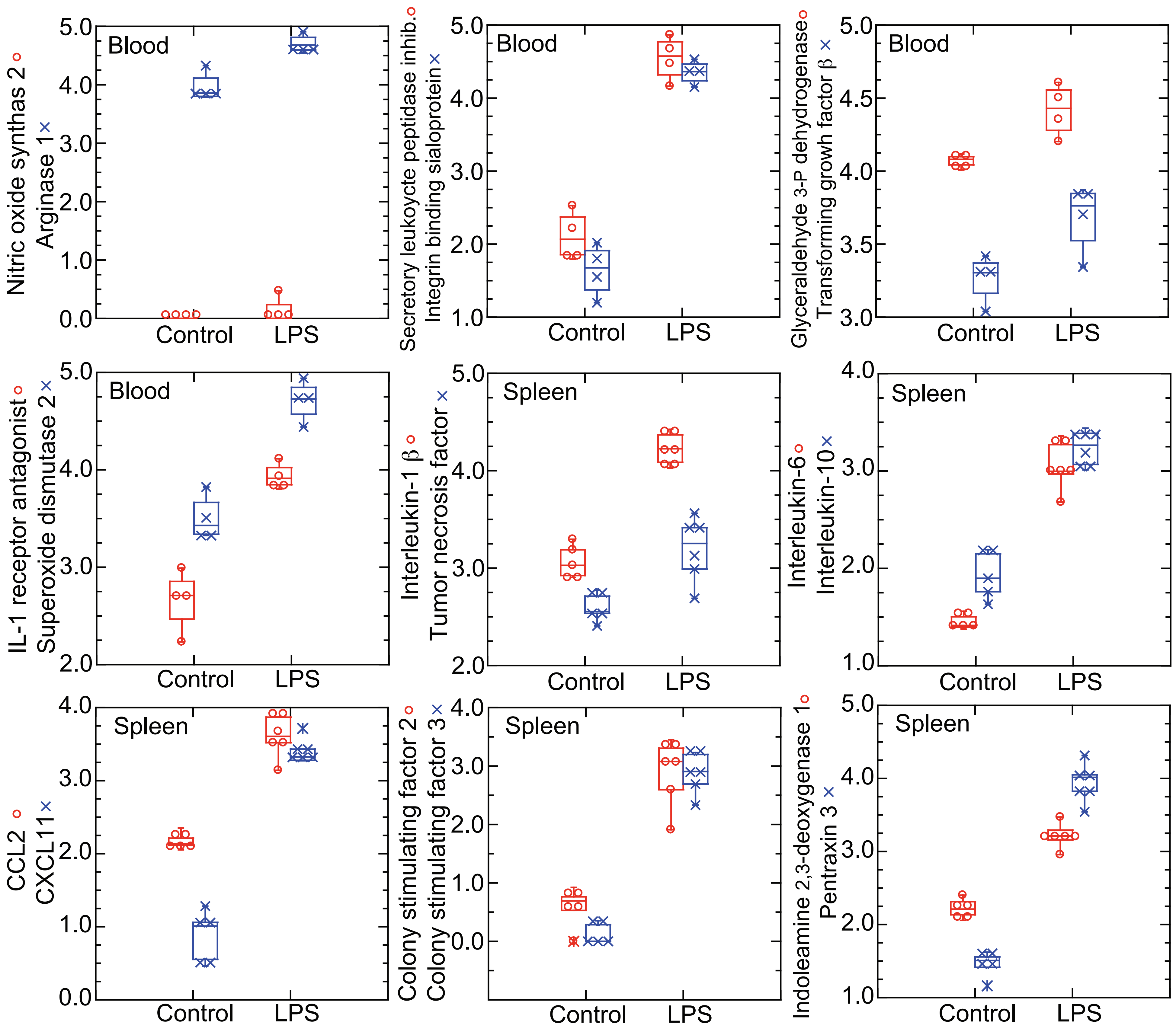

Blood
Nitric oxide synthas 2
Arginase 1
Control
LPS
Blood
Secretory leukoycte peptidase inhib.
Integrin binding sialoprotein
Control
LPS
Blood
Glyceraldehyde 3-P dehydrogenase
Transforming growth factor β
Control
LPS
Blood
IL-1 receptor antagonist
Superoxide dismutase 2
Control
LPS
Spleen
Interleukin-1 β
Tumor necrosis factor
Control
LPS
Spleen
Interleukin-6
Interleukin-10
Control
LPS
Spleen
CCL2
CXCL11
Control
LPS
Spleen
Colony stimulating factor 2
Colony stimulating factor 3
Control
LPS
Spleen
Indoleamine 2,3-deoxygenase 1
Pentraxin 3
Control
LPS

| Table S1. Line listing of animals by species and other characteristics for two LPS experiments | | | | | | | | | | |
|---|---|---|---|---|---|---|---|---|---|---|
| Species | Animal | Group | Sex | Age (weeks | Mass (g) | Conjunc-tivitis | Specimens | Microbiome alpha diversity | Cortico-sterone | Nitric oxide |
| *Peromyscus leucopus* | 22275 | Control | F | 15 | 20 | - | bld, liv, spl | 3.49 | 254 | 17.3 |
| | 22277* | Control | F | 15 | 14 | - | bld, liv, spl | 3.18 | 238 | 1.1 |
| | 22282 | Control | F | 14 | 15 | - | bld, liv, spl | 3.51 | 21 | nd |
| | 22279 | Control | F | 15 | 15 | - | bld, liv, spl | 3.50 | 78 | 3.9 |
| | 22321 | Control | M | 13 | 16 | - | bld, liv, spl | 3.49 | 214 | 9.8 |
| | 22335 | Control | M | 12 | 17 | - | bld, liv, spl | 3.42 | 84 | 7.8 |
| | 22341 | Control | M | 11 | 15 | - | bld, liv, spl | 3.80 | 100 | 3.7 |
| | 22315 | Control | M | 13 | 20 | - | bld, liv, spl | 3.83 | 497 | nd |
| | 22258 | LPS | F | 17 | 26 | - | bld, liv, spl | 3.54 | 681 | 7.7 |
| | 22281 | LPS | F | 15 | 21 | + | bld, liv, spl | 3.61 | 748 | 10.2 |
| | 22278* | LPS | F | 15 | 17 | + | bld, liv, spl | 3.31 | 740 | 11.5 |
| | 22289 | LPS | F | 15 | 15 | - | bld, liv, spl | 3.62 | 642 | nd |
| | 22292 | LPS | F | 15 | 20 | - | bld, liv, spl | 3.69 | 817 | 0.0 |
| | 22284 | LPS | F | 15 | 15 | + | bld, liv, spl | 3.69 | 671 | 0.0 |
| | 22319 | LPS | M | 13 | 18 | - | bld, liv, spl | 3.76 | 697 | 2.7 |
| | 22316 | LPS | M | 13 | 21 | + | bld, liv, spl | 3.74 | 713 | 14.7 |
| | 22325 | LPS | M | 12 | 27 | - | bld, liv, spl | 3.73 | 697 | nd |
| | 22332 | LPS | M | 12 | 25 | - | bld, liv, spl | 3.40 | 658 | nd |
| | 22327 | LPS | M | 12 | 15 | + | bld, liv, spl | 3.77 | 664 | 4.7 |
| | 22339 | LPS | M | 11 | 20 | + | bld, liv, spl | 3.87 | 658 | 11.3 |
| *Mus musculus* | 102 | Control | F | 10 | 20 | - | bld, liv, spl | 3.68 | nd | nd |
| | 120 | Control | F | 10 | 20 | - | bld, liv, spl | 3.52 | 21 | 7.8 |
| | F1_01 | Control | F | 10 | 20 | - | bld, liv, spl | 3.65 | 159 | 4.5 |
| | F3_02 | Control | F | 10 | 19 | - | bld, liv, spl | 3.65 | 109 | 2.7 |
| | 106 | Control | M | 12 | 24 | - | bld, liv, spl | 3.67 | 34 | 20.7 |
| | 115 | Control | M | 12 | 26 | - | bld, liv, spl | 3.59 | 137 | 4.7 |
| | 116 | Control | M | 12 | 29 | - | bld, liv, spl | 3.57 | 38 | 3.7 |
| | M4_04 | Control | M | 12 | 31 | - | bld, liv, spl | 3.62 | 40 | 7.8 |
| | 101 | LPS | F | 10 | 19 | - | bld, liv, spl | 2.98 | 709 | 52.1 |
| | 103 | LPS | F | 10 | 21 | - | bld, liv, spl | 3.57 | 699 | 45.6 |
| | 104 | LPS | F | 10 | 20 | - | bld, liv, spl | 3.14 | 662 | 29.3 |
| | 105 | LPS | F | 10 | 19 | - | bld, liv, spl | 3.47 | 647 | 42.6 |
| | 118 | LPS | F | 10 | 18 | - | bld, liv, spl | 3.63 | 709 | 18.3 |
| | F3_03 | LPS | F | 10 | 21 | - | bld, liv, spl | 3.65 | 687 | 23.5 |
| | 107 | LPS | M | 12 | 29 | - | bld, liv, spl | 3.69 | 566 | 8.1 |
| | 108 | LPS | M | 12 | 25 | - | bld, liv, spl | 3.59 | 490 | 30.3 |
| | 109 | LPS | M | 12 | 29 | - | bld, liv, spl | 3.70 | 634 | 25.5 |
| | M2_05 | LPS | M | 12 | 27 | - | bld, liv, spl | 3.66 | 587 | 12.2 |

| | | | | | | | | | | |
|---|---|---|---|---|---|---|---|---|---|---|
| | M4_06 | LPS | M | 12 | 28 | - | bld, liv, spl | 3.62 | 518 | nd |
| | M4_07 | LPS | M | 12 | 30 | - | bld, liv, spl | 3.62 | 577 | 2.5 |
| *Peromyscus leucopus* | 20694 | Control | F | 83 | 21 | - | spl | nd | nd | nd |
| | 20713 | Control | F | 73 | 23 | - | bld | nd | nd | nd |
| | 20733† | Control | F | 79 | 18 | - | spl | nd | nd | nd |
| | 20807# | Control | F | 69 | 18 | - | spl | nd | nd | nd |
| | 20949 | Control | F | 54 | 19 | - | spl | nd | nd | nd |
| | 20495 | Control | M | 93 | 22 | - | bld, spl | nd | nd | nd |
| | 20613 | Control | M | 84 | 22 | - | bld | nd | nd | nd |
| | 20688 | LPS | F | 83 | 18 | + | spl | nd | nd | nd |
| | 20695 | LPS | F | 83 | 25 | - | spl | nd | nd | nd |
| | 20696 | LPS | F | 74 | 18 | - | bld | nd | nd | nd |
| | 20702# | LPS | F | 83 | 17 | - | spl | nd | nd | nd |
| | 20704 | LPS | F | 74 | 19 | - | bld | nd | nd | nd |
| | 20732† | LPS | F | 71 | 23 | - | bld, spl | nd | nd | nd |
| | 20819 | LPS | F | 69 | 28 | + | spl | nd | nd | nd |
| | 20483 | LPS | M | 94 | 23 | - | bld, spl | nd | nd | nd |
| | 20501 | LPS | M | 93 | 24 | - | bld | nd | nd | nd |

* Mating pair H-106, same litter
† Mating pair H-1056, same litter
### Mating pair H-1055, different litter

Table S2. Pathway enrichments from metabolomics of plasma of Peromyscus leucopus and Mus musculus with or without LPS treatment

| Pathway | Pathway total | Mus hits | Mus significant | Mus expected | Mus enriched | Mus p value | Peromyscus hits | Peromyscus significant | Peromyscus expected | Peromyscus enriched | Peromyscus p value | Mus KEGG hits | Peromyscus KEGG hits |
|---|---|---|---|---|---|---|---|---|---|---|---|---|---|
| Alanine, aspartate and glutamate metabolism | 28 | 25 | 17 | 8.66 | 1.96 | 6.7E-04 | 25 | 15 | 12.30 | 1.22 | 2.0E-01 | C01042;C02362;C00049;C00152;C00402;C03406;C00041;C00232;C00025;C03912;C20776;C00158;C00438;C00026;C00169;C00352;C03090 | C01042;C02362;C00049;C00152;C00402;C00041;C00232;C00025;C00334;C00064;C20776;C00158;C00026;C00352;C03090 |
| alpha-Linolenic acid metabolism | 12 | 11 | 4 | 3.71 | 1.08 | 5.7E-01 | 11 | 2 | 5.27 | 0.38 | 9.9E-01 | C16331;C16327;C06427;C16336;C16300 | C06427;C16300 |
| Amino sugar and nucleotide sugar metabolism | 35 | 28 | 4 | 10.82 | 0.37 | 1.0E+00 | 35 | 13 | 15.37 | 0.85 | 9.5E-01 | C06241;C00270;C00352;C00190 | C00357;C04501;C00270;C00029;C00352;C00267;C00984;C01019;C03410;C00461;C00159;C02336;C00329;C04257;C00052;C03691 |
| Aminoacyl-tRNA biosynthesis | 22 | 21 | 11 | 6.80 | 1.62 | 7.4E-02 | 21 | 18 | 9.66 | 1.86 | 6.2E-04 | C00152;C00135;C00079;C00049;C00065;C00073;C00183;C00041;C00078;C00148;C00025 | C00152;C00079;C00062;C00064;C00037;C00049;C00065;C00073;C00183;C00041;C00047;C00407;C00123;C00188;C00078;C00082;C00148;C00025 |
| Arachidonic acid metabolism | 35 | 35 | 4 | 10.82 | 0.37 | 1.0E+00 | 35 | 13 | 15.37 | 0.85 | 9.5E-01 | C00219;C05956;C02166;C05951 | C14768;C14769;C14770;C14771;C00219;C14781;C14813;C05966;C05356;C05956;C00909;C02165;C00427;C00584;C14782;C14814;C14778;C14823;C14812;C14748;C04742;C14749;C04805;C05965;C04853;C01312;C02198;C00696;C00639 |
| Arginine and proline metabolism | 37 | 35 | 10 | 11.44 | 0.87 | 8.4E-01 | 35 | 21 | 16.25 | 1.29 | 1.4E-01 | C05933;C00300;C00019;C00763;C05946;C05938;C00148;C03912;C03287;C00025;C00077 | C00062;C00334;C00019;C00315;C02714;C05936;C00750;C05147;C00763;C05947;C05938;C04281;C01157;C00148;C03440;C03287;C00025;C01165;C00077;C02946;C04282 |
| Arginine biosynthesis | 14 | 13 | 9 | 4.33 | 2.08 | 1.2E-02 | 13 | 9 | 6.15 | 1.46 | 1.3E-01 | C00025;C03406;C00327;C00049;C00169;C00077;C00026;C00624;C00086 | C00025;C00062;C00437;C00327;C00049;C00077;C00064;C00026;C00624 |
| Ascorbate and aldarate metabolism | 10 | 9 | 2 | 3.09 | 0.65 | 8.8E-01 | 9 | 4 | 4.39 | 0.91 | 7.4E-01 | C01040;C00191;C00800 | C01040;C00137;C00029;C00191;C00800;C00818 |
| beta-Alanine metabolism | 21 | 16 | 5 | 6.49 | 0.77 | 7.1E-01 | 16 | 8 | 9.22 | 0.87 | 5.9E-01 | C00099;C00049;C02642;C05341;C00135 | C00083;C00099;C00049;C02642;C00386;C00750;C05341;C00315 |
| Biosynthesis of unsaturated fatty acids | 34 | 27 | 7 | 10.51 | 0.67 | 8.9E-01 | 27 | 10 | 14.94 | 0.67 | 9.4E-01 | C01595;C00219;C03242;C06426;C06429;C06428;C06427 | C00249;C01530;C00712;C01595;C00219;C03242;C06426;C06429;C06428;C06427 |
| Biotin metabolism | 4 | 4 | 1 | 1.24 | 0.81 | 8.2E-01 | 4 | 4 | 1.76 | 2.28 | 6.1E-02 | C00120 | C00120;C05921;C05552;C00047 |
| Butanoate metabolism | 15 | 14 | 8 | 4.64 | 1.72 | 7.3E-02 | 13 | 8 | 6.59 | 1.21 | 2.8E-01 | C01089;C00164;C00332;C00025;C00232;C00246;C00026;C02630 | C01089;C00164;C00334;C00025;C00232;C00246;C00026;C02630 |
| Caffeine metabolism | 12 | 12 | 4 | 3.71 | 1.08 | 6.5E-01 | 12 | 7 | 5.27 | 1.33 | 3.8E-01 | C13747;C07480;C16359;C16360;C16356;C16365;C16355 | C13747;C16358;C16361;C07480;C16353;C16362;C16360;C16356;C16365 |
| Citrate cycle (TCA cycle) | 16 | 14 | 4 | 4.95 | 0.81 | 7.8E-01 | 15 | 8 | 7.03 | 1.14 | 4.9E-01 | C00026;C05379;C00311;C00417;C00158 | C00026;C05381;C05379;C00311;C00149;C00417;C00158;C05125 |
| Cysteine and methionine metabolism | 33 | 28 | 14 | 10.20 | 1.37 | 7.0E-02 | 28 | 17 | 14.50 | 1.17 | 1.6E-01 | C01180;C04582;C04188;C00019;C00065;C00073;C00021;C15650;C00491;C11537;C00109;C01005;C02218;C05528 | C01180;C04582;C04188;C00170;C00019;C02291;C00065;C00155;C00073;C00021;C15650;C00491;C02356;C00109;C03232;C02218;C00197 |
| D-Arginine and D-ornithine metabolism | 4 | 4 | 1 | 1.24 | 0.81 | 8.2E-01 | 4 | 3 | 1.76 | 1.71 | 3.1E-01 | C00515 | C00515;C00792;C01110 |
| D-Glutamine and D-glutamate metabolism | 6 | 6 | 3 | 1.86 | 1.62 | 3.5E-01 | 6 | 5 | 2.64 | 1.90 | 1.1E-01 | C00217;C00025;C00026 | C00217;C00025;C00819;C00064;C02237;C00026 |
| Drug metabolism - cytochrome P450 | 21 | 19 | 10 | 6.49 | 1.54 | 8.4E-02 | 19 | 12 | 9.22 | 1.30 | 1.7E-01 | C07645;C16608;C16609;C16545;C16561;C16551;C07646;C16610;C16596;C16591 | C16608;C16609;C16586;C16595;C16587;C16545;C16561;C16577;C16582;C16584;C16596;C16591 |

| | | | | | | | | | | | | | |
|---|---|---|---|---|---|---|---|---|---|---|---|---|---|
| Drug metabolism - other enzymes | 27 | 20 | 5 | 8.35 | 0.60 | 8.8E-01 | 20 | 12 | 11.86 | 1.01 | 2.4E-01 | C04646;C16631;C16633;C11376;C21749 | C16619;C16614;C16618;C16630;C16631;C16633;C11173;C21748;C21750;C21751;C07585;C16620 |
| Ether lipid metabolism | 2 | 2 | 1 | 0.62 | 1.62 | 5.8E-01 | 2 | 1 | 0.88 | 1.14 | 7.5E-01 | C01233 | C01233 |
| Fatty acid biosynthesis | 10 | 9 | 2 | 3.09 | 0.65 | 8.8E-01 | 9 | 5 | 4.39 | 1.14 | 4.9E-01 | C02679;C06423 | C00083;C00249;C06424;C02679;C01571 |
| Fatty acid degradation | 36 | 29 | 8 | 11.13 | 0.72 | 8.5E-01 | 27 | 4 | 15.81 | 0.25 | 1.0E+00 | C05268;C05274;C02990;C05271;C00332;C05267;C05275;C05261 | C00249;C02990;C05269;C00010 |
| Fatty acid elongation | 30 | 22 | 5 | 9.28 | 0.54 | 9.3E-01 | 21 | 3 | 13.18 | 0.23 | 1.0E+00 | C05261;C05274;C05275;C05267;C05271;C05268 | C05269;C00083;C00249 |
| Folate biosynthesis | 24 | 24 | 7 | 7.42 | 0.94 | 7.9E-01 | 23 | 6 | 10.54 | 0.57 | 9.9E-01 | C00268;C06148;C05923;C00921;C04895;C19848;C15522;C00272;C04244;C02953;C21640;C04874 | C00504;C00921;C18239;C15522;C04874;C20239 |
| Galactose metabolism | 27 | 23 | 2 | 8.35 | 0.24 | 1.0E+00 | 27 | 8 | 11.86 | 0.67 | 9.9E-01 | C00089;C05402;C05401;C05400;C05399;C01235;C00243 | C00124;C05401;C05399;C00267;C00052;C00029;C00984;C00031;C00095;C01697;C00116;C00159;C00794;C00137 |
| Glutathione metabolism | 19 | 18 | 3 | 5.87 | 0.51 | 9.8E-01 | 17 | 12 | 8.35 | 1.44 | 6.7E-02 | C00669;C00025;C00077 | C00051;C00127;C00037;C00669;C00025;C01419;C01879;C00077;C00315;C01672;C02090;C00750 |
| Glycerolipid metabolism | 7 | 6 | 1 | 2.16 | 0.46 | 9.2E-01 | 6 | 4 | 3.07 | 1.30 | 3.4E-01 | C00093 | C00093;C00116;C00111;C00631 |
| Glycerophospholipid metabolism | 13 | 12 | 4 | 4.02 | 1.00 | 6.5E-01 | 12 | 7 | 5.71 | 1.23 | 3.8E-01 | C01996;C00346;C00093;C01233 | C00111;C00588;C00114;C01996;C00346;C00093;C01233 |
| Glycine, serine and threonine metabolism | 31 | 24 | 7 | 9.58 | 0.73 | 7.9E-01 | 24 | 16 | 13.62 | 1.18 | 7.0E-02 | C00065;C00719;C01005;C00213;C00740;C00300;C00109 | C00065;C00114;C00719;C00197;C01026;C02291;C00037;C00213;C00143;C00188;C00631;C00740;C03232;C05519;C00109;C03508;C00430 |
| Glycosaminoglycan biosynthesis - chondroitin sulfate / [illegible] | 8 | 6 | 1 | 2.47 | 0.40 | 9.2E-01 | 7 | 2 | 3.51 | 0.57 | 9.4E-01 | C00401 | G00159;G00154 |
| Glycosaminoglycan degradation | 21 | 19 | 3 | 6.49 | 0.46 | 9.8E-01 | 19 | 4 | 9.22 | 0.43 | 1.0E+00 | G13074;G00526;G02632 | G13035;G13037;G00526;G00711 |
| Glycosphingolipid biosynthesis - ganglio series | 21 | 16 | 1 | 6.49 | 0.15 | 1.0E+00 | 16 | 3 | 9.22 | 0.33 | 1.0E+00 | G00113 | G00109;G00108;G00092 |
| Glycosphingolipid biosynthesis - lacto and neolacto seri[illegible] | 43 | 39 | 3 | 13.30 | 0.23 | 1.0E+00 | 33 | 4 | 18.89 | 0.21 | 1.0E+00 | G00082;G00068;G00077 | G00051;G00092;G00038;G00063;G00428 |
| Glyoxylate and dicarboxylate metabolism | 31 | 24 | 7 | 9.58 | 0.73 | 7.9E-01 | 24 | 9 | 13.62 | 0.66 | 9.2E-01 | C00417;C01127;C00158;C00065;C00025;C00311;C00332 | C00417;C00158;C00149;C00065;C00037;C00025;C00311;C00064;C00631 |
| Histidine metabolism | 16 | 13 | 9 | 4.95 | 1.82 | 1.2E-02 | 11 | 9 | 7.03 | 1.28 | 3.1E-02 | C00025;C03680;C00785;C00135;C05130;C01152;C00439;C05828;C00049 | C00025;C03680;C00386;C05130;C01152;C01044;C00439;C05828;C00049 |
| Inositol phosphate metabolism | 21 | 17 | 2 | 6.49 | 0.31 | 9.9E-01 | 21 | 4 | 9.22 | 0.43 | 1.0E+00 | C01245;C01243;C00191 | C01245;C01243;C00137;C00111;C00191;C00118 |
| Linoleic acid metabolism | 4 | 4 | 3 | 1.24 | 2.43 | 1.3E-01 | 4 | 3 | 1.76 | 1.71 | 3.1E-01 | C01595;C14825;C14826 | C01595;C14825;C14826 |
| Lipoic acid metabolism | 2 | 2 | 1 | 0.62 | 1.62 | 5.8E-01 | 2 | 1 | 0.88 | 1.14 | 7.5E-01 | C16241 | C16241 |
| Lysine degradation | 19 | 19 | 7 | 5.87 | 1.19 | 5.2E-01 | 18 | 10 | 8.35 | 1.20 | 4.0E-01 | C01181;C03793;C00449;C00956;C00322;C00332;C03366 | C00047;C01181;C03793;C00449;C04076;C00956;C00322;C00408;C03366;C00450 |
| Mannose type O-glycan biosynthesis | 16 | 16 | 3 | 4.95 | 0.61 | 9.6E-01 | 16 | 4 | 7.03 | 0.57 | 9.9E-01 | G13091;C00789;C01068;G13028 | C00789;G13096;G13095;C01068 |
| Metabolism of xenobiotics by cytochrome P450 | 64 | 55 | 19 | 19.79 | 0.96 | 5.8E-01 | 56 | 24 | 28.11 | 0.85 | 8.8E-01 | C14852;C06899;C14800;C19604;C19586;C06800;C19561;C19489;C19490;C14040;C14853;C14866;C14871;C14856;C14855;C14801;C19566;C19607;C19595;C16756;C19559;C11148;C19563;C14802 | C14852;C14556;C14851;C14850;C14865;C06205;C14800;C19604;C19586;C19561;C19489;C19490;C14040;C14853;C14871;C14868;C14861;C14864;C14791;C14801;C19607;C11278;C19595;C16756;C19559;C14849;C14802 |
| N-Glycan biosynthesis | 38 | 24 | 3 | 11.75 | 0.26 | 1.0E+00 | 22 | 1 | 16.69 | 0.06 | 1.0E+00 | G00014;G00004;G00003 | C00621 |
| Nicotinate and nicotinamide metabolism | 15 | 13 | 7 | 4.64 | 1.51 | 1.3E-01 | 11 | 8 | 6.59 | 1.21 | 1.1E-01 | C00049;C03150;C00153;C02918;C05840;C05843;C05842 | C00049;C00455;C03150;C00153;C02918;C05840;C05843;C05842 |
| Nitrogen metabolism | 6 | 4 | 2 | 1.86 | 1.08 | 4.4E-01 | 4 | 2 | 2.64 | 0.76 | 6.8E-01 | C00025;C00169 | C00025;C00064 |

| | | | | | | | | | | | | | |
|---|---|---|---|---|---|---|---|---|---|---|---|---|---|
| Pantothenate and CoA biosynthesis | 17 | 14 | 8 | 5.26 | 1.52 | 7.3E-02 | 14 | 11 | 7.47 | 1.47 | 2.7E-02 | C00882;C01134;C00831;C04079;C02642;C00183;C00049;C00099 | C00831;C04352;C04079;C00864;C02642;C00183;C00049;C00010;C03492;C00099;C00054 |
| Pentose and glucuronate interconversions | 17 | 16 | 5 | 5.26 | 0.95 | 7.1E-01 | 17 | 10 | 7.47 | 1.34 | 3.1E-01 | C01068;C00310;C00532;C00379;C00800;C00181;C00191;C00789;C00259;C00312;C00618;C02266 | C01068;C00310;C00532;C00379;C00800;C00029;C00181;C00191;C00789;C00259;C00312;C00618;C02266 |
| Pentose phosphate pathway | 22 | 18 | 5 | 6.80 | 0.74 | 8.1E-01 | 21 | 6 | 9.66 | 0.62 | 9.9E-01 | C01801;C00121;C00345;C01236;C00257;C00198 | C00121;C05382;C00118;C00257;C00198;C00631 |
| Phenylalanine metabolism | 12 | 12 | 3 | 3.71 | 0.81 | 8.5E-01 | 12 | 8 | 5.27 | 1.52 | 1.9E-01 | C00079;C05852;C01586 | C00079;C05332;C00166;C07086;C02763;C01586;C00082;C05598 |
| Phenylalanine, tyrosine and tryptophan biosynthesis | 4 | 4 | 1 | 1.24 | 0.81 | 8.2E-01 | 4 | 3 | 1.76 | 1.71 | 3.1E-01 | C00079 | C00166;C00079;C00082 |
| Phosphatidylinositol signaling system | 17 | 14 | 1 | 5.26 | 0.19 | 1.0E+00 | 17 | 3 | 7.47 | 0.40 | 1.0E+00 | C01245;C01243 | C00137;C01245;C01243;C11174;C11526 |
| Porphyrin and chlorophyll metabolism | 27 | 26 | 5 | 8.35 | 0.60 | 9.8E-01 | 26 | 9 | 11.86 | 0.76 | 9.6E-01 | C06505;C00500;C03263;C00025;C05768 | C00032;C00037;C00853;C00931;C00430;C00025;C02191;C05770;C06508 |
| Primary bile acid biosynthesis | 46 | 41 | 20 | 14.22 | 1.41 | 4.4E-02 | 42 | 25 | 20.21 | 1.24 | 1.3E-01 | C17339;C05455;C05446;C01301;C17335;C06341;C15520;C04554;C05445;C04722;C17333;C15518;C05453;C05466;C05465;C17337;C17336;C17332;C05122;C17331 | C17339;C05455;C02528;C00037;C05337;C05446;C01301;C17335;C06341;C15520;C00695;C01794;C04554;C05445;C05447;C04722;C17333;C15518;C05453;C05466;C05465;C17337;C17336;C17332;C01921;C17331 |
| Propanoate metabolism | 19 | 17 | 6 | 5.87 | 1.02 | 5.8E-01 | 18 | 5 | 8.35 | 0.60 | 9.8E-01 | C06002;C05983;C00099;C05984;C00109;C00332 | C06002;C00099;C00083;C05984;C00109 |
| Purine metabolism | 66 | 61 | 20 | 20.41 | 0.98 | 6.9E-01 | 59 | 30 | 28.99 | 1.03 | 4.8E-01 | C00035;C00385;C03090;C04640;C04677;C04751;C00575;C00104;C00242;C00499;C00942;C00224;C00081;C00147;C01345;C12248;C00086;C00053;C06196;C02350 | C00035;C00385;C00064;C03090;C04677;C04751;C04734;C00575;C00020;C00130;C00212;C00360;C00559;C05512;C00104;C00144;C00655;C00242;C00330;C00499;C00366;C00362;C00361;C00224;C00147;C01344;C01345;C00131;C06196;C02350 |
| Pyrimidine metabolism | 39 | 31 | 14 | 12.06 | 1.16 | 1.5E-01 | 31 | 18 | 17.13 | 1.05 | 2.2E-01 | C00169;C00299;C02642;C00475;C00705;C00458;C00881;C00460;C01346;C00365;C00526;C00363;C00438;C00099 | C00015;C00064;C00337;C00075;C00299;C02642;C00112;C00475;C00705;C00239;C00881;C00365;C00526;C00214;C21028;C21029;C00099;C01205 |
| Pyruvate metabolism | 19 | 16 | 3 | 5.87 | 0.51 | 9.6E-01 | 17 | 3 | 8.35 | 0.36 | 1.0E+00 | C03451;C05993;C00332 | C05125;C00149;C00083 |
| Retinol metabolism | 15 | 15 | 2 | 4.64 | 0.43 | 9.9E-01 | 15 | 9 | 6.59 | 1.37 | 3.0E-01 | C00777;C15493 | C00376;C02110;C00777;C16681;C15492;C03455;C16677;C16679;C16680;C15493 |
| Selenocompound metabolism | 16 | 10 | 2 | 4.95 | 0.40 | 9.1E-01 | 10 | 2 | 7.03 | 0.28 | 9.9E-01 | C05684;C00041 | C05699;C00041 |
| Sphingolipid metabolism | 9 | 9 | 2 | 2.78 | 0.72 | 8.8E-01 | 9 | 6 | 3.95 | 1.52 | 2.5E-01 | C00065;C00346 | C06124;C01120;C00319;C00065;C02934;C00346 |
| Starch and sucrose metabolism | 12 | 9 | 1 | 3.71 | 0.27 | 9.8E-01 | 12 | 2 | 5.27 | 0.38 | 1.0E+00 | C00089;C01083;C00208;C00252 | C00095;C00029;C00031 |
| Steroid biosynthesis | 41 | 41 | 10 | 12.68 | 0.79 | 9.5E-01 | 41 | 10 | 18.01 | 0.56 | 1.0E+00 | C01561;C05107;C11508;C15783;C11455;C01673;C22112;C22136;C18231;C07712 | C01561;C05107;C01943;C15808;C01673;C00448;C22112;C22136;C18231;C07712 |

| | | | | | | | | | | | | | |
|---|---|---|---|---|---|---|---|---|---|---|---|---|---|
| Steroid hormone biosynthesis | 77 | 77 | 38 | 23.81 | 1.60 | 4.9E-03 | 77 | 61 | 33.82 | 1.80 | 3.6E-08 | C05501;C05499;C01953;C05485;C05138;C05487;C05489;C01227;C05139;C04295;C03205;C02140;C00410;C05476;C05498;C13712;C03681;C01176;C00735;C05488;C05497;C00535;C00280;C04373;C00523;C00468;C05141;C05299;C05301;C01124;C05297;C05294;C18043;C02373;C18038;C18044;C18040;C05473;C04042;C05490;C05478;C05479;C18041;C18075;C05471;C05469;C05285;C03772;C00674;C03917;C05293;C05503;C04518;C18045 | C05501;C05499;C01953;C05485;C05138;C05487;C05489;C05139;C04295;C03205;C02140;C01780;C00410;C05476;C05498;C13712;C03681;C01176;C00735;C05284;C05488;C00762;C05497;C00280;C04373;C00523;C00468;C00951;C05141;C05298;C05299;C05302;C05301;C02538;C01124;C05290;C05297;C05294;C05295;C18043;C18038;C05140;C18040;C05473;C04042;C05490;C05478;C05479;C18041;C18075;C05471;C05469;C05285;C05300;C03917;C05293;C11132;C11131;C05503;C04518;C18045 |
| Sulfur metabolism | 8 | 4 | 2 | 2.47 | 0.81 | 4.4E-01 | 3 | 2 | 3.51 | 0.57 | 5.0E-01 | C00224;C00053 | C00224;C17267 |
| Synthesis and degradation of ketone bodies | 5 | 5 | 3 | 1.55 | 1.94 | 2.3E-01 | 5 | 2 | 2.20 | 0.91 | 8.1E-01 | C00332;C01089;C00164 | C01089;C00164 |
| Taurine and hypotaurine metabolism | 8 | 5 | 2 | 2.47 | 0.81 | 5.7E-01 | 5 | 1 | 3.51 | 0.28 | 9.7E-01 | C05844;C05122 | C05844 |
| Terpenoid backbone biosynthesis | 15 | 15 | 4 | 4.64 | 0.86 | 8.3E-01 | 15 | 3 | 6.59 | 0.46 | 1.0E+00 | C00332;C01143;C00341;C17432 | C01143;C01107;C00448 |
| Thiamine metabolism | 5 | 2 | 1 | 1.55 | 0.65 | 5.8E-01 | 4 | 1 | 2.20 | 0.46 | 9.4E-01 | C01081 | C00378 |
| Tryptophan metabolism | 41 | 41 | 13 | 12.68 | 1.03 | 7.2E-01 | 40 | 30 | 18.01 | 1.67 | 8.3E-04 | C00078;C05635;C05651;C00332;C00322;C04409;C00632;C00328;C05653;C03227;C05647;C05660;C05639;C02220;C00108;C05645;C01252;C05637 | C00078;C00978;C00780;C05635;C05651;C00643;C00322;C03824;C04409;C00632;C00328;C05653;C03227;C00637;C05634;C00398;C00331;C05642;C05643;C05647;C05660;C05639;C02220;C00108;C05640;C05645;C01252;C05637;C00954;C06213 |
| Tyrosine metabolism | 42 | 42 | 16 | 12.99 | 1.23 | 3.9E-01 | 40 | 24 | 18.45 | 1.30 | 1.2E-01 | C05580;C04043;C05587;C00355;C02515;C02514;C01061;C01036;C05578;C05585;C05584;C00164;C00642;C05579;C00822;C01829;C00628;C17938 | C05583;C00547;C05577;C05576;C05588;C03758;C01161;C05581;C05587;C00355;C02515;C00082;C01061;C00544;C03765;C05578;C05584;C05594;C05582;C02465;C00164;C01829;C00628;C17938 |
| Ubiquinone and other terpenoid-quinone biosynthesis | 9 | 9 | 2 | 2.78 | 0.72 | 8.8E-01 | 9 | 4 | 3.95 | 1.01 | 7.4E-01 | C00828;C02059 | C00082;C20265;C05819;C00544 |
| Valine, leucine and isoleucine biosynthesis | 8 | 8 | 2 | 2.47 | 0.81 | 8.3E-01 | 7 | 6 | 3.51 | 1.71 | 6.0E-02 | C00109;C00183 | C00188;C00671;C00123;C00109;C00407;C00183 |
| Valine, leucine and isoleucine degradation | 35 | 26 | 6 | 10.82 | 0.55 | 9.4E-01 | 25 | 11 | 15.37 | 0.72 | 7.8E-01 | C00332;C00164;C00183;C06001;C06002;C00349 | C15976;C00164;C00671;C00183;C06001;C00407;C06002;C00349;C03284;C00123;C01205 |
| Vitamin B6 metabolism | 9 | 9 | 5 | 2.78 | 1.80 | 1.7E-01 | 9 | 8 | 3.95 | 2.02 | 1.8E-02 | C06055;C00647;C00018;C00250;C00847 | C06055;C00647;C00314;C00018;C00250;C00534;C06054;C00847 |

Table S3. Tryptophan metabolite peak areas for plasma of Mus musculus and Peromyscus leucopus with and without LPS

| compound | mean Mus control | mean Mus LPS | Mus fold change | Mus t-test | mean Pero. control | mean Pero. LPS | Pero. fold change | Pero. t-test |
|---|---|---|---|---|---|---|---|---|
| tryptophan | 19936259 | 13286783 | 0.67 | 3.1E-03 | 16296850 | 19901915 | 1.22 | 9.2E-02 |
| kynurenine | 212507 | 230423 | 1.08 | 6.7E-01 | 168712 | 205584 | 1.22 | 3.5E-01 |
| 5-hydroxytryptophan_01 | 8663 | 6018 | 0.69 | 3.5E-01 | 68500 | 10443 | 0.15 | 1.0E-01 |
| 5-hydroxytryptophan_02 | 28809 | 17354 | 0.60 | 7.9E-02 | 117706 | 41044 | 0.35 | 5.5E-02 |
| kynurenic acid | 40153 | 37879 | 0.94 | 8.8E-01 | 23057 | 28907 | 1.25 | 3.3E-01 |
| xanthurenic acid | 8705 | 5078 | 0.58 | 2.3E-01 | 14600 | 9595 | 0.66 | 1.6E-01 |
| anthranilic acid | 63189 | 45994 | 0.73 | 2.5E-02 | 13065 | 15479 | 1.18 | 4.3E-01 |
| serotonin | 161653 | 77062 | 0.48 | 4.0E-02 | 38255 | 8626 | 0.23 | 2.0E-01 |
| N-acetylserotonin | 269871 | 260972 | 0.97 | 7.4E-02 | 257324 | 268634 | 1.04 | 2.0E-01 |
| indoleacetic acid | 37875 | 10662 | 0.28 | 7.8E-06 | 23883 | 17960 | 0.75 | 3.1E-01 |
| indolepropionic acid | 325406 | 36214 | 0.11 | 7.8E-06 | 99493 | 69070 | 0.69 | 2.1E-01 |
| C-glycosyltryptophan | 192626 | 155506 | 0.81 | 2.1E-01 | 260063 | 139304 | 0.54 | 5.1E-04 |
| N-acetyl-tryptophan | 105384 | 234798 | 2.23 | 5.8E-03 | 138381 | 198587 | 1.44 | 9.2E-02 |
| N-acetyl kynurenine | 109013 | 223318 | 2.05 | 7.1E-03 | 57154 | 85204 | 1.49 | 2.1E-01 |
| 5-hydroxy indole acetate | 61949 | 22762 | 0.37 | 4.1E-06 | 36032 | 27053 | 0.75 | 4.9E-01 |
| melatonin | 14264 | 13848 | 0.97 | 5.5E-01 | 13622 | 14262 | 1.05 | 3.5E-01 |

**Table S4. DEGs of the blood of Peromyscus leucopus**

| rank | gene | log2 fold change | FDR |
|---|---|---|---|
| 1 | Ibsp | 11.50 | 1.8E-175 |
| 2 | Fpr1 | 7.35 | 1.5E-169 |
| 3 | Fpr2 | 7.09 | 2.2E-144 |
| 4 | Socs3 | 4.63 | 4.0E-117 |
| 5 | Fcgr2b | 4.64 | 4.2E-111 |
| 6 | Sphk1 | 6.68 | 5.9E-108 |
| 7 | Sod2 | 5.30 | 9.6E-108 |
| 8 | Paf1 | 4.27 | 9.0E-106 |
| 9 | Hk3 | 4.59 | 1.4E-104 |
| 10 | LOC114685652 | 11.48 | 7.1E-90 |
| 11 | Ets2 | 3.85 | 5.7E-88 |
| 12 | Tmem132a | 3.76 | 4.3E-86 |
| 13 | Nfil3 | 4.14 | 3.6E-85 |
| 14 | Gpr84 | 7.10 | 1.5E-84 |
| 15 | LOC114694224 | 3.97 | 5.3E-84 |
| 16 | Il1rap | 5.08 | 1.1E-83 |
| 17 | Srgn | 4.20 | 2.1E-81 |
| 18 | Serpinb1 | 5.30 | 9.2E-81 |
| 19 | Tnip1 | 3.04 | 1.1E-80 |
| 20 | Bcl3 | 3.66 | 1.2E-80 |
| 21 | Il1rn | 5.90 | 3.3E-79 |
| 22 | N4bp1 | 3.15 | 4.1E-78 |
| 23 | Ncf4 | 3.97 | 4.5E-78 |
| 24 | Myo1d | 6.21 | 1.0E-77 |
| 25 | Slc28a3 | 7.53 | 1.5E-77 |
| 26 | Plaur | 4.20 | 4.8E-77 |
| 27 | Il1r2 | 7.38 | 3.9E-76 |
| 28 | Gpr35 | 5.06 | 1.0E-74 |
| 29 | Sla | 2.93 | 1.2E-74 |
| 30 | Tnfaip3 | 3.89 | 1.4E-74 |
| 31 | H6pd | 3.71 | 5.5E-73 |
| 32 | Il1r1 | 4.60 | 7.8E-73 |
| 33 | Nfkbiz | 3.11 | 1.2E-72 |
| 34 | LOC114704488 | 4.52 | 2.9E-72 |
| 35 | Tp53inp2 | 4.54 | 3.1E-72 |
| 36 | Acod1 | 7.37 | 6.6E-72 |
| 37 | Pglyrp1 | 5.07 | 3.2E-71 |
| 38 | Osgin1 | 3.71 | 1.7E-70 |
| 39 | Smtnl2 | 5.98 | 1.1E-69 |
| 40 | Arhgap32 | 7.57 | 1.4E-69 |
| 41 | Sav1 | 2.72 | 1.4E-69 |
| 42 | Socs1 | 5.10 | 1.4E-68 |
| 43 | Anxa3 | 5.70 | 4.6E-68 |
| 44 | Klf5 | 5.08 | 9.8E-68 |
| 45 | Tgm1 | 6.57 | 3.0E-67 |
| 46 | Wipi1 | 3.77 | 5.7E-67 |
| 47 | Nfkbia | 3.27 | 1.1E-66 |
| 48 | Abca13 | 8.33 | 3.6E-66 |
| 49 | St3gal4 | 3.24 | 3.9E-66 |
| 50 | C5ar1 | 3.55 | 1.1E-64 |
| 51 | Xdh | 3.33 | 3.4E-64 |
| 52 | Batf | 3.24 | 4.8E-64 |
| 53 | Sbno2 | 3.37 | 8.1E-64 |
| 54 | Ptafr | 3.61 | 5.0E-63 |
| 55 | Retn | 5.40 | 7.2E-63 |
| 56 | Csf2rb | 4.31 | 7.2E-63 |
| 57 | Nlrc4 | 3.10 | 1.3E-62 |
| 58 | Trib1 | 3.46 | 4.5E-62 |
| 59 | LOC114690951 | 4.52 | 4.9E-62 |
| 60 | Cebpd | 5.09 | 5.7E-62 |
| 61 | Glrx | 3.87 | 5.7E-62 |
| 62 | LOC114689891 | 4.68 | 7.0E-62 |
| 63 | Agtrap | 4.04 | 4.9E-61 |
| 64 | LOC114707706 | 6.38 | 6.6E-61 |
| 65 | Adap2 | 3.34 | 1.6E-60 |
| 66 | Hmox1 | 3.63 | 2.6E-60 |
| 67 | LOC114687425 | 8.15 | 5.7E-60 |
| 68 | Tmem164 | 2.88 | 1.4E-59 |
| 69 | Hip1 | 3.56 | 1.5E-59 |
| 70 | Mmp9 | 5.27 | 2.2E-59 |
| 71 | Cish | 3.34 | 4.6E-59 |
| 72 | Acsl4 | 3.02 | 3.3E-58 |
| 73 | Pstpip2 | 3.88 | 3.4E-58 |
| 74 | Lamp2 | 2.54 | 9.3E-58 |
| 75 | Prdx6 | 3.41 | 1.1E-57 |
| 76 | Slpi | 10.32 | 1.4E-57 |
| 77 | Picalm | 2.98 | 1.7E-57 |
| 78 | Gadd45g | 5.48 | 2.1E-57 |
| 79 | Megf9 | 4.45 | 3.9E-57 |
| 80 | Slc15a3 | 3.67 | 8.5E-57 |
| 81 | Il13ra1 | 4.45 | 1.1E-56 |
| 82 | Marcks | 5.34 | 2.0E-56 |
| 83 | Junb | 2.86 | 4.8E-56 |
| 84 | Hp | 5.21 | 6.4E-56 |
| 85 | Mmp25 | 5.20 | 2.1E-55 |
| 86 | Anxa1 | 4.70 | 4.1E-55 |
| 87 | Pik3ap1 | 3.02 | 5.4E-54 |
| 88 | Inhba | 6.96 | 7.8E-54 |
| 89 | LOC114702149 | 3.86 | 7.8E-54 |
| 90 | Arid5a | 2.90 | 7.8E-54 |
| 91 | Fgr | 3.59 | 1.1E-53 |
| 92 | Tspan14 | 2.20 | 2.0E-53 |
| 93 | Ehd1 | 3.37 | 2.0E-53 |
| 94 | Plagl2 | 2.84 | 2.6E-53 |
| 95 | Chi3l1 | 5.44 | 2.7E-53 |
| 96 | Ier3 | 4.16 | 1.2E-52 |
| 97 | Pim2 | 2.81 | 1.2E-52 |
| 98 | LOC114684895 | 3.23 | 2.1E-52 |
| 99 | Tmem268 | 2.50 | 3.8E-52 |
| 100 | Msrb1 | 4.29 | 7.3E-52 |
| 101 | Olfm4 | 8.85 | 8.8E-52 |
| 102 | Alpk1 | 4.42 | 1.5E-51 |
| 103 | CUNH5orf30 | 3.09 | 1.6E-51 |
| 104 | Tfec | 4.46 | 6.6E-51 |
| 105 | LOC114701319 | 3.38 | 2.1E-50 |
| 106 | Flot1 | 2.87 | 2.1E-50 |
| 107 | LOC114706499 | 3.99 | 5.1E-50 |
| 108 | Cd53 | 2.28 | 5.6E-50 |
| 109 | LOC114688035 | 2.51 | 7.6E-50 |
| 110 | Gramd1a | 2.47 | 7.7E-50 |
| 111 | C1rl | 6.67 | 2.0E-49 |
| 112 | Id1 | 4.62 | 5.3E-49 |
| 113 | Ffar2 | 4.68 | 1.0E-48 |
| 114 | Nek6 | 3.08 | 1.2E-48 |
| 115 | Stx7 | 2.54 | 1.2E-48 |
| 116 | LOC114686723 | 3.45 | 1.4E-48 |
| 117 | LOC114683936 | 3.77 | 1.4E-48 |
| 118 | Arg1 | 4.36 | 1.9E-48 |
| 119 | LOC114686457 | 7.06 | 2.5E-48 |
| 120 | Glipr2 | 3.37 | 2.8E-48 |
| 121 | LOC114683921 | 4.50 | 1.9E-47 |
| 122 | Mmp8 | 9.51 | 2.7E-47 |
| 123 | CUNH1orf162 | 5.07 | 7.1E-47 |
| 501 | Catsper2 | 2.30 | 7.8E-14 |
| 502 | Dhrs13 | 2.08 | 9.8E-14 |
| 503 | Slc43a3 | 2.45 | 1.4E-13 |
| 504 | Siglecl1 | 3.99 | 1.6E-13 |
| 505 | Itgam | 2.64 | 1.6E-13 |
| 506 | Ctnna1 | 2.11 | 1.9E-13 |
| 507 | Adam30 | 2.91 | 2.0E-13 |
| 508 | Kirrel3 | 2.51 | 2.3E-13 |
| 509 | Dnah17 | 2.34 | 2.4E-13 |
| 510 | Bmx | 2.71 | 3.4E-13 |
| 511 | Camp | 7.51 | 4.1E-13 |
| 512 | Sdsl | 4.07 | 4.4E-13 |
| 513 | LOC114691549 | 8.26 | 4.6E-13 |
| 514 | Rnd3 | 3.41 | 4.8E-13 |
| 515 | Il10 | 4.75 | 8.8E-13 |
| 516 | Slc2a2 | 6.53 | 1.0E-12 |
| 517 | Ncam1 | 2.11 | 1.1E-12 |
| 518 | Scimp | 2.19 | 1.2E-12 |
| 519 | Scn2b | 3.37 | 1.2E-12 |
| 520 | LOC114688720 | 2.01 | 1.5E-12 |
| 521 | Uba7 | 2.09 | 1.9E-12 |
| 522 | Sptssb | 7.93 | 2.1E-12 |
| 523 | Lama2 | 3.38 | 2.1E-12 |
| 524 | LOC114688775 | 2.90 | 2.5E-12 |
| 525 | Dab2ip | 3.71 | 2.9E-12 |
| 526 | LOC114702847 | 2.33 | 3.1E-12 |
| 527 | LOC114706392 | 2.24 | 3.5E-12 |
| 528 | Rtp4 | 2.01 | 5.3E-12 |
| 529 | LOC114688398 | 5.65 | 5.3E-12 |
| 530 | Cdh23 | 2.13 | 5.7E-12 |
| 531 | Gtf2ird1 | 2.99 | 5.8E-12 |
| 532 | LOC114699623 | 2.80 | 6.7E-12 |
| 533 | Dab2 | 4.49 | 7.2E-12 |
| 534 | Pcdhgc3 | 2.34 | 7.3E-12 |
| 535 | Snx10 | 2.40 | 7.6E-12 |
| 536 | Il20ra | 4.87 | 1.2E-11 |
| 537 | LOC114700453 | 5.50 | 1.2E-11 |
| 538 | Irgc | 6.12 | 1.3E-11 |
| 539 | Tg | 3.35 | 1.3E-11 |
| 540 | LOC114705143 | 3.90 | 1.6E-11 |
| 541 | LOC114703775 | 2.25 | 2.2E-11 |
| 542 | LOC114686444 | 4.17 | 2.2E-11 |
| 543 | Sfxn5 | 2.73 | 2.2E-11 |
| 544 | Spry1 | 2.73 | 3.2E-11 |
| 545 | Dtx3l | 2.21 | 3.2E-11 |
| 546 | Gdpd1 | 3.14 | 3.2E-11 |
| 547 | Slc5a6 | 2.66 | 3.4E-11 |
| 548 | Csgalnact1 | 2.99 | 3.7E-11 |
| 549 | Mybpc3 | 2.99 | 3.9E-11 |
| 550 | LOC114703149 | 2.08 | 4.2E-11 |
| 551 | Herc6 | 3.30 | 5.4E-11 |
| 552 | LOC114706470 | 3.98 | 5.5E-11 |
| 553 | Csf1 | 2.16 | 5.7E-11 |
| 554 | Slc18a1 | 3.55 | 6.7E-11 |
| 555 | Cdk14 | 2.27 | 8.1E-11 |
| 556 | LOC114701075 | 4.95 | 8.6E-11 |
| 557 | Igf1r | 2.20 | 9.0E-11 |
| 558 | Lpo | 8.77 | 1.0E-10 |
| 559 | LOC114683579 | 2.89 | 1.4E-10 |
| 560 | LOC114680858 | 3.73 | 1.5E-10 |
| 561 | Gpat3 | 2.50 | 1.5E-10 |
| 562 | LOC114687126 | 3.07 | 1.6E-10 |
| 563 | Fam126b | 2.29 | 1.6E-10 |
| 564 | Rgs1 | 2.98 | 2.0E-10 |
| 565 | Tuft1 | 4.73 | 2.4E-10 |
| 566 | LOC114700347 | 2.83 | 2.4E-10 |
| 567 | Penk | 4.70 | 2.7E-10 |
| 568 | Itga7 | 4.30 | 2.7E-10 |
| 569 | LOC114694456 | 6.82 | 2.9E-10 |
| 570 | Olig2 | 3.66 | 3.3E-10 |
| 571 | LOC114691550 | 6.68 | 4.8E-10 |
| 572 | Pdzd4 | 2.85 | 5.2E-10 |
| 573 | Tnip3 | 3.27 | 5.2E-10 |
| 574 | Adgrg6 | 2.34 | 6.7E-10 |
| 575 | Nt5e | 2.81 | 9.0E-10 |
| 576 | LOC114708128 | 2.30 | 9.6E-10 |
| 577 | Pate1 | 6.59 | 1.0E-09 |
| 578 | Btbd3 | 2.02 | 1.6E-09 |
| 579 | Rnase10 | 6.29 | 1.9E-09 |
| 580 | Osmr | 4.78 | 2.0E-09 |
| 581 | Jag1 | 2.56 | 2.3E-09 |
| 582 | Irf7 | 2.19 | 2.3E-09 |
| 583 | LOC114688006 | 2.41 | 2.5E-09 |
| 584 | Smim3 | 2.18 | 2.7E-09 |
| 585 | LOC114683226 | 2.43 | 3.7E-09 |
| 586 | Lrrc3 | 4.35 | 3.9E-09 |
| 587 | Itga1 | 2.19 | 4.0E-09 |
| 588 | CUNH10orf105 | 2.72 | 4.0E-09 |
| 589 | LOC114683116 | 5.74 | 4.0E-09 |
| 590 | Orc1 | 5.39 | 4.4E-09 |
| 591 | LOC114681141 | 2.73 | 5.1E-09 |
| 592 | Slc13a5 | 8.60 | 5.8E-09 |
| 593 | Mst1r | 6.02 | 6.3E-09 |
| 594 | Pcdh19 | 4.22 | 6.7E-09 |
| 595 | Slamf1 | 2.05 | 6.8E-09 |
| 596 | Cpb2 | 5.52 | 1.0E-08 |
| 597 | Mmp10 | 6.34 | 1.1E-08 |
| 598 | CUNH9orf152 | 4.99 | 1.2E-08 |
| 599 | Kirrel1 | 3.78 | 1.3E-08 |
| 600 | LOC114681707 | 2.53 | 1.6E-08 |
| 601 | LOC114698905 | 6.57 | 1.7E-08 |
| 602 | LOC114707504 | 2.84 | 1.8E-08 |
| 603 | LOC114686188 | 2.16 | 2.0E-08 |
| 604 | LOC114707876 | 2.87 | 2.1E-08 |
| 605 | Muc1 | 3.80 | 2.4E-08 |
| 606 | Kcnk12 | 3.40 | 2.5E-08 |
| 607 | Cc2d2a | 2.41 | 2.7E-08 |
| 608 | Il6 | 5.51 | 2.8E-08 |
| 609 | Mrpl16 | 2.09 | 2.8E-08 |
| 610 | LOC114685662 | 3.34 | 2.9E-08 |
| 611 | Bicdl2 | 4.99 | 3.0E-08 |
| 612 | LOC114688611 | 2.86 | 3.1E-08 |
| 613 | Ncan | 2.55 | 3.3E-08 |
| 614 | Cpn1 | 2.20 | 3.3E-08 |
| 615 | C3 | 3.35 | 3.4E-08 |
| 616 | Emilin2 | 2.45 | 3.4E-08 |
| 617 | LOC114698011 | 2.21 | 4.0E-08 |
| 618 | LOC114686010 | 2.91 | 4.2E-08 |
| 619 | LOC114681897 | 2.09 | 4.2E-08 |
| 620 | LOC114706494 | 2.18 | 4.7E-08 |
| 621 | Trib3 | 2.44 | 4.9E-08 |
| 622 | LOC114684920 | 2.48 | 5.1E-08 |
| 623 | Alpk2 | 4.01 | 5.1E-08 |

Table S4.

| | | | |
|---|---|---|---|
| 124 | Pram1 | 4.41 | 7.8E-47 |
| 125 | Ppp1r3d | 3.89 | 8.2E-47 |
| 126 | Nadk | 2.04 | 1.3E-46 |
| 127 | Il1b | 4.08 | 1.5E-46 |
| 128 | Flot2 | 2.69 | 2.1E-46 |
| 129 | LOC114683484 | 4.09 | 2.2E-46 |
| 130 | Cxcr2 | 4.01 | 2.7E-46 |
| 131 | Kiaa0232 | 2.32 | 5.1E-46 |
| 132 | Mmp27 | 7.16 | 6.2E-46 |
| 133 | Aoah | 3.78 | 1.8E-45 |
| 134 | LOC114704122 | 4.16 | 3.3E-45 |
| 135 | Spi1 | 2.78 | 3.9E-45 |
| 136 | Crispld2 | 3.27 | 4.8E-45 |
| 137 | Steap1 | 8.73 | 5.5E-45 |
| 138 | Dhrs9 | 4.69 | 7.8E-45 |
| 139 | Lrg1 | 7.08 | 8.2E-45 |
| 140 | Zfp36 | 2.83 | 9.1E-45 |
| 141 | Adgrg3 | 4.21 | 1.4E-44 |
| 142 | Cd274 | 3.56 | 1.6E-44 |
| 143 | LOC114705177 | 5.41 | 3.4E-44 |
| 144 | Stxbp5 | 2.65 | 6.1E-44 |
| 145 | Hck | 2.89 | 6.6E-44 |
| 146 | Mapkapk2 | 2.08 | 6.6E-44 |
| 147 | Tlr4 | 3.58 | 7.5E-44 |
| 148 | LOC114707732 | 7.36 | 1.0E-43 |
| 149 | Capn12 | 8.08 | 1.8E-43 |
| 150 | Pim1 | 2.64 | 2.2E-43 |
| 151 | Cask | 2.89 | 3.5E-43 |
| 152 | Dennd5a | 2.46 | 6.1E-43 |
| 153 | Abtb2 | 2.83 | 8.0E-43 |
| 154 | LOC114703403 | 8.36 | 9.5E-43 |
| 155 | Alox5ap | 4.07 | 1.2E-42 |
| 156 | Snx18 | 3.17 | 2.6E-42 |
| 157 | Pgs1 | 2.53 | 3.1E-42 |
| 158 | Tbc1d2 | 3.86 | 3.7E-42 |
| 159 | Spidr | 2.46 | 7.1E-42 |
| 160 | Gas7 | 5.77 | 1.6E-41 |
| 161 | Atp11a | 2.74 | 2.0E-41 |
| 162 | Fosl2 | 2.61 | 2.2E-41 |
| 163 | Gadd45b | 3.31 | 2.6E-41 |
| 164 | Tspo | 3.26 | 2.9E-41 |
| 165 | Cd300lf | 3.76 | 3.0E-41 |
| 166 | Ubtd1 | 2.74 | 3.6E-41 |
| 167 | Pfkfb4 | 3.12 | 4.3E-41 |
| 168 | LOC114703402 | 7.74 | 4.9E-41 |
| 169 | S100a9 | 4.60 | 1.1E-40 |
| 170 | Sdr9c7 | 5.62 | 1.4E-40 |
| 171 | LOC114682475 | 2.78 | 1.5E-40 |
| 172 | Gca | 4.31 | 3.1E-40 |
| 173 | Litaf | 2.77 | 3.6E-40 |
| 174 | Ano10 | 2.38 | 4.0E-40 |
| 175 | Atp8b4 | 3.59 | 4.2E-40 |
| 176 | Upp1 | 4.32 | 4.5E-40 |
| 177 | LOC114689280 | 2.76 | 4.9E-40 |
| 178 | Sema6b | 6.31 | 6.5E-40 |
| 179 | Dusp16 | 2.69 | 6.9E-40 |
| 180 | Prdx5 | 3.29 | 1.1E-39 |
| 181 | Clec5a | 3.55 | 1.3E-39 |
| 182 | S100a8 | 4.82 | 1.8E-39 |
| 183 | Arid3a | 2.64 | 3.2E-39 |
| 184 | Tmem106a | 3.53 | 4.4E-39 |
| 185 | Ppp1r15b | 2.27 | 1.7E-38 |
| 186 | Slc8b1 | 2.33 | 1.7E-38 |
| 187 | Tyrobp | 2.82 | 2.1E-38 |
| 188 | Ppp1r3b | 4.63 | 3.1E-38 |
| 189 | Atp6v0c | 2.31 | 3.2E-38 |
| 190 | Cyfip1 | 2.21 | 3.9E-38 |
| 191 | Pfkfb3 | 2.98 | 4.2E-38 |
| 192 | Ugcg | 3.26 | 5.3E-38 |
| 193 | Pofut1 | 2.63 | 7.2E-38 |
| 194 | Ltb4r | 4.19 | 7.6E-38 |
| 195 | Tnfrsf1a | 2.64 | 7.7E-38 |
| 196 | Arhgap26 | 2.82 | 8.1E-38 |
| 197 | Irs2 | 2.59 | 1.4E-37 |
| 198 | Sds | 5.11 | 1.7E-37 |
| 199 | Samsn1 | 2.54 | 2.1E-37 |
| 200 | Tes | 2.33 | 2.9E-37 |
| 201 | LOC114701932 | 2.85 | 2.9E-37 |
| 202 | Antxr2 | 2.93 | 3.7E-37 |
| 203 | CUNH19orf38 | 3.13 | 5.6E-37 |
| 204 | LOC114687826 | 3.35 | 6.0E-37 |
| 205 | Ifngr2 | 2.29 | 7.4E-37 |
| 206 | LOC114691764 | 4.38 | 8.9E-37 |
| 207 | Osm | 3.97 | 1.9E-36 |
| 208 | Skap2 | 2.97 | 2.3E-36 |
| 209 | Sgms2 | 3.29 | 2.8E-36 |
| 210 | LOC114689150 | 2.90 | 3.3E-36 |
| 211 | Stat3 | 2.12 | 3.9E-36 |
| 212 | Il4r | 2.02 | 7.9E-36 |
| 213 | Msra | 2.94 | 8.4E-36 |
| 214 | Dach1 | 4.07 | 2.2E-35 |
| 215 | C5ar2 | 4.35 | 2.9E-35 |
| 216 | Basp1 | 3.55 | 3.1E-35 |
| 217 | Lyn | 2.03 | 3.2E-35 |
| 218 | Tmed5 | 2.13 | 6.7E-35 |
| 219 | Tns3 | 3.47 | 7.2E-35 |
| 220 | Tank | 2.04 | 7.9E-35 |
| 221 | Pnp | 3.12 | 9.9E-35 |
| 222 | LOC114702823 | 9.25 | 1.7E-34 |
| 223 | Pgd | 2.69 | 1.8E-34 |
| 224 | Jun | 2.89 | 2.0E-34 |
| 225 | Zc3h12a | 2.62 | 3.2E-34 |
| 226 | Gng10 | 2.67 | 4.0E-34 |
| 227 | Efhd2 | 2.08 | 4.0E-34 |
| 228 | Rab11fip5 | 2.94 | 4.5E-34 |
| 229 | Chst2 | 3.50 | 4.5E-34 |
| 230 | Bbs10 | 2.38 | 6.6E-34 |
| 231 | Batf3 | 4.82 | 7.4E-34 |
| 232 | Bst1 | 4.90 | 7.5E-34 |
| 233 | Irak3 | 3.72 | 8.1E-34 |
| 234 | Slc11a1 | 3.06 | 9.4E-34 |
| 235 | Mcemp1 | 4.42 | 1.7E-33 |
| 236 | Arap1 | 2.01 | 1.7E-33 |
| 237 | Fndc3b | 3.19 | 2.5E-33 |
| 238 | Selplg | 2.25 | 3.2E-33 |
| 239 | B4galt5 | 3.68 | 6.0E-33 |
| 240 | Fam160a1 | 3.65 | 6.5E-33 |
| 241 | Pacsin2 | 2.14 | 8.0E-33 |
| 242 | Cpne2 | 2.31 | 9.3E-33 |
| 243 | Dgat1 | 2.39 | 9.8E-33 |
| 244 | Plin5 | 6.09 | 1.0E-32 |
| 245 | Steap2 | 8.70 | 1.1E-32 |
| 246 | Ggt6 | 3.03 | 1.7E-32 |
| 247 | Lbp | 3.28 | 2.4E-32 |
| 248 | Tlr6 | 2.95 | 2.7E-32 |
| 249 | LOC114701921 | 2.73 | 3.3E-32 |
| 624 | Nkx2-3 | 3.60 | 5.1E-08 |
| 625 | Slc28a2 | 2.89 | 6.0E-08 |
| 626 | Tlr5 | 2.77 | 6.1E-08 |
| 627 | Vdr | 4.08 | 6.4E-08 |
| 628 | B4galt4 | 2.55 | 6.5E-08 |
| 629 | Tinagl1 | 4.15 | 6.8E-08 |
| 630 | LOC114685823 | 2.13 | 6.9E-08 |
| 631 | Kcnb1 | 2.32 | 7.0E-08 |
| 632 | LOC114691956 | 2.30 | 8.1E-08 |
| 633 | Gldn | 3.37 | 1.2E-07 |
| 634 | LOC114706443 | 2.99 | 1.2E-07 |
| 635 | LOC114687856 | 4.37 | 1.2E-07 |
| 636 | Snap23 | 2.04 | 1.2E-07 |
| 637 | LOC114685948 | 2.11 | 1.4E-07 |
| 638 | LOC114706456 | 3.59 | 1.6E-07 |
| 639 | Il1a | 5.06 | 2.0E-07 |
| 640 | LOC114689426 | 4.69 | 2.0E-07 |
| 641 | Lgals7 | 4.68 | 2.0E-07 |
| 642 | LOC114687827 | 2.07 | 2.2E-07 |
| 643 | LOC114710254 | 2.10 | 2.5E-07 |
| 644 | LOC114709510 | 3.32 | 2.8E-07 |
| 645 | C2 | 3.74 | 3.0E-07 |
| 646 | Ido1 | 6.04 | 3.3E-07 |
| 647 | Adcy10 | 3.66 | 3.3E-07 |
| 648 | Abcb11 | 4.77 | 3.6E-07 |
| 649 | Rec8 | 2.30 | 3.6E-07 |
| 650 | Pkd1l1 | 4.34 | 4.2E-07 |
| 651 | Cacna1i | 4.18 | 4.2E-07 |
| 652 | LOC114683922 | 4.89 | 4.4E-07 |
| 653 | Hhla2 | 2.38 | 5.1E-07 |
| 654 | LOC114686450 | 4.53 | 5.4E-07 |
| 655 | Krbox1 | 3.26 | 5.4E-07 |
| 656 | Lipg | 4.82 | 5.6E-07 |
| 657 | Mafb | 2.90 | 5.9E-07 |
| 658 | Prokr1 | 2.66 | 6.2E-07 |
| 659 | Tmem98 | 3.30 | 6.6E-07 |
| 660 | S1pr3 | 4.21 | 8.9E-07 |
| 661 | Igf2bp1 | 5.89 | 9.4E-07 |
| 662 | Dpysl5 | 3.27 | 9.4E-07 |
| 663 | Olr1 | 4.27 | 1.0E-06 |
| 664 | LOC114708750 | 2.62 | 1.1E-06 |
| 665 | Hspa6 | 2.93 | 1.1E-06 |
| 666 | Edn1 | 4.89 | 1.2E-06 |
| 667 | Gpr33 | 3.43 | 1.3E-06 |
| 668 | LOC114685411 | 5.07 | 1.4E-06 |
| 669 | LOC114692300 | 2.16 | 1.7E-06 |
| 670 | LOC114688405 | 3.94 | 1.8E-06 |
| 671 | LOC114689409 | 3.26 | 2.1E-06 |
| 672 | Msantd3 | 3.55 | 2.1E-06 |
| 673 | Neu3 | 2.51 | 2.2E-06 |
| 674 | Ckb | 2.38 | 2.3E-06 |
| 675 | LOC114701736 | 6.19 | 2.5E-06 |
| 676 | Pmch | 4.22 | 2.7E-06 |
| 677 | Hal | 3.99 | 3.7E-06 |
| 678 | Tex48 | 2.52 | 3.8E-06 |
| 679 | Cpa6 | 4.34 | 3.9E-06 |
| 680 | LOC114690363 | 4.79 | 4.1E-06 |
| 681 | Mepe | 4.98 | 4.1E-06 |
| 682 | Rnd1 | 3.34 | 4.5E-06 |
| 683 | LOC114688399 | 2.49 | 4.6E-06 |
| 684 | Tnfrsf9 | 3.49 | 5.0E-06 |
| 685 | LOC114697558 | 5.30 | 6.0E-06 |
| 686 | Mrc1 | 2.20 | 6.3E-06 |
| 687 | Serpine1 | 9.88 | 6.3E-06 |
| 688 | LOC114707727 | 3.16 | 6.4E-06 |
| 689 | Gramd1c | 2.40 | 7.1E-06 |
| 690 | Ptx3 | 5.68 | 7.2E-06 |
| 691 | Selp | 4.37 | 8.6E-06 |
| 692 | LOC114699997 | 3.39 | 8.9E-06 |
| 693 | Nnmt | 3.18 | 9.0E-06 |
| 694 | Scn3b | 3.02 | 9.1E-06 |
| 695 | Sele | 5.88 | 9.9E-06 |
| 696 | LOC114685691 | 3.33 | 1.0E-05 |
| 697 | LOC114707793 | 3.99 | 1.1E-05 |
| 698 | Coro2a | 3.08 | 1.3E-05 |
| 699 | Macc1 | 2.77 | 1.4E-05 |
| 700 | Oas3 | 2.01 | 1.6E-05 |
| 701 | Kif1a | 2.93 | 1.6E-05 |
| 702 | Col4a2 | 2.80 | 1.7E-05 |
| 703 | Parpbp | 2.67 | 1.9E-05 |
| 704 | Ap1s3 | 2.01 | 1.9E-05 |
| 705 | LOC114694459 | 5.68 | 2.1E-05 |
| 706 | Asb2 | 2.36 | 2.2E-05 |
| 707 | Shoc1 | 2.77 | 2.2E-05 |
| 708 | Capns2 | 2.23 | 2.3E-05 |
| 709 | Cttnbp2nl | 2.03 | 2.3E-05 |
| 710 | Stard13 | 2.05 | 2.3E-05 |
| 711 | Lrrc27 | 3.35 | 2.3E-05 |
| 712 | Zscan10 | 3.10 | 2.7E-05 |
| 713 | LOC114704705 | 2.32 | 2.8E-05 |
| 714 | Drc7 | 4.59 | 3.1E-05 |
| 715 | Tex101 | 4.50 | 3.1E-05 |
| 716 | LOC114708095 | 2.89 | 3.7E-05 |
| 717 | LOC114684566 | 3.10 | 3.9E-05 |
| 718 | Rph3a | 2.49 | 3.9E-05 |
| 719 | Csf2 | 4.65 | 3.9E-05 |
| 720 | LOC114689483 | 4.40 | 4.6E-05 |
| 721 | LOC114683925 | 3.07 | 4.9E-05 |
| 722 | Spred1 | 2.01 | 4.9E-05 |
| 723 | Cplx1 | 4.78 | 5.4E-05 |
| 724 | Pkhd1 | 2.74 | 6.6E-05 |
| 725 | Fbn2 | 5.02 | 7.9E-05 |
| 726 | Ccdc13 | 3.90 | 8.2E-05 |
| 727 | Ptgs2 | 2.20 | 8.5E-05 |
| 728 | Mybph | 4.59 | 8.9E-05 |
| 729 | Ubd | 5.20 | 9.2E-05 |
| 730 | Syt1 | 2.89 | 1.0E-04 |
| 731 | Spint1 | 3.45 | 1.0E-04 |
| 732 | Coch | 2.87 | 1.3E-04 |
| 733 | LOC114691127 | 2.24 | 1.3E-04 |
| 734 | Cep295nl | 2.43 | 1.4E-04 |
| 735 | Them5 | 4.89 | 1.4E-04 |
| 736 | Wnt6 | 3.24 | 1.4E-04 |
| 737 | LOC114684568 | 2.41 | 1.4E-04 |
| 738 | Crispld1 | 2.97 | 1.4E-04 |
| 739 | Cpa2 | 3.40 | 1.5E-04 |
| 740 | Efnb2 | 3.39 | 1.8E-04 |
| 741 | Ap1m2 | 3.54 | 1.8E-04 |
| 742 | Prex2 | 2.53 | 1.8E-04 |
| 743 | LOC114703340 | 3.56 | 2.0E-04 |
| 744 | Tgm2 | 3.75 | 2.1E-04 |
| 745 | LOC114700928 | 3.39 | 2.2E-04 |
| 746 | Trim16 | 2.58 | 2.2E-04 |
| 747 | LOC114701096 | 2.11 | 2.3E-04 |
| 748 | Nupr1 | 4.31 | 2.6E-04 |
| 749 | LOC114695394 | 3.34 | 2.6E-04 |

Table S4.

| | | | | | | | |
|---|---|---|---|---|---|---|---|
| 250 | Stxbp6 | 4.09 | 3.6E-32 | 750 | Col4a1 | 2.97 | 2.7E-04 |
| 251 | Tpd52 | 2.89 | 4.7E-32 | 751 | Bpi | 2.48 | 2.7E-04 |
| 252 | Cemip2 | 2.24 | 5.2E-32 | 752 | Nox1 | 2.48 | 2.7E-04 |
| 253 | Sdcbp | 2.62 | 6.5E-32 | 753 | Cxcl5 | 3.64 | 2.8E-04 |
| 254 | Atf6 | 2.79 | 6.7E-32 | 754 | LOC114696646 | 3.87 | 2.9E-04 |
| 255 | Prr13 | 2.06 | 6.7E-32 | 755 | Ppp1r17 | 2.26 | 2.9E-04 |
| 256 | Fez2 | 2.12 | 7.9E-32 | 756 | Tubb6 | 2.92 | 3.2E-04 |
| 257 | Lcn2 | 8.89 | 9.6E-32 | 757 | Creg2 | 5.52 | 3.3E-04 |
| 258 | Osbpl8 | 2.64 | 1.1E-31 | 758 | LOC114683802 | 3.10 | 3.4E-04 |
| 259 | S100a11 | 2.91 | 1.4E-31 | 759 | Hspb9 | 3.78 | 3.6E-04 |
| 260 | Tlr2 | 2.95 | 1.5E-31 | 760 | Adora2a | 2.05 | 3.7E-04 |
| 261 | Prok2 | 4.35 | 1.5E-31 | 761 | Pax8 | 4.24 | 4.0E-04 |
| 262 | Gyg1 | 2.53 | 2.3E-31 | 762 | Myo5b | 3.98 | 4.1E-04 |
| 263 | B3galt4 | 2.40 | 2.3E-31 | 763 | Prss23 | 4.58 | 4.3E-04 |
| 264 | Nfam1 | 3.40 | 2.5E-31 | 764 | C1r | 2.43 | 4.7E-04 |
| 265 | Fam217b | 2.88 | 2.7E-31 | 765 | Fam3b | 2.96 | 4.9E-04 |
| 266 | Erg | 4.30 | 2.7E-31 | 766 | Igf2bp3 | 2.10 | 5.4E-04 |
| 267 | Rnf144b | 3.25 | 2.7E-31 | 767 | LOC114695511 | 3.30 | 5.4E-04 |
| 268 | LOC114691130 | 3.55 | 4.3E-31 | 768 | Cxcl11 | 5.46 | 5.5E-04 |
| 269 | Csf3r | 3.58 | 4.6E-31 | 769 | LOC114709037 | 3.19 | 5.7E-04 |
| 270 | Fbxl5 | 2.05 | 4.7E-31 | 770 | C1s | 2.48 | 5.8E-04 |
| 271 | Lmnb1 | 2.53 | 4.7E-31 | 771 | LOC114700341 | 4.08 | 6.0E-04 |
| 272 | LOC114697855 | 5.03 | 5.4E-31 | 772 | Fam187b | 3.08 | 6.3E-04 |
| 273 | Hyal2 | 3.46 | 5.6E-31 | 773 | LOC114688010 | 2.27 | 6.3E-04 |
| 274 | Saa3 | 11.04 | 5.8E-31 | 774 | LOC114697034 | 4.46 | 6.3E-04 |
| 275 | Tirap | 2.26 | 6.4E-31 | 775 | LOC114695136 | 2.14 | 6.5E-04 |
| 276 | Net1 | 3.35 | 1.0E-30 | 776 | Arhgef25 | 2.08 | 6.6E-04 |
| 277 | Steap4 | 3.84 | 1.6E-30 | 777 | LOC114707719 | 3.00 | 6.7E-04 |
| 278 | Col4a3bp | 2.17 | 2.0E-30 | 778 | Hunk | 2.72 | 7.3E-04 |
| 279 | Cd14 | 3.97 | 3.2E-30 | 779 | Dagla | 2.80 | 7.3E-04 |
| 280 | Ccdc154 | 2.07 | 4.4E-30 | 780 | Pik3c2b | 2.33 | 7.5E-04 |
| 281 | Znf689 | 2.13 | 5.1E-30 | 781 | Zdhhc2 | 4.17 | 7.6E-04 |
| 282 | Fcer1g | 2.58 | 7.4E-30 | 782 | LOC114701896 | 3.20 | 7.7E-04 |
| 283 | Hes3 | 7.21 | 7.5E-30 | 783 | LOC114701362 | 3.35 | 7.8E-04 |
| 284 | St3gal5 | 2.66 | 1.1E-29 | 784 | Pgpep1l | 4.29 | 8.1E-04 |
| 285 | Map3k15 | 4.30 | 1.3E-29 | 785 | Plin4 | 3.47 | 8.2E-04 |
| 286 | LOC114705361 | 3.36 | 1.3E-29 | 786 | Il22ra2 | 3.65 | 8.4E-04 |
| 287 | Lst1 | 2.98 | 1.7E-29 | 787 | LOC114699080 | 3.53 | 8.5E-04 |
| 288 | LOC114683947 | 3.41 | 1.7E-29 | 788 | Pnoc | 2.58 | 8.9E-04 |
| 289 | Rin2 | 2.63 | 1.8E-29 | 789 | LOC114688409 | 2.43 | 9.3E-04 |
| 290 | Col15a1 | 3.80 | 2.0E-29 | 790 | Padi3 | 2.71 | 1.0E-03 |
| 291 | Nrros | 2.10 | 2.1E-29 | 791 | LOC114683149 | 2.39 | 1.1E-03 |
| 292 | Mapk13 | 2.75 | 2.3E-29 | 792 | Ppic | 4.70 | 1.1E-03 |
| 293 | Atrn | 2.60 | 2.3E-29 | 793 | Rab27b | 2.28 | 1.1E-03 |
| 294 | Nin | 2.60 | 3.0E-29 | 794 | LOC114687201 | 2.57 | 1.1E-03 |
| 295 | Hlx | 3.39 | 3.6E-29 | 795 | LOC114686649 | 4.88 | 1.1E-03 |
| 296 | Pih1d2 | 2.97 | 5.4E-29 | 796 | Msr1 | 2.94 | 1.2E-03 |
| 297 | Tnf | 2.88 | 6.8E-29 | 797 | Lin28a | 3.95 | 1.2E-03 |
| 298 | LOC114685334 | 4.49 | 8.3E-29 | 798 | LOC114687127 | 2.20 | 1.2E-03 |
| 299 | Parp9 | 2.29 | 9.4E-29 | 799 | Gpx2 | 3.07 | 1.3E-03 |
| 300 | Mafg | 2.12 | 1.0E-28 | 800 | LOC114686458 | 4.47 | 1.3E-03 |
| 301 | Rffl | 2.07 | 1.3E-28 | 801 | Cxcl16 | 2.32 | 1.3E-03 |
| 302 | Hs3st6 | 5.45 | 1.9E-28 | 802 | Nqo1 | 2.16 | 1.4E-03 |
| 303 | Pde2a | 2.65 | 2.2E-28 | 803 | Mertk | 3.14 | 1.4E-03 |
| 304 | Ifngr1 | 2.58 | 2.2E-28 | 804 | Ston1 | 2.85 | 1.5E-03 |
| 305 | Cdkn2d | 2.23 | 3.1E-28 | 805 | Tmem119 | 3.25 | 1.5E-03 |
| 306 | Mboat2 | 4.49 | 7.0E-28 | 806 | LOC114681143 | 2.45 | 1.6E-03 |
| 307 | Trpm2 | 2.21 | 7.2E-28 | 807 | Mmp14 | 3.25 | 1.6E-03 |
| 308 | Apobr | 2.79 | 8.1E-28 | 808 | LOC114683804 | 3.64 | 1.7E-03 |
| 309 | LOC114700344 | 5.75 | 9.0E-28 | 809 | Rhou | 2.93 | 1.7E-03 |
| 310 | Pdxk | 2.42 | 1.4E-27 | 810 | LOC114702828 | 3.54 | 1.7E-03 |
| 311 | LOC114686729 | 4.36 | 1.7E-27 | 811 | CUNH6orf132 | 2.55 | 1.7E-03 |
| 312 | CUNH15orf48 | 3.59 | 1.9E-27 | 812 | Procr | 2.34 | 1.8E-03 |
| 313 | Anxa2 | 2.16 | 2.6E-27 | 813 | LOC114681228 | 2.23 | 1.9E-03 |
| 314 | Samhd1 | 2.71 | 2.9E-27 | 814 | Has2 | 5.76 | 1.9E-03 |
| 315 | Rubcnl | 2.45 | 4.6E-27 | 815 | Dkk3 | 2.47 | 2.1E-03 |
| 316 | Hk2 | 2.61 | 4.9E-27 | 816 | Enpp3 | 3.42 | 2.2E-03 |
| 317 | Plbd1 | 3.15 | 5.1E-27 | 817 | Chst13 | 2.03 | 2.2E-03 |
| 318 | LOC114700968 | 2.25 | 7.4E-27 | 818 | LOC114706527 | 4.59 | 2.4E-03 |
| 319 | Mboat1 | 3.19 | 7.8E-27 | 819 | LOC114686717 | 2.22 | 2.4E-03 |
| 320 | LOC114706473 | 2.90 | 7.8E-27 | 820 | Apoa2 | 4.39 | 2.5E-03 |
| 321 | LOC114694962 | 2.47 | 9.1E-27 | 821 | Fam162b | 2.89 | 2.7E-03 |
| 322 | Gramd2a | 7.89 | 1.4E-26 | 822 | Angptl4 | 3.34 | 2.8E-03 |
| 323 | Hacd2 | 2.83 | 1.5E-26 | 823 | LOC114703218 | 2.28 | 2.8E-03 |
| 324 | Lpcat2 | 2.46 | 1.7E-26 | 824 | Plat | 2.72 | 2.9E-03 |
| 325 | Sort1 | 2.71 | 1.9E-26 | 825 | Myo1b | 2.18 | 3.0E-03 |
| 326 | Rab31 | 2.91 | 2.0E-26 | 826 | Dnajb3 | 3.77 | 3.0E-03 |
| 327 | Hif1a | 2.35 | 2.6E-26 | 827 | Cdh3 | 5.20 | 3.0E-03 |
| 328 | Rab20 | 3.94 | 2.7E-26 | 828 | Icam4 | 3.57 | 3.3E-03 |
| 329 | Foxred2 | 2.55 | 3.3E-26 | 829 | Plod2 | 3.18 | 3.3E-03 |
| 330 | Txnrd1 | 2.05 | 3.9E-26 | 830 | Tldc2 | 2.16 | 3.5E-03 |
| 331 | Slk | 2.28 | 6.0E-26 | 831 | Esr1 | 2.59 | 3.5E-03 |
| 332 | LOC114681131 | 2.95 | 7.9E-26 | 832 | Palld | 3.32 | 3.6E-03 |
| 333 | Jade1 | 2.43 | 9.3E-26 | 833 | LOC114689091 | 3.10 | 3.6E-03 |
| 334 | Gk | 2.38 | 1.2E-25 | 834 | Cpa4 | 2.80 | 3.9E-03 |
| 335 | Cpne8 | 4.83 | 1.2E-25 | 835 | LOC114681071 | 2.89 | 3.9E-03 |
| 336 | Fkbp5 | 2.37 | 1.5E-25 | 836 | Hhipl1 | 2.27 | 4.0E-03 |
| 337 | Agpat2 | 2.88 | 1.6E-25 | 837 | Iqgap3 | 2.21 | 4.1E-03 |
| 338 | Anpep | 4.81 | 2.0E-25 | 838 | Clvs2 | 2.84 | 4.1E-03 |
| 339 | Hcar2 | 4.83 | 2.2E-25 | 839 | Cdh26 | 3.99 | 4.2E-03 |
| 340 | Alox5 | 3.20 | 4.4E-25 | 840 | Sycp2 | 2.86 | 4.5E-03 |
| 341 | Lcp1 | 2.18 | 5.3E-25 | 841 | Kif21a | 3.84 | 4.7E-03 |
| 342 | Daglb | 2.23 | 5.8E-25 | 842 | C7 | 2.09 | 4.8E-03 |
| 343 | F2rl2 | 3.00 | 8.2E-25 | 843 | LOC114689300 | 2.27 | 4.9E-03 |
| 344 | LOC114685646 | 3.19 | 1.0E-24 | 844 | LOC114691605 | 3.83 | 5.0E-03 |
| 345 | Mefv | 2.90 | 2.0E-24 | 845 | Ccdc89 | 2.32 | 5.0E-03 |
| 346 | Nfe2l2 | 2.65 | 2.1E-24 | 846 | LOC114710170 | 2.10 | 5.5E-03 |
| 347 | Olig1 | 2.83 | 4.0E-24 | 847 | Garem2 | 3.04 | 5.5E-03 |
| 348 | Chit1 | 3.22 | 7.6E-24 | 848 | Rasa4b | 2.66 | 5.6E-03 |
| 349 | Ckap4 | 2.81 | 1.4E-23 | 849 | Ccl22 | 3.70 | 5.8E-03 |
| 350 | Tgfbr1 | 2.15 | 2.1E-23 | 850 | Vcam1 | 3.00 | 5.9E-03 |
| 351 | Tns4 | 3.24 | 2.7E-23 | 851 | CUNH6orf222 | 2.74 | 6.1E-03 |
| 352 | Tmem8a | 2.16 | 4.0E-23 | 852 | Rsph1 | 2.77 | 6.3E-03 |
| 353 | Gpr27 | 3.34 | 5.4E-23 | 853 | Scara5 | 4.25 | 6.5E-03 |
| 354 | Dgat2 | 4.66 | 6.5E-23 | 854 | Musk | 4.80 | 6.6E-03 |
| 355 | Csrnp1 | 2.44 | 1.2E-22 | 855 | LOC114691664 | 2.29 | 6.9E-03 |
| 356 | Pam | 3.15 | 1.3E-22 | 856 | Acvr1c | 4.19 | 7.0E-03 |
| 357 | Eml2 | 2.20 | 1.4E-22 | 857 | LOC114702066 | 3.07 | 7.0E-03 |
| 358 | Limk2 | 2.41 | 1.4E-22 | 858 | LOC114705271 | 2.08 | 7.1E-03 |
| 359 | LOC114685651 | 9.86 | 1.5E-22 | 859 | LOC114688415 | 3.10 | 7.5E-03 |
| 360 | Csf3 | 9.24 | 1.7E-22 | 860 | Vash2 | 2.38 | 7.7E-03 |
| 361 | LOC114694553 | 3.70 | 1.7E-22 | 861 | Npr3 | 3.71 | 8.0E-03 |
| 362 | Mxd1 | 2.73 | 2.4E-22 | 862 | Serping1 | 2.09 | 8.4E-03 |
| 363 | LOC114688764 | 4.63 | 3.6E-22 | 863 | Ccin | 2.20 | 8.7E-03 |
| 364 | Mpzl3 | 2.69 | 4.8E-22 | 864 | Fbxl13 | 2.04 | 9.0E-03 |
| 365 | Lyst | 2.35 | 5.9E-22 | 865 | LOC114696858 | 3.10 | 9.3E-03 |
| 366 | Ncf1 | 2.03 | 7.0E-22 | 866 | LOC114688043 | 3.15 | 9.3E-03 |
| 367 | Ada | 2.53 | 8.9E-22 | 867 | LOC114701719 | 5.45 | 9.4E-03 |
| 368 | LOC114688352 | 3.26 | 9.6E-22 | 868 | Slit2 | 2.40 | 9.5E-03 |
| 369 | Mgam | 3.72 | 1.1E-21 | 869 | Mfap2 | 2.42 | 9.7E-03 |
| 370 | Vcan | 3.55 | 1.3E-21 | 870 | LOC114682596 | 3.72 | 1.0E-02 |
| 371 | Timp1 | 6.17 | 1.4E-21 | 871 | Adig | 2.60 | 1.0E-02 |
| 372 | Pik3cg | 2.19 | 1.4E-21 | 872 | Ccm2l | 3.18 | 1.1E-02 |
| 373 | LOC114687526 | 3.27 | 1.6E-21 | 873 | S100a5 | 2.61 | 1.2E-02 |
| 374 | Vasp | 2.00 | 1.7E-21 | 874 | LOC114695086 | 2.11 | 1.2E-02 |
| 375 | Il18rap | 2.99 | 2.2E-21 | 875 | Prune2 | 2.16 | 1.2E-02 |

Table S4.

| | | | |
|---|---|---|---|
| 376 | Ms4a3 | 5.75 | 4.0E-21 |
| 377 | CUNH2orf69 | 2.43 | 4.4E-21 |
| 378 | Smpd1 | 2.06 | 5.0E-21 |
| 379 | LOC114699151 | 3.56 | 5.4E-21 |
| 380 | Snx7 | 5.87 | 8.5E-21 |
| 381 | Rassf4 | 4.06 | 1.1E-20 |
| 382 | Jdp2 | 2.44 | 1.2E-20 |
| 383 | Lrrk2 | 2.50 | 1.2E-20 |
| 384 | Cebpb | 3.88 | 1.2E-20 |
| 385 | LOC114683495 | 4.59 | 1.7E-20 |
| 386 | Ltf | 8.17 | 2.5E-20 |
| 387 | Slco4c1 | 3.57 | 2.8E-20 |
| 388 | Kcnj2 | 2.83 | 2.9E-20 |
| 389 | Alas1 | 2.33 | 3.3E-20 |
| 390 | LOC114697897 | 3.41 | 3.4E-20 |
| 391 | Ctsb | 2.02 | 4.9E-20 |
| 392 | Tkt | 2.01 | 5.3E-20 |
| 393 | Nhsl2 | 2.48 | 6.9E-20 |
| 394 | LOC114701735 | 9.84 | 9.4E-20 |
| 395 | Capn10 | 2.13 | 1.1E-19 |
| 396 | Stom | 2.59 | 1.2E-19 |
| 397 | Unc93b1 | 2.04 | 1.3E-19 |
| 398 | Galnt3 | 3.27 | 1.6E-19 |
| 399 | Cdc42ep2 | 3.18 | 1.7E-19 |
| 400 | Lhfpl2 | 5.24 | 1.9E-19 |
| 401 | Samd15 | 3.87 | 1.9E-19 |
| 402 | Gstcd | 2.27 | 2.1E-19 |
| 403 | Sh2b2 | 2.16 | 2.2E-19 |
| 404 | Pxylp1 | 5.90 | 2.5E-19 |
| 405 | Fcrlb | 3.92 | 2.7E-19 |
| 406 | Mtus1 | 2.31 | 2.8E-19 |
| 407 | Kiaa0513 | 2.10 | 2.8E-19 |
| 408 | Wnt5a | 6.63 | 6.2E-19 |
| 409 | Tiam2 | 3.03 | 7.0E-19 |
| 410 | LOC114707960 | 4.25 | 7.6E-19 |
| 411 | Ampd3 | 2.12 | 7.8E-19 |
| 412 | Mgarp | 9.44 | 1.0E-18 |
| 413 | Sqor | 2.68 | 1.4E-18 |
| 414 | Cxcl10 | 6.73 | 1.6E-18 |
| 415 | Cfb | 6.22 | 2.4E-18 |
| 416 | Eepd1 | 2.27 | 2.4E-18 |
| 417 | Nlrp3 | 2.34 | 2.8E-18 |
| 418 | Oasl | 3.52 | 3.1E-18 |
| 419 | Ostf1 | 2.02 | 4.0E-18 |
| 420 | Dusp1 | 2.87 | 4.2E-18 |
| 421 | LOC114698108 | 2.75 | 4.6E-18 |
| 422 | Dnajc25 | 2.71 | 4.7E-18 |
| 423 | Eif2ak2 | 2.94 | 4.7E-18 |
| 424 | Prss57 | 3.08 | 4.9E-18 |
| 425 | LOC114689119 | 2.39 | 5.2E-18 |
| 426 | LOC114687363 | 5.01 | 5.6E-18 |
| 427 | Slc31a2 | 2.52 | 5.8E-18 |
| 428 | Ctu1 | 2.43 | 6.5E-18 |
| 429 | Vegfa | 2.45 | 9.8E-18 |
| 430 | LOC114703874 | 3.81 | 9.8E-18 |
| 431 | Tmem45b | 3.34 | 1.0E-17 |
| 432 | Unc5a | 4.86 | 1.1E-17 |
| 433 | Egln3 | 2.88 | 1.1E-17 |
| 434 | C2cd2l | 2.09 | 1.3E-17 |
| 435 | Nod2 | 2.25 | 1.3E-17 |
| 436 | B3galnt1 | 6.10 | 1.6E-17 |
| 437 | Rgs17 | 3.43 | 1.8E-17 |
| 438 | Tnfrsf1b | 2.08 | 1.9E-17 |
| 439 | LOC114704110 | 2.57 | 2.0E-17 |
| 440 | LOC114696821 | 3.90 | 2.5E-17 |
| 441 | Icmt | 2.04 | 2.9E-17 |
| 442 | LOC114703401 | 5.13 | 4.2E-17 |
| 443 | Cd177 | 6.55 | 4.4E-17 |
| 444 | Fgd4 | 2.19 | 5.3E-17 |
| 445 | Rpl3l | 5.53 | 5.4E-17 |
| 446 | Dram1 | 4.38 | 5.7E-17 |
| 447 | Card9 | 2.05 | 6.0E-17 |
| 448 | Cstb | 2.22 | 6.5E-17 |
| 449 | Hbegf | 3.19 | 8.2E-17 |
| 450 | LOC114706804 | 2.27 | 9.6E-17 |
| 451 | LOC114689241 | 3.31 | 1.0E-16 |
| 452 | Clec4e | 2.91 | 1.2E-16 |
| 453 | Gstz1 | 2.33 | 1.3E-16 |
| 454 | LOC114707630 | 7.52 | 1.5E-16 |
| 455 | Hhip | 5.50 | 1.6E-16 |
| 456 | Itgb4 | 2.65 | 1.7E-16 |
| 457 | Tmem61 | 4.68 | 1.8E-16 |
| 458 | LOC114683944 | 5.59 | 1.8E-16 |
| 459 | Rtn4rl2 | 6.25 | 1.9E-16 |
| 460 | Cdc42ep1 | 4.46 | 2.9E-16 |
| 461 | LOC114687825 | 3.56 | 3.5E-16 |
| 462 | LOC114706495 | 2.32 | 5.3E-16 |
| 463 | LOC114709015 | 5.07 | 6.3E-16 |
| 464 | Rtp3 | 7.60 | 7.1E-16 |
| 465 | Tnfsf13b | 2.14 | 7.4E-16 |
| 466 | Pygl | 2.07 | 1.1E-15 |
| 467 | Manf | 2.23 | 1.1E-15 |
| 468 | Acpp | 2.10 | 1.6E-15 |
| 469 | Gpr179 | 3.16 | 1.8E-15 |
| 470 | LOC114693805 | 2.24 | 1.9E-15 |
| 471 | Itih5 | 2.05 | 2.0E-15 |
| 472 | Tnfaip2 | 3.08 | 2.0E-15 |
| 473 | LOC114694510 | 6.62 | 2.4E-15 |
| 474 | B4galt6 | 3.50 | 3.2E-15 |
| 475 | Mroh6 | 4.60 | 3.8E-15 |
| 476 | Hsd11b1 | 3.51 | 4.3E-15 |
| 477 | LOC114691981 | 2.19 | 5.8E-15 |
| 478 | Usp18 | 3.94 | 6.1E-15 |
| 479 | Ctsh | 2.15 | 7.9E-15 |
| 480 | Plk2 | 2.90 | 8.7E-15 |
| 481 | Lyz | 2.09 | 9.7E-15 |
| 482 | Icam1 | 3.39 | 1.5E-14 |
| 483 | Kif15 | 2.53 | 1.5E-14 |
| 484 | LOC114682065 | 2.41 | 1.6E-14 |
| 485 | Dusp23 | 2.38 | 1.7E-14 |
| 486 | Cmpk2 | 2.90 | 1.8E-14 |
| 487 | Thbs1 | 5.15 | 2.0E-14 |
| 488 | B4galnt2 | 3.96 | 2.0E-14 |
| 489 | LOC114685183 | 5.61 | 2.1E-14 |
| 490 | LOC114700507 | 7.38 | 2.2E-14 |
| 491 | Zc3h13 | 2.04 | 2.5E-14 |
| 492 | Dhx58 | 2.54 | 2.5E-14 |
| 493 | LOC114682211 | 2.18 | 3.2E-14 |
| 494 | LOC114681329 | 4.60 | 3.3E-14 |
| 495 | Pde6h | 3.60 | 3.3E-14 |
| 496 | Ca4 | 6.45 | 5.1E-14 |
| 497 | Cd33 | 5.24 | 5.3E-14 |
| 498 | LOC114691361 | 2.31 | 6.7E-14 |
| 499 | LOC114694511 | 7.27 | 6.9E-14 |
| 500 | Helz2 | 2.49 | 7.2E-14 |
| 876 | LOC114692999 | 2.32 | 1.3E-02 |
| 877 | LOC114681204 | 2.52 | 1.3E-02 |
| 878 | Slc1a3 | 3.22 | 1.4E-02 |
| 879 | Sdk2 | 2.11 | 1.4E-02 |
| 880 | LOC114689301 | 4.42 | 1.4E-02 |
| 881 | Abcc9 | 2.66 | 1.5E-02 |
| 882 | Antxr1 | 2.39 | 1.5E-02 |
| 883 | Spsb4 | 3.14 | 1.5E-02 |
| 884 | C1qa | 2.58 | 1.6E-02 |
| 885 | LOC114701409 | 4.00 | 1.6E-02 |
| 886 | LOC114688041 | 3.60 | 1.6E-02 |
| 887 | LOC114702196 | 2.73 | 1.6E-02 |
| 888 | Tacr1 | 3.54 | 1.6E-02 |
| 889 | LOC114695731 | 4.29 | 1.6E-02 |
| 890 | Arhgef39 | 2.02 | 1.7E-02 |
| 891 | LOC114693762 | 2.78 | 1.7E-02 |
| 892 | LOC114708552 | 4.11 | 1.7E-02 |
| 893 | Tln2 | 2.30 | 1.8E-02 |
| 894 | Dcstamp | 3.01 | 1.8E-02 |
| 895 | LOC114694347 | 3.63 | 1.9E-02 |
| 896 | Scube1 | 3.87 | 1.9E-02 |
| 897 | Soat2 | 3.73 | 1.9E-02 |
| 898 | LOC114683786 | 2.10 | 2.0E-02 |
| 899 | LOC114710178 | 3.00 | 2.0E-02 |
| 900 | LOC114681072 | 2.99 | 2.0E-02 |
| 901 | LOC114689864 | 3.05 | 2.0E-02 |
| 902 | Tbc1d30 | 2.17 | 2.0E-02 |
| 903 | Stra8 | 3.08 | 2.1E-02 |
| 904 | Cux2 | 2.39 | 2.2E-02 |
| 905 | Slc10a5 | 3.48 | 2.2E-02 |
| 906 | CUNH4orf19 | 2.07 | 2.2E-02 |
| 907 | Fads2 | 2.53 | 2.2E-02 |
| 908 | LOC114704476 | 2.47 | 2.2E-02 |
| 909 | LOC114681248 | 2.36 | 2.2E-02 |
| 910 | Ccdc80 | 2.36 | 2.2E-02 |
| 911 | Emcn | 2.16 | 2.3E-02 |
| 912 | Crppa | 2.09 | 2.4E-02 |
| 913 | Pla1a | 2.51 | 2.4E-02 |
| 914 | Txndc2 | 3.11 | 2.5E-02 |
| 915 | Foxg1 | 2.77 | 2.5E-02 |
| 916 | Dmrt3 | 2.43 | 2.5E-02 |
| 917 | LOC114695191 | 2.35 | 2.5E-02 |
| 918 | Itgb8 | 2.44 | 2.5E-02 |
| 919 | Galnt18 | 2.53 | 2.6E-02 |
| 920 | Mogat2 | 3.67 | 2.6E-02 |
| 921 | LOC114691141 | 3.93 | 2.7E-02 |
| 922 | Il27 | 2.59 | 2.8E-02 |
| 923 | LOC114709154 | 2.00 | 2.8E-02 |
| 924 | Cep126 | 2.24 | 2.9E-02 |
| 925 | CUNH3orf22 | 2.83 | 2.9E-02 |
| 926 | Ifng | 2.82 | 3.0E-02 |
| 927 | Acsbg1 | 2.80 | 3.0E-02 |
| 928 | Ace2 | 2.13 | 3.0E-02 |
| 929 | Olfml2b | 2.39 | 3.2E-02 |
| 930 | Kcnma1 | 3.53 | 3.4E-02 |
| 931 | LOC114699467 | 2.11 | 3.4E-02 |
| 932 | LOC114683803 | 4.31 | 3.4E-02 |
| 933 | Ccl19 | 3.94 | 3.6E-02 |
| 934 | LOC114700454 | 2.71 | 3.6E-02 |
| 935 | LOC114686921 | 3.15 | 3.6E-02 |
| 936 | Best1 | 2.43 | 3.7E-02 |
| 937 | Ism2 | 2.57 | 3.8E-02 |
| 938 | Flrt3 | 3.73 | 3.9E-02 |
| 939 | LOC114684575 | 2.42 | 3.9E-02 |
| 940 | LOC114683732 | 4.44 | 4.0E-02 |
| 941 | LOC114682982 | 3.56 | 4.1E-02 |
| 942 | Gfpt2 | 2.45 | 4.1E-02 |
| 943 | Adora1 | 2.54 | 4.1E-02 |
| 944 | Rab6b | 2.03 | 4.1E-02 |
| 945 | LOC114680593 | 2.43 | 4.2E-02 |
| 946 | LOC114705570 | 2.03 | 4.2E-02 |
| 947 | Ccl17 | 3.74 | 4.2E-02 |
| 948 | LOC114706467 | 3.13 | 4.5E-02 |
| 949 | CUNH6orf52 | 2.28 | 4.5E-02 |
| 950 | Foxa3 | 3.45 | 4.5E-02 |
| 951 | Sdcbp2 | 2.15 | 4.7E-02 |
| 952 | LOC114709229 | 2.31 | 4.7E-02 |
| 953 | Aplp1 | 2.25 | 4.7E-02 |
| 954 | Sdk1 | 2.11 | 4.9E-02 |
| 955 | Klf15 | 2.92 | 4.9E-02 |

**Table S5. DEGs in spleen of *P. leucopus*** Table S5

| rank | gene | log2 fold change | FDR |
|---|---|---|---|
| 1 | Sod2 | 4.44 | 0.00E+00 |
| 2 | Tnfaip3 | 3.66 | 1.49E-289 |
| 3 | Slc39a6 | 3.14 | 6.59E-256 |
| 4 | Socs3 | 3.49 | 7.55E-238 |
| 5 | Slc2a6 | 3.54 | 4.76E-234 |
| 6 | Pfkfb3 | 2.81 | 5.15E-234 |
| 7 | Slc15a3 | 2.38 | 1.09E-214 |
| 8 | Icam1 | 3.28 | 2.12E-214 |
| 9 | Dram1 | 4.73 | 3.91E-214 |
| 10 | LOC114709884 | 3.33 | 1.41E-191 |
| 11 | Sdc4 | 4.40 | 1.48E-183 |
| 12 | Il1rn | 5.79 | 5.85E-181 |
| 13 | Ccrl2 | 3.90 | 1.76E-175 |
| 14 | Tnip1 | 2.42 | 3.01E-175 |
| 15 | Trim16 | 2.57 | 2.27E-169 |
| 16 | Fpr2 | 6.66 | 2.49E-162 |
| 17 | Cxcl10 | 5.71 | 4.26E-162 |
| 18 | Nfkbiz | 2.99 | 2.29E-149 |
| 19 | Pla2g2a | 5.74 | 3.64E-148 |
| 20 | Cebpd | 3.51 | 2.41E-145 |
| 21 | LOC114681715 | 3.54 | 2.54E-145 |
| 22 | Rassf4 | 2.21 | 5.30E-145 |
| 23 | LOC114701735 | 8.18 | 1.88E-141 |
| 24 | Osmr | 3.42 | 3.17E-136 |
| 25 | Cx3cl1 | 5.48 | 4.60E-135 |
| 26 | Cfb | 3.58 | 2.79E-130 |
| 27 | LOC114707630 | 3.67 | 2.84E-130 |
| 28 | Nfkbia | 2.82 | 2.00E-129 |
| 29 | Nampt | 2.44 | 8.39E-128 |
| 30 | Pla1a | 3.90 | 5.14E-122 |
| 31 | Socs1 | 3.72 | 4.21E-120 |
| 32 | Tfpi | 3.04 | 7.98E-118 |
| 33 | LOC114693504 | 2.35 | 2.97E-114 |
| 34 | Ptpn13 | 2.41 | 4.02E-114 |
| 35 | Timp1 | 3.57 | 1.02E-110 |
| 36 | LOC114702542 | 2.14 | 2.50E-109 |
| 37 | Marcksl1 | 3.40 | 6.68E-109 |
| 38 | Batf2 | 2.90 | 3.60E-108 |
| 39 | Ifrd1 | 2.14 | 2.07E-107 |
| 40 | Fkbp5 | 2.16 | 2.60E-107 |
| 41 | LOC114694510 | 4.84 | 5.17E-107 |
| 42 | Bcl3 | 2.42 | 7.94E-106 |
| 43 | Trex1 | 2.32 | 1.40E-105 |
| 44 | Rnf19b | 2.12 | 1.63E-105 |
| 45 | Cdkn1a | 3.05 | 6.86E-105 |
| 46 | LOC114706392 | 4.09 | 1.62E-104 |
| 47 | LOC114681707 | 3.84 | 8.46E-104 |
| 48 | Cd274 | 2.65 | 2.08E-102 |
| 49 | Samhd1 | 3.03 | 4.54E-102 |
| 50 | Serpine1 | 4.87 | 2.25E-101 |
| 51 | Tlr2 | 2.82 | 2.00E-100 |
| 52 | Slpi | 7.37 | 3.25E-98 |
| 53 | Tnf | 3.56 | 3.54E-97 |
| 54 | Tfec | 2.43 | 2.66E-96 |
| 55 | Cd40 | 2.82 | 1.38E-95 |
| 56 | Srgn | 2.48 | 8.18E-95 |
| 57 | Ifih1 | 4.17 | 2.04E-94 |
| 58 | Cebpb | 2.64 | 1.95E-93 |
| 59 | Il10 | 5.17 | 7.40E-92 |
| 60 | Tanc1 | 2.08 | 7.61E-92 |
| 61 | Csrnp1 | 2.56 | 8.40E-92 |
| 62 | Wars | 2.76 | 3.94E-90 |
| 63 | Batf | 2.98 | 2.38E-89 |
| 64 | Tap1 | 2.19 | 4.77E-89 |
| 65 | Mmp3 | 7.51 | 6.89E-88 |
| 66 | Il6 | 5.50 | 1.77E-87 |
| 67 | Pi15 | 5.55 | 9.00E-87 |
| 68 | Tnfaip2 | 3.09 | 2.43E-86 |
| 69 | Zc3h12a | 2.25 | 1.58E-84 |
| 70 | Nfil3 | 3.63 | 2.18E-84 |
| 71 | Rab32 | 2.24 | 9.99E-84 |
| 72 | Pml | 2.19 | 1.46E-83 |
| 73 | LOC114708004 | 2.28 | 3.41E-83 |
| 74 | Arl5c | 2.56 | 1.17E-82 |
| 75 | Nnmt | 3.95 | 7.06E-82 |
| 76 | Atf3 | 3.71 | 3.56E-81 |
| 77 | LOC114701736 | 6.42 | 5.54E-81 |
| 78 | Ibsp | 7.43 | 1.49E-80 |
| 79 | Crispld2 | 3.19 | 3.95E-80 |
| 80 | Ppa1 | 2.18 | 1.74E-79 |
| 81 | Dtx3l | 2.60 | 2.44E-79 |
| 82 | Steap4 | 5.51 | 2.82E-79 |
| 83 | Stat2 | 2.16 | 6.80E-79 |
| 84 | Fbxo27 | 2.45 | 1.68E-78 |
| 85 | Sphk1 | 2.73 | 2.16E-78 |
| 86 | Cd44 | 2.01 | 9.59E-78 |
| 87 | Acod1 | 7.32 | 3.62E-77 |
| 88 | Snx10 | 2.63 | 1.52E-76 |
| 89 | LOC114688414 | 4.22 | 1.49E-75 |
| 90 | LOC114689091 | 3.60 | 2.34E-75 |
| 91 | Mefv | 3.47 | 4.72E-75 |
| 92 | Upp1 | 2.96 | 1.01E-73 |
| 281 | Chp2 | 2.13 | 3.73E-21 |
| 282 | LOC114683116 | 6.36 | 6.14E-21 |
| 283 | Mmp8 | 5.09 | 1.53E-20 |
| 284 | Mmp10 | 4.69 | 2.23E-20 |
| 285 | Chi3l1 | 3.52 | 2.72E-20 |
| 286 | Hhla2 | 2.32 | 3.29E-20 |
| 287 | LOC114704463 | 2.25 | 3.98E-20 |
| 288 | LOC114703406 | 3.14 | 5.25E-20 |
| 289 | Tgm1 | 2.96 | 5.69E-20 |
| 290 | LOC114707728 | 5.28 | 7.48E-20 |
| 291 | Zmat1 | 2.19 | 7.78E-20 |
| 292 | Izumo1 | 2.97 | 1.66E-19 |
| 293 | LOC114684570 | 4.00 | 2.26E-19 |
| 294 | Gfra1 | 2.18 | 3.24E-19 |
| 295 | Nox1 | 3.31 | 3.88E-19 |
| 296 | Dgat2 | 2.66 | 4.93E-19 |
| 297 | Ccr8 | 2.09 | 5.33E-19 |
| 298 | LOC114681329 | 4.00 | 1.02E-18 |
| 299 | Isg15 | 3.93 | 1.22E-18 |
| 300 | Il1a | 2.53 | 1.67E-18 |
| 301 | Ntrk1 | 2.18 | 1.97E-18 |
| 302 | LOC114687526 | 2.58 | 5.78E-18 |
| 303 | Rasl11b | 2.06 | 8.19E-18 |
| 304 | Lrg1 | 3.94 | 1.45E-17 |
| 305 | LOC114691641 | 2.60 | 1.46E-17 |
| 306 | Rec8 | 2.26 | 1.59E-17 |
| 307 | Klf15 | 2.08 | 2.00E-17 |
| 308 | LOC114706978 | 2.41 | 2.06E-17 |
| 309 | Loxl2 | 2.15 | 2.39E-17 |
| 310 | Selp | 2.10 | 2.99E-17 |
| 311 | Arg1 | 2.79 | 3.51E-17 |
| 312 | LOC114703392 | 2.07 | 3.62E-17 |
| 313 | CUNH15orf48 | 2.72 | 4.97E-17 |
| 314 | LOC114681897 | 3.33 | 5.98E-17 |
| 315 | LOC114685334 | 3.22 | 7.68E-17 |
| 316 | Gfpt2 | 2.96 | 1.22E-16 |
| 317 | Fam71a | 4.10 | 1.24E-16 |
| 318 | LOC114703402 | 3.31 | 1.62E-16 |
| 319 | Mamstr | 2.01 | 1.68E-16 |
| 320 | Cntfr | 2.49 | 2.11E-16 |
| 321 | Ccl17 | 4.19 | 2.12E-16 |
| 322 | Mlph | 2.83 | 2.26E-16 |
| 323 | LOC114701896 | 3.86 | 2.45E-16 |
| 324 | Clec4e | 2.53 | 2.87E-16 |
| 325 | Aoah | 2.27 | 3.59E-16 |
| 326 | Tnfsf18 | 4.75 | 4.09E-16 |
| 327 | Ppfia2 | 2.43 | 4.41E-16 |
| 328 | Ooep | 3.02 | 5.74E-16 |
| 329 | CUNH6orf222 | 3.05 | 9.54E-16 |
| 330 | LOC114698852 | 2.26 | 1.51E-15 |
| 331 | LOC114697821 | 2.34 | 2.61E-15 |
| 332 | LOC114684798 | 2.67 | 3.16E-15 |
| 333 | Ccl5 | 2.44 | 3.83E-15 |
| 334 | Klk9 | 3.46 | 4.12E-15 |
| 335 | Irx3 | 3.30 | 6.99E-15 |
| 336 | LOC114692262 | 3.49 | 9.32E-15 |
| 337 | LOC114688093 | 3.11 | 1.03E-14 |
| 338 | Csrp3 | 5.91 | 1.75E-14 |
| 339 | Sgms2 | 2.28 | 2.14E-14 |
| 340 | LOC114699623 | 2.75 | 2.53E-14 |
| 341 | LOC114690678 | 2.08 | 4.60E-14 |
| 342 | Ca4 | 3.97 | 5.26E-14 |
| 343 | Nppc | 2.78 | 6.13E-14 |
| 344 | Kcng3 | 2.01 | 6.81E-14 |
| 345 | LOC114703403 | 3.70 | 7.10E-14 |
| 346 | Sds | 3.27 | 8.50E-14 |
| 347 | LOC114708913 | 2.02 | 1.51E-13 |
| 348 | Olfm4 | 3.29 | 3.33E-13 |
| 349 | Tnc | 3.38 | 6.58E-13 |
| 350 | LOC114684565 | 5.61 | 6.64E-13 |
| 351 | LOC114708095 | 3.93 | 7.29E-13 |
| 352 | Mmp12 | 5.99 | 7.59E-13 |
| 353 | Col6a5 | 2.51 | 9.34E-13 |
| 354 | Oas2 | 2.63 | 1.27E-12 |
| 355 | LOC114707940 | 2.56 | 1.70E-12 |
| 356 | Ccl22 | 2.41 | 2.12E-12 |
| 357 | LOC114681726 | 3.77 | 2.48E-12 |
| 358 | Clstn3 | 2.67 | 2.54E-12 |
| 359 | LOC114681724 | 3.06 | 2.76E-12 |
| 360 | Nts | 4.30 | 5.22E-12 |
| 361 | Cst7 | 2.68 | 6.04E-12 |
| 362 | LOC114689300 | 3.50 | 6.39E-12 |
| 363 | LOC114704347 | 3.50 | 7.58E-12 |
| 364 | CUNH11orf91 | 2.15 | 9.43E-12 |
| 365 | LOC114707719 | 6.25 | 1.14E-11 |
| 366 | Meox1 | 2.38 | 1.50E-11 |
| 367 | Lipg | 3.78 | 1.89E-11 |
| 368 | Slc13a5 | 2.78 | 2.02E-11 |
| 369 | Fut1 | 3.48 | 5.21E-11 |
| 370 | Sdr9c7 | 3.47 | 6.31E-11 |
| 371 | S100a9 | 2.25 | 7.33E-11 |
| 372 | LOC114681565 | 3.08 | 7.45E-11 |

Table S5

| | | | |
|---|---|---|---|
| 93 | Rars | 2.83 | 2.77E-73 |
| 94 | Fn1 | 2.66 | 3.18E-73 |
| 95 | LOC114707400 | 4.60 | 8.12E-72 |
| 96 | Xaf1 | 2.55 | 1.09E-71 |
| 97 | Septin5 | 2.25 | 2.54E-71 |
| 98 | Cystm1 | 2.99 | 2.76E-71 |
| 99 | LOC114680957 | 2.35 | 9.06E-71 |
| 100 | Ccl20 | 5.91 | 2.38E-69 |
| 101 | LOC114691127 | 2.33 | 5.44E-69 |
| 102 | Pstpip2 | 2.01 | 7.31E-69 |
| 103 | LOC114682175 | 2.21 | 1.44E-68 |
| 104 | Cxcl16 | 2.65 | 4.32E-68 |
| 105 | Samd9 | 2.36 | 6.37E-68 |
| 106 | Vcan | 4.13 | 8.53E-68 |
| 107 | Agpat4 | 2.26 | 1.67E-67 |
| 108 | Il1b | 3.48 | 6.87E-67 |
| 109 | LOC114688413 | 5.40 | 2.28E-66 |
| 110 | Tnip3 | 4.71 | 4.32E-66 |
| 111 | Neurl3 | 3.71 | 4.90E-66 |
| 112 | Gadd45b | 2.57 | 6.48E-66 |
| 113 | Pnp | 2.87 | 7.78E-66 |
| 114 | Slc30a1 | 2.18 | 1.12E-65 |
| 115 | Isg20 | 4.07 | 1.62E-65 |
| 116 | Cgas | 2.64 | 1.63E-65 |
| 117 | Zbtb16 | 3.37 | 1.21E-64 |
| 118 | Parp14 | 2.40 | 2.45E-64 |
| 119 | Maff | 2.26 | 5.53E-64 |
| 120 | Pim1 | 2.49 | 6.99E-64 |
| 121 | LOC114686891 | 2.43 | 8.92E-64 |
| 122 | Xdh | 2.30 | 1.13E-63 |
| 123 | Stx11 | 2.28 | 1.74E-62 |
| 124 | Steap2 | 6.38 | 9.96E-62 |
| 125 | LOC114689864 | 3.61 | 1.31E-61 |
| 126 | Csf3 | 8.15 | 3.33E-61 |
| 127 | LOC114694511 | 6.32 | 3.67E-61 |
| 128 | Ptx3 | 8.12 | 5.90E-61 |
| 129 | Il27 | 6.11 | 6.60E-61 |
| 130 | Prss23 | 2.18 | 6.60E-61 |
| 131 | LOC114697855 | 4.10 | 1.43E-60 |
| 132 | Rsad2 | 6.78 | 2.28E-59 |
| 133 | Cflar | 2.19 | 2.29E-59 |
| 134 | Parp9 | 2.05 | 4.79E-59 |
| 135 | LOC114706381 | 2.38 | 3.09E-57 |
| 136 | Ddx58 | 3.22 | 1.71E-56 |
| 137 | Arid5a | 2.31 | 1.96E-56 |
| 138 | Icam4 | 3.54 | 2.99E-56 |
| 139 | Thbs1 | 2.52 | 1.10E-55 |
| 140 | Tnfaip6 | 4.32 | 4.26E-55 |
| 141 | Rtp3 | 4.57 | 5.00E-54 |
| 142 | LOC114681718 | 2.11 | 1.29E-53 |
| 143 | LOC114702823 | 6.87 | 5.19E-52 |
| 144 | LOC114706400 | 2.66 | 1.22E-51 |
| 145 | Slamf1 | 2.90 | 1.63E-51 |
| 146 | LOC114691550 | 4.98 | 2.07E-51 |
| 147 | LOC114698905 | 5.13 | 1.58E-50 |
| 148 | Parp12 | 2.42 | 2.34E-50 |
| 149 | Helz2 | 3.05 | 2.41E-50 |
| 150 | Plat | 2.61 | 1.26E-49 |
| 151 | Cemip2 | 2.26 | 2.22E-49 |
| 152 | Stra6 | 3.27 | 2.95E-48 |
| 153 | Cxcl11 | 8.37 | 5.20E-48 |
| 154 | Tubb6 | 2.88 | 9.09E-48 |
| 155 | Hp | 3.10 | 1.31E-47 |
| 156 | Slc28a2 | 3.60 | 1.33E-47 |
| 157 | Fjx1 | 3.22 | 1.83E-47 |
| 158 | Csf2 | 7.83 | 2.49E-47 |
| 159 | LOC114690079 | 3.54 | 2.62E-47 |
| 160 | LOC114697861 | 2.25 | 3.12E-47 |
| 161 | Hcar2 | 4.71 | 3.64E-47 |
| 162 | Eif2ak2 | 3.27 | 4.01E-47 |
| 163 | Nid2 | 2.72 | 8.28E-47 |
| 164 | Rnf213 | 3.27 | 1.60E-46 |
| 165 | Oasl | 5.78 | 8.27E-46 |
| 166 | LOC114681719 | 2.56 | 9.80E-46 |
| 167 | Lgals9 | 2.46 | 9.87E-46 |
| 168 | Dusp1 | 2.21 | 1.60E-45 |
| 169 | Uba7 | 2.93 | 7.34E-45 |
| 170 | Calhm6 | 2.08 | 5.87E-44 |
| 171 | Cmpk2 | 4.88 | 3.31E-43 |
| 172 | LOC114688405 | 5.77 | 5.98E-43 |
| 173 | LOC114707727 | 4.83 | 7.22E-43 |
| 174 | Sdc1 | 2.39 | 9.08E-43 |
| 175 | Gpr33 | 4.49 | 1.38E-42 |
| 176 | LOC114689891 | 2.06 | 2.09E-42 |
| 177 | LOC114684568 | 4.96 | 2.78E-42 |
| 178 | Rnd1 | 3.91 | 3.38E-42 |
| 179 | Fpr1 | 3.04 | 3.98E-42 |
| 180 | LOC114688398 | 4.20 | 6.20E-42 |
| 181 | Dhx58 | 3.46 | 2.73E-41 |
| 182 | LOC114687018 | 2.30 | 3.07E-41 |
| 183 | Ubd | 4.81 | 4.19E-41 |
| 184 | Sema3a | 3.46 | 7.29E-41 |
| 185 | LOC114701399 | 2.02 | 5.89E-40 |
| 186 | Nupr1 | 3.67 | 6.41E-40 |
| 187 | Peg10 | 3.52 | 3.26E-39 |
| 373 | LOC114692453 | 3.05 | 1.22E-10 |
| 374 | Ifng | 4.37 | 1.56E-10 |
| 375 | LOC114681904 | 3.21 | 1.67E-10 |
| 376 | Anxa2r | 5.74 | 2.09E-10 |
| 377 | Krt8 | 2.13 | 3.26E-10 |
| 378 | Smtnl1 | 2.18 | 4.09E-10 |
| 379 | Ms4a3 | 3.16 | 4.11E-10 |
| 380 | Hpx | 2.27 | 4.21E-10 |
| 381 | Medag | 3.34 | 4.26E-10 |
| 382 | LOC114688826 | 2.18 | 4.52E-10 |
| 383 | LOC114702286 | 2.49 | 6.97E-10 |
| 384 | LOC114684569 | 3.50 | 7.13E-10 |
| 385 | LOC114698011 | 2.18 | 7.33E-10 |
| 386 | Fgf21 | 4.06 | 1.05E-09 |
| 387 | Epha4 | 2.67 | 1.49E-09 |
| 388 | Prok2 | 2.71 | 2.48E-09 |
| 389 | LOC114681165 | 2.98 | 2.57E-09 |
| 390 | LOC114696384 | 2.31 | 2.71E-09 |
| 391 | Srxn1 | 2.43 | 3.47E-09 |
| 392 | LOC114681708 | 2.67 | 3.63E-09 |
| 393 | LOC114694347 | 3.03 | 3.72E-09 |
| 394 | Nexmif | 2.10 | 4.40E-09 |
| 395 | Rnase10 | 4.15 | 5.39E-09 |
| 396 | LOC114683802 | 3.52 | 5.91E-09 |
| 397 | Ddx43 | 2.05 | 6.32E-09 |
| 398 | Eppk1 | 2.06 | 7.08E-09 |
| 399 | LOC114684575 | 5.03 | 1.19E-08 |
| 400 | Prrg4 | 2.16 | 1.23E-08 |
| 401 | Ido2 | 3.51 | 1.46E-08 |
| 402 | Tph1 | 2.47 | 1.47E-08 |
| 403 | LOC114685652 | 5.02 | 1.51E-08 |
| 404 | Nim1k | 2.64 | 1.84E-08 |
| 405 | Dio3 | 2.35 | 2.22E-08 |
| 406 | LOC114706385 | 2.80 | 2.56E-08 |
| 407 | LOC114690782 | 3.15 | 3.64E-08 |
| 408 | LOC114685651 | 4.29 | 4.27E-08 |
| 409 | Stc1 | 2.13 | 5.13E-08 |
| 410 | Mcidas | 4.35 | 5.17E-08 |
| 411 | Gpr39 | 2.29 | 5.41E-08 |
| 412 | Slamf9 | 2.07 | 5.64E-08 |
| 413 | Zdhhc2 | 2.36 | 6.81E-08 |
| 414 | LOC114708750 | 2.27 | 9.41E-08 |
| 415 | LOC114681164 | 2.46 | 1.51E-07 |
| 416 | Btbd16 | 3.25 | 2.21E-07 |
| 417 | Il22 | 6.54 | 2.50E-07 |
| 418 | Ano2 | 4.02 | 2.52E-07 |
| 419 | Prune2 | 2.18 | 2.80E-07 |
| 420 | Tgm6 | 2.75 | 2.87E-07 |
| 421 | Dmp1 | 2.96 | 3.03E-07 |
| 422 | LOC114681905 | 3.09 | 3.98E-07 |
| 423 | Sct | 2.50 | 5.56E-07 |
| 424 | LOC114692731 | 2.10 | 6.29E-07 |
| 425 | Mgam | 2.77 | 7.70E-07 |
| 426 | LOC114710025 | 3.49 | 8.22E-07 |
| 427 | Igf2bp1 | 2.65 | 8.41E-07 |
| 428 | LOC114682933 | 2.55 | 8.83E-07 |
| 429 | LOC114697856 | 2.57 | 1.08E-06 |
| 430 | LOC114689273 | 2.70 | 1.11E-06 |
| 431 | LOC114683804 | 4.38 | 1.14E-06 |
| 432 | Oas3 | 2.41 | 1.16E-06 |
| 433 | LOC114685183 | 5.08 | 1.23E-06 |
| 434 | LOC114694553 | 2.04 | 1.52E-06 |
| 435 | LOC114680639 | 2.80 | 1.57E-06 |
| 436 | LOC114683786 | 2.34 | 1.66E-06 |
| 437 | LOC114701719 | 2.99 | 1.79E-06 |
| 438 | LOC114697185 | 2.46 | 1.80E-06 |
| 439 | LOC114684572 | 2.87 | 1.93E-06 |
| 440 | Il17rb | 2.37 | 2.15E-06 |
| 441 | Pkhd1 | 2.06 | 2.39E-06 |
| 442 | Flrt3 | 2.32 | 3.33E-06 |
| 443 | Dppa5 | 3.00 | 3.41E-06 |
| 444 | Musk | 2.66 | 3.70E-06 |
| 445 | LOC114690609 | 2.16 | 4.32E-06 |
| 446 | LOC114710254 | 2.15 | 4.44E-06 |
| 447 | Nos2 | 3.79 | 5.92E-06 |
| 448 | LOC114697129 | 4.06 | 6.53E-06 |
| 449 | Mst1r | 2.36 | 8.47E-06 |
| 450 | Chrna2 | 2.53 | 1.02E-05 |
| 451 | Cartpt | 4.84 | 1.24E-05 |
| 452 | LOC114696647 | 4.99 | 1.41E-05 |
| 453 | LOC114697570 | 3.58 | 1.41E-05 |
| 454 | Il31ra | 2.13 | 1.47E-05 |
| 455 | LOC114692696 | 2.52 | 1.52E-05 |
| 456 | LOC114685648 | 2.89 | 1.69E-05 |
| 457 | LOC114681470 | 2.70 | 1.70E-05 |
| 458 | CUNH3orf80 | 2.20 | 1.74E-05 |
| 459 | Dio2 | 2.53 | 2.10E-05 |
| 460 | Camk2b | 2.29 | 2.65E-05 |
| 461 | LOC114706384 | 2.94 | 2.71E-05 |
| 462 | Plekhs1 | 2.27 | 4.53E-05 |
| 463 | Fbxo39 | 2.18 | 4.94E-05 |
| 464 | LOC114687407 | 3.37 | 5.08E-05 |
| 465 | Lpo | 3.60 | 5.37E-05 |
| 466 | Il19 | 2.74 | 5.73E-05 |
| 467 | Shox2 | 2.61 | 5.85E-05 |

Table S5

| | | | |
|---|---|---|---|
| 188 | Adamts4 | 3.37 | 3.91E-39 |
| 189 | Cldn1 | 2.74 | 2.96E-38 |
| 190 | Ido1 | 3.92 | 3.17E-38 |
| 191 | LOC114691549 | 3.97 | 4.83E-38 |
| 192 | LOC114700540 | 7.92 | 1.34E-37 |
| 193 | Kif1a | 3.00 | 3.66E-37 |
| 194 | Mitd1 | 2.07 | 1.31E-36 |
| 195 | Il15 | 2.32 | 1.73E-36 |
| 196 | Apold1 | 3.98 | 2.00E-36 |
| 197 | Lyve1 | 3.58 | 2.30E-36 |
| 198 | LOC114686894 | 2.89 | 3.89E-36 |
| 199 | Edn1 | 3.49 | 4.98E-36 |
| 200 | Ccn4 | 2.40 | 7.69E-36 |
| 201 | Alpk1 | 2.33 | 3.38E-35 |
| 202 | Tfpi2 | 4.23 | 4.72E-35 |
| 203 | Il15ra | 2.83 | 6.33E-35 |
| 204 | Herc6 | 4.67 | 8.77E-35 |
| 205 | Fscn1 | 2.28 | 9.46E-35 |
| 206 | LOC114700341 | 4.07 | 1.74E-34 |
| 207 | C1rl | 2.40 | 2.54E-34 |
| 208 | Depp1 | 2.45 | 3.56E-34 |
| 209 | LOC114707732 | 5.58 | 8.50E-34 |
| 210 | LOC114697860 | 2.34 | 6.02E-33 |
| 211 | LOC114700347 | 3.66 | 6.54E-33 |
| 212 | LOC114698259 | 2.18 | 2.20E-32 |
| 213 | LOC114694372 | 5.45 | 2.68E-32 |
| 214 | Samd9l | 2.03 | 3.86E-32 |
| 215 | Ackr1 | 4.54 | 8.61E-32 |
| 216 | Ifi44 | 2.00 | 1.14E-31 |
| 217 | Plaur | 2.10 | 1.32E-31 |
| 218 | Olr1 | 4.32 | 1.66E-31 |
| 219 | Gramd2a | 4.20 | 1.76E-31 |
| 220 | Il1r2 | 4.20 | 1.96E-31 |
| 221 | LOC114690951 | 2.13 | 1.98E-31 |
| 222 | Cxcl5 | 2.99 | 2.18E-31 |
| 223 | Ube2l6 | 2.71 | 2.56E-31 |
| 224 | LOC114688409 | 3.59 | 3.07E-31 |
| 225 | Gem | 2.64 | 6.49E-30 |
| 226 | LOC114689426 | 3.82 | 7.49E-29 |
| 227 | Adora2a | 2.37 | 7.85E-29 |
| 228 | LOC114691764 | 2.32 | 8.98E-29 |
| 229 | Kcne4 | 3.13 | 1.21E-28 |
| 230 | LOC114684574 | 5.62 | 1.78E-28 |
| 231 | Dcbld1 | 2.01 | 2.03E-28 |
| 232 | Steap1 | 6.48 | 2.13E-28 |
| 233 | LOC114681711 | 2.10 | 2.17E-28 |
| 234 | Irf9 | 2.33 | 6.75E-28 |
| 235 | LOC114684563 | 5.94 | 7.66E-28 |
| 236 | Sytl3 | 2.54 | 1.21E-27 |
| 237 | Sele | 4.50 | 1.51E-27 |
| 238 | Ier3 | 2.06 | 3.16E-27 |
| 239 | Adm | 2.28 | 3.17E-27 |
| 240 | LOC114701096 | 3.76 | 4.84E-27 |
| 241 | Vgf | 4.94 | 5.98E-27 |
| 242 | Nlrp3 | 2.01 | 7.11E-27 |
| 243 | Rgs1 | 2.19 | 1.99E-26 |
| 244 | Tnfrsf12a | 2.22 | 2.54E-26 |
| 245 | Rtp4 | 3.10 | 3.69E-26 |
| 246 | LOC114688399 | 3.05 | 3.75E-26 |
| 247 | Aldh1a2 | 2.82 | 3.76E-26 |
| 248 | LOC114688415 | 5.11 | 1.23E-25 |
| 249 | Nlrc5 | 2.32 | 1.35E-25 |
| 250 | LOC114688407 | 5.14 | 1.80E-25 |
| 251 | Asb4 | 5.60 | 2.04E-25 |
| 252 | Usp18 | 4.41 | 3.04E-25 |
| 253 | Slc28a3 | 3.61 | 1.69E-24 |
| 254 | LOC114684566 | 5.75 | 1.93E-24 |
| 255 | Cd14 | 3.30 | 2.87E-24 |
| 256 | LOC114690363 | 3.60 | 3.17E-24 |
| 257 | LOC114688611 | 3.72 | 9.68E-24 |
| 258 | Pdcd1lg2 | 3.16 | 1.88E-23 |
| 259 | Pappa | 2.87 | 1.93E-23 |
| 260 | Irf7 | 3.38 | 3.08E-23 |
| 261 | Lcn2 | 2.79 | 3.39E-23 |
| 262 | Adamts1 | 2.27 | 3.40E-23 |
| 263 | Penk | 3.35 | 8.90E-23 |
| 264 | Macc1 | 2.28 | 1.19E-22 |
| 265 | LOC114708899 | 2.03 | 1.21E-22 |
| 266 | Unc45b | 3.24 | 1.82E-22 |
| 267 | LOC114690618 | 6.15 | 2.34E-22 |
| 268 | Nr4a3 | 2.14 | 3.09E-22 |
| 269 | Dennd4a | 2.01 | 3.40E-22 |
| 270 | Slc16a14 | 4.05 | 4.24E-22 |
| 271 | LOC114707726 | 6.17 | 4.92E-22 |
| 272 | LOC114691793 | 3.00 | 9.03E-22 |
| 273 | LOC114685662 | 3.74 | 9.07E-22 |
| 274 | LOC114689226 | 3.43 | 1.35E-21 |
| 275 | Ifnb1 | 7.64 | 1.44E-21 |
| 276 | LOC114690754 | 7.57 | 1.88E-21 |
| 277 | Bst1 | 2.50 | 2.44E-21 |
| 278 | Apobec1 | 2.72 | 2.88E-21 |
| 279 | LOC114702149 | 2.65 | 2.99E-21 |
| 280 | LOC114710246 | 2.11 | 3.43E-21 |

| | | | |
|---|---|---|---|
| 468 | LOC114685748 | 2.20 | 7.94E-05 |
| 469 | LOC114709118 | 4.17 | 9.04E-05 |
| 470 | LOC114691422 | 2.46 | 1.42E-04 |
| 471 | LOC114690679 | 2.24 | 1.44E-04 |
| 472 | LOC114704902 | 4.44 | 1.49E-04 |
| 473 | Ankrd2 | 2.52 | 1.97E-04 |
| 474 | Sim2 | 2.84 | 2.09E-04 |
| 475 | Ankrd1 | 2.14 | 2.20E-04 |
| 476 | LOC114703306 | 4.23 | 2.23E-04 |
| 477 | Slc9a2 | 2.69 | 2.32E-04 |
| 478 | Fam221a | 2.12 | 2.47E-04 |
| 479 | Hes3 | 3.22 | 2.86E-04 |
| 480 | LOC114700344 | 2.11 | 2.94E-04 |
| 481 | LOC114698246 | 2.01 | 3.12E-04 |
| 482 | Scg2 | 3.72 | 3.41E-04 |
| 483 | LOC114700346 | 4.52 | 3.96E-04 |
| 484 | LOC114689301 | 3.42 | 4.11E-04 |
| 485 | LOC114700507 | 3.18 | 4.97E-04 |
| 486 | LOC114700187 | 3.88 | 7.59E-04 |
| 487 | Foxi2 | 4.75 | 9.14E-04 |
| 488 | Dpysl5 | 2.64 | 9.14E-04 |
| 489 | Olig1 | 2.55 | 1.05E-03 |
| 490 | Cfap161 | 2.07 | 1.42E-03 |
| 491 | Nr1h4 | 2.38 | 1.79E-03 |
| 492 | Gjb4 | 2.67 | 1.97E-03 |
| 493 | Liph | 2.28 | 2.11E-03 |
| 494 | Mymk | 2.58 | 2.41E-03 |
| 495 | Krt16 | 2.66 | 2.42E-03 |
| 496 | Frem3 | 2.81 | 2.46E-03 |
| 497 | LOC114691953 | 2.33 | 3.27E-03 |
| 498 | Dgkk | 2.19 | 3.41E-03 |
| 499 | Itgb6 | 3.10 | 3.46E-03 |
| 500 | LOC114687677 | 2.43 | 3.92E-03 |
| 501 | LOC114703660 | 2.84 | 4.97E-03 |
| 502 | LOC114702545 | 2.46 | 5.65E-03 |
| 503 | Aipl1 | 2.06 | 5.94E-03 |
| 504 | Crnn | 3.68 | 7.74E-03 |
| 505 | Pnoc | 2.34 | 7.81E-03 |
| 506 | LOC114706607 | 2.63 | 8.31E-03 |
| 507 | Trim54 | 3.46 | 8.67E-03 |
| 508 | Prm1 | 4.55 | 9.25E-03 |
| 509 | LOC114690676 | 2.22 | 1.03E-02 |
| 510 | Anxa10 | 3.53 | 1.05E-02 |
| 511 | LOC114710804 | 2.24 | 1.08E-02 |
| 512 | Tmprss13 | 3.46 | 1.08E-02 |
| 513 | LOC114683803 | 3.25 | 1.08E-02 |
| 514 | LOC114683863 | 3.38 | 1.23E-02 |
| 515 | LOC114710130 | 2.21 | 1.32E-02 |
| 516 | Cnga1 | 2.82 | 1.37E-02 |
| 517 | Tmcc2 | 2.12 | 1.74E-02 |
| 518 | LOC114710822 | 2.37 | 1.89E-02 |
| 519 | LOC114695766 | 2.74 | 1.92E-02 |
| 520 | Btnl2 | 2.66 | 1.99E-02 |
| 521 | Erfe | 2.16 | 2.07E-02 |
| 522 | LOC114704434 | 4.57 | 2.08E-02 |
| 523 | Ccnb1ip1 | 2.73 | 2.14E-02 |
| 524 | LOC114700427 | 3.71 | 2.23E-02 |
| 525 | LOC114683008 | 2.05 | 2.26E-02 |
| 526 | LOC114691605 | 2.50 | 2.62E-02 |
| 527 | LOC114682671 | 2.34 | 2.72E-02 |
| 528 | Prss37 | 3.23 | 2.82E-02 |
| 529 | LOC114694350 | 2.15 | 2.85E-02 |
| 530 | LOC114688145 | 2.38 | 2.90E-02 |
| 531 | LOC114694376 | 2.08 | 2.92E-02 |
| 532 | Pate1 | 3.39 | 2.99E-02 |
| 533 | Eqtn | 2.41 | 3.03E-02 |
| 534 | LOC114695173 | 2.06 | 3.16E-02 |
| 535 | Slfn14 | 2.14 | 3.21E-02 |
| 536 | LOC114696646 | 2.43 | 3.49E-02 |
| 537 | LOC114681963 | 2.02 | 3.49E-02 |
| 538 | Zar1 | 2.68 | 3.54E-02 |
| 539 | LOC114702919 | 2.29 | 3.60E-02 |
| 540 | LOC114689274 | 2.80 | 3.65E-02 |
| 541 | CUNH9orf50 | 3.06 | 3.67E-02 |
| 542 | Cngb3 | 3.59 | 3.83E-02 |
| 543 | LOC114686303 | 2.02 | 4.00E-02 |
| 544 | Trim69 | 3.13 | 4.05E-02 |
| 545 | Cldn24 | 2.33 | 4.23E-02 |
| 546 | Wnt3 | 2.42 | 4.34E-02 |
| 547 | Trim10 | 2.05 | 4.86E-02 |
| 548 | LOC114687761 | 2.12 | 4.91E-02 |
| 549 | LOC114705900 | 2.22 | 4.92E-02 |

**Table S6. DEGs in liver of P. leucopus**

| rank | gene | log2 fold change | FDR | rank | gene | log2 fold change | FDR | rank | gene | log2 fold change | FDR |
|---|---|---|---|---|---|---|---|---|---|---|---|
| 1 | Saa3 | 9.40 | 3.49E-29 | 376 | Wnt5b | 4.83 | 1.03E-11 | 751 | Nlrp6 | 3.88 | 2.43E-05 |
| 2 | Saa3 | 9.57 | 5.93E-28 | 377 | Pram1 | 3.03 | 1.13E-11 | 752 | LOC114688988 | 3.17 | 2.53E-05 |
| 3 | Saa5 | 9.37 | 6.13E-26 | 378 | Mcemp1 | 4.88 | 1.24E-11 | 753 | LOC114696384 | 2.09 | 2.53E-05 |
| 4 | Tnfaip3 | 4.57 | 2.29E-25 | 379 | Ca13 | 2.79 | 1.31E-11 | 754 | Gpr149 | 2.69 | 2.66E-05 |
| 5 | Znfx1 | 3.63 | 2.29E-25 | 380 | Cabp2 | 5.72 | 1.35E-11 | 755 | Mcm10 | 2.22 | 2.82E-05 |
| 6 | Isg20 | 7.13 | 3.97E-25 | 381 | Tph2 | 4.90 | 1.37E-11 | 756 | Gzmk | 5.82 | 2.91E-05 |
| 7 | Saa1 | 8.52 | 1.10E-24 | 382 | LOC114693967 | 2.08 | 1.38E-11 | 757 | Kcnh1 | 5.28 | 2.95E-05 |
| 8 | Sp140 | 4.29 | 1.10E-24 | 383 | LOC114681711 | 2.55 | 1.44E-11 | 758 | CUNH16orf96 | 3.98 | 2.99E-05 |
| 9 | Rars | 3.83 | 1.41E-24 | 384 | LOC114703215 | 4.52 | 1.53E-11 | 759 | Pdcd1lg2 | 3.53 | 3.05E-05 |
| 10 | Timp1 | 6.59 | 2.80E-24 | 385 | Hsh2d | 3.96 | 1.53E-11 | 760 | LOC114700264 | 4.53 | 3.07E-05 |
| 11 | Rnf213 | 4.88 | 2.80E-24 | 386 | Gas7 | 2.83 | 1.53E-11 | 761 | Ddr1 | 2.21 | 3.09E-05 |
| 12 | Slc39a14 | 3.43 | 2.80E-24 | 387 | Marcksl1 | 2.58 | 1.58E-11 | 762 | LOC114688401 | 4.12 | 3.17E-05 |
| 13 | Helz2 | 3.41 | 2.80E-24 | 388 | Ccl3 | 6.91 | 1.65E-11 | 763 | Lepr | 2.39 | 3.20E-05 |
| 14 | Efna5 | 3.35 | 2.80E-24 | 389 | Adcy1 | 2.80 | 1.65E-11 | 764 | Batf3 | 2.90 | 3.26E-05 |
| 15 | Gtf2ird1 | 3.02 | 2.80E-24 | 390 | LOC114688533 | 2.10 | 1.65E-11 | 765 | LOC114687932 | 3.37 | 3.32E-05 |
| 16 | Slc28a2 | 3.82 | 4.27E-24 | 391 | Cdkn1a | 3.31 | 1.86E-11 | 766 | Tbx4 | 2.29 | 3.37E-05 |
| 17 | Ddx58 | 4.35 | 4.66E-24 | 392 | LOC114688399 | 2.34 | 1.86E-11 | 767 | LOC114687928 | 3.42 | 3.61E-05 |
| 18 | Selp | 6.31 | 7.93E-24 | 393 | Gata3 | 5.44 | 1.88E-11 | 768 | LOC114694372 | 3.85 | 3.71E-05 |
| 19 | Icam1 | 4.85 | 8.97E-24 | 394 | Fbxo27 | 3.03 | 2.34E-11 | 769 | LOC114695552 | 2.06 | 4.36E-05 |
| 20 | C2cd3 | 3.49 | 9.86E-24 | 395 | Mrap | 3.54 | 2.58E-11 | 770 | LOC114690953 | 2.52 | 4.44E-05 |
| 21 | Steap4 | 5.33 | 1.01E-23 | 396 | Ccdc116 | 3.49 | 2.67E-11 | 771 | LOC114681290 | 2.27 | 4.49E-05 |
| 22 | Rnf19b | 4.00 | 1.07E-23 | 397 | LOC114686901 | 2.30 | 2.75E-11 | 772 | Rgcc | 3.33 | 4.87E-05 |
| 23 | Tap1 | 4.66 | 1.21E-23 | 398 | Cd274 | 2.82 | 2.75E-11 | 773 | LOC114686717 | 3.37 | 5.35E-05 |
| 24 | Slc13a5 | 5.40 | 1.56E-23 | 399 | Cacna1c | 2.99 | 2.76E-11 | 774 | Vash2 | 3.18 | 5.43E-05 |
| 25 | Cxcl10 | 7.70 | 3.60E-23 | 400 | Nfkb1 | 2.09 | 2.86E-11 | 775 | Stmn3 | 2.19 | 5.55E-05 |
| 26 | Nlrc5 | 6.71 | 3.60E-23 | 401 | Maff | 3.29 | 2.91E-11 | 776 | Eqtn | 2.88 | 5.86E-05 |
| 27 | Ptpn1 | 3.35 | 3.60E-23 | 402 | Capn9 | 3.20 | 3.03E-11 | 777 | Prrg4 | 4.69 | 6.26E-05 |
| 28 | Trex1 | 4.45 | 4.71E-23 | 403 | Rec8 | 2.88 | 3.10E-11 | 778 | LOC114689897 | 2.18 | 6.93E-05 |
| 29 | Arfgap3 | 2.60 | 5.34E-23 | 404 | LOC114710250 | 4.55 | 3.59E-11 | 779 | LOC114704122 | 3.37 | 7.30E-05 |
| 30 | Nnmt | 4.87 | 8.12E-23 | 405 | Sowahc | 2.94 | 3.85E-11 | 780 | LOC114695309 | 3.06 | 7.52E-05 |
| 31 | LOC114686891 | 4.08 | 8.18E-23 | 406 | LOC114697855 | 4.81 | 4.29E-11 | 781 | LOC114695955 | 2.27 | 7.60E-05 |
| 32 | Xaf1 | 5.00 | 1.09E-22 | 407 | LOC114688409 | 3.02 | 4.39E-11 | 782 | LOC114695845 | 4.95 | 7.84E-05 |
| 33 | LOC114707630 | 6.56 | 1.28E-22 | 408 | Gls | 2.14 | 4.55E-11 | 783 | LOC114689250 | 2.55 | 8.15E-05 |
| 34 | Eif2ak2 | 4.15 | 1.28E-22 | 409 | Fam71f2 | 6.68 | 4.59E-11 | 784 | LOC114681141 | 2.63 | 8.73E-05 |
| 35 | Slc41a1 | 3.48 | 1.42E-22 | 410 | Alpk2 | 8.57 | 4.64E-11 | 785 | Cemip | 4.02 | 9.00E-05 |
| 36 | Pfkfb3 | 4.73 | 1.60E-22 | 411 | Itih6 | 3.70 | 4.77E-11 | 786 | LOC114694103 | 6.34 | 9.20E-05 |
| 37 | Snx10 | 3.40 | 1.60E-22 | 412 | Batf | 4.34 | 4.91E-11 | 787 | Dhrs9 | 2.73 | 9.88E-05 |
| 38 | Junb | 3.15 | 1.60E-22 | 413 | Ccdc138 | 2.22 | 4.97E-11 | 788 | Frmd7 | 2.92 | 1.02E-04 |
| 39 | LOC114681715 | 4.69 | 2.26E-22 | 414 | Angptl4 | 4.12 | 5.03E-11 | 789 | Slc2a4rg | 2.79 | 1.02E-04 |
| 40 | LOC114688405 | 3.82 | 2.26E-22 | 415 | LOC114694400 | 2.90 | 5.43E-11 | 790 | LOC114700453 | 2.41 | 1.02E-04 |
| 41 | Stat3 | 2.73 | 2.56E-22 | 416 | Cds1 | 2.48 | 5.61E-11 | 791 | Pde6a | 2.88 | 1.04E-04 |
| 42 | LOC114681707 | 4.48 | 2.80E-22 | 417 | Ccrl2 | 2.99 | 6.26E-11 | 792 | LOC114706762 | 2.46 | 1.05E-04 |
| 43 | Rnd1 | 8.02 | 2.87E-22 | 418 | LOC114691641 | 4.68 | 6.29E-11 | 793 | Opn1sw | 4.47 | 1.06E-04 |
| 44 | Arid5a | 5.82 | 3.27E-22 | 419 | Prdm1 | 3.36 | 6.74E-11 | 794 | LOC114708201 | 3.30 | 1.23E-04 |
| 45 | LOC114706392 | 6.48 | 4.32E-22 | 420 | Gpr108 | 2.36 | 6.98E-11 | 795 | LOC114688129 | 2.92 | 1.23E-04 |
| 46 | Cd14 | 7.66 | 5.53E-22 | 421 | LOC114708095 | 2.91 | 7.21E-11 | 796 | Cxcl14 | 4.90 | 1.23E-04 |
| 47 | Tubb6 | 4.22 | 5.53E-22 | 422 | Ccl20 | 6.92 | 7.22E-11 | 797 | LOC114688645 | 2.07 | 1.24E-04 |
| 48 | Sod2 | 3.08 | 5.57E-22 | 423 | Serpina7 | 2.38 | 7.33E-11 | 798 | Myot | 5.64 | 1.25E-04 |
| 49 | Sphk1 | 6.35 | 6.03E-22 | 424 | Cxcl9 | 4.46 | 8.13E-11 | 799 | Il23a | 2.34 | 1.27E-04 |
| 50 | Rbm28 | 2.62 | 8.82E-22 | 425 | LOC114688611 | 5.29 | 8.56E-11 | 800 | LOC114692262 | 2.91 | 1.28E-04 |
| 51 | Acod1 | 7.79 | 9.37E-22 | 426 | LOC114688415 | 5.65 | 9.08E-11 | 801 | Gfpt2 | 2.12 | 1.28E-04 |
| 52 | Fpr2 | 6.83 | 9.72E-22 | 427 | LOC114702823 | 6.36 | 9.22E-11 | 802 | Scube3 | 2.84 | 1.29E-04 |
| 53 | Phf11 | 4.20 | 9.72E-22 | 428 | LOC114707719 | 6.61 | 9.33E-11 | 803 | LOC114685171 | 2.54 | 1.50E-04 |
| 54 | Gnl3 | 2.12 | 1.27E-21 | 429 | Stat1 | 2.40 | 9.37E-11 | 804 | Msantd1 | 2.55 | 1.51E-04 |
| 55 | Nfxl1 | 2.46 | 1.64E-21 | 430 | Nlgn2 | 2.36 | 9.51E-11 | 805 | LOC114703841 | 2.12 | 1.51E-04 |
| 56 | Dram1 | 5.36 | 1.68E-21 | 431 | LOC114681718 | 2.02 | 9.51E-11 | 806 | Cdk3 | 2.28 | 1.59E-04 |
| 57 | Smad1 | 2.59 | 1.71E-21 | 432 | LOC114688014 | 2.38 | 9.91E-11 | 807 | LOC114701896 | 3.28 | 1.78E-04 |
| 58 | Pprc1 | 2.56 | 1.80E-21 | 433 | Morc3 | 2.15 | 9.93E-11 | 808 | LOC114708202 | 3.38 | 1.92E-04 |
| 59 | Tnfrsf1b | 3.70 | 2.44E-21 | 434 | Col9a3 | 4.67 | 1.03E-10 | 809 | Rmi2 | 2.63 | 1.92E-04 |
| 60 | Sdc4 | 3.12 | 2.54E-21 | 435 | LOC114695911 | 4.81 | 1.05E-10 | 810 | Cx3cl1 | 2.16 | 1.96E-04 |
| 61 | Slc12a1 | 6.73 | 2.91E-21 | 436 | LOC114697308 | 2.54 | 1.32E-10 | 811 | Higd1c | 2.79 | 2.06E-04 |
| 62 | Upp1 | 4.65 | 4.06E-21 | 437 | Asap3 | 2.14 | 1.41E-10 | 812 | Fam3b | 2.82 | 2.14E-04 |
| 63 | CUNH1orf226 | 2.11 | 4.85E-21 | 438 | Coq10b | 2.32 | 1.43E-10 | 813 | LOC114699545 | 2.41 | 2.15E-04 |
| 64 | Cpne8 | 3.68 | 5.34E-21 | 439 | Lgals9 | 2.88 | 1.43E-10 | 814 | LOC114685334 | 4.36 | 2.28E-04 |
| 65 | Tnip2 | 3.63 | 5.87E-21 | 440 | Ebi3 | 2.91 | 1.55E-10 | 815 | Mybl2 | 2.34 | 2.40E-04 |
| 66 | Myc | 4.35 | 7.19E-21 | 441 | Gan | 2.02 | 1.67E-10 | 816 | Tmem114 | 2.12 | 2.40E-04 |
| 67 | Ets2 | 2.85 | 8.69E-21 | 442 | Kctd19 | 4.77 | 1.72E-10 | 817 | LOC114707659 | 2.30 | 2.41E-04 |
| 68 | LOC114701736 | 6.81 | 9.13E-21 | 443 | Runx1 | 3.28 | 1.96E-10 | 818 | Ctxn2 | 4.97 | 2.42E-04 |
| 69 | Cxcl11 | 9.15 | 1.08E-20 | 444 | LOC114688414 | 2.27 | 1.97E-10 | 819 | LOC114689445 | 2.13 | 2.45E-04 |
| 70 | Dtx2 | 2.45 | 1.08E-20 | 445 | Gpr84 | 5.22 | 2.03E-10 | 820 | Slc7a1 | 2.23 | 2.47E-04 |
| 71 | Ifih1 | 4.88 | 1.14E-20 | 446 | Slamf7 | 4.87 | 2.24E-10 | 821 | Pdcd1 | 4.84 | 2.58E-04 |
| 72 | Pla1a | 3.42 | 1.24E-20 | 447 | LOC114689266 | 4.47 | 2.54E-10 | 822 | LOC114693045 | 4.22 | 2.66E-04 |
| 73 | S100a9 | 5.49 | 1.25E-20 | 448 | Zfand2a | 2.24 | 2.79E-10 | 823 | Gal | 4.88 | 2.71E-04 |
| 74 | Mmp8 | 8.50 | 1.48E-20 | 449 | Adora2b | 4.22 | 3.20E-10 | 824 | Vat1l | 2.82 | 2.96E-04 |
| 75 | Gdf9 | 4.98 | 1.55E-20 | 450 | Atp10d | 2.31 | 3.27E-10 | 825 | LOC114699221 | 4.96 | 3.00E-04 |
| 76 | Sp110 | 3.08 | 1.70E-20 | 451 | LOC114709884 | 2.29 | 3.46E-10 | 826 | Npffr2 | 8.79 | 3.07E-04 |
| 77 | Dhx58 | 4.09 | 2.37E-20 | 452 | Rnf186 | 4.17 | 3.59E-10 | 827 | LOC114688764 | 3.22 | 3.17E-04 |
| 78 | Rrp9 | 2.82 | 2.57E-20 | 453 | Frmpd2 | 5.88 | 3.67E-10 | 828 | Scn4a | 3.67 | 3.22E-04 |
| 79 | Amigo3 | 3.61 | 2.74E-20 | 454 | Trh | 4.91 | 4.17E-10 | 829 | Asb4 | 4.94 | 3.29E-04 |
| 80 | Rab20 | 2.80 | 3.17E-20 | 455 | Pik3r5 | 2.47 | 4.46E-10 | 830 | Ido1 | 3.87 | 3.31E-04 |
| 81 | Pim1 | 5.02 | 3.47E-20 | 456 | Lcn2 | 6.96 | 4.62E-10 | 831 | Dok7 | 3.79 | 3.33E-04 |
| 82 | Cnp | 2.99 | 3.68E-20 | 457 | LOC114694398 | 2.68 | 4.63E-10 | 832 | Ccer2 | 2.47 | 3.53E-04 |
| 83 | Btg2 | 4.21 | 3.82E-20 | 458 | Slc39a6 | 2.20 | 4.63E-10 | 833 | Gramd2a | 4.28 | 3.55E-04 |
| 84 | Samhd1 | 5.12 | 4.10E-20 | 459 | Mmp12 | 5.02 | 4.92E-10 | 834 | CUNH9orf131 | 2.60 | 3.65E-04 |
| 85 | Dexi | 2.90 | 4.27E-20 | 460 | Cmya5 | 2.08 | 5.01E-10 | 835 | LOC114701422 | 2.54 | 3.65E-04 |
| 86 | Bhlha15 | 5.70 | 4.38E-20 | 461 | Vcan | 4.61 | 5.12E-10 | 836 | Gpr33 | 4.34 | 3.76E-04 |
| 87 | Tasor2 | 2.33 | 5.12E-20 | 462 | Cenpk | 4.25 | 5.12E-10 | 837 | Crabp2 | 3.30 | 3.77E-04 |
| 88 | Sbno2 | 2.82 | 5.76E-20 | 463 | Jun | 2.06 | 5.30E-10 | 838 | LOC114710471 | 2.07 | 3.84E-04 |
| 89 | Pglyrp1 | 6.24 | 5.91E-20 | 464 | Aoah | 3.30 | 5.86E-10 | 839 | LOC114706817 | 3.13 | 3.92E-04 |
| 90 | Tfpi | 3.75 | 6.25E-20 | 465 | Ccl4 | 7.69 | 6.56E-10 | 840 | Col27a1 | 2.00 | 3.94E-04 |
| 91 | Hilpda | 5.11 | 6.64E-20 | 466 | LOC114685537 | 2.97 | 8.00E-10 | 841 | Tldc2 | 3.15 | 3.95E-04 |
| 92 | Relb | 5.45 | 6.67E-20 | 467 | Tent5a | 2.09 | 8.00E-10 | 842 | Prph2 | 5.82 | 4.05E-04 |

Table S6

| | | | |
|---|---|---|---|
| 93 | Csf3r | 4.47 | 8.03E-20 |
| 94 | Als2cl | 3.89 | 9.22E-20 |
| 95 | Sema4b | 2.30 | 9.80E-20 |
| 96 | Socs3 | 5.48 | 1.01E-19 |
| 97 | Slpi | 9.09 | 1.38E-19 |
| 98 | Cmpk2 | 6.79 | 1.69E-19 |
| 99 | Rassf1 | 2.34 | 1.78E-19 |
| 100 | Cebpd | 3.02 | 1.84E-19 |
| 101 | Pml | 2.79 | 1.85E-19 |
| 102 | LOC114684568 | 9.18 | 2.02E-19 |
| 103 | Mitd1 | 2.51 | 2.10E-19 |
| 104 | LOC114684574 | 9.28 | 2.13E-19 |
| 105 | Kctd10 | 2.21 | 2.34E-19 |
| 106 | Ugcg | 2.81 | 2.47E-19 |
| 107 | Gadd45b | 5.17 | 2.82E-19 |
| 108 | Unc93b1 | 3.13 | 3.02E-19 |
| 109 | Psmb8 | 4.43 | 3.18E-19 |
| 110 | LOC114688093 | 4.59 | 3.49E-19 |
| 111 | Nfkbib | 2.04 | 3.58E-19 |
| 112 | LOC114686894 | 4.90 | 4.18E-19 |
| 113 | LOC114694511 | 6.02 | 5.99E-19 |
| 114 | Fosl2 | 3.78 | 6.08E-19 |
| 115 | Ibsp | 9.48 | 6.15E-19 |
| 116 | Brwd1 | 2.22 | 6.99E-19 |
| 117 | LOC114707940 | 5.77 | 7.05E-19 |
| 118 | Rnd3 | 2.76 | 8.87E-19 |
| 119 | Osmr | 5.78 | 9.25E-19 |
| 120 | Septin5 | 6.42 | 9.86E-19 |
| 121 | Mettl1 | 2.10 | 9.86E-19 |
| 122 | Rrp12 | 2.42 | 1.07E-18 |
| 123 | Parp14 | 3.55 | 1.13E-18 |
| 124 | Il1b | 4.44 | 1.15E-18 |
| 125 | Usp18 | 6.89 | 1.22E-18 |
| 126 | Mab21l3 | 3.71 | 1.28E-18 |
| 127 | Adar | 2.62 | 1.36E-18 |
| 128 | Irf8 | 3.31 | 1.63E-18 |
| 129 | Crispld2 | 3.68 | 2.31E-18 |
| 130 | Ripk2 | 4.18 | 2.50E-18 |
| 131 | LOC114683452 | 2.57 | 2.50E-18 |
| 132 | Pla2g2a | 8.05 | 2.67E-18 |
| 133 | Hmgcr | 2.75 | 2.98E-18 |
| 134 | Samd9 | 4.80 | 2.99E-18 |
| 135 | Irak3 | 4.08 | 3.46E-18 |
| 136 | Psmb9 | 3.80 | 3.57E-18 |
| 137 | Socs1 | 6.67 | 3.62E-18 |
| 138 | Acvr1b | 2.46 | 4.98E-18 |
| 139 | Cxcl2 | 8.20 | 5.37E-18 |
| 140 | Xdh | 2.55 | 5.94E-18 |
| 141 | Ackr1 | 5.85 | 6.38E-18 |
| 142 | Txnip | 3.31 | 6.82E-18 |
| 143 | Zfp36 | 3.08 | 7.26E-18 |
| 144 | LOC114691127 | 4.95 | 7.28E-18 |
| 145 | Fbxo39 | 7.16 | 8.41E-18 |
| 146 | LOC114697860 | 4.42 | 8.84E-18 |
| 147 | Fgb | 2.11 | 1.04E-17 |
| 148 | Cd40 | 5.90 | 1.07E-17 |
| 149 | Gadd45g | 5.35 | 1.15E-17 |
| 150 | Tifa | 3.66 | 1.17E-17 |
| 151 | Tead1 | 3.30 | 1.25E-17 |
| 152 | Nek4 | 2.53 | 1.25E-17 |
| 153 | Ficd | 2.50 | 1.36E-17 |
| 154 | Nfkb2 | 2.79 | 1.39E-17 |
| 155 | Pim3 | 3.24 | 1.41E-17 |
| 156 | Rgs4 | 3.89 | 1.74E-17 |
| 157 | Il6 | 7.50 | 1.83E-17 |
| 158 | Steap1 | 7.54 | 1.96E-17 |
| 159 | Nuak2 | 3.95 | 1.96E-17 |
| 160 | Lyve1 | 2.78 | 2.00E-17 |
| 161 | Cfb | 2.22 | 2.16E-17 |
| 162 | Spty2d1 | 2.07 | 2.16E-17 |
| 163 | Itga7 | 6.51 | 2.19E-17 |
| 164 | Trim21 | 2.45 | 2.29E-17 |
| 165 | Batf2 | 4.84 | 2.52E-17 |
| 166 | Arfip2 | 2.29 | 2.52E-17 |
| 167 | Sftpd | 6.20 | 2.72E-17 |
| 168 | Rsad2 | 8.45 | 2.74E-17 |
| 169 | Aen | 3.45 | 2.90E-17 |
| 170 | Il1rn | 3.95 | 3.07E-17 |
| 171 | Dot1l | 2.13 | 3.26E-17 |
| 172 | Tnfrsf21 | 4.49 | 3.51E-17 |
| 173 | Trim25 | 2.04 | 3.57E-17 |
| 174 | LOC114683484 | 5.09 | 3.75E-17 |
| 175 | Fgg | 2.09 | 3.77E-17 |
| 176 | Mvp | 3.08 | 3.85E-17 |
| 177 | Icam4 | 7.84 | 3.93E-17 |
| 178 | Ube2l6 | 3.06 | 4.41E-17 |
| 179 | Uchl3 | 2.17 | 4.41E-17 |
| 180 | Cntfr | 2.12 | 4.76E-17 |
| 181 | LOC114697861 | 3.69 | 5.01E-17 |
| 182 | Dnajb9 | 2.26 | 5.34E-17 |
| 183 | LOC114708897 | 2.35 | 5.70E-17 |
| 184 | Mefv | 2.72 | 6.36E-17 |
| 185 | Leap2 | 3.21 | 6.44E-17 |
| 186 | LOC114685530 | 3.61 | 6.57E-17 |
| 187 | Clec5a | 6.33 | 7.03E-17 |
| 468 | Igfbp1 | 3.38 | 8.25E-10 |
| 469 | Mmp3 | 7.67 | 8.33E-10 |
| 470 | Tcim | 2.15 | 8.57E-10 |
| 471 | Hsbp1l1 | 3.90 | 8.92E-10 |
| 472 | Duox2 | 3.20 | 9.56E-10 |
| 473 | LOC114705906 | 2.26 | 1.02E-09 |
| 474 | LOC114699049 | 6.44 | 1.14E-09 |
| 475 | CUNH11orf96 | 2.53 | 1.19E-09 |
| 476 | LOC114682342 | 3.80 | 1.25E-09 |
| 477 | LOC114687526 | 4.17 | 1.27E-09 |
| 478 | Ccl4 | 7.44 | 1.31E-09 |
| 479 | LOC114684572 | 7.01 | 1.35E-09 |
| 480 | Arl14 | 3.69 | 1.36E-09 |
| 481 | LOC114698011 | 3.32 | 1.40E-09 |
| 482 | B4galt1 | 2.04 | 1.41E-09 |
| 483 | Creld2 | 2.59 | 1.43E-09 |
| 484 | Tlr3 | 2.31 | 1.72E-09 |
| 485 | LOC114692715 | 2.64 | 1.77E-09 |
| 486 | Adamts1 | 2.76 | 1.77E-09 |
| 487 | Edn1 | 3.98 | 1.99E-09 |
| 488 | Clec4e | 4.12 | 2.09E-09 |
| 489 | CUNH3orf62 | 3.56 | 2.23E-09 |
| 490 | Saal1 | 2.72 | 2.23E-09 |
| 491 | LOC114709504 | 4.87 | 2.28E-09 |
| 492 | Apold1 | 2.50 | 2.43E-09 |
| 493 | Kcnj8 | 2.46 | 2.51E-09 |
| 494 | Csf1 | 2.00 | 2.68E-09 |
| 495 | LOC114689262 | 2.39 | 2.69E-09 |
| 496 | LOC114694553 | 3.97 | 2.78E-09 |
| 497 | Tent5c | 2.65 | 2.80E-09 |
| 498 | LOC114690036 | 2.21 | 2.91E-09 |
| 499 | Tfpi2 | 2.47 | 3.11E-09 |
| 500 | Srgn | 2.02 | 3.26E-09 |
| 501 | Ooep | 4.78 | 3.42E-09 |
| 502 | Tmcc2 | 3.17 | 3.62E-09 |
| 503 | LOC114696379 | 2.93 | 3.72E-09 |
| 504 | LOC114687942 | 2.09 | 3.99E-09 |
| 505 | Cxcr2 | 4.75 | 4.02E-09 |
| 506 | LOC114704754 | 2.74 | 4.02E-09 |
| 507 | LOC114689864 | 2.76 | 4.25E-09 |
| 508 | Cpt1b | 2.15 | 4.35E-09 |
| 509 | Ripor2 | 2.10 | 4.42E-09 |
| 510 | Elmsan1 | 2.21 | 4.51E-09 |
| 511 | Tnip3 | 4.08 | 4.53E-09 |
| 512 | LOC114690039 | 2.15 | 5.28E-09 |
| 513 | LOC114702149 | 2.87 | 5.51E-09 |
| 514 | Oasl | 4.66 | 5.53E-09 |
| 515 | Il1rl2 | 3.09 | 5.55E-09 |
| 516 | Nek10 | 3.87 | 5.63E-09 |
| 517 | Slc28a3 | 6.39 | 5.63E-09 |
| 518 | Plaur | 3.49 | 5.80E-09 |
| 519 | LOC114708973 | 7.25 | 6.67E-09 |
| 520 | Ly96 | 2.36 | 6.68E-09 |
| 521 | Znf599 | 2.09 | 6.87E-09 |
| 522 | LOC114689273 | 5.01 | 7.25E-09 |
| 523 | S100a11 | 2.98 | 7.45E-09 |
| 524 | Samsn1 | 2.77 | 7.45E-09 |
| 525 | Mc5r | 2.48 | 7.52E-09 |
| 526 | Smtnl1 | 5.21 | 7.69E-09 |
| 527 | LOC114690041 | 2.27 | 8.14E-09 |
| 528 | LOC114692716 | 3.15 | 8.53E-09 |
| 529 | Rab44 | 4.42 | 8.73E-09 |
| 530 | LOC114696940 | 4.09 | 8.73E-09 |
| 531 | Gstcd | 2.29 | 8.80E-09 |
| 532 | Hk3 | 2.20 | 8.83E-09 |
| 533 | LOC114694439 | 5.86 | 8.86E-09 |
| 534 | Kcnj4 | 5.88 | 9.11E-09 |
| 535 | Irf7 | 4.26 | 9.34E-09 |
| 536 | Aoc2 | 2.58 | 9.85E-09 |
| 537 | Trip10 | 2.98 | 9.90E-09 |
| 538 | Stra6 | 5.07 | 1.00E-08 |
| 539 | LOC114681897 | 4.69 | 1.12E-08 |
| 540 | Slc22a15 | 2.03 | 1.24E-08 |
| 541 | Cdc6 | 3.98 | 1.42E-08 |
| 542 | Prok2 | 5.26 | 1.46E-08 |
| 543 | LOC114683652 | 2.05 | 1.48E-08 |
| 544 | Fam131a | 2.31 | 1.50E-08 |
| 545 | LOC114689150 | 2.66 | 1.56E-08 |
| 546 | Alox5ap | 3.43 | 1.64E-08 |
| 547 | Ms4a3 | 5.52 | 1.75E-08 |
| 548 | LOC114698905 | 3.46 | 1.80E-08 |
| 549 | Slc10a6 | 3.28 | 1.85E-08 |
| 550 | Mthfd2 | 2.71 | 1.95E-08 |
| 551 | LOC114690034 | 2.42 | 2.02E-08 |
| 552 | LOC114686457 | 4.44 | 2.05E-08 |
| 553 | Cxcl5 | 6.56 | 2.11E-08 |
| 554 | LOC114683353 | 2.19 | 2.23E-08 |
| 555 | Far2 | 2.65 | 2.29E-08 |
| 556 | Ncf1 | 2.58 | 2.29E-08 |
| 557 | LOC114694940 | 2.30 | 2.31E-08 |
| 558 | LOC114706400 | 3.20 | 2.33E-08 |
| 559 | Serpina3 | 2.05 | 2.33E-08 |
| 560 | Dnhd1 | 2.13 | 2.36E-08 |
| 561 | LOC114697823 | 2.77 | 2.38E-08 |
| 562 | Avpr2 | 3.58 | 2.41E-08 |
| 843 | Cd44 | 2.22 | 4.17E-04 |
| 844 | Pla2g5 | 3.40 | 4.19E-04 |
| 845 | LOC114690674 | 2.16 | 4.19E-04 |
| 846 | Sgip1 | 2.00 | 4.25E-04 |
| 847 | LOC114683290 | 4.45 | 4.25E-04 |
| 848 | LOC114685338 | 2.39 | 4.27E-04 |
| 849 | LOC114682343 | 2.81 | 4.38E-04 |
| 850 | LOC114708220 | 2.15 | 4.57E-04 |
| 851 | LOC114705894 | 2.86 | 4.77E-04 |
| 852 | Fam47e | 2.20 | 4.91E-04 |
| 853 | Pik3ip1 | 2.90 | 5.08E-04 |
| 854 | LOC114700935 | 2.00 | 5.28E-04 |
| 855 | LOC114687188 | 2.13 | 5.30E-04 |
| 856 | LOC114681726 | 2.82 | 5.36E-04 |
| 857 | LOC114687333 | 2.46 | 5.69E-04 |
| 858 | Shoc1 | 3.36 | 6.07E-04 |
| 859 | LOC114702837 | 3.48 | 6.26E-04 |
| 860 | LOC114700602 | 3.08 | 7.01E-04 |
| 861 | Rarres1 | 2.02 | 7.09E-04 |
| 862 | Kncn | 3.12 | 7.32E-04 |
| 863 | Ampd1 | 3.19 | 7.33E-04 |
| 864 | LOC114680822 | 2.78 | 7.37E-04 |
| 865 | Kcnj13 | 2.57 | 7.41E-04 |
| 866 | LOC114707585 | 4.39 | 7.42E-04 |
| 867 | Slc9a4 | 4.17 | 7.49E-04 |
| 868 | LOC114699544 | 2.07 | 7.56E-04 |
| 869 | Patl2 | 3.31 | 7.68E-04 |
| 870 | Elf3 | 3.19 | 7.85E-04 |
| 871 | Fitm1 | 3.03 | 8.12E-04 |
| 872 | Il18rap | 2.19 | 8.51E-04 |
| 873 | Adgrf1 | 3.35 | 8.77E-04 |
| 874 | Gcnt3 | 2.66 | 8.82E-04 |
| 875 | LOC114687425 | 3.04 | 8.87E-04 |
| 876 | LOC114697910 | 2.36 | 8.95E-04 |
| 877 | Orc1 | 4.06 | 9.13E-04 |
| 878 | Dkk4 | 4.34 | 9.53E-04 |
| 879 | Tlr5 | 4.13 | 9.63E-04 |
| 880 | LOC114691793 | 2.12 | 9.69E-04 |
| 881 | Fam71a | 4.52 | 9.71E-04 |
| 882 | LOC114707018 | 2.85 | 1.05E-03 |
| 883 | Pde6h | 4.75 | 1.06E-03 |
| 884 | LOC114708913 | 4.32 | 1.10E-03 |
| 885 | LOC114692112 | 2.81 | 1.11E-03 |
| 886 | Ecscr | 2.24 | 1.12E-03 |
| 887 | Ttll8 | 3.88 | 1.14E-03 |
| 888 | Ptch2 | 3.18 | 1.16E-03 |
| 889 | LOC114685691 | 2.77 | 1.18E-03 |
| 890 | Hcst | 2.09 | 1.19E-03 |
| 891 | LOC114682171 | 4.22 | 1.20E-03 |
| 892 | CUNH1orf162 | 2.89 | 1.22E-03 |
| 893 | LOC114681423 | 2.82 | 1.23E-03 |
| 894 | LOC114684208 | 2.75 | 1.37E-03 |
| 895 | Tubb1 | 2.25 | 1.38E-03 |
| 896 | LOC114707876 | 2.96 | 1.44E-03 |
| 897 | LOC114684236 | 2.14 | 1.46E-03 |
| 898 | LOC114708073 | 3.11 | 1.55E-03 |
| 899 | CUNH8orf58 | 2.13 | 1.57E-03 |
| 900 | LOC114691956 | 2.84 | 1.59E-03 |
| 901 | LOC114686010 | 2.60 | 1.63E-03 |
| 902 | Duoxa2 | 3.70 | 1.65E-03 |
| 903 | Bdkrb1 | 4.99 | 1.83E-03 |
| 904 | Pax9 | 4.66 | 1.86E-03 |
| 905 | LOC114696821 | 2.12 | 1.90E-03 |
| 906 | LOC114695442 | 2.08 | 1.90E-03 |
| 907 | LOC114694847 | 3.22 | 1.91E-03 |
| 908 | Cfap69 | 2.08 | 1.91E-03 |
| 909 | LOC114706978 | 4.43 | 2.01E-03 |
| 910 | LOC114703653 | 2.29 | 2.01E-03 |
| 911 | Chp2 | 2.08 | 2.02E-03 |
| 912 | Anxa2r | 4.33 | 2.03E-03 |
| 913 | Adgb | 2.41 | 2.06E-03 |
| 914 | LOC114695572 | 2.78 | 2.09E-03 |
| 915 | Gzmm | 2.19 | 2.17E-03 |
| 916 | CUNH11orf86 | 2.55 | 2.18E-03 |
| 917 | LOC114693720 | 2.13 | 2.41E-03 |
| 918 | Il36b | 2.84 | 2.45E-03 |
| 919 | LOC114697589 | 4.94 | 2.51E-03 |
| 920 | LOC114681904 | 2.72 | 2.69E-03 |
| 921 | Phyhip | 3.18 | 2.73E-03 |
| 922 | LOC114688285 | 2.37 | 2.88E-03 |
| 923 | LOC114687759 | 4.15 | 2.96E-03 |
| 924 | LOC114698551 | 2.93 | 3.01E-03 |
| 925 | Mt4 | 4.97 | 3.03E-03 |
| 926 | Mmp27 | 3.21 | 3.05E-03 |
| 927 | Sall4 | 2.11 | 3.12E-03 |
| 928 | LOC114694376 | 4.11 | 3.40E-03 |
| 929 | LOC114696388 | 3.56 | 3.57E-03 |
| 930 | Spink1 | 4.01 | 3.57E-03 |
| 931 | Dnd1 | 2.12 | 3.66E-03 |
| 932 | Tspo2 | 5.20 | 3.74E-03 |
| 933 | Gmip | 2.81 | 3.80E-03 |
| 934 | Myt1l | 2.23 | 3.80E-03 |
| 935 | LOC114707743 | 3.72 | 3.96E-03 |
| 936 | LOC114705490 | 4.37 | 3.96E-03 |
| 937 | St8sia3 | 3.01 | 3.99E-03 |

| | | | |
|---|---|---|---|
| 188 | Uba7 | 3.65 | 7.45E-17 |
| 189 | Tmem248 | 2.12 | 8.15E-17 |
| 190 | Gar1 | 2.29 | 1.01E-16 |
| 191 | Lcp1 | 2.49 | 1.25E-16 |
| 192 | LOC114688413 | 6.30 | 1.26E-16 |
| 193 | Map1lc3b | 2.32 | 1.37E-16 |
| 194 | Serpine1 | 4.16 | 1.37E-16 |
| 195 | Hyal2 | 2.35 | 1.37E-16 |
| 196 | Tdrd15 | 11.28 | 1.41E-16 |
| 197 | Slc11a2 | 2.31 | 1.52E-16 |
| 198 | Ddx21 | 2.26 | 1.69E-16 |
| 199 | Shisa3 | 5.84 | 2.38E-16 |
| 200 | Creb3l2 | 2.99 | 2.38E-16 |
| 201 | Ago2 | 2.39 | 2.48E-16 |
| 202 | Slc43a1 | 2.92 | 3.21E-16 |
| 203 | Parp10 | 2.38 | 3.32E-16 |
| 204 | Lypd1 | 2.72 | 3.39E-16 |
| 205 | Cgas | 5.24 | 3.52E-16 |
| 206 | Usp43 | 3.06 | 3.95E-16 |
| 207 | Ltf | 5.98 | 3.98E-16 |
| 208 | LOC114688398 | 3.03 | 4.13E-16 |
| 209 | Baz2b | 2.54 | 4.51E-16 |
| 210 | LOC114681719 | 3.85 | 4.89E-16 |
| 211 | Wars | 2.32 | 5.13E-16 |
| 212 | Dennd2d | 2.65 | 5.22E-16 |
| 213 | Itih3 | 4.14 | 5.47E-16 |
| 214 | Suco | 2.28 | 5.57E-16 |
| 215 | Ifngr2 | 3.64 | 6.47E-16 |
| 216 | LOC114685534 | 3.08 | 6.58E-16 |
| 217 | Fgr | 3.04 | 6.58E-16 |
| 218 | Zeb1 | 2.35 | 6.63E-16 |
| 219 | Sec23b | 2.14 | 8.89E-16 |
| 220 | LOC114685650 | 3.63 | 1.01E-15 |
| 221 | Spats2l | 4.54 | 1.06E-15 |
| 222 | Camp | 7.58 | 1.11E-15 |
| 223 | Tnfrsf14 | 2.74 | 1.37E-15 |
| 224 | Zc3h12a | 2.50 | 1.38E-15 |
| 225 | Adora2a | 3.25 | 1.81E-15 |
| 226 | Nfkbia | 2.49 | 1.88E-15 |
| 227 | Fga | 2.00 | 1.92E-15 |
| 228 | Hcar2 | 7.31 | 2.10E-15 |
| 229 | LOC114700341 | 4.85 | 2.16E-15 |
| 230 | Spx | 4.09 | 2.20E-15 |
| 231 | LOC114689891 | 2.51 | 2.42E-15 |
| 232 | LOC114694510 | 3.62 | 2.65E-15 |
| 233 | Rell1 | 2.85 | 2.87E-15 |
| 234 | Sdk2 | 3.44 | 3.02E-15 |
| 235 | Il2rb | 2.80 | 3.02E-15 |
| 236 | Map2k3 | 2.06 | 3.06E-15 |
| 237 | Trim15 | 2.87 | 3.08E-15 |
| 238 | Zc3hav1 | 2.34 | 3.08E-15 |
| 239 | Ddit4 | 3.59 | 3.59E-15 |
| 240 | Tmem173 | 3.20 | 4.24E-15 |
| 241 | LOC114710254 | 7.36 | 4.78E-15 |
| 242 | LOC114686850 | 4.95 | 5.08E-15 |
| 243 | LOC114688396 | 3.22 | 5.24E-15 |
| 244 | Ptk2b | 2.81 | 5.81E-15 |
| 245 | Meox1 | 6.14 | 6.39E-15 |
| 246 | Isg15 | 5.75 | 6.55E-15 |
| 247 | Fbxo31 | 2.26 | 6.55E-15 |
| 248 | Tnf | 4.39 | 6.91E-15 |
| 249 | Bst1 | 3.30 | 6.93E-15 |
| 250 | LOC114685662 | 5.83 | 8.33E-15 |
| 251 | Fpr1 | 2.83 | 9.33E-15 |
| 252 | LOC114691764 | 2.08 | 1.08E-14 |
| 253 | LOC114684566 | 8.47 | 1.08E-14 |
| 254 | Serpina10 | 2.01 | 1.18E-14 |
| 255 | Faslg | 3.06 | 1.20E-14 |
| 256 | Rhbdf2 | 2.04 | 1.26E-14 |
| 257 | Fap | 5.65 | 1.44E-14 |
| 258 | Bach1 | 2.94 | 1.46E-14 |
| 259 | Plek | 3.37 | 1.57E-14 |
| 260 | Chsy3 | 2.21 | 1.94E-14 |
| 261 | Mical2 | 2.16 | 2.11E-14 |
| 262 | Crygs | 5.94 | 2.34E-14 |
| 263 | LOC114705177 | 4.15 | 2.42E-14 |
| 264 | LOC114704463 | 4.18 | 2.46E-14 |
| 265 | LOC114692714 | 3.08 | 2.62E-14 |
| 266 | Alpk1 | 5.03 | 2.74E-14 |
| 267 | Slamf1 | 6.38 | 3.30E-14 |
| 268 | LOC114684563 | 8.58 | 3.35E-14 |
| 269 | LOC114701096 | 4.90 | 3.48E-14 |
| 270 | Stx11 | 3.75 | 3.85E-14 |
| 271 | Inhbb | 3.67 | 3.96E-14 |
| 272 | LOC114685528 | 2.52 | 4.02E-14 |
| 273 | Herc6 | 5.23 | 4.24E-14 |
| 274 | Csf2 | 9.44 | 4.31E-14 |
| 275 | Utp14a | 2.71 | 4.39E-14 |
| 276 | Gjb2 | 2.25 | 4.39E-14 |
| 277 | LOC114689426 | 4.79 | 4.58E-14 |
| 278 | Sla | 2.81 | 4.60E-14 |
| 279 | Paf1 | 2.63 | 4.65E-14 |
| 280 | Dlx4 | 2.43 | 4.79E-14 |
| 281 | Chek1 | 4.00 | 5.22E-14 |
| 282 | Sema4c | 2.56 | 5.45E-14 |
| 563 | LOC114680639 | 4.22 | 2.65E-08 |
| 564 | Kcne4 | 3.32 | 2.66E-08 |
| 565 | LOC114682712 | 4.67 | 2.82E-08 |
| 566 | Sybu | 2.64 | 2.84E-08 |
| 567 | LOC114696643 | 6.26 | 3.14E-08 |
| 568 | Ppm1j | 4.92 | 3.24E-08 |
| 569 | CUNH5orf34 | 2.31 | 3.25E-08 |
| 570 | Icam2 | 2.11 | 3.30E-08 |
| 571 | LOC114696387 | 2.20 | 3.72E-08 |
| 572 | Krtcap3 | 2.60 | 3.98E-08 |
| 573 | LOC114701656 | 2.94 | 4.08E-08 |
| 574 | Scimp | 2.01 | 4.33E-08 |
| 575 | Tp53tg5 | 2.64 | 4.45E-08 |
| 576 | Csf2rb | 2.48 | 4.62E-08 |
| 577 | Steap2 | 4.18 | 4.68E-08 |
| 578 | LOC114689226 | 3.13 | 4.84E-08 |
| 579 | Chi3l1 | 3.07 | 4.90E-08 |
| 580 | LOC114694396 | 4.58 | 4.96E-08 |
| 581 | B3galt1 | 3.39 | 5.23E-08 |
| 582 | LOC114691550 | 5.35 | 5.29E-08 |
| 583 | LOC114697821 | 3.27 | 5.71E-08 |
| 584 | Gp1bb | 5.92 | 6.21E-08 |
| 585 | Olr1 | 6.27 | 6.37E-08 |
| 586 | Rhob | 2.02 | 6.42E-08 |
| 587 | Dio3 | 3.96 | 6.72E-08 |
| 588 | Artn | 2.92 | 7.21E-08 |
| 589 | Egflam | 2.54 | 7.21E-08 |
| 590 | Adam8 | 2.41 | 7.37E-08 |
| 591 | Gpr39 | 2.10 | 7.41E-08 |
| 592 | Lmcd1 | 2.45 | 7.52E-08 |
| 593 | LOC114704713 | 4.34 | 7.55E-08 |
| 594 | Faxc | 2.53 | 7.62E-08 |
| 595 | Nos2 | 5.23 | 8.48E-08 |
| 596 | LOC114691549 | 3.78 | 8.49E-08 |
| 597 | Retn | 4.90 | 8.77E-08 |
| 598 | Ovgp1 | 2.36 | 8.80E-08 |
| 599 | LOC114692146 | 2.50 | 9.65E-08 |
| 600 | LOC114688564 | 8.98 | 9.98E-08 |
| 601 | B4galt5 | 2.66 | 1.01E-07 |
| 602 | Plin4 | 5.04 | 1.02E-07 |
| 603 | Slc6a4 | 3.14 | 1.15E-07 |
| 604 | LOC114703402 | 2.55 | 1.18E-07 |
| 605 | LOC114708899 | 2.72 | 1.19E-07 |
| 606 | Fscn1 | 2.59 | 1.21E-07 |
| 607 | Kif25 | 3.26 | 1.21E-07 |
| 608 | LOC114684798 | 2.66 | 1.30E-07 |
| 609 | Ttn | 2.30 | 1.34E-07 |
| 610 | LOC114688986 | 3.06 | 1.35E-07 |
| 611 | Herpud1 | 2.04 | 1.39E-07 |
| 612 | CUNH3orf20 | 2.03 | 1.40E-07 |
| 613 | LOC114681724 | 3.42 | 1.40E-07 |
| 614 | LOC114693317 | 2.07 | 1.44E-07 |
| 615 | Peg10 | 3.24 | 1.55E-07 |
| 616 | LOC114696258 | 2.01 | 1.57E-07 |
| 617 | LOC114707706 | 4.28 | 1.82E-07 |
| 618 | Tsc22d1 | 2.09 | 1.82E-07 |
| 619 | Nptx2 | 5.16 | 2.01E-07 |
| 620 | Ptges | 7.05 | 2.28E-07 |
| 621 | Col6a5 | 2.38 | 2.32E-07 |
| 622 | Nkain1 | 4.21 | 2.34E-07 |
| 623 | Tlr7 | 2.64 | 2.41E-07 |
| 624 | LOC114683448 | 2.30 | 2.44E-07 |
| 625 | Galnt6 | 2.18 | 2.48E-07 |
| 626 | LOC114694224 | 2.60 | 2.73E-07 |
| 627 | LOC114701467 | 3.46 | 2.81E-07 |
| 628 | LOC114701403 | 2.56 | 2.86E-07 |
| 629 | Bcl6 | 2.32 | 2.96E-07 |
| 630 | Nexn | 2.12 | 3.03E-07 |
| 631 | LOC114706751 | 2.08 | 3.05E-07 |
| 632 | Il10 | 3.63 | 3.19E-07 |
| 633 | LOC114693254 | 2.26 | 3.24E-07 |
| 634 | Penk | 4.26 | 3.37E-07 |
| 635 | Unc5cl | 3.67 | 3.55E-07 |
| 636 | LOC114698458 | 4.33 | 3.69E-07 |
| 637 | Il15 | 2.24 | 4.12E-07 |
| 638 | Creb3l1 | 2.09 | 4.27E-07 |
| 639 | Ttk | 2.24 | 4.65E-07 |
| 640 | LOC114699170 | 2.15 | 4.83E-07 |
| 641 | LOC114703718 | 2.28 | 5.09E-07 |
| 642 | Pstpip1 | 2.22 | 5.19E-07 |
| 643 | Inka1 | 3.68 | 5.52E-07 |
| 644 | Fos | 2.63 | 5.66E-07 |
| 645 | Tes | 2.35 | 5.74E-07 |
| 646 | Prss23 | 2.52 | 5.77E-07 |
| 647 | Col10a1 | 2.35 | 5.89E-07 |
| 648 | Plekha4 | 2.66 | 5.95E-07 |
| 649 | LOC114693253 | 2.56 | 5.98E-07 |
| 650 | LOC114703406 | 3.18 | 6.07E-07 |
| 651 | Clec4d | 4.27 | 6.16E-07 |
| 652 | LOC114701086 | 6.82 | 6.30E-07 |
| 653 | Nlrp12 | 3.30 | 6.36E-07 |
| 654 | Mmp10 | 3.40 | 6.68E-07 |
| 655 | Nod2 | 2.39 | 6.96E-07 |
| 656 | LOC114690032 | 2.02 | 7.36E-07 |
| 657 | LOC114699169 | 2.15 | 7.41E-07 |
| 938 | LOC114704434 | 5.57 | 4.39E-03 |
| 939 | LOC114700622 | 2.35 | 4.68E-03 |
| 940 | LOC114682671 | 2.34 | 4.69E-03 |
| 941 | Spock1 | 3.15 | 4.75E-03 |
| 942 | Pgr | 2.98 | 4.99E-03 |
| 943 | LOC114685758 | 2.81 | 5.25E-03 |
| 944 | Smc1b | 2.69 | 5.29E-03 |
| 945 | Plxna3 | 2.35 | 5.32E-03 |
| 946 | Amhr2 | 3.07 | 5.72E-03 |
| 947 | LOC114682677 | 2.54 | 5.86E-03 |
| 948 | LOC114701363 | 2.61 | 5.92E-03 |
| 949 | Arg2 | 2.32 | 6.38E-03 |
| 950 | LOC114683802 | 3.34 | 6.60E-03 |
| 951 | Scgb1c1 | 2.96 | 7.04E-03 |
| 952 | LOC114704590 | 2.05 | 7.04E-03 |
| 953 | Erp27 | 2.51 | 7.10E-03 |
| 954 | Calml3 | 4.42 | 7.31E-03 |
| 955 | LOC114690609 | 2.01 | 7.31E-03 |
| 956 | Sh2d4b | 3.14 | 7.39E-03 |
| 957 | Tmem236 | 3.69 | 7.39E-03 |
| 958 | Epo | 2.97 | 7.45E-03 |
| 959 | Mybpc3 | 2.70 | 7.46E-03 |
| 960 | LOC114704779 | 4.06 | 7.47E-03 |
| 961 | Ror2 | 2.18 | 7.53E-03 |
| 962 | Endou | 2.46 | 7.65E-03 |
| 963 | Ptpn14 | 2.02 | 8.06E-03 |
| 964 | LOC114690678 | 2.23 | 8.08E-03 |
| 965 | LOC114685950 | 2.16 | 8.11E-03 |
| 966 | Upp2 | 2.15 | 8.30E-03 |
| 967 | Ccl22 | 3.93 | 8.48E-03 |
| 968 | Ankrd2 | 2.38 | 8.48E-03 |
| 969 | LOC114689030 | 2.07 | 8.62E-03 |
| 970 | LOC114681259 | 4.83 | 9.14E-03 |
| 971 | LOC114707960 | 3.43 | 9.16E-03 |
| 972 | LOC114694343 | 5.09 | 9.18E-03 |
| 973 | Tmem252 | 2.85 | 9.29E-03 |
| 974 | LOC114683766 | 3.06 | 9.39E-03 |
| 975 | Me3 | 2.16 | 9.50E-03 |
| 976 | LOC114689292 | 4.12 | 9.55E-03 |
| 977 | Gem | 2.31 | 1.03E-02 |
| 978 | LOC114694036 | 4.36 | 1.03E-02 |
| 979 | LOC114689583 | 2.53 | 1.06E-02 |
| 980 | Dusp15 | 2.22 | 1.11E-02 |
| 981 | LOC114706260 | 2.52 | 1.12E-02 |
| 982 | Apoa4 | 2.15 | 1.13E-02 |
| 983 | LOC114683833 | 2.36 | 1.14E-02 |
| 984 | Mroh6 | 2.32 | 1.14E-02 |
| 985 | Hpgds | 2.35 | 1.18E-02 |
| 986 | LOC114686348 | 2.84 | 1.23E-02 |
| 987 | LOC114691343 | 2.22 | 1.25E-02 |
| 988 | LOC114687135 | 2.43 | 1.25E-02 |
| 989 | Hapln3 | 3.73 | 1.25E-02 |
| 990 | Fam217a | 3.23 | 1.26E-02 |
| 991 | LOC114706399 | 4.28 | 1.29E-02 |
| 992 | Ccdc179 | 2.74 | 1.30E-02 |
| 993 | LOC114688873 | 4.33 | 1.37E-02 |
| 994 | LOC114685408 | 5.45 | 1.38E-02 |
| 995 | LOC114687407 | 3.23 | 1.43E-02 |
| 996 | Upk1a | 2.08 | 1.47E-02 |
| 997 | LOC114685776 | 2.47 | 1.49E-02 |
| 998 | LOC114696933 | 2.95 | 1.52E-02 |
| 999 | Tmem95 | 3.33 | 1.52E-02 |
| 1000 | Il18r1 | 2.02 | 1.54E-02 |
| 1001 | Mak | 2.26 | 1.54E-02 |
| 1002 | LOC114708800 | 2.67 | 1.55E-02 |
| 1003 | LOC114710769 | 2.29 | 1.55E-02 |
| 1004 | Zc3h12b | 2.42 | 1.56E-02 |
| 1005 | LOC114709118 | 3.14 | 1.58E-02 |
| 1006 | Smim33 | 2.95 | 1.59E-02 |
| 1007 | Tcf23 | 2.12 | 1.66E-02 |
| 1008 | LOC114688041 | 3.56 | 1.71E-02 |
| 1009 | Tex33 | 2.30 | 1.72E-02 |
| 1010 | LOC114702990 | 4.31 | 1.73E-02 |
| 1011 | LOC114686332 | 2.50 | 1.83E-02 |
| 1012 | Hdc | 2.55 | 1.86E-02 |
| 1013 | Il20ra | 3.45 | 1.91E-02 |
| 1014 | LOC114688391 | 2.64 | 2.19E-02 |
| 1015 | LOC114705398 | 4.32 | 2.21E-02 |
| 1016 | Anxa10 | 2.96 | 2.23E-02 |
| 1017 | Plekhg4 | 3.41 | 2.28E-02 |
| 1018 | LOC114690987 | 3.57 | 2.30E-02 |
| 1019 | Gprc5a | 2.34 | 2.36E-02 |
| 1020 | Bdkrb2 | 2.54 | 2.47E-02 |
| 1021 | Lsmem1 | 2.49 | 2.57E-02 |
| 1022 | Nectin4 | 3.29 | 2.58E-02 |
| 1023 | LOC114682072 | 2.80 | 2.66E-02 |
| 1024 | LOC114697856 | 2.49 | 2.71E-02 |
| 1025 | LOC114705143 | 2.16 | 2.71E-02 |
| 1026 | Mlip | 3.10 | 2.73E-02 |
| 1027 | LOC114705361 | 2.23 | 2.80E-02 |
| 1028 | Nfasc | 2.35 | 2.81E-02 |
| 1029 | LOC114698355 | 2.94 | 2.84E-02 |
| 1030 | LOC114700898 | 3.28 | 2.84E-02 |
| 1031 | Stpg4 | 2.19 | 2.95E-02 |
| 1032 | Dpysl5 | 2.24 | 2.96E-02 |

Table S6

| | | | |
|---|---|---|---|
| 283 | Nfkbie | 3.57 | 6.18E-14 |
| 284 | Nfkbiz | 2.11 | 6.35E-14 |
| 285 | LOC114704347 | 4.78 | 6.66E-14 |
| 286 | 3-Mar | 2.86 | 6.78E-14 |
| 287 | Csf3 | 5.35 | 8.35E-14 |
| 288 | Btg3 | 2.44 | 8.99E-14 |
| 289 | Csrnp1 | 2.73 | 9.65E-14 |
| 290 | Mxd1 | 2.39 | 1.00E-13 |
| 291 | Fkbp14 | 3.62 | 1.01E-13 |
| 292 | Tnfrsf18 | 3.56 | 1.04E-13 |
| 293 | LOC114688984 | 2.94 | 1.23E-13 |
| 294 | LOC114692659 | 2.24 | 1.44E-13 |
| 295 | Rbm43 | 2.50 | 1.62E-13 |
| 296 | LOC114681329 | 3.69 | 1.65E-13 |
| 297 | LOC114681721 | 2.39 | 1.66E-13 |
| 298 | Tgm1 | 6.73 | 1.74E-13 |
| 299 | C1rl | 2.06 | 1.76E-13 |
| 300 | Arl5c | 5.60 | 1.82E-13 |
| 301 | Cyren | 2.09 | 1.82E-13 |
| 302 | Sh2b2 | 2.38 | 2.35E-13 |
| 303 | Il10rb | 3.03 | 2.36E-13 |
| 304 | Tap2 | 2.16 | 2.41E-13 |
| 305 | Ubd | 5.69 | 2.45E-13 |
| 306 | LOC114703403 | 5.47 | 2.46E-13 |
| 307 | Atf3 | 4.87 | 2.46E-13 |
| 308 | Rtn4rl2 | 7.08 | 2.59E-13 |
| 309 | Slc15a3 | 2.51 | 2.61E-13 |
| 310 | Cxcl16 | 2.92 | 2.75E-13 |
| 311 | Lasp1 | 2.00 | 2.80E-13 |
| 312 | LOC114687933 | 2.04 | 3.06E-13 |
| 313 | Tmem132a | 2.64 | 3.17E-13 |
| 314 | Jmjd1c | 2.84 | 3.45E-13 |
| 315 | Ubash3b | 2.29 | 3.56E-13 |
| 316 | Sele | 4.17 | 3.86E-13 |
| 317 | Icam5 | 6.10 | 4.25E-13 |
| 318 | Jdp2 | 2.37 | 4.36E-13 |
| 319 | LOC114684565 | 10.93 | 4.61E-13 |
| 320 | Tnfaip2 | 2.42 | 4.61E-13 |
| 321 | Ptx3 | 6.87 | 4.66E-13 |
| 322 | Ccnl1 | 2.23 | 4.75E-13 |
| 323 | LOC114706381 | 2.21 | 5.03E-13 |
| 324 | Ablim1 | 2.16 | 5.17E-13 |
| 325 | LOC114698684 | 2.19 | 5.31E-13 |
| 326 | Neurl3 | 4.92 | 5.37E-13 |
| 327 | Rtp4 | 2.37 | 5.58E-13 |
| 328 | LOC114701319 | 2.06 | 5.77E-13 |
| 329 | LOC114686895 | 2.02 | 5.85E-13 |
| 330 | Tmem140 | 2.59 | 6.33E-13 |
| 331 | Adamts4 | 3.35 | 6.51E-13 |
| 332 | Ncf4 | 2.34 | 6.76E-13 |
| 333 | Mmp19 | 2.17 | 6.76E-13 |
| 334 | Lbp | 3.75 | 8.58E-13 |
| 335 | Ifi16 | 3.27 | 9.38E-13 |
| 336 | LOC114694456 | 9.15 | 9.63E-13 |
| 337 | Fndc4 | 2.76 | 1.06E-12 |
| 338 | LOC114704785 | 2.24 | 1.08E-12 |
| 339 | Sys1 | 2.25 | 1.09E-12 |
| 340 | Mmp9 | 4.60 | 1.09E-12 |
| 341 | LOC114700347 | 5.21 | 1.17E-12 |
| 342 | CUNH15orf48 | 5.24 | 1.18E-12 |
| 343 | Tspan2 | 2.42 | 1.29E-12 |
| 344 | LOC114690952 | 2.08 | 1.31E-12 |
| 345 | Cfap299 | 2.71 | 1.40E-12 |
| 346 | LOC114707726 | 6.23 | 1.47E-12 |
| 347 | Syn1 | 4.36 | 1.62E-12 |
| 348 | LOC114697913 | 4.81 | 1.66E-12 |
| 349 | Dusp2 | 4.16 | 1.81E-12 |
| 350 | Rhoj | 3.00 | 1.90E-12 |
| 351 | Ralgds | 2.49 | 2.40E-12 |
| 352 | Ifi44 | 2.86 | 2.50E-12 |
| 353 | Irs2 | 3.81 | 2.61E-12 |
| 354 | Zc3h13 | 3.74 | 2.61E-12 |
| 355 | Pik3r1 | 2.05 | 2.62E-12 |
| 356 | LOC114690756 | 3.56 | 2.63E-12 |
| 357 | Ecel1 | 7.80 | 2.94E-12 |
| 358 | Trim56 | 2.16 | 3.42E-12 |
| 359 | Arl4d | 2.01 | 3.61E-12 |
| 360 | Spred3 | 4.14 | 4.01E-12 |
| 361 | Rnf122 | 4.58 | 4.25E-12 |
| 362 | Kif12 | 3.86 | 4.37E-12 |
| 363 | LOC114690680 | 4.14 | 5.05E-12 |
| 364 | Arrdc4 | 3.98 | 5.13E-12 |
| 365 | Nlrp3 | 3.80 | 5.32E-12 |
| 366 | Retreg1 | 2.24 | 6.08E-12 |
| 367 | LOC114698341 | 3.43 | 6.11E-12 |
| 368 | LOC114680819 | 2.36 | 6.72E-12 |
| 369 | Ciita | 2.94 | 7.42E-12 |
| 370 | LOC114701932 | 2.72 | 7.42E-12 |
| 371 | Vgf | 5.32 | 8.30E-12 |
| 372 | Il17rb | 3.50 | 8.54E-12 |
| 373 | Rftn1 | 2.20 | 9.15E-12 |
| 374 | LOC114699623 | 4.71 | 9.35E-12 |
| 375 | LOC114689091 | 2.97 | 1.00E-11 |
| 658 | Slc7a11 | 4.15 | 7.44E-07 |
| 659 | LOC114684570 | 6.14 | 7.99E-07 |
| 660 | Slc1a3 | 3.46 | 8.75E-07 |
| 661 | Fam43a | 2.38 | 1.02E-06 |
| 662 | LOC114708227 | 4.05 | 1.08E-06 |
| 663 | Serpinb10 | 2.46 | 1.18E-06 |
| 664 | LOC114706029 | 2.11 | 1.23E-06 |
| 665 | Gadd45a | 2.11 | 1.23E-06 |
| 666 | Casp3 | 2.14 | 1.24E-06 |
| 667 | LOC114683936 | 2.19 | 1.24E-06 |
| 668 | Traf1 | 3.41 | 1.28E-06 |
| 669 | Il1rl1 | 3.94 | 1.28E-06 |
| 670 | Mustn1 | 2.68 | 1.29E-06 |
| 671 | LOC114686085 | 3.46 | 1.50E-06 |
| 672 | LOC114693047 | 4.64 | 1.61E-06 |
| 673 | LOC114680745 | 2.04 | 1.66E-06 |
| 674 | LOC114698875 | 2.05 | 1.75E-06 |
| 675 | LOC114708624 | 2.74 | 1.75E-06 |
| 676 | LOC114687187 | 2.09 | 1.80E-06 |
| 677 | Lysmd2 | 2.70 | 2.09E-06 |
| 678 | Izumo1 | 3.26 | 2.15E-06 |
| 679 | Pla2g10 | 2.06 | 2.17E-06 |
| 680 | Slfn11 | 5.10 | 2.29E-06 |
| 681 | Cd86 | 2.13 | 2.33E-06 |
| 682 | Kcnj14 | 3.20 | 2.39E-06 |
| 683 | Ace | 2.22 | 2.41E-06 |
| 684 | Pnma1 | 3.15 | 2.43E-06 |
| 685 | LOC114709406 | 2.34 | 2.46E-06 |
| 686 | Mia | 2.18 | 2.70E-06 |
| 687 | LOC114694945 | 2.18 | 2.73E-06 |
| 688 | Cnn3 | 2.45 | 2.74E-06 |
| 689 | LOC114690754 | 4.04 | 2.78E-06 |
| 690 | LOC114708200 | 2.73 | 3.18E-06 |
| 691 | Klhl29 | 2.13 | 3.18E-06 |
| 692 | LOC114701389 | 2.19 | 3.30E-06 |
| 693 | LOC114682733 | 5.39 | 3.33E-06 |
| 694 | LOC114687186 | 2.10 | 3.49E-06 |
| 695 | LOC114685582 | 3.65 | 3.59E-06 |
| 696 | Kcnd1 | 2.42 | 3.67E-06 |
| 697 | LOC114706385 | 4.36 | 3.76E-06 |
| 698 | LOC114684575 | 9.28 | 3.86E-06 |
| 699 | LOC114708221 | 2.48 | 4.11E-06 |
| 700 | Calhm6 | 4.12 | 4.23E-06 |
| 701 | LOC114701719 | 4.31 | 4.49E-06 |
| 702 | Pcdh19 | 2.02 | 4.52E-06 |
| 703 | Ramp3 | 2.83 | 4.56E-06 |
| 704 | LOC114706405 | 2.14 | 4.62E-06 |
| 705 | Myo1g | 2.09 | 5.00E-06 |
| 706 | Nrn1l | 2.39 | 5.08E-06 |
| 707 | Noxo1 | 2.14 | 5.18E-06 |
| 708 | LOC114708299 | 3.87 | 5.34E-06 |
| 709 | Hes3 | 4.91 | 5.45E-06 |
| 710 | Taf4b | 2.05 | 5.64E-06 |
| 711 | Lbh | 2.10 | 6.39E-06 |
| 712 | LOC114700344 | 5.30 | 6.85E-06 |
| 713 | Cd83 | 2.02 | 7.28E-06 |
| 714 | Ddx43 | 5.47 | 7.56E-06 |
| 715 | LOC114701776 | 2.15 | 7.59E-06 |
| 716 | Plcd4 | 3.32 | 7.87E-06 |
| 717 | LOC114706384 | 4.66 | 8.35E-06 |
| 718 | LOC114687955 | 2.25 | 8.38E-06 |
| 719 | LOC114692131 | 3.45 | 8.43E-06 |
| 720 | Glipr2 | 2.43 | 8.51E-06 |
| 721 | LOC114687661 | 3.16 | 8.69E-06 |
| 722 | LOC114699541 | 2.05 | 9.04E-06 |
| 723 | LOC114684343 | 2.34 | 9.23E-06 |
| 724 | Il20rb | 2.41 | 9.76E-06 |
| 725 | LOC114684569 | 6.75 | 1.01E-05 |
| 726 | LOC114691891 | 2.20 | 1.05E-05 |
| 727 | Medag | 2.10 | 1.06E-05 |
| 728 | LOC114695310 | 2.86 | 1.10E-05 |
| 729 | Slc22a14 | 3.81 | 1.13E-05 |
| 730 | LOC114700346 | 4.30 | 1.13E-05 |
| 731 | Lrrc73 | 2.72 | 1.25E-05 |
| 732 | LOC114696386 | 2.20 | 1.26E-05 |
| 733 | Kcnc4 | 3.35 | 1.26E-05 |
| 734 | Agpat4 | 2.09 | 1.29E-05 |
| 735 | Oas3 | 3.52 | 1.36E-05 |
| 736 | LOC114694399 | 2.22 | 1.37E-05 |
| 737 | LOC114706404 | 2.40 | 1.50E-05 |
| 738 | LOC114688813 | 2.07 | 1.53E-05 |
| 739 | Stat4 | 3.89 | 1.71E-05 |
| 740 | Fgf23 | 7.29 | 1.86E-05 |
| 741 | LOC114690363 | 2.44 | 1.96E-05 |
| 742 | Mcoln2 | 2.91 | 1.97E-05 |
| 743 | Anxa1 | 2.03 | 2.01E-05 |
| 744 | LOC114688817 | 2.69 | 2.16E-05 |
| 745 | Col20a1 | 2.75 | 2.18E-05 |
| 746 | LOC114680820 | 2.43 | 2.23E-05 |
| 747 | Il27 | 3.28 | 2.26E-05 |
| 748 | Edem1 | 2.26 | 2.26E-05 |
| 749 | LOC114689754 | 2.65 | 2.32E-05 |
| 750 | Pou2f2 | 2.05 | 2.38E-05 |
| 1033 | Nlgn3 | 2.99 | 3.02E-02 |
| 1034 | Vwa3b | 4.71 | 3.19E-02 |
| 1035 | LOC114687326 | 2.41 | 3.24E-02 |
| 1036 | LOC114684495 | 2.78 | 3.25E-02 |
| 1037 | CUNHXorf65 | 3.55 | 3.33E-02 |
| 1038 | Htr2a | 3.61 | 3.33E-02 |
| 1039 | LOC114680936 | 3.55 | 3.33E-02 |
| 1040 | LOC114682413 | 3.55 | 3.33E-02 |
| 1041 | CUNH1orf185 | 2.58 | 3.35E-02 |
| 1042 | Lta | 4.29 | 3.36E-02 |
| 1043 | LOC114690782 | 4.16 | 3.38E-02 |
| 1044 | Znf502 | 2.28 | 3.43E-02 |
| 1045 | CUNH9orf57 | 2.25 | 3.44E-02 |
| 1046 | Padi4 | 2.08 | 3.57E-02 |
| 1047 | Rassf10 | 3.75 | 3.57E-02 |
| 1048 | Cpne6 | 3.32 | 3.61E-02 |
| 1049 | LOC114709676 | 3.85 | 3.74E-02 |
| 1050 | LOC114684606 | 2.17 | 3.76E-02 |
| 1051 | LOC114708476 | 2.38 | 3.83E-02 |
| 1052 | Ucp3 | 2.28 | 3.90E-02 |
| 1053 | LOC114699542 | 2.14 | 4.24E-02 |
| 1054 | Slc44a5 | 4.51 | 4.37E-02 |
| 1055 | LOC114694459 | 3.74 | 4.40E-02 |
| 1056 | Sim1 | 2.68 | 4.41E-02 |
| 1057 | LOC114702645 | 2.32 | 4.61E-02 |
| 1058 | Islr2 | 3.10 | 4.62E-02 |
| 1059 | Rgs1 | 2.21 | 4.68E-02 |
| 1060 | Fam83c | 2.12 | 4.72E-02 |
| 1061 | Epb42 | 2.89 | 4.80E-02 |
| 1062 | LOC114703139 | 2.57 | 4.83E-02 |
| 1063 | LOC114698397 | 2.53 | 4.90E-02 |
| 1064 | Osm | 2.76 | 4.90E-02 |
| 1065 | Pnmt | 3.45 | 4.92E-02 |
| 1066 | Gje1 | 2.43 | 4.94E-02 |
| 1067 | Olig1 | 2.13 | 4.96E-02 |
| 1068 | Hhla2 | 3.17 | 4.97E-02 |
| 1069 | Fcer1a | 2.08 | 4.99E-02 |

**Table S7. DEGs of the blood of M. musculus**

| rank | gene | log2 fold change | FDR | rank | gene | log2 fold change | FDR |
|---|---|---|---|---|---|---|---|
| 1 | Stat2 | 2.87 | 3.38E-116 | 251 | A330040F15Rik | 2.53 | 6.44E-10 |
| 2 | Parp14 | 3.28 | 8.50E-115 | 252 | Gm37347 | 4.29 | 6.62E-10 |
| 3 | Usp18 | 4.43 | 1.10E-113 | 253 | Mfsd7a | 2.78 | 8.61E-10 |
| 4 | Irgm1 | 3.20 | 3.28E-109 | 254 | Gm15056 | 6.47 | 1.16E-09 |
| 5 | Eif2ak2 | 3.98 | 2.01E-108 | 255 | Ankrd66 | 3.54 | 1.80E-09 |
| 6 | Gbp5 | 4.27 | 9.13E-108 | 256 | Sorbs1 | 2.06 | 2.45E-09 |
| 7 | Xaf1 | 3.05 | 1.02E-107 | 257 | Gm45418 | 3.24 | 4.35E-09 |
| 8 | Rnf213 | 3.50 | 4.66E-104 | 258 | Fnbp1l | 2.93 | 5.80E-09 |
| 9 | Zbp1 | 3.97 | 2.79E-101 | 259 | Rilpl1 | 2.04 | 6.13E-09 |
| 10 | Ifit1 | 4.51 | 5.64E-93 | 260 | Ifnlr1 | 2.62 | 6.94E-09 |
| 11 | Herc6 | 3.86 | 8.57E-92 | 261 | Mmp25 | 2.83 | 7.55E-09 |
| 12 | Dhx58 | 3.59 | 3.00E-90 | 262 | Mt2 | 5.85 | 9.05E-09 |
| 13 | Parp9 | 2.59 | 1.96E-89 | 263 | Carmil1 | 2.36 | 9.72E-09 |
| 14 | Gbp2 | 3.54 | 3.25E-89 | 264 | Cldn1 | 2.32 | 1.01E-08 |
| 15 | Slfn5 | 3.52 | 1.29E-86 | 265 | Siglece | 2.48 | 1.30E-08 |
| 16 | Parp12 | 4.18 | 9.47E-86 | 266 | Clec4e | 3.00 | 1.82E-08 |
| 17 | Lipg | 9.78 | 3.26E-84 | 267 | Gas7 | 2.47 | 2.05E-08 |
| 18 | Ifit3 | 4.55 | 1.73E-81 | 268 | Ltf | 4.07 | 2.09E-08 |
| 19 | Gbp7 | 3.03 | 2.22E-79 | 269 | Gm43694 | 2.01 | 2.44E-08 |
| 20 | Nos2 | 8.95 | 1.37E-78 | 270 | Gm28914 | 2.65 | 2.97E-08 |
| 21 | Gbp2b | 3.88 | 3.12E-76 | 271 | Hp | 2.33 | 3.62E-08 |
| 22 | Trim30c | 4.69 | 3.00E-75 | 272 | Rab20 | 2.46 | 4.23E-08 |
| 23 | Parp10 | 2.32 | 6.11E-74 | 273 | Gm10851 | 4.18 | 4.34E-08 |
| 24 | Ifih1 | 3.92 | 1.08E-73 | 274 | Mefv | 2.48 | 5.04E-08 |
| 25 | Slfn8 | 2.59 | 6.54E-73 | 275 | Gm5483 | 2.46 | 5.21E-08 |
| 26 | Ddx58 | 2.49 | 9.05E-73 | 276 | A930037H05Rik | 2.09 | 5.41E-08 |
| 27 | Dtx3l | 2.46 | 1.78E-72 | 277 | Il1f9 | 2.46 | 5.61E-08 |
| 28 | Ifi206 | 3.97 | 8.33E-72 | 278 | Id1 | 2.37 | 6.85E-08 |
| 29 | Znfx1 | 2.32 | 4.74E-71 | 279 | Mxd1 | 2.20 | 8.09E-08 |
| 30 | Trim30a | 2.94 | 5.77E-71 | 280 | Itgb8 | 3.95 | 9.89E-08 |
| 31 | Tor3a | 3.60 | 9.92E-71 | 281 | Hmgn3 | 2.49 | 1.06E-07 |
| 32 | Cxcl10 | 9.94 | 2.87E-70 | 282 | Sema4c | 2.85 | 1.81E-07 |
| 33 | Mx2 | 5.07 | 3.00E-69 | 283 | Maff | 2.90 | 2.54E-07 |
| 34 | Socs1 | 3.81 | 3.94E-66 | 284 | Col9a3 | 2.10 | 2.60E-07 |
| 35 | Ifi47 | 2.64 | 9.17E-66 | 285 | Hap1 | 2.52 | 2.78E-07 |
| 36 | Marcksl1 | 4.78 | 4.06E-65 | 286 | Anxa1 | 2.10 | 3.69E-07 |
| 37 | Irgm2 | 2.79 | 8.69E-65 | 287 | Ypel4 | 2.61 | 3.86E-07 |
| 38 | Daxx | 2.66 | 1.04E-64 | 288 | Mrgpra2a | 3.68 | 4.18E-07 |
| 39 | Mx1 | 4.95 | 4.19E-64 | 289 | Asb2 | 2.14 | 4.35E-07 |
| 40 | Iigp1 | 4.01 | 2.02E-63 | 290 | Il27 | 2.63 | 4.50E-07 |
| 41 | Ccl4 | 4.94 | 4.35E-62 | 291 | Gm21188 | 2.07 | 5.00E-07 |
| 42 | Cmpk2 | 4.34 | 1.03E-60 | 292 | Il10 | 4.23 | 5.49E-07 |
| 43 | Ifit3b | 4.65 | 2.18E-58 | 293 | Adora2b | 2.76 | 5.63E-07 |
| 44 | Trafd1 | 2.88 | 1.93E-57 | 294 | Gucy2g | 4.67 | 6.53E-07 |
| 45 | Cd274 | 2.58 | 2.64E-57 | 295 | Csf3 | 5.57 | 7.80E-07 |
| 46 | Pfkfb3 | 2.79 | 6.38E-57 | 296 | Tigit | 2.13 | 7.92E-07 |
| 47 | Gbp3 | 2.85 | 2.11E-56 | 297 | Cxcl3 | 3.44 | 8.47E-07 |
| 48 | Ifi44 | 4.43 | 4.83E-56 | 298 | Ms4a4a | 2.05 | 8.62E-07 |
| 49 | Nampt | 2.56 | 5.12E-56 | 299 | Ptges | 4.19 | 8.64E-07 |
| 50 | Ifi35 | 2.38 | 4.78E-55 | 300 | Tbc1d2 | 2.65 | 9.87E-07 |
| 51 | Tnfaip3 | 4.03 | 5.66E-54 | 301 | Fap | 3.20 | 1.13E-06 |
| 52 | Zup1 | 2.37 | 6.42E-54 | 302 | Gm43181 | 2.23 | 1.13E-06 |
| 53 | Helz2 | 2.17 | 1.58E-52 | 303 | Gpr27 | 4.34 | 1.21E-06 |
| 54 | Oas1b | 2.18 | 8.78E-51 | 304 | Mt1 | 4.23 | 1.78E-06 |
| 55 | Cish | 2.88 | 3.91E-50 | 305 | S100a8 | 2.50 | 1.95E-06 |
| 56 | Bst2 | 2.96 | 1.16E-49 | 306 | Mpo | 3.09 | 2.02E-06 |
| 57 | Ccl3 | 4.49 | 2.90E-49 | 307 | Gm6377 | 2.28 | 2.32E-06 |
| 58 | Mov10 | 2.32 | 3.27E-49 | 308 | Gm38554 | 2.37 | 2.38E-06 |
| 59 | Pik3ap1 | 2.11 | 1.71E-48 | 309 | Sult6b2 | 4.34 | 2.53E-06 |
| 60 | Ifi44l | 3.69 | 2.36E-48 | 310 | Spic | 2.14 | 2.53E-06 |
| 61 | Ifi209 | 2.37 | 4.89E-48 | 311 | Bst1 | 2.12 | 4.82E-06 |
| 62 | Sod2 | 2.65 | 9.46E-47 | 312 | Clnk | 2.15 | 5.06E-06 |
| 63 | Bcl2a1b | 3.26 | 1.52E-46 | 313 | Sphk1 | 2.79 | 5.27E-06 |
| 64 | Bcl2a1a | 4.38 | 8.17E-46 | 314 | Gm47243 | 4.38 | 5.56E-06 |
| 65 | Il1rn | 4.91 | 1.21E-45 | 315 | Gpr31b | 4.05 | 6.01E-06 |
| 66 | Fmnl2 | 5.02 | 1.73E-45 | 316 | Gm42462 | 2.34 | 6.78E-06 |
| 67 | Tet2 | 2.13 | 5.51E-45 | 317 | Tarm1 | 2.04 | 7.09E-06 |
| 68 | Phf11b | 2.85 | 2.07E-44 | 318 | Pnpla1 | 2.75 | 7.12E-06 |
| 69 | Gm43302 | 3.66 | 2.41E-44 | 319 | Kif1a | 3.29 | 8.40E-06 |
| 70 | Oas1a | 3.72 | 3.45E-44 | 320 | Rhov | 3.09 | 9.49E-06 |
| 71 | Ifi204 | 2.74 | 5.80E-44 | 321 | Gm43068 | 3.09 | 1.01E-05 |
| 72 | Bcl3 | 2.65 | 8.39E-44 | 322 | Tnfaip6 | 3.18 | 1.18E-05 |
| 73 | Lgals9 | 2.19 | 3.47E-43 | 323 | Cxcl9 | 3.37 | 1.56E-05 |
| 74 | Ifi214 | 2.85 | 3.71E-43 | 324 | Tlr3 | 2.05 | 1.81E-05 |
| 75 | Icam1 | 2.88 | 4.55E-43 | 325 | C3 | 2.15 | 1.86E-05 |
| 76 | Plscr1 | 3.19 | 5.66E-43 | 326 | Gm7457 | 2.96 | 1.90E-05 |
| 77 | Nfkbia | 2.68 | 6.88E-42 | 327 | P2ry14 | 2.14 | 1.92E-05 |
| 78 | Epsti1 | 2.09 | 8.35E-42 | 328 | Gm16175 | 3.63 | 2.09E-05 |
| 79 | Oasl1 | 3.64 | 1.45E-41 | 329 | Dhx58os | 4.41 | 2.14E-05 |
| 80 | Igtp | 2.21 | 2.26E-41 | 330 | Mlkl | 2.39 | 2.22E-05 |
| 81 | Gbp9 | 2.41 | 2.69E-41 | 331 | Pdcd1 | 2.45 | 2.34E-05 |
| 82 | Ly6a | 4.38 | 1.97E-40 | 332 | Hcar2 | 2.34 | 2.50E-05 |
| 83 | Trex1 | 2.58 | 2.26E-40 | 333 | Ly6g | 2.94 | 2.78E-05 |
| 84 | Slfn9 | 2.14 | 3.35E-40 | 334 | Ly6i | 2.62 | 2.84E-05 |
| 85 | Batf | 2.34 | 1.09E-38 | 335 | Ccm2l | 2.10 | 2.94E-05 |
| 86 | Trim30d | 4.79 | 2.61E-38 | 336 | Lif | 3.41 | 2.97E-05 |
| 87 | Oas3 | 3.20 | 2.65E-38 | 337 | Retnlg | 2.03 | 3.07E-05 |
| 88 | Casp4 | 3.26 | 8.94E-38 | 338 | Gm45193 | 3.43 | 3.54E-05 |
| 89 | Ifi213 | 2.91 | 1.06E-37 | 339 | Adap2 | 2.60 | 3.81E-05 |
| 90 | Gbp4 | 2.76 | 1.09E-37 | 340 | Anxa3 | 2.97 | 4.54E-05 |
| 91 | Fcgr1 | 3.27 | 2.35E-37 | 341 | Mmp13 | 2.03 | 4.66E-05 |
| 92 | Ikbke | 2.02 | 3.22E-37 | 342 | F730016J06Rik | 2.56 | 4.90E-05 |
| 93 | Gzmb | 4.36 | 7.07E-37 | 343 | S100a9 | 2.07 | 5.22E-05 |
| 94 | Ddx60 | 3.22 | 3.06E-36 | 344 | Mir155hg | 2.67 | 6.26E-05 |
| 95 | Irak3 | 3.03 | 3.06E-36 | 345 | Gm13822 | 2.02 | 6.84E-05 |
| 96 | Gbp10 | 4.24 | 3.61E-36 | 346 | Gm4951 | 2.46 | 6.99E-05 |
| 97 | Ifit2 | 2.26 | 1.88E-35 | 347 | Olfml2b | 3.21 | 7.32E-05 |
| 98 | Serpina3g | 3.34 | 3.82E-35 | 348 | Stfa2 | 2.80 | 8.27E-05 |
| 99 | Lcn2 | 5.77 | 4.18E-35 | 349 | Rasip1 | 2.18 | 8.38E-05 |
| 100 | Tifa | 2.30 | 5.47E-35 | 350 | Ear6 | 5.10 | 8.40E-05 |
| 101 | Tgtp2 | 2.36 | 8.36E-35 | 351 | Gm27167 | 4.90 | 9.32E-05 |
| 102 | Rgs1 | 4.35 | 2.19E-34 | 352 | Gm11626 | 2.63 | 1.09E-04 |
| 103 | Gpr84 | 6.65 | 1.58E-33 | 353 | Ccl7 | 4.06 | 1.23E-04 |
| 104 | Samsn1 | 2.76 | 7.48E-33 | 354 | Orm1 | 3.94 | 1.40E-04 |
| 105 | Rtp4 | 2.64 | 1.18E-32 | 355 | AC025794.1 | 3.72 | 1.46E-04 |
| 106 | Serpina3f | 4.26 | 2.12E-32 | 356 | Gm19719 | 2.22 | 1.60E-04 |
| 107 | Ifi208 | 2.35 | 5.09E-32 | 357 | Gm36186 | 2.59 | 2.09E-04 |
| 108 | Oas2 | 3.32 | 6.74E-32 | 358 | Gm15845 | 2.23 | 2.10E-04 |
| 109 | Map3k8 | 2.74 | 7.17E-32 | 359 | Ciart | 2.51 | 2.14E-04 |
| 110 | Phf11c | 2.64 | 2.35E-31 | 360 | Tgm1 | 2.25 | 2.39E-04 |
| 111 | Ppa1 | 3.16 | 1.50E-30 | 361 | 4930512H18Rik | 2.15 | 2.50E-04 |
| 112 | Ebi3 | 3.38 | 3.35E-30 | 362 | Vmn2r16 | 2.24 | 2.84E-04 |
| 113 | 1600014C10Rik | 2.25 | 4.31E-30 | 363 | Htra1 | 3.41 | 2.99E-04 |
| 114 | Tgtp1 | 2.32 | 4.68E-30 | 364 | Hsd3b2 | 2.42 | 3.36E-04 |
| 115 | AA467197 | 4.57 | 1.56E-29 | 365 | U90926 | 4.26 | 3.38E-04 |
| 116 | Dach1 | 6.02 | 6.14E-29 | 366 | 4930599N23Rik | 2.56 | 3.44E-04 |
| 117 | Atp10a | 2.27 | 2.38E-28 | 367 | F830208F22Rik | 2.02 | 3.66E-04 |
| 118 | Edn1 | 7.60 | 2.92E-28 | 368 | Trpm6 | 3.19 | 3.71E-04 |
| 119 | Gbp11 | 4.57 | 4.05E-28 | 369 | Gm37529 | 2.38 | 4.02E-04 |
| 120 | Serpinb9 | 2.36 | 8.40E-28 | 370 | 1700047M11Rik | 2.68 | 4.26E-04 |
| 121 | Isg15 | 2.19 | 1.09E-27 | 371 | Ccl22 | 2.28 | 4.41E-04 |
| 122 | Oasl2 | 3.82 | 1.19E-27 | 372 | Adtrp | 2.26 | 5.11E-04 |
| 123 | Mapkapk2 | 2.15 | 1.97E-27 | 373 | Cxcl1 | 4.57 | 5.53E-04 |
| 124 | Irf7 | 2.16 | 8.09E-27 | 374 | Gm20743 | 2.80 | 5.87E-04 |
| 125 | Mmp14 | 4.55 | 1.72E-26 | 375 | 2810403G07Rik | 3.63 | 6.06E-04 |
| 126 | Thbs1 | 2.76 | 2.54E-26 | 376 | Prg2 | 4.00 | 6.25E-04 |

Table S7

| | | | |
|---|---|---|---|
| 127 | Plac8 | 2.82 | 5.08E-26 |
| 128 | Il1a | 5.44 | 5.36E-26 |
| 129 | Ankrd22 | 5.14 | 6.27E-26 |
| 130 | Dusp16 | 3.24 | 9.10E-26 |
| 131 | Ets2 | 2.25 | 9.94E-26 |
| 132 | Nfkbiz | 3.04 | 1.02E-25 |
| 133 | Ifit1bl1 | 2.85 | 1.43E-25 |
| 134 | Schip1 | 4.15 | 1.49E-25 |
| 135 | Gadd45b | 2.85 | 1.87E-25 |
| 136 | Phf11a | 3.21 | 4.30E-25 |
| 137 | N4bp1 | 2.15 | 6.71E-25 |
| 138 | Rnd1 | 3.70 | 6.74E-25 |
| 139 | Fas | 2.23 | 7.02E-25 |
| 140 | Gm43197 | 2.04 | 1.14E-24 |
| 141 | Upp1 | 5.77 | 2.19E-24 |
| 142 | Gk | 3.23 | 3.00E-24 |
| 143 | Acod1 | 6.13 | 9.91E-24 |
| 144 | Olfm4 | 5.03 | 1.12E-23 |
| 145 | Gm47242 | 3.43 | 1.78E-23 |
| 146 | Ifi211 | 2.39 | 8.87E-23 |
| 147 | Ier3 | 4.06 | 1.21E-22 |
| 148 | Slc15a3 | 2.98 | 2.08E-22 |
| 149 | Tnfsf10 | 2.18 | 2.43E-22 |
| 150 | Prok2 | 7.49 | 3.06E-22 |
| 151 | Tmem45a2 | 6.71 | 6.04E-22 |
| 152 | Oas1g | 3.86 | 9.30E-22 |
| 153 | Ntng2 | 2.52 | 1.02E-21 |
| 154 | Gbp6 | 3.43 | 1.29E-21 |
| 155 | Xcl1 | 3.48 | 1.93E-20 |
| 156 | Hck | 2.15 | 1.95E-20 |
| 157 | Tnip3 | 3.85 | 2.47E-20 |
| 158 | AC119228.1 | 2.33 | 3.12E-20 |
| 159 | Ifi202b | 3.25 | 3.19E-20 |
| 160 | Tnf | 4.20 | 3.39E-20 |
| 161 | Aoah | 2.22 | 5.25E-20 |
| 162 | Nmral1 | 2.62 | 6.23E-20 |
| 163 | Jun | 2.19 | 1.16E-19 |
| 164 | Tlr2 | 3.28 | 2.47E-19 |
| 165 | Ccrl2 | 3.98 | 2.51E-19 |
| 166 | Ly6c2 | 2.05 | 2.95E-19 |
| 167 | Gadd45g | 2.78 | 5.41E-19 |
| 168 | Neb | 2.05 | 8.87E-19 |
| 169 | Phf11d | 2.72 | 9.50E-19 |
| 170 | Alpk1 | 2.76 | 9.85E-19 |
| 171 | Gm5431 | 2.24 | 2.07E-18 |
| 172 | Cd69 | 2.00 | 8.64E-18 |
| 173 | Cxcl11 | 5.56 | 9.05E-18 |
| 174 | Rhou | 3.39 | 1.95E-17 |
| 175 | Cgas | 2.21 | 2.54E-17 |
| 176 | Tnfrsf9 | 3.53 | 2.55E-17 |
| 177 | Trib1 | 2.00 | 3.08E-17 |
| 178 | Ifi205 | 3.74 | 3.80E-17 |
| 179 | Dstn | 2.66 | 5.66E-17 |
| 180 | Il12rb1 | 2.43 | 5.74E-17 |
| 181 | Snai1 | 6.65 | 7.09E-17 |
| 182 | Ccr5 | 2.43 | 9.91E-17 |
| 183 | Adora3 | 3.49 | 1.11E-16 |
| 184 | Socs3 | 2.78 | 1.34E-16 |
| 185 | Gm12185 | 2.17 | 1.59E-16 |
| 186 | Setdb2 | 2.05 | 1.68E-16 |
| 187 | Ly6c1 | 2.12 | 3.87E-16 |
| 188 | Gm12367 | 3.66 | 3.97E-16 |
| 189 | Ifit1bl2 | 2.56 | 6.07E-16 |
| 190 | Slfn4 | 3.68 | 6.48E-16 |
| 191 | Olfr56 | 3.43 | 1.45E-15 |
| 192 | Serpinb9b | 2.43 | 1.50E-15 |
| 193 | Chil1 | 3.76 | 2.05E-15 |
| 194 | Cd14 | 4.59 | 2.05E-15 |
| 195 | Fpr2 | 3.46 | 3.20E-15 |
| 196 | Dusp1 | 2.70 | 3.82E-15 |
| 197 | Itgb2l | 4.75 | 7.06E-15 |
| 198 | Hdc | 3.07 | 7.21E-15 |
| 199 | Pkib | 2.15 | 8.16E-15 |
| 200 | Chil3 | 3.51 | 8.27E-15 |
| 201 | Slc7a11 | 3.02 | 1.06E-14 |
| 202 | Gca | 2.93 | 1.54E-14 |
| 203 | Bcl2a1d | 2.30 | 2.34E-14 |
| 204 | Ifng | 5.16 | 3.14E-14 |
| 205 | Ifitm3 | 2.04 | 3.81E-14 |
| 206 | Met | 2.96 | 4.18E-14 |
| 207 | Ifitm6 | 3.42 | 8.71E-14 |
| 208 | Ifi207 | 2.48 | 1.27E-13 |
| 209 | Car13 | 3.95 | 1.36E-13 |
| 210 | Tfec | 2.75 | 1.38E-13 |
| 211 | Cxcl2 | 6.37 | 3.33E-13 |
| 212 | Gm47761 | 2.07 | 4.27E-13 |
| 213 | 6720489N17Rik | 2.60 | 5.84E-13 |
| 214 | Inhba | 4.46 | 6.44E-13 |
| 215 | Slco4c1 | 2.59 | 8.83E-13 |
| 216 | Gm16712 | 3.57 | 9.73E-13 |
| 217 | Il1r2 | 3.69 | 1.50E-12 |
| 218 | Clec5a | 3.26 | 2.07E-12 |
| 219 | Slc28a3 | 4.29 | 2.11E-12 |
| 220 | Fpr1 | 3.27 | 2.74E-12 |
| 221 | Il1rap | 2.45 | 3.40E-12 |
| 222 | Ccl2 | 5.70 | 9.64E-12 |
| 223 | Stfa3 | 4.86 | 1.03E-11 |
| 224 | Wfdc21 | 3.64 | 1.12E-11 |
| 225 | Gm26822 | 2.53 | 1.15E-11 |
| 226 | Abca13 | 4.86 | 1.26E-11 |
| 227 | Il18bp | 2.57 | 1.51E-11 |
| 228 | Ffar2 | 3.37 | 2.02E-11 |
| 229 | Atp11a | 2.07 | 2.31E-11 |
| 230 | Ms4a6d | 2.26 | 2.68E-11 |
| 231 | Nr4a3 | 2.48 | 4.53E-11 |
| 232 | Ngp | 5.18 | 4.93E-11 |
| 233 | Il12rb2 | 2.23 | 5.50E-11 |
| 234 | Vcan | 3.84 | 6.52E-11 |
| 235 | Timp1 | 5.08 | 8.27E-11 |
| 236 | Dgat2 | 2.86 | 9.07E-11 |
| 237 | Rag1 | 4.87 | 1.12E-10 |
| 238 | Camp | 4.65 | 1.65E-10 |
| 239 | Cd177 | 3.40 | 1.73E-10 |
| 240 | Eppk1 | 3.19 | 1.94E-10 |
| 241 | Wdfy3 | 2.36 | 2.33E-10 |
| 242 | Muc1 | 3.27 | 2.55E-10 |
| 243 | Mrgpra2b | 3.15 | 3.02E-10 |
| 244 | Gm2511 | 2.96 | 3.17E-10 |
| 245 | Gm42763 | 2.83 | 4.11E-10 |
| 246 | Klf5 | 2.60 | 4.41E-10 |
| 247 | Mmp8 | 3.28 | 4.43E-10 |
| 248 | Gm49751 | 4.35 | 5.07E-10 |
| 249 | Nek2 | 2.27 | 5.81E-10 |
| 250 | Zfp365 | 2.19 | 6.01E-10 |
| 377 | Fst | 3.01 | 6.55E-04 |
| 378 | Ston1 | 2.18 | 6.56E-04 |
| 379 | Gm44165 | 2.51 | 8.37E-04 |
| 380 | Six3os1 | 2.67 | 9.74E-04 |
| 381 | Msantd3 | 2.38 | 9.85E-04 |
| 382 | Adamts9 | 3.37 | 1.16E-03 |
| 383 | A930024N18Rik | 3.63 | 1.31E-03 |
| 384 | Gm12253 | 2.59 | 1.32E-03 |
| 385 | Plscr2 | 2.30 | 1.36E-03 |
| 386 | Ccl12 | 4.22 | 1.41E-03 |
| 387 | Gm15856 | 3.79 | 1.45E-03 |
| 388 | Cfb | 2.81 | 1.46E-03 |
| 389 | Pnpla1os | 3.29 | 1.67E-03 |
| 390 | 4833419F23Rik | 2.14 | 1.77E-03 |
| 391 | Ear1 | 5.60 | 1.84E-03 |
| 392 | Lilrb4a | 2.20 | 1.96E-03 |
| 393 | Gm26535 | 2.54 | 2.15E-03 |
| 394 | Gm49482 | 4.06 | 2.35E-03 |
| 395 | Gipc3 | 2.06 | 2.48E-03 |
| 396 | Bcat1 | 3.12 | 2.73E-03 |
| 397 | Gm39556 | 2.06 | 2.82E-03 |
| 398 | Gm49730 | 2.86 | 3.10E-03 |
| 399 | Gm43635 | 2.89 | 3.20E-03 |
| 400 | Hspa1a | 3.45 | 3.23E-03 |
| 401 | Mfsd6l | 2.75 | 3.40E-03 |
| 402 | Gm38575 | 4.28 | 3.63E-03 |
| 403 | Fbxo39 | 2.32 | 3.63E-03 |
| 404 | Gm33326 | 4.26 | 3.67E-03 |
| 405 | Serpine1 | 3.43 | 3.85E-03 |
| 406 | Mfsd2a | 2.49 | 3.90E-03 |
| 407 | Gzmk | 2.18 | 4.62E-03 |
| 408 | F3 | 3.26 | 4.70E-03 |
| 409 | Igll1 | 4.41 | 4.82E-03 |
| 410 | Ptgs2 | 2.19 | 5.06E-03 |
| 411 | A530046M15Rik | 4.39 | 5.27E-03 |
| 412 | Gm7598 | 2.06 | 5.30E-03 |
| 413 | P2rx3 | 3.70 | 5.43E-03 |
| 414 | Fam107a | 2.41 | 6.74E-03 |
| 415 | Gm5416 | 2.17 | 7.09E-03 |
| 416 | Apold1 | 3.40 | 7.31E-03 |
| 417 | Drp2 | 2.04 | 7.36E-03 |
| 418 | Gm49393 | 2.40 | 7.86E-03 |
| 419 | Ctcflos | 2.57 | 8.20E-03 |
| 420 | Prss34 | 2.20 | 8.47E-03 |
| 421 | Gm16897 | 2.43 | 8.57E-03 |
| 422 | Hspa1b | 2.57 | 9.13E-03 |
| 423 | E230013L22Rik | 2.99 | 9.85E-03 |
| 424 | Vmn1r227 | 2.32 | 1.10E-02 |
| 425 | Lbp | 2.05 | 1.13E-02 |
| 426 | Muc4 | 2.53 | 1.15E-02 |
| 427 | Sema6a | 2.36 | 1.19E-02 |
| 428 | Mybph | 4.04 | 1.25E-02 |
| 429 | Has1 | 2.90 | 1.38E-02 |
| 430 | Gm26748 | 4.20 | 1.53E-02 |
| 431 | Rab15 | 4.39 | 1.53E-02 |
| 432 | F830016B08Rik | 2.04 | 1.55E-02 |
| 433 | Fcrlb | 3.11 | 1.57E-02 |
| 434 | Gm44750 | 2.82 | 1.57E-02 |
| 435 | Tsx | 3.07 | 1.62E-02 |
| 436 | Pln | 2.35 | 1.62E-02 |
| 437 | Gm15684 | 4.02 | 1.76E-02 |
| 438 | Olfr105-ps | 2.57 | 1.80E-02 |
| 439 | Lrrc2 | 2.37 | 1.90E-02 |
| 440 | Slc7a2 | 2.78 | 1.91E-02 |
| 441 | Gm11772 | 2.74 | 1.92E-02 |
| 442 | Gm42917 | 3.76 | 1.93E-02 |
| 443 | Gzmc | 2.47 | 2.02E-02 |
| 444 | Gm16741 | 2.46 | 2.20E-02 |
| 445 | Il1f6 | 4.10 | 2.21E-02 |
| 446 | Zfp819 | 3.01 | 2.27E-02 |
| 447 | Gm19684 | 3.08 | 2.35E-02 |
| 448 | Pcdh17 | 3.72 | 2.49E-02 |
| 449 | Gm28321 | 3.02 | 2.59E-02 |
| 450 | 4921509O07Rik | 2.74 | 2.77E-02 |
| 451 | Gm43814 | 2.20 | 2.87E-02 |
| 452 | Olfr808 | 3.17 | 3.00E-02 |
| 453 | Gm45137 | 2.05 | 3.05E-02 |
| 454 | Gm46519 | 2.13 | 3.15E-02 |
| 455 | Gng4 | 2.73 | 3.24E-02 |
| 456 | Gm37975 | 4.12 | 3.37E-02 |
| 457 | Tnfaip8l3 | 3.90 | 3.44E-02 |
| 458 | Gm16796 | 2.85 | 3.52E-02 |
| 459 | Btbd16 | 2.01 | 3.67E-02 |
| 460 | 4930579C12Rik | 2.79 | 3.73E-02 |
| 461 | Gm38304 | 2.97 | 3.81E-02 |
| 462 | Gm37131 | 3.50 | 4.06E-02 |
| 463 | Gm26637 | 3.94 | 4.07E-02 |
| 464 | Wnk3 | 2.49 | 4.08E-02 |
| 465 | Gm47403 | 3.05 | 4.24E-02 |
| 466 | Slfn5os | 3.48 | 4.36E-02 |
| 467 | Prom1 | 2.35 | 4.39E-02 |
| 468 | Gm16758 | 2.73 | 4.45E-02 |
| 469 | Gm44836 | 2.20 | 4.55E-02 |
| 470 | Tshz2 | 2.21 | 4.57E-02 |
| 471 | Adamts15 | 2.33 | 4.58E-02 |
| 472 | Lsmem2 | 2.03 | 4.79E-02 |
| 473 | Cts8 | 3.01 | 4.97E-02 |
| 474 | Thbs4 | 3.63 | 4.99E-02 |

**Table S8. DEGs of spleen of M. musculus**

| rank | gene | log2 fold change | FDR | rank | gene | log2 fold change | FDR |
|---|---|---|---|---|---|---|---|
| 1 | Socs3 | 4.30 | 2.96E-159 | 451 | 2810403G07Rik | 3.11 | 2.42E-16 |
| 2 | Cxcl2 | 7.72 | 9.31E-138 | 452 | Neb | 3.08 | 3.84E-16 |
| 3 | Irgm1 | 3.67 | 6.37E-137 | 453 | Pln | 4.16 | 4.27E-16 |
| 4 | Parp9 | 3.21 | 3.19E-136 | 454 | Gm43635 | 2.90 | 4.39E-16 |
| 5 | Serpina3i | 3.55 | 9.37E-131 | 455 | Lilrb4a | 2.11 | 5.72E-16 |
| 6 | Parp12 | 3.82 | 4.66E-127 | 456 | Slfn5os | 6.15 | 6.23E-16 |
| 7 | Il1rn | 5.91 | 3.13E-126 | 457 | Gpr171 | 2.35 | 8.49E-16 |
| 8 | Stat2 | 3.03 | 2.74E-118 | 458 | Crem | 2.30 | 1.01E-15 |
| 9 | Ifi204 | 4.02 | 4.06E-115 | 459 | Gm43196 | 2.38 | 1.55E-15 |
| 10 | Dhx58 | 3.99 | 4.14E-112 | 460 | Slc4a4 | 2.60 | 1.62E-15 |
| 11 | Ms4a6d | 4.29 | 6.51E-112 | 461 | Vwa3b | 2.60 | 2.07E-15 |
| 12 | AW011738 | 2.44 | 1.04E-111 | 462 | Gm44013 | 2.36 | 2.20E-15 |
| 13 | Mmp3 | 4.79 | 2.66E-111 | 463 | Phox2a | 4.37 | 2.27E-15 |
| 14 | Oasl2 | 4.36 | 7.10E-110 | 464 | Gm49730 | 3.89 | 2.48E-15 |
| 15 | Cdkn1a | 3.19 | 2.12E-106 | 465 | Gm16096 | 2.19 | 2.55E-15 |
| 16 | Eif2ak2 | 3.74 | 4.41E-105 | 466 | Gm47800 | 4.92 | 3.63E-15 |
| 17 | Tor3a | 3.65 | 1.50E-103 | 467 | Bbx | 2.14 | 5.14E-15 |
| 18 | Ifit1 | 5.78 | 1.67E-103 | 468 | Ifitm5 | 2.23 | 5.64E-15 |
| 19 | Ifi47 | 3.43 | 1.14E-102 | 469 | Gm40685 | 3.73 | 7.13E-15 |
| 20 | Herc6 | 3.88 | 2.95E-102 | 470 | Gm37043 | 4.44 | 7.65E-15 |
| 21 | Rtp4 | 3.95 | 5.08E-101 | 471 | Gm15845 | 2.42 | 8.33E-15 |
| 22 | Cxcl5 | 5.49 | 1.32E-95 | 472 | AC161438.1 | 4.07 | 9.10E-15 |
| 23 | Adar | 2.16 | 5.63E-95 | 473 | Ddn | 3.92 | 9.20E-15 |
| 24 | Ifih1 | 4.11 | 2.81E-94 | 474 | Gm6902 | 2.93 | 9.72E-15 |
| 25 | Mx2 | 5.12 | 7.23E-94 | 475 | AC147026.1 | 2.54 | 9.80E-15 |
| 26 | Igtp | 3.24 | 9.50E-94 | 476 | Car4 | 4.86 | 1.26E-14 |
| 27 | Ifi202b | 4.09 | 2.00E-93 | 477 | Creb5 | 2.92 | 1.67E-14 |
| 28 | Phf11d | 3.75 | 2.66E-93 | 478 | Slfn4 | 3.82 | 1.92E-14 |
| 29 | Parp14 | 3.77 | 3.53E-93 | 479 | Gm26637 | 2.71 | 2.20E-14 |
| 30 | Stra6 | 4.00 | 1.50E-92 | 480 | Tmem266 | 2.35 | 2.63E-14 |
| 31 | Daxx | 3.58 | 1.98E-90 | 481 | A930037H05Rik | 2.28 | 2.91E-14 |
| 32 | Maff | 3.29 | 1.51E-89 | 482 | Gm33994 | 2.11 | 3.71E-14 |
| 33 | Usp18 | 4.85 | 1.66E-88 | 483 | Ifng | 4.08 | 4.22E-14 |
| 34 | Cxcl1 | 7.24 | 3.42E-88 | 484 | Rsad2 | 2.73 | 5.83E-14 |
| 35 | Oasl1 | 5.05 | 3.42E-88 | 485 | AC118695.1 | 6.67 | 6.24E-14 |
| 36 | Gm45774 | 5.67 | 4.04E-88 | 486 | Olfr1396 | 2.55 | 7.76E-14 |
| 37 | Tdrd7 | 3.03 | 5.77E-88 | 487 | Il2ra | 2.33 | 7.94E-14 |
| 38 | Gpr31b | 3.99 | 5.77E-88 | 488 | Mxd1 | 2.08 | 8.50E-14 |
| 39 | Gm47140 | 3.51 | 1.10E-87 | 489 | Vmn1r76 | 2.18 | 1.14E-13 |
| 40 | Oas1b | 2.68 | 2.14E-87 | 490 | 0610043K17Rik | 2.22 | 1.21E-13 |
| 41 | Trafd1 | 2.84 | 1.15E-86 | 491 | Tnc | 2.36 | 1.25E-13 |
| 42 | Gm47922 | 3.18 | 1.54E-86 | 492 | Alkal2 | 3.65 | 1.54E-13 |
| 43 | Pml | 2.37 | 2.54E-86 | 493 | Gm16094 | 2.10 | 1.58E-13 |
| 44 | Ccl3 | 6.08 | 2.74E-86 | 494 | F830208F22Rik | 2.35 | 1.93E-13 |
| 45 | Slc15a3 | 2.67 | 1.48E-85 | 495 | Gm16758 | 3.60 | 2.08E-13 |
| 46 | Sod2 | 2.62 | 2.67E-85 | 496 | Plet1 | 3.57 | 2.13E-13 |
| 47 | Creb3l1 | 2.86 | 1.89E-84 | 497 | Gm26809 | 2.36 | 2.87E-13 |
| 48 | Errfi1 | 3.41 | 4.31E-84 | 498 | Dthd1 | 2.39 | 3.12E-13 |
| 49 | Trim30a | 3.53 | 8.40E-84 | 499 | Prl7d1 | 3.90 | 3.42E-13 |
| 50 | Tnfaip3 | 3.29 | 1.85E-83 | 500 | Gm34471 | 3.87 | 4.71E-13 |
| 51 | Ifit3b | 5.29 | 2.25E-83 | 501 | A3galt2 | 2.10 | 4.83E-13 |
| 52 | Phf11c | 3.51 | 3.23E-83 | 502 | Gm2511 | 4.06 | 5.00E-13 |
| 53 | Ifi44 | 4.77 | 7.77E-83 | 503 | Spag17os | 2.57 | 5.07E-13 |
| 54 | Slfn5 | 4.26 | 7.77E-83 | 504 | Gm47034 | 2.67 | 5.33E-13 |
| 55 | Ifit1bl2 | 4.06 | 1.53E-82 | 505 | Ankrd66 | 2.43 | 6.91E-13 |
| 56 | Fam129b | 2.32 | 1.29E-81 | 506 | Has1 | 3.97 | 9.42E-13 |
| 57 | Cxcl10 | 6.24 | 5.80E-81 | 507 | Drd5 | 2.92 | 1.02E-12 |
| 58 | Cd14 | 4.26 | 1.82E-80 | 508 | Pnpla1os | 2.61 | 1.02E-12 |
| 59 | Ccl4 | 7.02 | 2.26E-80 | 509 | 1190001M18Rik | 2.74 | 1.85E-12 |
| 60 | Mitd1 | 2.22 | 2.92E-80 | 510 | Irx3 | 2.69 | 2.10E-12 |
| 61 | Slc25a22 | 2.95 | 6.32E-80 | 511 | Gm43069 | 2.85 | 2.64E-12 |
| 62 | Parp10 | 2.80 | 8.43E-79 | 512 | Gm17641 | 2.72 | 2.80E-12 |
| 63 | Hsh2d | 2.53 | 1.03E-78 | 513 | Spp1 | 2.10 | 3.01E-12 |
| 64 | Stat1 | 2.48 | 3.57E-78 | 514 | Trim69 | 2.76 | 3.53E-12 |
| 65 | Nfkbia | 2.49 | 1.12E-77 | 515 | Klk4 | 2.06 | 3.65E-12 |
| 66 | Rnd1 | 4.84 | 4.79E-77 | 516 | Slc6a14 | 6.60 | 3.69E-12 |
| 67 | Tmem184b | 2.16 | 1.00E-76 | 517 | Tmem100 | 2.47 | 4.38E-12 |
| 68 | Gm20186 | 3.17 | 1.08E-76 | 518 | Gm12764 | 2.01 | 5.23E-12 |
| 69 | Ccl7 | 7.46 | 1.85E-76 | 519 | Il1f9 | 3.03 | 5.39E-12 |
| 70 | Gbp3 | 3.84 | 1.86E-76 | 520 | Gm5416 | 4.42 | 5.47E-12 |
| 71 | Il15ra | 2.62 | 3.80E-76 | 521 | F830016B08Rik | 3.35 | 6.17E-12 |
| 72 | Steap4 | 3.67 | 4.22E-76 | 522 | Trpm6 | 2.52 | 7.12E-12 |
| 73 | Edn1 | 3.65 | 1.28E-75 | 523 | Astl | 2.39 | 8.05E-12 |
| 74 | Sdc4 | 3.55 | 4.10E-75 | 524 | Gm13305 | 2.91 | 1.33E-11 |
| 75 | Sp100 | 2.15 | 1.06E-74 | 525 | Gpr84 | 4.27 | 1.51E-11 |
| 76 | Tgtp2 | 3.95 | 2.13E-74 | 526 | Cnksr1 | 2.44 | 1.55E-11 |
| 77 | Trim21 | 2.61 | 7.96E-74 | 527 | Gzmc | 2.46 | 1.65E-11 |
| 78 | Zbp1 | 4.70 | 1.00E-73 | 528 | Dbx2 | 3.02 | 1.78E-11 |
| 79 | Mov10 | 2.93 | 1.04E-73 | 529 | B3gnt7 | 2.16 | 1.84E-11 |
| 80 | Car13 | 3.22 | 1.51E-72 | 530 | Saa2 | 6.05 | 2.45E-11 |
| 81 | Oas2 | 5.14 | 1.68E-72 | 531 | Gm29371 | 2.35 | 3.32E-11 |
| 82 | Rnf213 | 4.28 | 2.09E-71 | 532 | Tsx | 2.50 | 4.16E-11 |
| 83 | Cd274 | 3.43 | 3.01E-70 | 533 | Adam3 | 2.24 | 4.61E-11 |
| 84 | Ifit3 | 5.41 | 4.04E-70 | 534 | U90926 | 3.01 | 4.62E-11 |
| 85 | Acod1 | 7.21 | 7.81E-70 | 535 | Ccn4 | 2.37 | 5.34E-11 |
| 86 | Serpina3n | 2.78 | 3.36E-69 | 536 | Rhov | 3.17 | 5.58E-11 |
| 87 | Zup1 | 2.69 | 9.41E-69 | 537 | Smpdl3b | 2.09 | 7.06E-11 |
| 88 | Ifi206 | 4.65 | 1.37E-67 | 538 | AY761185 | 2.87 | 7.10E-11 |
| 89 | Tubb6 | 2.77 | 1.62E-67 | 539 | Gm44815 | 2.76 | 8.78E-11 |
| 90 | Gm38554 | 3.89 | 2.49E-67 | 540 | Cfap69 | 3.07 | 9.65E-11 |
| 91 | Gpr85 | 4.62 | 2.53E-67 | 541 | Qrfpr | 5.57 | 9.94E-11 |
| 92 | Gm10309 | 5.79 | 2.60E-67 | 542 | Gm16325 | 2.77 | 1.09E-10 |
| 93 | Ifi44l | 3.52 | 4.78E-67 | 543 | Il17f | 2.51 | 1.17E-10 |
| 94 | Ifit2 | 5.12 | 7.48E-67 | 544 | Gm15888 | 4.99 | 1.25E-10 |
| 95 | Csf3 | 7.49 | 9.39E-67 | 545 | CT009580.1 | 2.10 | 1.42E-10 |
| 96 | Phf11b | 3.68 | 2.80E-66 | 546 | Gm42726 | 2.63 | 1.53E-10 |
| 97 | Gbp5 | 4.77 | 5.24E-65 | 547 | Gm43694 | 3.35 | 1.91E-10 |
| 98 | Dusp1 | 3.02 | 5.62E-65 | 548 | Tktl1 | 2.22 | 2.18E-10 |
| 99 | Marcksl1 | 3.29 | 7.79E-65 | 549 | Cdhr5 | 2.02 | 2.19E-10 |
| 100 | Gm20559 | 2.73 | 1.40E-64 | 550 | Ccl20 | 3.91 | 2.30E-10 |
| 101 | Tnf | 3.55 | 1.55E-64 | 551 | Rhou | 2.54 | 2.56E-10 |
| 102 | Msr1 | 3.32 | 8.12E-64 | 552 | Pnp2 | 2.19 | 2.77E-10 |
| 103 | Trim30d | 4.88 | 1.61E-63 | 553 | Gprc5a | 2.06 | 3.44E-10 |
| 104 | Rhob | 2.01 | 2.06E-63 | 554 | Vnn3 | 2.71 | 3.57E-10 |
| 105 | Ms4a4a | 3.37 | 3.21E-63 | 555 | Bean1 | 2.10 | 4.24E-10 |
| 106 | Nampt | 2.84 | 6.12E-63 | 556 | Qrich2 | 2.35 | 4.36E-10 |
| 107 | Timp1 | 4.85 | 8.44E-63 | 557 | Gm17334 | 2.31 | 4.46E-10 |
| 108 | Icam1 | 2.60 | 1.25E-62 | 558 | Gm2065 | 2.01 | 4.46E-10 |
| 109 | Il1b | 3.89 | 3.98E-62 | 559 | Foxj1 | 2.31 | 4.59E-10 |
| 110 | Gbp2b | 4.34 | 1.21E-61 | 560 | Chrm4 | 2.55 | 7.03E-10 |

Table S8

| | | | |
|---|---|---|---|
| 111 | Ccl2 | 6.87 | 3.72E-61 |
| 112 | Irf7 | 3.95 | 4.32E-61 |
| 113 | Mx1 | 5.35 | 4.69E-61 |
| 114 | Ccrl2 | 3.47 | 5.53E-61 |
| 115 | Znfx1 | 2.43 | 2.33E-60 |
| 116 | Ddx58 | 3.04 | 3.78E-60 |
| 117 | Socs1 | 3.97 | 2.01E-59 |
| 118 | 1600014C10Rik | 2.04 | 3.03E-59 |
| 119 | Dusp16 | 2.07 | 4.00E-59 |
| 120 | Serpina3g | 3.75 | 5.72E-59 |
| 121 | Olfr56 | 5.05 | 1.24E-58 |
| 122 | Phf11a | 3.33 | 1.60E-58 |
| 123 | Ifi207 | 2.92 | 1.64E-58 |
| 124 | Tlr2 | 2.06 | 4.16E-58 |
| 125 | Trim34b | 2.68 | 4.38E-58 |
| 126 | Spry4 | 2.07 | 7.19E-58 |
| 127 | P2ry2 | 2.25 | 1.61E-57 |
| 128 | Mab21l3 | 2.98 | 1.98E-57 |
| 129 | Serpina3f | 4.29 | 6.56E-57 |
| 130 | Fgf23 | 5.71 | 8.36E-57 |
| 131 | Tnfrsf9 | 3.32 | 1.62E-56 |
| 132 | Slfn8 | 3.47 | 2.87E-56 |
| 133 | Mlkl | 3.13 | 3.93E-56 |
| 134 | Ccl22 | 2.85 | 3.93E-56 |
| 135 | Tent4a | 2.00 | 9.08E-56 |
| 136 | Gbp2 | 4.09 | 1.56E-55 |
| 137 | Nod2 | 2.07 | 2.06E-55 |
| 138 | Tmem132e | 4.99 | 4.98E-55 |
| 139 | Xaf1 | 3.36 | 1.90E-54 |
| 140 | Fjx1 | 3.27 | 2.15E-54 |
| 141 | Mt2 | 5.06 | 2.41E-54 |
| 142 | Il1a | 3.42 | 4.11E-54 |
| 143 | Il27 | 4.79 | 4.57E-54 |
| 144 | Bcl3 | 2.55 | 6.21E-54 |
| 145 | Nfkbiz | 2.86 | 1.90E-53 |
| 146 | Ldhd | 2.47 | 2.63E-53 |
| 147 | Lgals9 | 2.68 | 2.97E-53 |
| 148 | Gadd45b | 2.96 | 4.79E-53 |
| 149 | Slfn9 | 2.61 | 6.04E-53 |
| 150 | Pkib | 2.71 | 1.24E-52 |
| 151 | Irgm2 | 2.76 | 2.48E-52 |
| 152 | Atf3 | 4.48 | 5.05E-52 |
| 153 | Ifit1bl1 | 4.31 | 3.51E-51 |
| 154 | A330040F15Rik | 3.37 | 4.24E-51 |
| 155 | Samhd1 | 2.31 | 6.00E-51 |
| 156 | Casp4 | 2.89 | 7.84E-51 |
| 157 | Trim30b | 2.64 | 1.91E-50 |
| 158 | Hcar2 | 2.10 | 2.36E-50 |
| 159 | Gm15287 | 4.64 | 4.04E-50 |
| 160 | Ifi211 | 3.37 | 4.63E-50 |
| 161 | Setdb2 | 2.97 | 4.88E-50 |
| 162 | Ddx60 | 3.95 | 5.85E-50 |
| 163 | Ptgs2 | 6.27 | 9.62E-50 |
| 164 | Slfn1 | 2.44 | 9.86E-50 |
| 165 | Lipg | 5.42 | 2.84E-49 |
| 166 | Dtx3l | 3.01 | 1.10E-48 |
| 167 | Sema4c | 2.76 | 2.85E-48 |
| 168 | Gm11626 | 2.69 | 4.89E-48 |
| 169 | Cxcl11 | 6.61 | 5.54E-48 |
| 170 | Serpine1 | 4.05 | 9.80E-48 |
| 171 | Gm42763 | 3.89 | 1.70E-47 |
| 172 | Lgals3bp | 2.08 | 3.00E-47 |
| 173 | 4930512H18Rik | 4.03 | 4.51E-47 |
| 174 | Pfkfb3 | 2.64 | 5.00E-47 |
| 175 | Oas3 | 3.91 | 6.21E-47 |
| 176 | Snx10 | 2.22 | 8.73E-47 |
| 177 | Ms4a4d | 2.29 | 1.72E-46 |
| 178 | Csrnp1 | 2.05 | 1.95E-46 |
| 179 | Nr4a3 | 3.51 | 1.98E-46 |
| 180 | Tap1 | 2.10 | 2.88E-46 |
| 181 | Gzmb | 4.41 | 4.57E-46 |
| 182 | Cmpk2 | 4.35 | 4.57E-46 |
| 183 | Rgs1 | 3.45 | 4.70E-46 |
| 184 | Has2 | 3.64 | 4.80E-46 |
| 185 | Gm43302 | 3.97 | 1.73E-45 |
| 186 | Gbp6 | 4.10 | 1.85E-45 |
| 187 | Myc | 2.12 | 1.85E-45 |
| 188 | Ak4 | 3.11 | 3.70E-45 |
| 189 | Tnfaip6 | 4.06 | 4.50E-45 |
| 190 | Gm47571 | 4.62 | 5.88E-45 |
| 191 | Adamts4 | 6.85 | 7.44E-45 |
| 192 | Gbp7 | 4.00 | 1.03E-44 |
| 193 | Nlrp3 | 2.49 | 1.66E-44 |
| 194 | Pnpt1 | 2.24 | 5.22E-44 |
| 195 | Agrn | 2.07 | 5.59E-44 |
| 196 | Gm42503 | 2.23 | 6.47E-44 |
| 197 | Lif | 3.27 | 9.51E-44 |
| 198 | Fap | 3.02 | 1.04E-43 |
| 199 | Gm32296 | 7.13 | 1.07E-43 |
| 200 | Ms4a4c | 2.66 | 1.30E-43 |
| 201 | Helz2 | 2.93 | 1.45E-43 |
| 202 | Vcan | 4.77 | 1.63E-43 |
| 203 | Uba7 | 2.00 | 1.74E-43 |
| 204 | Tnip3 | 5.93 | 1.85E-43 |
| 205 | Kank4 | 2.72 | 2.52E-43 |
| 206 | Ifi203 | 2.66 | 3.11E-43 |
| 207 | Fos | 2.89 | 6.27E-43 |
| 208 | Fscn1 | 2.58 | 6.27E-43 |
| 209 | AW112010 | 2.44 | 7.56E-43 |
| 210 | Muc1 | 3.16 | 1.26E-42 |
| 211 | Ccl11 | 5.18 | 1.35E-42 |
| 212 | Abtb2 | 2.18 | 1.48E-42 |
| 213 | Cd80 | 2.42 | 2.02E-42 |
| 214 | Mt1 | 3.94 | 3.04E-42 |
| 215 | Tor1aip2 | 2.07 | 3.41E-42 |
| 216 | Iigp1 | 4.37 | 5.70E-42 |
| 217 | Ifi209 | 3.28 | 8.09E-42 |
| 218 | Oas1a | 4.04 | 2.05E-41 |
| 219 | Tmprss2 | 5.15 | 3.52E-41 |
| 220 | Trim30c | 4.94 | 3.93E-41 |
| 221 | Jun | 2.48 | 4.51E-41 |
| 222 | Htr7 | 3.06 | 5.04E-41 |
| 223 | Ifi214 | 3.47 | 5.06E-41 |
| 224 | Hck | 2.01 | 1.05E-40 |
| 561 | Gm48545 | 4.68 | 7.38E-10 |
| 562 | Csf2 | 3.43 | 7.95E-10 |
| 563 | Six3os1 | 2.24 | 1.13E-09 |
| 564 | 4933432I03Rik | 2.32 | 1.15E-09 |
| 565 | Rbm11 | 2.17 | 1.38E-09 |
| 566 | Plat | 2.53 | 1.40E-09 |
| 567 | A630012P03Rik | 5.61 | 1.42E-09 |
| 568 | Pglyrp3 | 4.96 | 1.43E-09 |
| 569 | Cd70 | 3.00 | 1.71E-09 |
| 570 | Vmn2r16 | 2.43 | 2.06E-09 |
| 571 | 2210412B16Rik | 2.51 | 2.08E-09 |
| 572 | Gm27167 | 3.96 | 2.09E-09 |
| 573 | Hc | 2.62 | 2.12E-09 |
| 574 | Erich6 | 2.49 | 2.21E-09 |
| 575 | Olfr76 | 2.29 | 2.70E-09 |
| 576 | 4930435F18Rik | 2.19 | 3.19E-09 |
| 577 | Gm5483 | 3.46 | 3.86E-09 |
| 578 | Gdap10 | 2.96 | 4.23E-09 |
| 579 | Gm49751 | 3.79 | 4.61E-09 |
| 580 | Gm31026 | 2.49 | 5.62E-09 |
| 581 | AC119228.1 | 3.09 | 5.80E-09 |
| 582 | Gm12505 | 2.15 | 6.39E-09 |
| 583 | Gm19299 | 2.07 | 6.44E-09 |
| 584 | Gm15241 | 3.44 | 7.57E-09 |
| 585 | Tarm1 | 2.34 | 8.37E-09 |
| 586 | Hamp | 3.89 | 9.80E-09 |
| 587 | Gm19684 | 2.05 | 1.03E-08 |
| 588 | Ptx3 | 3.94 | 1.10E-08 |
| 589 | Gm15512 | 2.14 | 1.13E-08 |
| 590 | Gm49507 | 2.68 | 1.20E-08 |
| 591 | Sprr1a | 5.98 | 1.21E-08 |
| 592 | Stc1 | 2.08 | 1.26E-08 |
| 593 | Tnp2 | 3.20 | 1.27E-08 |
| 594 | Fosl1 | 3.60 | 1.33E-08 |
| 595 | Gm47735 | 3.16 | 1.33E-08 |
| 596 | Fam83c | 2.52 | 1.50E-08 |
| 597 | Gm28347 | 3.50 | 1.61E-08 |
| 598 | Olfr1395 | 3.62 | 1.97E-08 |
| 599 | Olfr877 | 2.15 | 2.10E-08 |
| 600 | Olfr596 | 2.06 | 2.12E-08 |
| 601 | Gm42519 | 2.21 | 2.46E-08 |
| 602 | Ubd | 2.15 | 2.96E-08 |
| 603 | Gm45542 | 2.07 | 2.96E-08 |
| 604 | Gm29408 | 2.21 | 3.20E-08 |
| 605 | Gm20139 | 2.61 | 3.55E-08 |
| 606 | Gm15701 | 2.30 | 3.61E-08 |
| 607 | Gm16098 | 4.20 | 3.92E-08 |
| 608 | Irx6 | 3.56 | 4.39E-08 |
| 609 | AC124231.3 | 4.81 | 4.60E-08 |
| 610 | Gm12606 | 3.72 | 4.76E-08 |
| 611 | Gm15418 | 2.80 | 4.91E-08 |
| 612 | Olfr362 | 2.79 | 5.02E-08 |
| 613 | Gm45912 | 2.20 | 5.37E-08 |
| 614 | 1700019C18Rik | 2.83 | 5.84E-08 |
| 615 | Hdc | 2.89 | 6.15E-08 |
| 616 | 4933421A08Rik | 2.58 | 6.85E-08 |
| 617 | Cdx4 | 4.50 | 7.11E-08 |
| 618 | Gm15133 | 2.61 | 8.52E-08 |
| 619 | Itgb8 | 2.61 | 8.79E-08 |
| 620 | Fam110c | 3.42 | 9.18E-08 |
| 621 | 4930478E11Rik | 2.03 | 9.24E-08 |
| 622 | Ifna2 | 4.35 | 1.07E-07 |
| 623 | Trem1 | 2.51 | 1.25E-07 |
| 624 | Gm20412 | 2.33 | 1.28E-07 |
| 625 | Gm20705 | 2.84 | 1.36E-07 |
| 626 | Gdf15 | 2.25 | 1.42E-07 |
| 627 | Krt24 | 5.21 | 1.65E-07 |
| 628 | Arsj | 2.11 | 1.78E-07 |
| 629 | Gm44165 | 2.03 | 1.81E-07 |
| 630 | Gm40578 | 2.09 | 1.87E-07 |
| 631 | Gm32089 | 3.84 | 2.17E-07 |
| 632 | Vmn2r11 | 2.37 | 2.17E-07 |
| 633 | AC132253.2 | 2.88 | 2.67E-07 |
| 634 | Olfr318 | 2.10 | 2.86E-07 |
| 635 | Clec4d | 2.15 | 2.97E-07 |
| 636 | Gm29686 | 3.18 | 3.16E-07 |
| 637 | Serpina3b | 3.05 | 3.22E-07 |
| 638 | Adgrg7 | 2.52 | 3.24E-07 |
| 639 | Gm44672 | 2.32 | 3.28E-07 |
| 640 | Gm35028 | 2.61 | 4.10E-07 |
| 641 | Gm47595 | 2.72 | 4.33E-07 |
| 642 | Ereg | 2.49 | 4.97E-07 |
| 643 | Gm37639 | 2.35 | 5.40E-07 |
| 644 | Psg22 | 4.17 | 5.67E-07 |
| 645 | Il23a | 4.62 | 6.83E-07 |
| 646 | Gm17034 | 3.15 | 7.24E-07 |
| 647 | C230029F24Rik | 2.66 | 7.46E-07 |
| 648 | Tnfaip8l3 | 2.92 | 7.46E-07 |
| 649 | Gm12534 | 3.02 | 7.60E-07 |
| 650 | Tcam1 | 2.37 | 8.07E-07 |
| 651 | Gm44542 | 3.68 | 9.39E-07 |
| 652 | Hcn4 | 4.82 | 1.07E-06 |
| 653 | Cdh15 | 2.95 | 1.52E-06 |
| 654 | Gm13067 | 3.13 | 1.59E-06 |
| 655 | Misp | 2.32 | 1.94E-06 |
| 656 | Gm21149 | 3.74 | 1.99E-06 |
| 657 | Gm29461 | 2.14 | 2.20E-06 |
| 658 | Gm43900 | 2.06 | 2.45E-06 |
| 659 | Gm43198 | 3.01 | 2.63E-06 |
| 660 | Mmp20 | 3.50 | 2.70E-06 |
| 661 | Kcns1 | 2.64 | 2.75E-06 |
| 662 | Areg | 3.50 | 4.00E-06 |
| 663 | 4921509O07Rik | 2.45 | 4.04E-06 |
| 664 | Olfr93 | 2.84 | 4.41E-06 |
| 665 | Gm45528 | 2.25 | 5.35E-06 |
| 666 | Gm15283 | 2.04 | 5.56E-06 |
| 667 | Gm43464 | 2.33 | 5.76E-06 |
| 668 | Spata31d1b | 2.43 | 6.01E-06 |
| 669 | Gm42917 | 2.01 | 6.08E-06 |
| 670 | Oacyl | 2.63 | 6.19E-06 |
| 671 | Plagl1 | 2.27 | 6.57E-06 |
| 672 | Platr23 | 2.70 | 6.61E-06 |
| 673 | Gm4065 | 3.01 | 6.85E-06 |
| 674 | Unc93a | 2.27 | 7.00E-06 |

Table S8

| | | | |
|---|---|---|---|
| 225 | Inhbb | 2.90 | 2.19E-40 |
| 226 | Fbxo39 | 3.56 | 2.86E-40 |
| 227 | Upp1 | 3.82 | 5.27E-40 |
| 228 | Il6 | 6.51 | 5.40E-40 |
| 229 | Tnfsf10 | 3.22 | 5.65E-40 |
| 230 | Ly6a | 3.52 | 7.86E-40 |
| 231 | Il15 | 2.72 | 9.92E-40 |
| 232 | Cd69 | 3.84 | 1.12E-39 |
| 233 | Ifi205 | 2.84 | 1.39E-39 |
| 234 | Gm4841 | 3.47 | 1.41E-39 |
| 235 | Cybb | 2.02 | 1.61E-39 |
| 236 | Gm15056 | 4.85 | 3.13E-39 |
| 237 | Ifi208 | 3.13 | 4.98E-39 |
| 238 | Tgtp1 | 3.80 | 5.99E-39 |
| 239 | Ppa1 | 2.72 | 9.05E-39 |
| 240 | Ccl12 | 5.27 | 1.31E-38 |
| 241 | Vdr | 2.74 | 3.13E-38 |
| 242 | Clic4 | 2.47 | 3.79E-38 |
| 243 | Atp10a | 2.82 | 8.32E-38 |
| 244 | Il12b | 3.59 | 9.26E-38 |
| 245 | Gm47243 | 4.39 | 9.78E-38 |
| 246 | Map3k8 | 2.45 | 1.56E-37 |
| 247 | Gm39556 | 2.34 | 1.77E-37 |
| 248 | Cxcl3 | 6.12 | 1.84E-37 |
| 249 | Gm11772 | 3.23 | 1.96E-37 |
| 250 | Cebpb | 2.74 | 2.98E-37 |
| 251 | Cfb | 2.44 | 3.78E-37 |
| 252 | Gm38271 | 2.48 | 9.23E-37 |
| 253 | Jaml | 2.01 | 9.68E-37 |
| 254 | Rhoc | 2.15 | 9.86E-37 |
| 255 | Sp110 | 2.04 | 1.23E-36 |
| 256 | Wfdc17 | 3.73 | 3.24E-36 |
| 257 | Epsti1 | 2.03 | 7.32E-36 |
| 258 | Junb | 2.10 | 8.85E-36 |
| 259 | Gm34045 | 2.49 | 1.38E-35 |
| 260 | Gm45418 | 4.47 | 1.39E-35 |
| 261 | Tcp10b | 5.29 | 2.17E-35 |
| 262 | Asb13 | 2.26 | 2.37E-35 |
| 263 | Jdp2 | 2.09 | 2.55E-35 |
| 264 | Gfra1 | 3.49 | 2.62E-35 |
| 265 | Prss23 | 2.08 | 2.97E-35 |
| 266 | B4galt5 | 2.17 | 3.05E-35 |
| 267 | Gm13074 | 2.95 | 7.79E-35 |
| 268 | Adamts9 | 3.06 | 9.46E-35 |
| 269 | Plscr1 | 2.33 | 1.29E-34 |
| 270 | Slmapos2 | 2.51 | 1.64E-34 |
| 271 | Mndal | 2.34 | 2.34E-34 |
| 272 | Bcl2a1a | 2.72 | 2.36E-34 |
| 273 | Slc2a6 | 2.02 | 2.46E-34 |
| 274 | Slc39a14 | 2.10 | 2.90E-34 |
| 275 | Nfil3 | 2.74 | 3.61E-34 |
| 276 | Tnfaip2 | 2.29 | 4.15E-34 |
| 277 | Asb4 | 3.67 | 7.15E-34 |
| 278 | Sh3pxd2b | 2.10 | 7.89E-34 |
| 279 | Zcchc2 | 2.19 | 1.07E-33 |
| 280 | Gm37975 | 2.58 | 1.36E-33 |
| 281 | Il10 | 3.53 | 1.56E-33 |
| 282 | Slc10a6 | 3.01 | 1.92E-33 |
| 283 | Nupr1 | 2.80 | 2.25E-33 |
| 284 | Dll4 | 2.41 | 2.90E-33 |
| 285 | Fpr1 | 2.59 | 1.72E-32 |
| 286 | Isg15 | 4.37 | 1.88E-32 |
| 287 | Gm12185 | 3.62 | 2.72E-32 |
| 288 | Ifi213 | 4.35 | 4.86E-32 |
| 289 | Acot4 | 2.33 | 6.12E-32 |
| 290 | Apol9a | 3.89 | 6.90E-32 |
| 291 | Gbp4 | 3.46 | 7.76E-32 |
| 292 | Akap2 | 2.63 | 2.55E-31 |
| 293 | Rtn4rl2 | 2.31 | 3.91E-31 |
| 294 | Nmi | 2.44 | 5.15E-31 |
| 295 | Ptprh | 4.95 | 6.84E-31 |
| 296 | Gm45193 | 4.11 | 8.81E-31 |
| 297 | Gm37795 | 2.48 | 1.43E-30 |
| 298 | BC035947 | 3.34 | 1.77E-30 |
| 299 | D430020J02Rik | 2.42 | 2.06E-30 |
| 300 | 4930599N23Rik | 3.06 | 2.12E-30 |
| 301 | AC160336.1 | 2.10 | 2.30E-30 |
| 302 | Lamc2 | 2.35 | 3.15E-30 |
| 303 | Oas1g | 4.06 | 4.24E-30 |
| 304 | Il12rb1 | 2.43 | 4.39E-30 |
| 305 | Gm29094 | 2.70 | 5.77E-30 |
| 306 | Asap3 | 2.07 | 7.23E-30 |
| 307 | Gm16685 | 2.18 | 8.32E-30 |
| 308 | Fosl2 | 2.52 | 9.55E-30 |
| 309 | Cebpd | 2.60 | 9.62E-30 |
| 310 | Mir155hg | 3.05 | 1.05E-29 |
| 311 | Hk2 | 2.43 | 1.15E-29 |
| 312 | Il22 | 8.43 | 1.49E-29 |
| 313 | Saa1 | 6.60 | 1.70E-29 |
| 314 | Pcdh19 | 3.76 | 2.65E-29 |
| 315 | Gm4951 | 3.29 | 3.60E-29 |
| 316 | Gbp10 | 4.67 | 4.02E-29 |
| 317 | Gm5431 | 2.97 | 4.22E-29 |
| 318 | Gm49342 | 2.73 | 7.52E-29 |
| 319 | Bst2 | 2.82 | 7.64E-29 |
| 320 | Gm13889 | 2.96 | 8.13E-29 |
| 321 | Eppk1 | 3.76 | 8.98E-29 |
| 322 | Dusp2 | 2.09 | 9.12E-29 |
| 323 | Batf | 2.33 | 1.64E-28 |
| 324 | Bdkrb1 | 3.17 | 2.30E-28 |
| 325 | Bcl2a1d | 2.34 | 2.47E-28 |
| 326 | Slfn2 | 2.14 | 2.92E-28 |
| 327 | Bcl2a1b | 2.57 | 3.31E-28 |
| 328 | Zfp365 | 2.79 | 5.34E-28 |
| 329 | Gm48942 | 2.45 | 5.81E-28 |
| 330 | Nudt17 | 2.52 | 6.20E-28 |
| 331 | Cxcl9 | 3.57 | 6.45E-28 |
| 332 | Tnfrsf12a | 2.36 | 9.26E-28 |
| 333 | Mmp13 | 7.28 | 1.09E-27 |
| 334 | Ccl5 | 2.97 | 1.18E-27 |
| 335 | 1200007C13Rik | 3.12 | 1.40E-27 |
| 336 | Tnfsf9 | 2.87 | 2.50E-27 |
| 337 | Gm10851 | 2.41 | 2.77E-27 |
| 338 | Bcl2l14 | 3.04 | 2.96E-27 |
| 675 | Gm45717 | 2.21 | 7.18E-06 |
| 676 | Wfdc18 | 2.34 | 7.21E-06 |
| 677 | Foxc2 | 2.32 | 7.37E-06 |
| 678 | D6Ertd474e | 2.77 | 7.39E-06 |
| 679 | Calca | 4.64 | 8.55E-06 |
| 680 | Gm38220 | 2.06 | 8.96E-06 |
| 681 | Gm49442 | 2.61 | 1.00E-05 |
| 682 | Gm21833 | 2.61 | 1.41E-05 |
| 683 | Gm29010 | 2.72 | 1.42E-05 |
| 684 | Il12rb2 | 2.13 | 1.45E-05 |
| 685 | Gm43237 | 3.59 | 1.60E-05 |
| 686 | Gm47822 | 2.51 | 1.77E-05 |
| 687 | Sycp2l | 2.51 | 2.24E-05 |
| 688 | AC157784.1 | 4.21 | 2.38E-05 |
| 689 | Gm37042 | 2.50 | 2.83E-05 |
| 690 | Gfra3 | 2.92 | 2.84E-05 |
| 691 | Prss23os | 2.93 | 2.97E-05 |
| 692 | AA467197 | 2.37 | 2.98E-05 |
| 693 | Spem1 | 2.13 | 3.00E-05 |
| 694 | Nts | 2.28 | 3.14E-05 |
| 695 | Olfr231 | 4.38 | 3.25E-05 |
| 696 | 4930455M05Rik | 2.34 | 3.38E-05 |
| 697 | Spdef | 2.64 | 3.52E-05 |
| 698 | Asb5 | 2.26 | 3.65E-05 |
| 699 | Efcab3 | 3.68 | 3.71E-05 |
| 700 | Gucy2c | 2.07 | 3.73E-05 |
| 701 | Gm12065 | 2.13 | 3.99E-05 |
| 702 | Gm37560 | 2.09 | 4.13E-05 |
| 703 | Oas1e | 2.41 | 5.64E-05 |
| 704 | En2 | 2.02 | 6.22E-05 |
| 705 | 8030451A03Rik | 2.24 | 6.99E-05 |
| 706 | Stfa2 | 3.07 | 7.39E-05 |
| 707 | Prok2 | 3.90 | 7.72E-05 |
| 708 | Gm29064 | 4.79 | 8.92E-05 |
| 709 | Sct | 2.60 | 9.06E-05 |
| 710 | Peak1os | 3.31 | 9.14E-05 |
| 711 | Il19 | 2.78 | 9.89E-05 |
| 712 | Gm37347 | 3.63 | 1.11E-04 |
| 713 | Gm44696 | 2.15 | 1.30E-04 |
| 714 | Prm1 | 2.01 | 1.34E-04 |
| 715 | Gm13481 | 3.40 | 1.48E-04 |
| 716 | C2cd4b | 2.64 | 1.65E-04 |
| 717 | Olfr1167 | 2.09 | 1.77E-04 |
| 718 | Gm15990 | 3.18 | 1.90E-04 |
| 719 | D030024E09Rik | 5.25 | 1.93E-04 |
| 720 | Gm28055 | 2.38 | 2.06E-04 |
| 721 | Hpcal4 | 3.08 | 2.14E-04 |
| 722 | Gm40655 | 2.46 | 2.28E-04 |
| 723 | Gm44867 | 2.19 | 2.56E-04 |
| 724 | Gm8098 | 2.52 | 2.72E-04 |
| 725 | BC026762 | 2.63 | 2.82E-04 |
| 726 | Ifna4 | 4.05 | 2.85E-04 |
| 727 | Orm1 | 2.63 | 3.30E-04 |
| 728 | Lrrtm2 | 2.96 | 3.40E-04 |
| 729 | 9130230L23Rik | 2.15 | 3.45E-04 |
| 730 | Gjb4 | 4.82 | 3.49E-04 |
| 731 | Tex12 | 2.26 | 3.78E-04 |
| 732 | Prdx6b | 2.80 | 3.82E-04 |
| 733 | 1600002D24Rik | 3.54 | 3.94E-04 |
| 734 | Duoxa2 | 3.64 | 4.14E-04 |
| 735 | Adam32 | 2.12 | 4.56E-04 |
| 736 | Ccin | 3.18 | 4.73E-04 |
| 737 | Iqcf1 | 2.28 | 5.13E-04 |
| 738 | Nt5c1a | 3.39 | 5.41E-04 |
| 739 | Meikin | 4.00 | 5.42E-04 |
| 740 | Hp | 2.26 | 5.44E-04 |
| 741 | Chil1 | 2.60 | 5.47E-04 |
| 742 | Gm43884 | 3.55 | 5.89E-04 |
| 743 | Ube2ql1 | 2.31 | 6.26E-04 |
| 744 | Gm42963 | 4.13 | 6.44E-04 |
| 745 | Casr | 2.71 | 7.72E-04 |
| 746 | Gm38316 | 2.41 | 7.79E-04 |
| 747 | Tmem171 | 2.43 | 8.10E-04 |
| 748 | Gm20661 | 2.05 | 8.16E-04 |
| 749 | Slc26a4 | 2.76 | 8.22E-04 |
| 750 | 1700008H02Rik | 2.23 | 8.35E-04 |
| 751 | Il1f6 | 4.90 | 8.38E-04 |
| 752 | Gm31522 | 3.33 | 1.02E-03 |
| 753 | D530033B14Rik | 2.00 | 1.09E-03 |
| 754 | Gm37881 | 2.24 | 1.14E-03 |
| 755 | 1700022H01Rik | 2.40 | 1.15E-03 |
| 756 | Gm2270 | 3.36 | 1.30E-03 |
| 757 | Epgn | 2.10 | 1.40E-03 |
| 758 | Gm5152 | 4.09 | 1.42E-03 |
| 759 | Gm13264 | 2.40 | 1.48E-03 |
| 760 | Mrgpra2a | 2.06 | 1.80E-03 |
| 761 | Gm36107 | 2.74 | 1.95E-03 |
| 762 | Gm47480 | 2.17 | 1.97E-03 |
| 763 | Gm16349 | 2.43 | 2.05E-03 |
| 764 | Gm19744 | 2.40 | 2.15E-03 |
| 765 | Clec2e | 2.36 | 2.15E-03 |
| 766 | A530046M15Rik | 4.24 | 2.20E-03 |
| 767 | AW047730 | 2.05 | 2.23E-03 |
| 768 | E230025N22Rik | 2.31 | 2.37E-03 |
| 769 | Gm38177 | 2.61 | 2.59E-03 |
| 770 | Mrgpra2b | 2.04 | 2.62E-03 |
| 771 | 4930540M05Rik | 2.83 | 2.83E-03 |
| 772 | Wif1 | 4.32 | 2.84E-03 |
| 773 | Lcn2 | 2.32 | 2.87E-03 |
| 774 | Meig1 | 2.48 | 3.09E-03 |
| 775 | Tac1 | 4.24 | 3.16E-03 |
| 776 | A430028G04Rik | 2.04 | 3.22E-03 |
| 777 | Gm47043 | 3.07 | 3.32E-03 |
| 778 | Dmkn | 2.55 | 3.55E-03 |
| 779 | Gm37842 | 3.55 | 3.89E-03 |
| 780 | A230028O05Rik | 2.73 | 4.37E-03 |
| 781 | 2900011O08Rik | 2.09 | 4.39E-03 |
| 782 | Gm1559 | 4.25 | 4.52E-03 |
| 783 | Gm47118 | 3.95 | 4.56E-03 |
| 784 | 4930580E04Rik | 2.45 | 4.62E-03 |
| 785 | Gm20532 | 2.82 | 4.84E-03 |
| 786 | Garem2 | 2.24 | 4.85E-03 |
| 787 | Gm16014 | 3.23 | 4.98E-03 |
| 788 | Sycp3 | 3.07 | 5.06E-03 |

Table S8

| | | | |
|---|---|---|---|
| 339 | Gm43814 | 2.67 | 3.49E-27 |
| 340 | Batf2 | 3.64 | 3.80E-27 |
| 341 | Gm9920 | 2.10 | 4.33E-27 |
| 342 | Ier3 | 2.77 | 5.72E-27 |
| 343 | Ifi35 | 2.64 | 8.08E-27 |
| 344 | Nlrc5 | 2.53 | 1.28E-26 |
| 345 | Nos2 | 2.46 | 1.43E-26 |
| 346 | Trim34a | 2.67 | 1.48E-26 |
| 347 | Irf1 | 2.26 | 1.73E-26 |
| 348 | Cgas | 2.42 | 1.74E-26 |
| 349 | Olfr853 | 2.19 | 1.92E-26 |
| 350 | Dusp5 | 2.19 | 2.15E-26 |
| 351 | Htr1b | 2.37 | 2.44E-26 |
| 352 | Ifitm3 | 2.63 | 2.71E-26 |
| 353 | 1700011B04Rik | 6.31 | 2.74E-26 |
| 354 | Gm34585 | 2.48 | 5.48E-26 |
| 355 | Arid5a | 2.08 | 7.43E-26 |
| 356 | Skor1 | 4.19 | 9.85E-26 |
| 357 | Samd9l | 2.57 | 2.16E-25 |
| 358 | Apol9b | 4.63 | 2.24E-25 |
| 359 | Gm33056 | 3.02 | 2.83E-25 |
| 360 | Gm12840 | 3.02 | 3.01E-25 |
| 361 | Gm36186 | 2.58 | 3.01E-25 |
| 362 | Gm26535 | 3.83 | 3.03E-25 |
| 363 | Tlr3 | 2.25 | 4.34E-25 |
| 364 | Avpr1a | 3.22 | 4.76E-25 |
| 365 | Kcne4 | 2.66 | 4.85E-25 |
| 366 | Tlr7 | 2.22 | 7.86E-25 |
| 367 | Gm47242 | 4.23 | 8.45E-25 |
| 368 | Il33 | 2.33 | 8.51E-25 |
| 369 | Gm19412 | 2.61 | 2.50E-24 |
| 370 | A730049H05Rik | 2.96 | 3.32E-24 |
| 371 | Pik3ap1 | 2.06 | 3.34E-24 |
| 372 | Fcgr4 | 2.60 | 3.82E-24 |
| 373 | Lox | 3.20 | 4.03E-24 |
| 374 | Vgf | 4.82 | 4.15E-24 |
| 375 | Gbp9 | 3.08 | 4.50E-24 |
| 376 | Trex1 | 2.21 | 6.31E-24 |
| 377 | Sult6b1 | 2.56 | 7.46E-24 |
| 378 | Gm43068 | 5.27 | 1.24E-23 |
| 379 | Ehf | 2.22 | 1.81E-23 |
| 380 | Ccdc33 | 3.68 | 3.87E-23 |
| 381 | Ildr1 | 2.08 | 4.15E-23 |
| 382 | Slit1 | 2.89 | 4.45E-23 |
| 383 | Clec4e | 4.15 | 4.96E-23 |
| 384 | Rgs16 | 2.37 | 1.29E-22 |
| 385 | Pnp | 2.53 | 1.39E-22 |
| 386 | Gm6904 | 2.66 | 1.45E-22 |
| 387 | Dgkh | 2.27 | 1.51E-22 |
| 388 | Medag | 2.41 | 1.59E-22 |
| 389 | 4833419F23Rik | 2.12 | 1.89E-22 |
| 390 | Mfsd2a | 2.86 | 2.42E-22 |
| 391 | Fpr2 | 3.70 | 4.59E-22 |
| 392 | Steap1 | 2.94 | 5.18E-22 |
| 393 | Slc26a9 | 5.19 | 5.45E-22 |
| 394 | Ank2 | 2.21 | 7.04E-22 |
| 395 | 1110032F04Rik | 2.16 | 7.13E-22 |
| 396 | Plk2 | 2.52 | 9.52E-22 |
| 397 | Pmepa1 | 2.14 | 9.66E-22 |
| 398 | Cdh6 | 4.38 | 1.14E-21 |
| 399 | Calhm6 | 2.68 | 1.18E-21 |
| 400 | Serpina3c | 2.69 | 1.48E-21 |
| 401 | Ccn1 | 2.63 | 2.00E-21 |
| 402 | Ebi3 | 2.03 | 2.30E-21 |
| 403 | Gpr33 | 3.36 | 2.30E-21 |
| 404 | Chl1 | 2.97 | 2.45E-21 |
| 405 | Gem | 2.33 | 2.69E-21 |
| 406 | Gbp11 | 4.99 | 4.07E-21 |
| 407 | Ccl17 | 3.56 | 4.16E-21 |
| 408 | Cx3cl1 | 2.41 | 4.68E-21 |
| 409 | Lilr4b | 3.07 | 5.34E-21 |
| 410 | Ascc3 | 2.01 | 6.25E-21 |
| 411 | Kif1a | 2.02 | 8.64E-21 |
| 412 | BC147527 | 2.05 | 9.04E-21 |
| 413 | Il1r2 | 3.81 | 9.30E-21 |
| 414 | Oas1h | 3.39 | 1.08E-20 |
| 415 | H2-T24 | 2.46 | 1.21E-20 |
| 416 | Ch25h | 2.37 | 2.31E-20 |
| 417 | Adora2b | 2.78 | 2.82E-20 |
| 418 | Saa3 | 5.66 | 9.14E-20 |
| 419 | Gm13822 | 4.26 | 1.01E-19 |
| 420 | Bmp10 | 3.97 | 1.15E-19 |
| 421 | Gm7598 | 2.18 | 1.38E-19 |
| 422 | Gm4070 | 2.51 | 1.43E-19 |
| 423 | 4930512B01Rik | 2.24 | 1.57E-19 |
| 424 | Gm12253 | 2.60 | 2.55E-19 |
| 425 | Ogfr | 2.08 | 2.86E-19 |
| 426 | Elovl4 | 3.95 | 4.31E-19 |
| 427 | Htra4 | 2.60 | 5.14E-19 |
| 428 | AC167363.2 | 3.39 | 5.65E-19 |
| 429 | Gm16175 | 2.07 | 6.93E-19 |
| 430 | Vash1 | 2.60 | 8.39E-19 |
| 431 | Gm19951 | 3.20 | 1.02E-18 |
| 432 | Ppfia3 | 2.38 | 1.74E-18 |
| 433 | Gm28177 | 2.23 | 4.09E-18 |
| 434 | Gm15856 | 3.25 | 6.05E-18 |
| 435 | Cdh17 | 2.16 | 7.52E-18 |
| 436 | Tnfsf15 | 3.70 | 7.77E-18 |
| 437 | Btbd16 | 2.13 | 7.99E-18 |
| 438 | Crybb3 | 2.46 | 1.33E-17 |
| 439 | Gfpt2 | 2.16 | 1.50E-17 |
| 440 | Gm49339 | 2.03 | 1.57E-17 |
| 441 | Barx2 | 4.06 | 1.58E-17 |
| 442 | Tfpi2 | 3.20 | 3.49E-17 |
| 443 | Gng4 | 3.33 | 4.20E-17 |
| 444 | 1700025N23Rik | 2.52 | 5.50E-17 |
| 445 | A930024N18Rik | 2.06 | 6.83E-17 |
| 446 | Gm48878 | 3.45 | 9.56E-17 |
| 447 | Gm49482 | 5.36 | 1.05E-16 |
| 448 | Gm6377 | 2.09 | 1.45E-16 |
| 449 | Sele | 3.72 | 1.75E-16 |
| 450 | Akap12 | 2.62 | 2.11E-16 |
| 789 | Speer4b | 2.87 | 5.16E-03 |
| 790 | Ifnb1 | 4.67 | 5.58E-03 |
| 791 | 1700066B17Rik | 3.76 | 5.72E-03 |
| 792 | Gm14280 | 3.44 | 5.92E-03 |
| 793 | Gm44448 | 3.68 | 6.12E-03 |
| 794 | Gm3411 | 4.22 | 6.99E-03 |
| 795 | Gm14236 | 2.06 | 7.00E-03 |
| 796 | Gm26679 | 2.61 | 7.05E-03 |
| 797 | 4933409D19Rik | 4.15 | 7.51E-03 |
| 798 | Gm8110 | 2.02 | 7.73E-03 |
| 799 | Reg3g | 2.68 | 7.87E-03 |
| 800 | Gm5796 | 2.12 | 7.93E-03 |
| 801 | 4930478L05Rik | 3.92 | 8.25E-03 |
| 802 | Gpr12 | 2.39 | 8.43E-03 |
| 803 | Fam69c | 3.68 | 8.52E-03 |
| 804 | Acta1 | 3.19 | 8.96E-03 |
| 805 | Gm34322 | 2.34 | 8.97E-03 |
| 806 | Rasef | 2.55 | 9.68E-03 |
| 807 | Gm32540 | 2.27 | 1.03E-02 |
| 808 | Hephl1 | 3.52 | 1.05E-02 |
| 809 | AC165153.1 | 2.39 | 1.11E-02 |
| 810 | Gm43399 | 2.06 | 1.15E-02 |
| 811 | Gm29106 | 2.31 | 1.20E-02 |
| 812 | Gm26603 | 2.11 | 1.23E-02 |
| 813 | Gm11795 | 3.66 | 1.23E-02 |
| 814 | 1810065E05Rik | 2.10 | 1.37E-02 |
| 815 | Gm12480 | 2.27 | 1.43E-02 |
| 816 | Lrrc6 | 2.28 | 1.49E-02 |
| 817 | Fgf6 | 2.63 | 1.50E-02 |
| 818 | Il11ra2 | 2.30 | 1.52E-02 |
| 819 | Krt81 | 4.51 | 1.60E-02 |
| 820 | 2310043L19Rik | 2.16 | 1.62E-02 |
| 821 | Gm15684 | 2.45 | 1.68E-02 |
| 822 | Iltifb | 3.92 | 1.72E-02 |
| 823 | Gm3331 | 2.97 | 1.78E-02 |
| 824 | Gm38258 | 2.70 | 1.93E-02 |
| 825 | Gm48899 | 2.94 | 1.94E-02 |
| 826 | Gm37728 | 3.47 | 1.98E-02 |
| 827 | Chrdl2 | 3.35 | 2.03E-02 |
| 828 | Gm37026 | 2.11 | 2.10E-02 |
| 829 | Defb33 | 2.29 | 2.13E-02 |
| 830 | 4930546C10Rik | 2.01 | 2.14E-02 |
| 831 | 4930503B20Rik | 2.84 | 2.27E-02 |
| 832 | Gm45326 | 2.87 | 2.29E-02 |
| 833 | Gm30676 | 2.61 | 2.31E-02 |
| 834 | Slc9a4 | 2.56 | 2.54E-02 |
| 835 | Gm47214 | 2.70 | 2.59E-02 |
| 836 | Adgrf2 | 2.11 | 2.66E-02 |
| 837 | Eif2c5 | 3.35 | 2.69E-02 |
| 838 | Pom121l2 | 3.46 | 2.77E-02 |
| 839 | Rprml | 2.52 | 2.81E-02 |
| 840 | Gm42450 | 3.00 | 2.99E-02 |
| 841 | Gm15734 | 3.35 | 3.04E-02 |
| 842 | Gm15323 | 3.24 | 3.05E-02 |
| 843 | Klf14 | 2.71 | 3.09E-02 |
| 844 | 9230110F11Rik | 3.75 | 3.10E-02 |
| 845 | Gm15340 | 2.36 | 3.18E-02 |
| 846 | Dnase2b | 2.45 | 3.23E-02 |
| 847 | Mt4 | 3.71 | 3.40E-02 |
| 848 | Gm38377 | 3.62 | 3.49E-02 |
| 849 | Pkp1 | 5.53 | 3.49E-02 |
| 850 | Sox30 | 2.59 | 3.52E-02 |
| 851 | Fkbp6 | 4.96 | 3.68E-02 |
| 852 | Ifnab | 2.21 | 3.79E-02 |
| 853 | Gm33091 | 3.14 | 4.05E-02 |
| 854 | Gm27019 | 3.06 | 4.26E-02 |
| 855 | Gm11722 | 3.29 | 4.30E-02 |
| 856 | Il17a | 3.31 | 4.33E-02 |
| 857 | Gm28626 | 2.16 | 4.45E-02 |
| 858 | Prr30 | 3.13 | 4.45E-02 |
| 859 | Serpinb5 | 3.23 | 4.52E-02 |
| 860 | Gm47657 | 2.28 | 4.52E-02 |
| 861 | Gm49420 | 2.77 | 4.61E-02 |
| 862 | Gm12356 | 2.61 | 4.66E-02 |
| 863 | Ighv2-7 | 2.51 | 4.68E-02 |
| 864 | Olfr94 | 2.14 | 4.69E-02 |
| 865 | Gm16726 | 3.29 | 4.71E-02 |
| 866 | Fam25c | 5.08 | 4.79E-02 |

**Table S9. DEGs of liver of M. musculus**

| rank | gene | log2 fold change | FDR | rank | gene | log2 fold change | FDR | rank | gene | log2 fold change | FDR | rank | gene | log2 fold change | FDR |
|---|---|---|---|---|---|---|---|---|---|---|---|---|---|---|---|
| 1 | Irak3 | 5.85 | 2.34E-178 | 501 | AW011738 | 3.20 | 3.61E-33 | 1001 | 1600010M07Rik | 2.38 | 1.21E-12 | 1501 | Adig | 2.95 | 3.99E-04 |
| 2 | Herc6 | 5.08 | 1.11E-175 | 502 | Ripk2 | 4.15 | 5.09E-33 | 1002 | Olfr16 | 3.00 | 1.27E-12 | 1502 | Gm44843 | 2.05 | 4.15E-04 |
| 3 | Tifa | 8.25 | 4.85E-173 | 503 | Skil | 2.08 | 6.13E-33 | 1003 | Cdk5r1 | 2.41 | 1.34E-12 | 1503 | 4930550C14Rik | 3.06 | 4.15E-04 |
| 4 | Ifit1 | 8.02 | 1.16E-165 | 504 | 4931429L15Rik | 7.49 | 6.54E-33 | 1004 | Spata46 | 3.56 | 1.61E-12 | 1504 | Gm48627 | 2.65 | 4.16E-04 |
| 5 | Ccl2 | 8.18 | 1.29E-161 | 505 | Slc37a1 | 4.19 | 6.66E-33 | 1005 | Gm4951 | 2.08 | 1.62E-12 | 1505 | Calca | 4.86 | 4.17E-04 |
| 6 | Ifit3 | 6.77 | 4.24E-152 | 506 | Il2rg | 2.46 | 7.07E-33 | 1006 | Gm36245 | 4.73 | 1.70E-12 | 1506 | Gm47043 | 3.07 | 4.17E-04 |
| 7 | Sele | 8.31 | 2.52E-151 | 507 | Tmem87b | 2.32 | 7.49E-33 | 1007 | Ece2 | 3.34 | 1.90E-12 | 1507 | Slc13a1 | 2.72 | 4.25E-04 |
| 8 | Gm42763 | 6.16 | 2.52E-151 | 508 | Tle3 | 2.29 | 1.21E-32 | 1008 | 4921523L03Rik | 4.11 | 1.98E-12 | 1508 | Gp1bb | 2.46 | 4.28E-04 |
| 9 | Il1rn | 6.71 | 1.37E-150 | 509 | 0610043K17Rik | 2.63 | 1.26E-32 | 1009 | Dhrs13os | 2.79 | 2.11E-12 | 1509 | 4930517G19Rik | 2.72 | 4.29E-04 |
| 10 | Serpina3f | 8.58 | 3.00E-149 | 510 | Lcp2 | 2.43 | 1.35E-32 | 1010 | Kcne4 | 3.50 | 2.11E-12 | 1510 | Mipepos | 3.60 | 4.45E-04 |
| 11 | Ccl3 | 6.93 | 2.66E-145 | 511 | Gm14319 | 3.12 | 2.52E-32 | 1011 | Pglyrp3 | 6.85 | 2.32E-12 | 1511 | Olfr92 | 3.86 | 4.71E-04 |
| 12 | Xaf1 | 4.47 | 6.60E-143 | 512 | Rasip1 | 3.20 | 2.98E-32 | 1012 | Olfr462 | 3.29 | 2.39E-12 | 1512 | Gm49392 | 3.50 | 4.71E-04 |
| 13 | Gm43302 | 6.39 | 7.59E-141 | 513 | Ly6a | 4.73 | 3.31E-32 | 1013 | Olfr76 | 3.19 | 2.39E-12 | 1513 | 4933432I03Rik | 3.78 | 4.98E-04 |
| 14 | Ifi44 | 5.13 | 5.79E-137 | 514 | Gm20400 | 2.99 | 4.63E-32 | 1014 | Camp | 3.75 | 2.64E-12 | 1514 | Gm45828 | 2.09 | 5.01E-04 |
| 15 | Usp18 | 6.30 | 1.20E-134 | 515 | Jdp2 | 3.01 | 4.77E-32 | 1015 | Fbp2 | 3.15 | 2.89E-12 | 1515 | AC132455.3 | 3.51 | 5.05E-04 |
| 16 | Parp14 | 4.98 | 2.99E-133 | 516 | 4833419F23Rik | 3.94 | 5.32E-32 | 1016 | Gm43197 | 2.30 | 3.14E-12 | 1516 | Gm26706 | 2.47 | 5.06E-04 |
| 17 | Gbp5 | 7.19 | 7.97E-132 | 517 | Il6 | 6.11 | 6.50E-32 | 1017 | Hoxc8 | 2.31 | 3.23E-12 | 1517 | Dgkk | 4.24 | 5.09E-04 |
| 18 | Cd14 | 7.78 | 4.88E-129 | 518 | Ldha | 2.53 | 6.56E-32 | 1018 | Gm45131 | 2.03 | 3.76E-12 | 1518 | Slc28a3 | 2.81 | 5.19E-04 |
| 19 | Ifit3b | 6.55 | 4.88E-129 | 519 | Spsb1 | 2.63 | 6.71E-32 | 1019 | Gm40685 | 3.14 | 4.12E-12 | 1519 | Olfr1395 | 4.11 | 5.40E-04 |
| 20 | Cxcl9 | 8.10 | 6.10E-129 | 520 | D430020J02Rik | 3.46 | 8.96E-32 | 1020 | Gm16098 | 3.44 | 4.16E-12 | 1520 | Prss16 | 3.12 | 5.40E-04 |
| 21 | Marcksl1 | 3.80 | 9.58E-129 | 521 | Ifi44l | 2.99 | 9.10E-32 | 1021 | Gm13598 | 2.80 | 4.42E-12 | 1521 | Chl1 | 4.72 | 5.47E-04 |
| 22 | Fpr2 | 5.27 | 1.05E-127 | 522 | Fam49a | 2.09 | 1.21E-31 | 1022 | Atp10a | 3.21 | 4.68E-12 | 1522 | Gm28651 | 4.41 | 5.54E-04 |
| 23 | Eif2ak2 | 3.78 | 1.14E-127 | 523 | Gm16758 | 5.74 | 1.25E-31 | 1023 | Stk32b | 2.29 | 5.20E-12 | 1523 | 9830107B12Rik | 2.64 | 5.70E-04 |
| 24 | Tgtp2 | 5.56 | 1.05E-126 | 524 | Vars | 2.08 | 1.25E-31 | 1024 | Ly6c1 | 2.51 | 5.74E-12 | 1524 | Gm42917 | 2.24 | 5.74E-04 |
| 25 | Ccrl2 | 8.12 | 2.27E-126 | 525 | Phf11a | 2.57 | 1.46E-31 | 1025 | Ccdc24 | 2.65 | 6.11E-12 | 1525 | Gm48648 | 3.01 | 5.76E-04 |
| 26 | Ccl4 | 7.73 | 4.90E-125 | 526 | Mafk | 2.43 | 1.57E-31 | 1026 | Ccl11 | 5.47 | 6.63E-12 | 1526 | Gm5796 | 5.08 | 5.78E-04 |
| 27 | Tgtp1 | 7.34 | 1.45E-124 | 527 | Spic | 2.62 | 1.70E-31 | 1027 | Gm42566 | 2.54 | 7.26E-12 | 1527 | A730049H05Rik | 2.65 | 5.91E-04 |
| 28 | Icam1 | 4.79 | 1.61E-124 | 528 | Timp1 | 6.12 | 1.91E-31 | 1028 | Fam50b | 3.11 | 7.27E-12 | 1528 | 5830418P13Rik | 3.16 | 5.99E-04 |
| 29 | Mmp13 | 7.89 | 8.85E-121 | 529 | Itpkc | 3.61 | 2.05E-31 | 1029 | Ccl20 | 4.75 | 7.64E-12 | 1529 | Gm43815 | 2.37 | 6.30E-04 |
| 30 | Tap1 | 4.12 | 4.21E-120 | 530 | Tat | 2.48 | 2.16E-31 | 1030 | Gm42963 | 5.27 | 8.35E-12 | 1530 | Timp4 | 2.76 | 6.36E-04 |
| 31 | Slfn9 | 4.79 | 5.99E-119 | 531 | Elf1 | 2.09 | 2.57E-31 | 1031 | Gm17334 | 2.28 | 1.01E-11 | 1531 | Dio2 | 3.46 | 6.55E-04 |
| 32 | Il17ra | 3.93 | 4.53E-117 | 532 | Gm45717 | 3.83 | 2.85E-31 | 1032 | Olfr1156 | 2.07 | 1.06E-11 | 1532 | Hs3st1 | 2.45 | 6.89E-04 |
| 33 | Oasl1 | 6.57 | 1.01E-116 | 533 | Lgals3bp | 2.24 | 3.26E-31 | 1033 | Dpep2nb | 4.68 | 1.14E-11 | 1533 | Gm12703 | 4.02 | 7.00E-04 |
| 34 | Myd88 | 3.47 | 1.15E-114 | 534 | Olr1 | 4.20 | 3.34E-31 | 1034 | Gm41225 | 3.22 | 1.17E-11 | 1534 | Gm13266 | 2.24 | 7.40E-04 |
| 35 | Isg15 | 6.18 | 9.38E-113 | 535 | Gm11626 | 3.12 | 3.65E-31 | 1035 | Gm34152 | 5.67 | 1.21E-11 | 1535 | Krt86 | 3.14 | 7.48E-04 |
| 36 | Rtp4 | 3.66 | 1.27E-112 | 536 | C5ar1 | 2.74 | 4.53E-31 | 1036 | Slc1a1 | 3.35 | 1.37E-11 | 1536 | Gm39307 | 2.35 | 7.62E-04 |
| 37 | Dhx58 | 4.57 | 2.96E-111 | 537 | St5 | 2.38 | 4.97E-31 | 1037 | Gm15845 | 3.18 | 1.48E-11 | 1537 | Gm10532 | 2.49 | 7.75E-04 |
| 38 | Il1b | 5.57 | 5.06E-111 | 538 | Adgrl2 | 2.01 | 5.47E-31 | 1038 | Svopl | 3.32 | 1.48E-11 | 1538 | Erich6 | 2.73 | 7.78E-04 |
| 39 | Gm10309 | 6.21 | 2.43E-110 | 539 | Stx18 | 2.07 | 5.47E-31 | 1039 | Serpinb2 | 4.89 | 1.54E-11 | 1539 | Zcchc12 | 2.57 | 7.80E-04 |
| 40 | Ifi47 | 5.30 | 4.23E-110 | 540 | Dram1 | 3.60 | 5.97E-31 | 1040 | Gm15348 | 3.42 | 1.56E-11 | 1540 | Gm45326 | 2.21 | 7.81E-04 |
| 41 | Serpina3i | 6.63 | 1.78E-109 | 541 | Sema6b | 2.65 | 6.79E-31 | 1041 | Gm32089 | 3.20 | 1.61E-11 | 1541 | Prss34 | 3.18 | 8.35E-04 |
| 42 | Tnip1 | 3.99 | 3.77E-109 | 542 | Slamf1 | 5.45 | 6.89E-31 | 1042 | Fabp12 | 3.23 | 1.64E-11 | 1542 | Gm11738 | 4.45 | 8.45E-04 |
| 43 | Tnfaip3 | 5.64 | 8.36E-109 | 543 | Fgg | 2.59 | 7.39E-31 | 1043 | Gm9967 | 2.11 | 1.67E-11 | 1543 | Cdh9 | 2.95 | 8.65E-04 |
| 44 | Gbp6 | 6.32 | 2.89E-108 | 544 | Pim3 | 2.29 | 7.73E-31 | 1044 | Gm43834 | 5.74 | 1.73E-11 | 1544 | Pglyrp1 | 2.15 | 8.69E-04 |
| 45 | Apol10b | 5.64 | 9.48E-108 | 545 | 4933412E12Rik | 2.99 | 9.22E-31 | 1045 | Aoc2 | 2.03 | 1.75E-11 | 1545 | Gm43585 | 3.16 | 8.86E-04 |
| 46 | Tlr2 | 7.50 | 1.25E-107 | 546 | Snhg5 | 2.55 | 1.08E-30 | 1046 | Smpdl3b | 2.46 | 1.77E-11 | 1546 | 1700041G16Rik | 2.30 | 9.19E-04 |
| 47 | Tgm2 | 3.79 | 4.13E-107 | 547 | Fcgr4 | 2.45 | 1.33E-30 | 1047 | Egr1 | 2.74 | 1.77E-11 | 1547 | Tmprss7 | 2.34 | 9.35E-04 |
| 48 | Zc3h12a | 3.61 | 9.77E-107 | 548 | Apaf1 | 2.38 | 1.44E-30 | 1048 | Trim72 | 3.40 | 1.81E-11 | 1548 | Gm4791 | 2.27 | 9.52E-04 |
| 49 | Parp12 | 3.16 | 6.93E-105 | 549 | Il20rb | 3.81 | 1.44E-30 | 1049 | Mmp12 | 2.55 | 1.90E-11 | 1549 | Dok7 | 2.26 | 9.54E-04 |
| 50 | Irf7 | 4.26 | 7.43E-105 | 550 | Slc4a4 | 2.03 | 1.44E-30 | 1050 | Spib | 2.17 | 1.95E-11 | 1550 | Acta2 | 2.29 | 9.79E-04 |
| 51 | Akap12 | 5.82 | 2.24E-104 | 551 | Rasd1 | 4.58 | 1.70E-30 | 1051 | Gpr35 | 2.17 | 2.18E-11 | 1551 | Tph2 | 3.57 | 1.00E-03 |
| 52 | Acod1 | 9.42 | 2.26E-103 | 552 | Gramd1a | 2.41 | 4.04E-30 | 1052 | Ces1a | 3.25 | 2.22E-11 | 1552 | Pdcd1lg2 | 2.23 | 1.03E-03 |
| 53 | Ccl7 | 8.71 | 4.19E-103 | 553 | Tmem169 | 3.42 | 4.49E-30 | 1053 | Ncf1 | 2.02 | 2.24E-11 | 1553 | AI849053 | 3.30 | 1.04E-03 |
| 54 | Slc15a3 | 3.63 | 5.27E-103 | 554 | Hmox1 | 2.50 | 5.56E-30 | 1054 | Gm9833 | 3.27 | 2.32E-11 | 1554 | Ptx3 | 4.37 | 1.05E-03 |
| 55 | Ccl5 | 6.31 | 1.31E-102 | 555 | Ackr1 | 2.96 | 5.84E-30 | 1055 | Ak7 | 2.38 | 2.37E-11 | 1555 | Tekt4 | 3.26 | 1.07E-03 |
| 56 | Irgm1 | 4.11 | 5.48E-102 | 556 | Gm28347 | 5.34 | 6.08E-30 | 1056 | Arrdc4 | 2.45 | 2.40E-11 | 1556 | Gpr33 | 3.28 | 1.07E-03 |
| 57 | Slfn5 | 3.57 | 1.86E-101 | 557 | Gm44199 | 2.94 | 6.66E-30 | 1057 | AC109138.8 | 2.09 | 2.46E-11 | 1557 | Cngb3 | 4.20 | 1.08E-03 |
| 58 | Dxo | 4.43 | 2.71E-101 | 558 | Gm44658 | 3.32 | 6.67E-30 | 1058 | Sh3bp4 | 2.16 | 2.48E-11 | 1558 | Gm5082 | 4.39 | 1.09E-03 |
| 59 | Zbp1 | 5.32 | 4.73E-100 | 559 | Rhbdf2 | 2.00 | 8.39E-30 | 1059 | Dnah17 | 2.12 | 2.59E-11 | 1559 | Ret | 4.80 | 1.09E-03 |
| 60 | Oas1a | 4.33 | 2.07E-99 | 560 | Gm20172 | 4.53 | 9.36E-30 | 1060 | Sobp | 2.75 | 2.71E-11 | 1560 | Gm29094 | 3.85 | 1.10E-03 |
| 61 | Oasl2 | 5.35 | 1.20E-98 | 561 | Tnfaip2 | 2.12 | 1.04E-29 | 1061 | Ngfr | 2.02 | 2.75E-11 | 1561 | Hs6st2 | 3.05 | 1.12E-03 |
| 62 | Slc11a2 | 3.18 | 2.19E-98 | 562 | Sdsl | 2.43 | 2.39E-29 | 1062 | Bean1 | 6.06 | 2.93E-11 | 1562 | Retn | 3.38 | 1.21E-03 |
| 63 | Arid5a | 6.10 | 2.50E-98 | 563 | Syk | 2.59 | 2.57E-29 | 1063 | A230077H06Rik | 2.04 | 3.52E-11 | 1563 | Gm30198 | 3.29 | 1.22E-03 |
| 64 | Rnf213 | 4.88 | 3.42E-98 | 564 | Tnfrsf9 | 5.70 | 3.10E-29 | 1064 | Gm15344 | 2.36 | 4.11E-11 | 1564 | Muc2 | 2.08 | 1.23E-03 |
| 65 | Adar | 3.65 | 9.34E-98 | 565 | Pprc1 | 2.37 | 3.50E-29 | 1065 | Hrh2 | 3.36 | 4.50E-11 | 1565 | Ccl17 | 3.05 | 1.24E-03 |
| 66 | Ch25h | 8.62 | 2.33E-97 | 566 | Cers6 | 2.49 | 3.66E-29 | 1066 | Chac1 | 2.27 | 4.54E-11 | 1566 | Gm38316 | 2.48 | 1.25E-03 |
| 67 | Cpne8 | 4.54 | 2.41E-97 | 567 | Cdh3 | 3.84 | 3.96E-29 | 1067 | Gm15556 | 5.97 | 4.69E-11 | 1567 | Ppm1n | 2.70 | 1.26E-03 |
| 68 | Trex1 | 4.86 | 4.41E-97 | 568 | Kdm3b | 2.19 | 4.09E-29 | 1068 | Tnfaip6 | 2.59 | 4.71E-11 | 1568 | Prm1 | 4.26 | 1.27E-03 |
| 69 | Isg20 | 5.24 | 2.56E-95 | 569 | Map4k4 | 2.21 | 4.73E-29 | 1069 | Zfp9 | 2.94 | 5.10E-11 | 1569 | Fndc1 | 3.18 | 1.28E-03 |
| 70 | Phf11d | 4.33 | 4.02E-95 | 570 | Tut4 | 2.55 | 4.82E-29 | 1070 | Ceacam16 | 3.89 | 6.08E-11 | 1570 | Msh4 | 3.08 | 1.29E-03 |
| 71 | Cxcl2 | 7.80 | 1.64E-94 | 571 | Ms4a6c | 2.37 | 8.50E-29 | 1071 | Lat2 | 2.36 | 6.22E-11 | 1571 | Grrp1 | 2.17 | 1.31E-03 |
| 72 | Plek | 4.17 | 1.72E-94 | 572 | Gm2511 | 6.67 | 1.14E-28 | 1072 | Gm15133 | 4.33 | 6.31E-11 | 1572 | Vwa2 | 4.37 | 1.33E-03 |
| 73 | Tor3a | 5.20 | 2.02E-94 | 573 | Trib1 | 2.11 | 1.21E-28 | 1073 | Gm26535 | 4.69 | 6.79E-11 | 1573 | Lgsn | 3.03 | 1.33E-03 |
| 74 | Slc2a6 | 5.39 | 4.05E-94 | 574 | Espnl | 3.99 | 1.27E-28 | 1074 | Lrrc63 | 4.82 | 7.26E-11 | 1574 | Gm44241 | 2.25 | 1.34E-03 |
| 75 | Gbp7 | 5.00 | 5.00E-94 | 575 | Gm38271 | 3.66 | 1.31E-28 | 1075 | Olfr695 | 2.56 | 7.28E-11 | 1575 | Gm37222 | 3.57 | 1.36E-03 |
| 76 | Mxd1 | 3.87 | 7.69E-94 | 576 | Fkbp5 | 2.04 | 1.40E-28 | 1076 | Slfn3 | 2.10 | 7.64E-11 | 1576 | Arsi | 3.04 | 1.40E-03 |
| 77 | Gbp9 | 5.14 | 1.30E-93 | 577 | Itgb3 | 2.17 | 1.41E-28 | 1077 | Ypel4 | 2.63 | 8.03E-11 | 1577 | AC155941.1 | 2.59 | 1.40E-03 |
| 78 | Tacc2 | 4.57 | 2.79E-93 | 578 | Bud13 | 4.24 | 2.00E-28 | 1078 | Syn2 | 3.62 | 8.66E-11 | 1578 | 2610318N02Rik | 3.11 | 1.44E-03 |
| 79 | Sebox | 6.25 | 2.08E-92 | 579 | Gadd45g | 5.15 | 2.01E-28 | 1079 | Fst | 5.06 | 8.73E-11 | 1579 | Gm15347 | 3.26 | 1.45E-03 |
| 80 | Akap2 | 3.55 | 2.58E-92 | 580 | Trim29 | 7.10 | 2.27E-28 | 1080 | Schip1 | 2.18 | 9.10E-11 | 1580 | Gm5416 | 3.07 | 1.47E-03 |
| 81 | Mlkl | 3.80 | 2.68E-92 | 581 | Pgam2 | 3.28 | 2.37E-28 | 1081 | Gm42864 | 2.52 | 9.19E-11 | 1581 | Lrrc4 | 2.02 | 1.52E-03 |
| 82 | Serpina3n | 4.18 | 2.74E-92 | 582 | Fgb | 2.42 | 2.45E-28 | 1082 | Gm45259 | 2.08 | 9.97E-11 | 1582 | Hist1h4a | 2.38 | 1.64E-03 |
| 83 | Fancc | 4.39 | 1.03E-90 | 583 | Nav2 | 2.18 | 2.48E-28 | 1083 | Trpc2 | 6.03 | 1.06E-10 | 1583 | Hrh1 | 2.43 | 1.66E-03 |
| 84 | Lbp | 4.14 | 1.57E-90 | 584 | Stx11 | 2.60 | 2.51E-28 | 1084 | Xntrpc | 2.04 | 1.08E-10 | 1584 | Alox5 | 2.11 | 1.68E-03 |
| 85 | Znfx1 | 4.05 | 3.83E-90 | 585 | Arg2 | 2.58 | 3.86E-28 | 1085 | A530032D15Rik | 3.26 | 1.14E-10 | 1585 | Slc36a2 | 5.69 | 1.69E-03 |
| 86 | Pml | 3.15 | 5.04E-90 | 586 | Cd300lf | 3.24 | 4.70E-28 | 1086 | Gm17193 | 2.71 | 1.17E-10 | 1586 | Wfdc18 | 3.56 | 1.71E-03 |
| 87 | H3f3b | 3.38 | 8.04E-90 | 587 | Rnd1 | 4.85 | 5.23E-28 | 1087 | Grid2 | 4.50 | 1.31E-10 | 1587 | Olfr1406 | 2.27 | 1.73E-03 |
| 88 | Socs1 | 7.72 | 4.34E-89 | 588 | Cebpb | 2.37 | 5.48E-28 | 1088 | Hist1h2bh | 2.43 | 1.32E-10 | 1588 | Thbs4 | 3.12 | 1.73E-03 |
| 89 | Bcl2a1a | 5.90 | 8.84E-89 | 589 | Gbp8 | 2.86 | 5.49E-28 | 1089 | Vxn | 3.84 | 1.34E-10 | 1589 | Cplx3 | 2.81 | 1.74E-03 |
| 90 | Jun | 3.78 | 1.33E-88 | 590 | Ggt6 | 2.43 | 6.30E-28 | 1090 | Vmn1r203 | 2.51 | 1.44E-10 | 1590 | Gm45632 | 2.17 | 1.81E-03 |
| 91 | Ptpn1 | 3.28 | 7.89E-88 | 591 | Cxcl11 | 5.73 | 7.99E-28 | 1091 | Gm38070 | 2.26 | 1.46E-10 | 1591 | Gm14820 | 2.46 | 1.84E-03 |
| 92 | Ddx60 | 5.28 | 9.42E-88 | 592 | Gm15298 | 2.59 | 8.55E-28 | 1092 | Gdf15 | 3.25 | 1.58E-10 | 1592 | Ampd1 | 3.32 | 1.86E-03 |
| 93 | Dusp2 | 6.21 | 3.34E-87 | 593 | BB123696 | 2.94 | 8.55E-28 | 1093 | Gm37834 | 4.18 | 1.68E-10 | 1593 | Gm13070 | 3.06 | 1.88E-03 |
| 94 | Slfn8 | 4.75 | 1.73E-86 | 594 | Gm44873 | 2.42 | 9.60E-28 | 1094 | Ptgir | 3.25 | 1.69E-10 | 1594 | Gm17224 | 2.63 | 1.91E-03 |
| 95 | Skiv2l | 3.68 | 1.18E-85 | 595 | H2-T24 | 2.38 | 1.04E-27 | 1095 | Gm49730 | 2.84 | 1.77E-10 | 1595 | Scrg1 | 4.24 | 1.98E-03 |
| 96 | Hcar2 | 7.84 | 1.31E-85 | 596 | Apobec1 | 2.86 | 1.13E-27 | 1096 | Ccdc180 | 3.94 | 1.81E-10 | 1596 | 7420701I03Rik | 2.64 | 2.06E-03 |
| 97 | Msh2 | 3.70 | 1.50E-85 | 597 | Ccr1 | 3.65 | 1.27E-27 | 1097 | Ankrd22 | 3.57 | 2.02E-10 | 1597 | Spem1 | 3.13 | 2.09E-03 |
| 98 | Sowahc | 3.38 | 4.65E-85 | 598 | Sdc4 | 2.18 | 1.39E-27 | 1098 | Stfa2 | 6.50 | 2.10E-10 | 1598 | Olfr1290 | 2.02 | 2.20E-03 |
| 99 | Phf11b | 4.62 | 1.07E-84 | 599 | Gm21859 | 6.05 | 2.25E-27 | 1099 | Gm34585 | 2.14 | 2.20E-10 | 1599 | Zbed6 | 7.06 | 2.22E-03 |
| 100 | Cebpd | 3.79 | 1.11E-84 | 600 | Tcirg1 | 2.28 | 2.26E-27 | 1100 | Cd200r2 | 2.42 | 2.24E-10 | 1600 | Gm48917 | 4.30 | 2.27E-03 |
| 101 | Chil1 | 7.89 | 3.24E-84 | 601 | Gm10851 | 6.04 | 2.49E-27 | 1101 | Speer4e | 4.90 | 2.49E-10 | 1601 | Gal | 4.21 | 2.28E-03 |
| 102 | Olfr56 | 5.31 | 1.44E-83 | 602 | Ctrl | 4.37 | 2.77E-27 | 1102 | Gm9989 | 2.80 | 2.61E-10 | 1602 | Gm21854 | 4.77 | 2.29E-03 |
| 103 | Stk19 | 5.48 | 3.56E-83 | 603 | Slc41a2 | 5.54 | 2.79E-27 | 1103 | Prf1 | 3.04 | 2.66E-10 | 1603 | A630012P03Rik | 3.14 | 2.29E-03 |
| 104 | Trim30a | 4.34 | 1.76E-82 | 604 | Tnip2 | 2.57 | 2.91E-27 | 1104 | Creld2 | 2.14 | 2.68E-10 | 1604 | Oas1e | 2.53 | 2.41E-03 |
| 105 | Atp6v0a4 | 6.72 | 5.61E-82 | 605 | Mtfr2 | 5.00 | 2.91E-27 | 1105 | Gm16028 | 2.70 | 2.76E-10 | 1605 | Zfp365 | 2.10 | 2.43E-03 |
| 106 | H2-Q4 | 3.22 | 9.89E-82 | 606 | Gm32296 | 6.94 | 3.58E-27 | 1106 | Speer4d | 4.64 | 3.43E-10 | 1606 | Gm12027 | 2.04 | 2.43E-03 |
| 107 | Rsad2 | 8.09 | 7.16E-81 | 607 | Gja1 | 2.03 | 4.59E-27 | 1107 | Ptges | 2.51 | 3.44E-10 | 1607 | Madcam1 | 2.50 | 2.44E-03 |
| 108 | Iigp1 | 3.70 | 7.35E-81 | 608 | Psd | 2.84 | 5.48E-27 | 1108 | Nav3 | 4.40 | 3.51E-10 | 1608 | Fam177a | 2.43 | 2.47E-03 |
| 109 | Gm12840 | 3.43 | 8.70E-80 | 609 | Wfdc17 | 2.09 | 6.13E-27 | 1109 | Mthfd2 | 3.22 | 3.87E-10 | 1609 | Gm34643 | 2.06 | 2.53E-03 |
| 110 | Mgarp | 9.63 | 1.40E-79 | 610 | Tmem173 | 2.39 | 6.20E-27 | 1110 | Itk | 2.07 | 3.93E-10 | 1610 | Gm47863 | 2.48 | 2.54E-03 |
| 111 | Daxx | 3.85 | 1.66E-79 | 611 | Rnf122 | 4.21 | 6.30E-27 | 1111 | Gadd45a | 2.07 | 3.97E-10 | 1611 | 3110009F21Rik | 3.11 | 2.57E-03 |
| 112 | Igtp | 3.94 | 5.48E-79 | 612 | 1810063I02Rik | 4.52 | 9.45E-27 | 1112 | Ubd | 5.18 | 4.03E-10 | 1612 | Gm11924 | 2.81 | 2.61E-03 |
| 113 | Apol6 | 6.39 | 7.04E-79 | 613 | Rgs1 | 2.91 | 9.87E-27 | 1113 | Sirpb1b | 5.59 | 4.98E-10 | 1613 | Gm38278 | 2.93 | 2.63E-03 |
| 114 | Ifit1bl2 | 6.52 | 8.23E-79 | 614 | Pcgf5 | 2.04 | 1.44E-26 | 1114 | Cd3e | 3.05 | 5.08E-10 | 1614 | Gm38094 | 5.12 | 2.66E-03 |
| 115 | Ifit2 | 7.42 | 2.89E-78 | 615 | Ikzf1 | 2.34 | 1.61E-26 | 1115 | Gm20705 | 3.65 | 5.50E-10 | 1615 | Gm36745 | 3.00 | 2.69E-03 |
| 116 | Clp1 | 3.13 | 3.11E-78 | 616 | Gm36186 | 3.50 | 1.95E-26 | 1116 | Dnd1 | 2.05 | 5.91E-10 | 1616 | Slc16a8 | 2.47 | 2.69E-03 |
| 117 | Syt12 | 5.43 | 3.61E-78 | 617 | Fap | 4.73 | 2.02E-26 | 1117 | Synpo2 | 2.22 | 6.25E-10 | 1617 | Il12a | 2.22 | 2.70E-03 |
| 118 | Tiparp | 4.35 | 4.11E-78 | 618 | Adm | 3.89 | 2.08E-26 | 1118 | B4galnt4 | 2.52 | 6.43E-10 | 1618 | Gm7030 | 2.18 | 2.71E-03 |
| 119 | Ets2 | 3.35 | 4.65E-78 | 619 | S100a9 | 5.15 | 2.30E-26 | 1119 | AC163629.2 | 2.99 | 6.64E-10 | 1619 | Ccl28 | 2.12 | 2.83E-03 |
| 120 | Cldn14 | 4.66 | 1.00E-77 | 620 | Mt1 | 4.44 | 2.67E-26 | 1120 | Rnf186 | 2.03 | 6.91E-10 | 1620 | Lrrc75aos1 | 3.76 | 3.07E-03 |
| 121 | Csf3 | 8.87 | 1.89E-76 | 621 | Inpp1 | 2.01 | 2.78E-26 | 1121 | 3930402G23Rik | 3.28 | 6.92E-10 | 1621 | Tm4sf20 | 4.57 | 3.08E-03 |
| 122 | Fam83a | 5.20 | 2.34E-76 | 622 | Ifitm1 | 2.04 | 3.54E-26 | 1122 | Trpm8 | 4.06 | 6.99E-10 | 1622 | Syt14 | 2.36 | 3.33E-03 |
| 123 | Ptp4a1 | 3.14 | 2.34E-76 | 623 | Mrgpra2b | 6.53 | 3.73E-26 | 1123 | Nts | 4.54 | 7.42E-10 | 1623 | D930020B18Rik | 2.79 | 3.34E-03 |

Table S9

| | | | |
|---|---|---|---|
| 124 | Gm4841 | 5.67 | 7.80E-76 |
| 125 | Ctf1 | 4.29 | 9.51E-76 |
| 126 | Slc10a6 | 6.98 | 1.11E-75 |
| 127 | Gbp2b | 7.54 | 2.37E-75 |
| 128 | Parp10 | 3.06 | 3.36E-75 |
| 129 | Xdh | 3.68 | 3.92E-75 |
| 130 | Srgn | 3.11 | 1.21E-74 |
| 131 | Bcl3 | 3.57 | 1.57E-74 |
| 132 | Ddx58 | 2.99 | 3.19E-74 |
| 133 | Ifi202b | 3.68 | 1.34E-73 |
| 134 | Ptpn2 | 2.34 | 2.76E-73 |
| 135 | Mitd1 | 2.63 | 3.12E-73 |
| 136 | Parp9 | 2.71 | 3.80E-73 |
| 137 | Heatr1 | 2.92 | 4.97E-73 |
| 138 | Hk2 | 4.08 | 5.57E-73 |
| 139 | Plaur | 4.56 | 7.47E-73 |
| 140 | Steap4 | 6.32 | 1.33E-72 |
| 141 | Snx10 | 3.26 | 1.51E-72 |
| 142 | Batf | 3.98 | 3.96E-72 |
| 143 | Thbs1 | 5.77 | 8.05E-72 |
| 144 | Sgk1 | 3.36 | 9.39E-72 |
| 145 | Gm43062 | 3.94 | 9.54E-72 |
| 146 | Mefv | 5.95 | 1.46E-71 |
| 147 | Olfr1312 | 4.16 | 1.86E-71 |
| 148 | Fas | 2.81 | 1.92E-71 |
| 149 | Upp1 | 5.72 | 2.16E-71 |
| 150 | Jmjd1c | 4.26 | 2.31E-71 |
| 151 | Gbp4 | 4.23 | 4.57E-71 |
| 152 | Dusp8 | 5.52 | 6.97E-71 |
| 153 | Cxcl10 | 8.90 | 7.94E-71 |
| 154 | Ifih1 | 3.32 | 1.13E-70 |
| 155 | Trafd1 | 2.87 | 1.82E-70 |
| 156 | Mx1 | 7.79 | 2.81E-70 |
| 157 | Rnf31 | 2.85 | 5.16E-70 |
| 158 | Pik3ap1 | 3.09 | 3.28E-69 |
| 159 | Etv6 | 3.26 | 3.55E-69 |
| 160 | Nos2 | 9.85 | 5.74E-69 |
| 161 | Atf3 | 5.84 | 1.05E-68 |
| 162 | Ikbke | 4.51 | 1.10E-68 |
| 163 | Gm20186 | 2.98 | 1.34E-68 |
| 164 | Ifi205 | 5.51 | 1.58E-68 |
| 165 | Slfn2 | 3.89 | 2.31E-68 |
| 166 | Pnp | 2.76 | 4.49E-68 |
| 167 | Gm20546 | 5.55 | 5.52E-68 |
| 168 | Calhm6 | 7.17 | 7.37E-68 |
| 169 | Serpina3g | 5.70 | 8.16E-68 |
| 170 | Trim56 | 2.85 | 1.68E-67 |
| 171 | Serpine1 | 7.48 | 2.44E-67 |
| 172 | Nfkb2 | 2.99 | 1.53E-66 |
| 173 | Gbp2 | 6.45 | 1.02E-65 |
| 174 | Rab8b | 2.63 | 1.06E-65 |
| 175 | G3bp2 | 2.69 | 1.20E-65 |
| 176 | Susd6 | 3.53 | 1.37E-65 |
| 177 | Tap2 | 2.51 | 2.22E-65 |
| 178 | AW112010 | 3.03 | 4.51E-65 |
| 179 | 1200007C13Rik | 6.42 | 7.18E-65 |
| 180 | Sgms1 | 3.61 | 7.36E-65 |
| 181 | Dennd4a | 3.41 | 8.14E-65 |
| 182 | Gm47922 | 2.49 | 1.67E-64 |
| 183 | Cmpk2 | 7.47 | 1.74E-64 |
| 184 | Adamts4 | 6.35 | 2.73E-64 |
| 185 | Csrnp1 | 3.55 | 4.89E-64 |
| 186 | Osmr | 4.17 | 9.54E-64 |
| 187 | Gbp3 | 6.04 | 1.14E-63 |
| 188 | Tnf | 5.96 | 1.78E-63 |
| 189 | Arel1 | 2.76 | 7.32E-63 |
| 190 | B4galt1 | 2.81 | 7.69E-63 |
| 191 | Sp140 | 2.76 | 1.48E-62 |
| 192 | Cd274 | 5.15 | 2.04E-62 |
| 193 | Eif1a | 2.71 | 2.21E-62 |
| 194 | Lcn2 | 8.20 | 8.21E-62 |
| 195 | Tor1aip1 | 2.83 | 9.49E-62 |
| 196 | Gcnt2 | 3.38 | 1.08E-61 |
| 197 | Gm29371 | 4.72 | 1.50E-61 |
| 198 | Lrrc8a | 2.76 | 2.02E-61 |
| 199 | H2-Q6 | 2.84 | 3.56E-61 |
| 200 | Gm17300 | 3.35 | 3.78E-61 |
| 201 | Zc3h13 | 5.22 | 4.09E-61 |
| 202 | Gbp10 | 5.21 | 5.71E-61 |
| 203 | Pik3r5 | 4.12 | 9.97E-61 |
| 204 | Oas1b | 3.87 | 1.18E-60 |
| 205 | Casp4 | 3.94 | 1.41E-60 |
| 206 | Fosl2 | 2.76 | 1.62E-60 |
| 207 | Psmb8 | 2.51 | 1.75E-60 |
| 208 | Tbk1 | 2.35 | 2.04E-60 |
| 209 | Slfn1 | 4.40 | 3.02E-60 |
| 210 | Car13 | 4.74 | 4.09E-60 |
| 211 | Litaf | 3.20 | 4.45E-60 |
| 212 | Ifi204 | 4.07 | 5.46E-60 |
| 213 | Trim6 | 4.54 | 5.54E-60 |
| 214 | Gm32815 | 6.10 | 5.82E-60 |
| 215 | Acvr1b | 2.44 | 5.86E-60 |
| 216 | Fmnl2 | 2.46 | 7.20E-60 |
| 217 | Mx2 | 5.18 | 2.04E-59 |
| 218 | Igfbp1 | 6.72 | 2.59E-59 |
| 219 | Nfkbie | 3.50 | 2.73E-59 |
| 220 | Irgm2 | 3.96 | 3.71E-59 |
| 221 | Plac8 | 2.68 | 4.92E-59 |
| 222 | Birc3 | 3.48 | 2.19E-58 |
| 223 | Gm43814 | 5.29 | 2.24E-58 |
| 224 | Gadd45b | 5.34 | 2.49E-58 |
| 225 | Mvp | 2.44 | 3.86E-58 |
| 226 | Vcam1 | 3.11 | 5.12E-58 |
| 227 | Gm47571 | 3.74 | 8.16E-58 |
| 228 | Trim12c | 2.35 | 8.63E-58 |
| 229 | Ms4a6d | 3.69 | 1.41E-57 |
| 230 | Sntb2 | 2.14 | 1.58E-57 |
| 231 | Slc13a5 | 6.02 | 1.64E-57 |
| 232 | Tmem178b | 5.09 | 3.09E-57 |
| 233 | Cd44 | 2.96 | 3.36E-57 |
| 234 | Ifi35 | 2.51 | 4.15E-57 |
| 235 | Col4a1 | 3.06 | 5.69E-57 |
| 236 | Relb | 4.08 | 1.53E-56 |
| 237 | Mctp2 | 2.65 | 2.59E-56 |
| 238 | Inhbb | 5.85 | 2.96E-56 |
| 239 | Tlk2 | 2.16 | 3.19E-56 |
| 240 | Csf1 | 2.86 | 4.77E-56 |
| 241 | Socs3 | 5.52 | 5.69E-56 |
| 242 | Parp11 | 3.19 | 6.34E-56 |
| 243 | H2-Q7 | 2.90 | 6.64E-56 |
| 244 | Junb | 3.79 | 6.82E-56 |
| 245 | Nfkbia | 2.83 | 1.04E-55 |
| 246 | Samhd1 | 3.67 | 1.50E-55 |
| 247 | Tead1 | 4.26 | 1.82E-55 |
| 248 | Il1r2 | 5.99 | 2.56E-55 |
| 249 | Creb3l1 | 3.78 | 2.67E-55 |
| 624 | Gm44641 | 4.21 | 4.05E-26 |
| 625 | Il33 | 3.66 | 4.06E-26 |
| 626 | AC166172.1 | 3.14 | 6.22E-26 |
| 627 | Retreg1 | 2.18 | 6.56E-26 |
| 628 | Zup1 | 2.48 | 7.96E-26 |
| 629 | Traf1 | 3.26 | 7.99E-26 |
| 630 | Rhob | 2.74 | 8.36E-26 |
| 631 | Nos1ap | 3.10 | 8.80E-26 |
| 632 | Serpina10 | 2.15 | 9.15E-26 |
| 633 | Tent5a | 2.03 | 9.61E-26 |
| 634 | Arl15 | 2.10 | 9.94E-26 |
| 635 | Pnp2 | 2.66 | 1.37E-25 |
| 636 | Gm45193 | 4.71 | 1.46E-25 |
| 637 | Myo16 | 6.75 | 1.47E-25 |
| 638 | Gm8883 | 2.06 | 1.47E-25 |
| 639 | AC118695.1 | 3.25 | 1.48E-25 |
| 640 | S1pr3 | 2.10 | 1.63E-25 |
| 641 | Gm2379 | 4.35 | 1.91E-25 |
| 642 | Gm10631 | 2.88 | 2.29E-25 |
| 643 | C1qtnf9 | 2.92 | 2.62E-25 |
| 644 | Crp | 2.51 | 4.40E-25 |
| 645 | Gm44950 | 2.05 | 5.11E-25 |
| 646 | Artn | 6.04 | 5.12E-25 |
| 647 | Gm32592 | 2.60 | 5.45E-25 |
| 648 | Sla | 2.49 | 5.49E-25 |
| 649 | Cxcl3 | 8.75 | 5.96E-25 |
| 650 | Gm32856 | 2.48 | 7.15E-25 |
| 651 | Trim12a | 3.43 | 7.19E-25 |
| 652 | 9930111J21Rik2 | 2.52 | 7.47E-25 |
| 653 | Rab20 | 2.22 | 7.81E-25 |
| 654 | Gm11827 | 3.82 | 7.86E-25 |
| 655 | Ptafr | 2.87 | 9.24E-25 |
| 656 | A2m | 7.30 | 9.31E-25 |
| 657 | Ssx2ip | 2.18 | 9.45E-25 |
| 658 | 9930111J21Rik1 | 4.13 | 1.12E-24 |
| 659 | Dpp9 | 2.80 | 1.25E-24 |
| 660 | Slc16a1 | 2.40 | 1.26E-24 |
| 661 | Pgs1 | 2.43 | 1.29E-24 |
| 662 | AC102496.1 | 2.59 | 1.35E-24 |
| 663 | AU020206 | 2.17 | 2.15E-24 |
| 664 | Gm37305 | 2.02 | 2.26E-24 |
| 665 | Ifi214 | 3.58 | 2.52E-24 |
| 666 | Srrm4 | 2.69 | 2.52E-24 |
| 667 | Tmem132e | 2.21 | 3.36E-24 |
| 668 | Tubb6 | 2.71 | 3.83E-24 |
| 669 | Mir155hg | 4.45 | 5.02E-24 |
| 670 | Saa2 | 7.19 | 5.33E-24 |
| 671 | Fga | 2.67 | 7.08E-24 |
| 672 | Gm29685 | 8.89 | 7.42E-24 |
| 673 | Uck2 | 2.03 | 9.51E-24 |
| 674 | Slc12a7 | 2.46 | 1.10E-23 |
| 675 | Numbl | 5.67 | 1.19E-23 |
| 676 | Adcy1 | 6.42 | 1.44E-23 |
| 677 | Mmp8 | 5.65 | 1.50E-23 |
| 678 | Nnmt | 2.85 | 1.51E-23 |
| 679 | Tnfsf9 | 5.17 | 1.77E-23 |
| 680 | Gm13074 | 3.66 | 2.29E-23 |
| 681 | Hivep2 | 2.14 | 2.87E-23 |
| 682 | 9630002D21Rik | 4.34 | 2.91E-23 |
| 683 | Fcer1a | 3.71 | 2.92E-23 |
| 684 | Hpx | 2.31 | 3.07E-23 |
| 685 | Asb4 | 3.53 | 3.21E-23 |
| 686 | Ctps | 2.02 | 3.44E-23 |
| 687 | Gm13889 | 2.21 | 3.66E-23 |
| 688 | Procr | 3.02 | 3.88E-23 |
| 689 | Klhl30 | 8.44 | 3.99E-23 |
| 690 | Orm1 | 2.36 | 4.62E-23 |
| 691 | 4933405L10Rik | 4.80 | 4.87E-23 |
| 692 | Olfr509 | 2.35 | 5.02E-23 |
| 693 | Ptgs2 | 6.38 | 5.39E-23 |
| 694 | Golt1b | 2.37 | 6.17E-23 |
| 695 | Btbd19 | 2.66 | 6.80E-23 |
| 696 | Csf2rb | 2.15 | 7.13E-23 |
| 697 | Irx2 | 6.83 | 7.35E-23 |
| 698 | Gm42503 | 3.10 | 8.93E-23 |
| 699 | Mt2 | 5.04 | 8.98E-23 |
| 700 | Gm43622 | 2.23 | 9.28E-23 |
| 701 | Cep170 | 2.16 | 9.71E-23 |
| 702 | P2ry2 | 2.44 | 1.01E-22 |
| 703 | Cd177 | 5.31 | 1.08E-22 |
| 704 | Cd93 | 2.30 | 1.37E-22 |
| 705 | Gm29408 | 2.70 | 1.50E-22 |
| 706 | Gem | 2.73 | 2.06E-22 |
| 707 | Bcl6b | 2.58 | 2.76E-22 |
| 708 | Btg3 | 3.52 | 3.15E-22 |
| 709 | Gm13822 | 4.08 | 3.19E-22 |
| 710 | Slc7a11 | 5.11 | 3.24E-22 |
| 711 | Apbb3 | 2.45 | 3.67E-22 |
| 712 | Trim67 | 2.84 | 3.74E-22 |
| 713 | Psme2b | 4.16 | 5.10E-22 |
| 714 | Icosl | 2.41 | 5.17E-22 |
| 715 | BC030867 | 3.63 | 6.01E-22 |
| 716 | B3galt1 | 3.33 | 6.50E-22 |
| 717 | Ltb | 2.24 | 1.18E-21 |
| 718 | Il12b | 5.88 | 1.28E-21 |
| 719 | Slc6a6 | 2.30 | 1.28E-21 |
| 720 | Cd200 | 3.01 | 1.30E-21 |
| 721 | Slc9a3r1 | 2.26 | 1.41E-21 |
| 722 | Gm15287 | 3.78 | 1.42E-21 |
| 723 | Ankrd2 | 4.11 | 1.42E-21 |
| 724 | Trim34a | 2.88 | 1.49E-21 |
| 725 | Lypd8 | 6.79 | 1.54E-21 |
| 726 | Gm43136 | 3.80 | 1.59E-21 |
| 727 | Gm29684 | 7.09 | 1.60E-21 |
| 728 | Chek1 | 4.25 | 1.80E-21 |
| 729 | Dop1b | 2.02 | 1.80E-21 |
| 730 | Psors1c2 | 5.64 | 1.84E-21 |
| 731 | Taf7 | 2.01 | 2.62E-21 |
| 732 | Cd244a | 3.07 | 2.67E-21 |
| 733 | Retnlg | 5.23 | 2.86E-21 |
| 734 | Zbtb21 | 2.08 | 3.05E-21 |
| 735 | Gm44542 | 4.91 | 3.26E-21 |
| 736 | Milr1 | 2.10 | 3.40E-21 |
| 737 | Marco | 4.19 | 3.82E-21 |
| 738 | Gm31718 | 2.43 | 4.11E-21 |
| 739 | Maff | 2.29 | 4.15E-21 |
| 740 | Csf3r | 3.11 | 4.42E-21 |
| 741 | AC161438.1 | 4.58 | 4.98E-21 |
| 742 | Aldh1l2 | 4.57 | 6.99E-21 |
| 743 | 4930509G22Rik | 2.14 | 7.17E-21 |
| 744 | Gm49354 | 6.48 | 7.86E-21 |
| 745 | Apol9a | 2.24 | 8.28E-21 |
| 746 | Gm16685 | 3.70 | 9.52E-21 |
| 747 | Cxcr2 | 2.42 | 1.05E-20 |
| 748 | Myh7 | 5.34 | 1.05E-20 |
| 749 | Igsf6 | 2.25 | 1.08E-20 |
| 1124 | Reg3b | 8.33 | 7.85E-10 |
| 1125 | Spon1 | 2.06 | 8.19E-10 |
| 1126 | Ikzf4 | 2.36 | 8.71E-10 |
| 1127 | BC100530 | 4.67 | 1.06E-09 |
| 1128 | Noxred1 | 2.12 | 1.17E-09 |
| 1129 | Gm48857 | 2.19 | 1.22E-09 |
| 1130 | Oas1h | 3.81 | 1.25E-09 |
| 1131 | Styk1 | 2.96 | 1.29E-09 |
| 1132 | 1500035N22Rik | 4.82 | 1.30E-09 |
| 1133 | Slc6a14 | 5.27 | 1.31E-09 |
| 1134 | Cyp2j8 | 3.02 | 1.35E-09 |
| 1135 | Gm2061 | 2.11 | 1.38E-09 |
| 1136 | Gm49391 | 3.33 | 1.39E-09 |
| 1137 | Crispld2 | 2.07 | 1.44E-09 |
| 1138 | Gm36307 | 3.79 | 1.49E-09 |
| 1139 | Gm37101 | 2.79 | 1.52E-09 |
| 1140 | Gm28914 | 2.50 | 1.56E-09 |
| 1141 | Pdzd4 | 2.47 | 1.62E-09 |
| 1142 | BC035947 | 2.01 | 1.85E-09 |
| 1143 | Tmem252 | 2.08 | 1.87E-09 |
| 1144 | Gm31219 | 3.28 | 1.88E-09 |
| 1145 | 2010110K18Rik | 2.45 | 1.89E-09 |
| 1146 | Cd28 | 2.93 | 2.18E-09 |
| 1147 | Fam169a | 2.21 | 2.20E-09 |
| 1148 | Itga2 | 2.63 | 2.26E-09 |
| 1149 | Duoxa1 | 4.77 | 2.42E-09 |
| 1150 | 2210406H18Rik | 2.43 | 2.49E-09 |
| 1151 | Klrg2 | 2.41 | 2.56E-09 |
| 1152 | Mmp7 | 4.93 | 2.57E-09 |
| 1153 | Gucy2g | 3.38 | 3.01E-09 |
| 1154 | Tpbg | 3.06 | 3.02E-09 |
| 1155 | Arfgap3 | 2.20 | 3.09E-09 |
| 1156 | Batf3 | 2.87 | 3.09E-09 |
| 1157 | 2210414F02Rik | 4.34 | 3.11E-09 |
| 1158 | Apol7c | 2.71 | 3.30E-09 |
| 1159 | AB041806 | 2.60 | 3.58E-09 |
| 1160 | Gm48868 | 3.31 | 3.83E-09 |
| 1161 | Gpr171 | 2.44 | 4.17E-09 |
| 1162 | Gm9910 | 2.96 | 4.27E-09 |
| 1163 | Cyp1b1 | 3.28 | 4.57E-09 |
| 1164 | 2310043P16Rik | 2.26 | 4.92E-09 |
| 1165 | Cyp27b1 | 3.85 | 5.76E-09 |
| 1166 | Gm30173 | 4.96 | 5.86E-09 |
| 1167 | Gm6902 | 5.03 | 5.95E-09 |
| 1168 | Olfr1161 | 3.14 | 6.32E-09 |
| 1169 | Socs2 | 2.01 | 6.52E-09 |
| 1170 | Mab21l3 | 2.57 | 6.56E-09 |
| 1171 | Gm6034 | 2.04 | 7.23E-09 |
| 1172 | Kirrel3 | 2.27 | 8.00E-09 |
| 1173 | Gm43980 | 2.01 | 8.30E-09 |
| 1174 | Gpr37l1 | 4.58 | 8.32E-09 |
| 1175 | Cyp2t4 | 2.22 | 9.59E-09 |
| 1176 | Sult6b2 | 6.26 | 9.60E-09 |
| 1177 | Vmn1r79 | 2.55 | 1.01E-08 |
| 1178 | Ccdc154 | 2.03 | 1.01E-08 |
| 1179 | Gm43068 | 3.17 | 1.02E-08 |
| 1180 | Gm35028 | 4.52 | 1.08E-08 |
| 1181 | Gm43794 | 2.92 | 1.08E-08 |
| 1182 | Olfr877 | 2.96 | 1.11E-08 |
| 1183 | Gm37373 | 2.34 | 1.13E-08 |
| 1184 | Muc1 | 2.55 | 1.20E-08 |
| 1185 | Mybpc3 | 2.86 | 1.26E-08 |
| 1186 | Mmp9 | 2.62 | 1.30E-08 |
| 1187 | Olfr1404 | 2.33 | 1.36E-08 |
| 1188 | Acpp | 3.11 | 1.39E-08 |
| 1189 | Ppp1r3d | 2.07 | 1.51E-08 |
| 1190 | 2210009P08Rik | 5.47 | 1.54E-08 |
| 1191 | Ankrd66 | 4.48 | 1.66E-08 |
| 1192 | Gm49368 | 3.78 | 1.84E-08 |
| 1193 | AC103947.2 | 2.42 | 1.90E-08 |
| 1194 | Dsg1b | 2.96 | 1.93E-08 |
| 1195 | Foxs1 | 2.49 | 1.94E-08 |
| 1196 | Slc13a3 | 2.26 | 1.94E-08 |
| 1197 | Rhd | 2.74 | 1.94E-08 |
| 1198 | Msantd1 | 2.93 | 1.96E-08 |
| 1199 | Gm45675 | 2.53 | 2.01E-08 |
| 1200 | Ptger4 | 2.13 | 2.01E-08 |
| 1201 | Ifng | 4.83 | 2.06E-08 |
| 1202 | Il7 | 2.25 | 2.25E-08 |
| 1203 | 1700025B11Rik | 5.46 | 2.38E-08 |
| 1204 | Gm28177 | 4.03 | 2.43E-08 |
| 1205 | Vgf | 5.46 | 2.61E-08 |
| 1206 | Bhlha15 | 4.54 | 2.80E-08 |
| 1207 | Erfe | 3.47 | 2.85E-08 |
| 1208 | 4930513N20Rik | 2.75 | 3.31E-08 |
| 1209 | Gm34047 | 5.81 | 3.48E-08 |
| 1210 | 4921525O09Rik | 5.94 | 3.59E-08 |
| 1211 | Ffar2 | 3.70 | 3.65E-08 |
| 1212 | Gm10310 | 2.16 | 4.07E-08 |
| 1213 | Gm26623 | 3.35 | 4.30E-08 |
| 1214 | Gm3512 | 3.28 | 4.39E-08 |
| 1215 | Lman1l | 5.00 | 4.42E-08 |
| 1216 | Epgn | 6.41 | 4.42E-08 |
| 1217 | Slc6a19 | 3.98 | 4.58E-08 |
| 1218 | AC154912.3 | 2.30 | 4.81E-08 |
| 1219 | Fjx1 | 3.50 | 5.15E-08 |
| 1220 | Gcg | 5.72 | 5.47E-08 |
| 1221 | Scn8a | 2.23 | 5.87E-08 |
| 1222 | D330046F09Rik | 5.52 | 6.11E-08 |
| 1223 | Gm16267 | 3.71 | 6.17E-08 |
| 1224 | Heatr9 | 2.56 | 6.61E-08 |
| 1225 | 1700051A21Rik | 2.82 | 6.85E-08 |
| 1226 | Adam1a | 2.26 | 6.88E-08 |
| 1227 | Trim69 | 2.45 | 7.07E-08 |
| 1228 | Gm26902 | 3.99 | 7.25E-08 |
| 1229 | Kbtbd6 | 2.33 | 7.56E-08 |
| 1230 | Kcnn3 | 5.88 | 7.61E-08 |
| 1231 | Gm27167 | 4.29 | 8.21E-08 |
| 1232 | Gm36235 | 2.39 | 9.12E-08 |
| 1233 | Crispld1 | 4.58 | 9.17E-08 |
| 1234 | Gm34045 | 2.53 | 9.62E-08 |
| 1235 | Spx | 2.71 | 9.99E-08 |
| 1236 | Gm48199 | 2.40 | 1.02E-07 |
| 1237 | Olfr922 | 3.44 | 1.06E-07 |
| 1238 | Gm16070 | 2.28 | 1.16E-07 |
| 1239 | Gm11496 | 3.19 | 1.16E-07 |
| 1240 | Gm17017 | 2.40 | 1.20E-07 |
| 1241 | Hcn4 | 5.41 | 1.31E-07 |
| 1242 | 4933417E11Rik | 5.26 | 1.51E-07 |
| 1243 | H2-T3 | 2.14 | 1.54E-07 |
| 1244 | Trp53tg5 | 2.39 | 1.71E-07 |
| 1245 | Tmem212 | 3.33 | 1.73E-07 |
| 1246 | Myh6 | 4.55 | 1.79E-07 |
| 1247 | Gm43694 | 4.72 | 1.85E-07 |
| 1248 | Gm19866 | 2.11 | 2.16E-07 |
| 1249 | Begain | 3.78 | 2.16E-07 |
| 1624 | Gm39090 | 3.52 | 3.36E-03 |
| 1625 | 1700003D09Rik | 3.43 | 3.46E-03 |
| 1626 | Gm43731 | 2.14 | 3.57E-03 |
| 1627 | Gm29264 | 2.68 | 3.60E-03 |
| 1628 | Dnah3 | 2.66 | 3.78E-03 |
| 1629 | Gpr152 | 2.48 | 3.80E-03 |
| 1630 | 1700112J05Rik | 2.74 | 3.84E-03 |
| 1631 | Neurog2 | 3.89 | 3.86E-03 |
| 1632 | Gm37669 | 2.86 | 4.06E-03 |
| 1633 | Bdkrb1 | 2.43 | 4.07E-03 |
| 1634 | 4933425B07Rik | 2.94 | 4.26E-03 |
| 1635 | Gm49378 | 2.07 | 4.26E-03 |
| 1636 | Apod | 2.35 | 4.26E-03 |
| 1637 | Ism1 | 2.80 | 4.42E-03 |
| 1638 | Tdrd12 | 2.29 | 4.44E-03 |
| 1639 | Tmc5 | 3.40 | 4.68E-03 |
| 1640 | Gm36325 | 2.51 | 4.78E-03 |
| 1641 | Gm11559 | 2.49 | 4.83E-03 |
| 1642 | Gm16541 | 2.28 | 4.84E-03 |
| 1643 | Duox1 | 3.72 | 4.89E-03 |
| 1644 | Tent5b | 3.26 | 4.92E-03 |
| 1645 | Cryaa | 4.22 | 5.00E-03 |
| 1646 | Tmem45a2 | 2.30 | 5.03E-03 |
| 1647 | Peak1os | 3.53 | 5.06E-03 |
| 1648 | Cuzd1 | 4.28 | 5.10E-03 |
| 1649 | Ly6k | 2.09 | 5.19E-03 |
| 1650 | Glb1l2 | 2.42 | 5.20E-03 |
| 1651 | Cd163l1 | 2.81 | 5.21E-03 |
| 1652 | 4933436P19Rik | 2.42 | 5.30E-03 |
| 1653 | Gm49733 | 2.14 | 5.46E-03 |
| 1654 | Clec2l | 2.91 | 5.61E-03 |
| 1655 | Gm37011 | 3.13 | 5.65E-03 |
| 1656 | Gm36694 | 4.51 | 5.68E-03 |
| 1657 | Apobec4 | 4.02 | 5.73E-03 |
| 1658 | Ppp1r18os | 2.28 | 5.78E-03 |
| 1659 | 4930562C15Rik | 2.07 | 5.96E-03 |
| 1660 | Gm28730 | 3.03 | 6.04E-03 |
| 1661 | Slfn5os | 2.78 | 6.06E-03 |
| 1662 | Pappa | 2.14 | 6.06E-03 |
| 1663 | Plch1 | 2.91 | 6.12E-03 |
| 1664 | Tmem236 | 2.75 | 6.19E-03 |
| 1665 | Gm33340 | 2.14 | 6.20E-03 |
| 1666 | Gm15512 | 3.94 | 6.23E-03 |
| 1667 | A730089K16Rik | 4.34 | 6.36E-03 |
| 1668 | Hist1h4c | 2.02 | 6.41E-03 |
| 1669 | Bdkrb2 | 2.01 | 6.42E-03 |
| 1670 | AC138768.3 | 2.24 | 6.43E-03 |
| 1671 | Mt3 | 2.74 | 6.57E-03 |
| 1672 | Gm36987 | 4.59 | 6.58E-03 |
| 1673 | Tmem213 | 3.43 | 6.61E-03 |
| 1674 | Krt28 | 2.64 | 6.61E-03 |
| 1675 | 1500015O10Rik | 2.68 | 6.62E-03 |
| 1676 | Rpl39l | 2.35 | 6.70E-03 |
| 1677 | Zfp711 | 2.60 | 6.70E-03 |
| 1678 | Slc10a4 | 3.12 | 6.72E-03 |
| 1679 | AC164314.1 | 3.80 | 6.85E-03 |
| 1680 | Spink6 | 4.84 | 6.88E-03 |
| 1681 | Mybph | 2.54 | 6.90E-03 |
| 1682 | Dmbt1 | 8.61 | 7.06E-03 |
| 1683 | Olfr1428 | 2.92 | 7.39E-03 |
| 1684 | 1700011B04Rik | 4.77 | 7.45E-03 |
| 1685 | Gm16251 | 2.25 | 7.96E-03 |
| 1686 | Gm29825 | 3.23 | 8.03E-03 |
| 1687 | 4930597G03Rik | 4.17 | 8.03E-03 |
| 1688 | Txlnb | 3.24 | 8.07E-03 |
| 1689 | Gm37032 | 3.60 | 8.08E-03 |
| 1690 | Kctd19 | 3.73 | 8.16E-03 |
| 1691 | Cfd | 7.50 | 8.42E-03 |
| 1692 | Gdpd2 | 4.67 | 8.66E-03 |
| 1693 | Abca13 | 3.43 | 8.76E-03 |
| 1694 | Gm9945 | 2.28 | 9.00E-03 |
| 1695 | Fgf17 | 2.35 | 9.14E-03 |
| 1696 | Gm13619 | 2.23 | 9.33E-03 |
| 1697 | Muc4 | 3.03 | 9.64E-03 |
| 1698 | 1700016P03Rik | 3.09 | 9.70E-03 |
| 1699 | 1110032F04Rik | 3.09 | 9.71E-03 |
| 1700 | Spin2d | 4.60 | 9.77E-03 |
| 1701 | Gm41480 | 2.06 | 9.91E-03 |
| 1702 | Gm29025 | 5.28 | 1.01E-02 |
| 1703 | Gm3696 | 3.07 | 1.02E-02 |
| 1704 | Lvrn | 2.29 | 1.02E-02 |
| 1705 | Scnn1g | 3.67 | 1.05E-02 |
| 1706 | Gm49603 | 3.51 | 1.05E-02 |
| 1707 | Cpa3 | 2.75 | 1.08E-02 |
| 1708 | Adgrg7 | 7.37 | 1.09E-02 |
| 1709 | Dlx1 | 2.53 | 1.10E-02 |
| 1710 | U90926 | 2.66 | 1.10E-02 |
| 1711 | 1700019E08Rik | 3.96 | 1.10E-02 |
| 1712 | Gm19552 | 2.28 | 1.11E-02 |
| 1713 | Gm15562 | 4.11 | 1.11E-02 |
| 1714 | Chgb | 2.95 | 1.12E-02 |
| 1715 | Plin1 | 2.43 | 1.13E-02 |
| 1716 | Lhx3 | 3.73 | 1.18E-02 |
| 1717 | Asb5 | 2.90 | 1.18E-02 |
| 1718 | F530104D19Rik | 2.53 | 1.24E-02 |
| 1719 | Jph3 | 3.93 | 1.28E-02 |
| 1720 | Serpinb1c | 3.53 | 1.28E-02 |
| 1721 | Gm45715 | 2.22 | 1.31E-02 |
| 1722 | Fbxo39 | 2.68 | 1.32E-02 |
| 1723 | Muc5b | 2.03 | 1.32E-02 |
| 1724 | Syt2 | 2.33 | 1.37E-02 |
| 1725 | Muc15 | 2.14 | 1.38E-02 |
| 1726 | D730001G18Rik | 3.42 | 1.38E-02 |
| 1727 | Gm43195 | 2.38 | 1.39E-02 |
| 1728 | Gm6934 | 2.83 | 1.43E-02 |
| 1729 | Dnah10 | 2.38 | 1.47E-02 |
| 1730 | Togaram2 | 2.78 | 1.50E-02 |
| 1731 | Gm28710 | 3.55 | 1.52E-02 |
| 1732 | Ramp3 | 2.25 | 1.56E-02 |
| 1733 | Tac1 | 3.84 | 1.58E-02 |
| 1734 | Gm12227 | 2.92 | 1.58E-02 |
| 1735 | Gm47735 | 2.85 | 1.58E-02 |
| 1736 | Gm2832 | 3.72 | 1.59E-02 |
| 1737 | Nxph1 | 2.16 | 1.62E-02 |
| 1738 | Hsd11b2 | 2.36 | 1.64E-02 |
| 1739 | 1500009L16Rik | 2.51 | 1.66E-02 |
| 1740 | Gm30411 | 2.22 | 1.68E-02 |
| 1741 | Gm48904 | 3.60 | 1.68E-02 |
| 1742 | Prss22 | 2.88 | 1.69E-02 |
| 1743 | Gm28077 | 2.26 | 1.71E-02 |
| 1744 | Gm13860 | 3.61 | 1.73E-02 |
| 1745 | Tpsb2 | 3.13 | 1.75E-02 |
| 1746 | Gm48075 | 3.62 | 1.75E-02 |
| 1747 | Htr7 | 2.30 | 1.77E-02 |
| 1748 | Gm35048 | 2.75 | 1.81E-02 |
| 1749 | Capns2 | 2.81 | 1.82E-02 |

| | | | |
|---|---|---|---|
| 250 | Oas3 | 4.15 | 3.92E-55 |
| 251 | Clic4 | 3.20 | 5.13E-55 |
| 252 | Dusp1 | 3.47 | 5.28E-55 |
| 253 | Jak2 | 2.03 | 8.30E-55 |
| 254 | Helz2 | 3.39 | 1.30E-54 |
| 255 | Pnrc1 | 3.44 | 2.05E-54 |
| 256 | Spef2 | 5.73 | 2.14E-54 |
| 257 | Trp53inp1 | 4.12 | 2.47E-54 |
| 258 | Stat3 | 2.57 | 2.82E-54 |
| 259 | Clec4e | 7.16 | 3.61E-54 |
| 260 | Gm20559 | 3.59 | 4.75E-54 |
| 261 | Pnpt1 | 2.42 | 6.39E-54 |
| 262 | Sphk1 | 4.80 | 6.43E-54 |
| 263 | Eppk1 | 5.01 | 7.71E-54 |
| 264 | Olfr920 | 4.18 | 1.17E-53 |
| 265 | Tnfsf14 | 4.33 | 1.49E-53 |
| 266 | Psmb9 | 2.70 | 1.97E-53 |
| 267 | Gm47140 | 2.69 | 2.32E-53 |
| 268 | Spty2d1 | 2.47 | 4.29E-53 |
| 269 | Klf6 | 2.55 | 5.07E-53 |
| 270 | Irf9 | 3.08 | 5.22E-53 |
| 271 | Saa3 | 8.25 | 8.35E-53 |
| 272 | Slc41a1 | 2.71 | 9.24E-53 |
| 273 | Il1a | 3.55 | 2.38E-52 |
| 274 | Crybg1 | 2.92 | 3.12E-52 |
| 275 | Elf3 | 4.75 | 3.56E-52 |
| 276 | Dcp2 | 3.16 | 5.10E-52 |
| 277 | Glipr2 | 4.68 | 6.43E-52 |
| 278 | Ythdf1 | 2.13 | 9.48E-52 |
| 279 | Plscr1 | 5.01 | 2.58E-51 |
| 280 | Crem | 2.58 | 2.72E-51 |
| 281 | Slfn4 | 7.52 | 3.20E-51 |
| 282 | Rhoc | 2.30 | 3.45E-51 |
| 283 | Ms4a4a | 5.08 | 4.98E-51 |
| 284 | Samsn1 | 4.97 | 5.24E-51 |
| 285 | Slco3a1 | 3.20 | 5.79E-51 |
| 286 | Gm38554 | 6.30 | 9.59E-51 |
| 287 | Fam107a | 6.13 | 1.48E-50 |
| 288 | Nt5c3 | 3.16 | 2.94E-50 |
| 289 | Tdrd7 | 2.27 | 3.56E-50 |
| 290 | Stat1 | 2.86 | 3.61E-50 |
| 291 | Sbno2 | 3.06 | 4.03E-50 |
| 292 | Ddit4 | 5.07 | 4.40E-50 |
| 293 | Zc3hav1 | 2.78 | 4.42E-50 |
| 294 | Slc38a2 | 2.77 | 4.50E-50 |
| 295 | N4bp1 | 2.20 | 8.30E-50 |
| 296 | Runx1 | 3.79 | 1.53E-49 |
| 297 | Dbn1 | 3.73 | 4.38E-49 |
| 298 | Trim21 | 3.03 | 6.94E-49 |
| 299 | Ebi3 | 3.90 | 8.58E-49 |
| 300 | Uba7 | 2.97 | 1.07E-48 |
| 301 | Nr4a3 | 4.32 | 1.62E-48 |
| 302 | Sp110 | 2.70 | 1.75E-48 |
| 303 | Aida | 2.19 | 1.80E-48 |
| 304 | Ogfr | 2.39 | 3.54E-48 |
| 305 | Ms4a4c | 4.13 | 8.10E-48 |
| 306 | Olfr1328 | 4.26 | 8.19E-48 |
| 307 | Bcl2l1 | 2.62 | 9.02E-48 |
| 308 | Usb1 | 2.48 | 1.31E-47 |
| 309 | Tnfrsf23 | 3.25 | 2.18E-47 |
| 310 | Zfp36 | 3.12 | 2.27E-47 |
| 311 | Pfkfb3 | 5.08 | 2.69E-47 |
| 312 | Tmem67 | 2.78 | 3.48E-47 |
| 313 | Tacc1 | 2.37 | 4.14E-47 |
| 314 | Klf2 | 3.62 | 7.84E-47 |
| 315 | Cybb | 2.33 | 8.92E-47 |
| 316 | Psmb10 | 2.34 | 9.64E-47 |
| 317 | Nlrp12 | 6.51 | 1.31E-46 |
| 318 | Il10 | 6.99 | 1.62E-46 |
| 319 | Heatr6 | 3.86 | 1.87E-46 |
| 320 | 4930535L15Rik | 7.19 | 2.49E-46 |
| 321 | Smad1 | 2.07 | 2.97E-46 |
| 322 | Il1f9 | 4.88 | 2.99E-46 |
| 323 | Foxp4 | 2.25 | 5.78E-46 |
| 324 | Hdc | 2.77 | 6.12E-46 |
| 325 | Oas1g | 4.34 | 6.48E-46 |
| 326 | Fam208b | 2.72 | 8.32E-46 |
| 327 | Vcan | 5.45 | 1.31E-45 |
| 328 | Naa25 | 2.15 | 1.54E-45 |
| 329 | Coq8b | 3.16 | 1.81E-45 |
| 330 | Fam83c | 4.17 | 2.10E-45 |
| 331 | Elmod2 | 2.05 | 2.33E-45 |
| 332 | Mrtfa | 2.06 | 2.72E-45 |
| 333 | 2010003K11Rik | 3.00 | 2.97E-45 |
| 334 | Tbl2 | 2.01 | 2.97E-45 |
| 335 | Zbtb7c | 4.72 | 3.48E-45 |
| 336 | Itih4 | 2.25 | 3.49E-45 |
| 337 | Batf2 | 4.89 | 4.56E-45 |
| 338 | Bcl2a1b | 4.51 | 4.89E-45 |
| 339 | Tlr3 | 3.72 | 5.25E-45 |
| 340 | Nmi | 2.49 | 5.42E-45 |
| 341 | Hip1r | 2.11 | 7.65E-45 |
| 342 | Tbrg1 | 2.26 | 9.66E-45 |
| 343 | Gm48942 | 2.46 | 1.31E-44 |
| 344 | Selp | 3.88 | 2.15E-44 |
| 345 | Nfil3 | 2.49 | 2.16E-44 |
| 346 | Asap3 | 3.11 | 2.72E-44 |
| 347 | Siglece | 3.08 | 2.78E-44 |
| 348 | Insrr | 6.79 | 3.04E-44 |
| 349 | Sema4c | 3.54 | 3.67E-44 |
| 350 | Cdkn1a | 6.64 | 5.39E-44 |
| 351 | Gm10251 | 5.53 | 6.16E-44 |
| 352 | Nlrc5 | 2.98 | 6.38E-44 |
| 353 | Hck | 3.39 | 6.54E-44 |
| 354 | Tapbpl | 2.46 | 6.86E-44 |
| 355 | Ccl19 | 5.23 | 2.33E-43 |
| 356 | Slc39a14 | 2.77 | 2.54E-43 |
| 357 | Adprh | 2.89 | 3.15E-43 |
| 358 | Fcer1g | 2.15 | 4.80E-43 |
| 359 | Bag3 | 2.71 | 6.88E-43 |
| 360 | Slmapos2 | 2.42 | 9.65E-43 |
| 361 | Creb3l2 | 2.22 | 9.69E-43 |
| 362 | Btg2 | 3.22 | 9.83E-43 |
| 363 | Rnf19b | 2.99 | 1.06E-42 |
| 364 | Pawr | 2.68 | 1.11E-42 |
| 365 | Itprip | 3.09 | 1.11E-42 |
| 366 | Ifitm5 | 4.35 | 1.30E-42 |
| 367 | Rasgef1b | 3.07 | 1.64E-42 |
| 368 | Edn1 | 3.96 | 2.58E-42 |
| 369 | Clec4d | 5.68 | 2.69E-42 |
| 370 | Samd9l | 4.20 | 3.16E-42 |
| 371 | Gm19705 | 2.96 | 3.45E-42 |
| 372 | Gm12185 | 5.14 | 4.32E-42 |
| 373 | Nfkb1 | 2.02 | 4.58E-42 |
| 374 | Trim25 | 2.11 | 5.22E-42 |
| 375 | Chmp4b | 2.06 | 5.89E-42 |
| 750 | Col9a3 | 3.39 | 1.09E-20 |
| 751 | Soat2 | 2.43 | 1.50E-20 |
| 752 | Mamstr | 3.51 | 1.55E-20 |
| 753 | Sh3pxd2b | 2.21 | 1.59E-20 |
| 754 | Lyve1 | 2.93 | 1.71E-20 |
| 755 | Gm42493 | 3.02 | 1.91E-20 |
| 756 | Rbl1 | 2.11 | 2.02E-20 |
| 757 | Zc3h7a | 2.41 | 2.02E-20 |
| 758 | Btbd16 | 3.77 | 2.21E-20 |
| 759 | Gm21149 | 6.09 | 2.58E-20 |
| 760 | Sell | 2.54 | 2.84E-20 |
| 761 | Fam222a | 2.40 | 2.96E-20 |
| 762 | Gm45774 | 4.18 | 3.47E-20 |
| 763 | Bbx | 2.14 | 3.92E-20 |
| 764 | Gm20219 | 3.09 | 3.95E-20 |
| 765 | Gdap10 | 3.25 | 4.41E-20 |
| 766 | Amigo2 | 3.75 | 4.66E-20 |
| 767 | Gm11592 | 4.69 | 4.85E-20 |
| 768 | Ifit1bl1 | 4.56 | 4.91E-20 |
| 769 | Sec1 | 6.18 | 5.14E-20 |
| 770 | Samd4 | 3.11 | 5.26E-20 |
| 771 | Gm14010 | 5.01 | 5.39E-20 |
| 772 | Gm11963 | 3.69 | 6.77E-20 |
| 773 | A930037H05Rik | 2.64 | 7.11E-20 |
| 774 | Gm15998 | 2.77 | 7.59E-20 |
| 775 | Lrrc75a | 2.81 | 9.55E-20 |
| 776 | Olfr99 | 3.82 | 9.83E-20 |
| 777 | Adam9 | 2.13 | 1.17E-19 |
| 778 | Ccl22 | 5.03 | 1.26E-19 |
| 779 | Gm41144 | 3.55 | 1.29E-19 |
| 780 | Hsh2d | 2.50 | 1.31E-19 |
| 781 | Abcc1 | 2.18 | 1.39E-19 |
| 782 | Zpr1 | 3.15 | 1.50E-19 |
| 783 | Plekhd1 | 4.12 | 1.62E-19 |
| 784 | Cerkl | 2.24 | 1.83E-19 |
| 785 | Ngp | 4.97 | 2.13E-19 |
| 786 | Syvn1 | 2.96 | 2.16E-19 |
| 787 | Cilp2 | 5.05 | 2.36E-19 |
| 788 | Slc13a4 | 3.16 | 2.52E-19 |
| 789 | Ncf4 | 2.14 | 2.77E-19 |
| 790 | Dusp5 | 2.20 | 2.84E-19 |
| 791 | Tox4 | 2.17 | 3.04E-19 |
| 792 | Themis2 | 2.07 | 3.20E-19 |
| 793 | Rcan1 | 2.19 | 3.79E-19 |
| 794 | Gas6 | 2.44 | 4.37E-19 |
| 795 | S100a8 | 5.52 | 4.59E-19 |
| 796 | Asprv1 | 4.04 | 4.72E-19 |
| 797 | Gm33091 | 5.31 | 7.55E-19 |
| 798 | Prr7 | 4.24 | 7.65E-19 |
| 799 | Cxcl5 | 4.55 | 8.66E-19 |
| 800 | Fbxl22 | 2.84 | 9.44E-19 |
| 801 | Adora2b | 5.39 | 9.90E-19 |
| 802 | Gm34354 | 2.97 | 1.03E-18 |
| 803 | Ifitm6 | 3.57 | 1.06E-18 |
| 804 | Arfip2 | 2.19 | 1.24E-18 |
| 805 | Cd33 | 2.80 | 1.31E-18 |
| 806 | Sycp2 | 4.21 | 1.33E-18 |
| 807 | Gm11772 | 3.41 | 1.37E-18 |
| 808 | Lpar1 | 2.26 | 1.47E-18 |
| 809 | Gm31036 | 3.17 | 1.79E-18 |
| 810 | Gzmb | 4.68 | 2.54E-18 |
| 811 | Gm20496 | 3.53 | 2.59E-18 |
| 812 | Tcp10b | 2.82 | 2.79E-18 |
| 813 | Epsti1 | 2.47 | 3.02E-18 |
| 814 | Slc6a9 | 2.31 | 3.31E-18 |
| 815 | Il1r1 | 3.28 | 3.46E-18 |
| 816 | AC119228.1 | 4.05 | 4.17E-18 |
| 817 | Kcnj2 | 2.47 | 4.56E-18 |
| 818 | Sprr2a2 | 4.93 | 4.58E-18 |
| 819 | Etv4 | 3.09 | 4.96E-18 |
| 820 | Fgl1 | 2.33 | 5.00E-18 |
| 821 | Hbegf | 2.36 | 5.29E-18 |
| 822 | E030018B13Rik | 4.58 | 5.41E-18 |
| 823 | Fsd1 | 3.86 | 5.41E-18 |
| 824 | Pim2 | 2.03 | 6.20E-18 |
| 825 | Pced1b | 2.27 | 6.69E-18 |
| 826 | Csf2rb2 | 2.14 | 7.37E-18 |
| 827 | Dnajb9 | 2.23 | 7.49E-18 |
| 828 | Gfpt2 | 3.77 | 1.28E-17 |
| 829 | Gab2 | 2.39 | 1.35E-17 |
| 830 | Tmem171 | 4.69 | 1.36E-17 |
| 831 | Trem3 | 4.95 | 1.51E-17 |
| 832 | Vmn1r43 | 2.33 | 1.52E-17 |
| 833 | Gm49123 | 4.45 | 1.71E-17 |
| 834 | F630028O10Rik | 2.83 | 1.79E-17 |
| 835 | Saa1 | 5.58 | 2.29E-17 |
| 836 | Oas2 | 2.82 | 2.54E-17 |
| 837 | Ifi211 | 3.32 | 2.79E-17 |
| 838 | Nfxl1 | 2.45 | 3.61E-17 |
| 839 | Gm37880 | 3.22 | 4.39E-17 |
| 840 | Gm15787 | 2.60 | 4.72E-17 |
| 841 | Duox2 | 7.26 | 4.75E-17 |
| 842 | C1qtnf12 | 2.07 | 4.95E-17 |
| 843 | Itgam | 2.99 | 5.00E-17 |
| 844 | Gm45629 | 2.39 | 5.07E-17 |
| 845 | Shh | 2.47 | 6.67E-17 |
| 846 | Fos | 2.66 | 6.90E-17 |
| 847 | Gm15056 | 6.85 | 7.79E-17 |
| 848 | Gm49339 | 2.46 | 9.46E-17 |
| 849 | Apol11a | 5.44 | 9.84E-17 |
| 850 | Olfr1033 | 2.07 | 1.05E-16 |
| 851 | Cgas | 2.20 | 1.17E-16 |
| 852 | Gprc5b | 3.91 | 1.19E-16 |
| 853 | Gm49482 | 7.21 | 1.26E-16 |
| 854 | A530040E14Rik | 2.62 | 1.33E-16 |
| 855 | Ereg | 4.20 | 1.36E-16 |
| 856 | Dnah1 | 3.93 | 1.47E-16 |
| 857 | Aspm | 2.62 | 1.48E-16 |
| 858 | Phf11c | 2.01 | 1.73E-16 |
| 859 | Arhgap8 | 3.45 | 1.85E-16 |
| 860 | Rab37 | 3.00 | 1.85E-16 |
| 861 | Arhgap22 | 2.12 | 1.92E-16 |
| 862 | Adora2a | 2.12 | 1.93E-16 |
| 863 | Arntl2 | 3.79 | 1.96E-16 |
| 864 | Pf4 | 2.43 | 2.36E-16 |
| 865 | Nr4a1 | 3.38 | 2.49E-16 |
| 866 | Stfa2l1 | 4.40 | 2.59E-16 |
| 867 | Abcb1b | 3.00 | 2.62E-16 |
| 868 | Gm10388 | 2.22 | 2.68E-16 |
| 869 | AC167363.2 | 7.05 | 2.86E-16 |
| 870 | Olfm4 | 6.63 | 2.89E-16 |
| 871 | Gm42031 | 3.28 | 3.00E-16 |
| 872 | Gm37701 | 3.03 | 3.18E-16 |
| 873 | 4930527F14Rik | 4.01 | 3.79E-16 |
| 874 | Egr2 | 3.92 | 4.09E-16 |
| 875 | Plscr2 | 2.09 | 4.74E-16 |
| 1250 | Sytl3 | 2.78 | 2.17E-07 |
| 1251 | Gm37248 | 2.25 | 2.28E-07 |
| 1252 | Trim36 | 2.18 | 2.34E-07 |
| 1253 | Fgf23 | 3.55 | 2.50E-07 |
| 1254 | 4930540M05Rik | 3.23 | 2.59E-07 |
| 1255 | Gm14548 | 2.03 | 2.62E-07 |
| 1256 | Gm16094 | 2.78 | 2.63E-07 |
| 1257 | Amh | 2.80 | 2.94E-07 |
| 1258 | Gm48071 | 2.38 | 3.03E-07 |
| 1259 | Gm20663 | 2.46 | 3.30E-07 |
| 1260 | 4933413C19Rik | 3.36 | 3.34E-07 |
| 1261 | Gm40645 | 2.20 | 3.88E-07 |
| 1262 | Gm34907 | 3.61 | 3.88E-07 |
| 1263 | Mfap5 | 4.32 | 3.91E-07 |
| 1264 | Hopxos | 2.12 | 4.00E-07 |
| 1265 | Gm49342 | 2.62 | 4.04E-07 |
| 1266 | Orm2 | 3.00 | 4.35E-07 |
| 1267 | Sptbn5 | 2.59 | 4.46E-07 |
| 1268 | Pirt | 2.21 | 4.51E-07 |
| 1269 | Map1b | 2.25 | 4.69E-07 |
| 1270 | Gm17359 | 2.52 | 4.78E-07 |
| 1271 | Gzma | 2.53 | 5.02E-07 |
| 1272 | 4930580E04Rik | 4.39 | 5.20E-07 |
| 1273 | 9330121K16Rik | 2.34 | 5.35E-07 |
| 1274 | Serpinb9b | 2.93 | 5.48E-07 |
| 1275 | Gm44371 | 2.36 | 5.50E-07 |
| 1276 | Ly6d | 3.16 | 5.73E-07 |
| 1277 | Gm45684 | 2.09 | 5.88E-07 |
| 1278 | Gm37666 | 2.31 | 6.03E-07 |
| 1279 | Misp | 2.43 | 6.19E-07 |
| 1280 | Ccl8 | 3.66 | 6.27E-07 |
| 1281 | Tchh | 2.62 | 6.27E-07 |
| 1282 | Gm48203 | 2.76 | 6.31E-07 |
| 1283 | Pdcd1 | 4.01 | 6.58E-07 |
| 1284 | Frrs1l | 2.27 | 6.63E-07 |
| 1285 | Ccdc74a | 2.49 | 6.71E-07 |
| 1286 | Gm14817 | 3.88 | 6.75E-07 |
| 1287 | Gm12764 | 2.81 | 6.76E-07 |
| 1288 | 2610306O10Rik | 2.73 | 6.81E-07 |
| 1289 | Eno2 | 2.17 | 7.29E-07 |
| 1290 | Npy | 3.43 | 7.55E-07 |
| 1291 | Mfsd6l | 2.33 | 7.58E-07 |
| 1292 | Prps1l1 | 2.72 | 7.60E-07 |
| 1293 | Gm28863 | 5.16 | 7.79E-07 |
| 1294 | Prok2 | 5.60 | 7.89E-07 |
| 1295 | Pram1 | 2.93 | 7.90E-07 |
| 1296 | Gm47242 | 4.19 | 8.05E-07 |
| 1297 | Gm614 | 3.22 | 8.39E-07 |
| 1298 | Gm37347 | 5.24 | 8.80E-07 |
| 1299 | Trdn | 2.77 | 9.48E-07 |
| 1300 | Asns | 2.31 | 1.00E-06 |
| 1301 | Csf2 | 3.14 | 1.07E-06 |
| 1302 | 1700001P01Rik | 2.41 | 1.09E-06 |
| 1303 | Gm49695 | 2.25 | 1.19E-06 |
| 1304 | Gm26644 | 2.22 | 1.21E-06 |
| 1305 | Reg3a | 5.50 | 1.24E-06 |
| 1306 | Serpina7 | 2.77 | 1.25E-06 |
| 1307 | Ntn5 | 4.04 | 1.26E-06 |
| 1308 | Tmem63c | 4.18 | 1.30E-06 |
| 1309 | Gm11985 | 5.20 | 1.34E-06 |
| 1310 | Lepr | 2.84 | 1.44E-06 |
| 1311 | Tigd4 | 2.25 | 1.45E-06 |
| 1312 | Mybpc2 | 2.39 | 1.48E-06 |
| 1313 | 1700045H11Rik | 2.40 | 1.49E-06 |
| 1314 | Gm48853 | 2.56 | 1.51E-06 |
| 1315 | Gm29443 | 3.04 | 1.70E-06 |
| 1316 | Gm45137 | 2.44 | 1.72E-06 |
| 1317 | AC151478.3 | 3.99 | 1.87E-06 |
| 1318 | Lamc2 | 2.19 | 2.00E-06 |
| 1319 | Fut2 | 3.05 | 2.05E-06 |
| 1320 | Gm20234 | 2.87 | 2.08E-06 |
| 1321 | Gm15990 | 2.72 | 2.25E-06 |
| 1322 | Gm26545 | 2.10 | 2.34E-06 |
| 1323 | Cym | 2.03 | 2.47E-06 |
| 1324 | Gm45708 | 2.35 | 2.56E-06 |
| 1325 | Ms4a8a | 2.04 | 2.62E-06 |
| 1326 | Dach1 | 2.27 | 2.75E-06 |
| 1327 | Trim30c | 2.43 | 2.82E-06 |
| 1328 | Gm45564 | 3.06 | 2.85E-06 |
| 1329 | Slc7a15 | 3.56 | 3.01E-06 |
| 1330 | Tarm1 | 2.59 | 3.02E-06 |
| 1331 | Sirpb1a | 2.61 | 3.03E-06 |
| 1332 | Gm2396 | 2.69 | 3.24E-06 |
| 1333 | Fam229a | 3.31 | 3.36E-06 |
| 1334 | Shisa7 | 2.65 | 3.49E-06 |
| 1335 | Cyp21a1 | 2.48 | 3.71E-06 |
| 1336 | Dlx3 | 3.06 | 3.76E-06 |
| 1337 | Cdh23 | 2.96 | 3.85E-06 |
| 1338 | Apol10a | 2.30 | 3.97E-06 |
| 1339 | Clcn1 | 2.86 | 4.19E-06 |
| 1340 | Uchl1 | 4.04 | 4.31E-06 |
| 1341 | 1700029B24Rik | 5.89 | 4.34E-06 |
| 1342 | Gm43064 | 3.45 | 4.40E-06 |
| 1343 | Dnah9 | 3.28 | 4.49E-06 |
| 1344 | Mmp3 | 4.67 | 4.69E-06 |
| 1345 | Osr1 | 3.27 | 4.79E-06 |
| 1346 | AC132253.2 | 4.07 | 5.28E-06 |
| 1347 | Gm17089 | 2.77 | 5.39E-06 |
| 1348 | 1700071M16Rik | 2.98 | 5.52E-06 |
| 1349 | Cpt1b | 2.13 | 5.53E-06 |
| 1350 | Gm35501 | 2.19 | 5.53E-06 |
| 1351 | Gm49132 | 2.10 | 5.84E-06 |
| 1352 | Gm31518 | 2.19 | 6.28E-06 |
| 1353 | Ltf | 3.36 | 6.65E-06 |
| 1354 | Gm31683 | 2.28 | 6.65E-06 |
| 1355 | Gprc5a | 2.63 | 6.68E-06 |
| 1356 | Glp2r | 3.05 | 6.73E-06 |
| 1357 | Crtam | 2.82 | 6.81E-06 |
| 1358 | Gng3 | 3.26 | 6.98E-06 |
| 1359 | Slco4c1 | 3.21 | 7.48E-06 |
| 1360 | Gm10662 | 3.81 | 7.66E-06 |
| 1361 | Gm5891 | 3.81 | 7.66E-06 |
| 1362 | 9130015A21Rik | 5.53 | 7.68E-06 |
| 1363 | Has1 | 3.54 | 7.80E-06 |
| 1364 | Chst11 | 2.16 | 8.33E-06 |
| 1365 | Mrgprh | 6.02 | 8.34E-06 |
| 1366 | Fasl | 2.51 | 9.12E-06 |
| 1367 | Stc1 | 2.58 | 9.22E-06 |
| 1368 | 4930478E11Rik | 2.81 | 9.77E-06 |
| 1369 | Gm6592 | 2.66 | 9.77E-06 |
| 1370 | Cabyr | 2.17 | 9.87E-06 |
| 1371 | Rasl10b | 2.61 | 9.94E-06 |
| 1372 | Pgf | 2.70 | 1.07E-05 |
| 1373 | Slc51b | 2.67 | 1.08E-05 |
| 1374 | AC165072.1 | 2.16 | 1.12E-05 |
| 1375 | Plscr5 | 2.49 | 1.13E-05 |
| 1750 | Egr3 | 2.04 | 1.84E-02 |
| 1751 | Gm49427 | 3.43 | 1.85E-02 |
| 1752 | Il1f6 | 3.41 | 1.86E-02 |
| 1753 | Gm26626 | 2.31 | 1.96E-02 |
| 1754 | Nkain1 | 2.63 | 2.00E-02 |
| 1755 | Lrfn5 | 2.93 | 2.08E-02 |
| 1756 | Wnt10b | 2.12 | 2.12E-02 |
| 1757 | Gm13398 | 3.17 | 2.12E-02 |
| 1758 | Gm43748 | 4.00 | 2.12E-02 |
| 1759 | Cfap44 | 2.44 | 2.31E-02 |
| 1760 | 9930022D16Rik | 2.48 | 2.33E-02 |
| 1761 | Krt17 | 3.89 | 2.40E-02 |
| 1762 | Gm41219 | 2.59 | 2.43E-02 |
| 1763 | AC129217.1 | 4.82 | 2.45E-02 |
| 1764 | Gm15518 | 2.48 | 2.45E-02 |
| 1765 | Gm21083 | 2.81 | 2.46E-02 |
| 1766 | Psg19 | 3.95 | 2.47E-02 |
| 1767 | Gm3286 | 4.13 | 2.51E-02 |
| 1768 | Obox3 | 3.21 | 2.52E-02 |
| 1769 | Gm38025 | 2.21 | 2.55E-02 |
| 1770 | Ntrk1 | 2.12 | 2.56E-02 |
| 1771 | Fam83e | 3.26 | 2.57E-02 |
| 1772 | 1700047I17Rik2 | 4.56 | 2.58E-02 |
| 1773 | Gm47704 | 2.03 | 2.60E-02 |
| 1774 | Gm11842 | 2.03 | 2.66E-02 |
| 1775 | Slc9a3 | 4.00 | 2.75E-02 |
| 1776 | Gm15895 | 3.75 | 2.76E-02 |
| 1777 | E230025N22Rik | 2.08 | 2.76E-02 |
| 1778 | Gm12205 | 2.13 | 2.77E-02 |
| 1779 | Gm44012 | 3.20 | 2.77E-02 |
| 1780 | Gm37530 | 2.94 | 2.78E-02 |
| 1781 | Igkv4-55 | 3.73 | 2.81E-02 |
| 1782 | Hoxc9 | 2.07 | 2.83E-02 |
| 1783 | Adipoq | 3.79 | 2.87E-02 |
| 1784 | Gm33424 | 2.49 | 2.87E-02 |
| 1785 | Plekhg4 | 2.26 | 2.91E-02 |
| 1786 | AC132253.5 | 2.08 | 2.93E-02 |
| 1787 | Gm45238 | 3.70 | 2.94E-02 |
| 1788 | Gm28235 | 4.21 | 2.95E-02 |
| 1789 | Stra6 | 4.58 | 2.95E-02 |
| 1790 | Gm49182 | 3.14 | 2.96E-02 |
| 1791 | Cidea | 2.24 | 3.03E-02 |
| 1792 | Muc3 | 6.04 | 3.08E-02 |
| 1793 | Gm5627 | 3.29 | 3.16E-02 |
| 1794 | Olfr1016 | 3.10 | 3.22E-02 |
| 1795 | Guca2b | 2.55 | 3.25E-02 |
| 1796 | Gm49227 | 2.85 | 3.30E-02 |
| 1797 | Gm49776 | 3.54 | 3.31E-02 |
| 1798 | Nxf2 | 2.50 | 3.33E-02 |
| 1799 | Tent5d | 2.81 | 3.33E-02 |
| 1800 | Gm15218 | 2.62 | 3.42E-02 |
| 1801 | 4921528I07Rik | 4.48 | 3.44E-02 |
| 1802 | Gm29233 | 3.24 | 3.48E-02 |
| 1803 | Actg2 | 3.38 | 3.48E-02 |
| 1804 | Hesx1 | 3.44 | 3.52E-02 |
| 1805 | Gm21748 | 3.31 | 3.53E-02 |
| 1806 | Gm21860 | 3.31 | 3.53E-02 |
| 1807 | Dusp15 | 2.82 | 3.59E-02 |
| 1808 | Gm41043 | 3.09 | 3.61E-02 |
| 1809 | Areg | 3.09 | 3.67E-02 |
| 1810 | Ptger3 | 2.30 | 3.68E-02 |
| 1811 | Spdef | 2.82 | 3.73E-02 |
| 1812 | Olfr753-ps1 | 3.41 | 3.74E-02 |
| 1813 | Sema7a | 2.05 | 3.74E-02 |
| 1814 | Gm3636 | 2.52 | 3.75E-02 |
| 1815 | Gm19951 | 2.28 | 3.76E-02 |
| 1816 | Gm32939 | 3.33 | 3.80E-02 |
| 1817 | Acrv1 | 3.19 | 3.84E-02 |
| 1818 | Igkv14-126 | 2.08 | 3.90E-02 |
| 1819 | Cel | 2.23 | 3.96E-02 |
| 1820 | Gm20089 | 3.00 | 4.00E-02 |
| 1821 | 4930509H03Rik | 2.43 | 4.29E-02 |
| 1822 | Gm42800 | 3.87 | 4.29E-02 |
| 1823 | 9230109A22Rik | 3.40 | 4.30E-02 |
| 1824 | Bcl2l14 | 2.09 | 4.31E-02 |
| 1825 | Gm49618 | 3.37 | 4.35E-02 |
| 1826 | Gm15684 | 3.10 | 4.35E-02 |
| 1827 | Scel | 3.33 | 4.52E-02 |
| 1828 | Il20ra | 3.36 | 4.53E-02 |
| 1829 | Skor1 | 2.14 | 4.54E-02 |
| 1830 | Olfr549 | 3.66 | 4.56E-02 |
| 1831 | Gm48950 | 2.96 | 4.58E-02 |
| 1832 | Gm11273 | 2.11 | 4.62E-02 |
| 1833 | Ppp1r1a | 2.94 | 4.64E-02 |
| 1834 | Gm12117 | 2.36 | 4.71E-02 |
| 1835 | Mip | 3.63 | 4.73E-02 |
| 1836 | Ighv11-2 | 3.42 | 4.75E-02 |
| 1837 | Syt8 | 2.79 | 4.93E-02 |

Table S9

| | | | |
|---|---|---|---|
| 376 | Abca1 | 2.09 | 7.42E-42 |
| 377 | Gm4070 | 4.64 | 7.49E-42 |
| 378 | Tub | 6.06 | 9.28E-42 |
| 379 | Gm36161 | 4.02 | 1.68E-41 |
| 380 | Enpp4 | 3.27 | 1.68E-41 |
| 381 | Ifi207 | 3.20 | 1.71E-41 |
| 382 | Dhx58os | 4.63 | 1.82E-41 |
| 383 | Gm2564 | 4.69 | 2.02E-41 |
| 384 | Pcsk5 | 2.23 | 2.38E-41 |
| 385 | Tnfsf10 | 3.85 | 3.46E-41 |
| 386 | AA467197 | 7.46 | 6.83E-41 |
| 387 | Nub1 | 2.07 | 7.74E-41 |
| 388 | Birc2 | 2.84 | 8.98E-41 |
| 389 | Duoxa2 | 8.73 | 1.00E-40 |
| 390 | Rtn4rl2 | 5.64 | 1.16E-40 |
| 391 | Il27 | 5.36 | 1.43E-40 |
| 392 | Map3k8 | 2.97 | 1.73E-40 |
| 393 | Ifi209 | 3.73 | 1.78E-40 |
| 394 | Trim34b | 2.84 | 1.89E-40 |
| 395 | Plekha4 | 4.84 | 2.22E-40 |
| 396 | Adamts9 | 3.67 | 2.60E-40 |
| 397 | Tor1aip2 | 2.20 | 2.89E-40 |
| 398 | Cfap69 | 5.24 | 4.14E-40 |
| 399 | Tnip3 | 6.35 | 4.83E-40 |
| 400 | Nedd4l | 2.03 | 5.08E-40 |
| 401 | Gm47806 | 3.91 | 6.42E-40 |
| 402 | Nol10 | 2.10 | 7.63E-40 |
| 403 | Enc1 | 2.94 | 7.79E-40 |
| 404 | Ror1 | 3.21 | 8.17E-40 |
| 405 | Lilrb4a | 3.19 | 8.49E-40 |
| 406 | Tmem140 | 3.04 | 9.11E-40 |
| 407 | Arhgef26 | 2.70 | 1.19E-39 |
| 408 | Trim30b | 3.86 | 1.21E-39 |
| 409 | Lpgat1 | 2.35 | 1.43E-39 |
| 410 | Sarm1 | 3.46 | 1.61E-39 |
| 411 | Pim1 | 2.42 | 1.93E-39 |
| 412 | Alpk1 | 3.25 | 2.69E-39 |
| 413 | Fgr | 2.87 | 4.39E-39 |
| 414 | Steap1 | 5.24 | 4.84E-39 |
| 415 | Casz1 | 3.22 | 5.09E-39 |
| 416 | Fyb2 | 2.76 | 6.09E-39 |
| 417 | Gm27252 | 2.85 | 7.31E-39 |
| 418 | Gm5483 | 7.39 | 7.39E-39 |
| 419 | Cdhr4 | 5.72 | 9.65E-39 |
| 420 | Ehd4 | 2.40 | 1.49E-38 |
| 421 | Bcl2a1d | 5.45 | 1.95E-38 |
| 422 | Tnfrsf1b | 2.22 | 2.03E-38 |
| 423 | Gm43513 | 4.02 | 2.05E-38 |
| 424 | Ccdc6 | 2.06 | 2.08E-38 |
| 425 | Sav1 | 2.12 | 2.13E-38 |
| 426 | Gm35309 | 6.91 | 2.35E-38 |
| 427 | H2-Q2 | 2.11 | 2.52E-38 |
| 428 | Fbf1 | 3.18 | 2.71E-38 |
| 429 | Fpr1 | 3.52 | 3.25E-38 |
| 430 | Trim30d | 3.97 | 3.76E-38 |
| 431 | Hap1 | 3.70 | 4.11E-38 |
| 432 | Gm15567 | 4.25 | 5.70E-38 |
| 433 | Spata13 | 2.02 | 6.06E-38 |
| 434 | Spats2l | 4.85 | 6.11E-38 |
| 435 | Katna1 | 2.81 | 7.91E-38 |
| 436 | Marcks | 2.08 | 1.27E-37 |
| 437 | Ier3 | 5.69 | 1.34E-37 |
| 438 | Gm17139 | 2.70 | 2.61E-37 |
| 439 | Gm26809 | 2.51 | 3.08E-37 |
| 440 | Wdr43 | 2.20 | 3.22E-37 |
| 441 | Gna13 | 2.20 | 3.38E-37 |
| 442 | Lilr4b | 3.05 | 4.02E-37 |
| 443 | Tut7 | 2.05 | 5.31E-37 |
| 444 | Mcoln2 | 4.54 | 5.62E-37 |
| 445 | Oas1c | 3.08 | 5.90E-37 |
| 446 | Gm16175 | 4.00 | 8.04E-37 |
| 447 | Sh3bp2 | 2.32 | 8.13E-37 |
| 448 | S100a11 | 2.41 | 1.02E-36 |
| 449 | Nop58 | 2.73 | 1.04E-36 |
| 450 | Defb1 | 3.98 | 1.28E-36 |
| 451 | Rnf225 | 2.90 | 1.30E-36 |
| 452 | Bmper | 5.10 | 1.39E-36 |
| 453 | Adrb2 | 3.45 | 1.43E-36 |
| 454 | Ube2s | 2.56 | 2.09E-36 |
| 455 | Nlrp3 | 4.33 | 3.48E-36 |
| 456 | Apex1 | 2.46 | 3.69E-36 |
| 457 | Cyb561 | 4.80 | 4.53E-36 |
| 458 | Susd1 | 2.77 | 4.73E-36 |
| 459 | Tmco4 | 2.11 | 5.99E-36 |
| 460 | Rassf1 | 2.55 | 6.33E-36 |
| 461 | Cd40 | 2.98 | 7.07E-36 |
| 462 | AI506816 | 3.00 | 7.47E-36 |
| 463 | Gan | 2.59 | 7.50E-36 |
| 464 | Fam199x | 2.28 | 7.84E-36 |
| 465 | Nostrin | 2.02 | 8.09E-36 |
| 466 | Cxcl13 | 5.90 | 8.37E-36 |
| 467 | Fscn1 | 2.93 | 1.27E-35 |
| 468 | Phc2 | 2.60 | 1.51E-35 |
| 469 | 4930512H18Rik | 4.20 | 2.13E-35 |
| 470 | Slc7a1 | 2.93 | 2.27E-35 |
| 471 | Ralgds | 4.46 | 2.30E-35 |
| 472 | Nod2 | 4.06 | 3.07E-35 |
| 473 | Car4 | 9.68 | 3.32E-35 |
| 474 | Gm5431 | 4.29 | 3.64E-35 |
| 475 | Slc25a18 | 5.15 | 4.09E-35 |
| 476 | Slc7a6 | 4.15 | 4.59E-35 |
| 477 | Gm16196 | 4.97 | 4.67E-35 |
| 478 | Reep2 | 4.35 | 5.67E-35 |
| 479 | Ccl9 | 2.15 | 8.55E-35 |
| 480 | Lrrc25 | 2.46 | 8.72E-35 |
| 481 | Nampt | 2.72 | 9.39E-35 |
| 482 | Dtx3l | 2.18 | 1.13E-34 |
| 483 | Il18rap | 3.45 | 1.29E-34 |
| 484 | Gm7694 | 3.07 | 1.43E-34 |
| 485 | Gm42900 | 3.05 | 1.45E-34 |
| 486 | 4732465J04Rik | 2.31 | 1.46E-34 |
| 487 | Gm42764 | 3.11 | 1.53E-34 |
| 488 | Cd69 | 4.12 | 1.63E-34 |
| 489 | Nfkbib | 2.07 | 1.98E-34 |
| 490 | Ier5 | 2.26 | 3.09E-34 |
| 491 | 4930599N23Rik | 3.32 | 4.20E-34 |
| 492 | Ppp1r10 | 2.37 | 4.44E-34 |
| 493 | Trem1 | 6.15 | 4.48E-34 |
| 494 | Atxn1 | 2.45 | 6.63E-34 |
| 495 | Garem1 | 2.10 | 7.64E-34 |
| 496 | Tmem39a | 2.27 | 8.41E-34 |
| 497 | 3110001I22Rik | 2.62 | 8.89E-34 |
| 498 | Prg4 | 2.61 | 9.07E-34 |
| 499 | 9530077C05Rik | 6.38 | 2.21E-33 |
| 500 | Nupr1 | 3.98 | 2.64E-33 |
| 876 | Ube2v2 | 2.76 | 5.12E-16 |
| 877 | Aoah | 3.12 | 5.67E-16 |
| 878 | 5430427O19Rik | 2.55 | 6.91E-16 |
| 879 | Gm4262 | 3.45 | 7.35E-16 |
| 880 | Ceacam18 | 3.44 | 7.53E-16 |
| 881 | A730081D07Rik | 2.16 | 7.76E-16 |
| 882 | Sox9 | 2.69 | 8.62E-16 |
| 883 | Gm5420 | 3.15 | 8.79E-16 |
| 884 | Irs2 | 2.55 | 9.28E-16 |
| 885 | Gm29427 | 2.88 | 1.02E-15 |
| 886 | Gm15821 | 2.36 | 1.04E-15 |
| 887 | Gm35279 | 2.99 | 1.26E-15 |
| 888 | Gm42726 | 2.58 | 1.37E-15 |
| 889 | Dmtn | 3.03 | 1.40E-15 |
| 890 | Gbp11 | 3.12 | 1.46E-15 |
| 891 | Clec5a | 4.39 | 1.53E-15 |
| 892 | Sftpd | 2.12 | 1.59E-15 |
| 893 | Gm38259 | 4.59 | 1.79E-15 |
| 894 | Fosl1 | 5.47 | 1.91E-15 |
| 895 | Gm15418 | 4.23 | 2.23E-15 |
| 896 | Gm14168 | 2.71 | 2.36E-15 |
| 897 | Ms4a6b | 2.19 | 2.47E-15 |
| 898 | Platr22 | 2.64 | 2.62E-15 |
| 899 | Hp | 2.04 | 2.65E-15 |
| 900 | Orai2 | 2.61 | 2.70E-15 |
| 901 | Rdh12 | 4.02 | 2.80E-15 |
| 902 | Gm37043 | 4.50 | 2.93E-15 |
| 903 | Gm21188 | 3.23 | 2.94E-15 |
| 904 | Npepps | 2.69 | 3.01E-15 |
| 905 | Gm45418 | 3.19 | 3.15E-15 |
| 906 | Mcemp1 | 3.97 | 3.32E-15 |
| 907 | Lrg1 | 2.71 | 4.12E-15 |
| 908 | Adam8 | 2.76 | 4.31E-15 |
| 909 | Nat14 | 2.23 | 4.50E-15 |
| 910 | Unc79 | 4.08 | 4.64E-15 |
| 911 | Treml2 | 2.96 | 4.93E-15 |
| 912 | Creb5 | 2.83 | 5.55E-15 |
| 913 | Lilra6 | 2.54 | 5.65E-15 |
| 914 | Gm42863 | 2.52 | 5.85E-15 |
| 915 | Gpr84 | 2.41 | 6.26E-15 |
| 916 | Gm20661 | 5.35 | 6.53E-15 |
| 917 | Osm | 4.26 | 7.47E-15 |
| 918 | Plk3 | 2.31 | 7.74E-15 |
| 919 | Gm8075 | 6.08 | 7.90E-15 |
| 920 | Gm43305 | 2.22 | 7.98E-15 |
| 921 | Slpi | 2.95 | 8.42E-15 |
| 922 | Gm19719 | 3.66 | 9.27E-15 |
| 923 | Slc45a3 | 2.85 | 9.44E-15 |
| 924 | 9130230L23Rik | 2.84 | 9.46E-15 |
| 925 | Gdf9 | 3.15 | 9.66E-15 |
| 926 | Zc3hav1l | 2.22 | 9.77E-15 |
| 927 | Nudt17 | 2.51 | 1.13E-14 |
| 928 | Il4i1 | 2.41 | 1.15E-14 |
| 929 | Gm16712 | 2.89 | 1.20E-14 |
| 930 | Gm44017 | 2.67 | 1.22E-14 |
| 931 | Chil3 | 5.38 | 1.23E-14 |
| 932 | Mrgpra2a | 5.31 | 1.26E-14 |
| 933 | Apol11b | 3.47 | 1.34E-14 |
| 934 | Slamf7 | 3.12 | 1.35E-14 |
| 935 | Sapcd2 | 3.45 | 1.51E-14 |
| 936 | Efna3 | 2.69 | 1.51E-14 |
| 937 | Gm38171 | 3.25 | 1.57E-14 |
| 938 | Cxcl1 | 3.80 | 1.60E-14 |
| 939 | Gm4316 | 2.32 | 2.18E-14 |
| 940 | Reg3g | 10.15 | 2.26E-14 |
| 941 | Gpr85 | 4.74 | 2.77E-14 |
| 942 | Gm31026 | 2.02 | 2.80E-14 |
| 943 | A3galt2 | 4.12 | 3.42E-14 |
| 944 | Gm30414 | 2.30 | 3.74E-14 |
| 945 | Gm15758 | 2.64 | 4.60E-14 |
| 946 | Gm28286 | 4.16 | 4.72E-14 |
| 947 | Lpar2 | 2.55 | 6.36E-14 |
| 948 | Olfr1260 | 2.30 | 6.45E-14 |
| 949 | Ccnl1 | 2.16 | 6.48E-14 |
| 950 | Hspa2 | 2.01 | 6.52E-14 |
| 951 | Gm21962 | 2.04 | 6.74E-14 |
| 952 | Smim3 | 2.12 | 7.37E-14 |
| 953 | 1190001M18Rik | 3.96 | 8.26E-14 |
| 954 | Rasa4 | 2.27 | 8.86E-14 |
| 955 | Vmn2r95 | 2.13 | 9.08E-14 |
| 956 | Gpr31b | 5.16 | 9.53E-14 |
| 957 | Gm28893 | 2.40 | 9.60E-14 |
| 958 | Gm30097 | 3.21 | 1.07E-13 |
| 959 | Crybb3 | 3.30 | 1.10E-13 |
| 960 | Gm43462 | 2.23 | 1.13E-13 |
| 961 | Mmp25 | 2.43 | 1.45E-13 |
| 962 | Gm43196 | 2.58 | 1.46E-13 |
| 963 | Gm16907 | 2.30 | 1.47E-13 |
| 964 | Inka1 | 2.77 | 1.61E-13 |
| 965 | Piwil2 | 2.57 | 1.64E-13 |
| 966 | Gm47518 | 2.54 | 1.70E-13 |
| 967 | Nrg1 | 3.04 | 1.74E-13 |
| 968 | Gm37139 | 2.34 | 1.74E-13 |
| 969 | Hk3 | 2.19 | 1.79E-13 |
| 970 | Gm6904 | 2.19 | 1.98E-13 |
| 971 | Ccl12 | 5.52 | 2.04E-13 |
| 972 | Gm28913 | 2.67 | 2.04E-13 |
| 973 | Armc12 | 5.39 | 2.10E-13 |
| 974 | Gm19684 | 2.65 | 2.18E-13 |
| 975 | Dnhd1 | 2.13 | 2.21E-13 |
| 976 | Chrna9 | 4.83 | 2.69E-13 |
| 977 | Gm26841 | 2.91 | 3.01E-13 |
| 978 | Fam3b | 4.73 | 3.23E-13 |
| 979 | Gm36283 | 3.39 | 3.29E-13 |
| 980 | Kcnh4 | 4.12 | 3.87E-13 |
| 981 | Hspbap1 | 2.11 | 3.94E-13 |
| 982 | 9330121J05Rik | 2.30 | 4.09E-13 |
| 983 | Gm11795 | 4.72 | 4.37E-13 |
| 984 | Ccdc88b | 2.32 | 4.83E-13 |
| 985 | Ddn | 5.27 | 4.85E-13 |
| 986 | 4930579D07Rik | 5.06 | 5.07E-13 |
| 987 | Atp11a | 2.55 | 5.30E-13 |
| 988 | Ifi206 | 3.74 | 5.32E-13 |
| 989 | Gm47248 | 2.98 | 5.35E-13 |
| 990 | Gja4 | 2.35 | 5.41E-13 |
| 991 | Gm15987 | 3.20 | 5.53E-13 |
| 992 | Tssk3 | 4.33 | 5.83E-13 |
| 993 | Adamts6 | 2.46 | 6.76E-13 |
| 994 | Rab44 | 3.70 | 6.81E-13 |
| 995 | Padi4 | 3.81 | 7.01E-13 |
| 996 | Map3k6 | 2.26 | 9.04E-13 |
| 997 | Ppp1r32 | 2.60 | 9.14E-13 |
| 998 | B630019K06Rik | 2.42 | 9.91E-13 |
| 999 | Ctla2b | 2.12 | 9.97E-13 |
| 1000 | Gm44867 | 3.69 | 1.16E-12 |
| 1376 | Gm19817 | 3.87 | 1.15E-05 |
| 1377 | Majin | 2.53 | 1.15E-05 |
| 1378 | Gm33201 | 2.70 | 1.17E-05 |
| 1379 | 2010110E17Rik | 2.60 | 1.19E-05 |
| 1380 | Ccdc33 | 5.07 | 1.20E-05 |
| 1381 | F13a1 | 2.72 | 1.22E-05 |
| 1382 | Gvin1 | 2.90 | 1.24E-05 |
| 1383 | Gzmm | 2.06 | 1.37E-05 |
| 1384 | Vdr | 3.81 | 1.52E-05 |
| 1385 | Lox | 3.15 | 1.54E-05 |
| 1386 | Rasal1 | 2.44 | 1.55E-05 |
| 1387 | 4833427F10Rik | 3.32 | 1.58E-05 |
| 1388 | 4833407H14Rik | 2.98 | 1.76E-05 |
| 1389 | Fcrlb | 2.53 | 1.83E-05 |
| 1390 | 1700022H01Rik | 2.24 | 1.88E-05 |
| 1391 | A730090N16Rik | 2.97 | 1.91E-05 |
| 1392 | Gm16224 | 3.67 | 1.98E-05 |
| 1393 | Gm43040 | 2.87 | 1.98E-05 |
| 1394 | Gm13874 | 5.09 | 2.00E-05 |
| 1395 | Pdpn | 2.49 | 2.06E-05 |
| 1396 | Gm49590 | 2.20 | 2.06E-05 |
| 1397 | Tctex1d4 | 3.03 | 2.17E-05 |
| 1398 | Gm19439 | 2.77 | 2.25E-05 |
| 1399 | Gm48302 | 2.73 | 2.44E-05 |
| 1400 | Gm28134 | 6.39 | 2.44E-05 |
| 1401 | Gm37574 | 6.39 | 2.44E-05 |
| 1402 | BC048559 | 2.25 | 2.46E-05 |
| 1403 | Ehf | 2.24 | 2.52E-05 |
| 1404 | Gm15232 | 3.03 | 2.58E-05 |
| 1405 | Acot2 | 2.42 | 2.59E-05 |
| 1406 | AC125101.1 | 2.16 | 2.70E-05 |
| 1407 | Klra17 | 3.18 | 2.79E-05 |
| 1408 | Gm15856 | 2.78 | 3.03E-05 |
| 1409 | Slc34a2 | 2.35 | 3.17E-05 |
| 1410 | Sox1 | 2.58 | 3.17E-05 |
| 1411 | Ly6c2 | 2.28 | 3.20E-05 |
| 1412 | Gm43581 | 2.45 | 3.45E-05 |
| 1413 | B230311B06Rik | 4.73 | 3.45E-05 |
| 1414 | Ppfia3 | 3.64 | 3.49E-05 |
| 1415 | Gm28694 | 3.23 | 3.75E-05 |
| 1416 | Tldc2 | 3.60 | 3.88E-05 |
| 1417 | Ghrl | 2.61 | 4.02E-05 |
| 1418 | Pdk4 | 3.23 | 4.02E-05 |
| 1419 | Col25a1 | 2.20 | 4.18E-05 |
| 1420 | Il19 | 4.92 | 4.27E-05 |
| 1421 | Mapk13 | 2.44 | 4.37E-05 |
| 1422 | Otoa | 3.88 | 4.42E-05 |
| 1423 | Gm45912 | 3.11 | 4.61E-05 |
| 1424 | Gm21847 | 3.06 | 4.80E-05 |
| 1425 | Gm40765 | 2.18 | 5.10E-05 |
| 1426 | Gpha2 | 3.61 | 5.19E-05 |
| 1427 | Parm1 | 3.70 | 5.25E-05 |
| 1428 | Tmem119 | 2.62 | 5.56E-05 |
| 1429 | AU022793 | 2.93 | 6.17E-05 |
| 1430 | Gm12212 | 2.52 | 6.30E-05 |
| 1431 | Melk | 2.99 | 6.91E-05 |
| 1432 | Gm20412 | 3.07 | 7.29E-05 |
| 1433 | Gm29506 | 2.04 | 7.35E-05 |
| 1434 | Sh2d4b | 3.03 | 7.68E-05 |
| 1435 | Olfr96 | 3.37 | 7.78E-05 |
| 1436 | Trim38 | 3.20 | 8.11E-05 |
| 1437 | Gm19345 | 2.22 | 8.77E-05 |
| 1438 | H2-M2 | 3.09 | 8.90E-05 |
| 1439 | Pbx4 | 2.39 | 8.91E-05 |
| 1440 | Gm17103 | 2.30 | 9.32E-05 |
| 1441 | Gm48336 | 2.11 | 9.35E-05 |
| 1442 | Slc26a9 | 4.61 | 9.38E-05 |
| 1443 | Abra | 3.79 | 9.58E-05 |
| 1444 | Klk1 | 2.06 | 9.68E-05 |
| 1445 | Slc5a5 | 3.13 | 9.73E-05 |
| 1446 | Gm37964 | 2.51 | 9.96E-05 |
| 1447 | Tacstd2 | 4.92 | 1.03E-04 |
| 1448 | Fam71d | 2.29 | 1.04E-04 |
| 1449 | Tg | 3.60 | 1.04E-04 |
| 1450 | F2rl3 | 2.62 | 1.07E-04 |
| 1451 | Gm44981 | 2.41 | 1.09E-04 |
| 1452 | 3110099E03Rik | 2.20 | 1.09E-04 |
| 1453 | Gm3248 | 3.09 | 1.15E-04 |
| 1454 | Dpep3 | 4.95 | 1.15E-04 |
| 1455 | Gm12694 | 3.47 | 1.19E-04 |
| 1456 | Il17c | 3.04 | 1.24E-04 |
| 1457 | Gm49751 | 4.66 | 1.27E-04 |
| 1458 | Gm47416 | 2.19 | 1.29E-04 |
| 1459 | Olfr571 | 2.87 | 1.30E-04 |
| 1460 | Gp9 | 2.20 | 1.39E-04 |
| 1461 | Npas2 | 2.24 | 1.40E-04 |
| 1462 | Fam131b | 3.22 | 1.46E-04 |
| 1463 | Fhdc1 | 2.04 | 1.47E-04 |
| 1464 | Klf5 | 3.41 | 1.48E-04 |
| 1465 | Gm43198 | 4.98 | 1.51E-04 |
| 1466 | Egfros | 2.00 | 1.52E-04 |
| 1467 | Tmem196 | 2.02 | 1.56E-04 |
| 1468 | Dclk1 | 2.89 | 1.57E-04 |
| 1469 | Gm48662 | 2.69 | 1.80E-04 |
| 1470 | Fam205c | 2.83 | 1.83E-04 |
| 1471 | Gm5475 | 4.36 | 1.84E-04 |
| 1472 | 1600002D24Rik | 4.28 | 1.86E-04 |
| 1473 | Speer4c | 4.65 | 2.02E-04 |
| 1474 | Slit3 | 4.68 | 2.05E-04 |
| 1475 | Gm20394 | 2.68 | 2.13E-04 |
| 1476 | Gm28499 | 4.48 | 2.14E-04 |
| 1477 | Dppa3 | 2.87 | 2.15E-04 |
| 1478 | Gm41293 | 2.58 | 2.37E-04 |
| 1479 | Rhov | 2.46 | 2.39E-04 |
| 1480 | Gm45897 | 2.69 | 2.48E-04 |
| 1481 | Tnn | 3.10 | 2.55E-04 |
| 1482 | Trp63 | 2.79 | 2.56E-04 |
| 1483 | Gm4779 | 5.77 | 2.59E-04 |
| 1484 | Gm29585 | 2.20 | 2.62E-04 |
| 1485 | Gm44785 | 5.63 | 2.62E-04 |
| 1486 | Vmn1r83 | 2.13 | 2.74E-04 |
| 1487 | Stfa3 | 4.88 | 2.84E-04 |
| 1488 | Fndc3c1 | 3.53 | 2.87E-04 |
| 1489 | Gm45151 | 3.14 | 2.90E-04 |
| 1490 | Gm40634 | 2.03 | 2.91E-04 |
| 1491 | Nrip3 | 2.11 | 3.08E-04 |
| 1492 | Gm38211 | 2.02 | 3.29E-04 |
| 1493 | Gm47941 | 3.36 | 3.47E-04 |
| 1494 | Il23r | 4.76 | 3.50E-04 |
| 1495 | E330023G01Rik | 2.86 | 3.59E-04 |
| 1496 | Elfn1 | 2.36 | 3.62E-04 |
| 1497 | Xcl1 | 2.72 | 3.71E-04 |
| 1498 | Gm48719 | 4.30 | 3.78E-04 |
| 1499 | 4930594M17Rik | 5.25 | 3.89E-04 |
| 1500 | Gm13293 | 3.88 | 3.90E-04 |

Table S10. Genes differentially expressed after LPS treatment between female and male
P. leucopus in blood and spleen

Blood LPS

| accession | gene | blood LPS | Fold change | FDR p-value |
|---|---|---|---|---|
| XM_028888689.1 | Zzef1 | 1.7 | 1042.8 | 2.4E-03 |
| XM_028854130.1 | Kdm6a | 3.3 | 1021.5 | 1.6E-03 |
| XM_028877208.1 | Jmjd1c | 2.1 | 1017.3 | 2.6E-03 |
| XR_003735732.1 | Adipor2 | 3.8 | 826.3 | 2.0E-02 |
| XR_003736827.1 | Xist (LOC114708611) | 22.5 | 627.6 | 0.0E+00 |
| XM_028872572.1 | Septin7 | 4.6 | 622.6 | 1.0E-02 |
| XM_028856218.1 | LOC114682368 | 2.2 | 594.8 | 1.0E-02 |
| XM_028857194.1 | Ubr5 | 1.1 | 559.3 | 1.0E-02 |
| XM_028875728.1 | Mbtps1 | 2.1 | 514.6 | 2.0E-02 |
| XM_028880876.1 | Ankrd11 | 0.9 | 487.3 | 2.0E-02 |
| XM_028878248.1 | Mtmr4 | 1.2 | 477.1 | 4.0E-02 |
| XM_028885812.1 | Pitpnm2 | 1.1 | 469.6 | 2.0E-02 |
| XM_028873422.1 | Rasgrp3 | 1.5 | 455.1 | 3.0E-02 |
| XM_028893826.1 | Tomm40 | 5.0 | 431.8 | 2.0E-02 |
| XM_028859573.1 | Sipa1l3 | 1.0 | 430.1 | 3.0E-02 |
| XM_028853918.1 | Tsc2 | 1.3 | 400.1 | 3.0E-02 |
| XM_028892301.1 | Pex5 | 2.2 | 376.6 | 2.0E-02 |
| XM_028878946.1 | Ambra1 | 1.3 | 363.3 | 3.0E-02 |
| XM_028867184.1 | Golim4 | 1.4 | 342.2 | 4.0E-02 |
| XM_028889988.1 | Extl3 | 1.0 | 340.8 | 3.0E-02 |
| XM_028869682.1 | Heatr5a | 0.8 | 328.1 | 4.0E-02 |
| XM_028855906.1 | Kat5 | 2.3 | 301.1 | 4.0E-02 |
| XM_028875315.1 | Fam43b | 13.7 | 222.5 | 6.0E-05 |
| XM_028890997.1 | Tns3 | 2.0 | 212.4 | 2.4E-03 |
| XM_028882900.1 | Myof | 1.4 | 117.8 | 1.0E-02 |
| XM_028860252.1 | Dffb | 2.3 | 108.6 | 2.0E-02 |
| XM_028881051.1 | Clcn3 | 1.2 | 76.2 | 3.0E-02 |
| XM_028864513.1 | Ythdf3 | 1.3 | 71.4 | 7.2E-03 |
| XM_028874886.1 | Prune2 | 0.6 | 51.7 | 3.0E-02 |
| XM_028869660.1 | Lyz2 | 68.5 | 41.1 | 1.7E-04 |
| XM_028862879.1 | LOC114688257 | 1.7 | 38.3 | 3.0E-02 |
| XM_028873188.1 | Cxcl13 | 199.1 | 22.2 | 2.0E-02 |
| XM_028890581.1 | Ccl9 | 95.4 | 20.6 | 3.0E-02 |
| XM_028894435.1 | Serpine1 | 136.1 | 17.8 | 3.0E-02 |
| XM_028870030.1 | Fn1 | 454.6 | 14.4 | 2.0E-02 |
| XM_028873448.1 | Ltbp1 | 12.0 | 13.1 | 2.0E-02 |
| XM_028857646.1 | Cd5l | 308.2 | 12.0 | 4.0E-02 |
| XM_028890578.1 | Ccl6 | 429.4 | 10.1 | 3.0E-02 |
| XM_028889222.1 | Tgm2 | 40.1 | 6.6 | 1.0E-02 |
| XM_028870103.1 | Limk1 | 14.8 | 6.3 | 3.0E-02 |
| XM_028858147.1 | LOC114683802 | 102.5 | 3.9 | 3.0E-02 |
| XM_028885765.1 | Hcar2 | 104.4 | -2.5 | 3.0E-02 |
| XM_028881875.1 | LOC114701717 | 1332.7 | -2.7 | 2.0E-02 |
| XM_028884286.1 | Rab3il1 | 45.5 | -3.0 | 4.0E-02 |
| XM_028877289.1 | Tent5c | 105.0 | -3.2 | 3.0E-02 |
| XM_028860286.1 | Saa5 | 122.8 | -3.8 | 1.7E-04 |
| XM_028860283.1 | Saa3 | 204.2 | -5.3 | 9.4E-03 |
| XM_028880143.1 | Saa1 | 63.5 | -12.9 | 1.0E-02 |
| XM_028863833.1 | LOC114689504 | 9.1 | -19.6 | 2.0E-02 |
| XM_028884798.1 | Zbtb43 | 2.9 | -29.6 | 1.0E-02 |
| XR_003733219.1 | pseudogene | 270.6 | -36.1 | 3.6E-07 |
| XM_028865773.1 | Map4k4 | 1.9 | -42.1 | 3.0E-02 |
| XM_028854653.1 | Mapk8ip3 | 1.5 | -89.9 | 3.4E-03 |
| XM_028862666.1 | Akap9 | 0.4 | -100.3 | 4.0E-02 |
| XM_028884690.1 | Golga2 | 1.6 | -131.2 | 7.6E-03 |
| XM_028891094.1 | Ap1g1 | 1.0 | -142.8 | 8.1E-03 |
| XM_028894762.1 | Rnf38 | 2.1 | -156.7 | 3.4E-03 |
| XM_028865080.1 | Ulk3 | 1.3 | -242.3 | 4.0E-02 |
| XM_028867172.1 | Glce | 0.9 | -245.6 | 4.0E-02 |
| XM_028886382.1 | Atrx | 0.3 | -253.5 | 4.0E-02 |
| XM_028892298.1 | Pex5 | 1.2 | -264.3 | 3.0E-02 |
| XM_028888004.1 | Bag6 | 0.7 | -275.9 | 3.0E-02 |
| XM_028875262.1 | Ubr4 | 0.4 | -460.2 | 8.1E-03 |
| XM_028891866.1 | Adcy6 | 0.9 | -462.8 | 8.2E-03 |
| XM_028874496.1 | Znf740 | 1.4 | -495.1 | 1.0E-02 |
| XM_028876259.1 | Gigyf2 | 1.8 | -558.2 | 6.0E-03 |
| XM_028878824.1 | Taok3 | 2.5 | -585.8 | 7.0E-03 |
| XM_028872487.1 | Yipf2 | 3.3 | -630.9 | 1.0E-02 |
| XM_028878062.1 | Esd | 8.2 | -740.1 | 3.0E-03 |
| XM_028889000.1 | Cpne1 | 2.3 | -773.5 | 4.9E-04 |
| XM_028876722.1 | D1Pas1 (LOC114698204) | 6.9 | -849.5 | 2.0E-11 |

Blood control

| accession | gene | blood control | Fold change | FDR p-value |
|---|---|---|---|---|
| XM_028874737.1 | Prg4 | 42.9 | 6804.8 | 3.6E-04 |
| XM_028884676.1 | Prrc2b | 5.0 | 2934.7 | 1.0E-03 |
| XM_028876680.1 | Arel1 | 4.4 | 1318.8 | 1.0E-02 |
| XM_028863820.1 | Iars | 4.9 | 1185.8 | 2.0E-02 |
| XM_028879137.1 | Fubp1 | 2.8 | 1081.5 | 2.0E-02 |
| XM_028859872.1 | Hnrnpul1 | 5.1 | 988.0 | 4.0E-02 |
| XR_003736827.1 | Xist (LOC114708611) | 36.9 | 981.3 | 0.0E+00 |
| XM_028882900.1 | Myof | 2.7 | 939.4 | 4.0E-02 |
| XM_028886553.1 | Trappc8 | 3.0 | 881.9 | 4.0E-02 |
| XM_028887862.1 | Itgb1 | 3.8 | 818.5 | 3.0E-02 |
| XM_028876596.1 | Zc3h11a | 2.9 | 757.3 | 4.0E-02 |
| XM_028891460.1 | Klhl17 | 4.7 | 744.1 | 4.0E-02 |
| XM_028875315.1 | Fam43b | 5.8 | 111.7 | 1.0E-02 |
| XM_028872367.1 | Msr1 | 5.0 | 93.5 | 2.0E-02 |
| XR_003736344.1 | LOC114705024 | 10.6 | 72.6 | 4.0E-02 |
| XM_028861029.1 | H2-DMa | 14.1 | 60.6 | 4.0E-02 |
| XM_028867331.1 | Lrrc8b | 2.0 | 57.7 | 4.0E-02 |
| XM_028868565.1 | Ptgis | 8.0 | 43.5 | 1.0E-02 |
| XM_028882356.1 | Evc | 5.0 | 41.1 | 2.0E-02 |
| XM_028855524.1 | Rnf41 | 4.4 | 34.0 | 2.0E-02 |
| XM_028889747.1 | Gga3 | 7.2 | 26.0 | 4.2E-03 |
| XM_028874738.1 | Prg4 | 30.3 | 20.1 | 2.0E-02 |
| XM_028888879.1 | Vsig4 | 36.9 | 18.2 | 2.8E-04 |
| XM_028891607.1 | Serpinb2 | 38.3 | 16.4 | 9.2E-04 |
| XM_028862632.1 | LOC114688009 | 30.4 | 12.6 | 3.3E-03 |
| XM_028870030.1 | Fn1 | 238.7 | 8.0 | 2.0E-02 |
| XM_028863631.1 | LOC114689267 | 75.6 | -4.3 | 4.0E-02 |
| XM_028856593.1 | Fech | 484.7 | -5.3 | 3.0E-02 |
| XM_028855654.1 | Hectd4 | 5.5 | -5.4 | 3.3E-03 |
| XM_028889108.1 | Numa1 | 6.6 | -35.2 | 1.0E-03 |
| XM_028882612.1 | Gpatch8 | 3.6 | -73.9 | 1.0E-02 |
| XM_028870949.1 | Hsf1 | 3.5 | -99.9 | 2.0E-02 |
| XM_028861918.1 | Nsdhl | 1.9 | -235.8 | 3.3E-03 |
| XM_028865786.1 | Map4k4 | 3.4 | -905.0 | 4.0E-02 |
| XM_028856461.1 | Tbl1xr1 | 3.0 | -1381.1 | 1.0E-02 |
| XM_028878248.1 | Mtmr4 | 3.7 | -1475.2 | 2.0E-02 |
| XM_028878824.1 | Taok3 | 7.1 | -1677.1 | 1.0E-02 |
| XM_028876722.1 | D1Pas1 (LOC114698204) | 7.2 | -1762.6 | 3.3E-03 |
| XM_028886469.1 | Foxj3 | 4.8 | -1768.9 | 8.0E-03 |

Spleen LPS

| accession | gene | spleen LPS | Fold change | FDR p-value |
|---|---|---|---|---|
| XR_003735732.1 | Adipor2 | 3.1 | 4619.9 | 5.6E-05 |
| XM_028858810.1 | Tfip11 | 3.3 | 3696.3 | 1.1E-05 |
| XM_028857773.1 | Lpar1 | 1.1 | 1300.6 | 3.3E-04 |
| XM_028854407.1 | LOC114681120 | 1.2 | 836.7 | 9.7E-03 |
| XR_003736827.1 | Xist (LOC114708611) | 45.1 | 590.3 | 0.0E+00 |
| XM_028885014.1 | Tff2 | 9.5 | 14.1 | 1.0E-02 |
| XM_028880146.1 | Otog | 1.9 | 6.1 | 1.0E-02 |
| XM_028861007.1 | Gp1ba | 43.1 | 2.9 | 3.5E-03 |
| XM_028888744.1 | Fbxo39 | 11.6 | 2.6 | 2.0E-02 |
| XM_028890601.1 | Mbnl1 | 10.2 | 2.0 | 2.0E-02 |
| XM_028884620.1 | Rgs2 | 36.4 | -1.7 | 4.0E-02 |
| XM_028873432.1 | Picalm | 41.7 | -1.8 | 4.0E-02 |
| XM_028893766.1 | Anxa1 | 52.3 | -1.8 | 6.4E-03 |
| XM_028892582.1 | Birc5 | 31.2 | -2.7 | 2.0E-02 |
| XM_028859394.1 | Glrx5 | 100.2 | -2.9 | 4.0E-02 |
| XM_028867842.1 | Cdr2 | 29.6 | -3.1 | 3.0E-02 |
| XM_028884304.1 | Ms4a3 | 16.0 | -3.2 | 1.0E-02 |
| XM_028877913.1 | Cat | 111.1 | -3.3 | 3.0E-02 |
| XM_028875917.1 | Ddx3x | 29.7 | -3.3 | 0.0E+00 |
| XM_028854261.1 | Ranbp10 | 64.2 | -3.4 | 4.0E-02 |
| XM_028882602.1 | Slc25a39 | 151.3 | -3.6 | 1.0E-02 |
| XM_028883198.1 | LOC114702707 | 52.9 | -3.6 | 2.0E-02 |
| XM_028877289.1 | Tent5c | 31.1 | -3.7 | 4.0E-02 |
| XM_028889819.1 | Urod | 75.5 | -3.8 | 2.0E-02 |
| XM_028876534.1 | Tmcc2 | 14.5 | -3.9 | 3.0E-02 |
| XM_028883976.1 | Fam214b | 20.4 | -4.0 | 2.0E-02 |
| XM_028889307.1 | Mkrn1 | 177.3 | -4.0 | 2.0E-02 |
| XM_028869180.1 | Ppp6r2 | 1.6 | -4.2 | 4.0E-02 |
| XM_028856593.1 | Fech | 56.5 | -4.4 | 3.0E-02 |
| XM_028869007.1 | Phospho1 | 13.0 | -4.6 | 2.0E-02 |
| XM_028892812.1 | LOC114709142 | 8.2 | -4.6 | 4.0E-02 |
| XM_028869008.1 | Phospho1 | 20.8 | -4.6 | 1.0E-02 |
| XM_028887625.1 | Epb41 | 49.0 | -4.8 | 2.0E-02 |
| XM_028854967.1 | Prxl2a | 54.2 | -4.9 | 1.0E-02 |
| XM_028890537.1 | Dhrs11 | 55.4 | -5.1 | 2.0E-02 |
| XM_028857150.1 | LOC114682995 | 37.4 | -5.4 | 3.0E-02 |
| XM_028889268.1 | Snca | 56.3 | -5.5 | 3.5E-03 |
| XM_028863978.1 | Syn2 | 3.8 | -5.7 | 4.0E-02 |
| XM_028857149.1 | Trim58 | 13.0 | -6.4 | 4.0E-02 |
| XM_028866142.1 | Bpgm | 528.1 | -6.5 | 8.8E-04 |
| XM_028886826.1 | Smim33 | 4.3 | -7.0 | 4.0E-02 |
| XM_028854133.1 | Kdm6a | 5.1 | -7.1 | 3.0E-02 |
| XM_028891155.1 | Btrc | 1.8 | -7.2 | 8.9E-03 |
| XM_028889997.1 | LOC114707298 | 51.0 | -7.7 | 8.1E-04 |
| XM_028874127.1 | Mff | 4.3 | -10.2 | 4.0E-02 |
| XM_028878062.1 | Esd | 8.4 | -10.5 | 2.0E-02 |
| XM_028887202.1 | Ccnd3 | 5.9 | -11.6 | 4.0E-02 |
| XM_028874225.1 | Snrpd3 | 21.6 | -12.2 | 1.9E-03 |
| XR_003733219.1 | pseudogene | 40.3 | -12.9 | 1.1E-04 |
| XM_028876271.1 | Neu2 | 4.4 | -15.9 | 4.0E-02 |
| XM_028863833.1 | LOC114689504 | 39.0 | -16.7 | 4.7E-06 |
| XM_028883980.1 | Fam214b | 2.5 | -37.5 | 2.0E-02 |
| XM_028863966.1 | Hrh1 | 1.5 | -42.7 | 3.8E-06 |
| XM_028876722.1 | D1Pas1 (LOC114698204) | 5.8 | -328.3 | 0.0E+00 |
| XM_028873251.1 | Sh3kbp1 | 2.2 | -3157.8 | 1.1E-04 |
| XM_028856570.1 | Cit | 1.1 | -3633.2 | 1.1E-05 |

**Table S11. Enrichment GO terms and included genes for Figure 4**

***P. leucopus***

| | Group | regulation of neuron differentiation | negative regulation of cellular extravasation | negative regulation of epidermal cell differentiation | positive regulation of ERK1 and ERK2 cascade | regulation of MAPK cascade |
|---|---|---|---|---|---|---|
| Overlap | Blood_Down | 2/84 | 1/6 | 1/7 | 2/201 | 2/203 |
| P.value | | 1.32E-03 | 3.89E-03 | 4.54E-03 | 7.29E-03 | 7.43E-03 |
| Adjusted.P.value | | 1.00E+00 | 1.00E+00 | 1.00E+00 | 1.00E+00 | 1.00E+00 |
| Odds.Ratio | | 36.63 | 256.41 | 219.78 | 15.31 | 15.16 |
| Combined.Score | | 242.88 | 1422.64 | 1185.59 | 75.34 | 74.31 |
| Genes | | NCS1<br>DTX1 | CCL25 | GDF3 | CCL25<br>EPHA10 | EPHA10<br>GDF3 |

| | Group | indolalkylamine catabolic process | tryptophan catabolic process | tryptophan metabolic process | neutrophil degranulation | neutrophil activation involved in immune response |
|---|---|---|---|---|---|---|
| Overlap | Spleen_Up | 2/9 | 2/10 | 2/10 | 7/479 | 7/483 |
| P.value | | 3.14E-04 | 3.92E-04 | 3.92E-04 | 5.58E-04 | 5.86E-04 |
| Adjusted.P.value | | 1.00E+00 | 1.00E+00 | 6.67E-01 | 7.12E-01 | 5.98E-01 |
| Odds.Ratio | | 74.07 | 66.67 | 66.67 | 4.87 | 4.83 |
| Combined.Score | | 597.41 | 522.93 | 522.93 | 36.49 | 35.95 |
| Genes | | IDO1<br>IDO2 | IDO1<br>IDO2 | IDO1<br>IDO2 | CRISPLD2<br>LRG1<br>MMP8<br>MS4A3<br>OLFM4<br>OLR1<br>SLPI | CRISPLD2<br>LRG1<br>MMP8<br>MS4A3<br>OLFM4<br>OLR1<br>SLPI |

| | Group | neutrophil mediated immunity | neutrophil degranulation | neutrophil activation involved in immune response | inflammatory response | cellular response to cytokine stimulus |
|---|---|---|---|---|---|---|
| Overlap | Blood_Up | 28/487 | 26/479 | 26/483 | 19/252 | 24/456 |
| P.value | | 1.21E-11 | 2.40E-10 | 2.88E-10 | 3.24E-10 | 2.26E-09 |
| Adjusted.P.value | | 6.18E-08 | 6.14E-07 | 4.89E-07 | 4.14E-07 | 2.31E-06 |
| Odds.Ratio | | 4.69 | 4.43 | 4.39 | 6.15 | 4.30 |
| Combined.Score | | 117.98 | 98.14 | 96.54 | 134.48 | 85.53 |
| Genes | | ADGRG3<br>ALOX5<br>ANPEP<br>ATP8B4<br>C5AR1<br>CD33<br>CHIT1<br>CRISPLD2<br>CXCL5<br>CXCR2<br>FGR<br>HK3<br>IL6<br>LRG1<br>MCEMP1<br>MGAM<br>MMP9<br>MS4A3<br>NFAM1<br>OLR1<br>PGLYRP1<br>PLAUR<br>PNP<br>PRDX6<br>PTAFR<br>RETN<br>SLC11A1<br>SLPI | ADGRG3<br>ALOX5<br>ANPEP<br>ATP8B4<br>C5AR1<br>CD33<br>CHIT1<br>CRISPLD2<br>CXCR2<br>FGR<br>HK3<br>LRG1<br>MCEMP1<br>MGAM<br>MMP9<br>MS4A3<br>NFAM1<br>OLR1<br>PGLYRP1<br>PLAUR<br>PNP<br>PRDX6<br>PTAFR<br>RETN<br>SLC11A1<br>SLPI | ADGRG3<br>ALOX5<br>ANPEP<br>ATP8B4<br>C5AR1<br>CD33<br>CHIT1<br>CRISPLD2<br>CXCR2<br>FGR<br>HK3<br>LRG1<br>MCEMP1<br>MGAM<br>MMP9<br>MS4A3<br>NFAM1<br>OLR1<br>PGLYRP1<br>PLAUR<br>PNP<br>PRDX6<br>PTAFR<br>RETN<br>SLC11A1<br>SLPI | C5AR1<br>C5AR2<br>CCL17<br>CCL19<br>CEBPB<br>CXCL5<br>CXCR2<br>FCGR2B<br>GPR33<br>IL1B<br>IL6<br>NFAM1<br>NLRC4<br>PRDX5<br>SCUBE1<br>SELE<br>SLC11A1<br>TACR1<br>TLR4 | ALOX5<br>CCL17<br>CCL19<br>CEBPD<br>CSF2<br>CSF2RB<br>CSF3R<br>DAB2IP<br>DCSTAMP<br>HAS2<br>HMOX1<br>HYAL2<br>IL13RA1<br>IL1B<br>IL1R1<br>IL20RA<br>IL22RA2<br>IL6<br>MMP9<br>OSM<br>PTAFR<br>RTN4RL2<br>SBNO2<br>WNT5A |

| | Group | ion transport | response to retinoic acid | positive regulation of protein kinase C signaling | cellular response to retinoic acid | ion transmembrane transport |
|---|---|---|---|---|---|---|
| Overlap | Liver_Down | 8/289 | 4/57 | 2/7 | 3/42 | 5/219 |
| P.value | | 6.41E-07 | 1.34E-05 | 1.33E-04 | 1.67E-04 | 2.34E-04 |
| Adjusted.P.value | | 3.27E-03 | 3.42E-02 | 2.26E-01 | 2.13E-01 | 2.39E-01 |
| Odds.Ratio | | 10.86 | 27.52 | 112.04 | 28.01 | 8.95 |
| Combined.Score | | 154.80 | 308.77 | 1000.20 | 243.60 | 74.83 |
| Genes | | ATP2C2<br>CHRNB4<br>GABRA6<br>GLRB<br>SGK2<br>SLC15A2<br>SLC24A1<br>SLC26A3 | ASCL1<br>FZD10<br>WNT11<br>WNT5A | WNT11<br>WNT5A | FZD10<br>WNT11<br>WNT5A | ATP2C2<br>CHRNB4<br>GABRA6<br>GLRB<br>SGK2 |

| | Group | neuroepithelial cell differentiation | negative regulation of neurogenesis | regulation of neuron differentiation | skeletal system development | positive regulation of lipid localization |
|---|---|---|---|---|---|---|
| Overlap | Spleen_Down | 3/10 | 4/33 | 4/84 | 5/146 | 2/10 |
| P.value | | 1.94E-05 | 3.25E-05 | 1.23E-03 | 1.33E-03 | 1.33E-03 |
| Adjusted.P.value | | 9.90E-02 | 8.29E-02 | 1.00E+00 | 1.00E+00 | 1.00E+00 |
| Odds.Ratio | | 54.05 | 21.84 | 8.58 | 6.17 | 36.04 |
| Combined.Score | | 586.48 | 225.72 | 57.47 | 40.85 | 238.54 |
| Genes | | FGF8<br>HES5<br>WNT11 | BMP7<br>HES5<br>SOX2<br>WNT7A | BMP7<br>HES5<br>MMD2<br>SOX2 | BMP7<br>FGF8<br>HAPLN4<br>HES5<br>WNT11 | APOB<br>VSTM2A |

| | Group | cytokine-mediated signaling pathway | phosphatidylserine acyl-chain remodeling | regulation of interleukin-5 secretion | regulation of interleukin-13 secretion | phosphatidylserine metabolic process |
|---|---|---|---|---|---|---|
| Overlap | Liver_Up | 13/633 | 3/21 | 2/6 | 2/6 | 3/27 |
| P.value | | 3.36E-04 | 3.58E-04 | 6.57E-04 | 6.57E-04 | 7.65E-04 |
| Adjusted.P.value | | 1.00E+00 | 9.14E-01 | 1.00E+00 | 8.38E-01 | 7.80E-01 |
| Odds.Ratio | | 3.07 | 21.32 | 49.75 | 49.75 | 16.58 |
| Combined.Score | | 24.52 | 169.19 | 364.59 | 364.59 | 119.01 |
| Genes | | GATA3<br>IFNGR2<br>IL10RB<br>IL17RB<br>IL1RL1<br>IL1RL2<br>IRF8<br>LTA<br>MYC<br>SOD2<br>STAT4<br>TNFRSF18<br>TNFRSF21 | PLA2G2A<br>PLA2G5<br>PLA1A | GATA3<br>TNFRSF21 | GATA3<br>TNFRSF21 | PLA2G2A<br>PLA2G5<br>PLA1A |

Table S11. Enrichment GO terms and included genes for Figure 4

| ***M. musculus*** | Group | negative regulation of blood coagulation | regulation of fibrinolysis | negative regulation of fibrinolysis | regulation of blood coagulation | positive regulation of blood coagulation |
|---|---|---|---|---|---|---|
| Overlap | Blood_Down | 12/36 | 7/13 | 6/9 | 8/30 | 7/19 |
| P.value | | 1.43E-17 | 1.51E-12 | 1.08E-11 | 3.21E-11 | 4.26E-11 |
| Adjusted.P.value | | 7.30E-14 | 3.84E-09 | 1.84E-08 | 4.10E-08 | 4.35E-08 |
| Odds.Ratio | | 45.98 | 74.27 | 91.95 | 36.78 | 50.82 |
| Combined.Score | | 1783.28 | 2021.81 | 2322.02 | 888.69 | 1213.44 |
| Genes | | APOH | APOH | APOH | APOH | APOH |
| | | F12 | F12 | F2 | F12 | F12 |
| | | F2 | F2 | HRG | F2 | F2 |
| | | FGA | HRG | PLG | HRG | HRG |
| | | FGG | PLG | SERPINF2 | KNG1 | PLG |
| | | HRG | SERPINF2 | THBD | PROC | SERPINF2 |
| | | KNG1 | THBD | | THBD | THBD |
| | | PLG | | | VTN | |
| | | PROC | | | | |
| | | SERPINF2 | | | | |
| | | THBD | | | | |
| | | VTN | | | | |

| | Group | cytokine-mediated signaling pathway | positive regulation of myeloid leukocyte differentiation | regulation of granulocyte macrophage CSF production | positive regulation of osteoclast differentiation | regulation of cellular pH |
|---|---|---|---|---|---|---|
| Overlap | Spleen_Up | 17/633 | 4/32 | 3/13 | 3/14 | 4/38 |
| P.value | | 9.46E-07 | 5.65E-05 | 7.66E-05 | 9.70E-05 | 1.12E-04 |
| Adjusted.P.value | | 4.83E-03 | 1.44E-01 | 1.30E-01 | 1.24E-01 | 1.15E-01 |
| Odds.Ratio | | 4.07 | 18.94 | 34.97 | 32.47 | 15.95 |
| Combined.Score | | 56.44 | 185.26 | 331.37 | 300.03 | 145.03 |
| Genes | | BST2 | IL12B | IL12B | IL12B | SLC26A4 |
| | | CD70 | IL17A | IL17F | IL17A | SLC26A9 |
| | | CXCL9 | IL23A | IL23A | IL23A | SLC4A4 |
| | | FOS | LIF | | | SLC9A4 |
| | | IFI35 | | | | |
| | | IL12B | | | | |
| | | IL17A | | | | |
| | | IL17F | | | | |
| | | IL23A | | | | |
| | | LIF | | | | |
| | | MUC1 | | | | |
| | | PTGS2 | | | | |
| | | RHOU | | | | |
| | | TNFRSF9 | | | | |
| | | TNFSF15 | | | | |
| | | TNFSF9 | | | | |
| | | TRIM21 | | | | |

| | Group | cellular response to type I interferon | type I interferon signaling pathway | response to cytokine | positive regulation of cytokine-mediated signaling pathway | T-helper 1 type immune response |
|---|---|---|---|---|---|---|
| Overlap | Blood_Up | 4/65 | 4/65 | 5/138 | 3/33 | 2/7 |
| P.value | | 4.04E-05 | 4.04E-05 | 5.40E-05 | 1.25E-04 | 1.78E-04 |
| Adjusted.P.value | | 2.06E-01 | 1.03E-01 | 9.18E-02 | 1.59E-01 | 1.82E-01 |
| Odds.Ratio | | 20.86 | 20.86 | 12.28 | 30.82 | 96.85 |
| Combined.Score | | 211.02 | 211.02 | 120.69 | 276.97 | 836.22 |
| Genes | | BST2 | BST2 | BST2 | HSPA1A | EBI3 |
| | | OAS2 | OAS2 | IFNLR1 | PARP14 | IL18BP |
| | | STAT2 | STAT2 | IL18BP | ZBP1 | |
| | | XAF1 | XAF1 | TRAFD1 | | |
| | | | | XAF1 | | |

| | Group | negative regulation of androgen receptor signaling pathway | branching involved in blood vessel morphogenesis | cardiac ventricle development | spinal cord dorsal/ventral patterning | regulation of androgen receptor signaling pathway |
|---|---|---|---|---|---|---|
| Overlap | Liver_Down | 3/12 | 3/14 | 3/16 | 2/6 | 3/23 |
| P.value | | 5.11E-04 | 8.28E-04 | 1.25E-03 | 2.70E-03 | 3.68E-03 |
| Adjusted.P.value | | 1.00E+00 | 1.00E+00 | 1.00E+00 | 1.00E+00 | 1.00E+00 |
| Odds.Ratio | | 18.25 | 15.64 | 13.69 | 24.33 | 9.52 |
| Combined.Score | | 138.32 | 111.00 | 91.51 | 143.86 | 53.37 |
| Genes | | FOXH1 | DLL4 | DLL4 | DLL4 | FOXH1 |
| | | HEYL | GDF2 | HEYL | RFX4 | HEYL |
| | | NODAL | NRARP | HEY1 | | NODAL |

| | Group | skeletal muscle thin filament assembly | skeletal myofibril assembly | cardiac muscle tissue morphogenesis | cardiac myofibril assembly | myofibril assembly |
|---|---|---|---|---|---|---|
| Overlap | Spleen_Down | 3/10 | 3/12 | 4/36 | 3/17 | 4/47 |
| P.value | | 2.77E-05 | 5.03E-05 | 7.34E-05 | 1.52E-04 | 2.11E-04 |
| Adjusted.P.value | | 1.41E-01 | 1.28E-01 | 1.25E-01 | 1.94E-01 | 2.15E-01 |
| Odds.Ratio | | 48.00 | 40.00 | 17.78 | 28.24 | 13.62 |
| Combined.Score | | 503.72 | 395.89 | 169.25 | 248.23 | 115.27 |
| Genes | | ACTC1 | ACTC1 | ACTC1 | ACTC1 | ACTC1 |
| | | MYOM2 | MYOM2 | ISL1 | MYOM2 | LMOD1 |
| | | MYOM3 | MYOM3 | MYOM2 | MYOM3 | MYOM2 |
| | | | | MYOM3 | | MYOM3 |

| | Group | cytokine-mediated signaling pathway | regulation of inflammatory response | positive regulation of T cell proliferation | positive regulation of I-kappaB kinase/NF-kappaB signaling | positive regulation of intracellular signal transduction |
|---|---|---|---|---|---|---|
| Overlap | Liver_Up | 36/633 | 16/166 | 10/61 | 14/163 | 24/479 |
| P.value | | 9.64E-09 | 1.64E-07 | 2.57E-07 | 3.85E-06 | 2.51E-05 |
| Adjusted.P.value | | 4.92E-05 | 4.17E-04 | 4.38E-04 | 4.91E-03 | 2.56E-02 |
| Odds.Ratio | | 2.92 | 4.96 | 8.43 | 4.42 | 2.58 |
| Combined.Score | | 53.97 | 77.44 | 127.88 | 55.06 | 27.28 |
| Genes | | BCL2L1 | ADIPOQ | CCL19 | ADIPOQ | ADIPOQ |
| | | BIRC2 | BCL6B | CCL5 | BIRC2 | ADRB2 |
| | | CCL17 | BIRC2 | CD28 | CCL19 | BCL2L1 |
| | | CCL19 | CCL5 | CD3E | GPRC5B | BIRC2 |
| | | CCL5 | FFAR2 | IL12B | IKBKE | CCL19 |
| | | CD300LF | GHRL | IL23R | IL12B | CCL5 |
| | | CTF1 | GPRC5B | PNP | IL1A | CD28 |
| | | CXCL13 | HCK | SYK | LITAF | EREG |
| | | DUOX1 | IL12B | TNFSF9 | MYD88 | GPR37L1 |
| | | EREG | IL33 | VCAM1 | RNF31 | GPRC5B |
| | | HCK | MYD88 | | ROR1 | IKBKE |
| | | IFI35 | PIK3AP1 | | TLR2 | IL1A |
| | | IL12B | PTGS2 | | TNFSF10 | INSRR |
| | | IL17C | TLR2 | | TRIM38 | LITAF |
| | | IL17RA | TNFAIP6 | | | MYD88 |
| | | IL19 | TNIP1 | | | NRG1 |
| | | IL1A | | | | PIK3AP1 |
| | | IL1R1 | | | | RET |
| | | IL1R2 | | | | RNF31 |
| | | IL23R | | | | ROR1 |
| | | IL33 | | | | SOX9 |
| | | IRF9 | | | | THBS1 |
| | | ITGAM | | | | TNFSF10 |
| | | MUC1 | | | | TRIM38 |
| | | MYD88 | | | | |
| | | NUMBL | | | | |
| | | OAS2 | | | | |
| | | PTAFR | | | | |
| | | PTGS2 | | | | |
| | | SERPINB2 | | | | |
| | | SYK | | | | |
| | | TNFRSF9 | | | | |
| | | TNFSF14 | | | | |
| | | TNFSF9 | | | | |
| | | TRIM38 | | | | |
| | | VCAM1 | | | | |
| | | RTN4RL2 | | | | |
| | | SAMHD1 | | | | |
| | | SOCS1 | | | | |
| | | SOCS3 | | | | |
| | | STAT3 | | | | |
| | | TIMP1 | | | | |
| | | TNF | | | | |
| | | TNIP2 | | | | |

Table S11. Enrichment GO terms and included genes for Figure 4

**Shared** (No Blood_Down GO terms)

| | Group | cytokine-mediated signaling pathway | cellular response to cytokine stimulus | neutrophil degranulation | neutrophil activation involved in immune response | neutrophil mediated immunity |
|---|---|---|---|---|---|---|
| Overlap | Blood_Up | 24/633 | 19/456 | 18/479 | 18/483 | 18/487 |
| P.value | | 2.34E-16 | 9.12E-14 | 2.53E-12 | 2.90E-12 | 3.33E-12 |
| Adjusted.P.value | | 1.20E-12 | 2.33E-10 | 4.30E-09 | 3.70E-09 | 3.40E-09 |
| Odds.Ratio | | 8.62 | 9.47 | 8.54 | 8.47 | 8.40 |
| Combined.Score | | 310.13 | 284.33 | 228.07 | 225.01 | 222.00 |
| Genes | | CISH | IL10 | GCA | GCA | GCA |
| | | CSF3 | ACOD1 | ABCA13 | ABCA13 | ABCA13 |
| | | CXCL10 | CISH | ANXA3 | ANXA3 | ANXA3 |
| | | CXCL11 | CSF3 | CAMP | CAMP | CAMP |
| | | FPR1 | CXCL10 | CD14 | CD14 | CD14 |
| | | ICAM1 | FPR1 | CD177 | CD177 | CD177 |
| | | IFNG | ICAM1 | CLEC5A | CLEC5A | CLEC5A |
| | | IL10 | IL1A | FPR1 | FPR1 | FPR1 |
| | | IL1A | IL1R2 | FPR2 | FPR2 | FPR2 |
| | | IL1R2 | IL1RN | GPR84 | GPR84 | GPR84 |
| | | IL1RN | IRAK3 | LCN2 | LCN2 | LCN2 |
| | | IL27 | LCN2 | LTF | LTF | LTF |
| | | IRAK3 | MUC1 | MMP25 | MMP25 | MMP25 |
| | | LCN2 | OSMR | MMP8 | MMP8 | MMP8 |
| | | MUC1 | RHOU | OLFM4 | OLFM4 | OLFM4 |
| | | NFKBIA | SOCS1 | PTX3 | PTX3 | PTX3 |
| | | OSMR | SOCS3 | SLCO4C1 | SLCO4C1 | SLCO4C1 |
| | | RHOU | TIMP1 | TLR2 | TLR2 | TLR2 |
| | | SOCS1 | TNF | | | |
| | | SOCS3 | | | | |
| | | SOD2 | | | | |
| | | TIMP1 | | | | |
| | | TNF | | | | |
| | | TNFRSF9 | | | | |

| | Group | regulation of programmed cell death | negative regulation of cellular amide metabolic process | BMP signaling pathway | cellular response to BMP stimulus | negative regulation of translation |
|---|---|---|---|---|---|---|
| Overlap | Liver_Down | 4/268 | 2/75 | 2/81 | 2/87 | 2/88 |
| P.value | | 3.19E-04 | 3.94E-03 | 4.58E-03 | 5.26E-03 | 5.38E-03 |
| Adjusted.P.value | | 1.00E+00 | 1.00E+00 | 1.00E+00 | 1.00E+00 | 1.00E+00 |
| Odds.Ratio | | 11.94 | 21.33 | 19.75 | 18.39 | 18.18 |
| Combined.Score | | 96.11 | 118.13 | 106.41 | 96.51 | 95.01 |
| Genes | | KANK2 | GRB7 | LEFTY1 | LEFTY1 | GRB7 |
| | | GAS1 | TOB1 | TOB1 | TOB1 | TOB1 |
| | | LEFTY1 | | | | |
| | | MAP2K6 | | | | |

| | Group | G-protein coupled receptor signaling | regulation of cAMP-mediated signaling | adenylate cyclase-activating adrenergic receptor signaling pathway | adrenergic receptor signaling pathway | adenylate cyclase-activating G-protein coupled receptor signaling pathway |
|---|---|---|---|---|---|---|
| Overlap | Spleen_Down | 4/125 | 2/18 | 2/20 | 2/20 | 2/78 |
| P.value | | 2.28E-05 | 2.65E-04 | 3.28E-04 | 3.28E-04 | 4.95E-03 |
| Adjusted.P.value | | 1.16E-01 | 6.76E-01 | 5.59E-01 | 4.19E-01 | 1.00E+00 |
| Odds.Ratio | | 23.70 | 82.30 | 74.07 | 74.07 | 18.99 |
| Combined.Score | | 253.33 | 677.85 | 594.15 | 594.15 | 100.83 |
| Genes | | XCR1 | APLNR | ADRA1D | ADRA1D | ADRA1D |
| | | ADRA1D | GPR61 | ADRA2C | ADRA2C | ADRA2C |
| | | AGTR2 | | | | |
| | | ADRA2C | | | | |

| | Group | cytokine-mediated signaling pathway | inflammatory response | cellular response to cytokine stimulus | cellular response to lipopolysaccharide | positive regulation of cytokine production |
|---|---|---|---|---|---|---|
| Overlap | Spleen_Up | 34/633 | 21/252 | 24/456 | 11/91 | 14/220 |
| P.value | | 2.11E-27 | 9.13E-21 | 5.46E-19 | 4.17E-13 | 1.73E-12 |
| Adjusted.P.value | | 1.08E-23 | 2.33E-17 | 9.28E-16 | 5.32E-10 | 1.76E-09 |
| Odds.Ratio | | 11.55 | 17.92 | 11.32 | 26.00 | 13.69 |
| Combined.Score | | 709.49 | 826.94 | 475.98 | 741.03 | 370.66 |
| Genes | | CCL17 | CCL17 | ACOD1 | ACOD1 | CD14 |
| | | CCL20 | CCL20 | CCL17 | CD14 | CEBPB |
| | | CDKN1A | CCRL2 | CCL20 | CD274 | CSF2 |
| | | CEBPD | CEBPB | CDKN1A | CXCL10 | DDX58 |
| | | CSF2 | CXCL10 | CEBPD | IL10 | DHX58 |
| | | CSF3 | CXCL11 | CSF2 | IL1B | EIF2AK2 |
| | | CXCL10 | CXCL5 | CSF3 | IL6 | IFNG |
| | | CXCL11 | FPR1 | CXCL10 | SERPINE1 | IL10 |
| | | CXCL5 | FPR2 | FPR1 | TNF | IL1A |
| | | FPR1 | GPR33 | ICAM1 | TNFAIP3 | IL1B |
| | | ICAM1 | IL10 | IFNB1 | TNIP3 | IL6 |
| | | IFNB1 | IL1A | IL10 | | IRF7 |
| | | IFNG | IL1B | IL19 | | SERPINE1 |
| | | IL10 | IL22 | IL1A | | TNF |
| | | IL15RA | IL6 | IL1B | | |
| | | IL19 | KRT16 | IL1R2 | | |
| | | IL1A | PROK2 | IL1RN | | |
| | | IL1B | PTX3 | IL22 | | |
| | | IL1R2 | SELE | IL6 | | |
| | | IL1RN | TNF | MMP3 | | |
| | | IL22 | TNFAIP6 | SOCS1 | | |
| | | IL27 | | SOCS3 | | |
| | | IL6 | | TIMP1 | | |
| | | IRF7 | | TNF | | |
| | | ISG15 | | | | |
| | | MMP3 | | | | |
| | | OAS2 | | | | |
| | | RSAD2 | | | | |
| | | SOCS1 | | | | |
| | | SOCS3 | | | | |
| | | SOD2 | | | | |
| | | TIMP1 | | | | |
| | | TNF | | | | |
| | | XAF1 | | | | |

| | Group | cytokine-mediated signaling pathway | inflammatory response | cellular response to cytokine stimulus | response to lipopolysaccharide | cellular response to lipopolysaccharide |
|---|---|---|---|---|---|---|
| Overlap | Liver_Up | 53/633 | 30/252 | 37/456 | 23/155 | 16/91 |
| P.value | | 1.81E-32 | 7.47E-23 | 3.90E-22 | 6.20E-20 | 2.00E-15 |
| Adjusted.P.value | | 9.22E-29 | 1.91E-19 | 6.63E-19 | 7.91E-17 | 2.04E-12 |
| Odds.Ratio | | 7.86 | 11.18 | 7.62 | 13.93 | 16.51 |
| Combined.Score | | 574.63 | 569.51 | 375.58 | 616.22 | 558.77 |
| Genes | | ADAR | ACKR1 | ACOD1 | ACOD1 | ACOD1 |
| | | BATF | AOAH | BATF | CD14 | CD14 |
| | | CCL20 | CCL20 | CCL20 | CD274 | CD274 |
| | | CCL22 | CCL22 | CCL22 | CD40 | CXCL10 |
| | | CD40 | CCRL2 | CDKN1A | CXCL10 | IL10 |
| | | CDKN1A | CD40 | CEBPD | CXCL11 | IL1B |
| | | CEBPD | CXCL10 | CSF2 | CXCL5 | IL6 |
| | | CSF2 | CXCL11 | CSF3 | CXCL9 | LBP |
| | | CSF3 | CXCL5 | CSF3R | IL10 | NLRP3 |
| | | CSF3R | CXCL9 | CXCL10 | IL6 | SBNO2 |
| | | CXCL10 | ELF3 | DUOX2 | IRAK3 | SERPINE1 |
| | | CXCL11 | FOS | FOS | JUNB | TNF |
| | | CXCL5 | FPR1 | FPR1 | LBP | TNFAIP3 |
| | | CXCL9 | FPR2 | FSCN1 | NLRP3 | TNIP2 |
| | | DUOX2 | GPR33 | ICAM1 | SBNO2 | TNIP3 |
| | | EBI3 | IL10 | IL10 | SELE | ZFP36 |
| | | FOS | IL1B | IL1B | SELP | |
| | | FPR1 | IL6 | IL1RN | SERPINE1 | |
| | | FSCN1 | MEFV | IL20RA | SLPI | |
| | | ICAM1 | NFKB2 | IL6 | TNFAIP3 | |
| | | IL10 | NLRP12 | IRAK3 | TNIP2 | |
| | | IL1B | NLRP3 | JUNB | TNIP3 | |
| | | IL1RN | PROK2 | LBP | ZFP36 | |
| | | IL20RA | PTX3 | LCN2 | | |
| | | IL27 | RELB | MMP3 | | |
| | | IL6 | RIPK2 | MMP9 | | |
| | | IRAK3 | S100A9 | NLRP12 | | |
| | | IRF7 | SELE | OSM | | |
| | | IRS2 | STAT3 | OSMR | | |
| | | ISG15 | TNF | RTN4RL2 | | |
| | | ISG20 | | SBNO2 | | |
| | | JUNB | | SOCS1 | | |
| | | LBP | | SOCS3 | | |
| | | LCN2 | | STAT3 | | |
| | | MMP3 | | TIMP1 | | |
| | | MMP9 | | TNF | | |
| | | OAS3 | | ZFP36 | | |
| | | OSM | | | | |
| | | OSMR | | | | |
| | | PML | | | | |
| | | PSMB8 | | | | |
| | | PSMB9 | | | | |
| | | RIPK2 | | | | |
| | | RSAD2 | | | | |
| | | RTN4RL2 | | | | |
| | | SAMHD1 | | | | |
| | | SOCS1 | | | | |
| | | SOCS3 | | | | |
| | | STAT3 | | | | |
| | | TIMP1 | | | | |
| | | TNF | | | | |
| | | TNIP2 | | | | |
| | | XAF1 | | | | |

**Table S12. Enrichment GO terms and included genes of the four modules of Figure 5**

| **darkorange** | neutrophil mediated immunity | neutrophil activation involved in immune response | neutrophil degranulation | cellular response to cytokine stimulus | cytokine-mediated signaling pathway |
|---|---|---|---|---|---|
| Overlap | 154/487 | 151/483 | 150/479 | 138/456 | 170/633 |
| P.value | 9.7E-26 | 1.1E-24 | 1.3E-24 | 3.3E-21 | 8.8E-20 |
| Adjusted.P.value | 5.0E-22 | 2.8E-21 | 2.2E-21 | 4.2E-18 | 8.9E-17 |
| Odds.Ratio | 2.35 | 2.32 | 2.32 | 2.25 | 1.99 |
| Combined.Score | 135.20 | 128.05 | 127.86 | 105.94 | 87.49 |
| Genes | ABCA13 | ABCA13 | ABCA13 | ACOD1 | ADAM17 |
| | ACE | ACPP | ACPP | ADAMTS12 | ADAR |
| | ACPP | ACTR2 | ACTR2 | ALOX5 | ALOX5 |
| | ACTR2 | ADGRG3 | ADGRG3 | ANKRD1 | ANXA1 |
| | ADAM17 | AGPAT2 | AGPAT2 | ANXA1 | ANXA2 |
| | ADGRG3 | ALDOA | ALDOA | BATF | ARF1 |
| | AGPAT2 | ALDOC | ALDOC | BCL2L1 | B2M |
| | ALDOA | ALOX5 | ALOX5 | BCL6 | BATF |
| | ALDOC | AMPD3 | AMPD3 | CACTIN | BCL2L1 |
| | ALOX5 | ANPEP | ANPEP | CASP3 | BCL6 |
| | AMPD3 | ANXA2 | ANXA2 | CCL19 | BRWD1 |
| | ANPEP | ANXA3 | ANXA3 | CCL20 | CAPZA1 |
| | ANXA2 | AP2A2 | AP2A2 | CDK9 | CASP3 |
| | ANXA3 | APAF1 | APAF1 | CEBPD | CCL19 |
| | AP2A2 | ARG1 | ARG1 | CHUK | CCL20 |
| | APAF1 | ARMC8 | ARMC8 | CIB1 | CD300LF |
| | ARG1 | ARSA | ARSA | CISH | CD44 |
| | ARMC8 | ARSB | ARSB | CLCF1 | CDC42 |
| | ARSA | ATP11A | ATP11A | CNTFR | CEBPD |
| | ARSB | ATP6AP2 | ATP6AP2 | CRK | CFL1 |
| | ATP11A | ATP6V0A1 | ATP6V0A1 | CSF1 | CHUK |
| | ATP6AP2 | ATP6V0C | ATP6V0C | CSF2 | CIITA |
| | ATP6V0A1 | ATP8A1 | ATP8A1 | CSF2RB | CISH |
| | ATP6V0C | ATP8B4 | ATP8B4 | CSF3R | CLCF1 |
| | ATP8A1 | B2M | B2M | CX3CL1 | CNTFR |
| | ATP8B4 | BST1 | BST1 | DAB2IP | CRK |
| | B2M | C5AR1 | C5AR1 | FN1 | CSF1 |
| | BST1 | CAMP | CAMP | FOXH1 | CSF2 |
| | C5AR1 | CAP1 | CAP1 | FOXO3 | CSF2RB |
| | CAMP | CAPN1 | CAPN1 | FPR1 | CSF3R |
| | CAP1 | CD44 | CD44 | GAB2 | CX3CL1 |
| | CAPN1 | CD47 | CD47 | GAPDH | CXCL11 |
| | CD44 | CD53 | CD53 | GATA3 | CXCL5 |
| | CD47 | CD63 | CD63 | GPER1 | EPO |
| | CD53 | CD68 | CD68 | GSK3A | FBXW11 |
| | CD63 | CHRNB4 | CHRNB4 | HAS2 | FN1 |
| | CD68 | CKAP4 | CKAP4 | HCLS1 | FOXO3 |
| | CHRNB4 | CLEC5A | CLEC5A | HGF | FPR1 |
| | CKAP4 | CMTM6 | CMTM6 | HMOX1 | GAB2 |
| | CLEC5A | CPNE3 | CPNE3 | HSP90B1 | GATA3 |
| | CMTM6 | CRISPLD2 | CRISPLD2 | HSPA8 | GPR35 |
| | CPNE3 | CSTB | CSTB | HYAL2 | HGF |
| | CRISPLD2 | CTSB | CTSB | HYAL3 | HMOX1 |
| | CSTB | CTSH | CTSH | ICAM1 | HNRNPF |
| | CTSB | CTSS | CTSS | IFNAR2 | HSP90B1 |
| | CTSH | CXCR2 | CXCR2 | IFNB1 | HSPA8 |
| | CTSS | CYFIP1 | CYFIP1 | IFNGR1 | ICAM1 |
| | CXCL5 | CYSTM1 | CYSTM1 | IFNGR2 | IFI35 |
| | CXCR2 | DEGS1 | DEGS1 | IFNLR1 | IFNAR2 |
| | CYFIP1 | DGAT1 | DGAT1 | IL10 | IFNB1 |
| | CYSTM1 | DLC1 | DLC1 | IL10RB | IFNGR1 |
| | DEGS1 | DNAJC13 | DNAJC13 | IL13RA1 | IFNGR2 |
| | DGAT1 | DNASE1L3 | DOK3 | IL18BP | IFNLR1 |
| | DLC1 | DOK3 | DYNC1H1 | IL18R1 | IKBKG |
| | DNAJC13 | DYNC1H1 | ENPP4 | IL19 | IL10 |
| | DOK3 | ENPP4 | EPX | IL1B | IL10RB |
| | DYNC1H1 | EPX | FABP5 | IL1R1 | IL12RB2 |
| | ENPP4 | FABP5 | FCER1G | IL1R2 | IL13RA1 |
| | EPX | FCER1G | FGR | IL1RAP | IL15RA |
| | FABP5 | FGR | FPR1 | IL1RN | IL17RA |
| | FCER1G | FPR1 | FPR2 | IL20RA | IL18BP |
| | FGR | FPR2 | FUCA1 | IL20RB | IL18R1 |
| | FPR1 | FUCA1 | FUCA2 | IL22RA2 | IL19 |
| | FPR2 | FUCA2 | GCA | IL23R | IL1B |
| | FUCA1 | GCA | GMFG | IL2RB | IL1R1 |
| | FUCA2 | GMFG | GPR84 | IL31RA | IL1R2 |
| | GCA | GPR84 | HK3 | INPP5K | IL1RAP |
| | GMFG | HK3 | HSPA1A | IRAK2 | IL1RN |
| | GPR84 | HSPA1A | HSPA8 | IRAK4 | IL20RA |
| | HK3 | HSPA8 | IQGAP1 | IRF8 | IL20RB |
| | HSPA1A | IQGAP1 | IQGAP2 | JAK1 | IL22RA2 |
| | HSPA8 | IQGAP2 | ITGAV | JAK2 | IL23R |
| | IQGAP1 | ITGAV | KCNAB2 | JUNB | IL2RB |
| | IQGAP2 | KCNAB2 | LAMP2 | LBP | IL31RA |
| | IRAK4 | LAMP2 | LAMTOR1 | LCN2 | ILK |
| | ITGAV | LAMTOR1 | LAMTOR3 | LRP8 | IRAK2 |
| | KCNAB2 | LAMTOR3 | LCN2 | LRRC3 | IRAK4 |
| | LAMP2 | LCN2 | LGALS3 | MAPK11 | IRF2 |
| | LAMTOR1 | LGALS3 | LPCAT1 | MAPK13 | IRF6 |
| | LAMTOR3 | LPCAT1 | LRG1 | MCL1 | IRF8 |
| | LCN2 | LRG1 | LTF | MMP3 | IRF9 |
| | LGALS3 | LTF | MAN2B1 | MMP9 | ISG20 |

| neutrophil mediated immunity | neutrophil activation involved in immune response | neutrophil degranulation | cellular response to cytokine stimulus | cytokine-mediated signaling pathway |
|---|---|---|---|---|
| LPCAT1 | MAN2B1 | MAPK14 | MYC | JAK1 |
| LRG1 | MAPK14 | MCEMP1 | NFE2L2 | JAK2 |
| LTF | MCEMP1 | MGAM | NLRP12 | JUNB |
| MAN2B1 | MGAM | MMP25 | NPNT | KARS |
| MAPK14 | MMP25 | MMP8 | NUMBL | LBP |
| MCEMP1 | MMP8 | MMP9 | OCSTAMP | LCN2 |
| MGAM | MMP9 | MS4A3 | OSM | LCP1 |
| MMP25 | MS4A3 | MVP | PDE12 | LEP |
| MMP8 | MVP | NCKAP1L | PDE2A | LEPR |
| MMP9 | NCKAP1L | NCSTN | PID1 | LMNB1 |
| MS4A3 | NCSTN | NFAM1 | PIK3CB | LRP8 |
| MVP | NFAM1 | NFASC | PIM1 | LRRC3 |
| NCKAP1L | NFASC | OLFM4 | PTAFR | LTBR |
| NCSTN | OLFM4 | OLR1 | PTGIS | MAP3K14 |
| NFAM1 | OLR1 | OSTF1 | PTGS2 | MCL1 |
| NFASC | OSTF1 | PAFAH1B2 | PTPN12 | MMP3 |
| OLFM4 | PAFAH1B2 | PDXK | PTPN13 | MMP9 |
| OLR1 | PDXK | PGLYRP1 | PTPN14 | MSN |
| OSTF1 | PGLYRP1 | PGM2 | PTPN6 | MTAP |
| PAFAH1B2 | PGM2 | PKM | RELA | MYC |
| PDXK | PKM | PLAU | RHOU | MYD88 |
| PGLYRP1 | PLAU | PLAUR | RIPK1 | NCAM1 |
| PGM2 | PLAUR | PNP | RIPOR2 | NFKBIA |
| PKM | PNP | PRDX6 | RTN4RL2 | NGFR |
| PLAU | PRDX6 | PRG2 | SBNO2 | NOD1 |
| PLAUR | PRG2 | PRG3 | SDC1 | NUMBL |
| PNP | PRG3 | PSMA2 | SELPLG | OSM |
| PRDX6 | PSMA2 | PSMD1 | SOCS1 | PAK2 |
| PRG2 | PSMD1 | PSMD12 | SOCS3 | PDCD4 |
| PRG3 | PSMD12 | PTAFR | SOCS5 | PIK3CB |
| PSMA2 | PTAFR | PTGES2 | STAT1 | PIM1 |
| PSMD1 | PTGES2 | PTPN6 | STAT2 | PSMA1 |
| PSMD12 | PTPN6 | PTPRJ | STAT3 | PSMA2 |
| PTAFR | PTPRJ | PTX3 | STAT5B | PSMA3 |
| PTGES2 | PTX3 | RAB10 | TCF7 | PSMC1 |
| PTPN6 | RAB10 | RAB18 | TEC | PSMD1 |
| PTPRJ | RAB18 | RAB31 | TGFB1 | PSMD12 |
| PTX3 | RAB31 | RAB3D | TIMP1 | PSMD5 |
| RAB10 | RAB3D | RAB44 | TNF | PSMF1 |
| RAB18 | RAB44 | RAB4B | TNFRSF1A | PTAFR |
| RAB31 | RAB4B | RAB5C | TNFRSF1B | PTGS2 |
| RAB3D | RAB5C | RAB6A | TNFRSF21 | PTK2B |
| RAB44 | RAB6A | RAP1B | TYK2 | PTPN6 |
| RAB4B | RAP1B | RAP2B | UBC | RAP1B |
| RAB5C | RAP2B | RAP2C | UBE2G2 | RELA |
| RAB6A | RAP2C | RETN | UBE2K | RHOU |
| RAP1B | RETN | RHOG | VAV1 | RIPK1 |
| RAP2B | RHOG | S100A11 | VCAM1 | RTN4RL2 |
| RAP2C | S100A11 | S100A9 | VEGFA | SAMHD1 |
| RETN | S100A9 | SDCBP | VIM | SDC1 |
| RHOG | SDCBP | SELL | WNT5A | SOCS1 |
| S100A11 | SELL | SERPINB10 | XBP1 | SOCS3 |
| S100A9 | SERPINB10 | SIRPA | YBX3 | SOCS5 |
| SDCBP | SIRPA | SLC11A1 | YWHAZ | SOD2 |
| SELL | SLC11A1 | SLCO4C1 | ZEB1 | SQSTM1 |
| SERPINB10 | SLCO4C1 | SLPI | ZFP36 | STAT1 |
| SIRPA | SLPI | STOM | | STAT2 |
| SLC11A1 | STOM | STXBP2 | | STAT3 |
| SLCO4C1 | STXBP2 | TCIRG1 | | STAT5B |
| SLPI | TCIRG1 | TLR2 | | TALDO1 |
| STOM | TLR2 | TMBIM1 | | TEC |
| STXBP2 | TMBIM1 | TMEM173 | | TGFB1 |
| TCIRG1 | TMEM173 | TNFRSF1B | | TIMP1 |
| TLR2 | TNFRSF1B | TRPM2 | | TNF |
| TMBIM1 | TRPM2 | TSPAN14 | | TNFRSF14 |
| TMEM173 | TSPAN14 | TYROBP | | TNFRSF18 |
| TNFRSF1B | TYROBP | VCL | | TNFRSF1A |
| TRPM2 | VCL | VCP | | TNFRSF1B |
| TSPAN14 | VCP | | | TNFRSF21 |
| TYROBP | | | | TNFRSF4 |
| VCL | | | | TNFRSF8 |
| VCP | | | | TNFSF13B |
| | | | | TNFSF14 |
| | | | | TNFSF18 |
| | | | | TRIM21 |
| | | | | TRIM25 |
| | | | | TRIM31 |
| | | | | TRIM38 |
| | | | | TRIM8 |
| | | | | TSLP |
| | | | | TYK2 |
| | | | | UBC |
| | | | | VAV1 |
| | | | | VCAM1 |
| | | | | VEGFA |
| | | | | VIM |
| | | | | YWHAZ |
| | | | | ZEB1 |

| darkseagreen | plasma membrane bounded cell projection assembly | cilium assembly | transmembrane receptor protein tyrosine kinase signaling pathway | cellular protein modification process | protein phosphorylation | cilium assembly | transmembrane receptor protein tyrosine kinase signaling pathway | cellular protein modification process | protein phosphorylation | cellular protein modification process | cellular protein modification process |
|---|---|---|---|---|---|---|---|---|---|---|---|
| Overlap | 131/241 | 146/288 | 188/396 | 416/1001 | 215/470 | TMEM237 | PIK3C2A | FBXL4 | NRBP2 | PIGS | VRK2 |
| P.value | 3.2E-12 | 1.8E-10 | 6.1E-10 | 1.4E-09 | 2.2E-09 | TNPO1 | PIK3CB | FBXL5 | NTRK2 | PIGU | VWA1 |
| Adjusted.P.value | 1.6E-08 | 4.7E-07 | 1.0E-06 | 1.8E-06 | 2.2E-06 | TOGARAM1 | PIK3R1 | FBXL7 | NUAK1 | PIK3C3 | WDR45 |
| Odds.Ratio | 1.66 | 1.55 | 1.45 | 1.27 | 1.40 | TTBK2 | PIK3R2 | FBXO10 | PAK1 | PIK3CB | WDTC1 |
| Combined.Score | 43.91 | 34.69 | 30.74 | 25.85 | 27.85 | TTC21B | PIK3R3 | FBXO11 | PDPK1 | PIK3CD | WDYHV1 |
| Genes | ABCC4 | ABCC4 | ABL1 | AAK1 | AAK1 | TTC26 | PLCB1 | FBXO17 | PEAK1 | PIK3R4 | WIPI2 |
| | ABLIM2 | ABLIM3 | ACTR2 | ABL1 | ABL1 | TTLL3 | POLR2A | FBXO2 | PHKA1 | PINK1 | WNK1 |
| | ABLIM3 | ACTR1A | ACTR3 | ACVR1 | ACOD1 | TUBB4B | POLR2C | FBXO31 | PHKB | PKN2 | WNK2 |
| | ACTN2 | ACTR2 | AKT1 | ACVR1C | ACVR1 | TUBG1 | POLR2D | FBXO32 | PHKG1 | PKN3 | WNK3 |
| | ACTR2 | ACTR3 | ANGPT1 | ADAM10 | ACVR1C | UBXN10 | POLR2E | FBXO41 | PICK1 | PLK2 | WNK4 |
| | ACTR3 | AKAP9 | ANGPT2 | ADM2 | ADAM10 | UNC119B | POLR2G | FBXO6 | PIK3C3 | PLOD1 | WNT11 |
| | AIF1 | ARL3 | AP2A1 | AGA | ADM2 | WDR19 | POLR2H | FBXO9 | PIK3CB | PMM2 | YES1 |
| | ANO6 | ARL6 | AP2A2 | AKT1 | AKT1 | WDR35 | POLR2I | FBXW2 | PIK3CD | POMT1 | ZBTB16 |
| | ARFIP2 | ASAP1 | AP2M1 | ALG5 | APP | YWHAE | POLR2K | FBXW5 | PIK3R4 | POMT2 | |
| | ARHGEF6 | ATP6V0D1 | AP3S1 | ALOX12B | ATM | ZMYND10 | POLR2L | FBXW7 | PINK1 | PORCN | |
| | ARHGEF7 | ATP6V1D | APH1A | AMBN | AURKB | | PRKD1 | FBXW8 | PKD1 | PPM1A | |
| | ARL3 | B9D1 | ARHGEF28 | ANO8 | BCR | | PRLR | FBXW9 | PKN2 | PPM1D | |
| | ARL6 | BBOF1 | ARHGEF7 | ANXA1 | BMP10 | | PSEN1 | FEM1A | PKN3 | PPM1G | |
| | ASAP1 | BBS1 | ARPC1A | APOA1 | BMP2 | | PSEN2 | FEM1B | PLK2 | PPP1CA | |
| | ATP6V0D1 | BBS10 | ARPC3 | APOA2 | BMPR1A | | PTBP1 | FER | PNCK | PPP1CB | |
| | ATP6V1D | BBS2 | ARPC4 | APP | BRSK1 | | PTK2 | FES | PPP3CB | PPP2CA | |
| | B9D1 | BBS5 | ATP6AP1 | ARSA | BRSK2 | | PTPN11 | FRK | PRDX4 | PPP2R1A | |
| | BBS1 | BBS7 | ATP6V0A2 | ARSB | BTK | | PTPRG | FSTL1 | PRKAA1 | PPP2R5A | |
| | BBS2 | BBS9 | ATP6V0A4 | ARSJ | CAMK1D | | PTPRT | FSTL3 | PRKAB1 | PPP2R5D | |
| | BBS5 | C2CD3 | ATP6V0D1 | ARSK | CAMK2D | | PXN | FUCA2 | PRKACA | PPP3CB | |
| | BBS7 | CBY1 | ATP6V0E2 | ASB10 | CAMK2G | | RAB14 | FUT1 | PRKACB | PRKAA1 | |
| | BBS9 | CBY3 | ATP6V1A | ASB11 | CAMK4 | | RAPGEF2 | FUT7 | PRKAG1 | PRKAB1 | |
| | CBY1 | CC2D2A | ATP6V1C2 | ASB16 | CAMKK1 | | RASGRP4 | GAS6 | PRKAR1B | PRKACA | |
| | CBY3 | CCDC113 | ATP6V1D | ASB2 | CAMKV | | RGS14 | GGCX | PRKCB | PRKACB | |
| | CC2D2A | CCDC13 | ATP6V1E1 | ASB3 | CDC42BPA | | RHOA | GMFG | PRKCH | PRKAG1 | |
| | CCDC113 | CCDC39 | ATP6V1E2 | ASB7 | CDC7 | | ROCK1 | GOLGA2 | PRKCZ | PRKAR1B | |
| | CCDC65 | CCDC65 | ATP6V1G1 | ASB8 | CDK10 | | SDC2 | GPAA1 | PRKD1 | PRKCB | |
| | CCDC88A | CDC14A | ATP6V1H | ATE1 | CDK13 | | SETX | GPC3 | PRKDC | PRKCH | |
| | CDC14A | CDC14B | AXL | ATG10 | CDK14 | | SH2D6 | GSK3B | PSKH1 | PRKCSH | |
| | CDC14B | CEP126 | BAIAP2 | ATG4C | CDK15 | | SHC1 | HIF3A | PTK2 | PRKCZ | |
| | CDH13 | CEP131 | BAIAP2L1 | ATG7 | CDK16 | | SHC2 | HIPK2 | PTPN6 | PRKD1 | |
| | CEP126 | CEP135 | BAIAP2L2 | ATM | CDK18 | | SHC3 | HLCS | RAF1 | PRKDC | |
| | CEP131 | CEP164 | BCAR1 | AURKB | CDK8 | | SHOC2 | HRC | RARA | PSMA2 | |
| | CEP164 | CEP19 | BDKRB2 | B3GALNT1 | CDK9 | | SIRT2 | ICK | RET | PSMA3 | |
| | CEP19 | CEP192 | BDNF | B3GALT4 | CDKL2 | | SLA2 | ICMT | ROS1 | PSMB5 | |
| | CEP290 | CEP250 | BMPR2 | B3GALT5 | CREB1 | | SOGA1 | IGFBP3 | RPS6KA1 | PSMB7 | |
| | CEP41 | CEP290 | BRK1 | B3GNT2 | CRY1 | | SORT1 | IGFBP4 | RPS6KA2 | PSMC1 | |
| | CEP89 | CEP41 | BTK | B3GNT6 | CSF1R | | SOS1 | IGFBP5 | RPS6KA4 | PSMC3 | |
| | CFAP221 | CEP57 | CAV2 | B3GNT9 | CSK | | SPRED2 | IKBKB | RPS6KA5 | PSMC4 | |
| | CLUAP1 | CEP78 | CD3E | B4GALT7 | CSNK1A1 | | SRC | IRAK1 | RPS6KA6 | PSMC6 | |
| | CYFIP1 | CEP89 | CD4 | BAP1 | CSNK1G1 | | SULF2 | IVL | RPS6KB1 | PSMD1 | |
| | DNAAF1 | CFAP206 | CD7 | BCR | CSNK1G3 | | TEK | KCTD6 | SBK1 | PSMD11 | |
| | DNAH6 | CFAP221 | CD8A | BMP2 | CSNK2B | | TIA1 | KCTD7 | SMAD7 | PSMD12 | |
| | DYNC2H1 | CKAP5 | CDC42 | BMPR1A | CTBP1 | | TIAL1 | KEAP1 | SNRK | PSMD13 | |
| | DYNLL1 | CLASP1 | CEP57 | BPIFB2 | DAB2IP | | TSC2 | KLHL13 | SRC | PSMD2 | |
| | DYNLL2 | CLUAP1 | CHN1 | BRSK1 | DAPK1 | | TYK2 | KLHL20 | SRPK1 | PSMD3 | |
| | DZIP1 | DCTN1 | CHRNA3 | BRSK2 | DAPK2 | | VAV2 | KLHL21 | SRPK2 | PSMD5 | |
| | DZIP1L | DNAAF1 | CNKSR1 | BTBD1 | DAPK3 | | VEGFA | KLHL25 | STK10 | PSME3 | |
| | EMP2 | DNAH6 | CSF1R | BTBD6 | DGKQ | | VEGFC | KLHL3 | STK16 | PTAR1 | |
| | EMP3 | DYNC1I2 | CSK | BTK | DSTYK | | WASL | KLHL41 | STK24 | PTP4A2 | |
| | FAM161A | DYNC2H1 | CYFIP1 | CAMK2D | DYRK1A | | YES1 | KLHL5 | STK25 | PTPN14 | |
| | FAM92A | DYNC2LI1 | CYFIP2 | CAMK4 | DYRK1B | | ZFYVE27 | KRT2 | STK26 | PTPN5 | |
| | FBF1 | DYNLL1 | DNM1 | CAND1 | DYRK2 | | | KSR1 | STK32B | PTPN6 | |
| | FGD1 | DYNLL2 | DOCK1 | CCDC22 | EFEMP1 | | | LAMB1 | STK33 | PTPN7 | |
| | FGD3 | DYNLRB1 | EFNA1 | CCNF | EGFR | | | LAMB2 | STK36 | PTPN9 | |
| | FGD4 | DZIP1 | EFNA2 | CDC14B | EIF2AK1 | | | LARGE1 | STK38 | PTPRB | |
| | FGD5 | DZIP1L | EFNA3 | CDC42BPA | EIF2AK3 | | | LATS2 | STK38L | PTPRD | |
| | FGD6 | FAM161A | EFNA4 | CDH2 | EIF2AK4 | | | LCK | STK4 | PTPRN2 | |
| | FLNA | FAM92A | EFNB1 | CDK10 | EPHA3 | | | LGALS1 | TAOK2 | PTPRO | |
| | GALNT11 | FBF1 | EFS | CDK13 | EPHB3 | | | LHPP | TAOK3 | PTPRR | |
| | GSN | FLNA | EIF2AK3 | CDK14 | EPHB4 | | | LIMK1 | TEK | PTPRS | |
| | HDAC6 | GALNT11 | ELMO2 | CDK15 | ERBB2 | | | LOXL1 | TESK2 | PTPRT | |
| | ICK | GSN | EPHA10 | CDK16 | ERG | | | LOXL2 | TGFB2 | PTPRU | |
| | IFT122 | HAUS2 | EPHA2 | CDK18 | FAM20A | | | MACROD2 | TGFBR1 | PTPRZ1 | |
| | IFT172 | HAUS4 | EPHA3 | CDK8 | FAM20C | | | MAN2A1 | TGFBR3 | RAB11A | |
| | IFT20 | HAUS5 | EPHA7 | CDK9 | FASTK | | | MAN2B2 | TLR7 | RAB1B | |
| | IFT43 | HAUS7 | EPHB3 | CDKL2 | FER | | | MANBA | TNIK | RAB2A | |
| | IFT46 | HAUS8 | EPHB4 | CEL | FES | | | MAP3K11 | TTBK2 | RAB38 | |
| | IFT52 | HDAC6 | EPHB6 | CHGB | FGFR2 | | | MAP3K12 | TYK2 | RAB3A | |
| | IFT57 | HSPB11 | ERBB2 | COG7 | FGFR3 | | | MAP3K13 | UHMK1 | RAB8A | |
| | IFT74 | ICK | ESRP1 | COL3A1 | FGFR4 | | | MAP3K2 | ULK2 | RABGGTA | |
| | IFT81 | IFT122 | FER | COMMD1 | FLT1 | | | MAP3K20 | ULK3 | RAF1 | |
| | IFT88 | IFT140 | FES | COPS5 | FLT3 | | | MAP3K21 | USP15 | RARA | |
| | IQCB1 | IFT172 | FGF10 | COPS7A | FLT4 | | | MAP3K6 | VRK2 | RBBP5 | |

| | | | | | | | |
|---|---|---|---|---|---|---|---|
| KIF3A | IFT20 | FGF18 | COPS7B | FRK | MAP4K1 | WNK1 | RET |
| KIT | IFT43 | FGF22 | COPS8 | GAS6 | MAPK10 | WNK2 | RHBDD1 |
| MKKS | IFT46 | FGF3 | CPE | GMFG | MAPK3 | WNK3 | RNF7 |
| MKS1 | IFT52 | FGF5 | CREB1 | GSK3B | MAPK4 | WNK4 | ROS1 |
| MTSS1 | IFT57 | FGF9 | CSK | GTF2H1 | MAPK6 | WNT11 | RPS6KA1 |
| MYLK | IFT74 | FGFR2 | CSNK1A1 | GTF2H4 | MAPK8 | | RPS6KA2 |
| NPHP3 | IFT81 | FGFR3 | CSNK1G3 | HIPK2 | MAPK9 | | RPS6KA4 |
| NTN1 | IFT88 | FGFR4 | CTBP1 | ICK | MAPKAPK2 | | RPS6KA5 |
| NUDCD3 | IQCB1 | FGFRL1 | CTDNEP1 | IGFBP3 | MARK4 | | RPS6KA6 |
| OCRL | KIF17 | FLRT2 | CTDSP1 | IKBKB | MAST2 | | RPS6KB1 |
| OFD1 | KIF3A | FLT1 | CTDSP2 | IRAK1 | MBTPS1 | | SBF1 |
| OPHN1 | KIF3C | FLT3 | CTH | KDR | MELTF | | SDC2 |
| P2RX7 | KIFAP3 | FLT4 | CUL3 | KIT | MEN1 | | SDHAF2 |
| PARVB | LRRC6 | FOXO1 | CUL4A | KSR1 | MEX3B | | SENP8 |
| PCNT | MARK4 | FRK | CUL4B | LATS2 | MINK1 | | SERP2 |
| PMP22 | MKKS | FRS3 | CUL7 | LCK | MKNK1 | | SKP2 |
| POC1B | MKS1 | GAB1 | CUL9 | LIMK1 | MMP15 | | SLC35A1 |
| PPP1R9B | NDE1 | GHR | DAPK1 | MAP2K6 | MPI | | SNRK |
| PTPDC1 | NEK2 | GIGYF2 | DAPK2 | MAP3K10 | MSRA | | SOCS5 |
| PTPRO | NINL | GIT1 | DAPK3 | MAP3K11 | MTMR2 | | SPARCL1 |
| RAB17 | NPHP1 | GPLD1 | DCAF11 | MAP3K12 | MTMR4 | | SPSB2 |
| RAB23 | NPHP3 | HGF | DCAF17 | MAP3K13 | MTMR6 | | SPSB3 |
| RAB8A | NPHP4 | HRAS | DCAF4 | MAP3K2 | MTMR7 | | SPSB4 |
| RAPGEF2 | NUDCD3 | IDE | DCAF5 | MAP3K20 | MXRA8 | | SRPK1 |
| RFX3 | OCRL | IL31RA | DCAF6 | MAP3K21 | MYLK3 | | SRPK2 |
| RFX4 | OFD1 | ITGAV | DCAF7 | MAP3K3 | MYOD1 | | ST3GAL4 |
| RILP | PAFAH1B1 | KALRN | DCAF8 | MAP3K6 | NEK10 | | ST6GALNAC2 |
| RILPL1 | PCNT | KDR | DCUN1D1 | MAP4K1 | NEK2 | | ST8SIA2 |
| RILPL2 | POC1B | KIF16B | DDA1 | MAPK10 | NEK3 | | STC2 |
| S1PR1 | PPP2R1A | KIT | DDB1 | MAPK13 | NEK4 | | STK10 |
| SCLT1 | PRKACA | LCK | DDOST | MAPK14 | NEK7 | | STK16 |
| SHTN1 | PRKAR2B | MAPK13 | DMP1 | MAPK3 | NGLY1 | | STK24 |
| SLC9A3R1 | PTPDC1 | MAPK14 | DOLK | MAPK4 | NIM1K | | STK25 |
| SNAP29 | RAB11A | MAPK3 | DSTYK | MAPK6 | NLK | | STK26 |
| SPATA13 | RAB17 | MAPKAPK2 | DUPD1 | MAPK8 | NOTUM | | STK36 |
| SRGAP2 | RAB23 | MAPKAPK3 | DUSP6 | MAPK9 | NPC1 | | STK38 |
| SSX2IP | RAB3IP | MMP2 | DUSP7 | MAPKAPK2 | NRBP1 | | STK38L |
| STK26 | RAB8A | MPZL1 | DUSP8 | MAPKAPK3 | NRBP2 | | STK4 |
| STK36 | RFX3 | MST1R | DYRK1A | MAPKAPK5 | NUAK1 | | STYXL1 |
| TBC1D32 | RFX4 | MTSS1 | DYRK1B | MARK4 | NUCB1 | | SUMF1 |
| TCTN1 | RILP | NCBP1 | DYRK2 | MAST2 | OBSL1 | | SUMF2 |
| TCTN2 | RILPL1 | NCKAP1 | EGLN3 | MAST3 | P3H1 | | TAOK3 |
| TCTN3 | RILPL2 | NCKAP1L | EIF2AK3 | MAST4 | P4HB | | TESK2 |
| TMEM17 | SCLT1 | NCSTN | EIF2AK4 | MEX3B | PAK1 | | TGFB2 |
| TMEM216 | SDCCAG8 | NEDD9 | ERBB2 | MINK1 | PCSK9 | | TGFBR1 |
| TMEM231 | SNAP29 | NELL2 | ERG | MKNK1 | PDPK1 | | TIMM50 |
| TMEM237 | SPATA6 | NGEF | EVPL | MYLK2 | PDXP | | TNIK |
| TNIK | SSNA1 | NGFR | FAM20A | MYLK3 | PEAK1 | | TRIOBP |
| TTBK2 | SSX2IP | NTF3 | FAM20C | MYOD1 | PGP | | TULP4 |
| TTC26 | STK36 | NTRK2 | FASTK | NEK10 | PHEX | | TYK2 |
| TTLL3 | TBC1D32 | NTRK3 | FBN1 | NEK2 | PHKA1 | | UBA7 |
| UBXN10 | TCTN1 | PAK1 | FBXL14 | NEK3 | PHKB | | UBE2E1 |
| UNC119B | TCTN2 | PAK3 | FBXL16 | NEK4 | PHLPP1 | | UBE2M |
| VCL | TCTN3 | PDGFA | FBXL18 | NEK7 | PHPT1 | | UBL4A |
| VSTM5 | TMEM17 | PDGFD | FBXL19 | NIM1K | PICK1 | | UBLCP1 |
| WDR19 | TMEM216 | PDK2 | FBXL2 | NLK | PIGH | | USP25 |
| WDR35 | TMEM231 | PGF | FBXL22 | NRBP1 | PIGK | | VHL |

| brown | transmembrane receptor protein tyrosine kinase signaling pathway | protein autophosphorylation | phosphorylation | positive regulation of endothelial cell migration | response to insulin |
|---|---|---|---|---|---|
| Overlap | 38/396 | 21/175 | 35/386 | 12/70 | 12/71 |
| P.value | 6.8E-06 | 3.2E-05 | 5.1E-05 | 5.2E-05 | 6.0E-05 |
| Adjusted.P.value | 3.5E-02 | 8.3E-02 | 8.6E-02 | 6.7E-02 | 6.1E-02 |
| Odds.Ratio | 2.17 | 2.71 | 2.05 | 3.87 | 3.82 |
| Combined.Score | 25.81 | 28.03 | 20.27 | 38.20 | 37.11 |
| Genes | ABL1 | ABL1 | ABL1 | AAMP | FOXO1 |
| | ATP6V1C1 | ATP13A2 | BCKDK | ABL1 | GOT1 |
| | ATP6V1E1 | ATR | BIRC6 | BCAS3 | GPLD1 |
| | ATP6V1G2 | CAMKK2 | CAMKK2 | CD40 | IRS1 |
| | CSK | CSK | CDC25B | FLT4 | KAT2B |
| | ELMO2 | DAPK2 | CDC42BPB | GPLD1 | MARS |
| | EPHA1 | EIF2AK2 | CDC42BPG | ITGB1BP1 | OGT |
| | EPHB3 | EPHA1 | CHCHD10 | KDR | PDE3B |
| | ESRP2 | EPHB3 | CSK | NOS3 | PIK3R2 |
| | FGF23 | FGR | CSNK1G2 | PLCG1 | SLC2A4 |
| | FGR | FLT4 | DAPK2 | PRKD2 | TSC1 |
| | FIBP | KDR | DGKE | TEK | XBP1 |
| | FLT4 | MAPK3 | DGKQ | | |
| | FOXO1 | MATK | EIF2AK2 | | |
| | FRS3 | MEX3B | FGR | | |
| | GAB1 | MINK1 | GALK2 | | |
| | GPLD1 | PIM1 | GNPTAB | | |
| | IRS1 | PIM3 | HUNK | | |
| | KDR | PRKD2 | ICK | | |
| | KLB | SLA | MAP4K5 | | |
| | MAPK1 | TEK | MAPK1 | | |
| | MAPK3 | | MAPK3 | | |
| | MATK | | MATK | | |
| | PDK2 | | MEX3B | | |
| | PIK3R2 | | MINK1 | | |
| | POLR2A | | NEK1 | | |
| | POLR2B | | NEK3 | | |
| | POLR2J | | PIM1 | | |
| | PRKD2 | | PIM3 | | |
| | PSENEN | | PKN1 | | |
| | RAPGEF1 | | PNKP | | |
| | RASGRP4 | | PRKD2 | | |
| | SIPA1L1 | | SRPK2 | | |
| | SLA | | STAT3 | | |
| | STAT3 | | TEFM | | |
| | TEK | | | | |
| | TSC2 | | | | |
| | VAV3 | | | | |

| lightblue | cytokine-mediated signaling pathway | cellular response to cytokine stimulus | response to cytokine | response to lipopolysaccharide | response to molecule of bacterial origin | cytokine-mediated signaling pathway | cellular response to cytokine stimulus | cytokine-mediated signaling pathway |
|---|---|---|---|---|---|---|---|---|
| Overlap | 121/633 | 87/456 | 38/138 | 40/155 | 28/98 | IL12B | IL22 | PSMB10 |
| P.value | 5.3E-21 | 2.4E-15 | 2.6E-12 | 6.7E-12 | 7.6E-10 | IL17C | IL23R | PSMB8 |
| Adjusted.P.value | 2.7E-17 | 6.0E-12 | 4.4E-09 | 8.5E-09 | 7.8E-07 | IL17F | IL2RA | PSMB9 |
| Odds.Ratio | 2.47 | 2.46 | 3.56 | 3.33 | 3.69 | IL17RA | IL2RG | PSME1 |
| Combined.Score | 115.32 | 83.02 | 94.89 | 85.79 | 77.50 | IL18RAP | IRAK3 | PSME2 |
| Genes | ADAM17 | ACOD1 | ADAR | ACOD1 | ACOD1 | IL1A | ITGAM | PTAFR |
| | ADAR | BCL2L1 | BCL2L1 | ADAM17 | ADAM17 | IL1R1 | JAK3 | RAP1B |
| | BCL2L1 | CCL17 | BST2 | CD14 | CD14 | IL1RL2 | JUNB | RHOU |
| | BIRC2 | CCL19 | CASP8 | CD274 | CXCL10 | IL1RN | LBP | RIPK1 |
| | BST2 | CCL22 | CCR7 | CD80 | CXCL11 | IL20RB | LCN2 | SAMHD1 |
| | CCL17 | CCL5 | CD274 | CD86 | CXCL13 | IL22 | LGALS9 | SERPINB2 |
| | CCL19 | CCR7 | CSF3 | CXCL10 | CXCL5 | IL23R | LIF | SOCS1 |
| | CCL22 | CD80 | DPYSL3 | CXCL11 | CXCL9 | IL27 | LRCH1 | SOCS2 |
| | CCL5 | CD86 | EIF2AK2 | CXCL13 | FER | IL2RA | LRRC4 | SOCS3 |
| | CD300LF | CDKN1A | HCLS1 | CXCL5 | IL12A | IL2RG | MUC1 | STAT2 |
| | CD44 | CISH | IKBKE | CXCL9 | IL23R | IL33 | MYC | STAT4 |
| | CD80 | CSF1 | IL1R1 | FER | IRAK3 | IRAK3 | NLRP12 | SYK |
| | CD86 | CSF3 | IL1RL2 | GFI1 | JUN | IRF2 | NUMBL | TANK |
| | CDKN1A | CTF1 | IL23R | IL12A | JUNB | IRF9 | OCSTAMP | TNF |
| | CISH | CXCL10 | IRAK3 | IL23R | JUND | ISG15 | OSM | TNFRSF11A |
| | CSF1 | DHX9 | ISG15 | IRAK3 | LBP | ITGAM | PDE1B | TNFRSF1B |
| | CSF3 | DPYSL3 | ITIH4 | JUN | LGALS9 | JAK3 | PIK3CD | TNFRSF9 |
| | CTF1 | DUOX2 | JAK3 | JUNB | MAPKAPK2 | JUNB | PTAFR | TNFSF13B |
| | CXCL10 | EREG | JUN | JUND | NGFR | LBP | PTPN13 | TNFSF14 |
| | CXCL11 | FER | JUNB | LBP | OTUD5 | LCN2 | PTPN2 | TNFSF15 |
| | CXCL13 | FOS | JUND | LGALS9 | PTPN22 | LEPR | PTPN23 | TNFSF4 |
| | CXCL5 | FOXC1 | LGALS9 | MAPKAPK2 | SELE | LIF | RHOU | TNFSF8 |
| | CXCL9 | FOXO3 | MAPKAPK2 | NGFR | TIRAP | LRRC4 | RIPK1 | TNFSF9 |
| | DUOX2 | FSCN1 | MYD88 | NR1I3 | TNFRSF11A | LTA | SBNO2 | TNIP2 |
| | EBI3 | GAS6 | NLRP12 | OTUD5 | TNFRSF1B | MUC1 | SELPLG | TRIM26 |
| | EREG | GRB2 | PML | PTPN22 | TNFRSF9 | MYC | SOCS1 | UBB |
| | FER | HAS2 | PTPN13 | SBNO2 | TRIB1 | MYD88 | SOCS2 | VAV1 |
| | FOS | HCK | PTPN2 | SCARB1 | TRIM6 | NFKBIA | SOCS3 | VCAM1 |
| | FOXC1 | HCLS1 | PTPN23 | SELE | | NGFR | SOX9 | XAF1 |
| | FOXO3 | HMOX1 | REL | TIRAP | | NOD1 | STAT2 | |
| | FSCN1 | IFNK | RIPK1 | TNFRSF11A | | NOD2 | STAT4 | |
| | GPR35 | IL10RA | SELE | TNFRSF1B | | NUMBL | TANK | |
| | GRB2 | IL12A | SHMT2 | TNFRSF9 | | OAS2 | TBK1 | |
| | HCK | IL12B | TBK1 | TNFSF4 | | OAS3 | TNF | |
| | HFE | IL17F | TNFRSF11A | TNIP1 | | OSM | TNFRSF1B | |
| | HMOX1 | IL1A | TRAFD1 | TNIP2 | | PDGFB | TRIM6 | |
| | IFNG | IL1R1 | TRIM56 | TNIP3 | | PIK3CD | UBB | |
| | IFNK | IL1RL2 | XAF1 | TRIB1 | | PML | VAV1 | |
| | IL10RA | IL1RN | | TRIM6 | | PRKCD | VCAM1 | |
| | IL12A | IL20RB | | ZC3H12A | | PSMA4 | ZC3H12A | |
| | | | | | | PSMA5 | ZFP36L1 | |
| | | | | | | PSMA7 | ZFP36L2 | |

Table S13. RNA-seq of blood of P. leucopus and M. musculus with or without LPS and presented as log10 normalized unique reads

| name | genus | treatment | Alox5 | Arg1 | Gapdh | Ibsp | IL1rn | Lcn2 | Nos2 | Slpi | Sod2 | Tgfb |
|---|---|---|---|---|---|---|---|---|---|---|---|---|
| 102 | Mus | control | 1.80 | 0.00 | 3.26 | 1.04 | 1.53 | 1.85 | 0.38 | 1.96 | 2.18 | 2.53 |
| 106 | Mus | control | 2.78 | 2.76 | 3.82 | 1.50 | 1.63 | 2.33 | 0.65 | 3.92 | 2.66 | 2.91 |
| 115 | Mus | control | 2.33 | 0.00 | 3.36 | 1.02 | 1.98 | 2.60 | 0.00 | 2.38 | 2.20 | 2.68 |
| 116 | Mus | control | 2.22 | 0.00 | 3.49 | 1.40 | 1.77 | 2.25 | 0.00 | 2.29 | 2.38 | 2.79 |
| 120 | Mus | control | 2.01 | 0.30 | 3.33 | 0.85 | 1.67 | 2.15 | 0.00 | 1.96 | 2.41 | 2.68 |
| F1_01 | Mus | control | 1.63 | 0.05 | 3.32 | 0.89 | 1.30 | 2.14 | 0.00 | 1.60 | 2.24 | 2.64 |
| F3_02 | Mus | control | 1.91 | 0.00 | 3.39 | 1.19 | 1.47 | 2.22 | 0.00 | 2.03 | 2.27 | 2.59 |
| M4_04 | Mus | control | 2.51 | 0.00 | 3.54 | 0.81 | 2.01 | 2.92 | 0.00 | 2.58 | 2.29 | 2.72 |
| 101 | Mus | LPS | 2.00 | 0.00 | 3.28 | 0.81 | 2.99 | 4.01 | 2.48 | 2.15 | 2.80 | 2.51 |
| 103 | Mus | LPS | 1.69 | 0.00 | 3.05 | 0.53 | 2.87 | 3.64 | 2.31 | 2.24 | 2.66 | 2.17 |
| 104 | Mus | LPS | 1.33 | 0.00 | 3.13 | 0.97 | 2.74 | 3.58 | 2.18 | 1.75 | 2.62 | 2.23 |
| 105 | Mus | LPS | 1.52 | 0.00 | 3.12 | 0.80 | 2.91 | 3.77 | 2.38 | 1.97 | 2.64 | 2.20 |
| 107 | Mus | LPS | 1.63 | 0.33 | 3.28 | 0.03 | 2.94 | 3.96 | 2.72 | 2.16 | 2.87 | 2.49 |
| 108 | Mus | LPS | 1.56 | 0.00 | 3.29 | 0.53 | 2.94 | 3.91 | 2.59 | 1.78 | 2.82 | 2.61 |
| 109 | Mus | LPS | 1.36 | 0.00 | 3.04 | 0.82 | 2.67 | 3.67 | 2.27 | 1.58 | 2.62 | 2.24 |
| 118 | Mus | LPS | 1.70 | 0.00 | 3.16 | 0.03 | 3.05 | 3.92 | 2.43 | 1.97 | 2.82 | 2.28 |
| F3_03 | Mus | LPS | 1.26 | 0.00 | 3.15 | 1.03 | 2.81 | 3.62 | 2.09 | 2.09 | 2.69 | 2.48 |
| M2_05 | Mus | LPS | 1.94 | 0.00 | 3.44 | 1.28 | 3.08 | 4.15 | 2.86 | 2.20 | 3.01 | 2.50 |
| M4_06 | Mus | LPS | 2.08 | 0.00 | 3.31 | 0.65 | 2.85 | 3.86 | 2.43 | 2.16 | 2.83 | 2.34 |
| M4_07 | Mus | LPS | 1.74 | 0.00 | 3.19 | 0.04 | 2.78 | 3.80 | 2.39 | 1.77 | 2.78 | 2.38 |
| Mus | LPS v control | | | | | | | | | | | |
| | fold change | | 0.32 | 0.44 | 0.58 | 0.35 | 16.35 | 32.89 | 198.65 | 0.44 | 2.73 | 0.48 |
| | t test | | 3.0E-03 | 2.1E-01 | 2.4E-03 | 1.2E-02 | 1.7E-11 | 7.2E-11 | 2.3E-14 | 1.2E-01 | 1.3E-06 | 6.1E-05 |
| | FDR | | 4.3E-03 | 2.1E-01 | 4.0E-03 | 1.5E-02 | 8.5E-11 | 2.4E-10 | 2.3E-13 | 1.3E-01 | 3.3E-06 | 1.2E-04 |
| 22275 | Peromyscus | control | 1.96 | 2.97 | 3.67 | 0.21 | 1.61 | 1.85 | 0.00 | 0.76 | 2.72 | 2.75 |
| 22277 | Peromyscus | control | 2.32 | 2.80 | 3.88 | 0.61 | 1.75 | 1.77 | 0.00 | 0.39 | 2.75 | 2.98 |
| 22279 | Peromyscus | control | 1.87 | 3.05 | 3.77 | 0.68 | 1.76 | 1.64 | 0.00 | 0.68 | 2.67 | 2.79 |
| 22282 | Peromyscus | control | 2.50 | 3.38 | 3.72 | 0.01 | 2.11 | 1.73 | 0.00 | 0.31 | 3.06 | 2.97 |
| 22315 | Peromyscus | control | 2.09 | 2.98 | 3.97 | 0.00 | 1.56 | 1.43 | 0.00 | 0.32 | 2.70 | 2.84 |
| 22321 | Peromyscus | control | 1.89 | 2.78 | 3.74 | 0.75 | 1.59 | 2.01 | 0.00 | 1.42 | 2.59 | 2.69 |
| 22335 | Peromyscus | control | 2.59 | 3.44 | 4.02 | 0.70 | 2.21 | 1.93 | 0.00 | 1.32 | 3.12 | 3.23 |
| 22341 | Peromyscus | control | 2.24 | 2.76 | 3.67 | 1.02 | 1.70 | 2.68 | 0.00 | 1.48 | 2.52 | 2.67 |
| 22258 | Peromyscus | LPS | 2.72 | 4.03 | 3.94 | 3.76 | 3.22 | 4.49 | 0.27 | 3.73 | 4.09 | 3.18 |
| 22278 | Peromyscus | LPS | 3.19 | 4.30 | 4.13 | 3.70 | 3.36 | 4.92 | 0.00 | 4.27 | 4.35 | 3.40 |
| 22281 | Peromyscus | LPS | 3.30 | 4.45 | 4.18 | 4.10 | 3.71 | 4.86 | 0.00 | 4.24 | 4.35 | 3.43 |
| 22284 | Peromyscus | LPS | 3.13 | 4.40 | 4.27 | 3.91 | 3.70 | 4.76 | 0.30 | 4.02 | 4.51 | 3.70 |
| 22289 | Peromyscus | LPS | 3.13 | 4.42 | 4.18 | 4.10 | 3.46 | 4.98 | 0.00 | 4.28 | 4.48 | 3.51 |
| 22292 | Peromyscus | LPS | 3.05 | 4.19 | 4.01 | 4.00 | 3.32 | 4.90 | 0.26 | 4.22 | 4.34 | 3.35 |
| 22316 | Peromyscus | LPS | 3.00 | 4.35 | 4.17 | 3.84 | 3.55 | 4.77 | 0.32 | 4.23 | 4.37 | 3.30 |
| 22319 | Peromyscus | LPS | 3.34 | 4.38 | 4.15 | 3.77 | 3.50 | 4.96 | 0.00 | 4.15 | 4.39 | 3.40 |
| 22325 | Peromyscus | LPS | 2.90 | 4.23 | 4.03 | 3.83 | 3.58 | 4.73 | 0.00 | 4.02 | 4.30 | 3.39 |
| 22327 | Peromyscus | LPS | 2.73 | 3.82 | 3.92 | 3.22 | 2.99 | 4.28 | 0.00 | 3.56 | 3.89 | 2.80 |
| 22332 | Peromyscus | LPS | 3.25 | 4.49 | 4.15 | 3.84 | 3.68 | 4.88 | 0.90 | 4.26 | 4.45 | 3.51 |
| 22339 | Peromyscus | LPS | 2.95 | 4.13 | 4.03 | 3.73 | 3.38 | 4.73 | 0.00 | 3.86 | 4.16 | 3.14 |
| Peromyscus | LPS v control | | | | | | | | | | | |
| | fold change | | 7.53 | 17.63 | 1.96 | 2086.18 | 46.30 | 784.37 | 1.49 | 1720.15 | 34.58 | 3.00 |
| | t test | | 1.7E-07 | 4.1E-10 | 3.9E-05 | 2.6E-15 | 3.8E-12 | 1.1E-14 | 9.1E-02 | 1.6E-13 | 1.0E-12 | 1.1E-04 |
| | FDR | | 2.4E-07 | 6.8E-10 | 4.9E-05 | 2.6E-14 | 7.6E-12 | 5.5E-14 | 9.1E-02 | 5.3E-13 | 2.5E-12 | 1.2E-04 |

Table S14. RNA-seq of spleen of P. leucopus and M. musculus with or without LPS and presented as log10 of normalized unique reads

| animal | genus | treatment | Alox5 | Arg1 | Ccl2 | Cd86 | Csf2 | Csf3 | Cxcl11 | Gapdh | Gata3 | Hmox1 | Ibsp | Ido1 | Ifnb1 |
|---|---|---|---|---|---|---|---|---|---|---|---|---|---|---|---|
| 102 | Mus | control | 1.51 | 0.00 | 2.06 | 2.98 | 1.37 | 1.37 | 1.10 | 4.59 | 2.11 | 3.95 | 0.00 | 0.91 | 0.00 |
| 106 | Mus | control | 2.21 | 0.46 | 1.44 | 3.07 | 1.21 | 0.58 | 1.12 | 4.51 | 2.41 | 4.17 | 0.00 | 0.98 | 0.00 |
| 115 | Mus | control | 2.48 | 0.81 | 1.33 | 3.02 | 0.03 | 0.81 | 0.73 | 4.52 | 2.15 | 4.22 | 0.00 | 0.98 | 0.00 |
| 116 | Mus | control | 2.09 | 1.06 | 1.52 | 2.96 | 1.18 | 0.88 | 0.88 | 4.53 | 2.31 | 4.15 | 0.00 | 0.88 | 0.00 |
| 120 | Mus | control | 2.03 | 0.86 | 1.36 | 3.09 | 0.92 | 0.71 | 1.01 | 4.49 | 2.45 | 4.20 | 0.00 | 1.13 | 0.00 |
| F1_01 | Mus | control | 1.86 | 1.23 | 1.74 | 3.03 | 0.86 | 0.69 | 0.69 | 4.53 | 2.38 | 4.32 | 0.00 | 0.93 | 0.00 |
| F3_02 | Mus | control | 1.90 | 1.08 | 1.47 | 2.97 | 0.70 | 0.96 | 0.91 | 4.52 | 2.52 | 4.16 | 0.00 | 1.21 | 0.00 |
| M4_04 | Mus | control | 2.86 | 0.81 | 1.39 | 2.74 | 0.33 | 0.00 | 0.73 | 4.52 | 1.84 | 4.11 | 0.00 | 0.81 | 0.00 |
| 101 | Mus | LPS | 2.04 | 0.64 | 3.67 | 3.62 | 1.75 | 3.21 | 3.19 | 4.51 | 2.06 | 3.88 | 0.00 | 1.98 | 0.34 |
| 103 | Mus | LPS | 1.84 | 0.62 | 3.70 | 3.77 | 1.93 | 3.22 | 3.11 | 4.47 | 2.13 | 3.73 | 0.00 | 1.87 | 0.00 |
| 104 | Mus | LPS | 1.81 | 0.51 | 3.85 | 3.72 | 2.08 | 3.22 | 3.01 | 4.46 | 2.05 | 3.83 | 0.00 | 1.51 | 1.44 |
| 105 | Mus | LPS | 1.97 | 0.88 | 3.76 | 3.65 | 2.07 | 3.24 | 3.18 | 4.51 | 1.92 | 3.85 | 0.00 | 1.91 | 0.00 |
| 107 | Mus | LPS | 2.28 | 0.63 | 3.49 | 3.49 | 2.01 | 2.91 | 2.75 | 4.53 | 1.84 | 3.96 | 0.00 | 1.45 | 0.00 |
| 108 | Mus | LPS | 1.78 | 0.69 | 3.90 | 3.60 | 2.28 | 3.20 | 3.34 | 4.53 | 1.81 | 3.86 | 0.00 | 1.54 | 0.69 |
| 109 | Mus | LPS | 1.95 | 0.87 | 3.57 | 3.66 | 2.15 | 3.02 | 3.07 | 4.47 | 2.03 | 3.75 | 0.00 | 1.64 | 0.00 |
| 118 | Mus | LPS | 1.82 | 0.68 | 3.60 | 3.59 | 1.74 | 3.07 | 2.90 | 4.52 | 2.20 | 3.86 | 0.00 | 1.60 | 1.03 |
| F3_03 | Mus | LPS | 1.62 | 0.79 | 3.69 | 3.70 | 1.89 | 3.16 | 3.12 | 4.48 | 1.97 | 3.79 | 0.00 | 1.79 | 0.92 |
| M2_05 | Mus | LPS | 2.56 | 1.20 | 3.56 | 3.36 | 1.86 | 2.76 | 2.84 | 4.58 | 1.79 | 3.96 | 0.00 | 1.26 | 0.00 |
| M4_06 | Mus | LPS | 2.18 | 0.96 | 3.40 | 3.21 | 2.04 | 2.81 | 2.51 | 4.44 | 1.84 | 3.74 | 0.00 | 1.38 | 1.10 |
| M4_07 | Mus | LPS | 2.16 | 0.91 | 3.68 | 3.37 | 2.29 | 2.98 | 2.61 | 4.50 | 1.79 | 3.79 | 0.00 | 1.10 | 0.56 |
| Mus | LPS v control | | | | | | | | | | | | | | |
| | fold change | | 0.77 | 0.99 | 130.88 | 3.79 | 15.22 | 207.38 | 117.79 | 0.94 | 0.48 | 0.47 | na | 4.04 | 3.21 |
| | t test | | 4.5E-01 | 9.7E-01 | 3.1E-15 | 9.7E-08 | 2.0E-07 | 3.9E-13 | 7.7E-14 | 1.2E-01 | 1.0E-03 | 2.1E-07 | na | 1.4E-05 | 1.4E-02 |
| | FDR | | 5.3E-01 | 9.7E-01 | 6.0E-14 | 2.7E-07 | 5.2E-07 | 2.2E-12 | 5.0E-13 | 1.5E-01 | 1.6E-03 | 5.1E-07 | na | 2.6E-05 | 2.2E-02 |
| 22275 | Pero | control | 2.20 | 2.67 | 2.10 | 2.93 | 1.16 | 0.00 | 2.00 | 4.50 | 2.77 | 4.56 | 0.00 | 1.98 | 0.00 |
| 22277 | Pero | control | 2.37 | 2.12 | 2.11 | 3.07 | 0.92 | 0.00 | 1.17 | 4.49 | 2.69 | 4.23 | 0.00 | 2.13 | 0.32 |
| 22279 | Pero | control | 2.18 | 2.83 | 1.97 | 2.97 | 0.20 | 0.00 | 1.56 | 4.52 | 2.80 | 4.50 | 0.85 | 2.17 | 0.00 |
| 22282 | Pero | control | 2.63 | 2.65 | 2.13 | 3.05 | 0.60 | 0.00 | 1.11 | 4.55 | 2.76 | 4.35 | 0.00 | 2.03 | 0.00 |
| 22315 | Pero | control | 2.17 | 2.43 | 1.90 | 2.93 | 0.26 | 0.00 | 1.39 | 4.61 | 2.75 | 4.44 | 0.26 | 1.81 | 0.26 |
| 22321 | Pero | control | 2.28 | 2.80 | 2.16 | 3.15 | 0.93 | 0.00 | 1.39 | 4.54 | 2.65 | 4.18 | 0.28 | 2.26 | 0.28 |
| 22335 | Pero | control | 2.25 | 2.69 | 2.11 | 2.99 | 0.90 | 0.24 | 1.17 | 4.51 | 2.77 | 4.22 | 0.24 | 1.77 | 0.00 |
| 22341 | Pero | control | 2.18 | 2.46 | 2.25 | 3.07 | 0.71 | 0.47 | 1.45 | 4.48 | 2.88 | 4.38 | 0.17 | 1.97 | 0.00 |
| 22258 | Pero | LPS | 2.14 | 3.11 | 3.71 | 2.88 | 3.03 | 2.45 | 4.08 | 4.44 | 2.43 | 4.70 | 2.44 | 3.20 | 1.98 |
| 22278 | Pero | LPS | 2.44 | 3.44 | 3.66 | 3.29 | 3.04 | 2.20 | 3.82 | 4.50 | 2.51 | 4.52 | 2.64 | 2.86 | 2.04 |
| 22281 | Pero | LPS | 2.26 | 3.20 | 3.55 | 3.21 | 2.67 | 2.49 | 4.12 | 4.46 | 2.35 | 4.47 | 2.61 | 3.20 | 2.54 |
| 22284 | Pero | LPS | 2.29 | 3.37 | 3.81 | 3.38 | 2.48 | 2.66 | 4.18 | 4.53 | 2.51 | 4.45 | 2.74 | 3.42 | 1.56 |
| 22289 | Pero | LPS | 2.54 | 3.15 | 3.06 | 3.15 | 3.28 | 2.27 | 3.59 | 4.51 | 2.59 | 5.05 | 2.60 | 3.04 | 1.65 |
| 22292 | Pero | LPS | 2.18 | 3.12 | 3.28 | 3.23 | 3.06 | 2.44 | 3.60 | 4.48 | 2.57 | 4.58 | 2.43 | 3.14 | 2.12 |
| 22316 | Pero | LPS | 2.14 | 3.41 | 3.65 | 3.32 | 3.12 | 2.79 | 4.15 | 4.45 | 2.47 | 4.48 | 2.83 | 3.17 | 2.32 |
| 22319 | Pero | LPS | 2.39 | 3.67 | 3.30 | 3.20 | 3.34 | 2.20 | 3.69 | 4.51 | 2.46 | 4.74 | 2.78 | 3.04 | 2.32 |
| 22325 | Pero | LPS | 2.14 | 3.20 | 3.56 | 3.21 | 2.64 | 2.34 | 4.02 | 4.48 | 2.32 | 4.62 | 2.45 | 3.05 | 2.05 |
| 22327 | Pero | LPS | 2.08 | 3.22 | 3.28 | 2.83 | 2.99 | 2.34 | 3.63 | 4.49 | 2.25 | 4.84 | 2.52 | 2.99 | 1.32 |
| 22332 | Pero | LPS | 2.57 | 3.63 | 3.50 | 3.30 | 3.05 | 2.33 | 4.15 | 4.49 | 2.43 | 4.78 | 2.73 | 3.37 | 2.52 |
| 22339 | Pero | LPS | 2.47 | 3.56 | 3.26 | 3.28 | 2.68 | 2.40 | 3.51 | 4.56 | 2.61 | 4.64 | 2.86 | 2.85 | 1.76 |
| Peromyscus | LPS v control | | | | | | | | | | | | | | |
| | fold change | | 1.05 | 5.73 | 23.78 | 1.48 | 172.70 | 208.88 | 298.56 | 0.92 | 0.50 | 1.99 | 258.07 | 12.46 | 81.05 |
| | t test | | 8.0E-01 | 3.8E-07 | 5.3E-12 | 1.5E-02 | 3.1E-12 | 1.5E-16 | 1.0E-13 | 5.8E-02 | 2.3E-06 | 8.3E-04 | 2.1E-15 | 5.4E-11 | 9.0E-11 |
| | FDR | | 8.0E-01 | 7.1E-07 | 1.7E-11 | 1.9E-02 | 1.1E-11 | 2.0E-15 | 4.6E-13 | 6.4E-02 | 3.9E-06 | 1.2E-03 | 1.7E-14 | 1.6E-10 | 2.5E-10 |

| animal | genus | treatment | Ifng | IL10 | IL12 | IL13 | IL1a | IL1b | Il1r | IL1rn | IL17a | IL4 | IL5 | IL6 |
|---|---|---|---|---|---|---|---|---|---|---|---|---|---|---|
| 102 | Mus | control | 1.03 | 0.73 | 2.40 | 0.00 | 1.79 | 2.86 | 1.18 | 2.06 | 0.00 | 1.39 | 0.00 | 0.91 |
| 106 | Mus | control | 0.76 | 1.21 | 2.62 | 0.00 | 2.04 | 3.07 | 2.54 | 1.85 | 0.00 | 1.02 | 0.00 | 0.76 |
| 115 | Mus | control | 1.07 | 1.31 | 2.52 | 0.00 | 2.26 | 3.03 | 2.55 | 2.12 | 0.00 | 0.81 | 0.00 | 0.98 |
| 116 | Mus | control | 1.31 | 1.50 | 2.83 | 0.41 | 2.16 | 2.94 | 2.54 | 2.04 | 0.00 | 0.58 | 0.00 | 1.18 |
| 120 | Mus | control | 1.09 | 0.92 | 2.66 | 0.00 | 2.23 | 2.81 | 2.55 | 2.04 | 0.00 | 0.71 | 0.00 | 0.79 |
| F1_01 | Mus | control | 1.23 | 1.17 | 2.66 | 0.00 | 2.21 | 2.71 | 2.44 | 2.09 | 0.00 | 1.17 | 0.00 | 0.93 |
| F3_02 | Mus | control | 1.40 | 1.26 | 2.75 | 0.00 | 2.00 | 2.75 | 2.49 | 1.83 | 0.00 | 0.85 | 0.00 | 1.08 |
| M4_04 | Mus | control | 1.14 | 0.99 | 2.41 | 0.00 | 2.15 | 2.94 | 1.91 | 2.00 | 0.00 | 0.81 | 0.00 | 1.21 |
| 101 | Mus | LPS | 2.67 | 2.27 | 2.78 | 0.00 | 3.35 | 4.02 | 2.82 | 3.77 | 0.48 | 1.24 | 0.00 | 2.88 |
| 103 | Mus | LPS | 2.69 | 2.60 | 2.79 | 0.32 | 3.18 | 4.15 | 2.80 | 3.94 | 0.30 | 1.28 | 0.00 | 3.17 |
| 104 | Mus | LPS | 2.13 | 2.38 | 2.72 | 0.93 | 2.91 | 4.00 | 2.83 | 3.81 | 0.48 | 0.03 | 0.00 | 3.41 |
| 105 | Mus | LPS | 2.70 | 2.32 | 2.72 | 0.00 | 2.97 | 4.14 | 2.46 | 3.82 | 0.30 | 0.95 | 0.00 | 3.09 |
| 107 | Mus | LPS | 2.12 | 2.08 | 2.59 | 0.03 | 2.98 | 4.05 | 2.65 | 3.80 | 0.00 | 0.73 | 0.00 | 2.56 |
| 108 | Mus | LPS | 2.71 | 2.08 | 2.66 | 0.00 | 3.24 | 4.21 | 2.50 | 3.75 | 0.00 | 1.20 | 0.00 | 3.20 |
| 109 | Mus | LPS | 2.56 | 2.04 | 2.64 | 0.00 | 3.07 | 3.93 | 2.79 | 3.77 | 0.30 | 0.62 | 0.00 | 2.81 |
| 118 | Mus | LPS | 2.05 | 2.16 | 2.59 | 0.00 | 2.90 | 3.93 | 2.48 | 3.64 | 0.00 | 0.76 | 0.00 | 2.70 |
| F3_03 | Mus | LPS | 2.49 | 2.32 | 2.59 | 0.00 | 3.14 | 3.95 | 2.74 | 3.79 | 0.30 | 0.79 | 0.00 | 2.96 |
| M2_05 | Mus | LPS | 1.86 | 1.95 | 2.60 | 0.00 | 3.06 | 4.06 | 2.52 | 3.76 | 0.00 | 0.69 | 0.00 | 2.60 |
| M4_06 | Mus | LPS | 1.26 | 1.84 | 2.45 | 0.00 | 2.86 | 3.98 | 2.46 | 3.49 | 0.00 | 0.66 | 0.00 | 2.44 |
| M4_07 | Mus | LPS | 1.62 | 1.99 | 2.46 | 0.00 | 2.47 | 3.99 | 1.84 | 3.22 | 0.00 | 1.07 | 0.00 | 2.52 |
| Mus | LPS v control | | | | | | | | | | | | | |
| | fold change | | 12.88 | 10.78 | 1.06 | 1.14 | 8.03 | 13.97 | 1.99 | 50.89 | 1.51 | 0.83 | na | 75.85 |
| | t test | | 8.1E-06 | 9.2E-09 | 6.6E-01 | 6.0E-01 | 1.1E-08 | 6.4E-15 | 9.8E-02 | 8.7E-15 | 2.0E-02 | 5.8E-01 | na | 6.4E-12 |
| | FDR | | 1.7E-05 | 3.6E-08 | 7.2E-01 | 6.7E-01 | 3.9E-08 | 8.3E-14 | 1.3E-01 | 8.5E-14 | 2.9E-02 | 6.7E-01 | na | 3.1E-11 |
| 22275 | Pero | control | 0.26 | 1.88 | 1.42 | 0.00 | 1.48 | 3.17 | 3.37 | 1.86 | 0.00 | 1.16 | 0.00 | 1.19 |
| 22277 | Pero | control | 0.92 | 2.13 | 1.30 | 0.00 | 1.77 | 3.04 | 3.31 | 2.04 | 0.00 | 0.32 | 0.00 | 1.30 |
| 22279 | Pero | control | 0.67 | 1.98 | 1.33 | 0.00 | 1.75 | 3.21 | 3.34 | 1.86 | 0.00 | 0.20 | 0.00 | 1.29 |
| 22282 | Pero | control | 0.47 | 1.92 | 1.60 | 0.00 | 1.58 | 3.01 | 3.18 | 2.01 | 0.60 | 0.00 | 0.00 | 1.36 |
| 22315 | Pero | control | 0.91 | 2.09 | 1.52 | 0.00 | 1.45 | 2.91 | 3.16 | 2.05 | 0.00 | 0.26 | 0.00 | 1.19 |
| 22321 | Pero | control | 0.00 | 1.88 | 1.72 | 0.00 | 1.28 | 3.39 | 3.25 | 1.98 | 0.00 | 0.58 | 0.00 | 1.64 |
| 22335 | Pero | control | 1.24 | 1.95 | 1.49 | 0.00 | 1.76 | 3.12 | 3.25 | 1.82 | 0.00 | 0.24 | 0.00 | 1.32 |
| 22341 | Pero | control | 0.47 | 1.93 | 1.38 | 0.00 | 1.36 | 2.98 | 3.44 | 1.72 | 0.30 | 0.17 | 0.00 | 1.49 |
| 22258 | Pero | LPS | 1.99 | 3.25 | 1.37 | 0.37 | 2.13 | 4.12 | 3.40 | 3.36 | 0.30 | 0.00 | 0.00 | 3.11 |
| 22278 | Pero | LPS | 1.86 | 3.41 | 1.15 | 0.60 | 2.09 | 4.17 | 3.41 | 3.51 | 0.00 | 0.00 | 0.00 | 2.98 |
| 22281 | Pero | LPS | 1.64 | 3.30 | 1.16 | 0.64 | 2.18 | 4.01 | 3.53 | 3.39 | 0.60 | 0.00 | 0.00 | 3.02 |
| 22284 | Pero | LPS | 1.31 | 3.24 | 0.92 | 0.22 | 2.24 | 4.12 | 3.69 | 3.70 | 0.00 | 0.00 | 0.00 | 2.89 |
| 22289 | Pero | LPS | 2.25 | 3.63 | 1.15 | 0.00 | 2.23 | 4.19 | 3.43 | 3.66 | 0.00 | 0.00 | 0.00 | 2.55 |
| 22292 | Pero | LPS | 1.70 | 3.55 | 1.42 | 0.00 | 2.05 | 3.92 | 3.62 | 3.67 | 0.00 | 0.00 | 0.00 | 3.05 |
| 22316 | Pero | LPS | 2.52 | 3.36 | 1.61 | 0.00 | 2.36 | 4.13 | 3.63 | 3.69 | 0.30 | 0.21 | 0.00 | 3.15 |
| 22319 | Pero | LPS | 1.99 | 3.68 | 1.17 | 0.00 | 2.57 | 4.22 | 3.49 | 3.68 | 0.00 | 0.00 | 0.00 | 2.98 |
| 22325 | Pero | LPS | 2.03 | 3.52 | 1.02 | 0.36 | 2.66 | 4.18 | 3.46 | 3.66 | 0.00 | 0.36 | 0.00 | 2.98 |
| 22327 | Pero | LPS | 1.43 | 3.40 | 1.09 | 0.00 | 2.14 | 3.91 | 3.39 | 3.46 | 0.00 | 0.00 | 0.00 | 2.83 |
| 22332 | Pero | LPS | 2.11 | 3.60 | 1.24 | 0.00 | 2.35 | 4.05 | 3.53 | 3.66 | 0.60 | 0.00 | 0.00 | 2.97 |
| 22339 | Pero | LPS | 0.92 | 3.36 | 1.31 | 0.00 | 2.17 | 4.09 | 3.55 | 3.69 | 0.70 | 0.27 | 0.00 | 2.84 |
| Peromyscus | LPS v control | | | | | | | | | | | | | |
| | fold change | | 15.65 | 29.68 | 0.56 | 1.52 | 5.13 | 9.75 | 1.66 | 47.38 | 1.25 | 0.51 | na | 39.62 |
| | t test | | 8.4E-06 | 2.7E-15 | 4.6E-03 | 5.6E-02 | 1.9E-07 | 1.2E-12 | 9.5E-05 | 8.9E-17 | 4.3E-01 | 1.7E-02 | na | 1.3E-14 |
| | FDR | | 1.3E-05 | 1.8E-14 | 6.3E-03 | 6.4E-02 | 3.7E-07 | 4.5E-12 | 1.4E-04 | 1.8E-15 | 4.4E-01 | 2.1E-02 | na | 7.6E-14 |

| animal | genus | treatment | Irf5 | Lcn2 | Mrc1 | Mt_Co1 | Nfe2l2 | Nfkb | Nos2 | Ptx3 | Slpi | Sod2 | Stat1 | Stat2 | Stat3 | Stat4 | Tgfb1 | Tnf |
|---|---|---|---|---|---|---|---|---|---|---|---|---|---|---|---|---|---|---|
| 102 | Mus | control | 3.44 | 3.05 | 2.85 | 5.45 | 3.12 | 4.02 | 1.83 | 1.03 | 3.61 | 3.28 | 3.37 | 2.71 | 2.71 | 3.15 | 4.02 | 2.32 |
| 106 | Mus | control | 3.72 | 3.11 | 3.66 | 5.69 | 3.52 | 3.99 | 2.24 | 0.00 | 3.47 | 3.24 | 3.60 | 3.44 | 3.51 | 3.35 | 3.85 | 2.49 |
| 115 | Mus | control | 3.56 | 3.72 | 3.69 | 5.58 | 3.51 | 3.77 | 2.07 | 0.00 | 3.47 | 3.21 | 3.52 | 3.33 | 3.38 | 3.02 | 3.80 | 2.34 |
| 116 | Mus | control | 3.66 | 2.87 | 3.46 | 5.85 | 3.49 | 3.90 | 2.08 | 0.00 | 3.45 | 3.21 | 3.58 | 3.34 | 3.47 | 3.19 | 3.86 | 2.55 |
| 120 | Mus | control | 3.68 | 2.83 | 3.63 | 5.76 | 3.49 | 3.87 | 2.22 | 0.00 | 3.33 | 3.21 | 3.54 | 3.39 | 3.49 | 3.18 | 3.82 | 2.55 |
| F1_01 | Mus | control | 3.61 | 2.76 | 3.75 | 5.64 | 3.51 | 3.86 | 2.09 | 0.00 | 3.38 | 3.26 | 3.59 | 3.35 | 3.40 | 3.12 | 3.85 | 2.28 |
| F3_02 | Mus | control | 3.72 | 2.69 | 3.48 | 5.74 | 3.50 | 3.93 | 2.00 | 0.00 | 3.28 | 3.22 | 3.59 | 3.36 | 3.46 | 3.27 | 3.85 | 2.56 |
| M4_04 | Mus | control | 3.34 | 4.26 | 3.44 | 5.61 | 3.45 | 3.53 | 1.97 | 0.33 | 3.67 | 3.29 | 3.21 | 3.01 | 3.02 | 2.81 | 3.85 | 2.22 |
| 101 | Mus | LPS | 3.69 | 4.16 | 3.34 | 5.70 | 3.58 | 4.15 | 2.72 | 2.15 | 3.40 | 4.04 | 4.30 | 4.25 | 4.02 | 3.35 | 3.59 | 3.37 |
| 103 | Mus | LPS | 3.72 | 4.08 | 3.29 | 5.77 | 3.59 | 4.21 | 2.76 | 2.47 | 3.39 | 4.04 | 4.32 | 4.23 | 4.13 | 3.63 | 3.61 | 3.29 |
| 104 | Mus | LPS | 3.69 | 3.96 | 3.37 | 5.76 | 3.56 | 4.19 | 2.82 | 2.41 | 3.38 | 3.93 | 4.20 | 4.24 | 4.16 | 3.67 | 3.65 | 3.33 |
| 105 | Mus | LPS | 3.69 | 4.14 | 3.28 | 5.65 | 3.54 | 4.13 | 2.67 | 2.34 | 3.51 | 4.11 | 4.18 | 4.08 | 3.88 | 3.44 | 3.66 | 3.41 |
| 107 | Mus | LPS | 3.59 | 4.49 | 3.30 | 5.61 | 3.60 | 4.08 | 2.89 | 1.87 | 3.54 | 4.02 | 4.24 | 4.17 | 3.83 | 3.33 | 3.70 | 3.39 |
| 108 | Mus | LPS | 3.61 | 4.35 | 3.14 | 5.58 | 3.63 | 4.06 | 2.86 | 1.29 | 3.37 | 4.05 | 4.21 | 4.07 | 3.76 | 3.20 | 3.59 | 3.47 |
| 109 | Mus | LPS | 3.68 | 4.05 | 3.25 | 5.73 | 3.55 | 4.11 | 2.65 | 1.73 | 3.26 | 3.91 | 4.35 | 4.34 | 4.02 | 3.38 | 3.55 | 3.43 |
| 118 | Mus | LPS | 3.68 | 4.11 | 3.17 | 5.60 | 3.52 | 4.10 | 2.38 | 1.29 | 3.39 | 3.99 | 4.25 | 4.14 | 3.87 | 3.38 | 3.61 | 3.39 |
| F3_03 | Mus | LPS | 3.70 | 3.97 | 3.36 | 5.78 | 3.54 | 4.09 | 2.70 | 1.88 | 3.43 | 3.98 | 4.28 | 4.26 | 3.96 | 3.46 | 3.57 | 3.36 |
| M2_05 | Mus | LPS | 3.54 | 4.62 | 3.20 | 5.62 | 3.65 | 4.03 | 3.00 | 1.48 | 3.55 | 4.03 | 4.22 | 4.16 | 3.75 | 3.02 | 3.72 | 3.50 |
| M4_06 | Mus | LPS | 3.55 | 4.13 | 3.11 | 5.61 | 3.45 | 3.96 | 2.50 | 1.01 | 3.51 | 3.94 | 4.02 | 3.93 | 3.54 | 3.13 | 3.65 | 3.43 |
| M4_07 | Mus | LPS | 3.69 | 4.39 | 2.56 | 5.43 | 3.28 | 4.21 | 2.52 | 1.43 | 3.71 | 4.02 | 3.98 | 3.64 | 3.34 | 3.08 | 3.78 | 3.49 |
| Mus | LPS v control | | | | | | | | | | | | | | | | | |
| | fold change | | 1.15 | 11.04 | 0.50 | 0.98 | 1.24 | 1.78 | 4.41 | 40.63 | 0.99 | 5.84 | 5.15 | 7.63 | 3.56 | 1.60 | 0.60 | 9.78 |
| | t test | | 2.0E-01 | 1.1E-05 | 1.6E-02 | 8.5E-01 | 8.5E-02 | 1.2E-04 | 7.2E-08 | 2.9E-07 | 9.4E-01 | 1.2E-17 | 2.0E-10 | 4.5E-08 | 1.8E-04 | 2.9E-02 | 7.7E-07 | 2.2E-14 |
| | FDR | | 2.4E-01 | 2.1E-05 | 2.4E-02 | 9.0E-01 | 1.1E-01 | 2.1E-04 | 2.2E-07 | 6.7E-07 | 9.6E-01 | 4.7E-16 | 8.7E-10 | 1.5E-07 | 3.1E-04 | 4.0E-02 | 1.7E-06 | 1.7E-13 |
| 22275 | Pero | control | 3.82 | 3.14 | 4.04 | 5.59 | 3.62 | 3.89 | 1.36 | 2.20 | 1.21 | 3.37 | 4.08 | 3.69 | 3.87 | 2.92 | 3.84 | 2.54 |
| 22277 | Pero | control | 3.71 | 3.16 | 4.04 | 5.67 | 3.61 | 3.83 | 0.87 | 1.71 | 0.80 | 3.32 | 4.03 | 3.56 | 3.82 | 3.02 | 3.81 | 2.66 |
| 22279 | Pero | control | 3.87 | 3.39 | 4.13 | 5.66 | 3.60 | 3.91 | 1.28 | 1.79 | 0.94 | 3.37 | 4.04 | 3.64 | 3.85 | 3.00 | 3.84 | 2.56 |
| 22282 | Pero | control | 3.80 | 3.09 | 4.07 | 5.64 | 3.63 | 3.92 | 1.63 | 1.25 | 1.11 | 3.37 | 4.06 | 3.67 | 3.85 | 3.12 | 3.86 | 2.65 |
| 22315 | Pero | control | 3.76 | 3.03 | 4.24 | 5.57 | 3.60 | 3.80 | 0.73 | 1.82 | 0.80 | 3.38 | 4.38 | 3.84 | 3.81 | 3.12 | 3.77 | 2.50 |
| 22321 | Pero | control | 3.79 | 3.19 | 4.05 | 5.62 | 3.65 | 3.87 | 1.82 | 1.61 | 1.28 | 3.39 | 4.11 | 3.61 | 3.81 | 3.07 | 3.76 | 2.67 |
| 22335 | Pero | control | 3.81 | 3.26 | 4.02 | 5.65 | 3.66 | 3.92 | 1.67 | 1.65 | 0.85 | 3.30 | 4.22 | 3.69 | 3.90 | 3.02 | 3.84 | 2.61 |
| 22341 | Pero | control | 3.91 | 3.07 | 4.14 | 5.65 | 3.63 | 4.03 | 1.60 | 1.68 | 0.91 | 3.27 | 4.05 | 3.82 | 3.87 | 3.07 | 3.85 | 2.57 |
| 22258 | Pero | LPS | 3.46 | 3.64 | 4.05 | 5.48 | 3.97 | 4.09 | 1.57 | 4.37 | 2.96 | 4.57 | 4.53 | 4.21 | 4.14 | 2.90 | 3.85 | 3.57 |
| 22278 | Pero | LPS | 3.65 | 3.99 | 3.95 | 5.54 | 3.99 | 4.29 | 2.12 | 4.31 | 3.41 | 4.60 | 4.57 | 4.20 | 4.20 | 2.89 | 3.86 | 3.55 |
| 22281 | Pero | LPS | 3.35 | 3.75 | 3.84 | 5.52 | 4.06 | 4.23 | 1.88 | 4.20 | 3.08 | 4.56 | 4.52 | 4.25 | 4.21 | 3.00 | 3.89 | 3.47 |
| 22284 | Pero | LPS | 3.57 | 4.01 | 3.81 | 5.57 | 4.03 | 4.41 | 1.62 | 4.12 | 3.14 | 4.61 | 4.55 | 4.24 | 4.31 | 3.12 | 3.87 | 3.85 |
| 22289 | Pero | LPS | 3.61 | 3.81 | 4.06 | 5.51 | 3.99 | 4.17 | 2.84 | 3.87 | 2.97 | 4.58 | 4.54 | 4.22 | 4.20 | 2.83 | 3.84 | 3.58 |
| 22292 | Pero | LPS | 3.58 | 3.80 | 3.97 | 5.52 | 4.05 | 4.27 | 2.79 | 4.17 | 3.03 | 4.61 | 4.62 | 4.32 | 4.31 | 2.91 | 3.88 | 3.70 |
| 22316 | Pero | LPS | 3.52 | 3.91 | 3.97 | 5.54 | 4.03 | 4.36 | 1.62 | 4.52 | 3.18 | 4.66 | 4.62 | 4.37 | 4.36 | 2.88 | 3.84 | 3.70 |
| 22319 | Pero | LPS | 3.47 | 4.38 | 4.07 | 5.54 | 4.02 | 4.18 | 2.00 | 3.85 | 3.40 | 4.64 | 4.47 | 4.12 | 4.15 | 2.70 | 3.93 | 3.67 |
| 22325 | Pero | LPS | 3.47 | 3.87 | 3.71 | 5.50 | 3.96 | 4.17 | 2.05 | 3.89 | 3.04 | 4.57 | 4.49 | 4.15 | 4.18 | 2.89 | 3.86 | 3.52 |
| 22327 | Pero | LPS | 3.23 | 3.76 | 4.29 | 5.36 | 3.77 | 3.96 | 2.94 | 3.80 | 3.04 | 4.50 | 4.28 | 4.08 | 4.03 | 2.64 | 3.61 | 3.35 |
| 22332 | Pero | LPS | 3.59 | 4.06 | 3.90 | 5.67 | 4.08 | 4.27 | 2.94 | 4.09 | 3.42 | 4.60 | 4.56 | 4.25 | 4.22 | 2.91 | 3.94 | 3.58 |
| 22339 | Pero | LPS | 3.49 | 4.10 | 3.93 | 5.71 | 3.95 | 4.22 | 1.87 | 4.07 | 3.21 | 4.59 | 4.56 | 4.24 | 4.26 | 3.03 | 3.82 | 3.41 |
| Peromyscus | LPS v control | | | | | | | | | | | | | | | | | |
| | fold change | | 0.49 | 5.73 | 0.74 | 0.81 | 2.33 | 2.09 | 6.59 | 246.78 | 148.60 | 17.52 | 2.54 | 3.40 | 2.33 | 0.71 | 1.07 | 9.61 |
| | t test | | 2.6E-06 | 1.5E-08 | 3.8E-02 | 1.1E-02 | 2.9E-10 | 2.1E-06 | 1.7E-03 | 2.1E-14 | 5.9E-16 | 7.0E-23 | 6.4E-08 | 9.8E-11 | 1.7E-09 | 7.5E-03 | 3.9E-01 | 2.3E-13 |
| | FDR | | 4.3E-06 | 3.2E-08 | 4.5E-02 | 1.4E-02 | 7.0E-10 | 3.7E-06 | 2.4E-03 | 1.1E-13 | 6.0E-15 | 2.9E-21 | 1.3E-07 | 2.5E-10 | 3.9E-09 | 9.9E-03 | 4.1E-01 | 9.4E-13 |

Table S15. RNA-seq of liver of P. leucopus and M. musculus with or without LPS and presented as log10 normalized unique reads

| name | genus | treatment | Alb | Arg1 | Crp | Gapdh | Hp | MT_Cox1 | Nos2 | Ptx3 | Saa3 | Sod2 | Tlr2 | Tlr4 |
|---|---|---|---|---|---|---|---|---|---|---|---|---|---|---|
| 102 | Mus | control | 6.40 | 4.65 | 3.44 | 4.59 | 5.14 | 6.27 | 0.00 | 0.00 | 3.10 | 3.65 | 1.85 | 1.60 |
| 106 | Mus | control | 6.20 | 4.67 | 3.63 | 4.77 | 5.46 | 6.04 | 0.30 | 0.00 | 4.08 | 3.75 | 1.61 | 0.85 |
| 115 | Mus | control | 6.20 | 4.66 | 3.31 | 4.80 | 5.47 | 6.49 | 0.00 | 0.00 | 3.59 | 3.76 | 1.81 | 0.90 |
| 116 | Mus | control | 6.45 | 4.78 | 3.69 | 4.83 | 5.45 | 6.18 | 0.30 | 0.00 | 3.37 | 3.84 | 1.69 | 1.30 |
| 120 | Mus | control | 6.37 | 4.62 | 3.65 | 4.67 | 5.27 | 6.34 | 0.60 | 0.00 | 3.06 | 3.85 | 1.83 | 1.40 |
| F1_01 | Mus | control | 6.29 | 4.61 | 3.20 | 4.58 | 5.20 | 6.26 | 0.00 | 0.00 | 3.32 | 3.71 | 1.90 | 1.23 |
| F3_02 | Mus | control | 6.29 | 4.45 | 3.22 | 4.57 | 5.30 | 6.12 | 0.30 | 0.00 | 3.17 | 3.79 | 1.86 | 1.34 |
| M4_04 | Mus | control | 6.42 | 4.74 | 3.53 | 4.81 | 5.73 | 6.26 | 0.00 | 0.60 | 4.35 | 3.80 | 2.16 | 1.20 |
| 101 | Mus | LPS | 6.23 | 4.50 | 3.85 | 4.55 | 6.07 | 6.29 | 2.61 | 1.70 | 5.97 | 4.36 | 3.76 | 1.23 |
| 103 | Mus | LPS | 6.22 | 4.43 | 3.75 | 4.48 | 5.97 | 6.32 | 2.84 | 0.00 | 5.83 | 4.22 | 3.83 | 1.11 |
| 104 | Mus | LPS | 6.23 | 4.36 | 4.02 | 4.47 | 5.92 | 6.03 | 2.80 | 0.00 | 5.76 | 4.20 | 4.14 | 1.11 |
| 105 | Mus | LPS | 6.12 | 4.18 | 3.90 | 4.47 | 6.04 | 6.25 | 2.79 | 0.00 | 5.85 | 4.30 | 3.90 | 1.54 |
| 107 | Mus | LPS | 6.06 | 4.36 | 3.88 | 4.58 | 5.92 | 6.21 | 2.73 | 0.00 | 5.72 | 4.10 | 3.80 | 1.72 |
| 108 | Mus | LPS | 6.09 | 4.40 | 4.28 | 4.60 | 5.90 | 6.16 | 3.07 | 0.60 | 5.74 | 4.18 | 4.05 | 1.18 |
| 109 | Mus | LPS | 6.29 | 4.62 | 4.36 | 4.74 | 5.90 | 6.17 | 3.26 | 0.00 | 5.77 | 4.21 | 4.13 | 1.08 |
| 118 | Mus | LPS | 6.18 | 4.51 | 4.05 | 4.49 | 5.89 | 6.13 | 2.49 | 0.48 | 5.71 | 4.19 | 3.99 | 1.34 |
| F3_03 | Mus | LPS | 6.22 | 4.41 | 3.88 | 4.43 | 5.87 | 6.17 | 2.67 | 0.85 | 5.78 | 4.17 | 4.01 | 1.36 |
| M2_05 | Mus | LPS | 5.95 | 4.32 | 4.11 | 4.56 | 5.87 | 6.28 | 2.85 | 2.21 | 5.67 | 4.05 | 3.87 | 1.82 |
| M4_06 | Mus | LPS | 6.30 | 4.56 | 4.36 | 4.67 | 5.76 | 5.93 | 3.06 | 0.00 | 5.62 | 4.07 | 4.03 | 1.65 |
| M4_07 | Mus | LPS | 6.37 | 4.69 | 4.46 | 4.78 | 5.81 | 6.34 | 2.96 | 0.90 | 5.66 | 4.20 | 4.12 | 1.86 |
| Mus | LPS v control | | | | | | | | | | | | | |
| | fold change | | 0.73 | 0.63 | 4.12 | 0.74 | 3.41 | 0.88 | 453.13 | 3.06 | 177.42 | 2.62 | 134.99 | 1.55 |
| | t test | | 1.3E-02 | 2.4E-03 | 9.5E-06 | 1.8E-02 | 8.5E-08 | 3.5E-01 | 7.0E-16 | 9.2E-02 | 3.9E-12 | 1.1E-09 | 2.1E-17 | 1.5E-01 |
| | FDR | | 2.0E-02 | 4.1E-03 | 1.9E-05 | 2.4E-02 | 2.0E-07 | 3.5E-01 | 4.2E-15 | 1.1E-01 | 1.6E-11 | 3.3E-09 | 2.5E-16 | 1.6E-01 |
| 22275 | Peromyscus | control | 6.41 | 4.66 | 3.58 | 4.92 | 5.31 | 6.03 | 0.00 | 0.60 | 3.39 | 3.63 | 2.55 | 2.00 |
| 22277 | Peromyscus | control | 6.53 | 4.51 | 3.91 | 4.96 | 5.50 | 6.12 | 0.00 | 0.78 | 2.85 | 3.68 | 2.72 | 1.86 |
| 22279 | Peromyscus | control | 6.54 | 4.74 | 3.63 | 5.00 | 5.33 | 6.14 | 0.00 | 0.00 | 2.92 | 3.65 | 2.77 | 1.97 |
| 22282 | Peromyscus | control | 6.53 | 4.55 | 3.98 | 4.94 | 5.42 | 6.11 | 0.00 | 0.48 | 2.77 | 3.71 | 3.04 | 2.08 |
| 22315 | Peromyscus | control | 6.41 | 4.58 | 3.75 | 4.93 | 5.28 | 6.00 | 0.00 | 0.30 | 2.74 | 3.55 | 2.75 | 2.04 |
| 22321 | Peromyscus | control | 6.54 | 4.80 | 3.86 | 5.00 | 5.73 | 6.24 | 0.00 | 0.00 | 3.24 | 3.77 | 2.71 | 2.05 |
| 22335 | Peromyscus | control | 6.52 | 4.67 | 3.85 | 4.89 | 5.42 | 6.09 | 0.00 | 1.20 | 2.85 | 3.68 | 2.93 | 2.03 |
| 22341 | Peromyscus | control | 6.56 | 4.83 | 3.83 | 5.00 | 5.38 | 6.25 | 0.00 | 0.60 | 2.94 | 3.65 | 2.97 | 1.88 |
| 22258 | Peromyscus | LPS | 6.44 | 4.73 | 4.25 | 5.01 | 5.81 | 6.16 | 1.30 | 2.59 | 5.82 | 4.56 | 3.21 | 2.08 |
| 22281 | Peromyscus | LPS | 6.41 | 4.71 | 4.28 | 4.90 | 5.93 | 6.06 | 1.00 | 2.72 | 5.83 | 4.60 | 3.17 | 2.03 |
| 22284 | Peromyscus | LPS | 6.27 | 4.50 | 4.35 | 4.90 | 5.84 | 6.34 | 1.60 | 2.21 | 5.85 | 4.51 | 3.15 | 2.22 |
| 22289 | Peromyscus | LPS | 6.59 | 4.60 | 4.76 | 4.83 | 5.75 | 6.13 | 1.32 | 1.91 | 5.81 | 4.55 | 3.23 | 2.16 |
| 22292 | Peromyscus | LPS | 6.38 | 4.40 | 4.16 | 4.77 | 5.77 | 5.95 | 1.49 | 2.25 | 5.74 | 4.44 | 2.74 | 1.88 |
| 22316 | Peromyscus | LPS | 6.44 | 4.73 | 4.31 | 4.85 | 5.77 | 6.06 | 2.00 | 3.01 | 5.76 | 4.51 | 3.15 | 2.01 |
| 22319 | Peromyscus | LPS | 6.24 | 4.56 | 3.63 | 4.79 | 5.86 | 6.38 | 1.18 | 2.36 | 5.63 | 4.40 | 2.83 | 1.95 |
| 22325 | Peromyscus | LPS | 6.35 | 4.85 | 4.07 | 4.96 | 5.77 | 6.14 | 1.49 | 2.34 | 5.70 | 4.48 | 3.01 | 1.99 |
| 22327 | Peromyscus | LPS | 6.42 | 4.90 | 4.32 | 4.71 | 5.66 | 5.98 | 1.15 | 2.92 | 5.57 | 4.53 | 2.99 | 1.72 |
| 22332 | Peromyscus | LPS | 6.31 | 4.81 | 4.08 | 4.87 | 5.56 | 6.03 | 0.48 | 2.64 | 5.58 | 4.31 | 3.13 | 2.03 |
| 22339 | Peromyscus | LPS | 6.49 | 4.82 | 4.17 | 4.81 | 5.83 | 6.19 | 1.86 | 2.69 | 5.65 | 4.35 | 3.12 | 1.57 |
| Peromyscus | LPS v control | | | | | | | | | | | | | |
| | fold change | | 22.45 | 12.30 | 17.12 | 10.32 | 33.25 | 20.58 | 17.33 | 90.56 | 1529.26 | 35.95 | 7.85 | 2.32 |
| | t test | | 1.2E-01 | 8.0E-02 | 2.1E-02 | 1.2E-01 | 3.9E-02 | 8.7E-02 | 2.4E-06 | 1.5E-07 | 1.6E-06 | 3.7E-03 | 2.1E-02 | 1.5E-01 |
| | FDR | | 3.4E-01 | 2.9E-01 | 1.0E-01 | 3.3E-01 | 1.5E-01 | 2.8E-01 | 2.8E-05 | 7.4E-06 | 1.1E-05 | 1.7E-02 | 8.8E-02 | 3.6E-01 |

Table S16. Targeted RNA-seq of spleen and blood of older P. leucopus with or without LPS treatment and presented as log10 normalized unique reads

| name | tissue | treatment | Alox5 | Arg1 | Ccl2 | Csf2 | Csf3 | Cxcl11 | Gapdh | Hmox1 | Ibsp | Ido1 | Ifnb1 | Ifng | IL13 | IL1a | IL1b | IL1rn | IL6 | IL10 | IL12 | IL22 | Lcn2 | MTcox1 | Nfe2l2 | Nos2 | Ptx3 | Slpi | Sod2 | Tgfb1 | Tnf |
|---|---|---|---|---|---|---|---|---|---|---|---|---|---|---|---|---|---|---|---|---|---|---|---|---|---|---|---|---|---|---|---|
| 20949 | spleen | control | 2.28 | 2.52 | 2.35 | 0.69 | 0.00 | 1.06 | 4.56 | 4.24 | 1.56 | 2.13 | 0.00 | 0.82 | 0.00 | 1.45 | 2.90 | 1.78 | 1.38 | 2.15 | 1.64 | 0.21 | 3.49 | 5.65 | 3.59 | 1.24 | 1.56 | 1.84 | 3.48 | 3.82 | 2.55 |
| 20694 | spleen | control | 1.99 | 2.05 | 2.21 | 0.76 | 0.29 | 0.46 | 4.52 | 4.05 | 0.00 | 2.31 | 0.00 | 1.45 | 0.29 | 1.79 | 3.03 | 1.82 | 1.50 | 1.90 | 1.63 | 0.00 | 2.94 | 5.78 | 3.60 | 1.71 | 1.16 | 1.97 | 3.38 | 3.94 | 2.77 |
| 20733 | spleen | control | 2.08 | 2.50 | 2.13 | 0.53 | 0.35 | 1.28 | 4.56 | 4.25 | 0.00 | 2.06 | 0.00 | 1.45 | 0.00 | 1.51 | 3.19 | 1.65 | 1.41 | 2.19 | 1.57 | 0.00 | 3.30 | 5.74 | 3.65 | 1.56 | 1.41 | 0.35 | 3.35 | 3.82 | 2.41 |
| 20807 | spleen | control | 2.23 | 2.16 | 2.11 | 0.92 | 0.00 | 1.01 | 4.60 | 3.82 | 0.57 | 2.21 | 0.27 | 1.27 | 0.00 | 1.89 | 3.30 | 1.99 | 1.56 | 1.76 | 1.68 | 0.00 | 2.92 | 5.93 | 3.59 | 2.15 | 1.62 | 1.80 | 3.33 | 3.84 | 2.71 |
| 20688 | spleen | LPS | 1.97 | 3.04 | 3.67 | 3.09 | 2.33 | 3.28 | 4.57 | 4.81 | 2.20 | 3.25 | 1.31 | 1.61 | 0.00 | 2.27 | 4.09 | 3.54 | 2.98 | 3.34 | 1.08 | 0.00 | 3.84 | 5.65 | 3.94 | 1.70 | 3.82 | 2.95 | 4.53 | 3.97 | 3.13 |
| 20702 | spleen | LPS | 2.02 | 3.15 | 3.14 | 3.07 | 2.92 | 3.35 | 4.47 | 4.47 | 2.27 | 3.16 | 2.00 | 1.62 | 0.00 | 2.26 | 4.03 | 3.59 | 2.97 | 3.39 | 1.51 | 1.71 | 3.51 | 5.58 | 3.91 | 2.44 | 4.05 | 3.01 | 4.45 | 3.90 | 3.42 |
| 20695 | spleen | LPS | 2.22 | 3.42 | 3.87 | 3.45 | 3.23 | 3.71 | 4.45 | 4.38 | 2.90 | 3.18 | 2.33 | 1.95 | 0.00 | 2.72 | 4.37 | 3.63 | 3.27 | 3.07 | 1.28 | 2.81 | 4.07 | 5.54 | 4.04 | 1.99 | 4.04 | 3.24 | 4.59 | 3.94 | 3.56 |
| 20819 | spleen | LPS | 2.32 | 3.67 | 3.54 | 2.59 | 2.69 | 3.30 | 4.52 | 4.58 | 3.01 | 3.47 | 1.25 | 1.42 | 0.33 | 2.12 | 4.24 | 3.46 | 2.68 | 3.44 | 1.25 | 0.72 | 4.25 | 5.51 | 3.93 | 2.31 | 3.99 | 3.74 | 4.55 | 3.92 | 2.99 |
| 20483 | spleen | LPS | 2.08 | 3.16 | 3.52 | 1.91 | 3.20 | 3.43 | 4.65 | 4.02 | 1.16 | 3.29 | 0.76 | 2.70 | 0.00 | 2.03 | 4.21 | 3.51 | 3.02 | 3.01 | 1.42 | 3.14 | 3.13 | 5.88 | 3.80 | 1.99 | 3.54 | 3.03 | 4.39 | 4.18 | 2.69 |
| 20732 | spleen | LPS | 1.96 | 3.51 | 3.87 | 3.31 | 2.90 | 3.27 | 4.45 | 4.65 | 2.52 | 2.96 | 2.43 | 2.31 | 0.00 | 2.22 | 4.43 | 3.66 | 3.36 | 3.19 | 1.08 | 2.76 | 3.70 | 5.60 | 4.06 | 1.92 | 4.31 | 3.41 | 4.55 | 3.91 | 3.38 |
| fold-change | | | 0.89 | 10.4 | 25.1 | 150 | 522 | 274 | 0.91 | 2.47 | 64.9 | 10.9 | 41.1 | 4.89 | 0.96 | 4.07 | 13.3 | 56.7 | 38.2 | 17.3 | 0.44 | 63.7 | 3.87 | 0.71 | 2.19 | 2.47 | 333 | 54.6 | 13.4 | 1.30 | 3.83 |
| t test | | | 6.0E-01 | 1.9E-04 | 1.1E-05 | 8.0E-05 | 4.8E-07 | 3.9E-07 | 3.6E-01 | 4.0E-02 | 3.6E-03 | 5.0E-06 | 1.6E-03 | 3.7E-02 | 8.5E-01 | 3.3E-03 | 4.9E-06 | 5.4E-09 | 1.7E-06 | 7.3E-06 | 4.3E-03 | 2.4E-02 | 3.6E-02 | 1.1E-01 | 1.4E-04 | 9.1E-02 | 2.0E-07 | 9.0E-04 | 8.5E-09 | 8.6E-02 | 1.1E-02 |
| 20501 | blood | control | 3.06 | 4.33 | | | | | 4.12 | | 1.80 | | | | | | | 2.99 | | | | | 3.33 | | | 0.00 | | 2.22 | 3.82 | 3.42 | |
| 20613 | blood | control | 2.80 | 3.90 | | | | | 4.08 | | 1.55 | | | | | | | 2.70 | | | | | 2.74 | | | 0.00 | | 1.92 | 3.35 | 3.29 | |
| 20696 | blood | control | 2.51 | 3.79 | | | | | 4.08 | | 1.20 | | | | | | | 2.72 | | | | | 2.65 | | | 0.00 | | 1.79 | 3.32 | 3.32 | |
| 20704 | blood | control | 2.63 | 3.81 | | | | | 4.01 | | 2.02 | | | | | | | 2.23 | | | | | 3.52 | | | 0.00 | | 2.53 | 3.51 | 3.04 | |
| 20495 | blood | LPS | 3.32 | 4.65 | | | | | 4.36 | | 4.15 | | | | | | | 3.80 | | | | | 5.30 | | | 0.00 | | 4.47 | 4.70 | 3.82 | |
| 20713 | blood | LPS | 3.46 | 4.71 | | | | | 4.50 | | 4.40 | | | | | | | 3.89 | | | | | 5.38 | | | 0.00 | | 4.68 | 4.75 | 3.70 | |
| 20732 | blood | LPS | 3.15 | 4.55 | | | | | 4.20 | | 4.32 | | | | | | | 3.93 | | | | | 4.80 | | | 0.00 | | 4.16 | 4.44 | 3.34 | |
| 20483 | blood | LPS | 3.56 | 4.91 | | | | | 4.61 | | 4.53 | | | | | | | 4.11 | | | | | 5.51 | | | 0.48 | | 4.87 | 4.94 | 3.87 | |
| fold-change | | | 4.21 | 5.58 | | | | | 2.22 | | 511 | | | | | | | 18.8 | | | | | 155 | | | 1.3 | | 270 | 16.1 | 2.6 | |
| t test | | | 5.8E-03 | 2.3E-03 | | | | | 9.0E-03 | | 8.3E-06 | | | | | | | 2.9E-04 | | | | | 1.7E-04 | | | 3.6E-01 | | 3.5E-05 | 2.3E-04 | 2.7E-02 | |

Table S14. Systemic infection of *P. leucopus* with *Borrelia hermsii* MTW

| Animal | | 21019 | 20958 | 21030 | 20851 | 20921 | 20961 | 20846 | 20923 | fold change | t test |
|---|---|---|---|---|---|---|---|---|---|---|---|
| Infection | | no | no | no | yes | yes | yes | yes | yes | | |
| Age (weeks) | | 10 | 15 | 9 | 23 | 17 | 14 | 24 | 18 | | |
| Body weight (g) | | 22.5 | 16.7 | 19.4 | 25.9 | 18.4 | 19.3 | 23.5 | 27.6 | 1.2 | 0.22 |
| % spleen weight / body weight | | 0.119 | 0.161 | 0.180 | 0.702 | 0.318 | 0.369 | 0.290 | 0.269 | 1.7 | 0.04 |
| Log10 bacteria copies spleen (qPCR) | | 0.00 | 0.00 | 0.00 | 5.44 | 2.77 | 4.33 | 4.29 | 3.84 | na | na |
| Product | Gene | Log10 normalized unique reads | | | | | | | | | |
| Arginase 1 | Arg1 | 3.12 | 2.36 | 2.97 | 2.45 | 3.47 | 3.04 | 3.86 | 3.35 | 2.6 | 0.26 |
| Glyceraldehyde 3-P dehydrogenase | Gapdh | 3.78 | 3.73 | 3.76 | 3.62 | 3.93 | 3.88 | 4.33 | 4.06 | 1.6 | 0.15 |
| Integrin-binding sialoprotein | Ibsp | 0.00 | 0.33 | 0.73 | 2.10 | 2.15 | 2.68 | 3.66 | 3.11 | 243.8 | 0.0006 |
| Interleukin-1 receptor antagonist | Il1rn | 1.99 | 1.64 | 2.27 | 1.21 | 2.94 | 2.54 | 3.52 | 2.96 | 4.7 | 0.18 |
| Lipocalin 2 | Lcn2 | 2.03 | 1.41 | 2.36 | 2.25 | 2.67 | 3.48 | 4.20 | 3.62 | 20.7 | 0.026 |
| Nitric oxide synthase 2 | Nos2 | 0.00 | 0.00 | 0.00 | 0.00 | 0.00 | 0.00 | 0.05 | 0.00 | na | na |
| Secretory leukocyte peptidase inhibitor | Slpi | 0.00 | 0.03 | 0.26 | 1.28 | 1.93 | 2.37 | 2.99 | 2.45 | 129.0 | 0.0012 |
| Superoxide dismutase 2 | Sod2 | 2.73 | 2.23 | 2.59 | 3.07 | 3.40 | 3.59 | 4.06 | 3.71 | 11.2 | 0.0038 |
| Transforming growth factor beta | Tgfb | 2.98 | 2.91 | 3.21 | 2.73 | 3.46 | 3.38 | 3.72 | 3.41 | 2.0 | 0.15 |

Table S18. Log2 fold-changes DEGs of blood of LPS-treated P. leucopus and Borrelia hermsii infected P. leucopus

| accession | product | log2 LPS fold change | log2 infection fold change |
|---|---|---|---|
| XM_028880204.1 | serum amyloid A-3 protein (LOC114700540), transcript variant X2, mRNA | 10.048 | 10.882 |
| XM_028868627.1 | secretory leukocyte peptidase inhibitor (Slpi), mRNA | 10.003 | 8.914 |
| XM_028867455.1 | integrin binding sialoprotein (Ibsp), mRNA | 11.159 | 8.307 |
| XM_028859809.1 | formyl peptide receptor 1 (Fpr1), transcript variant X1, mRNA | 7.170 | 6.410 |
| XM_028890975.1 | matrix metallopeptidase 8 (Mmp8), mRNA | 9.300 | 6.159 |
| XM_028881893.1 | C-X-C motif chemokine 2-like (LOC114701735), mRNA | 10.271 | 6.345 |
| XM_028864652.1 | aconitate decarboxylase 1 (Acod1), mRNA | 7.111 | 5.882 |
| XM_028884808.1 | lipocalin 2 (Lcn2), transcript variant X2, mRNA | 8.960 | 5.205 |
| XM_028860285.1 | serum amyloid A-5 protein (LOC114685652), transcript variant X2, mRNA | 10.570 | 4.871 |
| XM_028890551.1 | C-C motif chemokine 3 (LOC114707732), mRNA | 6.929 | 4.355 |
| XM_028860034.1 | lactotransferrin (Ltf), mRNA | 8.214 | 4.149 |
| XM_028868677.1 | interleukin 1 receptor antagonist (Il1rn), transcript variant X1, mRNA | 6.553 | 4.019 |
| XM_028865841.1 | interleukin 1 receptor type 2 (Il1r2), mRNA | 7.172 | 3.790 |
| XM_028866678.1 | superoxide dismutase 2 (Sod2), mRNA | 5.363 | 3.763 |
| XM_028891075.1 | haptoglobin (Hp), mRNA | 5.341 | 3.120 |
| XM_028854887.1 | heme oxygenase 1 (Hmox1), mRNA | 3.518 | 2.868 |
| XM_028872948.1 | suppressor of cytokine signaling 1 (Socs1), mRNA | 5.122 | 2.451 |
| XM_028869645.1 | lysozyme C-2-like (LOC114693317), mRNA | 2.152 | 1.952 |
| XM_028877403.1 | arginase 1 (Arg1), mRNA | 4.373 | 1.723 |
| XM_028893401.1 | interleukin 1 beta (Il1b), mRNA | 4.092 | 1.674 |
| XM_028864037.1 | arachidonate 5-lipoxygenase (Alox5), mRNA | 3.158 | 1.313 |
| XM_028893355.1 | dual specificity phosphatase 1 (Dusp1), mRNA | 2.833 | -2.456 |
| XM_028887322.1 | interleukin 13 receptor subunit alpha 1 (Il13ra1), mRNA | 4.349 | 2.488 |
| XM_028878069.1 | arachidonate 5-lipoxygenase activating protein (Alox5ap), transcript variant X1, mRNA | 4.173 | 2.265 |
| XM_028893102.1 | C-X-C motif chemokine receptor 2 (Cxcr2), mRNA | 3.951 | 1.238 |
| XM_028887864.1 | toll like receptor 4 (Tlr4), mRNA | 3.441 | 2.040 |
| XM_028889246.1 | lipopolysaccharide binding protein (Lbp), mRNA | 3.274 | 2.479 |
| XM_028892294.1 | glyceraldehyde-3-phosphate dehydrogenase (Gapdh), mRNA | 1.304 | 0.918 |
| XM_028880203.1 | serum amyloid A-3 protein (LOC114700540), transcript variant X1, mRNA | 9.901 | 8.882 |
| XM_028860282.1 | serum amyloid A-3 protein-like (LOC114685651), transcript variant X1, mRNA | 9.575 | 6.528 |
| XM_028894435.1 | serpin family E member 1 (Serpine1), mRNA | 9.470 | 1.666 |
| XM_028884807.1 | lipocalin 2 (Lcn2), transcript variant X1, mRNA | 8.773 | 5.894 |
| XM_028860286.1 | serum amyloid A-5 protein (LOC114685652), transcript variant X3, mRNA | 8.515 | 9.491 |
| XM_028884227.1 | cobalamin binding intrinsic factor-like (LOC114703403), mRNA | 8.345 | 5.321 |
| XM_028889615.1 | sphingosine kinase 1 (Sphk1), transcript variant X4, mRNA | 8.273 | 3.533 |
| XM_028879198.1 | olfactomedin 4 (Olfm4), mRNA | 8.179 | 3.125 |
| XM_028887051.1 | cathelicidin antimicrobial peptide (Camp), mRNA | 7.909 | 4.322 |
| XM_028884226.1 | cobalamin binding intrinsic factor-like (LOC114703402), mRNA | 7.644 | 4.485 |
| XM_028863845.1 | solute carrier family 28 member 3 (Slc28a3), mRNA | 7.484 | 4.690 |
| XM_028869381.1 | growth arrest specific 7 (Gas7), transcript variant X3, mRNA | 7.236 | 6.108 |
| XM_028862085.1 | CD177 antigen-like (LOC114687425), mRNA | 7.208 | 4.046 |
| XM_028883061.1 | hes family bHLH transcription factor 3 (Hes3), mRNA | 7.163 | 5.227 |
| XM_028880026.1 | TNFAIP3 interacting protein 1 (Tnip1), transcript variant X5, mRNA | 7.162 | 5.928 |
| XM_028859718.1 | leucine rich alpha-2-glycoprotein 1 (Lrg1), mRNA | 7.139 | 3.028 |
| XM_028871479.1 | metallothionein-2 (LOC114694511), mRNA | 7.096 | -0.133 |
| XM_028859810.1 | formyl peptide receptor 1 (Fpr1), transcript variant X2, mRNA | 7.046 | 7.607 |
| XM_028859812.1 | formyl peptide receptor 2 (Fpr2), mRNA | 7.031 | 5.171 |
| XR_003735432.1 | metallothionein-1 pseudogene (LOC114698905), misc_RNA | 6.666 | 1.626 |
| XM_028892197.1 | transglutaminase 1 (Tgm1), mRNA | 6.626 | 4.174 |
| XM_028861133.1 | protein RoBo-1-like (LOC114686457), mRNA | 6.583 | 4.864 |
| XM_028890521.1 | protein Wfdc21 (LOC114707706), mRNA | 6.542 | 3.133 |
| XM_028871478.1 | metallothionein-1 (LOC114694510), mRNA | 6.460 | 1.164 |
| XM_028888721.1 | smoothelin like 2 (Smtnl2), mRNA | 5.970 | 3.747 |
| XM_028884304.1 | membrane spanning 4-domains A3 (Ms4a3), mRNA | 5.824 | 5.902 |
| XM_028873144.1 | annexin A3 (Anxa3), mRNA | 5.716 | 3.504 |
| XM_028863798.1 | growth arrest and DNA damage inducible gamma (Gadd45g), mRNA | 5.562 | 2.939 |
| XM_028886872.1 | neutrophilic granule protein-like (LOC114705177), mRNA | 5.461 | 3.089 |
| XM_028857368.1 | resistin (Retn), mRNA | 5.368 | 3.420 |
| XM_028882108.1 | serpin family B member 1 (Serpinb1), mRNA | 5.364 | 3.494 |
| XM_028876612.1 | chitinase 3 like 1 (Chi3l1), mRNA | 5.322 | 2.609 |
| XM_028868478.1 | matrix metallopeptidase 9 (Mmp9), transcript variant X1, mRNA | 5.296 | 2.366 |
| XM_028892932.1 | bone marrow stromal cell antigen 1 (Bst1), transcript variant X2, mRNA | 5.177 | 3.115 |
| XM_028874824.1 | myristoylated alanine rich protein kinase C substrate (Marcks), mRNA | 5.164 | 3.077 |
| XM_028893309.1 | CCAAT enhancer binding protein delta (Cebpd), mRNA | 5.132 | 3.312 |
| XM_028853955.1 | matrix metallopeptidase 25 (Mmp25), mRNA | 5.110 | 2.849 |
| XM_028856166.1 | peptidoglycan recognition protein 1 (Pglyrp1), mRNA | 5.092 | 2.815 |
| XM_028878785.1 | serine dehydratase (Sds), mRNA | 5.089 | 3.123 |
| XM_028875171.1 | interleukin 1 receptor accessory protein (Il1rap), transcript variant X4, mRNA | 5.016 | 1.800 |
| XM_028888075.1 | thrombospondin 1 (Thbs1), mRNA | 5.008 | 2.571 |
| XM_028890852.1 | protein S100-A8-like (LOC114707940), mRNA | 4.922 | 2.598 |
| XM_028857366.1 | mast cell expressed membrane protein 1 (Mcemp1), transcript variant X1, mRNA | 4.871 | 2.775 |
| XM_028877491.1 | diacylglycerol O-acyltransferase 2 (Dgat2), transcript variant X1, mRNA | 4.868 | 3.462 |
| XM_028892931.1 | bone marrow stromal cell antigen 1 (Bst1), transcript variant X1, mRNA | 4.847 | 1.270 |
| XM_028885765.1 | hydroxycarboxylic acid receptor 2 (Hcar2), mRNA | 4.828 | 2.398 |
| XM_028893766.1 | annexin A1 (Anxa1), mRNA | 4.761 | 2.326 |
| XM_028864324.1 | Fc fragment of IgG receptor IIb (Fcgr2b), transcript variant X1, mRNA | 4.702 | 3.974 |
| XM_028864329.1 | low affinity immunoglobulin gamma Fc region receptor IV-like (LOC114689891), mRNA | 4.695 | 3.203 |
| XM_028890855.1 | S100 calcium binding protein A9 (S100a9), mRNA | 4.687 | 2.645 |
| XM_028864327.1 | Fc fragment of IgG receptor IIb (Fcgr2b), transcript variant X4, mRNA | 4.645 | 3.499 |
| XM_028889627.1 | CD300 molecule like family member f (Cd300lf), transcript variant X2, mRNA | 4.639 | 2.953 |
| XM_028892581.1 | suppressor of cytokine signaling 3 (Socs3), mRNA | 4.617 | 2.951 |
| XM_028863733.1 | hexokinase 3 (Hk3), transcript variant X2, mRNA | 4.581 | 2.643 |
| XM_028854779.1 | 2'-5'-oligoadenylate synthase-like protein 2 (LOC114681329), transcript variant X1, mRNA | 4.543 | 0.260 |
| XM_028882006.1 | dehydrogenase/reductase 9 (Dhrs9), transcript variant X1, mRNA | 4.526 | 1.074 |
| XM_028863283.1 | triggering receptor expressed on myeloid cells 1-like (LOC114688764), mRNA | 4.500 | 2.876 |
| XM_028865982.1 | phospholipid scramblase 1 (LOC114690951), transcript variant X1, mRNA | 4.492 | 1.230 |
| XM_028885750.1 | cytochrome b reductase 1 (LOC114704488), mRNA | 4.481 | 3.468 |
| XM_028857367.1 | mast cell expressed membrane protein 1 (Mcemp1), transcript variant X2, mRNA | 4.428 | 3.382 |
| XM_028859919.1 | PML-RARA regulated adaptor molecule 1 (Pram1), mRNA | 4.405 | 1.695 |
| XM_028853958.1 | methionine sulfoxide reductase B1 (Msrb1), mRNA | 4.405 | 2.308 |
| XM_028867272.1 | placenta-specific gene 8 protein (LOC114691764), mRNA | 4.382 | 3.322 |
| XM_028894001.1 | glutaredoxin (Glrx), transcript variant X2, mRNA | 4.369 | 2.148 |
| XM_028888907.1 | tumor protein p53 inducible nuclear protein 2 (Tp53inp2), transcript varian | 4.353 | 1.229 |
| XM_028864484.1 | selectin P (Selp), mRNA | 4.350 | 3.998 |
| XM_028871068.1 | colony stimulating factor 2 receptor beta common subunit (Csf2rb), mRNA | 4.344 | 2.684 |
| XM_028864044.1 | Ras association domain family member 4 (Rassf4), transcript variant X2, mRNA | 4.341 | 3.863 |
| XM_028872738.1 | prokineticin 2 (Prok2), mRNA | 4.337 | 2.533 |
| XM_028885699.1 | multiple EGF like domains 9 (Megf9), mRNA | 4.323 | 2.485 |
| XR_003736535.1 | uncharacterized LOC114706499 (LOC114706499), transcript variant X2, ncRNA | 4.312 | 3.983 |
| XR_003736536.1 | uncharacterized LOC114706499 (LOC114706499), transcript variant X3, ncRNA | 4.303 | 3.002 |
| XM_028858293.1 | sialic acid-binding Ig-like lectin 12 (LOC114683921), mRNA | 4.301 | 2.432 |
| XM_028857815.1 | solute carrier family 2, facilitated glucose transporter member 3 (LOC114683484), transcript variant X1, mRNA | 4.282 | 1.784 |
| XM_028859123.1 | PAF1 homolog, Paf1/RNA polymerase II complex component (Paf1), mRNA | 4.280 | 2.478 |
| XM_028859947.1 | trem-like transcript 4 protein (LOC114685334), mRNA | 4.246 | 2.571 |
| XM_028864328.1 | Fc fragment of IgG receptor IIb (Fcgr2b), transcript variant X5, mRNA | 4.244 | 2.724 |
| XM_028891008.1 | uridine phosphorylase 1 (Upp1), transcript variant X1, mRNA | 4.221 | 10.244 |
| XM_028886190.1 | serglycin (Srgn), mRNA | 4.207 | 2.307 |
| XM_028892179.1 | leukotriene B4 receptor (Ltb4r), mRNA | 4.205 | 2.455 |
| XM_028893898.1 | plasminogen activator, urokinase receptor (Plaur), mRNA | 4.203 | 2.965 |
| XM_028864326.1 | Fc fragment of IgG receptor IIb (Fcgr2b), transcript variant X3, mRNA | 4.196 | 3.462 |
| XM_028885177.1 | tyrosine-protein phosphatase non-receptor type substrate 1-like (LOC1147041 | 4.189 | 2.984 |
| XM_028861705.1 | immediate early response 3 (Ier3), mRNA | 4.169 | 2.320 |
| XM_028868566.1 | beta-1,4-galactosyltransferase 5 (B4galt5), transcript variant X1, mRNA | 4.151 | 7.386 |
| XM_028871069.1 | cytokine receptor common subunit beta-like (LOC114694224), mRNA | 4.132 | 2.357 |
| XM_028857817.1 | solute carrier family 2, facilitated glucose transporter member 3 (LOC114683484), transcript variant X3, mRNA | 4.093 | 2.155 |
| XM_028871551.1 | adhesion G protein-coupled receptor G3 (Adgrg3), mRNA | 4.091 | 2.197 |
| XM_028863790.1 | nuclear factor, interleukin 3 regulated (Nfil3), transcript variant X1, mRNA | 4.090 | 2.104 |
| XM_028868678.1 | interleukin 1 receptor antagonist (Il1rn), transcript variant X2, mRNA | 4.080 | 7.897 |
| XM_028886840.1 | CD14 molecule (Cd14), mRNA | 4.044 | 0.200 |
| XM_028871067.1 | neutrophil cytosolic factor 4 (Ncf4), mRNA | 3.982 | 2.220 |
| XM_028880027.1 | TNFAIP3 interacting protein 1 (Tnip1), transcript variant X6, mRNA | 3.957 | 3.236 |
| XM_028893634.1 | angiotensin II receptor associated protein (Agtrap), mRNA | 3.951 | 2.117 |
| XM_028871982.1 | proline-serine-threonine phosphatase interacting protein 2 (Pstpip2), mRNA | 3.890 | 2.219 |
| XM_028879941.1 | ETS proto-oncogene 2, transcription factor (Ets2), mRNA | 3.836 | 2.231 |
| XM_028862398.1 | STEAP4 metalloreductase (Steap4), mRNA | 3.818 | 0.720 |
| XM_028871544.1 | liver carboxylesterase 1-like (LOC114694553), mRNA | 3.769 | 2.353 |
| XM_028860761.1 | TNF alpha induced protein 3 (Tnfaip3), transcript variant X2, mRNA | 3.767 | 1.511 |
| XM_028882480.1 | CMRF35-like molecule 7 (LOC114702149), mRNA | 3.730 | 2.084 |
| XM_028859043.1 | C-type lectin domain containing 5A (Clec5a), transcript variant X1, mRNA | 3.699 | 1.483 |
| XM_028860502.1 | acyloxyacyl hydrolase (Aoah), mRNA | 3.696 | 1.516 |
| XM_028893881.1 | BCL3 transcription coactivator (Bcl3), mRNA | 3.690 | 2.273 |
| XM_028887237.1 | triggering receptor expressed on myeloid cells 1-like (LOC114705361), mRNA | 3.670 | 2.840 |
| XM_028894000.1 | glutaredoxin (Glrx), transcript variant X1, mRNA | 3.621 | 2.747 |
| XM_028884235.1 | solute carrier family 15 member 3 (Slc15a3), mRNA | 3.607 | 2.162 |
| XM_028888173.1 | FGR proto-oncogene, Src family tyrosine kinase (Fgr), mRNA | 3.586 | 2.147 |
| XM_028873441.1 | phosphatidylinositol binding clathrin assembly protein (Picalm), transcript variant X19, mRNA | 3.557 | 3.830 |
| XM_028878655.1 | chromosome unknown C15orf48 homolog (CUNH15orf48), mRNA | 3.541 | 2.564 |

Table S18

| accession | product | log2 LPS fold change | log2 infection fold change |
|---|---|---|---|
| XM_028853721.1 | mitogen-activated protein kinase kinase kinase kinase 2 (Map4k2), mRN | 0.360 | -0.181 |
| XM_028878712.1 | huntingtin interacting protein K (Hypk), transcript variant X1, mRNA | 0.357 | -0.037 |
| XM_028879854.1 | cytochrome c oxidase subunit 7A2, mitochondrial (LOC114700271), mRN | 0.356 | 0.527 |
| XM_028892906.1 | calmodulin 2 (Calm2), mRNA | 0.353 | 0.399 |
| XM_028863348.1 | NADH:ubiquinone oxidoreductase subunit A11 (Ndufa11), mRNA | 0.350 | -0.067 |
| XM_028875462.1 | SYF2 pre-mRNA splicing factor (Syf2), mRNA | 0.349 | 0.133 |
| XM_028879761.1 | vinculin (Vcl), transcript variant X1, mRNA | 0.347 | -0.003 |
| XR_003734433.1 | uncharacterized LOC114692512 (LOC114692512), ncRNA | 0.347 | -0.143 |
| XM_028868796.1 | UBA domain containing 1 (Ubac1), mRNA | 0.346 | -0.186 |
| XM_028886928.1 | cytochrome b-c1 complex subunit 1, mitochondrial (LOC114705194), mF | 0.346 | 0.442 |
| XM_028878383.1 | oxoglutarate dehydrogenase like (Ogdhl), transcript variant X4, mRNA | 0.345 | 0.193 |
| XM_028892084.1 | peptidylprolyl cis/trans isomerase, NIMA-interacting 4 (Pin4), mRNA | 0.345 | -0.004 |
| XM_028889827.1 | uroporphyrinogen decarboxylase (Urod), transcript variant X2, mRNA | 0.344 | 0.000 |
| XM_028854184.1 | ribophorin I (Rpn1), mRNA | 0.344 | 0.702 |
| XM_028883031.1 | growth hormone inducible transmembrane protein (Ghitm), transcript v | 0.344 | 0.387 |
| XM_028870735.1 | tyrosine 3-monooxygenase/tryptophan 5-monooxygenase activation pro | 0.344 | -0.038 |
| XM_028868716.1 | tubulin beta 4B class IVb (Tubb4b), mRNA | 0.344 | 0.359 |
| XM_028877516.1 | signal peptidase complex subunit 2 (Spcs2), mRNA | 0.339 | 0.420 |
| XM_028870747.1 | lymphocyte specific protein 1 (Lsp1), transcript variant X2, mRNA | 0.339 | 0.423 |
| XM_028870105.1 | eukaryotic translation initiation factor 4H (Eif4h), mRNA | 0.338 | 0.334 |
| XM_028888763.1 | T-cell-specific guanine nucleotide triphosphate-binding protein 1-like (L | 0.336 | -0.251 |
| XR_003734705.1 | uncharacterized LOC114694442 (LOC114694442), ncRNA | 0.334 | 0.370 |
| XR_003735313.1 | small nucleolar RNA SNORA3/SNORA45 family (LOC114698186), ncRN | 0.333 | 0.399 |
| XM_028862160.1 | proteasome subunit beta 6 (Psmb6), transcript variant X1, mRNA | 0.331 | 1.024 |
| XR_003733830.1 | uncharacterized LOC114688979 (LOC114688979), ncRNA | 0.331 | -1.533 |
| XR_003734134.1 | uncharacterized LOC114690823 (LOC114690823), transcript variant X3, | 0.328 | -0.583 |
| XM_028869824.1 | paraoxonase 2 (Pon2), mRNA | 0.327 | -0.354 |
| XR_003735598.1 | small nucleolar RNA SNORA15 (LOC114699856), ncRNA | 0.325 | 0.834 |
| XM_028889608.1 | growth factor receptor bound protein 2 (Grb2), transcript variant X1, mF | 0.325 | 0.458 |
| XM_028889691.1 | aldo-keto reductase family 1 member A1 (Akr1a1), transcript variant X2 | 0.324 | 0.613 |
| XM_028857543.1 | adaptor related protein complex 2 subunit mu 1 (Ap2m1), transcript var | 0.324 | 0.683 |
| XM_028889819.1 | uroporphyrinogen decarboxylase (Urod), transcript variant X1, mRNA | 0.323 | 0.280 |
| XM_028862325.1 | ubiquitin conjugating enzyme E2 S (Ube2s), mRNA | 0.323 | 0.344 |
| XM_028855605.1 | myosin light chain 9 (Myl9), transcript variant X1, mRNA | 0.322 | -0.560 |
| XM_028862919.1 | ornithine decarboxylase antizyme 1 (Oaz1), mRNA | 0.319 | -0.255 |
| XM_028893583.1 | transmembrane protein 254 (Tmem254), transcript variant X1, mRNA | 0.319 | 0.089 |
| XM_028863632.1 | PRELI domain containing 1 (Prelid1), mRNA | 0.318 | 1.133 |
| XM_028872869.1 | calcium regulated heat stable protein 1 (Carhsp1), transcript variant X1, | 0.317 | -0.310 |
| XM_028872570.1 | septin 7 (Septin7), transcript variant X1, mRNA | 0.317 | -0.620 |
| XM_028870171.1 | coiled-coil-helix-coiled-coil-helix domain-containing protein 2 (LOC1146 | 0.316 | 0.970 |
| XM_028856958.1 | coatomer protein complex subunit epsilon (Cope), mRNA | 0.315 | 0.404 |
| XM_028858615.1 | R3H domain containing 4 (R3hdm4), mRNA | 0.315 | 0.014 |
| XM_028853755.1 | cytochrome c oxidase subunit 8A, mitochondrial (LOC114680703), mRN | 0.315 | 0.550 |
| XM_028859921.1 | membrane associated ring-CH-type finger 2 (March2), transcript variant | 0.313 | -0.264 |
| XM_028869563.1 | transmembrane protein 256 (Tmem256), transcript variant X1, mRNA | 0.308 | 0.113 |
| XM_028865204.1 | DEAD-box helicase 5 (Ddx5), transcript variant X2, mRNA | 0.306 | 0.051 |
| XM_028878856.1 | proteasome 26S subunit, ATPase 3 (Psmc3), mRNA | 0.306 | 0.479 |
| XM_028855029.1 | calnexin (Canx), transcript variant X1, mRNA | 0.303 | 0.433 |
| XM_028859906.1 | NADH:ubiquinone oxidoreductase subunit A7 (Ndufa7), mRNA | 0.299 | 0.085 |
| XM_028868438.1 | tyrosine 3-monooxygenase/tryptophan 5-monooxygenase activation pro | 0.298 | 0.216 |
| XM_028894523.1 | Janus kinase 2 (Jak2), mRNA | 0.297 | -0.129 |
| XM_028874405.1 | coatomer protein complex subunit zeta 1 (Copz1), transcript variant X2, | 0.295 | 0.573 |
| XM_028869104.1 | proteasome subunit beta 3 (Psmb3), mRNA | 0.295 | 0.841 |
| XM_028866582.1 | proteasome subunit beta 1 (Psmb1), mRNA | 0.292 | 0.478 |
| XR_003734734.1 | uncharacterized LOC114694674 (LOC114694674), ncRNA | 0.291 | 0.101 |
| XM_028856785.1 | actin related protein 2/3 complex subunit 5 like (Arpc5l), mRNA | 0.288 | 0.453 |
| XM_028870966.1 | VPS28 subunit of ESCRT-I (Vps28), mRNA | 0.287 | 0.501 |
| XM_028883701.1 | AKT serine/threonine kinase 1 (Akt1), mRNA | 0.287 | 0.420 |
| XM_028864186.1 | DnaJ heat shock protein family (Hsp40) member B6 (Dnajb6), transcript | 0.287 | 0.116 |
| XM_028865260.1 | protein kinase cAMP-dependent type I regulatory subunit alpha (Prkar1a | 0.285 | -0.181 |
| XM_028867591.1 | lysosomal protein transmembrane 4 alpha (Laptm4a), mRNA | 0.285 | 0.474 |
| XM_028894298.1 | ubiquitin C (Ubc), mRNA | 0.284 | 0.664 |
| XM_028860835.1 | iron-sulfur cluster assembly enzyme (Iscu), mRNA | 0.284 | 0.305 |
| XM_028891472.1 | TNF receptor superfamily member 14 (Tnfrsf14), transcript variant X2, | 0.283 | -0.390 |
| XM_028894408.1 | malate dehydrogenase 2 (Mdh2), mRNA | 0.283 | 0.330 |
| XM_028855698.1 | ubiquitin conjugating enzyme E2 N (Ube2n), mRNA | 0.281 | 0.569 |
| XM_028876074.1 | activator of HSP90 ATPase activity 1 (Ahsa1), mRNA | 0.279 | -0.054 |
| XM_028866212.1 | ATPase H+ transporting V1 subunit F (Atp6v1f), mRNA | 0.278 | 0.699 |
| XM_028878542.1 | DnaJ heat shock protein family (Hsp40) member C15 (Dnajc15), transcri | 0.278 | -0.084 |
| XM_028894422.1 | fission, mitochondrial 1 (Fis1), transcript variant X1, mRNA | 0.277 | -0.080 |
| XM_028860632.1 | protein phosphatase 1 catalytic subunit alpha (Ppp1ca), mRNA | 0.277 | 0.499 |
| XM_028861300.1 | eukaryotic translation initiation factor 3 subunit C (LOC114686662), tran | 0.276 | 0.513 |
| XR_003733775.1 | uncharacterized LOC114688353 (LOC114688353), ncRNA | 0.275 | 1.554 |
| XM_028870396.1 | mitochondrial calcium uptake 2 (Micu2), mRNA | 0.274 | 0.120 |
| XM_028874549.1 | DAZ associated protein 2 (Dazap2), transcript variant X1, mRNA | 0.273 | -0.610 |
| XM_028855700.1 | nudix hydrolase 4 (Nudt4), transcript variant X2, mRNA | 0.273 | -0.217 |
| XR_003735919.1 | uncharacterized LOC114702026 (LOC114702026), transcript variant X2, | 0.272 | -0.405 |
| XM_028879211.1 | interferon stimulated exonuclease gene 20 (Isg20), mRNA | 0.271 | 0.405 |
| XM_028861016.1 | proteasome subunit beta 8 (Psmb8), mRNA | 0.271 | 0.695 |
| XM_028866182.1 | ubiquitin conjugating enzyme E2 H (Ube2h), transcript variant X1, mRN | 0.270 | 0.301 |
| XM_028854604.1 | hydroxyacylglutathione hydrolase (Hagh), transcript variant X3, mRNA | 0.270 | 0.252 |
| XM_028891673.1 | cytochrome c oxidase subunit 7C, mitochondrial (LOC114708408), mRN | 0.268 | 0.680 |
| XR_003736363.1 | uncharacterized LOC114705193 (LOC114705193), ncRNA | 0.266 | -0.464 |
| XM_028875034.1 | Bruton tyrosine kinase (Btk), transcript variant X1, mRNA | 0.262 | -0.172 |
| XM_028854296.1 | ankyrin repeat and BTB domain containing 1 (Abtb1), transcript variant | 0.262 | -0.237 |
| XM_028870418.1 | crystallin lambda 1 (Cryl1), mRNA | 0.262 | -0.436 |
| XM_028880438.1 | ATPase H+ transporting V1 subunit E1 (Atp6v1e1), mRNA | 0.262 | 0.226 |
| XM_028855284.1 | presenilin enhancer, gamma-secretase subunit (Psenen), mRNA | 0.261 | 0.463 |
| XM_028884622.1 | regulator of G protein signaling 18 (Rgs18), mRNA | 0.261 | -0.028 |
| XM_028859920.1 | membrane associated ring-CH-type finger 2 (March2), transcript variant | 0.260 | -0.301 |
| XM_028858035.1 | aminolevulinate dehydratase (Alad), transcript variant X4, mRNA | 0.258 | -0.041 |
| XM_028878613.1 | Ras association domain family member 2 (Rassf2), transcript variant X1 | 0.257 | -0.288 |
| XM_028890289.1 | actin related protein 2/3 complex subunit 1A (Arpc1a), mRNA | 0.255 | 0.407 |
| XM_028869459.1 | eukaryotic translation initiation factor 4A1 (Eif4a1), mRNA | 0.254 | 0.696 |
| XM_028875318.1 | dolichyl-diphosphooligosaccharide--protein glycosyltransferase non-cata | 0.254 | 0.637 |
| XM_028853956.1 | elongin B (Elob), mRNA | 0.254 | 0.499 |
| XM_028859341.1 | homeodomain interacting protein kinase 1 (Hipk1), transcript variant X1 | 0.250 | 0.333 |
| XM_028856794.1 | proteasome subunit beta 7 (Psmb7), mRNA | 0.248 | 0.461 |
| XM_028889925.1 | ATP synthase peripheral stalk-membrane subunit b (Atp5pb), mRNA | 0.246 | 0.583 |
| XM_028871056.1 | myosin heavy chain 9 (Myh9), transcript variant X2, mRNA | 0.243 | -0.067 |
| XM_028868445.1 | serine/threonine kinase 4 (Stk4), mRNA | 0.243 | -0.326 |
| XM_028883184.1 | solute carrier family 25 member 37 (Slc25a37), transcript variant X1, m | 0.241 | 0.156 |
| XM_028855103.1 | biliverdin reductase B (Blvrb), mRNA | 0.240 | 0.021 |
| XM_028895314.1 | 5'-aminolevulinate synthase 1 (Alas1), mRNA | 0.238 | 0.714 |
| XM_028885909.1 | protein phosphatase 1 catalytic subunit gamma (Ppp1cc), mRNA | 0.237 | -0.178 |
| XM_028891377.1 | mitochondrial ribosomal protein L20 (Mrpl20), transcript variant X1, mR | 0.234 | 1.263 |
| XM_028855175.1 | methionine adenosyltransferase 2A (Mat2a), mRNA | 0.232 | -0.095 |
| XM_028884050.1 | BCL2 associated athanogene 1 (Bag1), transcript variant X1, mRNA | 0.231 | -0.048 |
| XM_028862604.1 | paired immunoglobulin-like type 2 receptor alpha (LOC114687970), mR | 0.230 | 1.838 |
| XM_028869008.1 | phosphoethanolamine/phosphocholine phosphatase (Phospho1), transc | 0.229 | -0.484 |
| XM_028877317.1 | cold shock domain containing E1 (Csde1), transcript variant X4, mRNA | 0.228 | 0.246 |
| XM_028887677.1 | protein tyrosine phosphatase 4A2 (Ptp4a2), transcript variant X3, mRNA | 0.227 | -0.262 |
| XM_028893048.1 | cyclin Pas1/PHO80 domain containing 1 (Cnppd1), transcript variant X2, | 0.225 | 0.568 |
| XM_028863942.1 | BRICK1 subunit of SCAR/WAVE actin nucleating complex (Brk1), mRNA | 0.225 | 0.591 |
| XM_028870106.1 | linker for activation of T cells family member 2 (Lat2), mRNA | 0.224 | 0.447 |
| XM_028871365.1 | AUP1 lipid droplet regulating VLDL assembly factor (Aup1), transcript va | 0.220 | 0.379 |
| XM_028884157.1 | kinesin family member 2A (Kif2a), mRNA | 0.220 | -1.449 |
| XM_028863258.1 | leukocyte-associated immunoglobulin-like receptor 1 (LOC114688727), | 0.218 | 2.094 |
| XM_028865666.1 | cytochrome c oxidase subunit 5B, mitochondrial (LOC114690777), mRN | 0.218 | 0.551 |
| XM_028891145.1 | NADH:ubiquinone oxidoreductase subunit B8 (Ndufb8), mRNA | 0.217 | 0.419 |
| XM_028859271.1 | epithelial membrane protein 3 (Emp3), mRNA | 0.215 | 0.979 |
| XM_028871062.1 | eukaryotic translation initiation factor 3 subunit D (Eif3d), mRNA | 0.214 | 0.097 |
| XM_028886390.1 | cytochrome c oxidase subunit 7B, mitochondrial (LOC114704883), mRN | 0.213 | 0.575 |
| XR_003736093.1 | uncharacterized LOC114703137 (LOC114703137), ncRNA | 0.211 | -0.742 |
| XM_028866320.1 | death associated protein kinase 2 (Dapk2), mRNA | 0.206 | 0.215 |
| XM_028865440.1 | histone H3.3 (LOC114690628), mRNA | 0.206 | 0.229 |
| XM_028857970.1 | syntaxin 5 (Stx5), mRNA | 0.206 | 0.431 |
| XM_028889996.1 | BCL2/adenovirus E1B 19 kDa protein-interacting protein 3-like (LOC1147 | 0.202 | 0.125 |
| XM_028873251.1 | SH3 domain containing kinase binding protein 1 (Sh3kbp1), transcript va | 0.202 | 1.302 |
| XM_028884803.1 | NADH:ubiquinone oxidoreductase subunit B11 (Ndufb11), mRNA | 0.201 | 0.117 |
| XM_028873577.1 | mitogen-activated protein kinase 1 interacting protein 1 like (Mapk1ip1l | 0.201 | -0.004 |
| XM_028881183.1 | centromere protein U (Cenpu), mRNA | 0.196 | -0.801 |
| XM_028861044.1 | H-2 class I histocompatibility antigen, Q10 alpha chain-like (LOC114686 | 0.193 | 0.511 |
| XM_028866028.1 | ATPase Na+/K+ transporting subunit beta 3 (Atp1b3), transcript variant | 0.192 | 0.339 |
| XM_028857069.1 | ubiquitin B (Ubb), mRNA | 0.192 | -0.031 |
| XM_028889663.1 | ATP synthase peripheral stalk subunit d (Atp5pd), transcript variant X1, | 0.189 | 0.387 |
| XM_028877529.1 | uncoupling protein 2 (Ucp2), mRNA | 0.189 | 0.307 |

Table S18

| | | | | | | | |
|---|---|---|---|---|---|---|---|
| XM_028859044.1 | C-type lectin domain containing 5A (Clec5a), transcript variant X2, mRNA | 3.527 | 1.889 | XM_028864100.1 | aquaporin 1 (Colton blood group) (Aqp1), mRNA | 0.188 | -0.006 |
| XM_028892728.1 | ATPase phospholipid transporting 8B4 (putative) (Atp8b4), mRNA | 3.499 | 1.867 | XM_028872420.1 | tumor necrosis factor receptor superfamily member 26-like (LOC11469[illegible] | 0.185 | -0.325 |
| XM_028869610.1 | interleukin 1 receptor associated kinase 3 (Irak3), mRNA | 3.498 | 2.656 | XM_028888492.1 | B cell receptor associated protein 31 (Bcap31), transcript variant X1, m[illegible] | 0.185 | 0.438 |
| XM_028885448.1 | colony stimulating factor 3 receptor (Csf3r), mRNA | 3.492 | 2.150 | XM_028880802.1 | cytochrome c oxidase subunit 4 isoform 1, mitochondrial (LOC11470094 | 0.182 | 1.229 |
| XM_028888153.1 | platelet activating factor receptor (Ptafr), mRNA | 3.470 | 2.838 | XM_028878347.1 | annexin A8 (LOC114699341), mRNA | 0.181 | -0.145 |
| XM_028864452.1 | peroxiredoxin 6 (Prdx6), mRNA | 3.468 | 1.597 | XM_028876876.1 | G protein subunit gamma 5 (Gng5), mRNA | 0.180 | 0.521 |
| XM_028854938.1 | complement C5a receptor 1 (C5ar1), mRNA | 3.460 | 1.646 | XM_028892569.1 | synaptogyrin 2 (Syngr2), transcript variant X1, mRNA | 0.179 | 0.753 |
| XM_028894546.1 | CD274 molecule (Cd274), mRNA | 3.435 | 2.375 | XM_028889501.1 | migration and invasion enhancer 1 (Mien1), mRNA | 0.177 | 0.262 |
| XM_028873188.1 | C-X-C motif chemokine ligand 13 (Cxcl13), mRNA | 3.429 | 0.823 | XR_003733876.1 | uncharacterized LOC114689225 (LOC114689225), ncRNA | 0.174 | 0.642 |
| XM_028856765.1 | translocator protein (Tspo), transcript variant X1, mRNA | 3.414 | 2.924 | XM_028889669.1 | WW domain binding protein 2 (Wbp2), transcript variant X4, mRNA | 0.174 | -0.515 |
| XM_028862915.1 | growth arrest and DNA damage inducible beta (Gadd45b), mRNA | 3.412 | 2.069 | XM_028864156.1 | heterogeneous nuclear ribonucleoprotein A2/B1 (Hnrnpa2b1), transcript | 0.173 | -0.525 |
| XM_028855869.1 | EH domain containing 1 (Ehd1), mRNA | 3.395 | 2.208 | XM_028881499.1 | GABA type A receptor associated protein like 1 (Gabarapl1), mRNA | 0.173 | 0.461 |
| XM_028889626.1 | CD300 molecule like family member f (Cd300lf), transcript variant X1, mRNA | 3.389 | 2.503 | XR_003734554.1 | uncharacterized LOC114693456 (LOC114693456), ncRNA | 0.170 | 1.034 |
| XM_028853710.1 | peroxiredoxin 5 (Prdx5), mRNA | 3.383 | 1.573 | XM_028854602.1 | hydroxyacylglutathione hydrolase (Hagh), transcript variant X1, mRNA | 0.167 | 0.010 |
| XM_028862946.1 | paired immunoglobulin-like type 2 receptor beta-2 (LOC114688352), mRNA | 3.351 | 1.532 | XM_028890347.1 | eukaryotic translation initiation factor 2 alpha kinase 1 (Eif2ak1), transc | 0.165 | -0.145 |
| XM_028861203.1 | complement C3 (C3), mRNA | 3.331 | 2.716 | XM_028886787.1 | zinc finger matrin-type 2 (Zmat2), mRNA | 0.164 | 0.253 |
| XM_028881407.1 | cytochrome b-245 heavy chain (LOC114701319), mRNA | 3.313 | 2.329 | XM_028869046.1 | CDK5 regulatory subunit associated protein 3 (Cdk5rap3), mRNA | 0.164 | 0.029 |
| XM_028869724.1 | NFKB inhibitor alpha (Nfkbia), mRNA | 3.300 | 1.906 | XM_028866516.1 | ornithine decarboxylase antizyme 2 (Oaz2), mRNA | 0.160 | -0.180 |
| XM_028894751.1 | GLI pathogenesis related 2 (Glipr2), mRNA | 3.298 | 1.866 | XM_028867415.1 | ATP synthase membrane subunit e (Atp5me), mRNA | 0.156 | 1.056 |
| XM_028861083.1 | NFAT activating protein with ITAM motif 1 (Nfam1), mRNA | 3.296 | 1.898 | XM_028887014.1 | RNA binding motif protein 5 (Rbm5), transcript variant X1, mRNA | 0.155 | -0.124 |
| XM_028877852.1 | basic leucine zipper ATF-like transcription factor (Batf), mRNA | 3.266 | 1.283 | XM_028881758.1 | glycophorin A (MNS blood group) (Gypa), mRNA | 0.154 | 0.197 |
| XM_028873674.1 | xanthine dehydrogenase (Xdh), mRNA | 3.262 | 2.052 | XM_028853916.1 | poly(rC) binding protein 1 (Pcbp1), mRNA | 0.153 | 0.216 |
| XM_028891125.1 | neuroepithelial cell transforming 1 (Net1), transcript variant X2, mRNA | 3.255 | 1.877 | XM_028855390.1 | guanylate-binding protein 4-like (LOC114681719), mRNA | 0.152 | -1.050 |
| XM_028862137.1 | T-cell-interacting, activating receptor on myeloid cells protein 1-like (LOC114687526), mRNA | 3.252 | 1.739 | XM_028873845.1 | zinc finger AN1-type containing 6 (Zfand6), transcript variant X4, mRNA | 0.151 | 0.923 |
| XM_028895342.1 | cytokine inducible SH2 containing protein (Cish), transcript variant X1, mR | 3.252 | 1.525 | XM_028893719.1 | MCTS1 re-initiation and release factor (Mcts1), transcript variant X2, m[illegible] | 0.149 | 0.629 |
| XM_028876606.1 | chitinase 1 (Chit1), mRNA | 3.218 | 0.306 | XM_028878301.1 | tumor protein, translationally-controlled 1 (Tpt1), mRNA | 0.147 | 0.216 |
| XM_028873528.1 | purine nucleoside phosphorylase (Pnp), mRNA | 3.187 | 3.188 | XM_028859916.1 | heterogeneous nuclear ribonucleoprotein M (Hnrnpm), transcript varian | 0.144 | -0.064 |
| XM_028877802.1 | GRAM domain containing 1A (Gramd1a), transcript variant X3, mRNA | 3.186 | 0.518 | XM_028859968.1 | proteasome subunit alpha 2 (Psma2), mRNA | 0.144 | 0.441 |
| XM_028888769.1 | hepatitis A virus cellular receptor 1 homolog (LOC114706385), mRNA | 3.172 | 4.766 | XM_028871297.1 | apolipoprotein L3-like (LOC114694378), mRNA | 0.138 | 0.350 |
| XM_028856420.1 | UDP-glucose ceramide glucosyltransferase (Ugcg), mRNA | 3.160 | 1.243 | XM_028858877.1 | mediator complex subunit 25 (Med25), transcript variant X5, mRNA | 0.137 | -1.178 |
| XM_028872166.1 | phospholipase B domain containing 1 (Plbd1), mRNA | 3.138 | 1.219 | XM_028853713.1 | HRAS-like suppressor 3 (LOC114680680), transcript variant X2, mRNA | 0.137 | -0.955 |
| XM_028884543.1 | sorting nexin 18 (Snx18), mRNA | 3.131 | 1.225 | XM_028863978.1 | synapsin II (Syn2), mRNA | 0.136 | 0.008 |
| XM_028863511.1 | hepatitis A virus cellular receptor 1 homolog (LOC114689091), mRNA | 3.126 | 4.903 | XM_028889269.1 | synuclein alpha (Snca), transcript variant X2, mRNA | 0.136 | 0.542 |
| XM_028861390.1 | putative killer cell immunoglobulin-like receptor like protein KIR3DP1 (LOC114686723), mRNA | 3.122 | 2.679 | XM_028877081.1 | ATP synthase peripheral stalk subunit OSCP (Atp5po), mRNA | 0.136 | 0.492 |
| XM_028888871.1 | C-type lectin domain family 4 member A-like (LOC114706473), mRNA | 3.077 | 1.939 | XM_028893365.1 | small nuclear ribonucleoprotein U11/U12 subunit 25 (Snrnp25), mRNA | 0.136 | 0.000 |
| XM_028887969.1 | leukocyte specific transcript 1 (Lst1), transcript variant X1, mRNA | 3.076 | 2.466 | XM_028856476.1 | FLII actin remodeling protein (Flii), mRNA | 0.136 | -0.042 |
| XM_028886974.1 | 6-phosphofructo-2-kinase/fructose-2,6-biphosphatase 4 (Pfkfb4), mRNA | 3.072 | 1.721 | XM_028855416.1 | citrate synthase (Cs), mRNA | 0.133 | 0.283 |
| XM_028891815.1 | NEDD4 binding protein 1 (N4bp1), mRNA | 3.060 | 1.576 | XM_028853669.1 | Rho GTPase activating protein 25 (Arhgap25), mRNA | 0.130 | 0.204 |
| XM_028893099.1 | solute carrier family 11 member 1 (Slc11a1), mRNA | 3.059 | 2.301 | XM_028855785.1 | ATP synthase F1 subunit epsilon (Atp5f1e), mRNA | 0.130 | -0.102 |
| XM_028858543.1 | strawberry notch homolog 2 (Sbno2), transcript variant X1, mRNA | 3.050 | 1.669 | XM_028885800.1 | cyclin dependent kinase 2 associated protein 1 (Cdk2ap1), mRNA | 0.126 | -0.265 |
| XM_028862630.1 | C-type lectin domain family 4 member E (Clec4e), mRNA | 3.012 | 1.577 | XM_028859394.1 | glutaredoxin 5 (Glrx5), mRNA | 0.125 | 0.084 |
| XM_028876009.1 | Src like adaptor (Sla), transcript variant X3, mRNA | 2.956 | -0.137 | XM_028887798.1 | voltage dependent anion channel 3 (Vdac3), transcript variant X1, mRN[illegible] | 0.125 | -0.314 |
| XM_028856428.1 | G protein subunit gamma 10 (Gng10), transcript variant X2, mRNA | 2.955 | 1.610 | XM_028868469.1 | cathepsin A (Ctsa), mRNA | 0.124 | 0.507 |
| XM_028861947.1 | S100 calcium binding protein A11 (S100a11), mRNA | 2.947 | 1.829 | XM_028858034.1 | aminolevulinate dehydratase (Alad), transcript variant X3, mRNA | 0.123 | -0.241 |
| XM_028864148.1 | src kinase associated phosphoprotein 2 (Skap2), mRNA | 2.932 | 1.939 | XM_028860251.1 | tumor protein p63 regulated 1 like (Tprg1l), mRNA | 0.121 | 0.037 |
| XM_028889392.1 | phosphoinositide-3-kinase adaptor protein 1 (Pik3ap1), mRNA | 2.931 | 1.550 | XM_028875981.1 | ATP synthase peripheral stalk subunit F6 (Atp5pf), transcript variant X1, | 0.121 | 0.254 |
| XM_028862233.1 | JunB proto-oncogene, AP-1 transcription factor subunit (Junb), mRNA | 2.926 | 0.336 | XM_028887629.1 | erythrocyte membrane protein band 4.1 (Epb41), transcript variant X16, | 0.119 | 0.666 |
| XM_028894284.1 | toll like receptor 2 (Tlr2), mRNA | 2.907 | 1.861 | XM_028869494.1 | eukaryotic translation initiation factor 5A (Eif5a), transcript variant X1, [illegible] | 0.118 | 0.376 |
| XM_028873434.1 | phosphatidylinositol binding clathrin assembly protein (Picalm), transcript variant X12, mRNA | 2.906 | 1.649 | XM_028869502.1 | CTD nuclear envelope phosphatase 1 (Ctdnep1), mRNA | 0.118 | -0.044 |
| XM_028887461.1 | HCK proto-oncogene, Src family tyrosine kinase (Hck), mRNA | 2.903 | 2.042 | XM_028887990.1 | proline rich coiled-coil 2A (Prrc2a), mRNA | 0.117 | -0.133 |
| XM_028861698.1 | flotillin 1 (Flot1), mRNA | 2.898 | 1.460 | XM_028857953.1 | UBX domain protein 1 (Ubxn1), mRNA | 0.116 | 0.032 |
| XR_003735902.1 | uncharacterized LOC114701932 (LOC114701932), ncRNA | 2.882 | 1.834 | XM_028891572.1 | NEDD4 family-interacting protein 1 (LOC114708344), transcript variant [illegible] | 0.114 | 0.071 |
| XM_028873986.1 | RAB31, member RAS oncogene family (Rab31), mRNA | 2.881 | 1.999 | XM_028865203.1 | DEAD-box helicase 5 (Ddx5), transcript variant X1, mRNA | 0.110 | -0.270 |
| XM_028879921.1 | interferon-induced GTP-binding protein Mx2 (LOC114700347), mRNA | 2.879 | -0.435 | XM_028854912.1 | NSF attachment protein alpha (Napa), mRNA | 0.110 | 0.315 |
| XM_028856766.1 | translocator protein (Tspo), transcript variant X2, mRNA | 2.879 | 8.586 | XM_028892089.1 | interleukin 2 receptor subunit gamma (Il2rg), mRNA | 0.110 | -0.278 |
| XM_028878860.1 | Spi-1 proto-oncogene (Spi1), transcript variant X1, mRNA | 2.870 | 1.813 | XM_028855516.1 | ADP ribosylation factor like GTPase 6 interacting protein 5 (Arl6ip5), m[illegible] | 0.109 | 0.526 |
| XM_028888360.1 | FKBP prolyl isomerase 5 (Fkbp5), transcript variant X1, mRNA | 2.869 | -0.243 | XM_028855424.1 | RAB5B, member RAS oncogene family (Rab5b), mRNA | 0.107 | 0.041 |
| XM_028861318.1 | apolipoprotein B receptor (Apobr), mRNA | 2.859 | 1.868 | XM_028880687.1 | chromosome unknown C6orf62 homolog (CUNH6orf62), transcript varia | 0.106 | -0.201 |
| XM_028880029.1 | TNFAIP3 interacting protein 1 (Tnip1), transcript variant X7, mRNA | 2.858 | 2.527 | XM_028882698.1 | integrin subunit beta 3 (Itgb3), mRNA | 0.104 | -0.802 |
| XM_028855239.1 | TYRO protein tyrosine kinase binding protein (Tyrobp), mRNA | 2.848 | 2.230 | XM_028888354.1 | serine and arginine rich splicing factor 3 (Srsf3), transcript variant X1, | 0.104 | 1.807 |
| XM_028858147.1 | granzyme B-like (LOC114683802), mRNA | 2.843 | 1.159 | XM_028892123.1 | NEDD8 ubiquitin like modifier (Nedd8), mRNA | 0.103 | 0.877 |
| XM_028859188.1 | ZFP36 ring finger protein (Zfp36), mRNA | 2.827 | -0.085 | XM_028870590.1 | drebrin like (Dbnl), transcript variant X2, mRNA | 0.100 | 0.687 |
| XM_028890404.1 | tumor protein D52 (Tpd52), transcript variant X3, mRNA | 2.826 | 1.748 | XM_028868577.1 | ubiquitin conjugating enzyme E2 V1 (Ube2v1), mRNA | 0.100 | 0.104 |
| XM_028873150.1 | ANTXR cell adhesion molecule 2 (Antxr2), mRNA | 2.823 | 1.395 | XM_028894904.1 | dedicator of cytokinesis 2 (Dock2), mRNA | 0.099 | 0.019 |
| XM_028878737.1 | solute carrier family 28 member 2 (Slc28a2), mRNA | 2.822 | 2.699 | XM_028869007.1 | phosphoethanolamine/phosphocholine phosphatase (Phospho1), transc[illegible] | 0.098 | -0.268 |
| XM_028873049.1 | oxysterol binding protein like 8 (Osbpl8), transcript variant X3, mRNA | 2.819 | 0.897 | XM_028887676.1 | protein tyrosine phosphatase 4A2 (Ptp4a2), transcript variant X2, mRNA | 0.096 | -0.462 |
| XM_028863547.1 | leukocyte immunoglobulin-like receptor subfamily B member 3 (LOC114689150), partial mRNA | 2.814 | 1.746 | XR_003735608.1 | uncharacterized LOC114699906 (LOC114699906), transcript variant X1, [illegible] | 0.095 | -0.431 |
| XM_028856355.1 | granulocyte-macrophage colony-stimulating factor receptor subunit alpha-like (LOC114682475), mRNA | 2.786 | 2.111 | XM_028870405.1 | Sin3A associated protein 18 (Sap18), mRNA | 0.094 | 0.118 |
| XM_028878524.1 | syndecan binding protein (Sdcbp), transcript variant X2, mRNA | 2.782 | 1.040 | XM_028886352.1 | splicing factor 3b subunit 1 (Sf3b1), transcript variant X1, mRNA | 0.094 | -0.112 |
| XM_028883622.1 | cytoskeleton associated protein 4 (Ckap4), mRNA | 2.778 | 1.287 | XM_028860381.1 | small ubiquitin like modifier 1 (Sumo1), transcript variant X2, mRNA | 0.093 | -0.172 |
| XM_028881371.1 | Pim-2 proto-oncogene, serine/threonine kinase (Pim2), mRNA | 2.763 | 0.336 | XM_028886865.1 | coiled-coil domain containing 12 (Ccdc12), mRNA | 0.093 | 0.177 |
| XM_028878742.1 | sulfide quinone oxidoreductase (Sqor), mRNA | 2.760 | 2.340 | XM_028886420.1 | small ArfGAP2 (Smap2), transcript variant X1, mRNA | 0.091 | -0.370 |
| XM_028880607.1 | nuclear factor, erythroid 2 like 2 (Nfe2l2), transcript variant X1, mRNA | 2.748 | 1.649 | XM_028881392.1 | RNA binding motif protein 3 (Rbm3), transcript variant X2, mRNA | 0.090 | 0.846 |
| XM_028853900.1 | MAX dimerization protein 1 (Mxd1), mRNA | 2.745 | 1.522 | XM_028880623.1 | ATP synthase membrane subunit c locus 3 (Atp5mc3), transcript variant | 0.090 | 0.683 |
| XM_028858621.1 | salvador family WW domain containing protein 1 (Sav1), mRNA | 2.735 | 0.350 | XM_028885852.1 | proteasome 26S subunit, non-ATPase 9 (Psmd9), mRNA | 0.088 | -0.302 |
| XM_028893599.1 | phosphogluconate dehydrogenase (Pgd), mRNA | 2.733 | 1.685 | XM_028853662.1 | centromere protein X (Cenpx), mRNA | 0.088 | 0.496 |
| XM_028876006.1 | Src like adaptor (Sla), transcript variant X1, mRNA | 2.728 | 0.788 | XM_028862937.1 | TSC22 domain family member 4 (Tsc22d4), transcript variant X1, mRNA | 0.087 | -0.343 |
| XM_028881290.1 | ATPase phospholipid transporting 11A (Atp11a), transcript variant X2, mRNA | 2.722 | -0.099 | XM_028880817.1 | microtubule associated protein 1 light chain 3 beta (Map1lc3b), mRNA | 0.086 | 0.554 |
| XM_028878765.1 | 2'-5'-oligoadenylate synthase 1A-like (LOC114699623), mRNA | 2.717 | 1.147 | XR_003737046.1 | uncharacterized LOC114710366 (LOC114710366), transcript variant X1, [illegible] | 0.085 | -0.050 |
| XM_028856427.1 | G protein subunit gamma 10 (Gng10), transcript variant X1, mRNA | 2.703 | 1.485 | XM_028855096.1 | proteasome 26S subunit, ATPase 4 (Psmc4), mRNA | 0.085 | 0.538 |
| XM_028877565.1 | phosphodiesterase 2A (Pde2a), mRNA | 2.692 | 1.051 | XR_003733780.1 | U2 spliceosomal RNA (LOC114688412), ncRNA | 0.083 | -0.625 |
| XM_028891607.1 | serpin family B member 2 (Serpinb2), mRNA | 2.680 | 0.651 | XM_028884040.1 | ubiquitin conjugating enzyme E2 R2 (Ube2r2), mRNA | 0.083 | 0.106 |
| XM_028883747.1 | creatine kinase B (Ckb), mRNA | 2.674 | 2.653 | XM_028877691.1 | ornithine decarboxylase 1 (Odc1), mRNA | 0.083 | -0.409 |
| XM_028882123.1 | leukocyte immunoglobulin-like receptor subfamily B member 3 (LOC114701921), | 2.673 | 2.025 | XM_028856558.1 | ribosomal protein lateral stalk subunit P0 (Rplp0), mRNA | 0.082 | 0.035 |
| XM_028860089.1 | Fc fragment of IgE receptor Ig (Fcer1g), mRNA | 2.668 | 2.224 | XM_028871057.1 | myosin heavy chain 9 (Myh9), transcript variant X3, mRNA | 0.082 | -1.277 |
| XM_028868425.1 | adenosine deaminase (Ada), transcript variant X2, mRNA | 2.660 | 0.342 | XM_028861334.1 | NADH:ubiquinone oxidoreductase subunit A13 (Ndufa13), mRNA | 0.080 | 0.582 |
| XM_028866853.1 | flotillin 2 (Flot2), transcript variant X1, mRNA | 2.654 | 0.200 | XM_028862230.1 | peroxiredoxin 2 (Prdx2), mRNA | 0.080 | -0.099 |
| XM_028864332.1 | activating transcription factor 6 (Atf6), transcript variant X1, mRNA | 2.651 | 1.186 | XM_028854747.1 | ring finger protein 10 (Rnf10), transcript variant X1, mRNA | 0.079 | 0.148 |
| XM_028885034.1 | Pim-1 proto-oncogene, serine/threonine kinase (Pim1), mRNA | 2.644 | 1.191 | XM_028888502.1 | ribosomal protein L10 (Rpl10), mRNA | 0.076 | 0.075 |
| XM_028872897.1 | lipopolysaccharide induced TNF factor (Litaf), mRNA | 2.637 | 1.656 | XM_028870737.1 | lymphocyte specific protein 1 (Lsp1), transcript variant X1, mRNA | 0.073 | 0.382 |
| XM_028885162.1 | signal-regulatory protein beta-1-like (LOC114704110), mRNA | 2.634 | 2.592 | XM_028878390.1 | nuclear receptor coactivator 4 (Ncoa4), transcript variant X1, mRNA | 0.071 | -0.593 |
| XM_028891388.1 | NAD kinase (Nadk), transcript variant X2, mRNA | 2.626 | 1.293 | XM_028867842.1 | cerebellar degeneration related protein 2 (Cdr2), transcript variant X1, [illegible] | 0.071 | -0.137 |
| XM_028855596.1 | SAM and HD domain containing deoxynucleoside triphosphate triphosphohydrolase 1 (Samhd1), mRNA | 2.608 | 1.408 | XM_028857150.1 | butyrophilin-like protein 10 (LOC114682995), mRNA | 0.071 | 0.001 |
| XM_028877410.1 | syntaxin 7 (Stx7), transcript variant X1, mRNA | 2.607 | 2.104 | XM_028861785.1 | RAD23 homolog A, nucleotide excision repair protein (Rad23a), transcri[illegible] | 0.069 | 0.326 |
| XM_028892550.1 | phosphatidylglycerophosphate synthase 1 (Pgs1), transcript variant X1, mRNA | 2.598 | 1.568 | XM_028878209.1 | serine carboxypeptidase 1 (Scpep1), mRNA | 0.067 | 0.630 |
| XM_028863639.1 | leukocyte immunoglobulin-like receptor subfamily B member 3 (LOC114689280), mRNA | 2.596 | 1.751 | XM_028865174.1 | LIM domain containing 2 (Limd2), transcript variant X2, mRNA | 0.066 | 0.066 |
| XM_028860519.1 | ST3 beta-galactoside alpha-2,3-sialyltransferase 5 (St3gal5), transcript variant X1, mRNA | 2.590 | 0.128 | XM_028891636.1 | transmembrane 7 superfamily member 3 (Tm7sf3), mRNA | 0.066 | -0.254 |
| XM_028878523.1 | syndecan binding protein (Sdcbp), transcript variant X1, mRNA | 2.586 | 0.625 | XM_028883718.1 | ATP synthase membrane subunit 6.8PL (Atp5mpl), mRNA | 0.065 | 0.702 |
| XM_028860726.1 | interferon gamma receptor 1 (Ifngr1), mRNA | 2.566 | 1.167 | XM_028892808.1 | CXXC repeat containing interactor of PDZ3 domain (Cript), mRNA | 0.065 | 0.402 |
| XM_028892331.1 | TNF receptor superfamily member 1A (Tnfrsf1a), mRNA | 2.560 | 1.673 | XM_028860633.1 | G protein-coupled receptor kinase 2 (Grk2), mRNA | 0.063 | 0.222 |
| XM_028888888.1 | C-type lectin domain family 4 member A-like (LOC114706495), mRNA | 2.547 | 2.235 | XM_028892800.1 | BRMS1 transcriptional repressor and anoikis regulator (Brms1), transcri | 0.063 | 0.114 |
| XM_028887447.1 | protein O-fucosyltransferase 1 (Pofut1), mRNA | 2.540 | 1.663 | XM_028878578.1 | DDRGK domain containing 1 (Ddrgk1), mRNA | 0.062 | 0.166 |
| XM_028873432.1 | phosphatidylinositol binding clathrin assembly protein (Picalm), transcript variant X11, mRNA | 2.536 | 0.685 | XM_028865550.1 | interferon gamma inducible protein 16 (Ifi16), transcript variant X3, mR | 0.062 | -0.586 |
| XM_028877107.1 | interferon gamma receptor 2 (Ifngr2), transcript variant X2, mRNA | 2.529 | 2.145 | XM_028861596.1 | prostaglandin E synthase 3 (Ptges3), mRNA | 0.061 | 0.268 |
| XM_028885647.1 | integrin subunit alpha M (Itgam), mRNA | 2.504 | 1.722 | XM_028870515.1 | vimentin (Vim), mRNA | 0.057 | 1.057 |
| XM_028874775.1 | SAM domain, SH3 domain and nuclear localization signals 1 (Samsn1), transcript variant X1, mRNA | 2.504 | 2.035 | XM_028861194.1 | cytochrome b-c1 complex subunit 10 (LOC114686525), mRNA | 0.057 | 0.589 |
| XM_028893723.1 | lysosomal associated membrane protein 2 (Lamp2), mRNA | 2.486 | 2.015 | XM_028870662.1 | splicing factor 3a subunit 1 (Sf3a1), mRNA | 0.056 | -0.389 |
| XM_028861099.1 | syntaxin binding protein 5 (Stxbp5), transcript variant X3, mRNA | 2.483 | 0.713 | XM_028872441.1 | cell division cycle 37 (Cdc37), mRNA | 0.055 | 0.166 |
| XM_028871500.1 | lysophosphatidylcholine acyltransferase 2 (Lpcat2), mRNA | 2.463 | 1.957 | XM_028857546.1 | proteasome 26S subunit, non-ATPase 2 (Psmd2), mRNA | 0.053 | 0.176 |
| XM_028891610.1 | lamin B1 (Lmnb1), mRNA | 2.458 | 1.520 | XM_028885900.1 | CDC like kinase 1 (Clk1), transcript variant X2, mRNA | 0.051 | 0.486 |
| XM_028877411.1 | syntaxin 7 (Stx7), transcript variant X2, mRNA | 2.457 | 1.303 | XM_028880019.1 | CDC42 small effector 2 (Cdc42se2), mRNA | 0.051 | -0.119 |
| XM_028860900.1 | glycogenin 1 (Gyg1), transcript variant X2, mRNA | 2.455 | 1.211 | XM_028853604.1 | Rho GDP dissociation inhibitor alpha (Arhgdia), transcript variant X1, m[illegible] | 0.050 | 0.591 |
| XM_028868424.1 | adenosine deaminase (Ada), transcript variant X1, mRNA | 2.450 | -0.251 | XM_028884055.1 | DnaJ heat shock protein family (Hsp40) member A1 (Dnaja1), transcript | 0.047 | -0.242 |
| XM_028886095.1 | stomatin (Stom), mRNA | 2.431 | 0.920 | XM_028872617.1 | SEC62 homolog, preprotein translocation factor (Sec62), mRNA | 0.045 | 0.049 |
| XM_028874065.1 | elastin microfibril interfacer 2 (Emilin2), mRNA | 2.428 | 2.539 | XM_028876356.1 | chaperonin containing TCP1 subunit 4 (Cct4), mRNA | 0.043 | 0.362 |
| XM_028862659.1 | leukocyte immunoglobulin-like receptor subfamily A member 6 (LOC114688035), mRNA | 2.422 | 2.051 | XM_028880613.1 | heterogeneous nuclear ribonucleoprotein A3 (Hnrnpa3), transcript varia[illegible] | 0.043 | 0.094 |
| XM_028893862.1 | vasodilator stimulated phosphoprotein (Vasp), transcript variant X2, mRNA | 2.390 | -3.207 | XM_028857457.1 | proteasome subunit beta 4 (Psmb4), mRNA | 0.041 | 0.463 |
| XM_028885276.1 | helicase with zinc finger 2 (Helz2), mRNA | 2.381 | -0.779 | XM_028881978.1 | small RNA binding exonuclease protection factor La (Ssb), mRNA | 0.037 | -0.094 |
| XM_028880842.1 | cytochrome b-245 light chain (LOC114700968), mRNA | 2.375 | 1.942 | XM_028862562.1 | RT1 class I histocompatibility antigen, AA alpha chain-like (LOC1146879 | 0.036 | 0.457 |
| XM_028894620.1 | DENN domain containing 5A (Dennd5a), transcript variant X1, mRNA | 2.362 | 0.784 | XM_028863163.1 | heterogeneous nuclear ribonucleoprotein K (Hnrnpk), transcript variant [illegible] | 0.032 | -0.275 |
| XM_028853878.1 | ATPase H+ transporting V0 subunit c (Atp6v0c), mRNA | 2.354 | 1.710 | XM_028885925.1 | TOM1-like protein 1 (LOC114704593), mRNA | 0.030 | -0.108 |
| XM_028863528.1 | leukocyte immunoglobulin-like receptor subfamily B member 3 (LOC114689119), mRNA | 2.330 | 1.604 | XM_028880646.1 | corepressor interacting with RBPJ 1 (LOC114700834), mRNA | 0.030 | 0.300 |
| XM_028895348.1 | mesencephalic astrocyte derived neurotrophic factor (Manf), mRNA | 2.325 | 1.975 | XM_028870936.1 | cytochrome c1, heme protein, mitochondrial (LOC114694169), mRNA | 0.030 | -0.033 |
| XM_028871771.1 | testin LIM domain protein (Tes), mRNA | 2.322 | 0.736 | XM_028887619.1 | erythrocyte membrane protein band 4.1 (Epb41), transcript variant X7, [illegible] | 0.030 | 0.676 |
| XM_028861280.1 | cystatin B (Cstb), mRNA | 2.302 | 2.502 | XM_028877177.1 | store-operated calcium entry associated regulatory factor (Saraf), mRN[illegible] | 0.026 | -0.293 |
| XM_028872450.1 | cyclin dependent kinase inhibitor 2D (Cdkn2d), mRNA | 2.256 | 1.172 | XM_028866877.1 | KIAA0100 ortholog (Kiaa0100), mRNA | 0.025 | -0.140 |
| XM_028877106.1 | interferon gamma receptor 2 (Ifngr2), transcript variant X1, mRNA | 2.249 | 1.136 | XM_028888179.1 | transmembrane protein 59 (Tmem59), mRNA | 0.024 | 0.301 |
| XM_028874469.1 | proline rich 13 (Prr13), transcript variant X1, mRNA | 2.241 | 1.040 | XM_028888449.1 | inflammation and lipid regulator with UBA-like and NBR1-like domains | 0.023 | -0.499 |
| XM_028879067.1 | ST3 beta-galactoside alpha-2,3-sialyltransferase 4 (St3gal4), transcript va | 2.237 | 1.009 | XM_028886488.1 | Y-box binding protein 1 (Ybx1), transcript variant X1, mRNA | 0.023 | -0.463 |
| XM_028865941.1 | annexin A2 (Anxa2), transcript variant X1, mRNA | 2.229 | 0.777 | XM_028874082.1 | lipin 2 (Lpin2), transcript variant X3, mRNA | 0.021 | -0.247 |
| XM_028894336.1 | LYN proto-oncogene, Src family tyrosine kinase (Lyn), transcript variant X2 | 2.206 | 1.193 | XM_028893688.1 | iron-sulfur cluster assembly 1 (Isca1), mRNA | 0.020 | 0.241 |
| XM_028857838.1 | C-type lectin domain family 4 member A-like (LOC114683505), mRNA | 2.201 | 1.602 | XM_028861485.1 | Rho guanine nucleotide exchange factor 1 (Arhgef1), transcript variant [illegible] | 0.020 | 0.178 |
| XM_028879436.1 | CD53 molecule (Cd53), mRNA | 2.180 | 0.896 | XM_028855194.1 | thymosin beta 10 (Tmsb10), mRNA | 0.016 | 1.667 |
| XM_028854964.1 | tetraspanin 14 (Tspan14), transcript variant X2, mRNA | 2.172 | 1.447 | XM_028874547.1 | bridging integrator 2 (Bin2), transcript variant X3, mRNA | 0.016 | -0.201 |
| XM_028865956.1 | cathepsin H (Ctsh), mRNA | 2.169 | 1.066 | XM_028885643.1 | FUS RNA binding protein (Fus), transcript variant X1, mRNA | 0.015 | 0.147 |
| XM_028858602.1 | glycogen phosphorylase L (Pygl), mRNA | 2.159 | 1.838 | XM_028858594.1 | glutathione peroxidase 4 (Gpx4), transcript variant X1, mRNA | 0.014 | 0.272 |
| XM_028860839.1 | selectin P ligand (Selplg), mRNA | 2.141 | 0.749 | XM_028887863.1 | integrin subunit beta 1 (Itgb1), transcript variant X2, mRNA | 0.012 | 0.409 |
| XM_028867469.1 | CD47 molecule (Cd47), transcript variant X2, mRNA | 2.139 | 0.584 | XM_028886730.1 | IK cytokine (Ik), mRNA | 0.012 | 0.099 |
| XR_003733219.1 | ferritin heavy chain pseudogene (LOC114683928), misc_RNA | 2.137 | 1.846 | XM_028890578.1 | C-C motif chemokine 6-like (LOC114707745), mRNA | 0.010 | -0.309 |
| XM_028888276.1 | Ras and Rab interactor 3 (Rin3), transcript variant X3, mRNA | 2.136 | 0.494 | XM_028878644.1 | erythrocyte membrane protein band 4.2 (Epb42), mRNA | 0.010 | 0.503 |
| XM_028876569.1 | protein phosphatase 1 regulatory subunit 15B (Ppp1r15b), mRNA | 2.131 | 0.616 | XM_028864136.1 | Tax1 binding protein 1 (Tax1bp1), transcript variant X1, mRNA | 0.008 | 0.975 |
| XM_028861006.1 | SLP adaptor and CSK interacting membrane protein (Scimp), mRNA | 2.128 | 0.913 | XM_028866142.1 | bisphosphoglycerate mutase (Bpgm), transcript variant X1, mRNA | 0.008 | 0.083 |
| XM_028859848.1 | leukocyte surface antigen CD53-like (LOC114685266), mRNA | 2.122 | 0.282 | XM_028876147.1 | serine and arginine rich splicing factor 5 (Srsf5), transcript variant X4, m[illegible] | 0.005 | -0.081 |
| XM_028873576.1 | galectin 3 (Lgals3), mRNA | 2.115 | 2.174 | XM_028865299.1 | family with sequence similarity 104 member A (Fam104a), transcript v[illegible] | 0.004 | -0.033 |

| | | | | | | | | |
|---|---|---|---|---|---|---|---|---|
| XM_028878161.1 | lymphocyte cytosolic protein 1 (Lcp1), transcript variant X1, mRNA | 2.114 | 0.629 | Table S18 | XM_028860198.1 | chaperonin containing TCP1 subunit 7 (Cct7), mRNA | 0.004 | 0.227 |
| XM_028876075.1 | MYC associated factor X (Max), transcript variant X1, mRNA | 2.102 | 0.550 | | XM_028854748.1 | ring finger protein 10 (Rnf10), transcript variant X2, mRNA | 0.002 | 0.200 |
| XM_028870335.1 | cathepsin B-like (LOC114693805), mRNA | 2.097 | 1.611 | | XM_028863340.1 | schlafen family member 12-like (LOC114688842), mRNA | 0.001 | 0.008 |
| XM_028865640.1 | signaling lymphocytic activation molecule family member 1 (Slamf1), mRNA | 2.090 | 0.273 | | XM_028875963.1 | chaperonin containing TCP1 subunit 8 (Cct8), mRNA | -0.001 | 0.348 |
| XM_028878171.1 | lymphocyte cytosolic protein 1 (Lcp1), transcript variant X2, mRNA | 2.088 | 0.950 | | XM_028857431.1 | selenium binding protein 1 (Selenbp1), mRNA | -0.002 | 0.085 |
| XM_028862653.1 | deltex E3 ubiquitin ligase 3L (Dtx3l), mRNA | 2.085 | -0.002 | | XM_028857931.1 | metastasis associated 1 family member 2 (Mta2), transcript variant X1, | -0.002 | -0.750 |
| XM_028891273.1 | transketolase (Tkt), mRNA | 2.085 | 0.979 | | XM_028892763.1 | GABA type A receptor associated protein like 2 (Gabarapl2), mRNA | -0.005 | 0.039 |
| XM_028894103.1 | negative regulator of reactive oxygen species (Nrros), transcript variant X | 2.082 | 1.414 | | XM_028894655.1 | NDUFA4 mitochondrial complex associated (Ndufa4), mRNA | -0.005 | 0.463 |
| XM_028863247.1 | leukocyte-associated immunoglobulin-like receptor 2 (LOC114688720), mRNA | 2.074 | 1.183 | | XM_028869647.1 | lysozyme C-2-like (LOC114693318), mRNA | -0.007 | 1.902 |
| XM_028876496.1 | MAPK activated protein kinase 2 (Mapkapk2), mRNA | 2.073 | 0.877 | | XM_028867919.1 | RAD21 cohesin complex component (Rad21), mRNA | -0.007 | -0.038 |
| XM_028866422.1 | junction adhesion molecule like (Jaml), transcript variant X1, mRNA | 2.066 | 0.439 | | XM_028868882.1 | target of myb1 like 1 membrane trafficking protein (Tom1l1), mRNA | -0.011 | 0.107 |
| XM_028870099.1 | interferon regulatory factor 7 (Irf7), transcript variant X1, mRNA | 2.057 | 0.201 | | XM_028891466.1 | TNF receptor superfamily member 18 (Tnfrsf18), transcript variant X1, r | -0.012 | -0.220 |
| XM_028861387.1 | leukocyte immunoglobulin-like receptor subfamily B member 3 (LOC114686718), partial mRNA | 2.056 | 1.717 | | XM_028854746.1 | ankyrin repeat domain 13A (Ankrd13a), transcript variant X2, mRNA | -0.016 | -0.128 |
| XM_028884166.1 | cathepsin B (Ctsb), mRNA | 2.051 | 2.282 | | XM_028892146.1 | short chain dehydrogenase/reductase family 39U member 1 (Sdr39u1), | -0.016 | -0.452 |
| XM_028860146.1 | TNF receptor superfamily member 1B (Tnfrsf1b), mRNA | 2.050 | 1.626 | | XM_028888164.1 | DnaJ heat shock protein family (Hsp40) member C8 (Dnajc8), mRNA | -0.020 | -0.252 |
| XM_028877562.1 | ArfGAP with RhoGAP domain, ankyrin repeat and PH domain 1 (Arap1), transcript variant X3, mRNA | 2.049 | 1.205 | | XM_028886768.1 | poly(A) binding protein interacting protein 2 (Paip2), mRNA | -0.020 | 0.070 |
| XM_028889281.1 | ERV-BabFcenv provirus ancestral Env polyprotein (LOC114706804), mRNA | 2.046 | 0.538 | | XM_028883198.1 | BCL2/adenovirus E1B 19 kDa protein-interacting protein 3-like (LOC1147 | -0.021 | -0.292 |
| XM_028874893.1 | osteoclast stimulating factor 1 (Ostf1), mRNA | 2.034 | 1.495 | | XM_028872762.1 | proline rich nuclear receptor coactivator 1 (Pnrc1), mRNA | -0.021 | 0.137 |
| XM_028892930.1 | F-box and leucine rich repeat protein 5 (Fbxl5), mRNA | 2.027 | 1.134 | | XM_028888103.1 | SH3 domain binding glutamate rich protein like 3 (Sh3bgrl3), mRNA | -0.022 | 0.621 |
| XM_028871737.1 | unc-93 homolog B1, TLR signaling regulator (Unc93b1), mRNA | 2.027 | 1.511 | | XM_028892106.1 | proteasome activator subunit 2 (Psme2), mRNA | -0.027 | 0.219 |
| XM_028880668.1 | EF-hand domain family member D2 (Efhd2), mRNA | 2.025 | 1.666 | | XM_028870941.1 | MAF1 homolog, negative regulator of RNA polymerase III (Maf1), mRN | -0.030 | -0.198 |
| XM_028880790.1 | KIAA0513 ortholog (Kiaa0513), transcript variant X2, mRNA | 2.021 | 1.101 | | XM_028854098.1 | CCHC-type zinc finger nucleic acid binding protein (Cnbp), transcript vari | -0.031 | 0.876 |
| XM_028867903.1 | interleukin 4 receptor (Il4r), mRNA | 1.999 | 1.127 | | XM_028859870.1 | heterogeneous nuclear ribonucleoprotein U like 1 (Hnrnpul1), transcript | -0.032 | -0.383 |
| XM_028893864.1 | vasodilator stimulated phosphoprotein (Vasp), transcript variant X3, mRNA | 1.966 | 0.805 | | XM_028857223.1 | S-phase kinase associated protein 1 (Skp1), mRNA | -0.035 | 0.124 |
| XM_028855812.1 | ring finger protein 19B (Rnf19b), mRNA | 1.966 | 0.520 | | XM_028878628.1 | glycerophosphocholine phosphodiesterase 1 (Gpcpd1), transcript variant | -0.036 | 0.239 |
| XM_028873761.1 | FES proto-oncogene, tyrosine kinase (Fes), transcript variant X1, mRNA | 1.962 | 1.512 | | XM_028861615.1 | ATP synthase F1 subunit beta (Atp5f1b), mRNA | -0.036 | 0.272 |
| XM_028870218.1 | neutrophil cytosolic factor 1 (Ncf1), mRNA | 1.958 | 1.790 | | XM_028874242.1 | macrophage migration inhibitory factor (Mif), mRNA | -0.037 | 0.681 |
| XM_028880022.1 | TNFAIP3 interacting protein 1 (Tnip1), transcript variant X1, mRNA | 1.940 | 1.450 | | XM_028890891.1 | HCLS1 associated protein X-1 (Hax1), mRNA | -0.038 | 0.223 |
| XM_028880653.1 | peptidyl arginine deiminase 4 (Padi4), mRNA | 1.940 | 0.474 | | XM_028863039.1 | NADH:ubiquinone oxidoreductase subunit A3 (Ndufa3), mRNA | -0.040 | 0.196 |
| XM_028878709.1 | protein disulfide isomerase family A member 3 (Pdia3), mRNA | 1.930 | 1.304 | | XR_003736307.1 | uncharacterized LOC114704708 (LOC114704708), ncRNA | -0.040 | -0.566 |
| XM_028856266.1 | cathepsin S (Ctss), mRNA | 1.920 | 1.667 | | XM_028888579.1 | family with sequence similarity 50 member A (Fam50a), mRNA | -0.041 | -0.256 |
| XM_028876076.1 | MYC associated factor X (Max), transcript variant X2, mRNA | 1.916 | -0.096 | | XM_028866673.1 | t-complex 1 (Tcp1), mRNA | -0.042 | 0.252 |
| XM_028859890.1 | transforming growth factor beta 1 (Tgfb1), mRNA | 1.900 | 1.165 | | XM_028858593.1 | RNA polymerase II subunit E (Polr2e), mRNA | -0.043 | -0.222 |
| XM_028863506.1 | NLR family pyrin domain containing 12 (Nlrp12), partial mRNA | 1.900 | -0.032 | | XR_003734001.1 | uncharacterized LOC114689979 (LOC114689979), transcript variant X1, r | -0.044 | -0.629 |
| XM_028877112.1 | interleukin 10 receptor subunit beta (Il10rb), mRNA | 1.897 | 0.991 | | XM_028862222.1 | deoxyribonuclease 2, lysosomal (Dnase2), mRNA | -0.048 | 0.992 |
| XM_028856401.1 | MYD88 innate immune signal transduction adaptor (Myd88), mRNA | 1.889 | 1.110 | | XM_028877794.1 | glucose-6-phosphate isomerase (Gpi), transcript variant X1, mRNA | -0.049 | 0.029 |
| XM_028893861.1 | vasodilator stimulated phosphoprotein (Vasp), transcript variant X1, mRNA | 1.887 | 1.398 | | XM_028887214.1 | ribosomal protein L7 like 1 (Rpl7l1), mRNA | -0.049 | -0.061 |
| XM_028862627.1 | C-type lectin domain family 4 member D (Clec4d), partial mRNA | 1.886 | 1.297 | | XM_028876534.1 | transmembrane and coiled-coil domain family 2 (Tmcc2), transcript var | -0.051 | -0.127 |
| XM_028885171.1 | copine 3 (Cpne3), transcript variant X1, mRNA | 1.880 | -0.473 | | XM_028875447.1 | proline rich nuclear receptor coactivator 2 (Pnrc2), mRNA | -0.051 | 0.160 |
| XM_028886072.1 | heat shock protein family A (Hsp70) member 5 (Hspa5), transcript variant X1 | 1.878 | 1.344 | | XM_028859367.1 | U2 small nuclear RNA auxiliary factor 2 (U2af2), transcript variant X2, n | -0.052 | 0.125 |
| XM_028869109.1 | LIM and SH3 protein 1 (Lasp1), mRNA | 1.877 | 0.864 | | XM_028869049.1 | pyridoxamine 5'-phosphate oxidase (Pnpo), mRNA | -0.052 | 0.445 |
| XM_028862227.1 | mannosidase alpha class 2B member 1 (Man2b1), mRNA | 1.872 | 1.296 | | XM_028876482.1 | cathepsin E (Ctse), mRNA | -0.052 | -0.227 |
| XM_028874737.1 | proteoglycan 4 (Prg4), transcript variant X7, mRNA | 1.871 | -6.919 | | XM_028883012.1 | thioredoxin (Txn), transcript variant X1, mRNA | -0.053 | 0.436 |
| XM_028874470.1 | proline rich 13 (Prr13), transcript variant X2, mRNA | 1.866 | 1.668 | | XM_028858548.1 | basigin (Ok blood group) (Bsg), transcript variant X3, mRNA | -0.054 | 0.190 |
| XM_028862833.1 | adhesion G protein-coupled receptor E5 (Adgre5), transcript variant X2, mRNA | 1.859 | 0.682 | | XM_028861786.1 | RAD23 homolog A, nucleotide excision repair protein (Rad23a), transcri | -0.057 | 0.703 |
| XM_028888858.1 | oxysterol binding protein like 9 (Osbpl9), transcript variant X8, mRNA | 1.855 | 1.060 | | XM_028892172.1 | abhydrolase domain containing 4 (Abhd4), transcript variant X3, mRNA | -0.057 | -0.528 |
| XM_028867976.1 | calcineurin B homologous protein 1 (LOC114692301), mRNA | 1.847 | 0.950 | | XM_028867790.1 | glycerophosphodiester phosphodiesterase 1 (Gde1), transcript variant X | -0.059 | 0.263 |
| XM_028868580.1 | protein tyrosine phosphatase non-receptor type 1 (Ptpn1), mRNA | 1.847 | 1.456 | | XM_028875715.1 | c-Maf inducing protein (Cmip), transcript variant X1, mRNA | -0.060 | -0.288 |
| XM_028856799.1 | signal transducer and activator of transcription 3 (Stat3), transcript variant X1, mRNA | 1.840 | 0.541 | | XM_028857969.1 | nuclear RNA export factor 1 (Nxf1), mRNA | -0.061 | -0.119 |
| XM_028861651.1 | CMRF35-like molecule 8 (LOC114686954), mRNA | 1.838 | 1.312 | | XM_028880246.1 | methionyl aminopeptidase 2 (Metap2), mRNA | -0.061 | 0.256 |
| XM_028863405.1 | core histone macro-H2A.1 (LOC114688930), transcript variant X2, mRNA | 1.836 | 0.261 | | XM_028874224.1 | small nuclear ribonucleoprotein D3 polypeptide (Snrpd3), transcript vari | -0.062 | 0.577 |
| XM_028859915.1 | myosin IF (Myo1f), mRNA | 1.832 | 1.103 | | XM_028892348.1 | chromosome unknown C12orf57 homolog (CUNH12orf57), mRNA | -0.068 | 0.676 |
| XM_028873763.1 | furin, paired basic amino acid cleaving enzyme (Furin), transcript variant X1, mRNA | 1.831 | 0.608 | | XM_028888245.1 | NADH:ubiquinone oxidoreductase subunit B1 (Ndufb1), mRNA | -0.068 | 0.796 |
| XM_028888608.1 | filamin A (Flna), transcript variant X1, mRNA | 1.811 | 0.782 | | XM_028863691.1 | schlafen family member 12-like (LOC114689363), mRNA | -0.069 | -0.142 |
| XM_028886393.1 | phosphoglycerate kinase 1 (Pgk1), mRNA | 1.800 | 1.369 | | XM_028887419.1 | SEC61 translocon gamma subunit (Sec61g), transcript variant X1, mRNA | -0.069 | 0.585 |
| XM_028866914.1 | WD repeat and SOCS box containing 1 (Wsb1), transcript variant X1, mRNA | 1.796 | 1.241 | | XM_028861277.1 | SS18 subunit of BAF chromatin remodeling complex (Ss18), mRNA | -0.071 | 0.155 |
| XM_028894335.1 | LYN proto-oncogene, Src family tyrosine kinase (Lyn), transcript variant X1 | 1.794 | 1.440 | | XM_028877621.1 | exportin 7 (Xpo7), transcript variant X1, mRNA | -0.071 | -0.523 |
| XM_028860275.1 | L-lactate dehydrogenase A chain (LOC114685644), mRNA | 1.782 | 1.332 | | XM_028854074.1 | CCHC-type zinc finger nucleic acid binding protein (Cnbp), transcript vari | -0.073 | 0.478 |
| XM_028874400.1 | integrin subunit alpha 5 (Itga5), mRNA | 1.778 | 0.839 | | XM_028859669.1 | mitochondrial contact site and cristae organizing system subunit 13 (Mi | -0.075 | 0.551 |
| XM_028873201.1 | spermidine/spermine N1-acetyltransferase 1 (Sat1), transcript variant X1, mRNA | 1.777 | 1.327 | | XM_028856470.1 | zinc finger CCHC domain-containing protein 3-like (LOC114682549), mR | -0.075 | -1.812 |
| XM_028889719.1 | UBA like domain containing 2 (Ubald2), mRNA | 1.767 | 0.602 | | XM_028883976.1 | family with sequence similarity 214 member B (Fam214b), transcript va | -0.076 | -0.592 |
| XM_028892317.1 | CD9 molecule (Cd9), mRNA | 1.767 | 0.001 | | XM_028871458.1 | ADP ribosylation factor like GTPase 2 binding protein (Arl2bp), mRNA | -0.077 | -0.264 |
| XM_028876501.1 | inhibitor of nuclear factor kappa B kinase subunit epsilon (Ikbke), mRNA | 1.760 | 1.339 | | XM_028854088.1 | CCHC-type zinc finger nucleic acid binding protein (Cnbp), transcript vari | -0.079 | 0.738 |
| XM_028863251.1 | immunoglobulin alpha Fc receptor-like (LOC114688723), mRNA | 1.746 | 0.728 | | XM_028870191.1 | RAN, member RAS oncogene family (Ran), transcript variant X1, mRNA | -0.079 | 0.225 |
| XM_028882412.1 | Rho guanine nucleotide exchange factor 3 (Arhgef3), transcript variant X3, | 1.741 | -0.085 | | XM_028886024.1 | protein tyrosine phosphatase 4A3 (Ptp4a3), transcript variant X1, mRNA | -0.080 | -0.185 |
| XM_028862383.1 | sorcin (Sri), transcript variant X1, mRNA | 1.736 | 0.899 | | XM_028891847.1 | solute carrier family 48 member 1 (Slc48a1), mRNA | -0.081 | 0.026 |
| XM_028886159.1 | V-set immunoregulatory receptor (Vsir), mRNA | 1.735 | 0.903 | | XM_028865564.1 | transgelin 2 (Tagln2), mRNA | -0.082 | -0.032 |
| XM_028888040.1 | chloride intracellular channel 1 (Clic1), mRNA | 1.730 | 1.288 | | XM_028883623.1 | t-complex 11 like 2 (Tcp11l2), mRNA | -0.083 | -0.573 |
| XM_028861273.1 | syntaxin 11 (Stx11), mRNA | 1.729 | -0.072 | | XM_028885911.1 | basic leucine zipper and W2 domains 1 (Bzw1), transcript variant X1, m | -0.084 | -0.235 |
| XM_028889400.1 | phosphoglycerate mutase 1 (Pgam1), mRNA | 1.723 | 2.005 | | XM_028877910.1 | cell cycle associated protein 1 (Caprin1), mRNA | -0.084 | -0.027 |
| XM_028860315.1 | dynein light chain Tctex-type 1 (Dynlt1), mRNA | 1.719 | 1.203 | | XM_028885891.1 | CDC like kinase 1 (Clk1), transcript variant X1, mRNA | -0.084 | -1.052 |
| XM_028865789.1 | mitogen-activated protein kinase kinase kinase kinase 4 (Map4k4), transcript variant X21, mRNA | 1.701 | -0.445 | | XM_028876687.1 | RIO kinase 3 (Riok3), mRNA | -0.086 | 0.820 |
| XM_028890747.1 | zinc finger and BTB domain containing 7B (Zbtb7b), mRNA | 1.698 | 0.757 | | XM_028864159.1 | heterogeneous nuclear ribonucleoprotein A2/B1 (Hnrnpa2b1), transcript | -0.089 | -0.012 |
| XM_028861094.1 | WD repeat-containing protein 26-like (LOC114686409), mRNA | 1.689 | 0.422 | | XM_028865188.1 | SWI/SNF related, matrix associated, actin dependent regulator of chron | -0.091 | 0.072 |
| XM_028891603.1 | serpin family B member 10 (Serpinb10), mRNA | 1.686 | 1.766 | | XM_028884346.1 | transmembrane protein 258 (Tmem258), mRNA | -0.092 | 0.421 |
| XM_028892933.1 | ADP-ribosyl cyclase/cyclic ADP-ribose hydrolase 1-like (LOC114709225), mRNA | 1.686 | -0.041 | | XM_028857265.1 | H/ACA ribonucleoprotein complex subunit 3 (LOC114683088), mRNA | -0.093 | 0.383 |
| XM_028858198.1 | dehydrogenase/reductase SDR family member 7 (LOC114683839), mRNA | 1.660 | 0.459 | | XM_028855811.1 | RB binding protein 4, chromatin remodeling factor (Rbbp4), mRNA | -0.095 | -0.105 |
| XM_028887935.1 | lymphotoxin beta (Ltb), transcript variant X1, mRNA | 1.657 | 0.458 | | XM_028866415.1 | ATP synthase membrane subunit g (Atp5mg), mRNA | -0.098 | 0.382 |
| XM_028859534.1 | RAS guanyl releasing protein 4 (Rasgrp4), mRNA | 1.654 | 1.684 | | XM_028889268.1 | synuclein alpha (Snca), transcript variant X1, mRNA | -0.099 | -0.210 |
| XM_028855193.1 | capping actin protein, gelsolin like (Capg), mRNA | 1.652 | 0.876 | | XM_028861048.1 | H-2 class I histocompatibility antigen, Q10 alpha chain-like (LOC11468 | -0.099 | -0.008 |
| XM_028894410.1 | NADPH--cytochrome P450 reductase (LOC114710307), transcript variant X2, mRN | 1.647 | 0.769 | | XM_028891283.1 | protein kinase C delta (Prkcd), transcript variant X2, mRNA | -0.101 | 0.083 |
| XM_028890794.1 | S100 calcium binding protein A6 (S100a6), transcript variant X1, mRNA | 1.642 | 1.451 | | XM_028874959.1 | C-X-C motif chemokine receptor 4 (Cxcr4), transcript variant X1, mRNA | -0.101 | -0.357 |
| XM_028862603.1 | paired immunoglobulin-like type 2 receptor alpha (LOC114687969), mRNA | 1.635 | 1.632 | | XM_028861504.1 | Rab acceptor 1 (Rabac1), mRNA | -0.101 | 0.755 |
| XM_028892346.1 | protein tyrosine phosphatase non-receptor type 6 (Ptpn6), transcript varian | 1.634 | 0.402 | | XM_028863861.1 | chaperonin containing TCP1 subunit 3 (Cct3), mRNA | -0.101 | 0.069 |
| XM_028863253.1 | leukocyte-associated immunoglobulin-like receptor 2 (LOC114688724), mRNA | 1.633 | 0.708 | | XM_028869483.1 | ArfGAP with coiled-coil, ankyrin repeat and PH domains 1 (Acap1), mRN | -0.102 | -0.302 |
| XM_028892345.1 | protein tyrosine phosphatase non-receptor type 6 (Ptpn6), transcript varian | 1.623 | 1.081 | | XM_028890537.1 | dehydrogenase/reductase 11 (Dhrs11), mRNA | -0.102 | 0.041 |
| XM_028890476.1 | zinc finger DHHC-type containing 3 (Zdhhc3), transcript variant X2, mRNA | 1.619 | 1.096 | | XM_028884481.1 | cytochrome c oxidase subunit 6C-2 (LOC114703635), mRNA | -0.103 | 0.524 |
| XM_028861207.1 | adhesion G protein-coupled receptor E1 (Adgre1), mRNA | 1.613 | 2.100 | | XM_028877148.1 | superoxide dismutase 1 (Sod1), mRNA | -0.108 | -0.059 |
| XM_028886267.1 | beta-1,4-N-acetyl-galactosaminyltransferase 1 (B4galnt1), transcript varian | 1.609 | 0.326 | | XM_028868234.1 | serine/threonine kinase 24 (Stk24), transcript variant X2, mRNA | -0.109 | -0.086 |
| XM_028869603.1 | glucosamine (N-acetyl)-6-sulfatase (Gns), mRNA | 1.605 | 1.592 | | XM_028886290.1 | CTD small phosphatase 2 (Ctdsp2), transcript variant X1, mRNA | -0.111 | -0.215 |
| XM_028856869.1 | interferon induced protein 35 (Ifi35), mRNA | 1.601 | 0.903 | | XM_028891952.1 | tubulin alpha-1B chain (LOC114708578), mRNA | -0.112 | 0.400 |
| XM_028865588.1 | CD48 molecule (Cd48), mRNA | 1.585 | 1.087 | | XM_028862540.1 | DENN domain containing 1C (Dennd1c), mRNA | -0.112 | -0.168 |
| XM_028866150.1 | RAB8B, member RAS oncogene family (Rab8b), mRNA | 1.564 | 0.866 | | XM_028860991.1 | ring finger protein 167 (Rnf167), transcript variant X2, mRNA | -0.114 | -0.251 |
| XM_028856555.1 | paxillin (Pxn), transcript variant X3, mRNA | 1.556 | 0.607 | | XR_003733960.1 | heterogeneous nuclear ribonucleoprotein A2/B1 (Hnrnpa2b1), transcript | -0.114 | -0.048 |
| XM_028894899.1 | lymphocyte cytosolic protein 2 (Lcp2), transcript variant X1, mRNA | 1.553 | 0.618 | | XM_028859107.1 | interferon-induced protein with tetratricopeptide repeats 1-like (LOC11 | -0.118 | 0.996 |
| XM_028867399.1 | transmembrane p24 trafficking protein 5 (Tmed5), transcript variant X1, mRNA | 1.549 | 0.588 | | XM_028864295.1 | methionine adenosyltransferase 2B (Mat2b), transcript variant X1, mRN | -0.124 | 0.416 |
| XM_028884971.1 | complement factor properdin (Cfp), transcript variant X2, mRNA | 1.546 | 1.790 | | XM_028881670.1 | eukaryotic translation initiation factor 2 subunit gamma (Eif2s3), mRNA | -0.127 | 0.223 |
| XM_028870604.1 | X-box binding protein 1 (Xbp1), mRNA | 1.545 | 1.317 | | XM_028873926.1 | protein phosphatase 1 catalytic subunit beta (Ppp1cb), mRNA | -0.128 | 0.114 |
| XM_028878056.1 | ubiquitin like 3 (Ubl3), mRNA | 1.539 | 1.012 | | XM_028861919.1 | centrin-2 (LOC114687209), mRNA | -0.130 | -0.308 |
| XM_028880448.1 | interleukin 17 receptor A (Il17ra), transcript variant X1, mRNA | 1.534 | 0.800 | | XM_028887601.1 | BSD domain containing 1 (Bsdc1), mRNA | -0.131 | 0.211 |
| XM_028893115.1 | tubulin alpha-4A chain-like (LOC114709335), mRNA | 1.527 | 0.699 | | XM_028878551.1 | small nuclear ribonucleoprotein polypeptides B and B1 (Snrpb), mRNA | -0.131 | 0.203 |
| XM_028892991.1 | CKLF like MARVEL transmembrane domain containing 6 (Cmtm6), mRNA | 1.525 | 0.920 | | XM_028873590.1 | cytohesin 1 interacting protein (Cytip), mRNA | -0.131 | -0.457 |
| XM_028857646.1 | CD5 molecule like (Cd5l), mRNA | 1.521 | -0.983 | | XM_028858282.1 | signal peptidase complex subunit 3 (Spcs3), mRNA | -0.131 | 0.523 |
| XM_028861559.1 | tripartite motif-containing protein 30A-like (LOC114686891), transcript variant X3, mRNA | 1.511 | 0.410 | | XM_028881248.1 | RAS p21 protein activator 3 (Rasa3), transcript variant X2, mRNA | -0.133 | -0.387 |
| XM_028886310.1 | poly(ADP-ribose) polymerase family member 14 (Parp14), mRNA | 1.505 | -0.097 | | XM_028886098.1 | N-acetyllactosaminide alpha-1,3-galactosyltransferase-like (LOC114704 | -0.133 | -0.068 |
| XM_028892472.1 | pyruvate kinase M1/2 (Pkm), transcript variant X3, mRNA | 1.501 | 0.394 | | XM_028880044.1 | G3BP stress granule assembly factor 1 (G3bp1), transcript variant X1, m | -0.136 | 0.029 |
| XM_028879641.1 | protein disulfide isomerase family A member 4 (Pdia4), mRNA | 1.500 | 2.266 | | XM_028874879.1 | nuclear receptor binding protein 1 (Nrbp1), transcript variant X2, mRNA | -0.136 | 0.251 |
| XM_028884228.1 | membrane-spanning 4-domains subfamily A member 6B-like (LOC114703404), mRNA | 1.494 | 1.334 | | XM_028871151.1 | DEAD-box helicase 17 (Ddx17), transcript variant X1, mRNA | -0.137 | -0.359 |
| XM_028883834.1 | ATPase H+ transporting V1 subunit B2 (Atp6v1b2), mRNA | 1.484 | 1.207 | | XM_028875464.1 | transmembrane protein 50A (Tmem50a), mRNA | -0.138 | 0.178 |
| XM_028885483.1 | disintegrin and metalloproteinase domain-containing protein 19 (LOC11470432 | 1.479 | -0.436 | | XM_028890510.1 | TATA-box binding protein associated factor 15 (Taf15), mRNA | -0.140 | -0.044 |
| XM_028874143.1 | uncharacterized LOC114696446 (LOC114696446), mRNA | 1.470 | -0.873 | | XM_028887397.1 | LFNG O-fucosylpeptide 3-beta-N-acetylglucosaminyltransferase (Lfng), | -0.140 | 1.125 |
| XM_028863767.1 | uncharacterized LOC114689465 (LOC114689465), mRNA | 1.468 | -0.844 | | XM_028857149.1 | tripartite motif containing 58 (Trim58), mRNA | -0.143 | -0.016 |
| XM_028887201.1 | cyclin D3 (Ccnd3), transcript variant X2, mRNA | 1.462 | -0.218 | | XM_028888711.1 | ATPase sarcoplasmic/endoplasmic reticulum Ca2+ transporting 3 (Atp2 | -0.144 | -0.396 |
| XM_028888459.1 | glucose-6-phosphate dehydrogenase (G6pd), mRNA | 1.456 | 0.824 | | XM_028861884.1 | aldehyde dehydrogenase 16 family member A1 (Aldh16a1), mRNA | -0.144 | 0.127 |
| XM_028853687.1 | reticulon 3 (Rtn3), transcript variant X4, mRNA | 1.453 | 0.876 | | XM_028892836.1 | splicing factor 3b subunit 2 (Sf3b2), transcript variant X1, mRNA | -0.144 | -0.151 |
| XM_028878543.1 | signal regulatory protein alpha (Sirpa), transcript variant X2, mRNA | 1.449 | 1.413 | | XM_028889014.1 | eukaryotic translation initiation factor 6 (Eif6), transcript variant X1, m | -0.144 | 0.567 |
| XM_028884620.1 | regulator of G protein signaling 2 (Rgs2), mRNA | 1.443 | -0.180 | | XM_028866356.1 | hydroxymethylbilane synthase (Hmbs), transcript variant X1, mRNA | -0.145 | -0.130 |
| XM_028854937.1 | adaptor related protein complex 2 subunit sigma 1 (Ap2s1), mRNA | 1.432 | 0.867 | | XM_028866369.1 | peptidylprolyl isomerase B (Ppib), mRNA | -0.145 | 0.542 |
| XM_028874408.1 | nuclear factor, erythroid 2 (Nfe2), mRNA | 1.430 | 0.559 | | XM_028854081.1 | CCHC-type zinc finger nucleic acid binding protein (Cnbp), transcript vari | -0.147 | 0.645 |
| XM_028866098.1 | myotrophin (Mtpn), mRNA | 1.419 | 0.947 | | XM_028888513.1 | membrane palmitoylated protein 1 (Mpp1), mRNA | -0.149 | 0.451 |
| XM_028866776.1 | refilin B (Rflnb), mRNA | 1.418 | -0.661 | | XM_028867001.1 | schlafen family member 5 (Slfn5), transcript variant X1, mRNA | -0.151 | -6.623 |
| XM_028861210.1 | vav guanine nucleotide exchange factor 1 (Vav1), mRNA | 1.408 | 0.739 | | XM_028861880.1 | CD37 molecule (Cd37), transcript variant X1, mRNA | -0.152 | -0.700 |
| XM_028888175.1 | ribosomal protein S6 kinase A1 (Rps6ka1), transcript variant X2, mRNA | 1.404 | 0.945 | | XM_028885354.1 | proteasome subunit alpha 7 (Psma7), mRNA | -0.154 | 0.520 |
| XM_028891829.1 | sorting nexin 20 (Snx20), mRNA | 1.403 | 0.626 | | XM_028889018.1 | RALY heterogeneous nuclear ribonucleoprotein (Raly), transcript variant | -0.156 | -0.123 |
| XM_028888713.1 | NACHT, LRR and PYD domains-containing protein 1b allele 4-like (LOC11470633 | 1.402 | -0.130 | | XM_028892582.1 | baculoviral IAP repeat containing 5 (Birc5), mRNA | -0.156 | -2.313 |
| XM_028871084.1 | Rac family small GTPase 2 (Rac2), mRNA | 1.399 | 0.184 | | XM_028884598.1 | integrin subunit alpha 4 (Itga4), mRNA | -0.158 | 0.063 |
| XM_028862635.1 | NACHT, LRR and PYD domains-containing protein 1a-like (LOC114688014), mRNA | 1.395 | 0.231 | | XM_028863433.1 | glutaryl-CoA dehydrogenase, mitochondrial-like (LOC114688976), mRNA | -0.159 | 0.100 |
| XM_028855174.1 | trans-golgi network protein 2 (Tgoln2), transcript variant X2, mRNA | 1.386 | 0.203 | | XM_028882040.1 | chaperonin containing TCP1 subunit 5 (Cct5), mRNA | -0.161 | 0.062 |
| XM_028864728.1 | grancalcin (Gca), mRNA | 1.384 | 0.362 | | XM_028893479.1 | SIN3 transcription regulator family member B (Sin3b), transcript variant | -0.162 | 0.572 |
| XM_028863402.1 | core histone macro-H2A.1 (LOC114688930), transcript variant X1, mRNA | 1.384 | 0.516 | | XM_028877289.1 | terminal nucleotidyltransferase 5C (Tent5c), mRNA | -0.165 | 0.313 |
| XM_028863332.1 | leukocyte immunoglobulin-like receptor subfamily A member 6 (LOC114688828), mRNA | 1.382 | 1.381 | | XM_028893533.1 | Kruppel like factor 2 (Klf2), mRNA | -0.166 | -0.694 |
| XM_028858757.1 | uncharacterized LOC114684244 (LOC114684244), mRNA | 1.381 | -0.981 | | XM_028879867.1 | cytidine monophosphate N-acetylneuraminic acid synthetase (Cmas), tr | -0.166 | 0.549 |
| XM_028887200.1 | cyclin D3 (Ccnd3), transcript variant X1, mRNA | 1.374 | -0.454 | | XM_028886764.1 | ubiquitin conjugating enzyme E2 D2 (Ube2d2), mRNA | -0.166 | -0.042 |
| XM_028863524.1 | docking protein 3 (Dok3), mRNA | 1.369 | 1.249 | | XM_028890695.1 | heat shock protein 90 alpha family class B member 1 (Hsp90ab1), mRN | -0.167 | 0.281 |
| XM_028874204.1 | RAB10, member RAS oncogene family (Rab10), mRNA | 1.366 | 0.502 | | XM_028880612.1 | heterogeneous nuclear ribonucleoprotein A3 (Hnrnpa3), transcript varia | -0.170 | 0.230 |
| XM_028856274.1 | extracellular matrix protein 1 (Ecm1), transcript variant X1, mRNA | 1.364 | 0.364 | | XM_028886213.1 | signal transducer and activator of transcription 6 (Stat6), transcript vari | -0.170 | -0.326 |
| XM_028882605.1 | granulin precursor (Grn), mRNA | 1.358 | 1.809 | | XM_028892105.1 | proteasome activator subunit 1 (Psme1), transcript variant X2, mRNA | -0.175 | 0.281 |

| | | | |
|---|---|---|---|
| XM_028885179.1 | carbonic anhydrase 1 (Ca1), mRNA | 1.355 | -0.417 |
| XM_028863929.1 | actin related protein 2/3 complex subunit 4 (Arpc4), mRNA | 1.352 | 1.432 |
| XM_028871110.1 | TRIO and F-actin binding protein (Triobp), transcript variant X1, mRNA | 1.338 | 0.831 |
| XM_028893563.1 | annexin A11 (Anxa11), transcript variant X1, mRNA | 1.333 | 0.779 |
| XM_028888232.1 | interferon alpha-inducible protein 27-like protein 2A (LOC114706032), mRNA | 1.330 | 0.995 |
| XM_028864498.1 | selectin L (Sell), mRNA | 1.326 | 0.845 |
| XM_028874319.1 | integrin subunit beta 2 (Itgb2), transcript variant X1, mRNA | 1.314 | -0.215 |
| XM_028890795.1 | S100 calcium binding protein A6 (S100a6), transcript variant X2, mRNA | 1.313 | 1.397 |
| XM_028870030.1 | fibronectin 1 (Fn1), mRNA | 1.300 | 2.731 |
| XM_028878732.1 | beta-2-microglobulin (B2m), mRNA | 1.294 | 0.301 |
| XM_028885899.1 | actin related protein 2/3 complex subunit 3 (Arpc3), mRNA | 1.287 | 1.110 |
| XM_028875596.1 | FYN binding protein 1 (Fyb1), mRNA | 1.280 | 0.078 |
| XM_028860500.1 | Y-box binding protein 3 (Ybx3), transcript variant X1, mRNA | 1.278 | -0.310 |
| XR_003735222.1 | glyceraldehyde-3-phosphate dehydrogenase pseudogene (LOC114697622), misc_RN | 1.269 | 1.298 |
| XM_028860217.1 | lysine rich coiled-coil 1 (Krcc1), mRNA | 1.261 | 0.768 |
| XM_028857674.1 | interferon regulatory factor 2 (Irf2), transcript variant X1, mRNA | 1.261 | -0.235 |
| XM_028893120.1 | transmembrane BAX inhibitor motif containing 1 (Tmbim1), transcript variant | 1.257 | 0.373 |
| XM_028890701.1 | phospholipase A2 group VII (Pla2g7), mRNA | 1.254 | 1.350 |
| XM_028867603.1 | ras homolog family member B (Rhob), mRNA | 1.238 | -0.903 |
| XM_028865102.1 | signal sequence receptor subunit 2 (Ssr2), mRNA | 1.236 | 1.443 |
| XM_028890269.1 | CD164 molecule (Cd164), transcript variant X1, mRNA | 1.227 | 0.405 |
| XM_028877920.1 | CD44 molecule (Indian blood group) (Cd44), transcript variant X1, mRNA | 1.223 | 0.764 |
| XM_028857418.1 | nuclear factor kappa B subunit 1 (Nfkb1), mRNA | 1.221 | 0.227 |
| XR_003733729.1 | uncharacterized LOC114687921 (LOC114687921), ncRNA | 1.220 | 0.360 |
| XM_028870723.1 | cathepsin D (Ctsd), mRNA | 1.212 | 1.455 |
| XM_028863284.1 | trem-like transcript 2 protein (LOC114688765), transcript variant X1, mRNA | 1.199 | -0.094 |
| XM_028856257.1 | pleckstrin homology domain containing O1 (Plekho1), mRNA | 1.199 | 0.987 |
| XM_028854772.1 | serine and arginine rich splicing factor 9 (Srsf9), mRNA | 1.199 | 0.849 |
| XM_028886473.1 | serine/threonine kinase 17b (Stk17b), transcript variant X1, mRNA | 1.198 | 0.191 |
| XM_028886842.1 | transmembrane protein 173 (Tmem173), mRNA | 1.191 | 0.484 |
| XM_028888108.1 | high mobility group nucleosomal binding domain 2 (Hmgn2), mRNA | 1.187 | 1.385 |
| XM_028855494.1 | CD63 molecule (Cd63), transcript variant X1, mRNA | 1.187 | 1.396 |
| XM_028869392.1 | phosphoinositide-3-kinase regulatory subunit 5 (Pik3r5), mRNA | 1.183 | 0.567 |
| XM_028870926.1 | glutamate ionotropic receptor NMDA type subunit associated protein 1 (Grina), mRNA | 1.176 | 0.925 |
| XM_028887957.1 | H-2 class I histocompatibility antigen, L-D alpha chain-like (LOC114705846) | 1.173 | 0.389 |
| XM_028866931.1 | ecotropic viral integration site 2B (Evi2b), mRNA | 1.168 | 0.281 |
| XM_028871183.1 | activating transcription factor 4 (Atf4), mRNA | 1.166 | 1.119 |
| XM_028885238.1 | granzyme A-like (LOC114704161), mRNA | 1.161 | 0.517 |
| XM_028864549.1 | ADP ribosylation factor guanine nucleotide exchange factor 1 (Arfgef1), mRNA | 1.158 | -0.040 |
| XM_028885499.1 | aldolase, fructose-bisphosphate A (Aldoa), transcript variant X1, mRNA | 1.148 | 0.998 |
| XM_028855439.1 | signal transducer and activator of transcription 2 (Stat2), mRNA | 1.143 | -0.380 |
| XM_028861345.1 | GEM-interacting protein-like (LOC114686693), mRNA | 1.136 | 0.503 |
| XM_028889955.1 | capping actin protein of muscle Z-line subunit alpha 1 (Capza1), mRNA | 1.136 | 0.513 |
| XM_028874912.1 | actin related protein 3 (Actr3), transcript variant X2, mRNA | 1.132 | 0.610 |
| XM_028887383.1 | actin, cytoplasmic 1 (LOC114705499), transcript variant X1, mRNA | 1.128 | 0.517 |
| XM_028892332.1 | triosephosphate isomerase 1 (Tpi1), mRNA | 1.119 | 1.789 |
| XM_028881507.1 | C-type lectin domain family 2 member D11-like (LOC114701420), mRNA | 1.111 | 0.099 |
| XM_028863331.1 | leukocyte immunoglobulin-like receptor subfamily A member 6 (LOC114688825), mRNA | 1.108 | 1.790 |
| XM_028877113.1 | interferon alpha and beta receptor subunit 2 (Ifnar2), mRNA | 1.103 | 0.720 |
| XM_028871240.1 | sterol regulatory element binding transcription factor 2 (Srebf2), transcript variant X1, mRNA | 1.102 | 0.647 |
| XM_028861428.1 | G protein subunit beta 2 (Gnb2), mRNA | 1.100 | 1.077 |
| XM_028854475.1 | cyclase associated actin cytoskeleton regulatory protein 1 (Cap1), transcript variant X1, mRNA | 1.099 | -0.309 |
| XM_028875931.1 | ATPase H+ transporting accessory protein 2 (Atp6ap2), mRNA | 1.077 | 1.406 |
| XM_028872176.1 | Rho GDP dissociation inhibitor beta (Arhgdib), transcript variant X1, mRNA | 1.077 | 0.389 |
| XM_028881876.1 | platelet factor 4-like (LOC114701718), mRNA | 1.062 | 1.462 |
| XM_028888609.1 | filamin A (Flna), transcript variant X2, mRNA | 1.059 | 0.808 |
| XM_028885543.1 | major vault protein (Mvp), mRNA | 1.057 | 0.887 |
| XM_028888444.1 | mitogen-activated protein kinase 14 (Mapk14), transcript variant X1, mRNA | 1.053 | 0.351 |
| XM_028880649.1 | RHO family interacting cell polarization regulator 2 (Ripor2), transcript v | 1.051 | -0.310 |
| XM_028859977.1 | glycogen synthase kinase 3 alpha (Gsk3a), mRNA | 1.049 | 0.143 |
| XM_028882770.1 | SEC61 translocon beta subunit (Sec61b), mRNA | 1.049 | 1.146 |
| XM_028856275.1 | extracellular matrix protein 1 (Ecm1), transcript variant X2, mRNA | 1.045 | 0.393 |
| XM_028884739.1 | cyclin dependent kinase 9 (Cdk9), mRNA | 1.043 | 0.648 |
| XR_003733081.1 | leucine-rich repeat-containing protein 74B-like (LOC114682722), transcript | 1.042 | 0.500 |
| XM_028855798.1 | guanine nucleotide-binding protein G(s) subunit alpha isoforms short (LOC114682029), transcript variant X7, mRNA | 1.040 | -0.018 |
| XM_028895088.1 | family with sequence similarity 107 member B (Fam107b), transcript variant | 1.039 | 0.112 |
| XM_028869784.1 | sterile alpha motif domain containing 9 (Samd9), transcript variant X1, mRNA | 1.039 | -0.391 |
| XM_028876455.1 | zyxin (Zyx), transcript variant X5, mRNA | 1.038 | 0.579 |
| XM_028874911.1 | actin related protein 3 (Actr3), transcript variant X1, mRNA | 1.036 | 0.183 |
| XM_028861714.1 | protein phosphatase 1 regulatory subunit 18 (Ppp1r18), transcript variant X1, mRNA | 1.033 | 0.080 |
| XM_028862158.1 | arrestin beta 2 (Arrb2), mRNA | 1.033 | 0.293 |
| XM_028889264.1 | high mobility group box 2 (Hmgb2), mRNA | 1.031 | 0.377 |
| XM_028892656.1 | ATPase H+ transporting V1 subunit G1 (Atp6v1g1), mRNA | 1.024 | 0.916 |
| XM_028859175.1 | glia maturation factor gamma (Gmfg), transcript variant X2, mRNA | 1.024 | 0.497 |
| XM_028862612.1 | F-box protein 9 (Fbxo9), mRNA | 1.019 | 0.428 |
| XM_028864383.1 | mitochondrial pyruvate carrier 2 (Mpc2), mRNA | 1.014 | 1.011 |
| XM_028888273.1 | interferon alpha-inducible protein 27-like protein 2A (LOC114706052), mRNA | 1.009 | 0.907 |
| XM_028856814.1 | RAB5C, member RAS oncogene family (Rab5c), mRNA | 1.007 | 0.337 |
| XM_028891462.1 | ISG15 ubiquitin like modifier (Isg15), mRNA | 1.006 | 0.786 |
| XM_028865090.1 | secretory carrier membrane protein 2 (Scamp2), transcript variant X2, mRNA | 0.999 | 0.647 |
| XM_028874016.1 | mitochondrial ribosomal protein L33 (Mrpl33), mRNA | 0.996 | 1.756 |
| XM_028854262.1 | ATPase H+ transporting V0 subunit d1 (Atp6v0d1), mRNA | 0.995 | 1.353 |
| XM_028880639.1 | WAS/WASL interacting protein family member 1 (Wipf1), transcript variant X4 | 0.992 | -0.182 |
| XM_028870213.1 | transaldolase 1 (Taldo1), mRNA | 0.991 | 0.702 |
| XM_028888864.1 | moesin (Msn), mRNA | 0.985 | 0.406 |
| XM_028863799.1 | semaphorin 4D (Sema4d), transcript variant X1, mRNA | 0.985 | 0.146 |
| XM_028880288.1 | heat shock protein 90 beta family member 1 (Hsp90b1), mRNA | 0.983 | 1.018 |
| XM_028894085.1 | p21 (RAC1) activated kinase 2 (Pak2), transcript variant X2, mRNA | 0.982 | -0.003 |
| XM_028893196.1 | ATPase H+ transporting V0 subunit b (Atp6v0b), mRNA | 0.970 | 1.448 |
| XM_028888539.1 | ATPase H+ transporting accessory protein 1 (Atp6ap1), mRNA | 0.964 | 0.714 |
| XM_028890280.1 | Rac family small GTPase 1 (Rac1), mRNA | 0.962 | 0.771 |
| XM_028876500.1 | Ras association domain family member 5 (Rassf5), mRNA | 0.961 | 0.070 |
| XM_028855901.1 | calpain 1 (Capn1), mRNA | 0.957 | 0.341 |
| XM_028872846.1 | Kruppel like factor 13 (Klf13), mRNA | 0.956 | -0.175 |
| XM_028886979.1 | ras homolog family member A (Rhoa), transcript variant X1, mRNA | 0.955 | 0.136 |
| XM_028874399.1 | NCK associated protein 1 like (Nckap1l), mRNA | 0.950 | 0.572 |
| XM_028890637.1 | arginyl aminopeptidase like 1 (Rnpepl1), transcript variant X1, mRNA | 0.950 | 0.043 |
| XM_028877815.1 | FXYD domain containing ion transport regulator 5 (Fxyd5), transcript variant X4, mRNA | 0.948 | 0.418 |
| XM_028869503.1 | GABA type A receptor-associated protein (Gabarap), mRNA | 0.942 | 0.671 |
| XM_028854476.1 | cyclase associated actin cytoskeleton regulatory protein 1 (Cap1), transcript variant X2, mRNA | 0.940 | 1.618 |
| XM_028895001.1 | hematopoietic cell-specific Lyn substrate 1 (Hcls1), mRNA | 0.939 | 0.759 |
| XM_028886666.1 | signal transducer and activator of transcription 1 (Stat1), mRNA | 0.936 | -0.307 |
| XM_028859787.1 | resistin-like gamma (LOC114685171), mRNA | 0.931 | 0.830 |
| XM_028879886.1 | exportin 6 (Xpo6), mRNA | 0.930 | -0.104 |
| XM_028854140.1 | RAB7A, member RAS oncogene family (Rab7a), mRNA | 0.930 | 0.426 |
| XM_028884538.1 | actin related protein 2 (Actr2), transcript variant X2, mRNA | 0.923 | 0.007 |
| XM_028883945.1 | talin 1 (Tln1), mRNA | 0.920 | 0.480 |
| XM_028857400.1 | ornithine decarboxylase antizyme 3 (Oaz3), mRNA | 0.918 | 0.210 |
| XM_028868268.1 | transmembrane 9 superfamily member 2 (Tm9sf2), mRNA | 0.916 | 0.736 |
| XM_028876003.1 | N-myc downstream regulated 1 (Ndrg1), mRNA | 0.911 | 0.419 |
| XM_028880299.1 | histone H2B type 1 (LOC114700621), mRNA | 0.906 | 0.195 |
| XM_028891487.1 | G protein subunit beta 1 (Gnb1), transcript variant X1, mRNA | 0.905 | 0.558 |
| XM_028893125.1 | angio associated migratory cell protein (Aamp), mRNA | 0.903 | 0.414 |
| XR_003736138.1 | uncharacterized LOC114703406 (LOC114703406), ncRNA | 0.891 | -0.379 |
| XM_028873771.1 | IQ motif containing GTPase activating protein 1 (Iqgap1), mRNA | 0.890 | 0.350 |
| XM_028860501.1 | Y-box binding protein 3 (Ybx3), transcript variant X2, mRNA | 0.888 | -0.211 |
| XM_028878280.1 | clathrin heavy chain (Cltc), transcript variant X4, mRNA | 0.886 | 0.862 |
| XM_028855656.1 | aldehyde dehydrogenase 2 family member (Aldh2), mRNA | 0.886 | -0.490 |
| XM_028889599.1 | MSL complex subunit 1 (Msl1), transcript variant X1, mRNA | 0.884 | -0.147 |
| XM_028856894.1 | 2',3'-cyclic nucleotide 3' phosphodiesterase (Cnp), mRNA | 0.884 | -0.263 |
| XM_028877179.1 | dynactin subunit 6 (Dctn6), mRNA | 0.884 | 0.466 |
| XM_028888389.1 | ral guanine nucleotide dissociation stimulator like 2 (Rgl2), transcript va | 0.880 | 0.654 |
| XM_028881480.1 | C-type lectin domain family 1 member B (Clec1b), transcript variant X1, mRN | 0.880 | 0.400 |
| XM_028886198.1 | Rho GTPase activating protein 9 (Arhgap9), transcript variant X2, mRNA | 0.878 | 0.815 |
| XM_028893352.1 | ATPase H+ transporting V0 subunit e1 (Atp6v0e1), mRNA | 0.876 | 0.286 |
| XM_028860338.1 | ferritin heavy chain (LOC114685692), mRNA | 0.874 | -0.611 |
| XM_028884536.1 | RAB1A, member RAS oncogene family (Rab1a), mRNA | 0.873 | 0.896 |
| XM_028887418.1 | pleckstrin (Plek), mRNA | 0.868 | 0.725 |
| XM_028890007.1 | clusterin (Clu), transcript variant X2, mRNA | 0.864 | 0.220 |
| XM_028862608.1 | paired immunoglobulin-like type 2 receptor beta (LOC114687976), mRNA | 0.863 | 1.706 |
| XM_028890324.1 | actin related protein 2/3 complex subunit 1B (Arpc1b), transcript variant X | 0.861 | 0.952 |
| XM_028887541.1 | histocompatibility minor 13 (Hm13), transcript variant X2, mRNA | 0.859 | 0.815 |
| XM_028893040.1 | actin related protein 2/3 complex subunit 2 (Arpc2), transcript variant X2, | 0.858 | 0.502 |
| XM_028883204.1 | ficolin-2 (LOC114702718), mRNA | 0.858 | 3.029 |
| XM_028892879.1 | RAB1B, member RAS oncogene family (Rab1b), mRNA | 0.857 | 0.778 |
| XM_028895302.1 | G protein subunit alpha i2 (Gnai2), mRNA | 0.854 | 0.563 |
| XM_028854486.1 | akirin 1 (Akirin1), mRNA | 0.853 | 0.224 |
| XM_028860317.1 | ezrin (Ezr), mRNA | 0.850 | 0.081 |
| XM_028887958.1 | H-2 class I histocompatibility antigen, L-D alpha chain-like (LOC114705846) | 0.848 | 0.532 |
| XM_028858785.1 | ADP ribosylation factor 6 (Arf6), mRNA | 0.847 | 0.579 |
| XM_028882169.1 | stress associated endoplasmic reticulum protein 1 (Serp1), mRNA | 0.847 | 0.719 |

Table S18

| | | | |
|---|---|---|---|
| XR_003737047.1 | uncharacterized LOC114710366 (LOC114710366), transcript variant X2, | -0.177 | -0.190 |
| XM_028871187.1 | GRB2 related adaptor protein 2 (Grap2), transcript variant X1, mRNA | -0.181 | 0.464 |
| XM_028888954.1 | ring finger protein 11 (Rnf11), mRNA | -0.182 | 0.170 |
| XM_028879784.1 | voltage dependent anion channel 2 (Vdac2), mRNA | -0.184 | 0.402 |
| XM_028861490.1 | Rho guanine nucleotide exchange factor 1 (Arhgef1), transcript variant | -0.187 | -0.737 |
| XM_028865899.1 | DEAH-box helicase 15 (Dhx15), mRNA | -0.188 | -0.381 |
| XM_028884960.1 | cytochrome b-245 chaperone 1 (LOC114703918), mRNA | -0.188 | 0.024 |
| XM_028860826.1 | ABRA C-terminal like (Abracl), transcript variant X2, mRNA | -0.189 | 0.975 |
| XM_028887649.1 | eukaryotic translation initiation factor 3 subunit I (Eif3i), mRNA | -0.189 | 0.142 |
| XM_028870242.1 | Janus kinase 1 (Jak1), mRNA | -0.190 | -0.472 |
| XM_028894283.1 | transmembrane protein 154 (Tmem154), transcript variant X2, mRNA | -0.192 | -0.056 |
| XM_028878565.1 | VPS16 core subunit of CORVET and HOPS complexes (Vps16), mRNA | -0.193 | 0.036 |
| XM_028879896.1 | ataxin-2-like protein (LOC114700310), mRNA | -0.195 | -0.497 |
| XM_028854261.1 | RAN binding protein 10 (Ranbp10), mRNA | -0.196 | -0.133 |
| XR_003732817.1 | serine/arginine repetitive matrix 2 (Srrm2), transcript variant X4, misc_ | -0.199 | -0.462 |
| XM_028886978.1 | glutathione peroxidase 1 (Gpx1), mRNA | -0.200 | 0.668 |
| XM_028856186.1 | small nuclear ribonucleoprotein Sm D2 (LOC114682345), mRNA | -0.202 | 0.487 |
| XM_028889307.1 | makorin ring finger protein 1 (Mkrn1), mRNA | -0.203 | 0.439 |
| XM_028885960.1 | peptidyl-prolyl cis-trans isomerase A-like (LOC114704621), mRNA | -0.204 | 1.530 |
| XM_028892192.1 | defender against cell death 1 (Dad1), mRNA | -0.205 | 0.458 |
| XM_028869464.1 | RNA polymerase II subunit A (Polr2a), mRNA | -0.207 | -0.545 |
| XM_028876149.1 | serine and arginine rich splicing factor 5 (Srsf5), transcript variant X6, m | -0.209 | -0.268 |
| XM_028886482.1 | serine/threonine kinase 17b (Stk17b), transcript variant X2, mRNA | -0.211 | -0.206 |
| XM_028893580.1 | ribosomal protein S24 (Rps24), transcript variant X5, mRNA | -0.215 | 0.071 |
| XM_028889721.1 | peroxiredoxin 1 (Prdx1), mRNA | -0.216 | 0.910 |
| XM_028881343.1 | proteolipid protein 2 (Plp2), mRNA | -0.217 | 0.523 |
| XM_028882787.1 | acidic nuclear phosphoprotein 32 family member B (Anp32b), mRNA | -0.219 | 0.228 |
| XM_028870807.1 | adducin 1 (Add1), transcript variant X4, mRNA | -0.219 | -0.216 |
| XM_028859224.1 | heterogeneous nuclear ribonucleoprotein L (Hnrnpl), transcript variant X | -0.220 | -0.044 |
| XM_028862828.1 | protein kinase N1 (Pkn1), transcript variant X2, mRNA | -0.222 | -0.002 |
| XM_028862553.1 | H-2 class I histocompatibility antigen, Q10 alpha chain-like (LOC114687 | -0.229 | -0.501 |
| XM_028869002.1 | prohibitin (Phb), mRNA | -0.231 | 0.058 |
| XM_028881940.1 | cyclin I (Ccni), transcript variant X1, mRNA | -0.231 | -0.444 |
| XM_028884447.1 | zinc finger CCCH-type containing 15 (Zc3h15), mRNA | -0.234 | 0.520 |
| XM_028885151.1 | carbonic anhydrase 2 (Ca2), mRNA | -0.238 | 0.427 |
| XM_028868545.1 | phosphatidylinositol-3,4,5-trisphosphate dependent Rac exchange facto | -0.239 | -0.012 |
| XM_028871981.1 | ATP synthase F1 subunit alpha (Atp5f1a), mRNA | -0.239 | 0.155 |
| XM_028876604.1 | BTG anti-proliferation factor 2 (Btg2), mRNA | -0.242 | -0.368 |
| XM_028892476.1 | cytochrome c oxidase subunit 7A-related protein, mitochondrial (LOC11 | -0.244 | 0.301 |
| XM_028870317.1 | ribosomal protein lateral stalk subunit P2 (Rplp2), mRNA | -0.245 | 0.255 |
| XM_028869568.1 | arachidonate 15-lipoxygenase (LOC114693249), mRNA | -0.248 | 0.082 |
| XM_028887625.1 | erythrocyte membrane protein band 4.1 (Epb41), transcript variant X12, | -0.249 | 0.849 |
| XM_028854968.1 | peroxiredoxin like 2A (Prxl2a), transcript variant X2, mRNA | -0.251 | 0.343 |
| XM_028889997.1 | BCL2/adenovirus E1B 19 kDa protein-interacting protein 3-like (LOC114 | -0.252 | -0.155 |
| XM_028871706.1 | basic transcription factor 3 (Btf3), transcript variant X2, mRNA | -0.255 | 0.518 |
| XM_028885440.1 | thyroid hormone receptor associated protein 3 (Thrap3), transcript varia | -0.256 | -0.436 |
| XM_028889679.1 | Jupiter microtubule associated homolog 1 (Jpt1), mRNA | -0.259 | 0.452 |
| XM_028885690.1 | sialophorin (Spn), mRNA | -0.264 | 0.204 |
| XM_028880854.1 | adenine phosphoribosyltransferase (Aprt), mRNA | -0.269 | 0.954 |
| XM_028891536.1 | ATP synthase F1 subunit gamma (Atp5f1c), transcript variant X1, mRNA | -0.271 | 0.528 |
| XM_028854299.1 | proteasome subunit beta 10 (Psmb10), mRNA | -0.279 | 0.198 |
| XM_028878340.1 | glutamate dehydrogenase 1 (Glud1), mRNA | -0.283 | 0.065 |
| XM_028872451.1 | solute carrier family 44 member 2 (Slc44a2), mRNA | -0.284 | 0.127 |
| XM_028861517.1 | 5'-nucleotidase, cytosolic IIIA (Nt5c3a), transcript variant X1, mRNA | -0.285 | 1.375 |
| XM_028889380.1 | cytochrome b-c1 complex subunit 6, mitochondrial (LOC114706860), mR | -0.292 | 0.271 |
| XM_028866322.1 | Thy-1 cell surface antigen (Thy1), transcript variant X1, mRNA | -0.297 | -0.335 |
| XM_028857792.1 | serine/threonine kinase 10 (Stk10), mRNA | -0.299 | -0.289 |
| XM_028880224.1 | solute carrier family 25 member 3 (Slc25a3), transcript variant X2, mRN | -0.302 | 0.362 |
| XR_003735117.1 | eukaryotic translation initiation factor 4A2 (Eif4a2), transcript variant X | -0.305 | -0.966 |
| XR_003735131.1 | arginine and serine rich protein 1 (Rsrp1), transcript variant X2, misc_RI | -0.306 | -0.404 |
| XM_028884758.1 | ribosomal protein L12 (Rpl12), mRNA | -0.306 | -0.169 |
| XM_028893647.1 | ribosomal protein S12 (Rps12), mRNA | -0.308 | 0.115 |
| XM_028875254.1 | Rh blood group CcEe antigens (Rhce), mRNA | -0.308 | 0.082 |
| XM_028861430.1 | leucine rich repeats and calponin homology domain containing 4 (Lrch4) | -0.308 | 0.181 |
| XM_028892347.1 | TAP binding protein like (Tapbpl), mRNA | -0.308 | -0.337 |
| XM_028887371.1 | SAM and SH3 domain containing 3 (Sash3), transcript variant X2, mRNA | -0.309 | -0.451 |
| XM_028878063.1 | high mobility group box 1 (Hmgb1), transcript variant X1, mRNA | -0.311 | -0.260 |
| XM_028863431.1 | IFI30 lysosomal thiol reductase (Ifi30), mRNA | -0.312 | 1.997 |
| XM_028877701.1 | tyrosine 3-monooxygenase/tryptophan 5-monooxygenase activation pro | -0.316 | -0.305 |
| XM_028866750.1 | pre-mRNA processing factor 8 (Prpf8), mRNA | -0.318 | -0.523 |
| XM_028856593.1 | ferrochelatase (Fech), mRNA | -0.321 | 0.380 |
| XM_028861593.1 | nascent polypeptide associated complex subunit alpha (Naca), transcrip | -0.321 | -0.622 |
| XM_028876229.1 | nucleolin (Ncl), mRNA | -0.324 | -0.222 |
| XM_028880021.1 | histidine triad nucleotide binding protein 1 (Hint1), mRNA | -0.326 | 0.240 |
| XM_028882342.1 | MORF4 family-associated protein 1 (LOC114702067), mRNA | -0.329 | 0.304 |
| XM_028870456.1 | phosphatidylinositol-5-phosphate 4-kinase type 2 alpha (Pip4k2a), trans | -0.330 | -0.377 |
| XM_028888778.1 | alpha-1,3-mannosyl-glycoprotein 2-beta-N-acetylglucosaminyltransfera | -0.331 | -0.225 |
| XM_028882595.1 | solute carrier family 4 member 1 (Diego blood group) (Slc4a1), mRNA | -0.332 | 0.116 |
| XM_028861659.1 | 60S ribosomal protein L38 (LOC114686962), transcript variant X1, mRN | -0.332 | 0.252 |
| XM_028860759.1 | eukaryotic translation initiation factor 3 subunit A (Eif3a), mRNA | -0.334 | -0.339 |
| XM_028871117.1 | histone H1.0 (LOC114694248), mRNA | -0.336 | 0.096 |
| XM_028859291.1 | eukaryotic translation elongation factor 1 delta (Eef1d), transcript varia | -0.340 | -0.162 |
| XM_028861945.1 | S100 calcium binding protein A10 (S100a10), mRNA | -0.340 | 1.148 |
| XM_028871690.1 | NSA2 ribosome biogenesis factor (Nsa2), mRNA | -0.341 | 0.113 |
| XM_028862307.1 | protein phosphatase 6 regulatory subunit 1 (Ppp6r1), transcript variant | -0.342 | 0.247 |
| XR_003736604.1 | serine and arginine rich splicing factor 2 (Srsf2), transcript variant X1, | -0.343 | -0.727 |
| XM_028854063.1 | RAB43, member RAS oncogene family (Rab43), mRNA | -0.343 | 0.506 |
| XM_028882214.1 | ras homolog family member H (Rhoh), mRNA | -0.344 | -0.762 |
| XM_028881391.1 | RNA binding motif protein 3 (Rbm3), transcript variant X1, mRNA | -0.346 | 0.725 |
| XM_028858537.1 | polypyrimidine tract binding protein 1 (Ptbp1), mRNA | -0.351 | 0.121 |
| XM_028861257.1 | RNA transcription, translation and transport factor (Rtraf), mRNA | -0.353 | 0.317 |
| XM_028877936.1 | chaperonin containing TCP1 subunit 2 (Cct2), mRNA | -0.357 | 0.100 |
| XR_003735035.1 | heterogeneous nuclear ribonucleoprotein A1 (Hnrnpa1), transcript varia | -0.357 | -0.221 |
| XM_028871108.1 | galectin 1 (Lgals1), mRNA | -0.358 | 1.262 |
| XM_028893248.1 | sorting nexin 5 (Snx5), mRNA | -0.359 | 0.200 |
| XM_028874427.1 | ATP synthase membrane subunit c locus 2 (Atp5mc2), mRNA | -0.360 | 0.191 |
| XM_028854967.1 | peroxiredoxin like 2A (Prxl2a), transcript variant X1, mRNA | -0.362 | 0.575 |
| XM_028860872.1 | proteasome activator subunit 4 (Psme4), mRNA | -0.364 | 0.917 |
| XM_028854762.1 | dynein light chain LC8-type 1 (Dynll1), mRNA | -0.368 | 0.446 |
| XM_028855444.1 | extended synaptotagmin 1 (Esyt1), mRNA | -0.369 | 0.153 |
| XM_028878680.1 | cyclin D1 binding protein 1 (Ccndbp1), transcript variant X1, mRNA | -0.372 | 1.076 |
| XM_028859139.1 | fibrillarin (Fbl), mRNA | -0.372 | 0.620 |
| XM_028878038.1 | RNA polymerase I and III subunit D (Polr1d), mRNA | -0.374 | 0.329 |
| XM_028887534.1 | BCL2 like 1 (Bcl2l1), transcript variant X3, mRNA | -0.375 | 0.112 |
| XM_028883119.1 | vesicle associated membrane protein 3 (Vamp3), mRNA | -0.375 | -0.077 |
| XM_028872425.1 | eukaryotic translation initiation factor 3 subunit G (Eif3g), mRNA | -0.378 | -0.010 |
| XM_028876127.1 | ZFP36 ring finger protein like 1 (Zfp36l1), mRNA | -0.380 | -0.815 |
| XM_028889956.1 | DENN domain containing 2D (Dennd2d), transcript variant X1, mRNA | -0.381 | -0.357 |
| XM_028878001.1 | PNN interacting serine and arginine rich protein (Pnisr), transcript varia | -0.384 | -0.711 |
| XM_028861518.1 | 5'-nucleotidase, cytosolic IIIA (Nt5c3a), transcript variant X2, mRNA | -0.386 | 0.512 |
| XM_028858263.1 | EP300 interacting inhibitor of differentiation 1 (Eid1), mRNA | -0.391 | 0.461 |
| XM_028871019.1 | ribosomal protein L8 (Rpl8), mRNA | -0.391 | -0.202 |
| XM_028874407.1 | heterogeneous nuclear ribonucleoprotein A1 (Hnrnpa1), transcript varia | -0.393 | -0.171 |
| XM_028886197.1 | Rho GTPase activating protein 9 (Arhgap9), transcript variant X1, mRNA | -0.394 | -0.798 |
| XM_028885498.1 | yippee like 3 (Ypel3), transcript variant X2, mRNA | -0.395 | 0.322 |
| XM_028880807.1 | interferon regulatory factor 8 (Irf8), mRNA | -0.397 | 0.186 |
| XM_028858267.1 | deoxyuridine triphosphatase (Dut), mRNA | -0.399 | -0.356 |
| XM_028883979.1 | family with sequence similarity 214 member B (Fam214b), transcript va | -0.403 | -0.591 |
| XM_028865496.1 | heterogeneous nuclear ribonucleoprotein U (Hnrnpu), mRNA | -0.403 | -0.160 |
| XM_028865104.1 | cytochrome c oxidase subunit 5A, mitochondrial (LOC114690344), mRN | -0.407 | 0.265 |
| XM_028871120.1 | eukaryotic translation initiation factor 3 subunit L (Eif3l), transcript vari | -0.407 | -0.353 |
| XM_028885840.1 | myosin light polypeptide 6-like (LOC114704518), mRNA | -0.409 | 1.302 |
| XM_028872191.1 | microsomal glutathione S-transferase 1 (Mgst1), mRNA | -0.409 | 0.808 |
| XM_028861402.1 | leukocyte receptor cluster member 8 (Leng8), transcript variant X4, mR | -0.409 | -0.274 |
| XM_028869557.1 | trafficking protein particle complex 1 (Trappc1), mRNA | -0.409 | 0.540 |
| XM_028892376.1 | chromodomain helicase DNA binding protein 4 (Chd4), mRNA | -0.412 | -0.444 |
| XM_028877622.1 | exportin 7 (Xpo7), transcript variant X2, mRNA | -0.412 | 0.440 |
| XM_028861399.1 | leukocyte receptor cluster member 8 (Leng8), transcript variant X1, mR | -0.414 | -2.421 |
| XM_028866946.1 | cytokine receptor like factor 3 (Crlf3), mRNA | -0.415 | -0.567 |
| XM_028878554.1 | isocitrate dehydrogenase (NAD(+)) 3 beta (Idh3b), transcript variant X2, | -0.424 | -0.079 |
| XM_028882970.1 | IKAROS family zinc finger 1 (Ikzf1), transcript variant X2, mRNA | -0.427 | -0.889 |
| XM_028859565.1 | eukaryotic translation initiation factor 3 subunit K (Eif3k), mRNA | -0.429 | 0.119 |
| XM_028870559.1 | peptidylprolyl isomerase A (Ppia), mRNA | -0.434 | 0.516 |
| XM_028884603.1 | ubiquitin conjugating enzyme E2 E3 (Ube2e3), transcript variant X3, mR | -0.434 | 1.254 |
| XM_028874477.1 | prefoldin subunit 5 (Pfdn5), mRNA | -0.434 | 0.142 |
| XM_028884489.1 | poly(A) binding protein cytoplasmic 1 (Pabpc1), mRNA | -0.434 | 0.009 |
| XM_028859125.1 | small nuclear ribonucleoprotein U1 subunit 70 (Snrnp70), transcript vari | -0.435 | -0.636 |
| XM_028857232.1 | transcription factor 7 (Tcf7), transcript variant X2, mRNA | -0.436 | -1.352 |
| XM_028860320.1 | T cell activation RhoGTPase activating protein (Tagap), transcript varian | -0.441 | -1.086 |
| XM_028892654.1 | AT-hook transcription factor (Akna), transcript variant X3, mRNA | -0.442 | -0.488 |
| XR_003733413.1 | uncharacterized LOC114684882 (LOC114684882), ncRNA | -0.442 | -1.051 |

| | | | | | | | | |
|---|---|---|---|---|---|---|---|---|
| XM_028881875.1 | platelet basic protein-like (LOC114701717), mRNA | 0.845 | 0.017 | Table S18 | XM_028889611.1 | ubiquitin conjugating enzyme E2 O (Ube2o), mRNA | -0.443 | 0.650 |
| XM_028887916.1 | G protein signaling modulator 3 (Gpsm3), transcript variant X1, mRNA | 0.843 | 0.367 | | XM_028853968.1 | ribosomal protein S2 (Rps2), mRNA | -0.445 | 0.036 |
| XM_028875281.1 | capping actin protein of muscle Z-line subunit beta (Capzb), transcript variant X2, mRNA | 0.843 | 0.620 | | XM_028892043.1 | non-POU domain containing octamer binding (Nono), mRNA | -0.445 | -0.272 |
| XM_028865632.1 | C-reactive protein (Crp), mRNA | 0.842 | 0.015 | | XM_028887892.1 | DExD-box helicase 39B (Ddx39b), mRNA | -0.446 | -0.092 |
| XM_028855872.1 | EH domain binding protein 1 like 1 (Ehbp1l1), transcript variant X2, mRNA | 0.837 | 0.718 | | XM_028887304.1 | solute carrier family 25 member 5 (Slc25a5), mRNA | -0.448 | 0.402 |
| XM_028862602.1 | paired immunoglobulin-like type 2 receptor alpha (LOC114687967), mRNA | 0.835 | 2.042 | | XM_028875584.1 | ribosomal protein L37 (Rpl37), mRNA | -0.451 | -0.097 |
| XM_028868628.1 | WAP four-disulfide core domain 2 (Wfdc2), mRNA | 0.833 | -0.291 | | XM_028855889.1 | FAU ubiquitin like and ribosomal protein S30 fusion (Fau), mRNA | -0.451 | 0.103 |
| XM_028895090.1 | family with sequence similarity 107 member B (Fam107b), transcript variant | 0.832 | -0.508 | | XM_028857796.1 | nucleophosmin 1 (Npm1), mRNA | -0.454 | 0.461 |
| XM_028857494.1 | mitogen-activated protein kinase kinase 3 (Map2k3), mRNA | 0.831 | 0.050 | | XM_028855210.1 | serine and arginine rich splicing factor 7 (Srsf7), transcript variant X1, m | -0.458 | -0.237 |
| XM_028884810.1 | ubiquitin like modifier activating enzyme 1 (Uba1), transcript variant X1, | 0.828 | -0.094 | | XM_028880035.1 | GM2 ganglioside activator (Gm2a), mRNA | -0.461 | 0.605 |
| XM_028875380.1 | cell division cycle 42 (Cdc42), transcript variant X1, mRNA | 0.827 | 1.300 | | XM_028883453.1 | apurinic/apyrimidinic endodeoxyribonuclease 2 (Apex2), transcript varia | -0.466 | 0.401 |
| XM_028876351.1 | UDP-GlcNAc:betaGal beta-1,3-N-acetylglucosaminyltransferase 2 (B3gnt2), mRNA | 0.824 | -0.526 | | XM_028863040.1 | ribosomal protein S9 (Rps9), mRNA | -0.468 | 0.004 |
| XR_003734371.1 | uncharacterized LOC114692242 (LOC114692242), ncRNA | 0.821 | 0.044 | | XM_028871808.1 | glutathione S-transferase P (LOC114694764), mRNA | -0.470 | 0.177 |
| XM_028874611.1 | transmembrane BAX inhibitor motif containing 6 (Tmbim6), mRNA | 0.820 | 0.189 | | XM_028863637.1 | RAB24, member RAS oncogene family (Rab24), transcript variant X1, m | -0.472 | 0.045 |
| XM_028860073.1 | Rho GTPase activating protein 30 (Arhgap30), mRNA | 0.818 | 0.396 | | XM_028854883.1 | HOP homeobox (Hopx), transcript variant X1, mRNA | -0.472 | 1.006 |
| XM_028862291.1 | eukaryotic translation initiation factor 1 (Eif1), mRNA | 0.818 | 0.689 | | XM_028857460.1 | acid phosphatase 1 (Acp1), mRNA | -0.473 | 0.773 |
| XM_028861660.1 | 60S ribosomal protein L38 (LOC114686962), transcript variant X2, mRNA | 0.816 | 0.433 | | XM_028865082.1 | C-terminal Src kinase (Csk), transcript variant X2, mRNA | -0.473 | -0.055 |
| XM_028859979.1 | death effector domain containing 2 (Dedd2), mRNA | 0.813 | 0.298 | | XM_028883123.1 | Parkinsonism associated deglycase (Park7), mRNA | -0.482 | 0.566 |
| XM_028865035.1 | proline-serine-threonine phosphatase interacting protein 1 (Pstpip1), transcript variant X1, mRNA | 0.802 | 0.066 | | XM_028861733.1 | H-2 class I histocompatibility antigen, Q10 alpha chain-like (LOC114687 | -0.483 | -0.133 |
| XM_028855572.1 | RAB5 interacting factor (Rab5if), mRNA | 0.796 | 0.836 | | XM_028895120.1 | TSC22 domain family member 3 (Tsc22d3), transcript variant X3, mRNA | -0.485 | -0.916 |
| XM_028890028.1 | protein tyrosine kinase 2 beta (Ptk2b), transcript variant X2, mRNA | 0.795 | 0.628 | | XM_028858892.1 | ribosomal protein L36a like (Rpl36al), mRNA | -0.490 | 0.147 |
| XM_028868574.1 | ring finger protein 114 (Rnf114), mRNA | 0.791 | 0.002 | | XM_028858596.1 | Rho GTPase activating protein 45 (Arhgap45), transcript variant X1, mR | -0.493 | -0.368 |
| XM_028888530.1 | podocan (Podn), transcript variant X2, mRNA | 0.789 | -1.635 | | XM_028882116.1 | Enah/Vasp-like (Evl), transcript variant X4, mRNA | -0.496 | -0.930 |
| XM_028888178.1 | zinc finger DHHC-type containing 18 (Zdhhc18), mRNA | 0.787 | 0.186 | | XM_028893941.1 | diazepam binding inhibitor, acyl-CoA binding protein (Dbi), mRNA | -0.501 | 1.579 |
| XM_028875036.1 | heterogeneous nuclear ribonucleoprotein H2 (Hnrnph2), mRNA | 0.786 | 0.554 | | XM_028889678.1 | small ubiquitin like modifier 2 (Sumo2), mRNA | -0.502 | 0.030 |
| XM_028861626.1 | heat shock cognate 71 kDa protein-like (LOC114686934), mRNA | 0.782 | 0.657 | | XM_028857620.1 | heparin binding growth factor (Hdgf), mRNA | -0.502 | 0.298 |
| XM_028857136.1 | ADP ribosylation factor 1 (Arf1), transcript variant X1, mRNA | 0.779 | 0.195 | | XM_028868351.1 | sphingomyelin phosphodiesterase acid like 3A (Smpdl3a), mRNA | -0.516 | 0.162 |
| XM_028853633.1 | anaphase promoting complex subunit 11 (Anapc11), transcript variant X2, mRNA | 0.779 | 0.478 | | XM_028888341.1 | ribosomal protein S10 (Rps10), mRNA | -0.517 | -0.206 |
| XM_028880236.1 | leukotriene A4 hydrolase (Lta4h), transcript variant X1, mRNA | 0.770 | 1.078 | | XM_028858655.1 | interleukin 27 receptor subunit alpha (Il27ra), mRNA | -0.524 | -0.502 |
| XM_028858998.1 | lamin B receptor (Lbr), mRNA | 0.761 | 0.247 | | XM_028880241.1 | small nuclear ribonucleoprotein polypeptide F (Snrpf), mRNA | -0.526 | 1.962 |
| XM_028855749.1 | glycoprotein IX platelet (Gp9), mRNA | 0.759 | -0.150 | | XM_028894461.1 | cyclin dependent kinase inhibitor 1B (Cdkn1b), transcript variant X2, mR | -0.531 | -0.563 |
| XM_028862148.1 | complement component receptor 1-like protein (LOC114687533), transcript variant X2, mRNA | 0.756 | 0.377 | | XM_028867497.1 | ribosomal protein L24 (Rpl24), mRNA | -0.536 | 0.206 |
| XM_028853582.1 | prolyl 4-hydroxylase subunit beta (P4hb), mRNA | 0.754 | 1.292 | | XM_028857231.1 | transcription factor 7 (Tcf7), transcript variant X1, mRNA | -0.552 | -1.095 |
| XM_028862605.1 | paired immunoglobulin-like type 2 receptor beta-2 (LOC114687973), mRNA | 0.753 | 1.533 | | XM_028879073.1 | Fli-1 proto-oncogene, ETS transcription factor (Fli1), mRNA | -0.555 | -0.305 |
| XM_028869794.1 | G protein subunit gamma 11 (Gng11), mRNA | 0.752 | -0.303 | | XM_028878391.1 | nuclear receptor coactivator 4 (Ncoa4), transcript variant X2, mRNA | -0.556 | 0.535 |
| XM_028880389.1 | histone H4 (LOC114700697), mRNA | 0.750 | -2.243 | | XM_028875207.1 | eukaryotic translation initiation factor 4A2 (Eif4a2), transcript variant X | -0.557 | -0.687 |
| XM_028877652.1 | integral membrane protein 2B (Itm2b), mRNA | 0.748 | 0.385 | | XM_028867766.1 | transmembrane protein 14C (Tmem14c), transcript variant X1, mRNA | -0.558 | 0.719 |
| XM_028872172.1 | histone H2A.J (LOC114695027), mRNA | 0.748 | 0.074 | | XR_003735543.1 | small nucleolar RNA SNORD86 (LOC114699541), ncRNA | -0.558 | -0.491 |
| XM_028889073.1 | ras homolog family member G (Rhog), mRNA | 0.747 | 0.799 | | XM_028865081.1 | C-terminal Src kinase (Csk), transcript variant X1, mRNA | -0.560 | -0.549 |
| XM_028884903.1 | ubiquitin conjugating enzyme E2 M (Ube2m), mRNA | 0.744 | -0.138 | | XM_028867921.1 | eukaryotic translation initiation factor 3 subunit H (Eif3h), mRNA | -0.565 | -0.094 |
| XM_028880043.1 | antioxidant 1 copper chaperone (Atox1), mRNA | 0.743 | 0.846 | | XM_028875671.1 | DNA topoisomerase II beta (Top2b), mRNA | -0.565 | -0.365 |
| XM_028857402.1 | CDC42 small effector 1 (Cdc42se1), mRNA | 0.741 | 0.194 | | XM_028877887.1 | eukaryotic translation initiation factor 3 subunit M (Eif3m), mRNA | -0.570 | 0.153 |
| XM_028855986.1 | SH3 domain binding glutamate rich protein like (Sh3bgrl), mRNA | 0.737 | 0.598 | | XM_028885488.1 | bolA family member 2B (Bola2b), transcript variant X2, mRNA | -0.580 | 0.293 |
| XM_028855795.1 | guanine nucleotide-binding protein G(s) subunit alpha isoforms short (LOC114682029), transcript variant X4, mRNA | 0.730 | -0.268 | | XM_028869018.1 | ATP synthase membrane subunit c locus 1 (Atp5mc1), transcript variant | -0.580 | 1.405 |
| XM_028862615.1 | glutathione S-transferase alpha-4-like (LOC114687986), mRNA | 0.730 | 0.229 | | XM_028878552.1 | NOP56 ribonucleoprotein (Nop56), transcript variant X1, mRNA | -0.582 | -0.177 |
| XM_028890929.1 | Kruppel like factor 6 (Klf6), mRNA | 0.724 | 0.119 | | XM_028870020.1 | ribosomal protein L37a (Rpl37a), mRNA | -0.590 | -0.183 |
| XM_028869016.1 | SNF8 subunit of ESCRT-II (Snf8), mRNA | 0.722 | 0.774 | | XM_028873821.1 | interleukin 16 (Il16), transcript variant X4, mRNA | -0.595 | -0.405 |
| XM_028855794.1 | guanine nucleotide-binding protein G(s) subunit alpha isoforms short (LOC114682029), transcript variant X3, mRNA | 0.721 | -0.442 | | XM_028892110.1 | poly(A) binding protein nuclear 1 (Pabpn1), mRNA | -0.595 | -0.105 |
| XM_028856974.1 | ubiquitin A-52 residue ribosomal protein fusion product 1 (Uba52), mRNA | 0.719 | -0.097 | | XR_003734007.1 | small nucleolar RNA SNORD24 (LOC114690032), ncRNA | -0.596 | -1.697 |
| XM_028885495.1 | protein phosphatase 4 catalytic subunit (Ppp4c), transcript variant X1, mRN | 0.717 | 0.634 | | XM_028868904.1 | nucleoside diphosphate kinase B (LOC114692876), mRNA | -0.598 | 0.533 |
| XM_028856743.1 | parvin beta (Parvb), mRNA | 0.716 | 0.179 | | XM_028862909.1 | BTG anti-proliferation factor 1 (Btg1), mRNA | -0.601 | -0.696 |
| XM_028873740.1 | calcium and integrin binding 1 (Cib1), mRNA | 0.713 | 0.537 | | XM_028881486.1 | early activation antigen CD69-like (LOC114701399), mRNA | -0.603 | -1.392 |
| XM_028866148.1 | aldose reductase-related protein 1 (LOC114691036), mRNA | 0.712 | 0.107 | | XM_028862557.1 | H-2 class I histocompatibility antigen, Q10 alpha chain-like (LOC114687 | -0.609 | 0.565 |
| XM_028856053.1 | heterogeneous nuclear ribonucleoprotein C (C1/C2) (Hnrnpc), transcript variant X7, mRNA | 0.709 | 0.670 | | XM_028854995.1 | heterogeneous nuclear ribonucleoprotein H1 (Hnrnph1), transcript varia | -0.614 | -1.136 |
| XM_028885568.1 | integrin subunit alpha L (Itgal), transcript variant X2, mRNA | 0.709 | 0.080 | | XM_028862335.1 | ribosomal protein L28 (Rpl28), mRNA | -0.619 | -0.146 |
| XM_028885047.1 | zinc finger AN1-type containing 3 (Zfand3), mRNA | 0.708 | -0.503 | | XM_028883681.1 | cysteine rich protein 1 (Crip1), mRNA | -0.624 | 0.571 |
| XM_028866633.1 | ribonuclease T2 (Rnaset2), mRNA | 0.704 | 0.707 | | XM_028859691.1 | ribosomal protein L36 (Rpl36), transcript variant X1, mRNA | -0.629 | 0.152 |
| XM_028892841.1 | cofilin 1 (Cfl1), mRNA | 0.703 | 0.728 | | XM_028882808.1 | TRAF3 interacting protein 3 (Traf3ip3), mRNA | -0.642 | -0.571 |
| XR_003734932.1 | uncharacterized LOC114695908 (LOC114695908), ncRNA | 0.703 | -1.022 | | XM_028861502.1 | ribosomal protein S19 (Rps19), transcript variant X1, mRNA | -0.644 | -1.017 |
| XM_028882373.1 | WD repeat domain 1 (Wdr1), mRNA | 0.702 | 0.310 | | XM_028885003.1 | CD7 molecule (Cd7), transcript variant X1, mRNA | -0.648 | -0.427 |
| XM_028891911.1 | tubulin alpha-1C chain (LOC114708557), mRNA | 0.701 | 0.232 | | XM_028860224.1 | required for meiotic nuclear division 5 homolog A (Rmnd5a), mRNA | -0.653 | -0.507 |
| XM_028886154.1 | prosaposin (Psap), transcript variant X4, mRNA | 0.699 | 1.445 | | XM_028872274.1 | interferon-induced transmembrane protein 1-like (LOC114695076), tran | -0.654 | -1.092 |
| XM_028890761.1 | tropomyosin 3 (Tpm3), transcript variant X5, mRNA | 0.698 | 0.108 | | XM_028857934.1 | beta-1,3-glucuronyltransferase 3 (B3gat3), mRNA | -0.655 | 0.036 |
| XM_028876192.1 | SP110 nuclear body protein (Sp110), transcript variant X1, mRNA | 0.687 | -0.404 | | XM_028892068.1 | O-linked N-acetylglucosamine (GlcNAc) transferase (Ogt), transcript va | -0.656 | -1.582 |
| XM_028860993.1 | profilin 1 (Pfn1), mRNA | 0.686 | 0.485 | | XM_028858546.1 | ribosomal protein S15 (Rps15), mRNA | -0.657 | 0.166 |
| XM_028881505.1 | killer cell lectin-like receptor subfamily B member 1F (LOC114701418), mRNA | 0.685 | -0.622 | | XM_028891307.1 | eukaryotic translation elongation factor 1 beta 2 (Eef1b2), mRNA | -0.657 | 0.036 |
| XR_003736179.1 | uncharacterized LOC114703746 (LOC114703746), ncRNA | 0.683 | 0.383 | | XM_028890603.1 | muscleblind like splicing regulator 1 (Mbnl1), transcript variant X17, mR | -0.661 | -0.508 |
| XM_028862548.1 | platelet glycoprotein VI-like (LOC114687928), mRNA | 0.681 | -0.368 | | XM_028861810.1 | immunoglobulin lambda-1 light chain-like (LOC114687105), mRNA | -0.668 | -0.733 |
| XM_028861196.1 | MOB kinase activator 3A (Mob3a), transcript variant X1, mRNA | 0.680 | -2.371 | | XM_028857949.1 | AHNAK nucleoprotein (Ahnak), mRNA | -0.669 | 0.317 |
| XM_028857385.1 | ubiquitin conjugating enzyme E2 D3 (Ube2d3), transcript variant X4, mRNA | 0.679 | 0.340 | | XM_028872314.1 | arginine and glutamate rich 1 (Arglu1), mRNA | -0.675 | -0.350 |
| XM_028888541.1 | GDP dissociation inhibitor 1 (Gdi1), mRNA | 0.679 | 0.145 | | XR_003735818.1 | uncharacterized LOC114701400 (LOC114701400), ncRNA | -0.682 | -0.380 |
| XM_028858452.1 | enolase 1 (Eno1), mRNA | 0.678 | 1.382 | | XM_028856775.1 | ribosomal protein L35 (Rpl35), mRNA | -0.682 | 0.075 |
| XM_028890580.1 | C-C motif chemokine ligand 5 (Ccl5), mRNA | 0.677 | -0.001 | | XM_028892277.1 | prohibitin 2 (Phb2), mRNA | -0.684 | 0.060 |
| XM_028889665.1 | unc-13 homolog D (Unc13d), mRNA | 0.676 | -0.165 | | XM_028870556.1 | myosin IG (Myo1g), mRNA | -0.693 | -0.116 |
| XM_028872757.1 | ubiquitin conjugating enzyme E2 J1 (Ube2j1), mRNA | 0.675 | 0.016 | | XM_028862569.1 | ribosomal protein L18a (Rpl18a), mRNA | -0.694 | 0.157 |
| XM_028858539.1 | calponin 2 (Cnn2), mRNA | 0.673 | 0.050 | | XM_028872259.1 | interferon-induced transmembrane protein 2 (LOC114695064), mRNA | -0.698 | -0.907 |
| XM_028857362.1 | protein PET100 homolog, mitochondrial (LOC114683173), mRNA | 0.672 | 0.139 | | XM_028867635.1 | ribosomal protein lateral stalk subunit P1 (Rplp1), mRNA | -0.702 | -0.120 |
| XM_028881324.1 | WASP actin nucleation promoting factor (Was), mRNA | 0.671 | 0.568 | | XM_028886961.1 | inosine monophosphate dehydrogenase 2 (Impdh2), transcript variant X | -0.702 | 0.484 |
| XM_028865959.1 | mortality factor 4 like 1 (Morf4l1), transcript variant X3, mRNA | 0.667 | 0.242 | | XM_028883160.1 | ribosomal protein S7 (Rps7), mRNA | -0.703 | -0.085 |
| XM_028893737.1 | radical S-adenosyl methionine domain containing 2 (Rsad2), mRNA | 0.667 | 0.294 | | XM_028859835.1 | ELOVL fatty acid elongase 5 (Elovl5), mRNA | -0.706 | -0.002 |
| XM_028855046.1 | RAB4B, member RAS oncogene family (Rab4b), mRNA | 0.666 | 0.580 | | XM_028875683.1 | ribosomal protein L15 (Rpl15), mRNA | -0.707 | 0.085 |
| XM_028870393.1 | PHD finger protein 11 (Phf11), transcript variant X1, mRNA | 0.666 | -0.534 | | XM_028894586.1 | eukaryotic translation initiation factor 3 subunit F (Eif3f), mRNA | -0.710 | -0.054 |
| XM_028866768.1 | tyrosine 3-monooxygenase/tryptophan 5-monooxygenase activation protein epsilon (Ywhae), transcript variant X1, mRNA | 0.665 | 0.652 | | XM_028873802.1 | BTB domain containing 1 (Btbd1), mRNA | -0.711 | -0.376 |
| XM_028859786.1 | resistin-like alpha (LOC114685170), mRNA | 0.664 | 0.000 | | XM_028858907.1 | ribosomal protein S29 (Rps29), mRNA | -0.711 | 0.162 |
| XM_028883087.1 | acyl-CoA thioesterase 7 (Acot7), mRNA | 0.664 | 0.173 | | XM_028891024.1 | ribosomal protein S23 (Rps23), mRNA | -0.711 | 0.004 |
| XM_028891406.1 | stromal cell derived factor 4 (Sdf4), mRNA | 0.650 | 0.265 | | XM_028864618.1 | ribosomal protein L7 (Rpl7), mRNA | -0.713 | -0.099 |
| XM_028885567.1 | integrin subunit alpha L (Itgal), transcript variant X1, mRNA | 0.648 | 0.490 | | XM_028890957.1 | transmembrane protein 123 (Tmem123), mRNA | -0.715 | -0.754 |
| XM_028867784.1 | ADP ribosylation factor like GTPase 6 interacting protein 1 (Arl6ip1), mRNA | 0.642 | 0.570 | | XM_028888353.1 | serine/threonine kinase 38 (Stk38), mRNA | -0.716 | -0.434 |
| XR_003733566.1 | cellular nucleic acid-binding protein pseudogene (LOC114686303), misc_RNA | 0.640 | 2.072 | | XR_003735008.1 | small nucleolar RNA U2-19 (LOC114696382), ncRNA | -0.717 | -0.490 |
| XM_028877913.1 | catalase (Cat), mRNA | 0.633 | 0.769 | | XM_028868399.1 | serine and arginine rich splicing factor 6 (Srsf6), mRNA | -0.717 | -0.369 |
| XM_028855792.1 | guanine nucleotide-binding protein G(s) subunit alpha isoforms short (LOC114682029), transcript variant X1, mRNA | 0.633 | -0.114 | | XM_028864182.1 | translocase of outer mitochondrial membrane 7 (Tomm7), mRNA | -0.718 | 0.285 |
| XM_028893497.1 | tropomyosin 4 (Tpm4), transcript variant X2, mRNA | 0.633 | 0.246 | | XM_028878424.1 | tetraspanin 13 (Tspan13), mRNA | -0.722 | -0.184 |
| XM_028866296.1 | heat shock protein family A (Hsp70) member 8 (Hspa8), mRNA | 0.633 | 0.025 | | XM_028894319.1 | ribosomal protein S20 (Rps20), mRNA | -0.723 | -0.152 |
| XM_028865328.1 | CMRF35-like molecule (LOC114690500), mRNA | 0.632 | 0.663 | | XM_028876638.1 | protein tyrosine phosphatase non-receptor type 7 (Ptpn7), mRNA | -0.723 | -0.395 |
| XM_028870519.1 | Ras suppressor protein 1 (Rsu1), mRNA | 0.632 | 0.044 | | XM_028866873.1 | ribosomal protein L23a (Rpl23a), mRNA | -0.726 | -0.175 |
| XM_028882481.1 | triggering receptor expressed on myeloid cells like 1 (Treml1), mRNA | 0.628 | -0.523 | | XM_028864489.1 | Fc receptor like A (Fcrla), mRNA | -0.727 | -0.397 |
| XM_028855415.1 | myosin light chain 6 (Myl6), transcript variant X2, mRNA | 0.624 | 0.013 | | XM_028882239.1 | small integral membrane protein 14 (Smim14), mRNA | -0.736 | 0.148 |
| XM_028890006.1 | clusterin (Clu), transcript variant X1, mRNA | 0.624 | -0.357 | | XM_028859693.1 | ribosomal protein L36 (Rpl36), transcript variant X2, mRNA | -0.737 | 0.363 |
| XM_028861007.1 | platelet glycoprotein Ib alpha chain-like (LOC114686345), mRNA | 0.619 | 0.327 | | XM_028890796.1 | interleukin enhancer binding factor 2 (Ilf2), mRNA | -0.747 | -0.096 |
| XM_028882606.1 | integrin subunit alpha 2b (Itga2b), mRNA | 0.618 | 0.206 | | XM_028875566.1 | selenoprotein P (Selenop), mRNA | -0.748 | 0.457 |
| XM_028861784.1 | calreticulin (Calr), mRNA | 0.616 | 1.299 | | XM_028892567.1 | ZFP36 ring finger protein like 2 (Zfp36l2), mRNA | -0.751 | -0.942 |
| XM_028886925.1 | shisa family member 5 (Shisa5), transcript variant X2, mRNA | 0.614 | 0.094 | | XM_028862171.1 | misshapen like kinase 1 (Mink1), mRNA | -0.751 | -0.527 |
| XM_028859182.1 | ferritin light chain (Ftl), mRNA | 0.611 | -0.286 | | XM_028854992.1 | heterogeneous nuclear ribonucleoprotein H1 (Hnrnph1), transcript varia | -0.755 | -0.343 |
| XM_028889695.1 | histone H3.3 (LOC114707068), mRNA | 0.609 | 0.217 | | XM_028892545.1 | septin 9 (Septin9), transcript variant X5, mRNA | -0.756 | -0.072 |
| XM_028860755.1 | peroxiredoxin 3 (Prdx3), mRNA | 0.604 | 0.550 | | XM_028879594.1 | GTPase, IMAP family member 4 (Gimap4), transcript variant X1, mRNA | -0.765 | -0.670 |
| XM_028855781.1 | cathepsin Z (Ctsz), mRNA | 0.604 | 1.937 | | XM_028890107.1 | ribosomal protein S14 (Rps14), mRNA | -0.767 | -0.034 |
| XM_028863328.1 | H-2 class I histocompatibility antigen, D-37 alpha chain-like (LOC114688820), mRNA | 0.602 | 0.177 | | XM_028893577.1 | ribosomal protein S24 (Rps24), transcript variant X2, mRNA | -0.768 | -0.080 |
| XM_028856245.1 | MCL1 apoptosis regulator, BCL2 family member (Mcl1), mRNA | 0.599 | 0.427 | | XM_028854724.1 | ribosomal protein SA (Rpsa), mRNA | -0.771 | -0.084 |
| XM_028871324.1 | MOB kinase activator 1A (Mob1a), mRNA | 0.596 | -0.288 | | XM_028863591.1 | heterogeneous nuclear ribonucleoprotein H-like (LOC114689210), mRN | -0.772 | -1.285 |
| XM_028877686.1 | protein disulfide isomerase family A member 6 (Pdia6), mRNA | 0.595 | 1.284 | | XM_028858749.1 | ribosomal protein L7a (Rpl7a), mRNA | -0.773 | -0.993 |
| XM_028884900.1 | charged multivesicular body protein 2A (Chmp2a), transcript variant X3, mRN | 0.594 | 0.267 | | XM_028876154.1 | 40S ribosomal protein S4, X isoform-like (LOC114697806), mRNA | -0.777 | 1.230 |
| XM_028882187.1 | purinergic receptor P2Y12 (P2ry12), transcript variant X1, mRNA | 0.594 | -1.643 | | XM_028879127.1 | interferon induced protein 44 (Ifi44), mRNA | -0.779 | -1.200 |
| XM_028893508.1 | RAB8A, member RAS oncogene family (Rab8a), mRNA | 0.593 | 0.286 | | XM_028867956.1 | eukaryotic translation initiation factor 3 subunit E (Eif3e), mRNA | -0.785 | -0.187 |
| XM_028853724.1 | fermitin family member 3 (Fermt3), transcript variant X3, mRNA | 0.593 | 0.241 | | XM_028879072.1 | ETS proto-oncogene 1, transcription factor (Ets1), transcript variant X2, | -0.787 | -1.276 |
| XR_003733396.1 | uncharacterized LOC114684668 (LOC114684668), ncRNA | 0.592 | -0.395 | | XM_028868164.1 | OTU deubiquitinase with linear linkage specificity like (Otulinl), mRNA | -0.788 | -0.255 |
| XM_028861017.1 | proteasome subunit beta 9 (Psmb9), mRNA | 0.590 | 0.923 | | XM_028879596.1 | GTPase, IMAP family member 8 (Gimap8), mRNA | -0.790 | -0.919 |
| XM_028887657.1 | lysosomal protein transmembrane 5 (Laptm5), mRNA | 0.588 | 1.268 | | XM_028859907.1 | ribosomal protein S28 (Rps28), mRNA | -0.792 | -0.147 |
| XM_028885781.1 | transmembrane p24 trafficking protein 2 (Tmed2), transcript variant X2, mRN | 0.586 | 0.675 | | XM_028895291.1 | ribosomal protein L29 (Rpl29), mRNA | -0.800 | -0.046 |
| XM_028881831.1 | cystatin-C-like (LOC114701688), mRNA | 0.585 | 0.567 | | XM_028861454.1 | eukaryotic translation elongation factor 1 alpha 1 (Eef1a1), mRNA | -0.803 | -0.201 |
| XM_028859909.1 | RAB11B, member RAS oncogene family (Rab11b), mRNA | 0.581 | 0.123 | | XM_028879645.1 | GTPase IMAP family member 7-like (LOC114700138), mRNA | -0.805 | -0.898 |
| XM_028863572.1 | G protein-coupled receptor kinase 6 (Grk6), transcript variant X1, mRNA | 0.581 | 0.089 | | XM_028865359.1 | cell cycle progression 1 (Ccpg1), transcript variant X4, mRNA | -0.806 | -0.515 |
| XM_028854943.1 | transmembrane protein 160 (Tmem160), mRNA | 0.580 | 0.043 | | XM_028856842.1 | ribosomal protein L34 (Rpl34), mRNA | -0.807 | -0.041 |
| XM_028859954.1 | trem-like transcript 1 protein (LOC114685340), mRNA | 0.577 | 0.263 | | XM_028878064.1 | high mobility group box 1 (Hmgb1), transcript variant X2, mRNA | -0.808 | 0.258 |
| XM_028878402.1 | TSC22 domain family member 1 (Tsc22d1), transcript variant X6, mRNA | 0.576 | -0.461 | | XM_028879892.1 | linker for activation of T cells (Lat), transcript variant X2, mRNA | -0.809 | -1.082 |
| XM_028871875.1 | NADH:ubiquinone oxidoreductase subunit B9 (Ndufb9), mRNA | 0.574 | 0.774 | | XM_028859535.1 | mitogen-activated protein kinase kinase kinase kinase 1 (Map4k1), tran | -0.812 | 0.202 |
| XM_028881483.1 | C-type lectin domain family 12 member B (Clec12b), mRNA | 0.574 | 0.387 | | XM_028863695.1 | leukocyte immunoglobulin-like receptor subfamily A member 5 (LOC11 | -0.812 | -0.337 |
| XM_028890426.1 | HIG1 hypoxia inducible domain family member 1A (Higd1a), mRNA | 0.571 | -0.419 | | XM_028867904.1 | interleukin 21 receptor (Il21r), mRNA | -0.823 | -0.871 |
| XM_028853723.1 | fermitin family member 3 (Fermt3), transcript variant X2, mRNA | 0.568 | -0.202 | | XM_028859253.1 | coiled-coil domain containing 114 (Ccdc114), mRNA | -0.824 | -0.237 |
| XM_028877301.1 | ATPase Na+/K+ transporting subunit alpha 1 (Atp1a1), mRNA | 0.565 | 0.514 | | XM_028880881.1 | ribosomal protein L13 (Rpl13), mRNA | -0.827 | -0.118 |
| XM_028858474.1 | proteasome subunit alpha 3 (Psma3), transcript variant X1, mRNA | 0.564 | 0.749 | | XM_028882234.1 | ribosomal protein L9 (Rpl9), mRNA | -0.836 | 0.008 |
| XM_028884049.1 | charged multivesicular body protein 5 (Chmp5), mRNA | 0.563 | 0.072 | | XM_028892082.1 | ribosomal protein S4 X-linked (Rps4x), mRNA | -0.839 | 0.048 |
| XM_028886633.1 | calmodulin 1 (Calm1), mRNA | 0.552 | 0.490 | | XM_028865759.1 | ribosomal protein L31 (Rpl31), transcript variant X1, mRNA | -0.839 | -0.165 |
| XM_028890241.1 | sorting nexin 3 (Snx3), mRNA | 0.551 | 0.693 | | XM_028866903.1 | ribosomal protein L4 (Rpl4), mRNA | -0.839 | -0.003 |
| XM_028868680.1 | pleckstrin and Sec7 domain containing 4 (Psd4), mRNA | 0.549 | 0.294 | | XM_028893197.1 | ribosomal protein S8 (Rps8), mRNA | -0.840 | -0.129 |
| XM_028878810.1 | tescalcin (Tesc), mRNA | 0.548 | -0.747 | | XM_028867398.1 | ribosomal protein L5 (Rpl5), transcript variant X2, mRNA | -0.843 | 0.359 |
| XM_028857213.1 | protein phosphatase 2 catalytic subunit alpha (Ppp2ca), mRNA | 0.548 | 0.232 | | XM_028894598.1 | ribosomal protein L27a (Rpl27a), mRNA | -0.844 | 0.096 |
| XM_028882650.1 | formin like 1 (Fmnl1), mRNA | 0.547 | 0.110 | | XM_028871801.1 | NADH:ubiquinone oxidoreductase core subunit V1 (Ndufv1), mRNA | -0.847 | 0.027 |
| XM_028887936.1 | lymphotoxin beta (Ltb), transcript variant X2, mRNA | 0.545 | -0.232 | | XM_028876231.1 | prothymosin alpha (Ptma), mRNA | -0.850 | 0.293 |
| XM_028861535.1 | protein phosphatase 2 scaffold subunit Aalpha (Ppp2r1a), mRNA | 0.545 | 0.394 | | XM_028859214.1 | ribosomal protein L18 (Rpl18), mRNA | -0.852 | 0.167 |
| XM_028865173.1 | LIM domain containing 2 (Limd2), transcript variant X1, mRNA | 0.538 | 0.305 | | XM_028856867.1 | ribosomal protein L27 (Rpl27), mRNA | -0.857 | -0.056 |
| XM_028880703.1 | NECAP endocytosis associated 2 (Necap2), transcript variant X1, mRNA | 0.537 | -0.242 | | XM_028854936.1 | NOP53 ribosome biogenesis factor (Nop53), mRNA | -0.860 | -0.333 |
| XM_028881270.1 | lysosomal associated membrane protein 1 (Lamp1), mRNA | 0.537 | 0.922 | | XM_028885113.1 | pinin, desmosome associated protein (Pnn), mRNA | -0.863 | -0.827 |
| XM_028888386.1 | TAP binding protein (Tapbp), mRNA | 0.534 | 0.090 | | XM_028869409.1 | ribosomal protein L26 (Rpl26), mRNA | -0.866 | -0.084 |
| XM_028885687.1 | chromosome unknown C16orf54 homolog (CUNH16orf54), transcript variant X2, m | 0.533 | 1.024 | | XM_028873788.1 | ribosomal protein S17 (Rps17), mRNA | -0.871 | -0.214 |
| XM_028887332.1 | NADH:ubiquinone oxidoreductase subunit A1 (Ndufa1), mRNA | 0.530 | 0.144 | | XM_028894706.1 | mitotic-spindle organizing protein 2-like (LOC114710540), mRNA | -0.876 | 0.029 |
| XM_028888277.1 | interferon alpha-inducible protein 27, mitochondrial-like (LOC114706055), t | 0.530 | 0.042 | | XM_028862728.1 | eukaryotic translation elongation factor 2 (Eef2), mRNA | -0.878 | -0.169 |

| | | | |
|---|---|---|---|
| XM_028863485.1 | transmembrane p24 trafficking protein 9 (Tmed9), mRNA | 0.528 | 1.132 |
| XM_028894746.1 | clathrin light chain A (Clta), transcript variant X4, mRNA | 0.528 | 1.163 |
| XM_028855783.1 | tubulin beta 1 class VI (Tubb1), mRNA | 0.526 | -0.180 |
| XM_028861013.1 | bromodomain containing 2 (Brd2), transcript variant X2, mRNA | 0.525 | -0.011 |
| XM_028856434.1 | prostaglandin reductase 1 (Ptgr1), mRNA | 0.519 | -0.242 |
| XM_028894964.1 | cathepsin C (Ctsc), transcript variant X1, mRNA | 0.516 | 1.500 |
| XM_028883986.1 | valosin containing protein (Vcp), transcript variant X1, mRNA | 0.516 | 0.521 |
| XM_028855254.1 | cytochrome c oxidase subunit 6B1 (LOC114681644), mRNA | 0.516 | 0.419 |
| XM_028893783.1 | RNA binding motif protein 38 (Rbm38), mRNA | 0.515 | 0.289 |
| XM_028858199.1 | integrin linked kinase (Ilk), mRNA | 0.515 | 0.150 |
| XM_028884491.1 | tyrosine 3-monooxygenase/tryptophan 5-monooxygenase activation protein zeta (Ywhaz) | 0.512 | 0.403 |
| XM_028878073.1 | mesenteric estrogen dependent adipogenesis (Medag), mRNA | 0.506 | -1.113 |
| XM_028855626.1 | ribophorin II (Rpn2), transcript variant X3, mRNA | 0.506 | 0.686 |
| XM_028873067.1 | GLI pathogenesis related 1 (Glipr1), mRNA | 0.505 | 1.206 |
| XM_028861049.1 | H-2 class I histocompatibility antigen, Q10 alpha chain-like (LOC114686366), transcript variant X16, mRNA | 0.504 | -0.027 |
| XM_028883807.1 | heat shock protein 90 alpha family class A member 1 (Hsp90aa1), mRNA | 0.503 | 0.246 |
| XM_028867833.1 | cytochrome b-c1 complex subunit 2, mitochondrial (LOC114692176), mRNA | 0.500 | 0.323 |
| XM_028883549.1 | thymosin beta 4 X-linked (Tmsb4x), mRNA | 0.498 | 0.215 |
| XM_028856755.1 | parvin gamma (Parvg), mRNA | 0.498 | 0.033 |
| XM_028867707.1 | guanosine monophosphate reductase (Gmpr), mRNA | 0.497 | -0.521 |
| XM_028881399.1 | tubulin beta 2A class IIa (Tubb2a), mRNA | 0.496 | 0.243 |
| XM_028878239.1 | SPT4 homolog, DSIF elongation factor subunit (Supt4h1), mRNA | 0.494 | 0.532 |
| XM_028876053.1 | histone H3.3-like (LOC114697740), mRNA | 0.494 | 0.240 |
| XM_028878715.1 | small EDRK-rich factor 2 (Serf2), transcript variant X2, mRNA | 0.493 | -0.089 |
| XM_028855189.1 | vesicle associated membrane protein 8 (Vamp8), mRNA | 0.493 | 0.134 |
| XM_028891463.1 | aurora kinase A interacting protein 1 (Aurkaip1), mRNA | 0.491 | 0.530 |
| XM_028856557.1 | protein kinase AMP-activated non-catalytic subunit beta 1 (Prkab1), mRNA | 0.491 | -0.038 |
| XM_028861732.1 | H-2 class I histocompatibility antigen, D-37 alpha chain-like (LOC114687019), mRNA | 0.489 | 0.554 |
| XM_028859357.1 | epsin 1 (Epn1), transcript variant X2, mRNA | 0.487 | -0.899 |
| XM_028853566.1 | actin gamma 1 (Actg1), mRNA | 0.487 | 0.842 |
| XM_028869630.1 | RAP1B, member of RAS oncogene family (Rap1b), mRNA | 0.485 | 0.339 |
| XM_028884898.1 | charged multivesicular body protein 2A (Chmp2a), transcript variant X1, mRN | 0.484 | -0.123 |
| XM_028856446.1 | polypyrimidine tract binding protein 3 (Ptbp3), transcript variant X1, mRNA | 0.479 | -0.410 |
| XM_028855191.1 | vesicle associated membrane protein 5 (Vamp5), mRNA | 0.478 | 0.077 |
| XM_028868279.1 | gamma-glutamylamine cyclotransferase (Ggact), transcript variant X1, mRNA | 0.475 | 0.060 |
| XM_028857532.1 | histone H2A.Z (LOC114683296), mRNA | 0.474 | 0.822 |
| XM_028866102.1 | short transmembrane mitochondrial protein 1 (Stmp1), mRNA | 0.473 | -0.928 |
| XM_028878995.1 | CD82 molecule (Cd82), mRNA | 0.473 | -0.478 |
| XM_028893767.1 | cytochrome b-c1 complex subunit 7 (LOC114709796), mRNA | 0.471 | -0.079 |
| XM_028861730.1 | H-2 class I histocompatibility antigen, Q10 alpha chain-like (LOC114687018), transcript variant X2, mRNA | 0.471 | 0.168 |
| XM_028861032.1 | H-2 class I histocompatibility antigen, Q10 alpha chain-like (LOC114686366), transcript variant X1, mRNA | 0.470 | 0.323 |
| XM_028871763.1 | capping actin protein of muscle Z-line subunit alpha 2 (Capza2), mRNA | 0.469 | 0.521 |
| XM_028862563.1 | H-2 class I histocompatibility antigen, Q10 alpha chain-like (LOC114687936), mRNA | 0.469 | 0.280 |
| XM_028856832.1 | beclin 1 (Becn1), transcript variant X2, mRNA | 0.467 | -0.495 |
| XR_003734151.1 | uncharacterized LOC114690910 (LOC114690910), ncRNA | 0.466 | 0.214 |
| XM_028888380.1 | prefoldin subunit 6 (Pfdn6), mRNA | 0.464 | 0.035 |
| XM_028861176.1 | MAPK interacting serine/threonine kinase 2 (Mknk2), transcript variant X2, mRNA | 0.463 | -0.087 |
| XM_028875919.1 | DEAD-box helicase 3 X-linked (Ddx3x), transcript variant X3, mRNA | 0.462 | -0.282 |
| XM_028889382.1 | PDZ and LIM domain 1 (Pdlim1), mRNA | 0.462 | -0.632 |
| XM_028860653.1 | DNA polymerase delta 4, accessory subunit (Pold4), mRNA | 0.462 | 0.246 |
| XM_028885491.1 | mitogen-activated protein kinase 3 (Mapk3), mRNA | 0.461 | 0.468 |
| XM_028859477.1 | neurogranin (Nrgn), mRNA | 0.458 | -0.079 |
| XM_028864376.1 | cellular repressor of E1A stimulated genes 1 (Creg1), transcript variant X1, mRNA | 0.457 | 0.537 |
| XM_028887799.1 | voltage dependent anion channel 3 (Vdac3), transcript variant X2, mRNA | 0.456 | -0.322 |
| XM_028875283.1 | mitochondrial contact site and cristae organizing system subunit 10 (Micos10), mRNA | 0.455 | 0.736 |
| XM_028880766.1 | growth arrest and DNA damage inducible alpha (Gadd45a), mRNA | 0.453 | 0.819 |
| XM_028855285.1 | calpain small subunit 1 (Capns1), mRNA | 0.451 | -0.212 |
| XM_028869723.1 | proteasome subunit alpha 6 (Psma6), mRNA | 0.450 | 0.739 |
| XM_028893074.1 | zinc finger AN1-type containing 2B (Zfand2b), mRNA | 0.450 | 0.010 |
| XM_028866047.1 | tropomyosin 1 (Tpm1), transcript variant X2, mRNA | 0.447 | 0.860 |
| XR_003733047.1 | 5S ribosomal RNA (LOC114682481), rRNA | 0.444 | 0.940 |
| XM_028876619.1 | adiponectin receptor 1 (Adipor1), transcript variant X1, mRNA | 0.443 | 0.509 |
| XM_028891951.1 | tubulin alpha-1A chain-like (LOC114708577), mRNA | 0.442 | 0.421 |
| XM_028891197.1 | GDP dissociation inhibitor 2 (Gdi2), mRNA | 0.441 | 0.544 |
| XM_028888920.1 | dynein light chain roadblock-type 1 (Dynlrb1), mRNA | 0.439 | 0.206 |
| XM_028866402.1 | archain 1 (Arcn1), mRNA | 0.439 | 0.085 |
| XM_028862558.1 | H-2 class I histocompatibility antigen, Q10 alpha chain-like (LOC114687933), transcript variant X5, mRNA | 0.437 | 1.142 |
| XM_028856538.1 | coagulation factor XIII A chain-like (LOC114682592), mRNA | 0.437 | -0.344 |
| XM_028871253.1 | NADH:ubiquinone oxidoreductase subunit A6 (Ndufa6), mRNA | 0.435 | 0.009 |
| XM_028854998.1 | sequestosome 1 (Sqstm1), mRNA | 0.434 | 0.238 |
| XM_028888094.1 | signal recognition particle 14 (Srp14), mRNA | 0.428 | 0.569 |
| XM_028876655.1 | troponin T2, cardiac type (Tnnt2), mRNA | 0.425 | 0.170 |
| XM_028857334.1 | CDGSH iron sulfur domain 2 (Cisd2), mRNA | 0.421 | -0.019 |
| XM_028888172.1 | ATP synthase inhibitory factor subunit 1 (Atp5if1), mRNA | 0.419 | 0.143 |
| XR_003735476.1 | uncharacterized LOC114699149 (LOC114699149), ncRNA | 0.418 | -0.691 |
| XM_028890144.1 | joining chain of multimeric IgA and IgM (Jchain), mRNA | 0.418 | -1.333 |
| XM_028872417.1 | ubiquitin like 5 (Ubl5), transcript variant X2, mRNA | 0.418 | 0.661 |
| XM_028868860.1 | interleukin 36 beta (Il36b), mRNA | 0.417 | -0.816 |
| XM_028855606.1 | myosin light chain 9 (Myl9), transcript variant X2, mRNA | 0.417 | -0.355 |
| XM_028888233.1 | interferon alpha-inducible protein 27-like protein 1 (LOC114706033), mRNA | 0.415 | 0.523 |
| XM_028855813.1 | adenylate kinase 2 (Ak2), transcript variant X1, mRNA | 0.415 | -1.574 |
| XM_028855814.1 | adenylate kinase 2 (Ak2), transcript variant X2, mRNA | 0.414 | -2.284 |
| XM_028888901.1 | charged multivesicular body protein 4B (Chmp4b), mRNA | 0.411 | -0.141 |
| XM_028870889.1 | CD81 molecule (Cd81), mRNA | 0.410 | 0.800 |
| XM_028886975.1 | ubiquitin specific peptidase 4 (Usp4), mRNA | 0.409 | 0.153 |
| XM_028857243.1 | ubiquitin conjugating enzyme E2 B (Ube2b), mRNA | 0.408 | 0.820 |
| XM_028873756.1 | yippee like 5 (Ypel5), transcript variant X1, mRNA | 0.408 | 0.154 |
| XM_028862787.1 | sphingosine-1-phosphate receptor 4 (S1pr4), mRNA | 0.407 | -0.118 |
| XM_028889855.1 | protein tyrosine phosphatase receptor type C (Ptprc), mRNA | 0.407 | -0.264 |
| XM_028885489.1 | coronin 1A (Coro1a), transcript variant X1, mRNA | 0.403 | 0.083 |
| XM_028874078.1 | myosin regulatory light polypeptide 9 (LOC114696416), transcript variant X2, mRNA | 0.403 | 0.589 |
| XM_028874519.1 | oligosaccharyltransferase complex subunit 4, non-catalytic (Ost4), transcript variant X1, mRNA | 0.402 | -0.466 |
| XM_028892994.1 | CKLF like MARVEL transmembrane domain containing 7 (Cmtm7), mRNA | 0.402 | 0.286 |
| XM_028890834.1 | protein S100-A4 (LOC114707922), mRNA | 0.402 | 1.933 |
| XM_028868771.1 | endothelial differentiation related factor 1 (Edf1), mRNA | 0.401 | -0.049 |
| XM_028861226.1 | SH3 domain containing GRB2 like, endophilin B1 (Sh3glb1), transcript variant X3, mRNA | 0.398 | 0.278 |
| XM_028854913.1 | calmodulin 3 (Calm3), mRNA | 0.397 | -0.263 |
| XM_028874073.1 | myosin regulatory light chain 12B (LOC114696413), mRNA | 0.396 | 0.164 |
| XM_028868423.1 | serine incorporator 3 (Serinc3), mRNA | 0.395 | 0.036 |
| XM_028880781.1 | coactosin like F-actin binding protein 1 (Cotl1), mRNA | 0.392 | 0.704 |
| XR_003734526.1 | uncharacterized LOC114693285 (LOC114693285), ncRNA | 0.388 | -0.019 |
| XM_028890334.1 | ATP synthase membrane subunit f (Atp5mf), mRNA | 0.387 | 0.261 |
| XM_028882581.1 | coagulation factor XIII A chain-like (LOC114702221), mRNA | 0.387 | -0.365 |
| XM_028869999.1 | coxsackievirus and adenovirus receptor homolog (LOC114693581), mRNA | 0.386 | 0.737 |
| XM_028876620.1 | adiponectin receptor 1 (Adipor1), transcript variant X2, mRNA | 0.379 | 0.841 |
| XM_028889938.1 | RAP1A, member of RAS oncogene family (Rap1a), mRNA | 0.378 | 0.007 |
| XM_028864058.1 | heterogeneous nuclear ribonucleoprotein F (Hnrnpf), transcript variant X1, mRNA | 0.377 | 0.189 |
| XR_003733828.1 | uncharacterized LOC114688977 (LOC114688977), ncRNA | 0.376 | -1.548 |
| XM_028875821.1 | eukaryotic translation initiation factor 4 gamma 2 (Eif4g2), mRNA | 0.375 | 0.108 |
| XR_003735045.1 | uncharacterized LOC114696652 (LOC114696652), ncRNA | 0.375 | -1.264 |
| XM_028882602.1 | solute carrier family 25 member 39 (Slc25a39), transcript variant X1, mRNA | 0.374 | -0.140 |
| XR_003733829.1 | uncharacterized LOC114688978 (LOC114688978), ncRNA | 0.374 | -1.554 |
| XR_003735752.1 | 5S ribosomal RNA (LOC114701005), rRNA | 0.371 | 0.773 |
| XR_003735525.1 | probable ATP-dependent RNA helicase DDX5 pseudogene (LOC114699416), misc_RN | 0.370 | -0.528 |
| XM_028855661.1 | SH2B adaptor protein 3 (Sh2b3), mRNA | 0.370 | 0.116 |
| XM_028854101.1 | thioredoxin interacting protein (Txnip), mRNA | 0.369 | -1.053 |
| XM_028891430.1 | protein kinase C theta (Prkcq), transcript variant X2, mRNA | 0.369 | -0.857 |
| XM_028870869.1 | tetraspanin 32 (Tspan32), transcript variant X2, mRNA | 0.369 | 0.214 |
| XM_028888793.1 | nardilysin convertase (Nrdc), mRNA | 0.368 | -0.228 |
| XM_028878045.1 | proteasome maturation protein (Pomp), mRNA | 0.368 | 0.715 |
| XM_028861050.1 | H-2 class I histocompatibility antigen, Q10 alpha chain-like (LOC114686366), transcript variant X17, mRNA | 0.368 | 0.113 |
| XM_028880258.1 | histone H2B type 1 (LOC114700585), mRNA | 0.367 | -0.485 |
| XM_028854784.1 | cytochrome c oxidase subunit 6A1, mitochondrial (LOC114681332), mRNA | 0.366 | 0.586 |
| XM_028893521.1 | family with sequence similarity 32 member A (Fam32a), mRNA | 0.365 | -0.205 |
| XM_028855414.1 | myosin light chain 6 (Myl6), transcript variant X1, mRNA | 0.364 | 0.557 |
| XM_028871233.1 | desumoylating isopeptidase 1 (Desi1), mRNA | 0.363 | 0.017 |
| XM_028861903.1 | protein phosphatase 2 regulatory subunit B'alpha (Ppp2r5a), mRNA | 0.363 | 0.016 |
| XM_028876468.1 | Kell metallo-endopeptidase (Kell blood group) (Kel), mRNA | 0.361 | 0.253 |

Table S18

| | | | |
|---|---|---|---|
| XM_028871942.1 | 60S ribosomal protein L17 (LOC114694874), mRNA | -0.878 | 0.006 |
| XM_028875816.1 | ribosomal protein L22 like 1 (Rpl22l1), transcript variant X1, mRNA | -0.878 | 2.129 |
| XM_028874009.1 | 40S ribosomal protein S6 (LOC114696334), mRNA | -0.881 | -0.066 |
| XM_028868992.1 | family with sequence similarity 117 member A (Fam117a), mRNA | -0.882 | 0.058 |
| XM_028858575.1 | cold inducible RNA binding protein (Cirbp), mRNA | -0.882 | -0.319 |
| XM_028891549.1 | ribosomal protein S27a (Rps27a), mRNA | -0.884 | -0.047 |
| XM_028868981.1 | protein phosphatase 1 regulatory subunit 9B (Ppp1r9b), mRNA | -0.884 | -0.262 |
| XM_028865190.1 | intercellular adhesion molecule 2 (Icam2), mRNA | -0.890 | -0.100 |
| XM_028887334.1 | ribosomal protein L39 (Rpl39), mRNA | -0.892 | 0.023 |
| XM_028867140.1 | acidic nuclear phosphoprotein 32 family member A (Anp32a), transcript | -0.892 | -0.158 |
| XM_028874506.1 | eukaryotic translation initiation factor 4B (Eif4b), mRNA | -0.894 | -0.388 |
| XM_028866355.1 | sorting nexin 1 (Snx1), mRNA | -0.896 | -0.347 |
| XM_028863998.1 | ribosomal protein L32 (Rpl32), mRNA | -0.897 | -0.018 |
| XM_028879128.1 | interferon induced protein 44 like (Ifi44l), mRNA | -0.899 | -1.166 |
| XM_028857939.1 | eukaryotic translation elongation factor 1 gamma (Eef1g), mRNA | -0.908 | -0.532 |
| XM_028884576.1 | ribosomal protein S13 (Rps13), mRNA | -0.909 | -0.241 |
| XM_028884902.1 | ribosomal protein S5 (Rps5), mRNA | -0.909 | 0.107 |
| XM_028888388.1 | ribosomal protein S18 (Rps18), mRNA | -0.912 | -0.134 |
| XM_028855663.1 | ribosomal protein L6 (Rpl6), mRNA | -0.913 | 0.003 |
| XM_028888762.1 | receptor for activated C kinase 1 (Rack1), mRNA | -0.916 | -0.070 |
| XM_028890844.1 | ribosomal protein S27 (Rps27), mRNA | -0.920 | -0.172 |
| XM_028875430.1 | ribosomal protein L11 (Rpl11), mRNA | -0.920 | -0.051 |
| XM_028885333.1 | ribosomal protein S21 (Rps21), mRNA | -0.922 | -0.240 |
| XM_028887783.1 | RB binding protein 7, chromatin remodeling factor (Rbbp7), mRNA | -0.923 | -0.407 |
| XM_028866372.1 | ribosomal protein S25 (Rps25), mRNA | -0.923 | -0.042 |
| XM_028863215.1 | 60S ribosomal protein L35a (LOC114688674), mRNA | -0.928 | 0.111 |
| XM_028879581.1 | GTPase IMAP family member 1-like (LOC114700095), transcript variant | -0.930 | -1.161 |
| XR_003735494.1 | 40S ribosomal protein S19 pseudogene (LOC114699272), misc_RNA | -0.930 | -0.035 |
| XM_028876589.1 | small nuclear ribonucleoprotein polypeptide E (Snrpe), mRNA | -0.937 | -0.251 |
| XM_028854008.1 | small nuclear ribonucleoprotein polypeptide G (Snrpg), mRNA | -0.941 | 0.196 |
| XM_028869108.1 | ribosomal protein L23 (Rpl23), mRNA | -0.944 | -0.097 |
| XM_028861747.1 | 60S ribosomal protein L7a (LOC114687032), mRNA | -0.946 | 0.753 |
| XM_028878028.1 | ribosomal protein L21 (Rpl21), mRNA | -0.946 | -0.041 |
| XM_028889523.1 | C-C motif chemokine receptor 7 (Ccr7), mRNA | -0.947 | -0.737 |
| XM_028854714.1 | ribosomal protein L14 (Rpl14), mRNA | -0.947 | -0.160 |
| XM_028858704.1 | ribosomal protein S3A (Rps3a), mRNA | -0.952 | -0.155 |
| XM_028883079.1 | ribosomal protein L22 (Rpl22), transcript variant X1, mRNA | -0.952 | -0.554 |
| XM_028889552.1 | ribosomal protein L19 (Rpl19), mRNA | -0.953 | 0.091 |
| XM_028861697.1 | tubulin beta class I (Tubb), mRNA | -0.956 | -0.198 |
| XM_028871173.1 | ribosomal protein L3 (Rpl3), mRNA | -0.957 | 0.160 |
| XM_028861900.1 | ribosomal protein S11 (Rps11), mRNA | -0.975 | -0.202 |
| XM_028876015.1 | transmembrane protein 71 (Tmem71), mRNA | -0.977 | -0.453 |
| XM_028877501.1 | ribosomal protein S3 (Rps3), mRNA | -0.979 | -0.148 |
| XM_028867397.1 | ribosomal protein L5 (Rpl5), transcript variant X1, mRNA | -0.982 | -0.200 |
| XM_028855499.1 | ribosomal protein S26 (Rps26), mRNA | -0.989 | -0.045 |
| XM_028860607.1 | SEC11 homolog C, signal peptidase complex subunit (Sec11c), transcript | -0.989 | -0.024 |
| XM_028884342.1 | membrane spanning 4-domains A1 (Ms4a1), mRNA | -0.990 | -0.639 |
| XM_028861811.1 | immunoglobulin lambda-1 light chain-like (LOC114687106), mRNA | -0.992 | -0.761 |
| XM_028884463.1 | ribosomal protein L30 (Rpl30), transcript variant X2, mRNA | -0.999 | -0.318 |
| XM_028867782.1 | ribosomal protein S15a (Rps15a), mRNA | -1.000 | 0.030 |
| XM_028879589.1 | GTPase IMAP family member 6-like (LOC114700097), mRNA | -1.001 | -1.312 |
| XM_028875039.1 | 60S ribosomal protein L36a-like (LOC114697018), mRNA | -1.007 | 0.093 |
| XM_028879593.1 | GTPase, IMAP family member 6 (Gimap6), mRNA | -1.015 | -1.948 |
| XM_028859132.1 | ribosomal protein S16 (Rps16), mRNA | -1.017 | -0.092 |
| XM_028895082.1 | interleukin 7 receptor (Il7r), mRNA | -1.018 | -0.884 |
| XM_028883169.1 | ADAM like decysin 1 (Adamdec1), mRNA | -1.032 | 1.233 |
| XM_028885692.1 | apoptosis-associated speck-like protein containing a CARD (LOC114704 | -1.046 | -0.226 |
| XM_028885545.1 | PAXIP1 associated glutamate rich protein 1 (Pagr1), mRNA | -1.052 | 0.066 |
| XM_028854941.1 | selenoprotein W (Selenow), mRNA | -1.063 | 0.204 |
| XM_028873742.1 | LBH regulator of WNT signaling pathway (Lbh), mRNA | -1.064 | -1.113 |
| XM_028894173.1 | selenoprotein H (Selenoh), mRNA | -1.065 | 0.146 |
| XM_028854718.1 | nucleoporin 210 (Nup210), mRNA | -1.065 | -0.649 |
| XM_028863729.1 | NACHT, LRR and PYD domains-containing protein 1a-like (LOC11468944 | -1.065 | -0.794 |
| XM_028884031.1 | DDB1 and CUL4 associated factor 12 (Dcaf12), mRNA | -1.089 | 0.692 |
| XM_028858994.1 | calpain 2 (Capn2), mRNA | -1.100 | -0.330 |
| XM_028858466.1 | phosphatidylinositol-4,5-bisphosphate 3-kinase catalytic subunit delta (I | -1.103 | -0.729 |
| XM_028875823.1 | switching B cell complex subunit SWAP70 (Swap70), mRNA | -1.112 | -0.238 |
| XM_028888381.1 | ribosomal protein L10a (Rpl10a), mRNA | -1.114 | -0.097 |
| XM_028861878.1 | ribosomal protein L13a (Rpl13a), mRNA | -1.126 | -0.163 |
| XM_028855797.1 | guanine nucleotide-binding protein G(s) subunit alpha isoforms short (L | -1.149 | 1.126 |
| XM_028885560.1 | septin 1 (Septin1), transcript variant X4, mRNA | -1.156 | -0.348 |
| XM_028860649.1 | protein tyrosine phosphatase receptor type C associated protein (Ptprca | -1.179 | -0.280 |
| XM_028880033.1 | annexin A6 (Anxa6), mRNA | -1.198 | -0.382 |
| XM_028861500.1 | CD79a molecule (Cd79a), mRNA | -1.200 | -0.507 |
| XM_028878598.1 | cell division cycle 25B (Cdc25b), mRNA | -1.210 | -1.000 |
| XM_028877269.1 | CD2 molecule (Cd2), mRNA | -1.211 | -0.886 |
| XM_028864239.1 | cytoplasmic FMR1 interacting protein 2 (Cyfip2), transcript variant X1, n | -1.218 | -0.924 |
| XM_028862784.1 | TLE family member 5, transcriptional modulator (Tle5), transcript variar | -1.230 | -0.272 |
| XM_028856303.1 | sphingosine-1-phosphate receptor 1 (S1pr1), mRNA | -1.232 | -0.771 |
| XM_028875051.1 | adducin 3 (Add3), transcript variant X1, mRNA | -1.242 | -0.804 |
| XM_028885764.1 | hydrogen voltage gated channel 1 (Hvcn1), mRNA | -1.245 | -0.628 |
| XM_028879592.1 | GTPase IMAP family member 7-like (LOC114700099), mRNA | -1.291 | -2.025 |
| XM_028862726.1 | immunoglobulin lambda-1 light chain-like (LOC114688112), mRNA | -1.306 | -0.059 |
| XM_028887940.1 | H-2 class II histocompatibility antigen, A-K alpha chain (LOC114705822) | -1.320 | 0.750 |
| XM_028874486.1 | integrin subunit beta 7 (Itgb7), mRNA | -1.323 | -0.615 |
| XM_028869029.1 | src kinase associated phosphoprotein 1 (Skap1), mRNA | -1.331 | -0.812 |
| XM_028857459.1 | B cell scaffold protein with ankyrin repeats 1 (Bank1), mRNA | -1.345 | -0.872 |
| XM_028855951.1 | niban apoptosis regulator 3 (Niban3), mRNA | -1.388 | -0.454 |
| XM_028855469.1 | diacylglycerol kinase alpha (Dgka), transcript variant X1, mRNA | -1.397 | -1.260 |
| XM_028855470.1 | diacylglycerol kinase alpha (Dgka), transcript variant X2, mRNA | -1.414 | -0.808 |
| XM_028866557.1 | CD3e molecule (Cd3e), mRNA | -1.417 | -1.008 |
| XM_028890086.1 | CD74 molecule (Cd74), transcript variant X1, mRNA | -1.435 | 0.232 |
| XM_028872539.1 | acid phosphatase 5, tartrate resistant (Acp5), transcript variant X3, mRI | -1.440 | -1.445 |
| XM_028872901.1 | stannin (Snn), mRNA | -1.478 | -0.621 |
| XM_028893335.1 | neuronal vesicle trafficking associated 2 (Nsg2), mRNA | -1.490 | -0.550 |
| XM_028869325.1 | chromodomain helicase DNA binding protein 3 (Chd3), mRNA | -1.493 | -0.848 |
| XM_028880970.1 | actin binding LIM protein 1 (Ablim1), transcript variant X16, mRNA | -1.500 | -0.647 |
| XM_028887662.1 | LCK proto-oncogene, Src family tyrosine kinase (Lck), mRNA | -1.528 | -0.719 |
| XM_028864375.1 | CD247 molecule (Cd247), mRNA | -1.531 | -0.967 |
| XM_028868389.1 | phospholipase C gamma 1 (Plcg1), mRNA | -1.532 | -1.128 |
| XM_028876399.1 | uncharacterized LOC114697972 (LOC114697972), mRNA | -1.536 | -0.934 |
| XM_028890087.1 | CD74 molecule (Cd74), transcript variant X2, mRNA | -1.541 | 0.287 |
| XM_028856225.1 | 60S ribosomal protein L23-like (LOC114682374), mRNA | -1.541 | 0.466 |
| XM_028855288.1 | B-cell receptor CD22-like (LOC114681663), mRNA | -1.560 | -0.571 |
| XM_028892340.1 | CD27 molecule (Cd27), mRNA | -1.591 | -1.016 |
| XM_028887934.1 | H-2 class II histocompatibility antigen, I-E beta chain-like (LOC11470581 | -1.595 | 0.552 |
| XM_028879590.1 | GTPase IMAP family member 7-like (LOC114700098), transcript variant | -1.613 | -3.559 |
| XM_028861331.1 | B-lymphocyte antigen CD19-like (LOC114686676), mRNA | -1.636 | -0.647 |
| XM_028857667.1 | Fc receptor-like protein 1 (LOC114683366), transcript variant X1, mRNA | -1.641 | -1.186 |
| XM_028874905.1 | Fc fragment of IgM receptor (Fcmr), mRNA | -1.652 | -1.053 |
| XM_028856533.1 | lymphocyte antigen 86 (Ly86), transcript variant X1, mRNA | -1.670 | -0.373 |
| XM_028863530.1 | rano class II histocompatibility antigen, B-1 beta chain-like (LOC114689 | -1.707 | -0.206 |
| XR_003735908.1 | uncharacterized LOC114701976 (LOC114701976), ncRNA | -1.711 | -1.410 |
| XM_028863726.1 | NACHT, LRR and PYD domains-containing protein 1-like (LOC114689440 | -1.754 | -1.408 |
| XM_028887939.1 | H-2 class II histocompatibility antigen, A-K alpha chain (LOC114705822) | -1.761 | 0.278 |
| XM_028887956.1 | H-2 class II histocompatibility antigen, A beta chain-like (LOC114705843 | -1.766 | 0.387 |
| XM_028875068.1 | programmed cell death 4 (Pdcd4), transcript variant X1, mRNA | -1.783 | -0.858 |
| XM_028865312.1 | CD79b molecule (Cd79b), mRNA | -1.784 | -0.473 |
| XM_028887941.1 | H-2 class II histocompatibility antigen, A-F alpha chain (LOC114705823) | -1.785 | 0.238 |
| XM_028857340.1 | Rho/Rac guanine nucleotide exchange factor 18 (Arhgef18), transcript v | -1.826 | -0.949 |
| XM_028892353.1 | CD4 molecule (Cd4), mRNA | -1.827 | -1.068 |
| XM_028884237.1 | CD6 molecule (Cd6), mRNA | -1.830 | -0.654 |
| XM_028873346.1 | forkhead box O1 (Foxo1), mRNA | -1.843 | -1.379 |
| XM_028861027.1 | class II histocompatibility antigen, M beta 1 chain (LOC114686362), mR | -1.942 | 0.968 |
| XM_028888056.1 | H-2 class II histocompatibility antigen, E-K alpha chain (LOC114705896), | -1.942 | 0.454 |
| XM_028872537.1 | acid phosphatase 5, tartrate resistant (Acp5), transcript variant X1, mRI | -1.958 | -2.003 |
| XM_028866553.1 | CD3g molecule (Cd3g), transcript variant X1, mRNA | -2.027 | -1.235 |
| XM_028861026.1 | HLA class II histocompatibility antigen, DO alpha chain (LOC114686361) | -2.116 | -0.076 |
| XM_028866556.1 | CD3d molecule (Cd3d), mRNA | -2.135 | -0.903 |
| XM_028857121.1 | GRB2-related adapter protein-like (LOC114682967), mRNA | -2.157 | -1.121 |
| XM_028884079.1 | signaling threshold regulating transmembrane adaptor 1 (Sit1), mRNA | -2.212 | -1.477 |
| XM_028861028.1 | class II histocompatibility antigen, M alpha chain (LOC114686363), tran: | -2.340 | 1.250 |
| XM_028863532.1 | HLA class II histocompatibility antigen, DO beta chain (LOC114689125), | -2.650 | -0.668 |
| XM_028866554.1 | CD3g molecule (Cd3g), transcript variant X2, mRNA | -2.951 | -1.368 |
| XM_028857356.1 | Fc fragment of IgE receptor II (Fcer2), mRNA | -3.395 | -0.534 |

Table S19

**Table S19. DEGs of *P. leucopus* fibroblast cultures treated with LPS or buffer alone**

| accession | gene | product | Max group mean | fold change | neg log10 FDR |
|---|---|---|---|---|---|
| XM_028874703.1 | Nfkb2 | nuclear factor kappa B subunit 2 (Nfkb2), transcript variant X1, mRNA | 1.80 | 2569.4 | 4.3 |
| XM_028877165.1 | Dis3 | DIS3 homolog, exosome endoribonuclease and 3'-5' exoribonuclease (Dis3), transcript variant X2, mRNA | 1.39 | 2297.3 | 5.7 |
| XM_028880935.1 | Macc1 | MET transcriptional regulator MACC1 (Macc1), transcript variant X2, mRNA | 1.36 | 1861.7 | 4.8 |
| XM_028867321.1 | LOC114691793 | guanylate-binding protein 6-like (LOC114691793), transcript variant X4, mRNA | 1.19 | 1384.4 | 4.6 |
| XM_028875302.1 | Pla2g2a | phospholipase A2 group IIA (Pla2g2a), mRNA | 695.45 | 968.6 | ≤13.5 |
| XM_028860283.1 | Saa3 | serum amyloid A-3 protein-like (LOC114685651), transcript variant X2, mRNA | 166.21 | 940.7 | ≤13.5 |
| XM_028866904.1 | Nos2 | nitric oxide synthase 2 (Nos2), mRNA | 16.90 | 938.0 | ≤13.5 |
| XM_028874061.1 | Ccl20 | C-C motif chemokine ligand 20 (Ccl20), transcript variant X1, mRNA | 40.94 | 929.9 | ≤13.5 |
| XM_028855796.1 | Gnas | guanine nucleotide-binding protein G(s) subunit alpha isoforms short (LOC114682029), transcript variant X5, mRNA | 1.05 | 650.3 | 2.7 |
| XM_028856652.1 | Ubd | ubiquitin D (Ubd), mRNA | 168.45 | 519.6 | ≤13.5 |
| XM_028890580.1 | Ccl5 | C-C motif chemokine ligand 5 (Ccl5), mRNA | 45.75 | 412.5 | ≤13.5 |
| XM_028889425.1 | Csf3 | colony stimulating factor 3 (Csf3), transcript variant X1, mRNA | 11.86 | 380.8 | ≤13.5 |
| XM_028855399.1 | LOC114681724 | guanylate-binding protein 1-like (LOC114681724), transcript variant X2, mRNA | 24.33 | 295.4 | ≤13.5 |
| XR_003736681.1 | LOC114707630 | uncharacterized LOC114707630 (LOC114707630), ncRNA | 7.02 | 271.8 | 7.5 |
| XM_028860282.1 | Saa3 | serum amyloid A-3 protein-like (LOC114685651), transcript variant X1, mRNA | 90.38 | 261.0 | ≤13.5 |
| XM_028863716.1 | Mx2 | interferon-induced GTP-binding protein Mx2-like (LOC114689426), mRNA | 52.93 | 242.7 | ≤13.5 |
| XM_028868484.1 | Cd40 | CD40 molecule (Cd40), mRNA | 44.91 | 209.0 | ≤13.5 |
| XM_028880144.1 | Saa1 | serum amyloid A-1 protein-like (LOC114700507), transcript variant X2, mRNA | 3.37 | 200.5 | 5.9 |
| XM_028855370.1 | LOC114681711 | guanylate-binding protein 1-like (LOC114681711), transcript variant X4, mRNA | 1.14 | 198.3 | 6.7 |
| XM_028855398.1 | LOC114681724 | guanylate-binding protein 1-like (LOC114681724), transcript variant X1, mRNA | 24.58 | 196.5 | ≤13.5 |
| XM_028881923.1 | Cxcl11 | C-X-C motif chemokine ligand 11 (Cxcl11), mRNA | 2.62 | 174.2 | 5.4 |
| XM_028874062.1 | Ccl20 | C-C motif chemokine ligand 20 (Ccl20), transcript variant X2, mRNA | 15.87 | 160.4 | 9.9 |
| XM_028860309.1 | Mx2 | interferon-induced GTP-binding protein Mx2-like (LOC114685662), transcript variant X1, mRNA | 47.65 | 158.5 | 8.8 |
| XM_028859104.1 | Ifit1 | interferon-induced protein with tetratricopeptide repeats 1-like (LOC114684570), mRNA | 5.95 | 144.9 | ≤13.5 |
| XM_028854779.1 | Oasl2 | 2'-5'-oligoadenylate synthase-like protein 2 (LOC114681329), transcript variant X1, mRNA | 15.32 | 143.4 | ≤13.5 |
| XM_028888017.1 | Cfb | complement factor B (Cfb), mRNA | 693.04 | 139.0 | ≤13.5 |
| XM_028859096.1 | Ifit1 | interferon-induced protein with tetratricopeptide repeats 1-like (LOC114684563), mRNA | 13.11 | 136.6 | ≤13.5 |
| XM_028866979.1 | Ccl2 | C-C motif chemokine 2-like (LOC114691550), mRNA | 712.11 | 135.6 | ≤13.5 |
| XM_028860285.1 | Saa5 | serum amyloid A-5 protein (LOC114685652), transcript variant X2, mRNA | 39.47 | 135.5 | ≤13.5 |
| XM_028859102.1 | Ifit1 | interferon-induced protein with tetratricopeptide repeats 1-like (LOC114684568), transcript variant X2, mRNA | 6.98 | 123.9 | 6.7 |
| XM_028858723.1 | Slc2a6 | solute carrier family 2 member 6 (Slc2a6), transcript variant X2, mRNA | 9.39 | 114.8 | ≤13.5 |
| XM_028880204.1 | Saa3 | serum amyloid A-3 protein (LOC114700540), transcript variant X2, mRNA | 76.85 | 114.7 | ≤13.5 |
| XM_028860284.1 | Saa5 | serum amyloid A-5 protein (LOC114685652), transcript variant X1, mRNA | 36.37 | 114.4 | ≤13.5 |
| XM_028883763.1 | Tnfaip2 | TNF alpha induced protein 2 (Tnfaip2), transcript variant X1, mRNA | 78.50 | 104.4 | ≤13.5 |
| XM_028859107.1 | Ifit1 | interferon-induced protein with tetratricopeptide repeats 1-like (LOC114684574), mRNA | 79.00 | 102.9 | ≤13.5 |
| XM_028859097.1 | Ifit1 | interferon-induced protein with tetratricopeptide repeats 1-like (LOC114684565), mRNA | 14.05 | 97.5 | ≤13.5 |
| XR_003735255.1 | LOC114697860 | nuclear autoantigen Sp-100-like (LOC114697860), transcript variant X3, misc | 1.38 | 97.2 | 4.3 |
| XM_028854790.1 | Oasl | 2'-5'-oligoadenylate synthetase like (Oasl), mRNA | 4.61 | 96.0 | ≤13.5 |
| XM_028879921.1 | Mx2 | interferon-induced GTP-binding protein Mx2 (LOC114700347), mRNA | 329.49 | 95.0 | ≤13.5 |
| XM_028859101.1 | LOC114684568 | interferon-induced protein with tetratricopeptide repeats 1-like (LOC114684568), transcript variant X1, mRNA | 6.67 | 92.6 | 5.5 |
| XM_028880143.1 | Saa1 | serum amyloid A-1 protein-like (LOC114700507), transcript variant X1, mRNA | 1.62 | 91.8 | 3.4 |
| XM_028867318.1 | LOC114691793 | guanylate-binding protein 6-like (LOC114691793), transcript variant X1, mRNA | 11.54 | 91.6 | 9.6 |
| XM_028857290.1 | Pglyrp3 | peptidoglycan recognition protein 3-like (LOC114683116), mRNA | 2.12 | 88.7 | 9.0 |
| XM_028869882.1 | Col6a5 | collagen type VI alpha 5 chain (Col6a5), mRNA | 2.44 | 87.7 | ≤13.5 |
| XM_028855361.1 | Gbp1 | guanylate-binding protein 1-like (LOC114681707), transcript variant X1, mRNA | 29.23 | 87.2 | ≤13.5 |
| XM_028867721.1 | Cd83 | CD83 molecule (Cd83), transcript variant X1, mRNA | 6.63 | 85.8 | ≤13.5 |
| XR_003735669.1 | ncRNA | uncharacterized LOC114700341 (LOC114700341), ncRNA | 60.04 | 84.9 | ≤13.5 |
| XM_028859103.1 | Ifit1 | interferon-induced protein with tetratricopeptide repeats 1-like (LOC114684569), mRNA | 8.94 | 83.9 | ≤13.5 |
| XM_028859108.1 | Ifit1 | interferon-induced protein with tetratricopeptide repeats 1-like (LOC114684575), mRNA | 3.88 | 81.7 | 12.2 |
| XM_028855362.1 | LOC114681707 | guanylate-binding protein 1-like (LOC114681707), transcript variant X2, mRNA | 5.35 | 81.7 | 10.4 |
| XM_028870120.1 | Irf7 | interferon regulatory factor 7 (Irf7), transcript variant X4, mRNA | 2.71 | 78.1 | 8.4 |
| XM_028881924.1 | Cxcl10 | C-X-C motif chemokine ligand 10 (Cxcl10), mRNA | 291.46 | 76.7 | ≤13.5 |
| XM_028863000.1 | Iigp1b | interferon-inducible GTPase 1-like (LOC114688415), mRNA | 4.03 | 75.0 | 9.6 |
| XM_028862993.1 | Iigp1b | interferon-inducible GTPase 1-like (LOC114688407), mRNA | 2.52 | 74.9 | 10.0 |
| XM_028859098.1 | Ifit1 | interferon-induced protein with tetratricopeptide repeats 1-like (LOC114684566), mRNA | 8.75 | 72.6 | ≤13.5 |
| XM_028880031.1 | Tnip1 | TNFAIP3 interacting protein 1 (Tnip1), transcript variant X9, mRNA | 7.23 | 72.4 | ≤13.5 |
| XM_028880027.1 | Tnip1 | TNFAIP3 interacting protein 1 (Tnip1), transcript variant X6, mRNA | 72.46 | 71.9 | ≤13.5 |
| XM_028882383.1 | Bbox1 | gamma-butyrobetaine hydroxylase 1 (Bbox1), transcript variant X1, mRNA | 1.21 | 71.3 | 9.3 |
| XM_028859105.1 | Ifit1 | interferon-induced protein with tetratricopeptide repeats 1-like (LOC114684572), mRNA | 8.55 | 63.0 | ≤13.5 |
| XM_028862994.1 | Iigp1b | interferon-inducible GTPase 1-like (LOC114688409), mRNA | 43.44 | 62.4 | ≤13.5 |
| XM_028883764.1 | Tnfaip2 | TNF alpha induced protein 2 (Tnfaip2), transcript variant X2, mRNA | 42.36 | 58.1 | ≤13.5 |
| XM_028868908.1 | LOC114692807 | CD302 antigen (LOC114692807), transcript variant X2, mRNA | 2.29 | 57.4 | ≤13.5 |
| XM_028891462.1 | Isg15 | ISG15 ubiquitin like modifier (Isg15), mRNA | 446.08 | 56.8 | ≤13.5 |
| XM_028865648.1 | Neurl3 | neuralized E3 ubiquitin protein ligase 3 (Neurl3), mRNA | 9.49 | 55.5 | ≤13.5 |
| XM_028862398.1 | Steap4 | STEAP4 metalloreductase (Steap4), mRNA | 18.52 | 54.7 | ≤13.5 |
| XM_028855369.1 | LOC114681711 | guanylate-binding protein 1-like (LOC114681711), transcript variant X3, mRNA | 2.56 | 53.9 | 11.7 |
| XM_028888782.1 | Irgm1 | immunity-related GTPase family M protein 1-like (LOC114706392), mRNA | 35.36 | 52.5 | ≤13.5 |
| XM_028881894.1 | Cxcl1 | growth-regulated alpha protein (LOC114701736), mRNA | 290.79 | 49.3 | ≤13.5 |
| XM_028864484.1 | Selp | selectin P (Selp), mRNA | 273.58 | 48.3 | ≤13.5 |
| XM_028866978.1 | Ccl7 | C-C motif chemokine 7-like (LOC114691549), mRNA | 296.13 | 47.5 | ≤13.5 |
| XM_028880203.1 | Saa3 | serum amyloid A-3 protein (LOC114700540), transcript variant X1, mRNA | 38.85 | 45.6 | 10.5 |
| XM_028880416.1 | Usp18 | ubiquitin specific peptidase 18 (Usp18), mRNA | 45.49 | 43.9 | ≤13.5 |
| XM_028881874.1 | Cxcl5 | C-X-C motif chemokine ligand 5 (Cxcl5), mRNA | 819.66 | 42.9 | ≤13.5 |
| XM_028862995.1 | Iigp1b | interferon-inducible GTPase 1-like (LOC114688413), mRNA | 30.61 | 42.8 | ≤13.5 |
| XM_028860310.1 | LOC114685662 | interferon-induced GTP-binding protein Mx2-like (LOC114685662), transcript variant X2, mRNA | 5.31 | 40.0 | ≤13.5 |
| XM_028855378.1 | Gbp2 | guanylate-binding protein 2-like (LOC114681715), mRNA | 352.82 | 39.7 | ≤13.5 |
| XM_028870218.1 | Ncf1 | neutrophil cytosolic factor 1 (Ncf1), mRNA | 4.72 | 37.7 | 10.9 |
| XM_028864183.1 | Il6 | interleukin 6 (Il6), mRNA | 8.83 | 35.3 | ≤13.5 |
| XM_028860286.1 | Saa5 | serum amyloid A-5 protein (LOC114685652), transcript variant X3, mRNA | 8.91 | 35.1 | 5.8 |
| XM_028880026.1 | Tnip1 | TNFAIP3 interacting protein 1 (Tnip1), transcript variant X5, mRNA | 11.67 | 34.6 | ≤13.5 |
| XM_028855364.1 | Gbp1 | guanylate-binding protein 1-like (LOC114681710), transcript variant X1, mRNA | 2.00 | 34.3 | 9.5 |
| XM_028874788.1 | Calhm6 | calcium homeostasis modulator family member 6 (Calhm6), mRNA | 16.37 | 34.1 | ≤13.5 |
| XM_028855067.1 | Coq8b | coenzyme Q8B (Coq8b), transcript variant X2, mRNA | 1.31 | 33.2 | 2.4 |
| XM_028885765.1 | Hcar2 | hydroxycarboxylic acid receptor 2 (Hcar2), mRNA | 6.28 | 32.9 | ≤13.5 |
| XM_028855388.1 | Gbp3 | guanylate-binding protein 4-like (LOC114681718), mRNA [renamed as Gbp3] | 60.82 | 32.2 | ≤13.5 |
| XM_028881893.1 | Cxcl2 | C-X-C motif chemokine 2-like (LOC114701735), mRNA | 155.28 | 32.2 | ≤13.5 |
| XM_028878765.1 | Oas1g | 2'-5'-oligoadenylate synthase 1A-like (LOC114699623), mRNA | 147.31 | 30.1 | ≤13.5 |
| XM_028893737.1 | Rsad2 | radical S-adenosyl methionine domain containing 2 (Rsad2), mRNA | 25.08 | 29.6 | ≤13.5 |
| XM_028886048.1 | Traf1 | TNF receptor associated factor 1 (Traf1), mRNA | 44.94 | 26.5 | ≤13.5 |
| XM_028875192.1 | Rtp4 | receptor transporter protein 4 (Rtp4), mRNA | 25.79 | 26.3 | ≤13.5 |
| XR_003733498.1 | LOC114685800 | uncharacterized LOC114685800 (LOC114685800), ncRNA | 3.87 | 25.1 | 7.8 |
| XM_028872433.1 | Icam4 | intercellular adhesion molecule 4 (Landsteiner-Wiener blood group) (Icam4), mRNA | 9.00 | 25.0 | 13.2 |
| XM_028856185.1 | LOC114682343 | hereditary hemochromatosis protein-like (LOC114682343), mRNA | 2.68 | 24.0 | 6.9 |
| XM_028895244.1 | Herc6 | HECT and RLD domain containing E3 ubiquitin protein ligase family member 6 | 8.03 | 24.0 | 5.1 |
| XM_028879127.1 | Ifi44 | interferon induced protein 44 (Ifi44), mRNA | 1.05 | 23.7 | 3.4 |
| XM_028894284.1 | Tlr2 | toll like receptor 2 (Tlr2), mRNA | 11.71 | 23.7 | ≤13.5 |
| XM_028891388.1 | Nadk | NAD kinase (Nadk), transcript variant X2, mRNA | 16.63 | 23.7 | ≤13.5 |
| XM_028865122.1 | Plet1 | placenta-expressed transcript 1 protein-like (LOC114690363), mRNA | 1.90 | 23.4 | 4.4 |
| XM_028858458.1 | H6pd | hexose-6-phosphate dehydrogenase/glucose 1-dehydrogenase (H6pd), transcript variant X3, mRNA | 2.80 | 23.3 | 4.6 |
| XM_028873581.1 | Gch1 | GTP cyclohydrolase 1 (Gch1), mRNA | 7.83 | 22.6 | ≤13.5 |
| XM_028863178.1 | LOC114688611 | T-cell-specific guanine nucleotide triphosphate-binding protein 1-like (LOC114688611), mRNA | 3.23 | 22.3 | 5.3 |
| XM_028867319.1 | LOC114691793 | guanylate-binding protein 6-like (LOC114691793), transcript variant X2, mRNA | 10.02 | 22.0 | 8.6 |
| XM_028880030.1 | Tnip1 | TNFAIP3 interacting protein 1 (Tnip1), transcript variant X8, mRNA | 3.42 | 21.7 | 3.9 |
| XM_028855390.1 | LOC114681719 | guanylate-binding protein 4-like (LOC114681719), mRNA | ≤13.54 | 21.5 | ≤13.5 |
| XM_028888273.1 | LOC114706052 | interferon alpha-inducible protein 27-like protein 2A (LOC114706052), mRNA | 36.65 | 21.3 | 11.6 |
| XM_028883765.1 | Exoc3l4 | exocyst complex component 3 like 4 (Exoc3l4), mRNA | 7.85 | 20.9 | ≤13.5 |
| XM_028862997.1 | Iigp1b | interferon-inducible GTPase 1-like (LOC114688414), transcript variant X2, mRNA | 14.86 | 20.9 | ≤13.5 |
| XM_028871456.1 | Cx3cl1 | C-X3-C motif chemokine ligand 1 (Cx3cl1), mRNA [fractaline] | 276.40 | 20.8 | ≤13.5 |
| XM_028891304.1 | Il15ra | interleukin 15 receptor subunit alpha (Il15ra), mRNA | 5.54 | 20.8 | 8.9 |
| XM_028872432.1 | Icam1 | intercellular adhesion molecule 1 (Icam1), mRNA | 649.58 | 20.6 | ≤13.5 |
| XM_028862648.1 | Parp9 | poly(ADP-ribose) polymerase family member 9 (Parp9), transcript variant X2, mRNA | 1.71 | 19.7 | 2.7 |
| XM_028870101.1 | Irf7 | interferon regulatory factor 7 (Irf7), transcript variant X2, mRNA | 4.69 | 19.7 | 5.0 |
| XM_028883166.1 | Cmpk2 | cytidine/uridine monophosphate kinase 2 (Cmpk2), mRNA | 5.90 | 19.5 | ≤13.5 |
| XM_028890852.1 | S100a8 | protein S100-A8-like (LOC114707940), mRNA | 18.33 | 19.2 | 6.7 |
| XM_028872430.1 | Icam5 | intercellular adhesion molecule 5 (Icam5), mRNA | 2.20 | 19.0 | 9.9 |
| XM_028855662.1 | Oas1a | 2'-5'-oligoadenylate synthase 1A-like (LOC114681897), mRNA | 29.00 | 18.8 | 10.8 |
| XM_028884822.1 | Mast4 | microtubule associated serine/threonine kinase family member 4 (Mast4), tra | 5.61 | 18.7 | ≤13.5 |
| XM_028862990.1 | Serpina3n | serine protease inhibitor A3N-like (LOC114688399), mRNA | 146.24 | 18.6 | ≤13.5 |
| XM_028893795.1 | Zbp1 | Z-DNA binding protein 1 (Zbp1), transcript variant X3, mRNA | 2.05 | 18.4 | 5.1 |
| XM_028895246.1 | Herc6 | HECT and RLD domain containing E3 ubiquitin protein ligase family member 6 | 2.56 | 18.1 | 4.0 |
| XM_028881036.1 | LOC114701096 | probable ATP-dependent RNA helicase DDX60 (LOC114701096), mRNA | 10.83 | 17.6 | ≤13.5 |
| XM_028870110.1 | Irf7 | interferon regulatory factor 7 (Irf7), transcript variant X3, mRNA | 26.52 | 17.5 | ≤13.5 |
| XR_003736255.1 | LOC114704347 | uncharacterized LOC114704347 (LOC114704347), ncRNA | 4.28 | 17.3 | 9.3 |
| XM_028865555.1 | LOC114690680 | interferon-inducible protein AIM2-like (LOC114690680), mRNA | 2.94 | 17.0 | 5.5 |
| XM_028870868.1 | Lif | LIF interleukin 6 family cytokine (Lif), mRNA | 28.06 | 16.8 | ≤13.5 |
| XM_028881877.1 | Cxcl3 | C-X-C motif chemokine 3-like (LOC114701719), mRNA | 316.05 | 16.4 | ≤13.5 |
| XM_028878797.1 | Med13l | mediator complex subunit 13L (Med13l), transcript variant X1, mRNA | 1.01 | 16.0 | 2.9 |
| XM_028860432.1 | Icoslg | inducible T cell costimulator ligand (Icoslg), mRNA | 14.38 | 15.5 | ≤13.5 |
| XM_028874699.1 | Mfsd13a | major facilitator superfamily domain containing 13A (Mfsd13a), mRNA | 48.19 | 15.5 | ≤13.5 |
| XM_028876612.1 | Chi3l1 | chitinase 3 like 1 (Chi3l1), mRNA | 1724.53 | 15.1 | ≤13.5 |
| XM_028866678.1 | Sod2 | superoxide dismutase 2 (Sod2), mRNA | 575.23 | 14.8 | ≤13.5 |
| XM_028891607.1 | Serpinb2 | serpin family B member 2 (Serpinb2), mRNA | 1.12 | 14.7 | 2.0 |
| XM_028870099.1 | Irf7 | interferon regulatory factor 7 (Irf7), transcript variant X1, mRNA | 47.93 | 14.7 | ≤13.5 |
| XM_028888232.1 | LOC114706032 | interferon alpha-inducible protein 27-like protein 2A (LOC114706032), mRNA | 94.34 | 14.2 | ≤13.5 |
| XM_028886531.1 | Tmeff2 | transmembrane protein with EGF like and two follistatin like domains 2 (Tme | 10.86 | 14.2 | 6.9 |

Table S19

| | | | | | |
|---|---|---|---|---|---|
| XM_028880298.1 | Dram1 | DNA damage regulated autophagy modulator 1 (Dram1), mRNA | 52.30 | 14.1 | ≤13.5 |
| XM_028892758.1 | Epas1 | endothelial PAS domain protein 1 (Epas1), mRNA | 91.32 | 13.7 | ≤13.5 |
| XM_028891828.1 | Nod2 | nucleotide binding oligomerization domain containing 2 (Nod2), mRNA | 8.23 | 13.7 | ≤13.5 |
| XM_028884633.1 | Ptges | prostaglandin E synthase (Ptges), mRNA | 33.11 | 13.6 | ≤13.5 |
| XM_028875298.1 | Pla2g5 | phospholipase A2 group V (Pla2g5), transcript variant X3, mRNA | 16.77 | 13.5 | 11.5 |
| XM_028879128.1 | Ifi44l | interferon induced protein 44 like (Ifi44l), mRNA | 4.78 | 13.4 | 6.1 |
| XM_028890892.1 | Kcnn3 | potassium calcium-activated channel subfamily N member 3 (Kcnn3), transcrip | 1.85 | 13.3 | 3.4 |
| XM_028871479.1 | Mt2 | metallothionein-2 (LOC114694511), mRNA | 346.02 | 13.3 | ≤13.5 |
| XM_028861564.1 | Trim30a | tripartite motif-containing protein 30A-like (LOC114686894), mRNA | 25.75 | 13.3 | ≤13.5 |
| XM_028862489.1 | Steap2 | STEAP2 metalloreductase (Steap2), transcript variant X4, mRNA | 1.79 | 13.1 | 2.4 |
| XM_028883007.1 | LOC114702542 | paralemmin-2 (LOC114702542), transcript variant X4, mRNA | 14.69 | 13.0 | 5.5 |
| XM_028855950.1 | Bst2 | bone marrow stromal cell antigen 2 (Bst2), mRNA | 87.40 | 13.0 | ≤13.5 |
| XM_028884064.1 | Ddx58 | DExD/H-box helicase 58 (Ddx58), mRNA | 40.80 | 12.7 | ≤13.5 |
| XM_028876201.1 | Sp100 | nuclear autoantigen Sp-100-like (LOC114697861), transcript variant X5, mRNA | 3.48 | 12.5 | 2.8 |
| XM_028876203.1 | LOC114697861 | nuclear autoantigen Sp-100-like (LOC114697861), transcript variant X7, mRNA | 1.90 | 12.4 | 3.7 |
| XM_028865223.1 | Ifih1 | interferon induced with helicase C domain 1 (Ifih1), mRNA | 27.76 | 12.2 | ≤13.5 |
| XM_028886310.1 | Parp14 | poly(ADP-ribose) polymerase family member 14 (Parp14), mRNA | 38.23 | 12.1 | ≤13.5 |
| XM_028880025.1 | Tnip1 | TNFAIP3 interacting protein 1 (Tnip1), transcript variant X4, mRNA | 1.87 | 12.0 | 3.0 |
| XM_028881797.1 | Il15 | interleukin 15 (Il15), transcript variant X2, mRNA | 2.73 | 12.0 | 2.8 |
| XM_028867006.1 | Slfn11 | schlafen family member 11 (Slfn11), mRNA | 2.45 | 11.5 | 3.1 |
| XM_028894207.1 | Ube2l6 | ubiquitin conjugating enzyme E2 L6 (Ube2l6), mRNA | 10.96 | 11.2 | 9.0 |
| XM_028893121.1 | Tmbim1 | transmembrane BAX inhibitor motif containing 1 (Tmbim1), transcript variant | 17.76 | 11.1 | ≤13.5 |
| XM_028877852.1 | Batf | basic leucine zipper ATF-like transcription factor (Batf), mRNA | 4.79 | 10.9 | 3.8 |
| XM_028881796.1 | Il15 | interleukin 15 (Il15), transcript variant X1, mRNA | 3.87 | 10.7 | 4.9 |
| XM_028858080.1 | Wars | tryptophanyl-tRNA synthetase (Wars), transcript variant X2, mRNA | 9.71 | 10.5 | 8.6 |
| XM_028887051.1 | Camp | cathelicidin antimicrobial peptide (Camp), mRNA | 5.23 | 10.4 | 2.9 |
| XM_028856840.1 | Dhx58 | DExH-box helicase 58 (Dhx58), mRNA | 11.25 | 10.2 | ≤13.5 |
| XM_028885451.1 | Zc3h12a | zinc finger CCCH-type containing 12A (Zc3h12a), mRNA | 25.29 | 10.2 | ≤13.5 |
| XM_028860050.1 | Ccrl2 | C-C motif chemokine receptor like 2 (Ccrl2), mRNA | 3.95 | 10.1 | 4.2 |
| XM_028871630.1 | Ripk2 | receptor interacting serine/threonine kinase 2 (Ripk2), transcript variant X2, mRNA | 8.29 | 10.0 | ≤13.5 |
| XM_028862987.1 | Serpina3f | serine protease inhibitor A3F-like (LOC114688398), transcript variant X1, mRNA | 248.00 | 9.8 | 10.3 |
| XM_028892102.1 | Irf9 | interferon regulatory factor 9 (Irf9), transcript variant X3, mRNA | 7.90 | 9.7 | 5.0 |
| XM_028863598.1 | LOC114689226 | guanylate-binding protein 1-like (LOC114689226), mRNA | 5.56 | 9.7 | 4.1 |
| XR_003735432.1 | Mt1 | metallothionein-1 pseudogene (LOC114698905), misc_RNA | 108.86 | 9.6 | ≤13.5 |
| XM_028853654.1 | Rnf213 | ring finger protein 213 (Rnf213), mRNA | 18.06 | 9.6 | ≤13.5 |
| XM_028858722.1 | Slc2a6 | solute carrier family 2 member 6 (Slc2a6), transcript variant X1, mRNA | 29.55 | 9.6 | ≤13.5 |
| XM_028862698.1 | LOC114688093 | E3 ubiquitin-protein ligase RNF213-like (LOC114688093), mRNA | 3.66 | 9.5 | 7.9 |
| XM_028890979.1 | Mmp3 | matrix metallopeptidase 3 (Mmp3), mRNA | 390.30 | 9.5 | 11.2 |
| XM_028885944.1 | Tnfaip6 | TNF alpha induced protein 6 (Tnfaip6), mRNA | 199.26 | 9.4 | ≤13.5 |
| XM_028892100.1 | Irf9 | interferon regulatory factor 9 (Irf9), transcript variant X1, mRNA | 14.59 | 9.4 | 9.8 |
| XM_028880498.1 | Wnt5b | Wnt family member 5B (Wnt5b), transcript variant X2, mRNA | 7.25 | 9.4 | 3.6 |
| XM_028880432.1 | Bid | BH3 interacting domain death agonist (Bid), mRNA | 21.19 | 9.3 | 9.2 |
| XM_028856869.1 | Ifi35 | interferon induced protein 35 (Ifi35), mRNA | 84.93 | 9.3 | ≤13.5 |
| XM_028869610.1 | Irak3 | interleukin 1 receptor associated kinase 3 (Irak3), mRNA | 10.45 | 9.3 | ≤13.5 |
| XM_028884083.1 | Ccl19 | C-C motif chemokine ligand 19 (Ccl19), mRNA | 40.83 | 9.2 | 12.3 |
| XM_028855667.1 | LOC114681904 | 2'-5'-oligoadenylate synthase 1A-like (LOC114681904), partial mRNA | 3.75 | 9.1 | 2.1 |
| XM_028865550.1 | Ifi16 | interferon gamma inducible protein 16 (Ifi16), transcript variant X3, mRNA | 6.77 | 9.1 | 5.6 |
| XM_028879211.1 | Isg20 | interferon stimulated exonuclease gene 20 (Isg20), mRNA | 26.80 | 8.8 | ≤13.5 |
| XM_028875830.1 | Ampd3 | adenosine monophosphate deaminase 3 (Ampd3), transcript variant X3, mRNA | 5.08 | 8.5 | 4.3 |
| XM_028888791.1 | Tgtp2 | T-cell-specific guanine nucleotide triphosphate-binding protein 1-like (LOC | 11.20 | 8.3 | 9.0 |
| XM_028894546.1 | Cd274 | CD274 molecule (Cd274), mRNA | 3.66 | 8.2 | 5.6 |
| XM_028871629.1 | Ripk2 | receptor interacting serine/threonine kinase 2 (Ripk2), transcript variant X1, mRNA | 42.99 | 8.2 | ≤13.5 |
| XM_028861016.1 | Psmb8 | proteasome subunit beta 8 (Psmb8), mRNA | 104.44 | 8.1 | ≤13.5 |
| XM_028891472.1 | Tnfrsf14 | TNF receptor superfamily member 14 (Tnfrsf14), transcript variant X2, mRNA | 5.22 | 8.1 | 4.3 |
| XM_028862988.1 | Serpina3f | serine protease inhibitor A3F-like (LOC114688398), transcript variant X2, mRNA | 9.30 | 8.0 | 7.1 |
| XM_028862433.1 | Steap1 | STEAP family member 1 (Steap1), mRNA | 96.90 | 8.0 | ≤13.5 |
| XM_028863946.1 | Irak2 | interleukin 1 receptor associated kinase 2 (Irak2), transcript variant X3, mRNA | 2.18 | 7.8 | 4.2 |
| XM_028893872.1 | Relb | RELB proto-oncogene, NF-kB subunit (Relb), transcript variant X2, mRNA | 7.64 | 7.8 | 9.5 |
| XM_028891936.1 | Rnd1 | Rho family GTPase 1 (Rnd1), mRNA | 39.11 | 7.8 | 6.9 |
| XM_028871478.1 | Mt1 | metallothionein-1 (LOC114694510), mRNA | 454.08 | 7.7 | ≤13.5 |
| XM_028868449.1 | Sdc4 | syndecan 4 (Sdc4), mRNA | 1221.44 | 7.6 | ≤13.5 |
| XM_028876195.1 | LOC114697860 | nuclear autoantigen Sp-100-like (LOC114697860), transcript variant X1, mRNA | 16.25 | 7.5 | 5.6 |
| XM_028885276.1 | Helz2 | helicase with zinc finger 2 (Helz2), mRNA | 28.39 | 7.5 | ≤13.5 |
| XM_028891888.1 | Cacnb3 | calcium voltage-gated channel auxiliary subunit beta 3 (Cacnb3), transcript | 13.77 | 7.4 | ≤13.5 |
| XM_028893871.1 | Relb | RELB proto-oncogene, NF-kB subunit (Relb), transcript variant X1, mRNA | 60.94 | 7.3 | ≤13.5 |
| XM_028871803.1 | Sema6a | semaphorin 6A (Sema6a), mRNA | 12.58 | 7.3 | ≤13.5 |
| XM_028862653.1 | Dtx3l | deltex E3 ubiquitin ligase 3L (Dtx3l), mRNA | 51.25 | 7.3 | ≤13.5 |
| XM_028874705.1 | Nfkb2 | nuclear factor kappa B subunit 2 (Nfkb2), transcript variant X3, mRNA | 59.69 | 7.3 | ≤13.5 |
| XM_028870393.1 | Phf11 | PHD finger protein 11 (Phf11), transcript variant X1, mRNA | 28.50 | 7.2 | 12.8 |
| XM_028869786.1 | Samd9l | sterile alpha motif domain containing 9 like (Samd9l), transcript variant X1, mRNA | 3.81 | 7.1 | 8.0 |
| XM_028864149.1 | Snx10 | sorting nexin 10 (Snx10), transcript variant X1, mRNA | 1.44 | 7.1 | 2.9 |
| XM_028870394.1 | Phf11 | PHD finger protein 11 (Phf11), transcript variant X2, mRNA | 5.63 | 7.1 | 2.9 |
| XM_028881371.1 | Pim2 | Pim-2 proto-oncogene, serine/threonine kinase (Pim2), mRNA | 6.63 | 7.0 | 8.4 |
| XM_028862647.1 | Parp9 | poly(ADP-ribose) polymerase family member 9 (Parp9), transcript variant X1, mRNA | 36.56 | 7.0 | ≤13.5 |
| XM_028890716.1 | Nfkbie | NFKB inhibitor epsilon (Nfkbie), mRNA | 19.69 | 7.0 | ≤13.5 |
| XM_028871647.1 | Lhfpl2 | LHFPL tetraspan subfamily member 2 (Lhfpl2), transcript variant X1, mRNA | 16.15 | 7.0 | ≤13.5 |
| XM_028862159.1 | Cxcl16 | C-X-C motif chemokine ligand 16 (Cxcl16), mRNA | 36.92 | 7.0 | ≤13.5 |
| XM_028876205.1 | Sp100 | nuclear autoantigen Sp-100-like (LOC114697861), transcript variant X9, mRNA | 8.07 | 7.0 | 6.2 |
| XM_028869390.1 | Ntn1 | netrin 1 (Ntn1), transcript variant X1, mRNA | 11.67 | 6.9 | 6.7 |
| XM_028889227.1 | Kiaa1755 | KIAA1755 ortholog (Kiaa1755), mRNA | 3.82 | 6.9 | 5.7 |
| XM_028871372.1 | Hk2 | hexokinase 2 (Hk2), transcript variant X2, mRNA | 5.11 | 6.9 | 5.5 |
| XM_028865553.1 | LOC114690678 | interferon-activable protein 203-like (LOC114690678), mRNA | 4.19 | 6.8 | 2.9 |
| XM_028892101.1 | Irf9 | interferon regulatory factor 9 (Irf9), transcript variant X2, mRNA | 45.57 | 6.8 | ≤13.5 |
| XM_028887028.1 | Uba7 | ubiquitin like modifier activating enzyme 7 (Uba7), transcript variant X3, | 10.78 | 6.8 | 11.7 |
| XM_028886190.1 | Srgn | serglycin (Srgn), mRNA | 22.62 | 6.7 | 3.2 |
| XM_028878370.1 | Nmi | N-myc and STAT interactor (Nmi), mRNA | 39.54 | 6.6 | ≤13.5 |
| XM_028867490.1 | Nfkbiz | NFKB inhibitor zeta (Nfkbiz), transcript variant X2, mRNA | 14.12 | 6.6 | ≤13.5 |
| XM_028890165.1 | Vcam1 | vascular cell adhesion molecule 1 (Vcam1), mRNA | 100.86 | 6.5 | ≤13.5 |
| XM_028877590.1 | Slc39a14 | solute carrier family 39 member 14 (Slc39a14), transcript variant X2, mRNA | 13.66 | 6.4 | 10.3 |
| XM_028860761.1 | Tnfaip3 | TNF alpha induced protein 3 (Tnfaip3), transcript variant X2, mRNA | 19.91 | 6.3 | ≤13.5 |
| XM_028855439.1 | Stat2 | signal transducer and activator of transcription 2 (Stat2), mRNA | 52.16 | 6.2 | 11.4 |
| XM_028890122.1 | Iigp1b | interferon-inducible GTPase 1-like (LOC114707400), mRNA | 4.30 | 6.2 | 3.5 |
| XM_028868958.1 | Cacna1g | calcium voltage-gated channel subunit alpha1 G (Cacna1g), transcript variant X22, mRNA | 2.75 | 6.2 | 2.6 |
| XM_028890898.1 | Adar | adenosine deaminase RNA specific (Adar), transcript variant X1, mRNA | 8.96 | 6.2 | 3.2 |
| XM_028867757.1 | Hivep1 | human immunodeficiency virus type I enhancer binding protein 1 (Hivep1), transcript variant X3, mRNA | 2.08 | 6.1 | 2.2 |
| XM_028882785.1 | Trim14 | tripartite motif containing 14 (Trim14), mRNA | 4.24 | 6.1 | 2.8 |
| XM_028881169.1 | Ankrd37 | ankyrin repeat domain 37 (Ankrd37), transcript variant X2, mRNA | 11.89 | 6.1 | 4.3 |
| XM_028861017.1 | Psmb9 | proteasome subunit beta 9 (Psmb9), mRNA | 89.39 | 6.1 | ≤13.5 |
| XM_028869724.1 | Nfkbia | NFKB inhibitor alpha (Nfkbia), mRNA | 104.62 | 6.1 | ≤13.5 |
| XM_028887029.1 | Uba7 | ubiquitin like modifier activating enzyme 7 (Uba7), transcript variant X4, | 1.83 | 6.0 | 2.6 |
| XM_028861015.1 | Tap2 | transporter 2, ATP binding cassette subfamily B member (Tap2), mRNA | 119.06 | 6.0 | ≤13.5 |
| XM_028876199.1 | LOC114697861 | nuclear autoantigen Sp-100-like (LOC114697861), transcript variant X3, mRNA | 15.27 | 5.9 | 3.8 |
| XM_028872888.1 | Dexi | Dexi homolog (Dexi), mRNA | 30.99 | 5.8 | ≤13.5 |
| XM_028862479.1 | Steap2 | STEAP2 metalloreductase (Steap2), transcript variant X2, mRNA | 10.22 | 5.8 | 10.8 |
| XM_028876992.1 | LOC114698354 | cyclin-dependent kinase 4 inhibitor B (LOC114698354), mRNA | 63.55 | 5.8 | ≤13.5 |
| XM_028861014.1 | Tap1 | transporter 1, ATP binding cassette subfamily B member (Tap1), mRNA | 61.75 | 5.8 | ≤13.5 |
| XM_028868933.1 | Abcc3 | ATP binding cassette subfamily C member 3 (Abcc3), mRNA | 6.92 | 5.7 | 11.8 |
| XM_028858281.1 | Vegfc | vascular endothelial growth factor C (Vegfc), mRNA | 7.28 | 5.7 | 5.0 |
| XM_028859927.1 | Angptl4 | angiopoietin like 4 (Angptl4), mRNA | 467.01 | 5.7 | ≤13.5 |
| XM_028861134.1 | Rab32 | RAB32, member RAS oncogene family (Rab32), mRNA | 48.31 | 5.7 | ≤13.5 |
| XM_028875975.1 | Dio2 | iodothyronine deiodinase 2 (Dio2), transcript variant X1, mRNA | 14.21 | 5.7 | 3.5 |
| XM_028877491.1 | Dgat2 | diacylglycerol O-acyltransferase 2 (Dgat2), transcript variant X1, mRNA | 23.43 | 5.7 | ≤13.5 |
| XM_028871464.1 | Nlrc5 | NLR family CARD domain containing 5 (Nlrc5), mRNA | 1.68 | 5.6 | 4.0 |
| XM_028887027.1 | Uba7 | ubiquitin like modifier activating enzyme 7 (Uba7), transcript variant X2, | 2.56 | 5.6 | 2.8 |
| XM_028861570.1 | LOC114686901 | tripartite motif-containing protein 34A-like (LOC114686901), mRNA | 19.30 | 5.5 | 10.8 |
| XM_028886652.1 | Gpr68 | G protein-coupled receptor 68 (Gpr68), transcript variant X2, mRNA | 7.34 | 5.4 | 6.2 |
| XM_028869381.1 | Gas7 | growth arrest specific 7 (Gas7), transcript variant X3, mRNA | 6.55 | 5.4 | 3.5 |
| XM_028890959.1 | LOC114708004 | baculoviral IAP repeat-containing protein 3-like (LOC114708004), mRNA | 29.56 | 5.4 | ≤13.5 |
| XM_028858184.1 | Hpx | hemopexin (Hpx), mRNA | 12.12 | 5.4 | 6.8 |
| XM_028876194.1 | Sp140 | SP140 nuclear body protein (Sp140), mRNA | 30.26 | 5.2 | ≤13.5 |
| XM_028875599.1 | Osmr | oncostatin M receptor (Osmr), transcript variant X1, mRNA | 100.25 | 5.2 | ≤13.5 |
| XM_028877781.1 | Chst8 | carbohydrate sulfotransferase 8 (Chst8), transcript variant X1, mRNA | 4.37 | 5.2 | 2.3 |
| XM_028877925.1 | Slc1a2 | solute carrier family 1 member 2 (Slc1a2), transcript variant X3, mRNA | 3.85 | 5.2 | 11.7 |
| XM_028876193.1 | Sp110 | SP110 nuclear body protein (Sp110), transcript variant X2, mRNA | 3.24 | 5.2 | 2.2 |
| XM_028871162.1 | Sun2 | Sad1 and UNC84 domain containing 2 (Sun2), transcript variant X3, mRNA | 4.46 | 5.1 | 2.2 |
| XM_028888719.1 | Xaf1 | XIAP associated factor 1 (Xaf1), transcript variant X1, mRNA | 31.27 | 5.1 | 12.5 |
| XM_028889081.1 | Trim21 | tripartite motif containing 21 (Trim21), mRNA | 19.71 | 5.0 | ≤13.5 |
| XM_028869035.1 | Nfe2l1 | nuclear factor, erythroid 2 like 1 (Nfe2l1), transcript variant X1, mRNA | 35.00 | 5.0 | 7.3 |
| XM_028857437.1 | Slc39a8 | solute carrier family 39 member 8 (Slc39a8), transcript variant X2, mRNA | 4.38 | 4.9 | 2.7 |
| XM_028887655.1 | Nkain1 | sodium/potassium transporting ATPase interacting 1 (Nkain1), transcript var | 38.40 | 4.9 | ≤13.5 |
| XM_028875600.1 | Osmr | oncostatin M receptor (Osmr), transcript variant X2, mRNA | 11.83 | 4.9 | 6.7 |
| XM_028861273.1 | Stx11 | syntaxin 11 (Stx11), mRNA | 8.11 | 4.9 | 5.1 |
| XM_028880029.1 | Tnip1 | TNFAIP3 interacting protein 1 (Tnip1), transcript variant X7, mRNA | 14.59 | 4.8 | 12.3 |
| XM_028871052.1 | LOC114694211 | apolipoprotein L3-like (LOC114694211), mRNA | 23.11 | 4.8 | 7.2 |
| XM_028870923.1 | Parp10 | poly(ADP-ribose) polymerase family member 10 (Parp10), transcript variant X1, mRNA | 33.49 | 4.8 | ≤13.5 |
| XM_028887654.1 | Nkain1 | sodium/potassium transporting ATPase interacting 1 (Nkain1), transcript var | 3.32 | 4.8 | 2.8 |
| XM_028872422.1 | Col5a3 | collagen type V alpha 3 chain (Col5a3), mRNA | 19.93 | 4.8 | 9.7 |
| XM_028868448.1 | Rbpjl | recombination signal binding protein for immunoglobulin kappa J region like (Rbpjl), mRNA | 8.47 | 4.8 | 3.5 |
| XM_028889718.1 | Rhbdf2 | rhomboid 5 homolog 2 (Rhbdf2), mRNA | 21.13 | 4.7 | ≤13.5 |

Table S19

| | | | | | |
|---|---|---|---|---|---|
| XM_028885253.1 | Esm1 | endothelial cell specific molecule 1 (Esm1), mRNA | 11.02 | 4.7 | 5.0 |
| XM_028880373.1 | Mlph | melanophilin (Mlph), transcript variant X2, mRNA | 2.67 | 4.7 | 4.9 |
| XM_028886666.1 | Stat1 | signal transducer and activator of transcription 1 (Stat1), mRNA | 57.85 | 4.7 | 10.3 |
| XM_028882074.1 | Bdkrb1 | bradykinin receptor B1 (Bdkrb1), mRNA | 44.96 | 4.7 | 11.7 |
| XM_028864517.1 | LOC114690079 | 25-hydroxycholesterol 7-alpha-hydroxylase (LOC114690079), transcript variant X1, mRNA | 51.49 | 4.6 | 10.3 |
| XM_028869961.1 | Nck1 | NCK adaptor protein 1 (Nck1), transcript variant X6, mRNA | 3.29 | 4.6 | 7.0 |
| XR_003735254.1 | LOC114697860 | nuclear autoantigen Sp-100-like (LOC114697860), transcript variant X2, misc | 2.91 | 4.6 | 3.0 |
| XM_028857418.1 | Nfkb1 | nuclear factor kappa B subunit 1 (Nfkb1), mRNA | 62.55 | 4.5 | ≤13.5 |
| XM_028876192.1 | Sp110 | SP110 nuclear body protein (Sp110), transcript variant X1, mRNA | 74.25 | 4.5 | 10.3 |
| XM_028862991.1 | Serpina3n | serine protease inhibitor A3N-like (LOC114688401), mRNA | 14.17 | 4.4 | 4.9 |
| XM_028867009.1 | LOC114691567 | breakpoint cluster region protein-like (LOC114691567), mRNA | 11.90 | 4.4 | 4.6 |
| XM_028872354.1 | Slc7a2 | solute carrier family 7 member 2 (Slc7a2), mRNA | 147.36 | 4.3 | 12.2 |
| XM_028871053.1 | Apol8 | apolipoprotein L3-like (LOC114694212), transcript variant X1, mRNA [low sequence identity; just as much like other ApoL proteins; not L3 in the blastp list] | 7.02 | 4.3 | 2.9 |
| XM_028883006.1 | LOC114702542 | paralemmin-2 (LOC114702542), transcript variant X3, mRNA | 2.73 | 4.3 | 3.9 |
| XM_028863944.1 | Irak2 | interleukin 1 receptor associated kinase 2 (Irak2), transcript variant X1, mRNA | 23.94 | 4.2 | 10.4 |
| XM_028880795.1 | Gse1 | Gse1 coiled-coil protein (Gse1), transcript variant X3, mRNA | 3.76 | 4.2 | 5.4 |
| XM_028877203.1 | Rnf122 | ring finger protein 122 (Rnf122), transcript variant X2, mRNA | 5.43 | 4.2 | 2.7 |
| XM_028871588.1 | Gem | GTP binding protein overexpressed in skeletal muscle (Gem), mRNA | 18.28 | 4.1 | 5.9 |
| XM_028866128.1 | Tmem140 | transmembrane protein 140 (Tmem140), transcript variant X2, mRNA | 6.79 | 4.1 | 2.1 |
| XM_028861705.1 | Ier3 | immediate early response 3 (Ier3), mRNA | 161.94 | 4.1 | 12.8 |
| XM_028877200.1 | Rnf122 | ring finger protein 122 (Rnf122), transcript variant X1, mRNA | 23.49 | 4.0 | 6.3 |
| XM_028861203.1 | C3 | complement C3 (C3), mRNA | 102.56 | 4.0 | 4.8 |
| XM_028860146.1 | Tnfrsf1b | TNF receptor superfamily member 1B (Tnfrsf1b), mRNA | 44.26 | 4.0 | 12.8 |
| XM_028890700.1 | Enpp4 | ectonucleotide pyrophosphatase/phosphodiesterase 4 (Enpp4), transcript vari | 12.71 | 4.0 | 6.7 |
| XM_028885721.1 | Cflar | CASP8 and FADD like apoptosis regulator (Cflar), transcript variant X4, mRN | 6.35 | 4.0 | 2.1 |
| XM_028870924.1 | Parp10 | poly(ADP-ribose) polymerase family member 10 (Parp10), transcript variant X2, mRNA | 11.36 | 4.0 | 9.6 |
| XM_028890108.1 | Smim3 | small integral membrane protein 3 (Smim3), transcript variant X1, mRNA | 36.37 | 4.0 | 6.7 |
| XM_028877107.1 | Ifngr2 | interferon gamma receptor 2 (Ifngr2), transcript variant X2, mRNA | 27.20 | 3.9 | 8.7 |
| XM_028869700.1 | Egln3 | egl-9 family hypoxia inducible factor 3 (Egln3), mRNA | 23.62 | 3.9 | 8.5 |
| XM_028863195.1 | LOC114688648 | MICAL-like protein 2 (LOC114688648), mRNA | 5.66 | 3.9 | 4.2 |
| XM_028873528.1 | Pnp | purine nucleoside phosphorylase (Pnp), mRNA | 230.12 | 3.9 | 9.5 |
| XM_028886842.1 | Tmem173 | transmembrane protein 173 (Tmem173), mRNA | 70.91 | 3.8 | 10.7 |
| XM_028861559.1 | Trim30a | tripartite motif-containing protein 30A-like (LOC114686891), transcript variant X3, mRNA | 68.92 | 3.8 | 6.0 |
| XM_028855596.1 | Samhd1 | SAM and HD domain containing deoxynucleoside triphosphate triphosphohydrolase 1 (Samhd1), mRNA | 35.63 | 3.8 | 5.8 |
| XM_028889308.1 | Parp12 | poly(ADP-ribose) polymerase family member 12 (Parp12), mRNA | 18.12 | 3.8 | 8.8 |
| XM_028872602.1 | Skil | SKI like proto-oncogene (Skil), transcript variant X2, mRNA | 2.60 | 3.8 | 3.1 |
| XM_028878485.1 | Rbm43 | RNA binding motif protein 43 (Rbm43), mRNA | 11.32 | 3.8 | 7.4 |
| XM_028882412.1 | Arhgef3 | Rho guanine nucleotide exchange factor 3 (Arhgef3), transcript variant X3, | 4.03 | 3.8 | 3.5 |
| XM_028871371.1 | Hk2 | hexokinase 2 (Hk2), transcript variant X1, mRNA | 13.24 | 3.8 | 8.3 |
| XM_028894523.1 | Jak2 | Janus kinase 2 (Jak2), mRNA | 120.54 | 3.7 | 8.5 |
| XM_028877106.1 | Ifngr2 | interferon gamma receptor 2 (Ifngr2), transcript variant X1, mRNA | 42.29 | 3.7 | ≤13.5 |
| XM_028877801.1 | Gramd1a | GRAM domain containing 1A (Gramd1a), transcript variant X2, mRNA | 71.40 | 3.7 | 12.0 |
| XM_028854299.1 | Psmb10 | proteasome subunit beta 10 (Psmb10), mRNA | 63.91 | 3.7 | ≤13.5 |
| XM_028893122.1 | Tmbim1 | transmembrane BAX inhibitor motif containing 1 (Tmbim1), transcript variant | 9.70 | 3.7 | 3.7 |
| XM_028865714.1 | Mitd1 | microtubule interacting and trafficking domain containing 1 (Mitd1), transcript variant X2, mRNA | 8.08 | 3.6 | 2.4 |
| XM_028868955.1 | Cacna1g | calcium voltage-gated channel subunit alpha1 G (Cacna1g), transcript variant X19, mRNA | 11.96 | 3.6 | 10.8 |
| XM_028867374.1 | Lrrc49 | leucine rich repeat containing 49 (Lrrc49), transcript variant X6, mRNA | 4.90 | 3.6 | 3.4 |
| XM_028873276.1 | Eif2ak2 | eukaryotic translation initiation factor 2 alpha kinase 2 (Eif2ak2), mRNA | 34.47 | 3.6 | 7.4 |
| XM_028885543.1 | Mvp | major vault protein (Mvp), mRNA | 98.24 | 3.6 | 10.9 |
| XM_028863301.1 | H2-Q10 | H-2 class I histocompatibility antigen, Q10 alpha chain-like (LOC114688790), transcript variant X1, mRNA | 8.35 | 3.6 | 2.7 |
| XM_028858102.1 | Prkch | protein kinase C eta (Prkch), mRNA | 3.97 | 3.5 | 2.7 |
| XM_028892501.1 | Pml | promyelocytic leukemia (Pml), transcript variant X1, mRNA | 16.23 | 3.5 | 6.7 |
| XM_028861045.1 | Ginm1 | glycoprotein integral membrane 1 (Ginm1), transcript variant X2, mRNA | 164.59 | 3.5 | 10.0 |
| XM_028875970.1 | Adamts5 | ADAM metallopeptidase with thrombospondin type 1 motif 5 (Adamts5), mRNA | 14.33 | 3.5 | 6.1 |
| XM_028893881.1 | Bcl3 | BCL3 transcription coactivator (Bcl3), mRNA | 18.04 | 3.5 | 4.8 |
| XM_028861557.1 | LOC114686891 | tripartite motif-containing protein 30A-like (LOC114686891), transcript variant X1, mRNA | 59.53 | 3.5 | 8.0 |
| XM_028879416.1 | Csf1 | colony stimulating factor 1 (Csf1), transcript variant X1, mRNA | 192.28 | 3.5 | 5.9 |
| XM_028854998.1 | Sqstm1 | sequestosome 1 (Sqstm1), mRNA | 320.56 | 3.5 | 5.6 |
| XM_028888663.1 | Ptx3 | pentraxin 3 (Ptx3), mRNA | 246.45 | 3.4 | 4.7 |
| XM_028866788.1 | Abr | ABR activator of RhoGEF and GTPase (Abr), transcript variant X1, mRNA | 12.76 | 3.4 | 5.3 |
| XM_028867535.1 | Dcbld2 | discoidin, CUB and LCCL domain containing 2 (Dcbld2), transcript variant X1, mRNA | 11.62 | 3.4 | 7.9 |
| XM_028888763.1 | Tgtp2 | T-cell-specific guanine nucleotide triphosphate-binding protein 1-like (LOC | 13.22 | 3.4 | 5.0 |
| XM_028891357.1 | Pfkfb3 | 6-phosphofructo-2-kinase/fructose-2,6-biphosphatase 3 (Pfkfb3), transcript | 9.48 | 3.4 | 6.9 |
| XM_028876449.1 | LOC114698011 | TRPM8 channel-associated factor 2-like (LOC114698011), mRNA | 4.04 | 3.4 | 3.3 |
| XM_028888767.1 | LOC114706382 | uncharacterized LOC114706382 (LOC114706382), mRNA | 1.66 | 3.4 | 2.4 |
| XM_028892347.1 | Tapbpl | TAP binding protein like (Tapbpl), mRNA | 72.12 | 3.3 | 9.5 |
| XM_028889615.1 | Sphk1 | sphingosine kinase 1 (Sphk1), transcript variant X4, mRNA | 13.35 | 3.3 | 3.8 |
| XM_028870483.1 | Nebl | nebulette (Nebl), transcript variant X2, mRNA | 5.42 | 3.3 | 3.9 |
| XM_028891804.1 | Camk1d | calcium/calmodulin dependent protein kinase ID (Camk1d), transcript variant | 3.20 | 3.3 | 5.1 |
| XM_028876501.1 | Ikbke | inhibitor of nuclear factor kappa B kinase subunit epsilon (Ikbke), mRNA | 33.70 | 3.3 | 12.4 |
| XM_028879417.1 | Csf1 | colony stimulating factor 1 (Csf1), transcript variant X2, mRNA | 2.84 | 3.3 | 2.4 |
| XM_028880765.1 | Gng12 | G protein subunit gamma 12 (Gng12), mRNA | 115.08 | 3.2 | 7.4 |
| XM_028855365.1 | Gbp1 | guanylate-binding protein 1-like (LOC114681710), transcript variant X2, mRNA | 13.15 | 3.2 | 6.2 |
| XM_028886751.1 | Cystm1 | cysteine rich transmembrane module containing 1 (Cystm1), transcript varian | 46.19 | 3.2 | 2.6 |
| XM_028862989.1 | Serpina3f | serine protease inhibitor A3F-like (LOC114688398), transcript variant X3, mRNA | 18.39 | 3.2 | 2.9 |
| XM_028874704.1 | Nfkb2 | nuclear factor kappa B subunit 2 (Nfkb2), transcript variant X2, mRNA | 43.09 | 3.2 | 6.5 |
| XM_028889222.1 | Tgm2 | transglutaminase 2 (Tgm2), mRNA | 34.37 | 3.2 | 5.2 |
| XM_028893120.1 | Tmbim1 | transmembrane BAX inhibitor motif containing 1 (Tmbim1), transcript variant | 218.33 | 3.1 | 7.4 |
| XM_028877588.1 | Slc39a14 | solute carrier family 39 member 14 (Slc39a14), transcript variant X1, mRNA | 65.15 | 3.1 | 5.7 |
| XM_028867536.1 | Dcbld2 | discoidin, CUB and LCCL domain containing 2 (Dcbld2), transcript variant X2, mRNA | 5.42 | 3.1 | 4.2 |
| XM_028865713.1 | Mitd1 | microtubule interacting and trafficking domain containing 1 (Mitd1), transcript variant X1, mRNA | 22.11 | 3.1 | 4.0 |
| XM_028862557.1 | H2-Q10 | H-2 class I histocompatibility antigen, Q10 alpha chain-like (LOC114687933), transcript variant X4, mRNA | 14.07 | 3.1 | 3.7 |
| XM_028866374.1 | Bcl9l | BCL9 like (Bcl9l), transcript variant X2, mRNA | 5.41 | 3.1 | 5.3 |
| XM_028859831.1 | B3galnt2 | beta-1,3-N-acetylgalactosaminyltransferase 2 (B3galnt2), mRNA | 23.30 | 3.1 | 10.5 |
| XM_028883005.1 | LOC114702542 | paralemmin-2 (LOC114702542), transcript variant X2, mRNA | 5.34 | 3.1 | 4.6 |
| XM_028879527.1 | Nub1 | negative regulator of ubiquitin like proteins 1 (Nub1), transcript variant | 29.88 | 3.1 | 8.6 |
| XM_028886019.1 | Dennd3 | DENN domain containing 3 (Dennd3), transcript variant X1, mRNA | 8.12 | 3.1 | 4.5 |
| XM_028890900.1 | Adar | adenosine deaminase RNA specific (Adar), transcript variant X3, mRNA | 12.22 | 3.0 | 5.5 |
| XM_028892339.1 | C1s | complement C1s (C1s), mRNA | 321.93 | 3.0 | 4.5 |
| XM_028861732.1 | H2-T23 | H-2 class I histocompatibility antigen, D-37 alpha chain-like (LOC114687019), mRNA | 175.45 | 3.0 | 6.8 |
| XM_028879528.1 | Nub1 | negative regulator of ubiquitin like proteins 1 (Nub1), transcript variant | 47.76 | 3.0 | 6.3 |
| XM_028874727.1 | Ptgs2 | prostaglandin-endoperoxide synthase 2 (Ptgs2), mRNA | 75.70 | 3.0 | 5.6 |
| XM_028861561.1 | LOC114686891 | tripartite motif-containing protein 30A-like (LOC114686891), transcript variant X5, mRNA | 15.70 | 3.0 | 3.3 |
| XM_028861049.1 | H2-Q10 | H-2 class I histocompatibility antigen, Q10 alpha chain-like (LOC114686366), transcript variant X16, mRNA | 13.97 | 3.0 | 2.1 |
| XM_028861531.1 | LOC114686853 | caspase-1-like (LOC114686853), mRNA | 15.23 | 3.0 | 4.7 |
| XM_028880797.1 | Gse1 | Gse1 coiled-coil protein (Gse1), transcript variant X4, mRNA | 3.33 | 3.0 | 3.0 |
| XM_028884825.1 | Mast4 | microtubule associated serine/threonine kinase family member 4 (Mast4), tra | 6.38 | 3.0 | 6.1 |
| XM_028856845.1 | LOC114682811 | signal transducer and activator of transcription 5A (LOC114682811), mRNA | 43.09 | 2.9 | 8.0 |
| XM_028879418.1 | Csf1 | colony stimulating factor 1 (Csf1), transcript variant X3, mRNA | 11.05 | 2.9 | 3.4 |
| XM_028867162.1 | Mocos | molybdenum cofactor sulfurase (Mocos), mRNA | 10.16 | 2.9 | 4.3 |
| XM_028866790.1 | Abr | ABR activator of RhoGEF and GTPase (Abr), transcript variant X2, mRNA | 9.66 | 2.9 | 2.5 |
| XM_028877703.1 | Adam17 | ADAM metallopeptidase domain 17 (Adam17), transcript variant X1, mRNA | 70.76 | 2.9 | 7.0 |
| XM_028868561.1 | Znfx1 | zinc finger NFX1-type containing 1 (Znfx1), transcript variant X4, mRNA | 25.49 | 2.9 | 4.3 |
| XM_028861528.1 | LOC114686852 | caspase-12-like (LOC114686852), transcript variant X1, mRNA | 18.99 | 2.9 | 5.5 |
| XM_028868770.1 | Traf2 | TNF receptor associated factor 2 (Traf2), mRNA | 46.21 | 2.9 | 10.9 |
| XM_028861328.1 | Nupr1 | nuclear protein 1, transcriptional regulator (Nupr1), mRNA | 405.53 | 2.9 | 4.4 |
| XM_028886462.1 | Hivep3 | human immunodeficiency virus type I enhancer binding protein 3 (Hivep3), tr | 1.44 | 2.9 | 2.4 |
| XM_028891396.1 | Pfkfb3 | 6-phosphofructo-2-kinase/fructose-2,6-biphosphatase 3 (Pfkfb3), transcript | 36.07 | 2.8 | 6.0 |
| XM_028887322.1 | Il13ra1 | interleukin 13 receptor subunit alpha 1 (Il13ra1), mRNA | 24.01 | 2.8 | 5.5 |
| XM_028861033.1 | H2-Q10 | H-2 class I histocompatibility antigen, Q10 alpha chain-like (LOC114686366), transcript variant X2, mRNA | 16.64 | 2.8 | 3.9 |
| XM_028856571.1 | Alpk1 | alpha kinase 1 (Alpk1), mRNA | 4.58 | 2.8 | 2.6 |
| XM_028879990.1 | Pdlim4 | PDZ and LIM domain 4 (Pdlim4), transcript variant X1, mRNA | 117.08 | 2.8 | 7.4 |
| XM_028857055.1 | LOC114682933 | PTB-containing, cubilin and LRP1-interacting protein (LOC114682933), transcript variant X1, mRNA | 39.37 | 2.8 | 4.3 |
| XM_028864204.1 | Insig1 | insulin induced gene 1 (Insig1), transcript variant X1, mRNA | 24.46 | 2.8 | 5.3 |
| XM_028877920.1 | Cd44 | CD44 molecule (Indian blood group) (Cd44), transcript variant X1, mRNA | 597.91 | 2.8 | 5.2 |
| XM_028867672.1 | Rnf144b | ring finger protein 144B (Rnf144b), transcript variant X1, mRNA | 14.84 | 2.8 | 4.5 |
| XM_028877459.1 | Ttc9 | tetratricopeptide repeat domain 9 (Ttc9), mRNA | 6.73 | 2.8 | 2.2 |
| XM_028892341.1 | C1r | complement C1r (C1r), mRNA | 273.81 | 2.8 | 4.8 |
| XM_028877921.1 | Cd44 | CD44 molecule (Indian blood group) (Cd44), transcript variant X2, mRNA | 25.41 | 2.7 | 3.8 |
| XM_028856234.1 | Ciart | circadian associated repressor of transcription (Ciart), mRNA | 10.28 | 2.7 | 3.0 |
| XM_028869784.1 | Samd9 | sterile alpha motif domain containing 9 (Samd9), transcript variant X1, mRNA | 26.73 | 2.7 | 3.5 |
| XM_028871898.1 | Zhx2 | zinc fingers and homeoboxes 2 (Zhx2), transcript variant X1, mRNA | 9.81 | 2.7 | 5.2 |
| XM_028855812.1 | Rnf19b | ring finger protein 19B (Rnf19b), mRNA | 53.28 | 2.7 | 5.5 |
| XM_028858724.1 | Slc2a6 | solute carrier family 2 member 6 (Slc2a6), transcript variant X3, mRNA | 15.41 | 2.7 | 3.7 |
| XM_028888277.1 | LOC114706055 | interferon alpha-inducible protein 27, mitochondrial-like (LOC114706055), t | 34.39 | 2.7 | 2.4 |
| XM_028857047.1 | Fbrsl1 | fibrosin like 1 (Fbrsl1), transcript variant X9, mRNA | 4.53 | 2.7 | 2.3 |
| XM_028894046.1 | Tjp2 | tight junction protein 2 (Tjp2), mRNA | 17.93 | 2.7 | 5.4 |
| XM_028890776.1 | Dap3 | death associated protein 3 (Dap3), transcript variant X3, mRNA | 13.73 | 2.7 | 2.9 |
| XM_028886497.1 | Slc2a1 | solute carrier family 2 member 1 (Slc2a1), mRNA | 108.28 | 2.7 | 5.1 |
| XM_028891336.1 | Pfkfb3 | 6-phosphofructo-2-kinase/fructose-2,6-biphosphatase 3 (Pfkfb3), transcript | 7.69 | 2.7 | 4.9 |
| XM_028881359.1 | Nqo2 | N-ribosyldihydronicotinamide:quinone reductase 2 (Nqo2), mRNA | 19.37 | 2.6 | 6.0 |
| XM_028891860.1 | Optn | optineurin (Optn), transcript variant X2, mRNA | 23.98 | 2.6 | 5.2 |
| XM_028880660.1 | Pdpn | podoplanin (Pdpn), transcript variant X1, mRNA | 60.61 | 2.6 | 5.0 |
| XM_028881507.1 | LOC114701420 | C-type lectin domain family 2 member D11-like (LOC114701420), mRNA | 60.87 | 2.6 | 4.3 |
| XM_028878605.1 | Smox | spermine oxidase (Smox), transcript variant X1, mRNA | 12.72 | 2.6 | 4.1 |
| XM_028892349.1 | C1rl | complement C1r subcomponent like (C1rl), mRNA | 16.39 | 2.6 | 3.0 |
| XM_028857430.1 | Slc39a8 | solute carrier family 39 member 8 (Slc39a8), transcript variant X1, mRNA | 49.43 | 2.6 | 2.5 |
| XM_028884433.1 | Il6st | interleukin 6 signal transducer (Il6st), mRNA | 203.63 | 2.6 | 3.3 |

Table S19

| | | | | | |
|---|---|---|---|---|---|
| XM_028875057.1 | Mxi1 | MAX interactor 1, dimerization protein (Mxi1), transcript variant X1, mRNA | 19.19 | 2.6 | 5.0 |
| XM_028884941.1 | Timp1 | TIMP metallopeptidase inhibitor 1 (Timp1), transcript variant X1, mRNA | 974.00 | 2.6 | 3.1 |
| XM_028875358.1 | Ece1 | endothelin converting enzyme 1 (Ece1), transcript variant X2, mRNA | 19.99 | 2.6 | 5.2 |
| XM_028862833.1 | Adgre5 | adhesion G protein-coupled receptor E5 (Adgre5), transcript variant X2, mRNA | 35.00 | 2.6 | 2.8 |
| XM_028861565.1 | LOC114686895 | tripartite motif-containing protein 30A-like (LOC114686895), mRNA | 22.22 | 2.6 | 7.1 |
| XM_028866905.1 | Lgals9 | galectin 9 (Lgals9), transcript variant X1, mRNA | 33.92 | 2.6 | 3.0 |
| XM_028884949.1 | Timp1 | TIMP metallopeptidase inhibitor 1 (Timp1), transcript variant X2, mRNA | 68.31 | 2.6 | 2.7 |
| XM_028867654.1 | Mrps25 | mitochondrial ribosomal protein S25 (Mrps25), transcript variant X1, mRNA | 73.12 | 2.5 | 4.6 |
| XM_028862207.1 | Abcb1 | ATP binding cassette subfamily B member 1 (Abcb1), transcript variant X2, mRNA | 6.35 | 2.5 | 2.1 |
| XM_028861733.1 | H2-Q10 | H-2 class I histocompatibility antigen, Q10 alpha chain-like (LOC114687020), mRNA | 17.35 | 2.5 | 3.0 |
| XM_028892105.1 | Psme1 | proteasome activator subunit 1 (Psme1), transcript variant X2, mRNA | 143.92 | 2.5 | 5.5 |
| XM_028892776.1 | Znrf1 | zinc and ring finger 1 (Znrf1), transcript variant X2, mRNA | 9.98 | 2.5 | 2.4 |
| XM_028875357.1 | Ece1 | endothelin converting enzyme 1 (Ece1), transcript variant X1, mRNA | 7.24 | 2.5 | 2.4 |
| XM_028858457.1 | H6pd | hexose-6-phosphate dehydrogenase/glucose 1-dehydrogenase (H6pd), transcript variant X2, mRNA | 22.31 | 2.5 | 5.0 |
| XM_028872603.1 | Skil | SKI like proto-oncogene (Skil), transcript variant X3, mRNA | 6.62 | 2.5 | 2.6 |
| XM_028872438.1 | Tyk2 | tyrosine kinase 2 (Tyk2), transcript variant X2, mRNA | 4.41 | 2.5 | 2.3 |
| XM_028870888.1 | LOC114694150 | gasdermin-D-like (LOC114694150), mRNA | 33.80 | 2.5 | 3.6 |
| XM_028866759.1 | Slc43a2 | solute carrier family 43 member 2 (Slc43a2), transcript variant X6, mRNA | 21.58 | 2.4 | 5.2 |
| XM_028889309.1 | Zc3hav1 | zinc finger CCCH-type containing, antiviral 1 (Zc3hav1), mRNA | 49.93 | 2.4 | 3.9 |
| XM_028871813.1 | Tnfaip8 | TNF alpha induced protein 8 (Tnfaip8), transcript variant X5, mRNA | 23.28 | 2.4 | 3.8 |
| XM_028868557.1 | Znfx1 | zinc finger NFX1-type containing 1 (Znfx1), transcript variant X1, mRNA | 7.03 | 2.4 | 3.1 |
| XM_028887606.1 | Marcksl1 | MARCKS like 1 (Marcksl1), mRNA | 56.63 | 2.4 | 5.5 |
| XM_028866786.1 | Nxn | nucleoredoxin (Nxn), mRNA | 48.40 | 2.4 | 3.7 |
| XM_028886628.1 | Myo1b | myosin IB (Myo1b), transcript variant X5, mRNA | 13.78 | 2.4 | 3.9 |
| XM_028892796.1 | Rhoq | ras homolog family member Q (Rhoq), mRNA | 40.61 | 2.4 | 6.7 |
| XM_028867165.1 | Slc39a6 | solute carrier family 39 member 6 (Slc39a6), mRNA | 72.37 | 2.4 | 4.2 |
| XM_028871818.1 | Lox | lysyl oxidase (Lox), transcript variant X1, mRNA | 166.42 | 2.4 | 3.9 |
| XM_028869871.1 | Tfpi2 | tissue factor pathway inhibitor 2 (Tfpi2), mRNA | 48.35 | 2.4 | 2.6 |
| XM_028857764.1 | LOC114683452 | ATP-binding cassette sub-family A member 1-like (LOC114683452), mRNA | 4.50 | 2.4 | 2.1 |
| XM_028865755.1 | Npas2 | neuronal PAS domain protein 2 (Npas2), transcript variant X3, mRNA | 13.11 | 2.4 | 5.2 |
| XM_028881946.1 | Ccng2 | cyclin G2 (Ccng2), mRNA | 30.00 | 2.4 | 5.1 |
| XM_028876181.1 | Pid1 | phosphotyrosine interaction domain containing 1 (Pid1), transcript variant X2, mRNA | 105.48 | 2.4 | 2.5 |
| XM_028891743.1 | Nampt | nicotinamide phosphoribosyltransferase (Nampt), mRNA | 57.21 | 2.3 | 4.0 |
| XM_028869881.1 | Ifitm3 | interferon-induced transmembrane protein 3 (LOC114693504), mRNA | 642.21 | 2.3 | 2.8 |
| XM_028891851.1 | Optn | optineurin (Optn), transcript variant X1, mRNA | 17.14 | 2.3 | 5.6 |
| XM_028886647.1 | Myo1b | myosin IB (Myo1b), transcript variant X7, mRNA | 18.33 | 2.3 | 4.1 |
| XM_028876604.1 | Btg2 | BTG anti-proliferation factor 2 (Btg2), mRNA | 51.70 | 2.3 | 4.2 |
| XM_028858071.1 | Hif1a | hypoxia inducible factor 1 subunit alpha (Hif1a), transcript variant X1, mRNA | 34.53 | 2.3 | 3.9 |
| XM_028863302.1 | H2-Q10 | H-2 class I histocompatibility antigen, Q10 alpha chain-like (LOC114688790), transcript variant X2, mRNA | 59.81 | 2.3 | 3.4 |
| XM_028870241.1 | Ak4 | adenylate kinase 4 (Ak4), mRNA | 37.08 | 2.3 | 2.9 |
| XM_028871819.1 | Lox | lysyl oxidase (Lox), transcript variant X2, mRNA | 82.12 | 2.3 | 4.3 |
| XM_028886602.1 | Myo1b | myosin IB (Myo1b), transcript variant X2, mRNA | 8.19 | 2.3 | 2.5 |
| XM_028867903.1 | Il4r | interleukin 4 receptor (Il4r), mRNA | 12.63 | 2.2 | 2.9 |
| XM_028875051.1 | Add3 | adducin 3 (Add3), transcript variant X1, mRNA | 8.92 | 2.2 | 3.4 |
| XM_028884444.1 | Itgav | integrin subunit alpha V (Itgav), transcript variant X2, mRNA | 19.81 | 2.2 | 3.2 |
| XM_028893220.1 | St3gal3 | ST3 beta-galactoside alpha-2,3-sialyltransferase 3 (St3gal3), transcript va | 12.45 | 2.2 | 2.6 |
| XM_028892106.1 | Psme2 | proteasome activator subunit 2 (Psme2), mRNA | 281.03 | 2.2 | 3.4 |
| XM_028874574.1 | Slc11a2 | solute carrier family 11 member 2 (Slc11a2), transcript variant X2, mRNA | 18.96 | 2.2 | 3.3 |
| XM_028878402.1 | Tsc22d1 | TSC22 domain family member 1 (Tsc22d1), transcript variant X6, mRNA | 246.99 | 2.2 | 2.4 |
| XM_028887991.1 | Usp24 | ubiquitin specific peptidase 24 (Usp24), transcript variant X1, mRNA | 8.39 | 2.2 | 3.6 |
| XM_028884443.1 | Itgav | integrin subunit alpha V (Itgav), transcript variant X1, mRNA | 45.77 | 2.2 | 2.4 |
| XM_028875628.1 | Arrdc3 | arrestin domain containing 3 (Arrdc3), transcript variant X1, mRNA | 17.11 | 2.2 | 4.9 |
| XM_028878205.1 | Trim25 | tripartite motif containing 25 (Trim25), mRNA | 44.19 | 2.1 | 3.0 |
| XM_028868578.1 | Cebpb | CCAAT enhancer binding protein beta (Cebpb), mRNA | 52.21 | 2.1 | 2.5 |
| XM_028882800.1 | Tdrd7 | tudor domain containing 7 (Tdrd7), mRNA | 15.31 | 2.1 | 3.5 |
| XM_028887804.1 | Plat | plasminogen activator, tissue type (Plat), mRNA | 37.64 | 2.1 | 2.7 |
| XM_028887958.1 | H2-L | H-2 class I histocompatibility antigen, L-D alpha chain-like (LOC114705846) | 62.89 | 2.1 | 3.2 |
| XM_028888386.1 | Tapbp | TAP binding protein (Tapbp), mRNA | 193.15 | 2.1 | 2.4 |
| XM_028894512.1 | Rcl1 | RNA terminal phosphate cyclase like 1 (Rcl1), mRNA | 25.42 | 2.1 | 2.1 |
| XM_028859505.1 | Shb | SH2 domain containing adaptor protein B (Shb), mRNA | 10.32 | 2.1 | 2.2 |
| XM_028872437.1 | Tyk2 | tyrosine kinase 2 (Tyk2), transcript variant X1, mRNA | 9.02 | 2.1 | 2.9 |
| XM_028885326.1 | Ogfr | opioid growth factor receptor (Ogfr), mRNA | 74.30 | 2.1 | 2.8 |
| XM_028881499.1 | Gabarapl1 | GABA type A receptor associated protein like 1 (Gabarapl1), mRNA | 159.61 | 2.1 | 2.6 |
| XM_028861618.1 | Sash1 | SAM and SH3 domain containing 1 (Sash1), transcript variant X8, mRNA | 8.98 | 2.1 | 2.3 |
| XM_028859710.1 | Ticam1 | toll like receptor adaptor molecule 1 (Ticam1), mRNA | 15.47 | 2.1 | 3.8 |
| XM_028882204.1 | Trib1 | tribbles pseudokinase 1 (Trib1), mRNA | 20.81 | 2.1 | 2.7 |
| XM_028861050.1 | H2-Q10 | H-2 class I histocompatibility antigen, Q10 alpha chain-like (LOC114686366), transcript variant X17, mRNA | 44.79 | 2.1 | 2.3 |
| XM_028862909.1 | Btg1 | BTG anti-proliferation factor 1 (Btg1), mRNA | 72.75 | 2.0 | 2.5 |
| XM_028872601.1 | Skil | SKI like proto-oncogene (Skil), transcript variant X1, mRNA | 12.31 | 2.0 | 2.3 |
| XM_028860214.1 | Kdm3a | lysine demethylase 3A (Kdm3a), mRNA | 15.25 | 2.0 | 3.7 |
| XM_028893750.1 | Lmo4 | LIM domain only 4 (Lmo4), transcript variant X3, mRNA | 30.17 | 2.0 | 3.0 |
| XM_028894449.1 | Trim56 | tripartite motif containing 56 (Trim56), mRNA | 24.79 | 2.0 | 2.1 |
| XM_028865763.1 | Rnf149 | ring finger protein 149 (Rnf149), mRNA | 20.85 | 2.0 | 2.4 |
| XM_028867446.1 | Fgfrl1 | fibroblast growth factor receptor like 1 (Fgfrl1), mRNA | 62.54 | -2.0 | 3.2 |
| XM_028854353.1 | Mcm2 | minichromosome maintenance complex component 2 (Mcm2), mRNA | 21.63 | -2.0 | 2.4 |
| XM_028886325.1 | Mylk | myosin light chain kinase (Mylk), mRNA | 12.75 | -2.0 | 2.1 |
| XM_028854848.1 | Slc6a6 | solute carrier family 6 member 6 (Slc6a6), transcript variant X1, mRNA | 19.64 | -2.0 | 2.2 |
| XM_028883856.1 | Psd3 | pleckstrin and Sec7 domain containing 3 (Psd3), mRNA | 6.22 | -2.0 | 2.1 |
| XM_028885699.1 | Megf9 | multiple EGF like domains 9 (Megf9), mRNA | 16.88 | -2.1 | 3.1 |
| XM_028871489.1 | Gnao1 | G protein subunit alpha o1 (Gnao1), transcript variant X3, mRNA | 7.11 | -2.1 | 2.4 |
| XM_028881158.1 | Sorbs2 | sorbin and SH3 domain containing 2 (Sorbs2), transcript variant X19, mRNA | 12.79 | -2.1 | 2.2 |
| XM_028879075.1 | Arhgap32 | Rho GTPase activating protein 32 (Arhgap32), mRNA | 5.02 | -2.1 | 3.1 |
| XM_028882911.1 | Plce1 | phospholipase C epsilon 1 (Plce1), mRNA | 14.22 | -2.2 | 3.5 |
| XM_028870088.1 | Map2 | microtubule associated protein 2 (Map2), transcript variant X14, mRNA | 11.15 | -2.2 | 2.3 |
| XM_028867566.1 | LOC114691978 | inositol 1,4,5-trisphosphate receptor type 1 (LOC114691978), transcript variant X1, mRNA | 7.29 | -2.2 | 2.9 |
| XM_028867534.1 | Col8a1 | collagen type VIII alpha 1 chain (Col8a1), mRNA | 213.68 | -2.2 | 2.2 |
| XM_028877408.1 | Sipa1l1 | signal induced proliferation associated 1 like 1 (Sipa1l1), transcript variant X11, mRNA | 8.56 | -2.2 | 2.3 |
| XM_028867781.1 | Xylt1 | xylosyltransferase 1 (Xylt1), mRNA | 50.32 | -2.2 | 2.3 |
| XM_028867579.1 | LOC114691978 | inositol 1,4,5-trisphosphate receptor type 1 (LOC114691978), transcript variant X13, mRNA | 19.42 | -2.3 | 3.2 |
| XM_028884976.1 | Metrnl | meteorin like, glial cell differentiation regulator (Metrnl), mRNA | 24.22 | -2.3 | 2.7 |
| XM_028873388.1 | Pcdh18 | protocadherin 18 (Pcdh18), transcript variant X2, mRNA | 11.72 | -2.3 | 4.4 |
| XM_028881161.1 | Sorbs2 | sorbin and SH3 domain containing 2 (Sorbs2), transcript variant X22, mRNA | 10.33 | -2.4 | 3.4 |
| XM_028869258.1 | Slc2a13 | solute carrier family 2 member 13 (Slc2a13), mRNA | 4.54 | -2.4 | 2.2 |
| XM_028880793.1 | Gse1 | Gse1 coiled-coil protein (Gse1), transcript variant X1, mRNA | 5.41 | -2.4 | 3.7 |
| XM_028868003.1 | Ogfrl1 | opioid growth factor receptor like 1 (Ogfrl1), transcript variant X2, mRNA | 8.12 | -2.5 | 3.4 |
| XM_028861276.1 | Plagl1 | PLAG1 like zinc finger 1 (Plagl1), transcript variant X2, mRNA | 19.79 | -2.5 | 2.5 |
| XM_028873742.1 | Lbh | LBH regulator of WNT signaling pathway (Lbh), mRNA | 26.99 | -2.5 | 2.0 |
| XM_028888075.1 | Thbs1 | thrombospondin 1 (Thbs1), mRNA | 1364.09 | -2.5 | 2.5 |
| XM_028892992.1 | Susd5 | sushi domain containing 5 (Susd5), transcript variant X1, mRNA | 42.95 | -2.5 | 4.3 |
| XM_028894973.1 | Prss23 | serine protease 23 (Prss23), transcript variant X2, mRNA | 13.97 | -2.5 | 2.5 |
| XM_028877339.1 | Sipa1l1 | signal induced proliferation associated 1 like 1 (Sipa1l1), transcript variant X3, mRNA | 5.53 | -2.5 | 3.3 |
| XM_028873810.1 | Adamtsl3 | ADAMTS like 3 (Adamtsl3), transcript variant X2, mRNA | 6.38 | -2.5 | 5.0 |
| XM_028871913.1 | Deptor | DEP domain containing MTOR interacting protein (Deptor), mRNA | 24.03 | -2.6 | 3.3 |
| XM_028881160.1 | Sorbs2 | sorbin and SH3 domain containing 2 (Sorbs2), transcript variant X21, mRNA | 6.73 | -2.6 | 2.9 |
| XM_028890459.1 | Tmem158 | transmembrane protein 158 (Tmem158), mRNA | 5.29 | -2.6 | 2.4 |
| XM_028876759.1 | Kcnk2 | potassium two pore domain channel subfamily K member 2 (Kcnk2), transcript variant X3, mRNA | 3.57 | -2.8 | 2.1 |
| XM_028873007.1 | Cped1 | cadherin like and PC-esterase domain containing 1 (Cped1), transcript variant X4, mRNA | 9.01 | -2.9 | 3.9 |
| XM_028882012.1 | Cers6 | ceramide synthase 6 (Cers6), transcript variant X2, mRNA | 8.55 | -2.9 | 7.1 |
| XM_028878353.1 | Gdf10 | growth differentiation factor 10 (Gdf10), mRNA | 47.00 | -2.9 | 3.6 |
| XM_028875611.1 | Ranbp3l | RAN binding protein 3 like (Ranbp3l), mRNA | 13.48 | -3.0 | 5.0 |
| XM_028855107.1 | Adamts9 | ADAM metallopeptidase with thrombospondin type 1 motif 9 (Adamts9), mRNA | 8.72 | -3.1 | 3.4 |
| XM_028875980.1 | Jam2 | junctional adhesion molecule 2 (Jam2), mRNA | 9.15 | -3.2 | 2.3 |
| XM_028878990.1 | Tspan18 | tetraspanin 18 (Tspan18), transcript variant X1, mRNA | 5.64 | -3.4 | 2.8 |
| XM_028889275.1 | Epha4 | EPH receptor A4 (Epha4), mRNA | 9.26 | -3.5 | 2.5 |
| XM_028861297.1 | Plagl1 | PLAG1 like zinc finger 1 (Plagl1), transcript variant X5, mRNA | 3.38 | -3.5 | 2.1 |
| XM_028869779.1 | Adamts18 | ADAM metallopeptidase with thrombospondin type 1 motif 18 (Adamts18), transcript variant X1, mRNA | 6.96 | -3.6 | 2.1 |
| XM_028867958.1 | Rspo2 | R-spondin 2 (Rspo2), transcript variant X2, mRNA | 18.88 | -3.6 | 7.8 |
| XM_028891994.1 | Kiaa1217 | KIAA1217 ortholog (Kiaa1217), transcript variant X8, mRNA | 2.42 | -3.7 | 2.3 |
| XM_028876756.1 | Kcnk2 | potassium two pore domain channel subfamily K member 2 (Kcnk2), transcript variant X1, mRNA | 21.66 | -3.9 | 8.4 |
| XM_028882805.1 | Plxna2 | plexin A2 (Plxna2), mRNA | 13.63 | -4.0 | 10.2 |
| XM_028871222.1 | Tef | TEF transcription factor, PAR bZIP family member (Tef), transcript variant X2, mRNA | 3.09 | -4.0 | 3.2 |
| XM_028875674.1 | Thrb | thyroid hormone receptor beta (Thrb), transcript variant X1, mRNA | 3.09 | -4.1 | 2.4 |
| XM_028876760.1 | Kcnk2 | potassium two pore domain channel subfamily K member 2 (Kcnk2), transcript variant X4, mRNA | 10.53 | -4.1 | 7.0 |
| XM_028892000.1 | Kiaa1217 | KIAA1217 ortholog (Kiaa1217), transcript variant X13, mRNA | 2.42 | -4.3 | 2.5 |
| XM_028869352.1 | Myocd | myocardin (Myocd), transcript variant X2, mRNA | 1.78 | -4.4 | 3.0 |
| XM_028878029.1 | Rasl11a | RAS like family 11 member A (Rasl11a), mRNA | 6.67 | -4.4 | 2.0 |
| XM_028873005.1 | Cped1 | cadherin like and PC-esterase domain containing 1 (Cped1), transcript variant X2, mRNA | 2.20 | -4.6 | 2.5 |
| XM_028867957.1 | Rspo2 | R-spondin 2 (Rspo2), transcript variant X1, mRNA | 8.62 | -4.7 | 7.3 |
| XM_028872057.1 | Phldb2 | pleckstrin homology like domain family B member 2 (Phldb2), transcript variant X14, mRNA | 1.26 | -5.3 | 2.5 |
| XM_028865057.1 | Ptpn9 | protein tyrosine phosphatase non-receptor type 9 (Ptpn9), transcript variant X2, mRNA | 1.90 | -28.7 | 5.2 |
| XM_028893827.1 | Tomm40 | translocase of outer mitochondrial membrane 40 (Tomm40), transcript variant | 3.41 | -178.4 | 4.9 |

Table S20. Accession numbers for limited set of protein coding sequences of mRNAs

| gene | product | P. leucopus accession | M. musculus accession |
|---|---|---|---|
| Acod1 | aconitate decarboxylase 1 | XM_028864652.1 | NM_008392.1 |
| Alb | albumin | XM_028887861.1 | NM_009654.4 |
| Alox5 | arachidonate 5-lipoxygenase | XM_028864037.1 | NM_009662.2 |
| Arg1 | arginase 1 | XM_028877403.1 | NM_007482.3 |
| Ccl2 | C-C motif ligand chemokine 2 | XM_028866979.1 | NM_011333.3 |
| Cd86 | CD86 antigen | XM_028862269.1 | NM_019388.3 |
| Crp | C-reactive protein | XM_028865632.1 | NM_007768.4 |
| Csf2 | colony stimulating factor 2 (granulocyte-macrophage) | XM_028879989.1 | NM_009969.4 |
| Csf3 | colony stimulating factor 3 (granulocyte) | XM_028889425.1 | NM_009971.1 |
| Cxcl11 | C-X-C motif chemokine ligand chemokine 11 | XM_028881923.1 | NR_038116.1 |
| Fth1 | ferritin heavy polypeptide 1 | XM_028860338.1 | NM_010239.2 |
| Gapdh | glyceraldehyde-3-phosphate dehydrogenase | XM_028892294.1 | NM_008084.3 |
| Gata3 | GATA binding protein 3 | XM_028891561.1 | XM_030247532.1 |
| Hmox1 | heme oxygenase 1 | XM_028854887.1 | NM_010442.2 |
| Hp | haptoglobin | XM_028891075.1 | NM_017370.2 |
| Ibsp | integrin binding sialoprotein | XM_028867455.1 | NM_008318.3: |
| Ido1 | indoleamine 2,3-dioxygenase 1 | XM_028887820.1 | NM_008324.2 |
| Ifnb1 | interferon beta 1 | XM_028859756.1 | NM_010510.1 |
| Ifng | interferon gamma | XM_028869658.1 | NM_008337.4 |
| Il10 | interleukin 10 | XM_028876480.1 | NM_010548.2 |
| Il12 | interleukin 12 | XM_028867243.1 | NM_001159424.2 |
| Il13 | interleukin 13 | XM_006995904.1 | NM_008355.3 |
| Il17a | interleukin 17A | XM_028868119.1 | NM_010552.3 |
| Il1a | interleukin 1 alpha | XM_028893405.1 | NM_010554.4 |
| Il1b | interleukin 1 beta | XM_028893401.1 | NM_008361.4 |
| Il1r | interleukin 1 receptor | XM_028865796.1 | XM_006495713.4 |
| Il1rn | interleukin 1 receptor antagonist | XM_028868677.1 | NM_001039701.3 |
| Il22 | interleukin 22 | XM_028869659.1 | NM_016971.2 |
| Il4 | interleukin 4 | XM_028855689.1 | NM_021283.2 |
| Il5 | interleukin 5 | XM_015989306.1 | NM_010558.1 |
| Il6 | interleukin 6 | XM_028864183.1 | NM_031168.2 |
| Irf5 | interferon regulatory factor 5 | XM_028866208.1 | NM_012057.4 |
| Lcn2 | lipocalin 2 | XM_028884808.1 | NM_008491.1 |
| Mpo | myeloperoxidase | XM_028878233.1 | NM_010824.2 |
| Mrc1 | mannose receptor C-type 1 | XM_028870508.1 | NM_008625.2 |
| MTcox1 | mitochondrial cytochrome oxidase 1 | MG674647 | AP014886 |
| Nfe2l2 | nuclear factor, erythroid 2 like 2 | XM_028880611.1 | NM_010902.4 |
| Nfkb | nuclear factor kappa B subunit 2 | XM_028874704.1 | XM_006526743.4 |
| Nos2 | nitric oxide synthase 2 | XM_028866904.1 | AF065919.1 |
| Ptx3 | pentraxin 3 | XM_028888663.1 | NM_008987.3 |
| Saa3 | serum amyloid A 3 | XM_028880204.1 | NM_011315.3 |
| Slpi | secretory leukocyte peptidase inhibitor | XM_028868627.1 | NM_011414.3 |
| Sod2 | superoxide dismutase 2 | XM_028866678.1 | NM_013671.3 |
| Stat1 | signal transducer and activator of transcription 1 | KY451962.1 | XM_017319463.3 |
| Stat2 | signal transducer and activator of transcription 2 | XM_028855439.1 | NM_019963.2 |
| Stat3 | signal transducer and activator of transcription 3 | XM_028856799.1 | NM_213659.3 |
| Stat4 | signal transducer and activator of transcription 4 | XM_028887336.1 | NM_011487.5 |
| Tgfb1 | transforming growth factor beta 1 | XM_028859890.1 | NM_011577.2 |
| Tlr2 | toll like receptor 2 | XM_028894284.1 | NM_011905.3 |
| Tlr4 | toll like receptor 4 | XM_028887864.1 | NM_021297.3 |
| Tnf | tumor necrosis factor | XM_028887933.1 | NM_013693.3 |
